# Steppe Ancestry in Western Eurasia and the Spread of the Germanic Languages

**DOI:** 10.1101/2024.03.13.584607

**Authors:** Hugh McColl, Guus Kroonen, J. Víctor Moreno-Mayar, Frederik Valeur Seersholm, Gabriele Scorrano, Thomaz Pinotti, Tharsika Vimala, Søren M. Sindbæk, Per Ethelberg, Ralph Fyfe, Marie-José Gaillard, Hanne M. Ellegård Larsen, Morten Fischer Mortensen, Fabrice Demeter, Marie Louise S. Jørkov, Sophie Bergerbrant, Peter de Barros Damgaard, Morten E. Allentoft, Lasse Vinner, Charleen Gaunitz, Abigail Ramsøe, Isin Altinkaya, Rasmus Amund Henriksen, Evan K. Irving-Pease, KG Sjögren, Serena Sabatini, Anders Fischer, William Barrie, Andrés Ingason, Anders Rosengren, Andrew Vaughn, Jialu Cao, Jacqueline Staring, Jesper Stenderup, Fulya Eylem Yediay, Torbjörn Ahlström, Irakli Akhvlediani, Sofie Laurine Albris, Biyaslan Atabiev, E.V. Balanovskaya, Pernille Bangsgaard, Maria Giovanna Belcastro, Nick Card, Philippe Charlier, Elizaveta Chernykh, Torben Trier Christiansen, Alfredo Coppa, Maura De Coster, Sean Dexter Denham, Sophie Desenne, Jane Downes, Karin Margarita Frei, Olivér Gábor, U.B. Gadiev, Johan Zakarias Gårdsvoll, Zanette Tsigaridas Glørstad, Jesper Hansen, Stijn Heeren, Merete Henriksen, Volker Heyd, Mette Høj, Mads Kähler Holst, Rimantas Jankauskas, Henrik Janson, Mads Dengsø Jessen, Jens Winther Johannsen, Torkel Johansen, Ole Thirup Kastholm, Anton Kern♰, Ruslan Khaskhanov, Katrine Ipsen Kjær, Vladimir Kolosov, Lisette M. Kootker, Klaudia Kyselicová, Anne Christine Larsen, Thierry Lejars, Mette Løvschal, Niels Lynnerup, Yvonne Magnusson, V. Yu. Malashev, Kristiina Mannermaa, Vyacheslav Masyakin, Anne Lene Melheim, Inga Merkyte, Vyacheslav Moiseyev, Stig Bergmann Møller, Erika Molnár, Nadja Mortensen, Eileen Murphy, Bjarne Henning Nielsen, Doris Pany-Kucera, Bettina Schulz Paulsson, Gertjan Plets, Marcia S Ponce de León, Håkon Reiersen, Walter Reinhard, Antti Sajantila, Birgitte Skar, Vladimir Slavchev, Václav Smrčka, Lasse Sørensen, Georg Tiefengraber, Otto Christian Uldum, Helle Vandkilde, Jorge Vega, Daniele Vitali, Alexey Voloshinov, Sidsel Wåhlin, Holger Wendling, Anna Wessman, Helene Wilhelmson, Karin Wiltschke, João Zilhão, Christoph PE Zollikofer, Thorfinn Sand Korneliussen, Bruno Chaume, Jean-Paul Demoule, Thomas Werge, Line Olsen, Rasmus Nielsen, Lotte Hedeager, Kristian Kristiansen, Martin Sikora, Eske Willerslev

## Abstract

Today, Germanic languages, including German, English, Frisian, Dutch and the Nordic languages, are widely spoken in northwest Europe. However, key aspects of the assumed arrival and diversification of this linguistic group remain contentious^1–3^. By adding 712 new ancient human genomes we find an archaeologically elusive population entering Sweden from the Baltic region by around 4000 BP. This population became widespread throughout Scandinavia by 3500 BP, matching the contemporaneous distribution of Palaeo-Germanic, the Bronze Age predecessor of Proto-Germanic^4–6^. These Baltic immigrants thus offer a new potential vector for the first Germanic speakers to arrive in Scandinavia, some 800 years later than traditionally assumed^7–12^. Following the disintegration of Proto-Germanic^13–16^, we find by 1650 BP a southward push from Southern Scandinavia into presumed Celtic-speaking areas, including Germany, Poland and the Netherlands. During the Migration Period (1575–1375 BP), we see this ancestry representing West Germanic Anglo-Saxons in Britain, and Langobards in southern Europe. We find a related large-scale northward migration into Denmark and South Sweden corresponding with historically attested Danes and the expansion of Old Norse. These movements have direct implications for multiple linguistic hypotheses. Our findings show the power of combining genomics with historical linguistics and archaeology in creating a unified, integrated model for the emergence, spread and diversification of a linguistic group.

## Main

The arrival of Steppe-related groups in Europe around 5000 BP (calibrated years before present/1950 CE) is commonly known as the last major prehistoric migration into the region^17,18^. Archaeologically, two major European groupings of this time are the Corded Ware and the Bell Beaker complexes. Linguistically, these are widely accepted as being connected with the dispersals of multiple Indo-European language groups^19^. In Scandinavia, both complexes have been specifically associated with the introduction of the Germanic language group^7–10,20^. However, this association is complicated by the significant time gap of ∼2-3 millennia that exists between these first waves of Steppe-related ancestry (∼5000**–**4500 BP) and the appearance of the earliest Germanic written evidence (∼2000–1800 BP)^21,22^.

In Scandinavia, the arrival of Steppe ancestry (∼4800 BP) coincides with the archaeological transition from the Funnel Beaker Culture to the Corded Ware Complex^23^. By ∼4600 BP, cultural boundaries emerged separating the Scandinavian Peninsula’s Battle Axe culture from the Jutlandic Single Grave culture, with these showing connections respectively to the east and the south^24^. In addition, Scandinavia had cultural links associated with Bell Beaker groups of western and central Europe^25^. At this time, no linguistic data is available on the distribution of the Germanic language group. Only by the Late Bronze Age do linguistic interactions with Celtic-speaking groups in the south^4,5,26^ and Finno-Saamic-speaking groups in the east^6,27^ suggest that an early form of Germanic was present in the area between northern Germany and the East Baltic. Despite some preliminary evidence of genetic structure at this time in Scandinavia^28,29^, its formation as well as its potential for tracing the spread of Germanic remain unexplored.

By ∼2000 BP, the common Germanic linguistic ancestor was diverging into several subgroups, probably involving an initial East vs Northwest Germanic split^13–16,30^. This linguistic process was associated with various migrations during the Roman Iron Age (1950– 1575 BP) and the Migration Period (1575–1375 BP). However, at present, our understanding of the relevant population dynamics is limited. Genetic studies have detected a northern European origin for Late Iron Age and Migration Period individuals often ascribed by post-Classical authors as Goths, Anglo-Saxons and Langobards^31–35^, consistent with Late Antique historical sources. However, these studies have not confirmed their specific regions of origin. Consequently, no comprehensive model for the spread and diversification of the Germanic languages currently exists.

Ancient genomics have proven a means to address historical linguistic hypotheses^19^ but have been constrained by limited sample sizes and an inability to distinguish between closely related populations. However, recent studies reveal that with dense ancient DNA sampling, at sufficient sequencing depth for imputation, the detection of fine-scale population structure is now possible^28,29,36^.

To investigate the formation and diversification of Germanic-speaking populations and link the findings to historical linguistic theory, we sequenced the genomes of 712 ancient individuals to above 0.01X (Supplementary Note S4, Supplementary Table S1.1), with a focus on the Northern European Iron Age and the bordering Celtic-speaking region of western Europe^26^ (Fig. 1). Together with published ancient genomes from around the world, we selected samples with suitable average depth of coverage for imputation (∼0.1X for whole genomes)^28,29,37^. After overlapping with publicly available ancient SNP capture data suitable for imputation (∼1X on targeted SNPs), removing close relatives and applying quality filters, the final imputed dataset contained 578 new and 4,009 published individuals covering 690,211 SNPs (Supplementary Note S4).

**Fig. 1.**
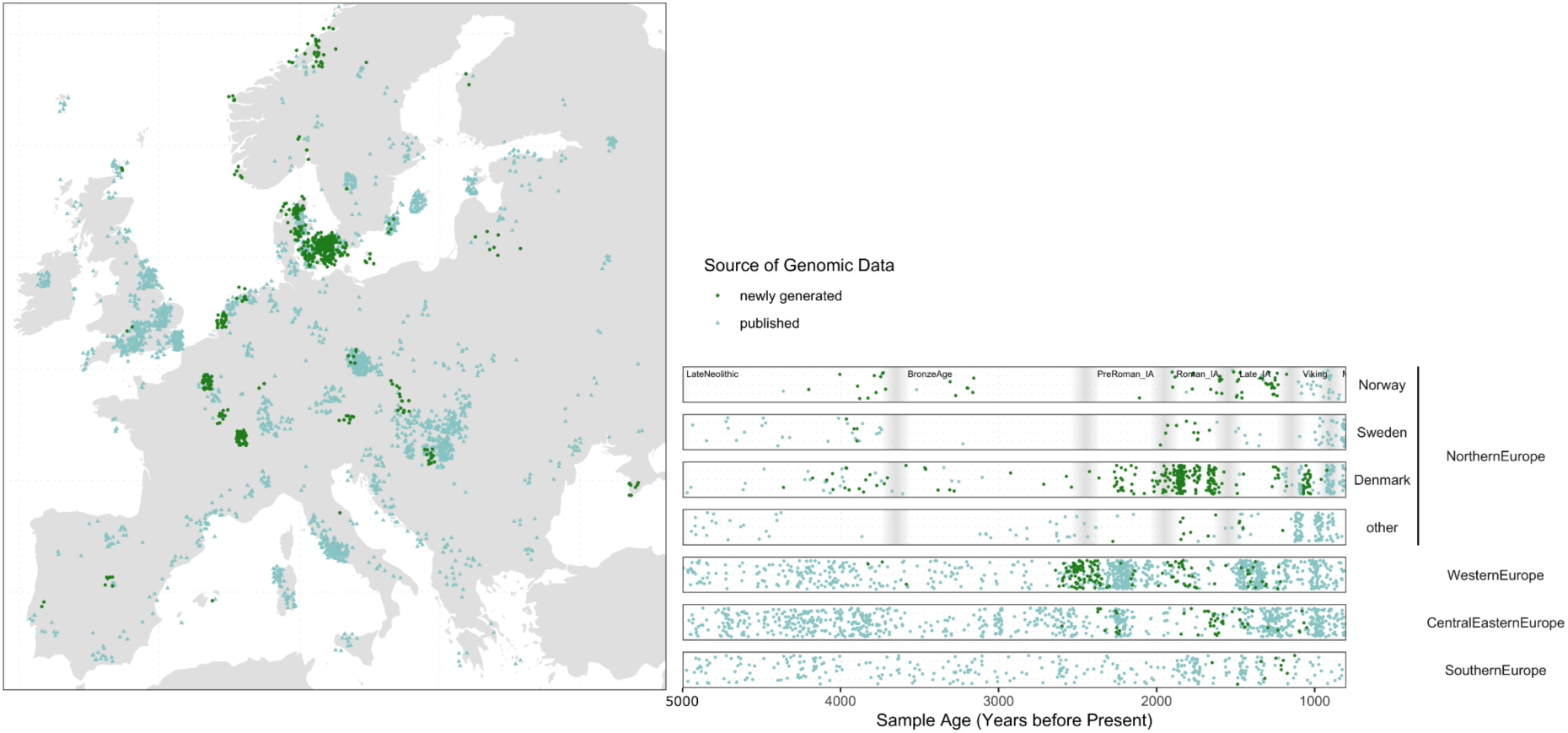
Geographic and temporal sampling of a subset of ancient individuals included in the final dataset. Newly generated (green) and published (light blue) ancient individuals from the Late Neolithic / Early Bronze Age to the Viking Age. Grey bars on the timeline represent the boundary between historical periods denoted in the top panel.

### Fine-scale resolution of early Steppe ancestry in Europe (5000–4000 BP)

We explored the genomic affinities between all individuals in the dataset using the identity-by-descent (IBD) hierarchical clustering method (Supplementary Note S5.2) and mixture modelling (Supplementary Note S5.3) to discern the closely related genomic ancestries^28^. Here, clusters form on the basis of the long shared genomic segments between all pairs of individuals within the dataset, rather than by proportions of the deeply diverging ancestries they carry. As discrete clustering does not display the complexities of admixture, potentially giving false impressions of continuity, we applied IBD mixture modelling to assess the genetic structure within the clusters. In brief, we created a ‘palette’ for every individual, based on the total length of IBD segments shared between that individual and all 386 clusters in the dataset. We then define sets of individuals from specific clusters as ‘sources’, and modelled the palettes of ‘target’ individuals as a mixture of all possible source palettes, using an non-negative least squares approach, similar to chromosome painting^38^. As a result, even though a temporally distant source and target individual may share very few segments, the shared segments with common intermediate populations are informative in the modelling. We find this method to be effective for the sampling density and temporal distances relevant to this study (Supplementary Note S5.3). To understand the general trends through time, we applied ordinary spatio-temporal kriging^39^ to the mixture modelling results (Supplementary Note S5.3, Extended Data Figures 1-7).

We focused our efforts within the last ∼5000 years, commonly considered the hallmark for introduction of Steppe ancestry across Europe and widely acknowledged as a likely terminus post quem for the spread of the Indo-European language family^17,18^. We find the majority of European individuals over the last 5,000 years fall within four main IBD clusters, with a varying geographical and cultural distinction for each (Supplementary Note S5.2.1). Based on this close correspondence with individuals assigned to various archaeological groups, we refer to these clusters as Yamnaya-, Corded Ware (East)-, Corded Ware (North)-, and Bell Beaker-related. Notably, individuals from each cluster are placed adjacent to each other in a standard western Eurasian PCA (Fig. 2, Supplementary Note S5.1), and each cluster occupies different positions along the well-established cline of Steppe-Farmer ancestry that formed in Europe from the Bronze Age.

**Fig. 2.**
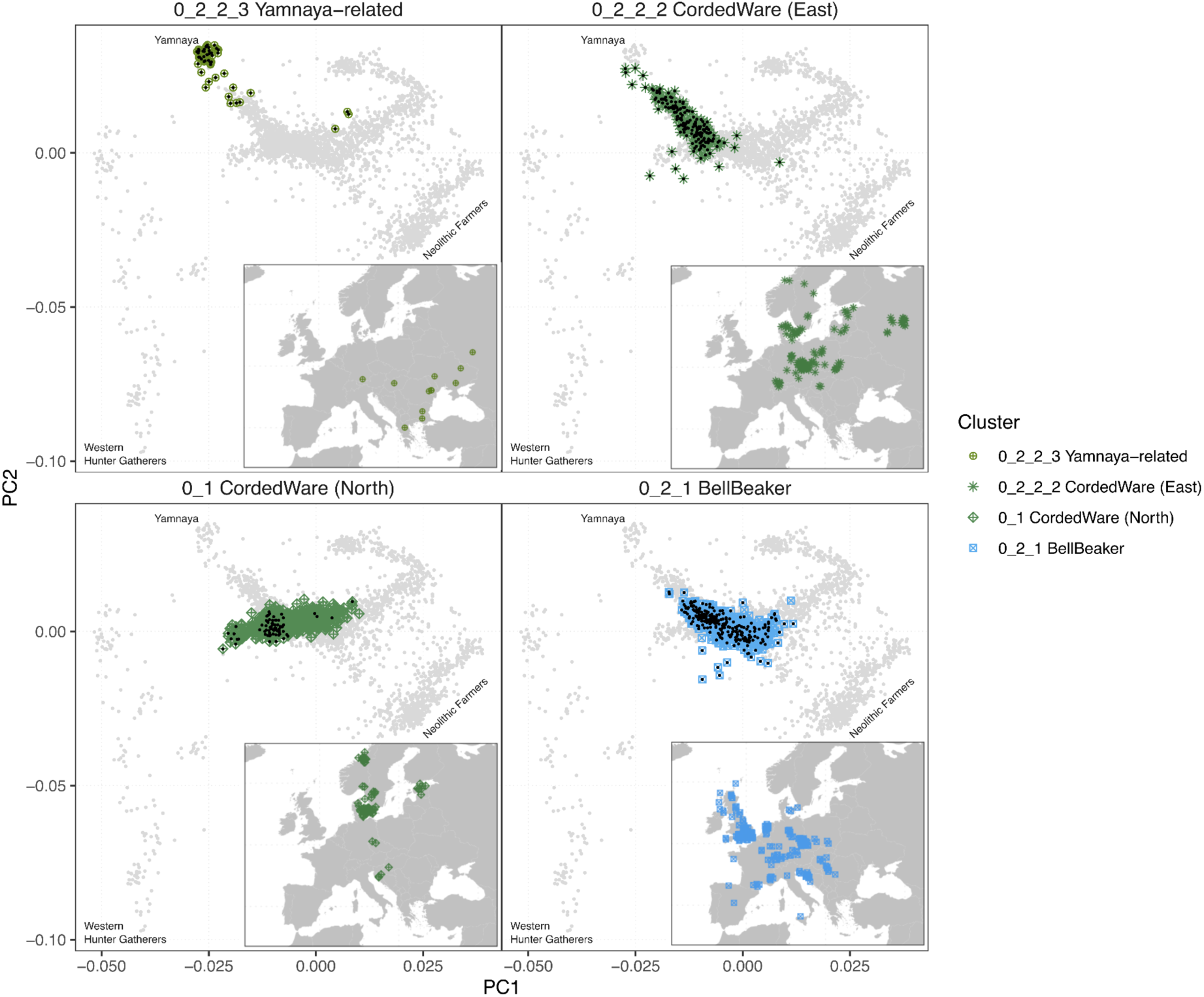
Distribution of the main four Steppe-related clusters in Geographical (inset) and western Eurasian PCA space. Samples older than 2800 BP are indicated with a black point on the PCA. On the map, only samples older than 2800 BP are shown.

Importantly, our mixture modelling results show that only samples that are modelled with a high proportion of Steppe ancestry fall within these clusters (Supplementary Fig. S5.16, Supplementary Note S5.2.3), meaning the Steppe ancestry of many individuals is not represented by the clustering. Of particular importance are many Bronze Age and later southern European individuals, whose clustering is informative of their Farmer rather than their Steppe ancestry. To understand how the Steppe ancestry of individuals of primarily Farming ancestry are related to the structure mentioned above, we applied IBD mixture modelling with sources representative of the Yamnaya, Corded Ware and Bell Beaker clusters (Supplementary Note S5.3).

We find the Steppe ancestry of the majority of the more southern individuals to be modelled as Bell Beaker-related (Fig. 3, Supplementary Note S5.2.3, Supplementary Fig. S5.17), together with a series of previously unidentified admixture clines (Fig. 4, Extended Data Fig.- 8, Supplementary Note S5.3.3). For the Yamnaya, Corded Ware and Bell Beaker-related source clusters, we found the difference in the mean proportion of Steppe ancestry to be significantly higher for other individuals archaeologically assigned to the corresponding culture, than to the other cultures (Supplementary Note S5.3.7). For many samples, an archaeological assignment was not present, however we found that samples with Steppe ancestry modelled primarily as Corded Ware-, Bell Beaker- or Yamnaya-related tend to correspond closely with the regions ascribed to each culture in the archaeological record (Fig. 3).

**Fig. 3.**
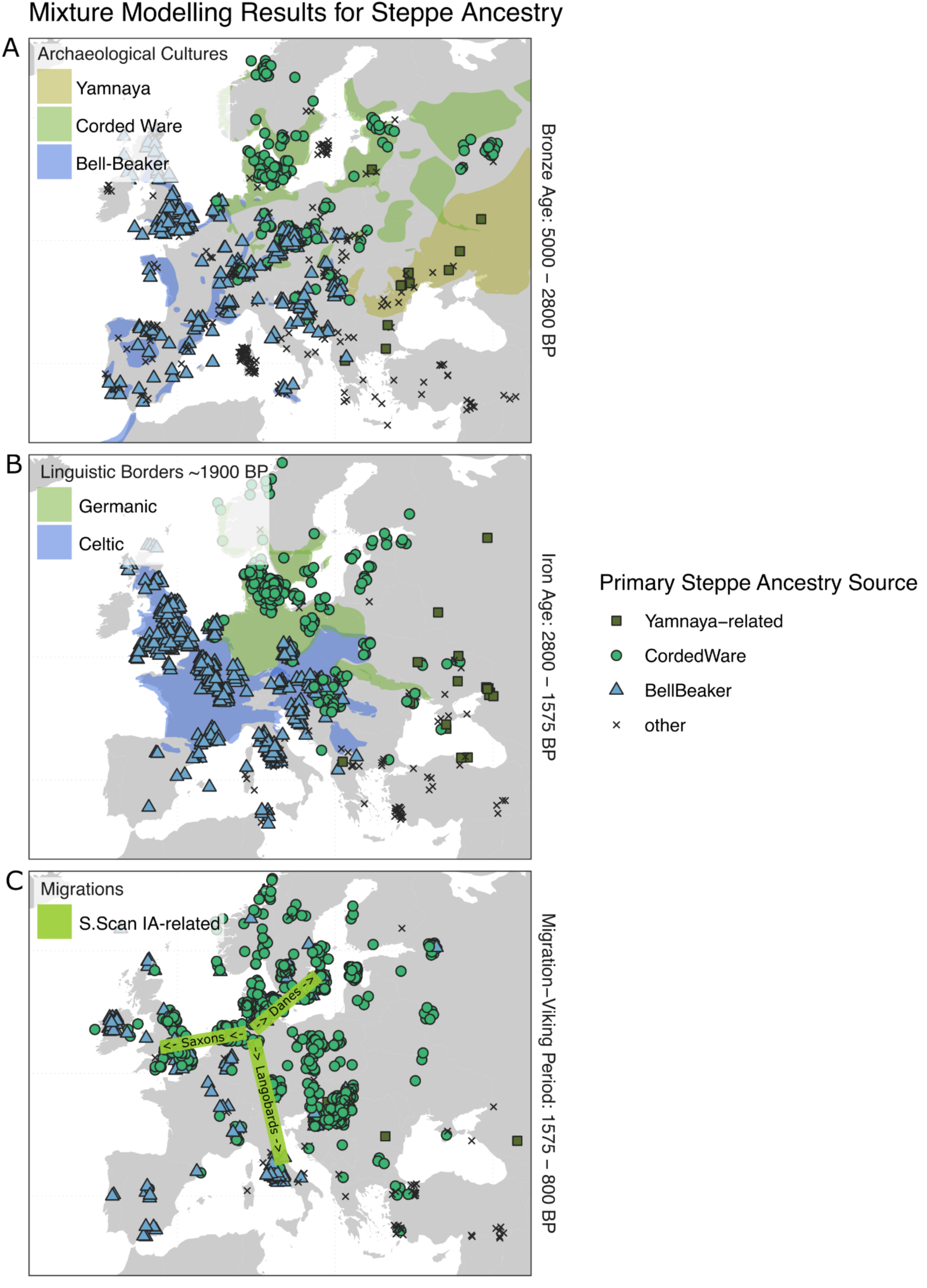
Geographical distributions of major archaeological cultures, language families and Steppe ancestry sources. Individuals modelled with less than 10% Steppe ancestry or less than 66% from one of the source groups are indicated with an ‘x’. Archaeological boundaries reproduced from Furholt et al.^40^.

**Fig. 4.**
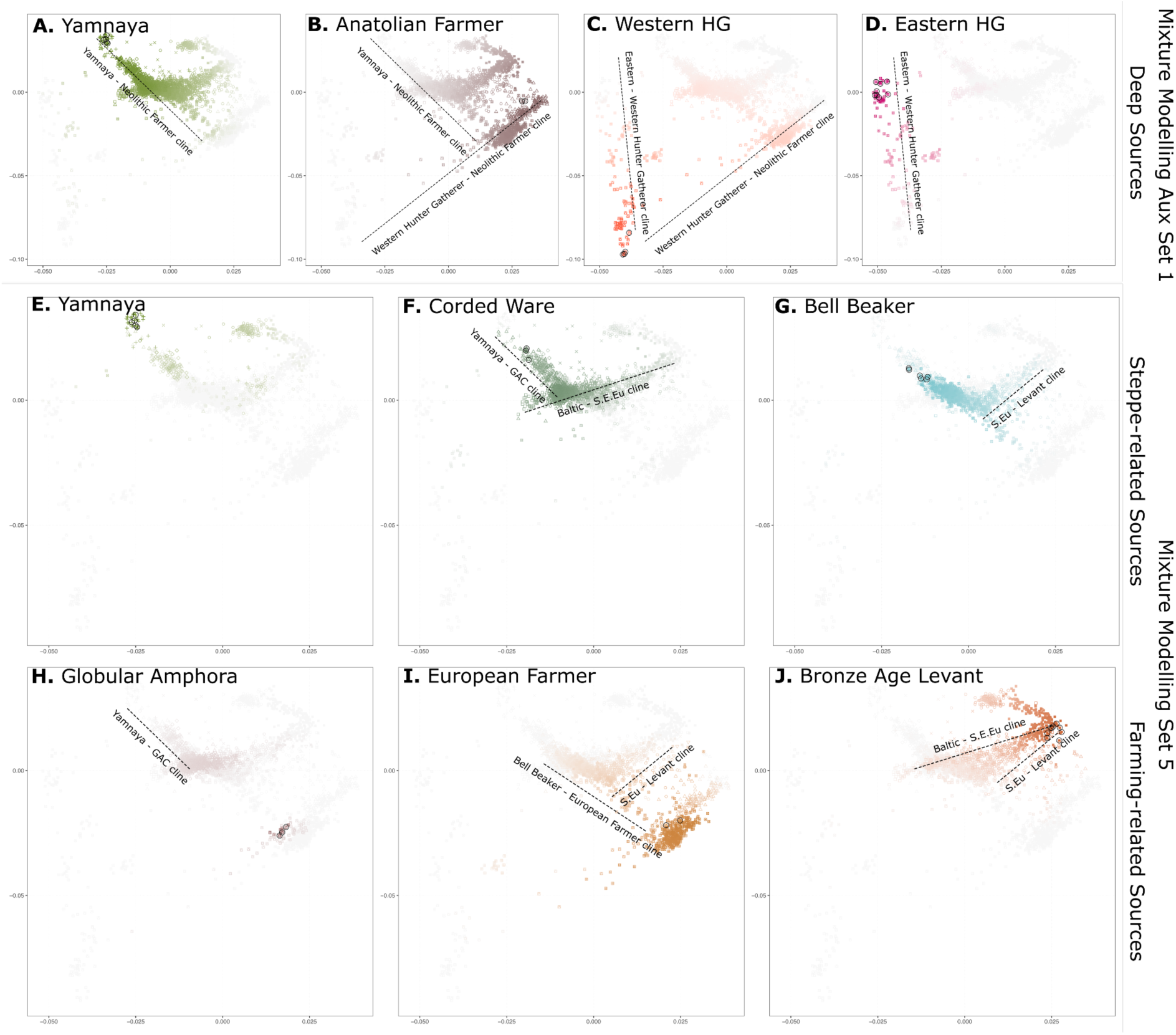
A subset of mixture modelling results highlighting admixture between various Steppe and Farming-related sources displayed on the western Eurasian PCA (Supplementary Note S5.3.1, Supplementary Fig. S5.20). Auxiliary Set 1 highlights the well-established clines representing the diversity of western Eurasian Hunter-Gatherers (C and D), the arrival of Neolithic Farmers (B) in Europe admixing with the local Hunter-Gatherers (C), and the arrival of Yamnaya-related ancestry (A) admixing with European Farmers. Mixture modelling Set 5 further highlights the clines between various Steppe-related ancestry sources: E) Yamnaya-related, F) Corded Ware-related and G) Bell Beaker-related, and Farmer ancestry-related sources: H) Globular Amphora Culture-related, I) European Farmer-related and J) Levant/Bronze Age Anatolia-related. Source individuals are circled here and detailed in Supplementary Table S5.3 (Set 5) and S5.4 (Aux Set 1). Admixture proportions follow a gradient from full colour (100%) to grey (0%).

During the Late Bronze Age (3150–2450 BP), the linguistically inferred presence of Palaeo-Germanic in Southern and Eastern Scandinavia falls within the geographic boundaries of the Corded Ware-related ancestries and cultures (Fig. 3). By the Roman Iron Age, Germanic-speaking regions continue to fall within the broader range of individuals of Corded Ware-related ancestry, bordered to the south by Celtic-speaking regions and Bell Beaker-related ancestries (Fig. 3). For Iron Age regions assigned as likely Germanic- or Celtic-speaking, we similarly find in all instances highly significant correlation with Corded Ware and Bell Beaker ancestries (Supplementary Note S5.3.7). This correspondence and continuity suggest that Germanic emerged and primarily continued to be spoken by populations of Corded Ware, rather than Bell Beaker-related ancestry.

From the Late Bronze Age onwards, irrespective of clusters, the Steppe ancestry in almost all Europeans is modelled by either Corded Ware (East) or the Bell Beaker sources (Fig. 3, Extended Data Fig.-4). Notable exceptions appear in the coastal region of the Netherlands and in Bohemia, where the two complexes overlap (Supplementary Note S5.7.2). In the Netherlands, we identify an ‘Eastern North Sea’ cluster that persists from 3700 BP to 1700 BP and is modelled with equal proportions of Corded Ware- and Bell Beaker-related ancestry (Supplementary Fig. S5.172, Supplementary Note S5.7.2). The existence of this genetic outlier has a linguistic analogue in the so-called ‘Nordwestblock’ hypothesis, which claims that the Netherlands harboured an Indo-European language distinct from both Celtic and Germanic^41^.

### Scandinavian population dynamics from the Late Neolithic/Bronze Age to the Iron Age (4000–2500 BP)

Although Corded Ware-related ancestry persisted in northern Europe from the Bronze Age to the Iron Age (Fig. 3), the extent to which migrations occurred within this region is poorly understood. Linguistically, this period encompasses the prehistoric evolution of the Germanic branch following its split from the Indo-European proto-language and comprises two phases: Palaeo- and Proto-Germanic. Lexical exchanges with Celtic in the south^4,5^ and Finno-Saamic in the east^6,27^ indicate that Palaeo-Germanic was present between northern Germany and the East Baltic by the Late Bronze Age. The timing of these lexical exchanges is evident in loans predating the so-called Germanic sound shifts, which had happened by the beginning of the Iron Age, which in Scandinavia started around 2500 BP. The entry of Germanic loans into the Finnic Uralic group cannot be earlier than the putative arrival of their speakers in the Baltic, which according to the latest studies coincides with the arrival of Siberian ancestry after 3500 BP and likely with the cist graves from around 3150-3050 BP^42^. Some early Celtic loans likewise predate the Germanic sound shifts. As a result Palaeo-Germanic can be said to have been in contact with both Finnic in the east and Celtic in the south during the period 3200-2500 BP, a period corresponding to the Late Bronze Age. To investigate relevant demographic shifts at this time, we first established the population substructure within Bronze Age Scandinavia by reclustering all ancient samples older than 2800 BP (Supplementary Note S5.2.2, Supplementary Table S5.2).

Within Scandinavia, three clusters are apparent (Extended Data Fig.-9): 1) an Early Scandinavian cluster, including individuals from the Swedish Battle Axe Culture, some of the oldest Danish individuals, and nearly all Norwegians; 2) a later ‘Southern Scandinavian’ cluster restricted to Denmark and the southern tip of Sweden; and 3) an even later ‘Eastern Scandinavian’ cluster, spanning Sweden and geographically overlapping with the Southern Scandinavia cluster. In all three clusters, we see a striking correspondence between Y-haplogroups and IBD clusters (Extended Data Fig.-9A), largely driven by different frequencies of haplogroups I1a (I1a-DF29), R1a1a1b1a3a (R1a-Z284) and R1b1a1b1a1a1 (R1b-U106), all being strongly associated with Scandinavian ancestry (Supplementary Note S5.2.2). We find a large degree of geographic overlap between the Early, Southern and Eastern Scandinavian clusters of the pre-2800 BP individuals and three subclusters detected in the original northern Corded Ware cluster: Western, Southern and Eastern Scandinavian, respectively (Extended Data Fig.-9B, Supplementary Note S5.2.2). The presence of genetic structure in the Bronze Age has previously been detected^28^, however, its formation and relation to the Bronze Age distribution of Palaeo-Germanic, and to the genetic structure of Iron Age Scandinavia, have not been addressed.

The three pre-2800 BP clusters are found across the region in which Palaeo-Germanic is assumed to have been spoken, and understanding their formation may be informative for the introduction of the Germanic branch of the Indo-European language family to Scandinavia. To investigate this, we applied mixture modelling, using a variety of deep Hunter-Gatherer sources. We find the Yamnaya-related ancestry in the three Scandinavian clusters to be modelled as Eastern Hunter-Gatherer and Caucasus Hunter-Gatherer (Supplementary Fig. S5.27, Supplementary Fig. S5.23: Set 2). This is in agreement with expectations for Steppe ancestry in Europe^43^. However, the early (4100–2800 BP) Eastern Scandinavians are distinct in their relatively high proportion of Eastern Hunter-Gatherer ancestry, compared to Western and Southern Scandinavians (Supplementary Note S5.3.4). To identify the specific source of this Hunter-Gatherer ancestry, we included additional Hunter-Gatherer sources from the region (Norway, Sweden, Denmark, Latvia and Lithuania) together with Yamnaya. We find that the Eastern Scandinavians’ Hunter-Gatherer ancestry is modelled entirely by the Latvian/Lithuanian Hunter-Gatherer source from across the Baltic, rather than the local Scandinavian Hunter-Gatherers (Supplementary Note S5.3.4, Supplementary Fig. S5.23: Set 3). In contrast, the Southern and Western Scandinavians are modelled with additional Western Hunter-Gatherer ancestry. These mixture modelling results are consistent with the subtle differences in the distribution of individuals older than 2800 BP in the Scandinavian clusters in the western Eurasian PCA (Supplementary Fig. S5.7, Supplementary Fig. S5.8, Supplementary Fig. S5.9). To confirm these patterns, we ran rotating qpAdm models with 10 different Hunter-Gatherer clusters as sources, including representatives of the previously identified genetic structure of Mesolithic Scandinavia^44–46^. While an early Corded Ware cluster could be modelled as only Yamnaya and Globular Amphora, the Southern and Western Scandinavians each required a contribution by Hunter Gatherers from continental Europe and Scandinavia respectively. Only when including a second Hunter-gatherer source could the Eastern Scandinavians be well modelled, in many combinations, suggesting the true source has not yet been sampled. The presence of a non-local Hunter-Gatherer ancestry suggests the three clusters did not diversify from a single migration into the region. Although individuals from all three clusters likely spoke Indo-European languages, the Corded-Ware related Battle Axe Culture of the Scandinavian Peninsula and the Single Grave Culture of the Jutlandic Peninsular are traditionally assumed to be the vectors for Indo-European dialects ancestral to Germanic (Supplementary Note S7.1.3), fitting closely with the early distribution of Western and Southern Scandinavian clusters respectively. However, the Eastern Scandinavian cluster must also be considered as a potential vector for the arrival of Germanic.

The Eastern Scandinavians first detection 6-800 years after the earliest Corded Ware populations in Scandinavia (Extended Data Fig.-9A), and the presence of a Hunter-Gatherer ancestry, not well represented by the three waves of Hunter-Gatherers previously identified in Scandinavia, points to an additional, late arrival into Scandinavia by the ancestors of the Eastern Scandinavians. The Hunter-Gatherer ancestry suggests a link across the Baltic or from the northeast along the Baltic coastline. With regards to their Steppe-related ancestry, the Corded Ware individuals from Lithuania (4842–4496 BP), Latvia (4833 BP) and Estonia (4638–4400 BP) are not well-modelled by the Eastern Scandinavians, suggesting a source region further north (Supplementary Fig. S5.28). Notably, Corded Ware and Hunter-Gatherer genomes from Finland and the northeast coast of Sweden are not represented in the dataset, and may also be a suitable source for the Hunter-Gatherer ancestry in Eastern Scandinavians. Such a location would be consistent with strontium isotopes of the Late Neolithic Swedes of Central Sweden which link with eastern and northern Sweden, Finland and possibly Karelia and with similarities in pottery styles between Late Neolithic Sweden and the Kiukainen culture (4500–3800 BP) of southwestern Finland and the Åland Islands^47–50^. Combined, the results point to the presence of an unsampled hunter-gatherer population, likely carrying I1 haplogroups, admixing with a Corded Ware-related population, similar to those of Scandinavia, to form the Eastern Scandinavians somewhere between Finland and Northeast Sweden. At present, no genomes from this region and time period exist.

Interestingly, the arrival of this first Eastern Scandinavian ancestry coincides with a new burial rite of gallery graves in South Sweden, the Danish Isles^25^ and Norway^51^, a new house type^52–54^ and the first durative bronze networks^55^. These events culminated at the onset of the Nordic Bronze Age (3550 BP) with cultural homogenisation and the end of the east-west divide in northern Europe^53^.

The Nordic Bronze Age is key in linking the three genetic clusters of the Late Neolithic with the demographic structure of the Iron Age. Archaeologically, the Nordic Bronze Age is a period of strong cultural homogenisation in southern Scandinavia, starting around 3600 BP, forming the so-called Nordic Cultural Zone, which lasted until the start of the Iron Age (2450 BP)^56,57^. This homogenisation is mirrored genetically. At the beginning of the period, we observe a number of admixed Norwegian and Danish Bronze Age outliers, each carrying local and Eastern Scandinavian ancestry (Supplementary Fig. S5.29). By the Iron Age, however, the population structure has become more homogenised by the widespread presence of Eastern Scandinavian ancestry, the trend of which is visible in the spatio-temporal kriging results produced from the mixture modelling output (Fig. 5, Supplementary Note S5.3.8). The impact of this expansion is most apparent on the Danish Isles, followed by Norway and finally the Jutlandic Peninsula. The least impact is seen in Jutland, where a large degree of genetic structure distinguishes it from the rest of Scandinavia (Fig. 5).

**Fig. 5.**
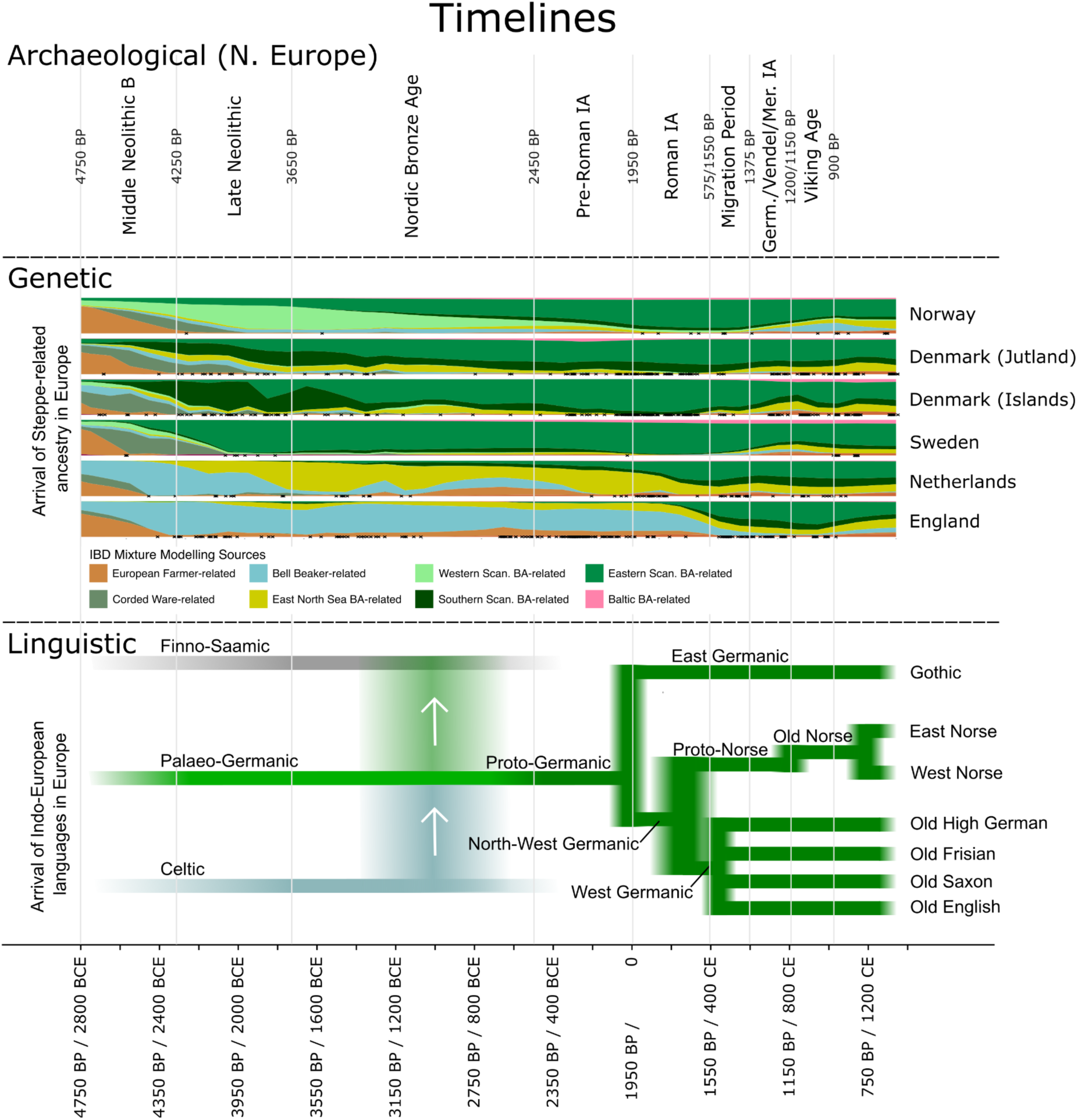
A timeline showing Archaeological Periods, Spatiotemporal Kriging results with the general trends of the proportion of ancestries using Bronze Age European sources (Set 5), and relevant events in the formation of Germanic languages. During the Late Neolithic, the highest proportions of Eastern, Western and Southern Scandinavian Bronze Age ancestry are in the local region, however some ancestry remains modelled as Corded Ware (East)-related. Throughout the Bronze Age and into the Iron Age, Eastern Scandinavian ancestry becomes widespread, coinciding with loan words passing from Celtic to Germanic, and Germanic to Finno-Saamic (indicated by the arrows). Samples within 250 km of the coordinates assigned to each country are indicated with an ‘x’, providing insight into the confidence with which we can assess the timing of the transition at these coordinates. The coordinates representing each country can be found in Supplementary Fig. S5.134.

To assess whether the spread of Eastern Scandinavian ancestry is compatible with being a vector for Palaeo-Germanic, understanding the timing of this migration is crucial. Linguistically, the Late Bronze Age is the period during which Palaeo-Germanic donated vocabulary to Finno-Saamic in the east^6,27^ and adopted vocabulary from Celtic in the south^4,5^ suggesting that Palaeo-Germanic was spoken within Sweden and Denmark^4,5^. However, no demographic vector has yet been identified to match this distribution. Although genomes are scarce for this period due to prevalent cremation practices, we used DATES^58^ to date the admixture time between the Eastern Scandinavians and the Southern Scandinavians for various Bronze Age and Iron Age clusters in Denmark, to confirm the formation of the Iron Age populations matched that of the earliest admixed individuals we detect directly (Supplementary Note S5.5). The timing of the earliest admixed individuals fits closely with the point estimates from the DATES results, suggesting an admixture time of 3750–3250 BP, consistent with widespread Eastern Scandinavian ancestry within the spatio-temporal range proposed for Palaeo-Germanic. The admixture time for Eastern and local Bronze Age ancestry in Norway (4200–2600 BP) presents a similar range, with point estimates again closely matching the first direct evidence of admixed Western Scandinavian and Eastern Scandinavian individuals. Conversely, at the time of the incorporation of loanwords (3200– 2500 BP), Western Scandinavian and Southern Scandinavian Bronze Age ancestry had decreased both in geographical range and proportion. Even at its most expansive distribution, Southern Scandinavian ancestry did not border the region in which Finno-Saamic was spoken, and by 4200-4000, Western Scandinavian bordered neither Finno-Saamic nor Celtic. Combined we find Eastern Scandinavian to be the most parsimonious vector for Palaeo-Germanic.

The link between Eastern Scandinavian ancestry and Palaeo-Germanic, and the arrival of Eastern Scandinavian ancestry in Norway generates the hypothesis of a dialect of Palaeo-Germanic being introduced to Norway, however we note that no linguistic evidence of a presence in Norway at this time exists. Moreover, the apparent formation of Eastern Scandinavians from the Baltic reigon potentially sheds new light on the well-known problem of the similarities shared between the Germanic and Balto-Slavic branches of the Indo-European language family, pointing to prehistoric borrowing, a linguistic subclade or a combination of both^59^.

### Early Iron Age Scandinavia and the formation of Proto-Germanic (2450– 1950 BP)

By the onset of the Northern European Iron Age (2450 BP), Palaeo-Germanic had transitioned into Proto-Germanic, the last common ancestor of the Germanic languages, through a series of defining sound changes^60–63^. This linguistic transition has been ascribed to the assimilation of speakers of an unknown language^21,64^, potentially indicative of admixture. The Proto-Germanic speech community is assumed to have been present in southern Scandinavia and northern Germany throughout the Pre-Roman Iron Age (2500–1950 BP)^65,66^, with the Nordic Iron Age and the Jastorf culture being likely archaeological contexts^3,67^. To date, it remains untested whether such admixture occurred.

Based on IBD mixture modelling using the admixed Bronze Age individuals as sources (Fig. 5, Supplementary Fig. S5.32) and the DATES results (Supplementary Note S5.5), we find evidence of genetic continuity in Jutland throughout this period. In the Danish Isles, additional Eastern Scandinavian ancestry is present relative to Jutland; however, the DATES analysis indicates that the formation of the structure of the two regions occurred at a similar time. The absence of major admixture events around the supposed transition from Palaeo- to Proto-Germanic suggests this linguistic phase shift may have been induced by other factors at the onset of the Iron Age, such as changes in social organisation or language-internal developments.

Between 1950 and 1550 BP, the earliest runic carvings attest to relatively undiverged Germanic dialects being spoken across Denmark, Norway and Sweden. At this time, although Eastern Scandinavian BA ancestry was widespread across the region, the genetic structure that formed during the Bronze Age still persisted between Jutland, the Danish Isles, Sweden and Norway. In Iron Age Jutland, and to a lesser degree the Danish Isles and Norway, individuals continue to be modelled with a minor proportion of the local Scandinavian Bronze Age ancestry (Fig. 5, Supplementary Fig. S5.29: Set 5). The combined genetic and linguistic insights reveal that these genetically distinct Iron Age populations were united by a Germanic dialect continuum.

### Roman Iron Age and the Migration Period expansions from Scandinavia

For the Roman Iron Age and Migration Period (1950–1375 BP), Late Antique historical sources suggest that multiple migrations of Germanic-speaking groups occurred out of Scandinavia, with precise but often contradictory homelands^68^. While genetic studies have confirmed a northern European origin for some individuals associated with these populations^31–34^, ascertaining precise source regions has proven challenging. To obtain more precise geographical sources of migration, we included a series of the admixed Iron Age clusters in the mixture modelling. These included subclusters from northern Jutland, the Danish Isles, Sweden, and Norway (Supplementary Note S5.3.1, Set 6). At a broader scale, we find a pattern south of the Nordic region that indicates independent migrations from Jutland and the Scandinavian Peninsula. In western Europe, including present-day Germany, the Netherlands and England, we find the Jutlandic Iron Age source to be the primary Scandinavian ancestry (Fig 6A, Extended Data Fig.-10). In contrast, in the coastal region south and east of the Baltic Sea, we find that populations are primarily mixtures of Swedish Eastern Scandinavian and Baltic Bronze Age ancestries (Extended Data Fig.-10).

**Fig. 6.**
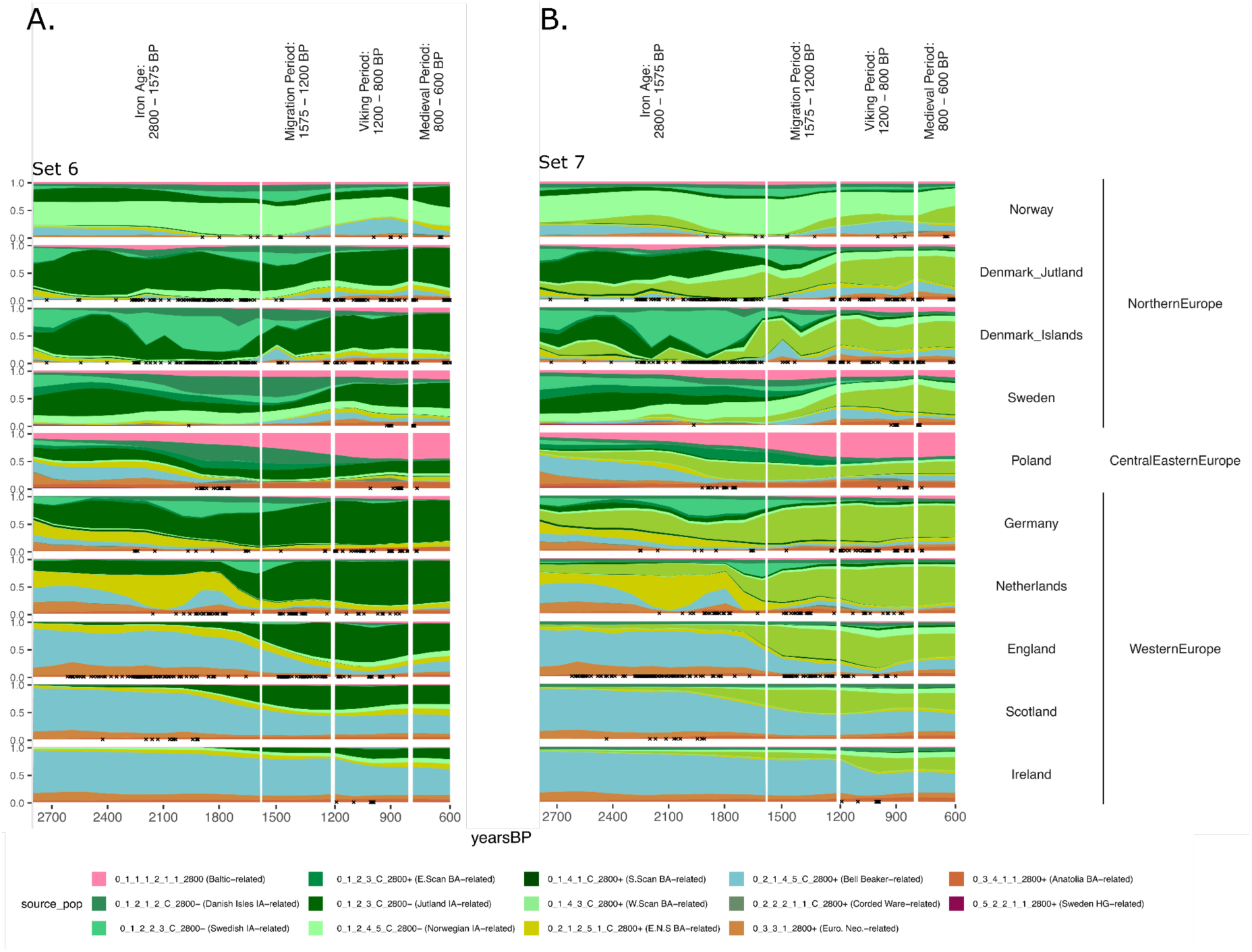
Spatiotemporal Kriging results showing the general trends of the proportion of ancestries modelled for each Iron Age individual from across Europe, including Iron Age European sources from Jutland, the Danish Isles, Sweden and Norway (Panel A, Set 6), and also including northern Germany (Panel B, Set 7). Samples with 250 km of these coordinates assigned to each country are indicated with an ‘x’. The coordinates representing each country can be found in Supplementary Fig. S5.134.

Historical and archaeological evidence suggest various regions along the Eastern North Sea coast, from the Netherlands to Jutland, as the sources of the migrations bringing the West Germanic predecessor of Old English to Britain^69^. Although genetic studies have indicated that a large-scale migration indeed occurred at this time, the precise source region has remained unclear, with candidates ranging from the Netherlands to southern Sweden^33^. We therefore added an additional Iron Age source from Mecklenburg, Northern Germany (Supplementary Note S5.3.1, Set 7), to the mixture modelling. This showed ancestry in Anglo-Saxons from England, Frisians from the Netherlands and Iron Age Germans to be modelled as the northern German source (Fig. 6B, Extended Data Fig.-11). The observed heterogeneity is consistent with the various homelands of the Angles, Saxons and Jutes along the Eastern North Sea coast migrating to Britain during this period (Extended Data Fig.-12, Supplementary Note S5.7.5). Prior to these expansions, individuals from northern Jutland, the Danish Isles and Sweden on the contrary, are modelled primarily as the local Iron Age ancestry. Therefore, we can reject Northern Jutland, the Danish Isles and Sweden as the main source areas for the Anglo-Saxon migrations to Britain.

A similar cultural transition to that in Britain occurred in the Netherlands. Following the Roman period, Dutch coastal areas saw a habitation hiatus around 1600 BP and the subsequent appearance of a new material culture that is often referred to as Anglo-Saxon in nature, and which may be related to the emergence of the Frisians^70^. As in Britain, we see a genetic transition occurring from local to northern German/southern Jutlandic ancestry. Here, the incoming ancestry becomes dominant in the area previously dominated by the distinct Eastern North Sea population (Supplementary Note S5.7.2). Although no unadmixed Eastern North Sea populations are found during the Migration Period, the population replacement was not complete: many later individuals from Northern and Western Europe from this time onward carry small proportions of Eastern North Sea ancestry.

A third West Germanic-speaking population, the Langobards^71^, are historically documented in the regions of today’s Czech Republic, Hungary and Italy and have genetically been linked to northern Europe^32^. However, according to post-classical origin legends, their homeland is in Southern Scandinavia specifically^72^. We find that the majority of the Langobards are indeed modelled by the Jutlandic Iron Age source (set 6), supporting a homeland around Jutland or northern Germany (Supplementary Note S5.7.4). Notably, there are three outlier Langobards from the Czech Republic and Hungary being of Eastern Scandinavian origin, and a number of individuals who genetically appear to be local. This close link between Southern Scandinavian ancestry and linguistically West Germanic groups is also seen in Germany (Extended Data Fig.-14). Here, 56 of 76 individuals between 1650 and 775 BP fall within the 0_1_3_x clusters, who tend to be modelled by the Northern German source (Supplementary Table S5.1).

While West Germanic groups appear closely linked with Northern German/Southern Jutland ancestry, consistent with historical evidence, the genetic affiliation of East Germanic groups is currently unknown. A 6th-century origin legend of the East Germanic Goths claims a homeland on the Scandinavian Peninsula^73^. The earliest evidence of plausibly East Germanic-speaking populations out of Scandinavia are the individuals from the Wielbark culture of present-day Poland, dated to around 1950 BP^34^. We find these individuals to be modelled primarily by Eastern Scandinavian BA and Swedish Iron Age ancestry (Supplementary Note S5.7.4), supporting a migration from a region and population distinct from those of the West Germanic groups – a scenario consistent with Gothic oral history^68^. In contrast, later individuals associated with the East Germanic-speaking groups, the Ukrainian Ostrogoths and the Iberian Visigoths, are modelled with local ancestries (Supplementary Note S5.7.4). Two exceptions are the Iberian Visigoths, who genetically fall on the Baltic cline, suggesting an origin in North East Europe, but not specifically in Eastern Scandinavia. The genetic evidence here is consistent with a two-step model on the ethnogenesis of the Goths: an initial migration from Scandinavia, followed by the acculturalization of others, including Baltic and Slavic groups^74^.

### Migration Period expansions into Denmark and Sweden

Unlike West and East Germanic, the distribution of North Germanic across Scandinavia is not typically associated with population migrations. Runic inscriptions from across this area testify to Germanic dialects that remained relatively similar to Proto-Germanic between 2000 and 1500 BP. However, from the beginning of the Migration Period to the Viking Period (1575–1200 BP), accelerated linguistic change led to the formation of Old Norse^75^, the common language of Viking-Age Scandinavians. Following this linguistic transition, the original Germanic runic script, known as the Elder Futhark, was fundamentally remodelled, giving rise to the Younger Futhark, a script exclusively used to document Old Norse^76,77^.

Interestingly, we observe a striking genetic transition in the Danish Isles and southern Sweden around the time of this linguistic transition. Prior to 1575 BP, individuals in these areas are modelled almost exclusively with Eastern Scandinavian Bronze Age ancestry, whereas by 1250 BP, i.e. approximately 100 years before the Viking Period, and increase in Southern Scandinavian and ENS ancestry is detected (Fig. 6, Extended Data Fig.-13). By using the Iron Age Scandinavian sources, the extent of the impact can be seen (Extended Data Fig.-10). On the Danish Isles, the local ancestry that was widespread from 2200 to 1600 BP essentially disappears by 1250 BP, however the rate at which this transition occurs is not clear. This contrasts with nearby regions, where Jutlandic Iron Age Southern Scandinavian and Swedish Iron Age Eastern Scandinavian ancestry persists through to the Viking Age, despite being less prominent than before.

To understand the source of these migrations, we turned to the period from 1250 BP onwards, when sampling improves as inhumation burials become more common. Here, most individuals can be modelled with small proportions of ancestries that prior to 1575 BP were primarily found south of Scandinavia: Eastern North Sea coast ancestry, Celtic ancestry from Britain, Ireland and France, Slavic-related ancestry from Northeastern Europe (Supplementary Note S5.7.6), and European Farming ancestry from western Europe (Extended Data Fig.-10, Extended Data Fig.-14). While previous studies have documented an influx of these more southern ancestries^78,79^, continuity of the local Scandinavian ancestry has generally been assumed.

To further investigate the transition within Denmark and southern Sweden, we again applied the mixture modelling with the Southern Scandinavian Iron Age sources from northern Jutland and from northern Germany (Extended Data Fig.-11). We find the Scandinavian ancestry of the later Danish Isles and southern Sweden individuals to be modelled as the northern German IA source ancestry rather than the more local northern Jutland IA source. This suggests large-scale population movements into Denmark and southern Sweden. In Jutland, we also see an influx of northern German IA ancestry. However, the impact is much less pronounced than for the Danish Isles. Due to the absence of samples between northern Jutland and northern Germany, we cannot rule out a southern Jutland origin as a source.

The above findings of a large-scale demographic shift have potential relevance for understanding the linguistic dynamics of Iron Age Scandinavia, suggesting that the emergence of Old Norse coincides with a period of social and demographic instability^80^. In addition, the spread of Southern Scandinavian ancestry may partially explain why the Old Norse language was referred to by its speakers as *dǫnsk tunga*, i.e. the Danish tongue, even in Norway, Iceland and Sweden^81,82^. However, the close correspondence between Northern German/Southern Jutlandic IA ancestry and Old Norse linguistic identity in Southern Scandinavia is not seen in Norway. Here, similar social changes as in South Scandinavia occurred during the same period, however, the Iron Age to Viking Age transition in Norway is one of genetic continuity (Supplementary Note S5.7.3). In Norway, the cultural changes occurred with limited genetic impact from populations from continental Europe. With the exception of a single early Viking sample, the majority of Viking Age Norwegians display either local ancestry or ancestry reflecting back-migrations from Celtic regions of Britain and Ireland. Of note, the linguistic boundary between the East and West Norse dialect areas^75,83^ roughly corresponds to the border between Southern Scandinavian and Western Scandinavian ancestry during the Viking Age (Figures S5.108 and S5.110).

Due to cremation practices, few genomes exist from the transition period (1575–1200 BP), requiring us to rely on additional lines of evidence for population dynamics during this period to assess when these major transitions in Denmark and Southern Sweden may have occurred (Extended Data Fig.-15). Volcanic activity (1414 and 1411 BP) and the Justinian Plague (1409 BP) caused a population decline in Scandinavia, from which the populations did not fully recover until around 1300 BP (Supplementary Note S7.1.4)^84^. However, despite some abandonment or depopulation of marginal subsistence areas^85^, there was continuity in more fertile and southern areas (Supplementary Note S6.4). For those who survived, the subsequent improving conditions and relative abundance of resources due to a lower population size would have created the opportunity for rapid expansion, as attested to in historical sources in other areas (Supplementary Note S7.1.4). Despite these dramatic events, the major cultural transitions began earlier, between 1550 and 1450 BP, and persisted through the Viking Age (Supplementary Note S7.1.4), according to antique sources mentioning the Danes living in South Scandinavia by 1450 BP^86,87^ and oral histories indicating the South Scandinavian royal lineage of the Danes, Swedes and Norwegians being initiated between 1550 and 1500 BP^88^. Based on the present archaeological and historical evidence (Supplementary Note S7.1.4), we may thus conclude that the major population shift in South Scandinavia between the Roman and the Viking periods was not solely driven by the climate events or plague of 1450–1350 BP, but instead likely took hold between 1550 and 1450 BP, coinciding with the Anglo-Saxon migrations and corresponding to a major cultural change with the introduction of a new elite culture. Historical texts and legends suggest various origins of the Danes ^68,86–88^, discussed in detail in Supplementary Note S7.1.4. Here we show genetically that the Migration Period population influx in South Scandinavia had origins around Northern Germany, with close similarity to that of the Anglo-Saxons.

This result has impacts on the interpretations of Gretzinger et al.^33^ in relation to the source region for Saxon migrations, and Margaryan et al.^78^ in relation to the major influx of Danish Viking Age ancestry into England (Supplementary Note S5.7.5.). Furthermore, the dense sampling and high resolution demographic inference have allowed us to establish a baseline ancestry for various regions and subsequently identify outliers (Supplementary Note S5.7.1).

### Conclusions

Our findings constitute a fundamental revision to the formation of West Eurasian ancestry in Europe. By additionally correlating these genetic findings with archaeological and linguistic evidence, we complement previous interpretations of the archaeological record after the Scandinavian Middle Neolithic B (4800 BP), identifying multiple populations movements, and propose a new model for the emergence and spread of the Germanic languages.

Following the arrival of Steppe ancestry in Scandinavia during the Middle Neolithic B (4800 BP), we detect the arrival of the closely related Eastern Scandinavian ancestry during the Late Neolithic with links to the North East Baltic region (4000 BP), offering a previously unknown vector for the introduction of an Indo-European language to Scandinavia. We cannot rule out more complex scenarios, for example, in which Palaeo-Germanic spread through culture diffusion, with no genetic impact, in the opposite direction to the spread of Eastern Scandinavian ancestry. However, the fact that Eastern Scandinavian ancestry was widespread by the Nordic Bronze Age, encompassing the broad region where Palaeo-Germanic is hypothesised to have been spoken, bordering Finno-Saamic in the east and Celtic in the south, makes Eastern Scandinavians the most parsimonious vector for the spread of Palaeo-Germanic. By 3500 BP, Eastern Scandinavian ancestry admixes with West and South Scandinavians in Norway and Denmark, suggesting that the linguistic phase shift between Palaeo- and Proto-Germanic postdated the admixture event. The genetic structure that forms at this time persists until the end of the Roman Iron Age (1600 BP), encompassing the Palaeo- to Proto-Germanic periods as well as the subsequent period of linguistic disintegration into East, North and West Germanic. Thus, we reveal that by around 2000 BP, Germanic dialects unified a region divided by clear genetic borders that had existed for a thousand years. In the Polish Wielbark culture, we find support for the Late Antique Gothic origin myth of an exodus from Scandinavia. At the same time, however, we observe that the sampled southern European Goths carry little Eastern Scandinavian ancestry, suggesting a largely cultural adoption of Gothic language and identity for these individuals. Consistent with linguistic consensus on a Proto-Germanic homeland in northern Germany and Southern Scandinavia during the Early Iron Age, we find the descendant North and West Germanic languages across Europe to be closely linked to Southern Scandinavian ancestry, in support of historical documentation of the West Germanic Anglo-Saxons and Langobards. Finally, the linguistic changes leading to the formation of Old Norse appear to coincide with a back-migration of Southern Scandinavians to the Danish Isles and southern Sweden, but not in the more northern parts of Scandinavia, suggesting that a combination of demographic and cultural mechanisms drove the evolution and transmission of this language. We later find the dialectal division between Old West and Old East Norse to correspond to that of the Western Scandinavian Vikings and the Southern Scandinavian Vikings.

These findings underline the potential of IBD approaches to unravel genetic, archaeological and linguistic prehistories, at the same time revealing marked differences in the mechanisms behind the proliferation of cultural and biological features. With the increasingly fine-scale resolution of population genetics, additional sampling from along the Baltic coastline will allow several unanswered questions to be addressed: 1) confirmation of the proposed Bronze Age source of the Eastern Scandinavians along the Baltic coast; 2) identification of the Iron Age genetic boundary between East Scandinavian and Jutlandic Iron Age ancestry in continental Europe between Mecklenburg and Gdansk, potentially corresponding to the linguistic border between East and Northwest Germanic; 3) determining whether the spread of East Germanic dialects with Eastern Scandinavian ancestry extends beyond Poland; 4) determining the more localised regions both along the Eastern North Sea coast and within Britain representing each of the Angles, Saxons and Jutes; and finally 5) identifying the regions in Northeast Europe related to the source of Baltic- and Slavic-speaking populations.

## Methods

### Data Generation

All steps of generating the ancient genomes were undertaken using well established protocols within ancient DNA, as described in Supplementary Note S1. In brief, samples were inspected visually to identify suitable candidates. Teeth and petrous bone have previously been identified as the optimal skeletal material for recovery of ancient DNA and were hence selected where possible. All drilling, extractions and library builds were carried out in dedicated ancient DNA clean lab facilities at the University of Copenhagen, both manually and with automatisation. Double and single stranded libraries^89–91^ were built and sequenced on the Illumina Hiseq 4000 and the Novaseq 6000. For the purpose of authenticating ancient reads as human, non-USER treated libraries were built to allow for the detection of post-mortem damage^92^. Where possible, USER treated libraries were built from the authenticated extracts to minimise the effects of post-mortem damage on downstream analysis^93^. All reads were mapped to the human reference genomes (build hs37d5) using bwa aln (0.7.17)^94^, and duplicates were removed using picard MarkDuplicates (2.25.0). Contaminated libraries were identified using contamix^95^ and excluded. The final 712 samples included in this study ranged from 0.01X to 32.83X in autosomal coverage.

### Imputation

A set of 4,009 published ancient individuals with coverages suitable for imputation^28,37^ was curated, including whole genomes and SNP capture data. The published data were merged with the 712 newly generated genomes and imputed with GLIMPSE v.1.1.1^96^ (Supplementary Note S4). As off-target capture sites have been shown to impute poorly, only on-target sites were considered. In addition, sites that imputed poorly (imputation info score < 0.5) or did not pass the 1000 Genomes strict mappability mask were excluded. Individuals with coverage lower than recommended (0.1–0.5X for whole genome, 1.0X for captured samples^28,37^) and average genotype probability resulting from the imputation (<0.9) were also excluded. First and second degree relatives were identified using ngsRelate^97^ (Supplementary Note S5.6), and were then excluded from later analyses. This resulted in a final set of 4517 individuals (including 578 new) covering 697,179 SNPs.

### IBD based analyses

Segments of identity-by-descent (IBD) shared between all individuals in the final imputed set of 4517 individuals were detected using IBDseq^98^ (Supplementary Note S4). Potentially spurious IBD segments with low LOD scores or in hotspot regions were removed as described in Allentoft et al.^28^.

To cluster ancient samples, a network-based hierarchical clustering^99^ based on total shared IBD lengths was performed, as described in Allentoft et al.^28^. As short segments are less informative on recent ancestry, single segments less than 2 cM in length or total shared segment lengths of less than 5cM were excluded. Clusters were then curated as described in Allentoft et al.^28^. For the deeper clusters, individuals were plotted on a map of western Eurasia, showing clear correspondences through time and space. To better visualise IBD clusters and finer-scale patterns in the data, we carried out PCA analysis using GCTA (v1.94.1) showing the out-of-Africa and western Eurasian diversity and highlighted positions of individuals from the IBD clusters within PCA space. A second clustering run was performed containing only samples from the Bronze Age and earlier (2800+ BP), as later admixed individuals can bias clustering of the Bronze Age structure (Supplementary Note S5.2.2).

To understand the relationship between and diversity within the clustering, we performed IBD mixture modelling as described in Allentoft et al.^28^, allowing us to model target individuals as mixtures of source clusters. For every individual in the dataset, a palette is created, formed by the normalised proportion of total IBD lengths that each individual shares with all 386 clusters in the dataset. As such, the different ways in which the target and source individuals are related to non-source or target individuals are leveraged when modelling the target palette with the source palettes. Here, the single segment length cutoff was 1 cM and no total shared segment length cutoff was applied, as deeper relationships between samples were of more relevance than to clustering. We first replicated previous results for the formation of European populations before systematically including more proximal sources in a series of sets to reveal insight into the genetic structure of the ancient individuals. We then plotted the mixture modelling proportions on the PCA showing western Eurasian diversity to identify a series of novel admixture clines within the well established PCA structure (Supplementary Note S5.3.3). For some specific questions, auxiliary source sets were used (Supplementary Note S5.3.2.). To understand how the results of the IBD mixture modelling were reflected on a regional level, we applied spatio-temporal ordinary kriging ^39^, as detailed in Supplementary Note S5.3.8. Spatiotemporal variograms were fit via a metric covariance model using the R package gstat. We used two sets of parameters, one similar to the original publication for ancestries relevant to Europe more broadly (IBD mixture modelling sets 1-4), and a second allowing for the closer proximity between populations of relevance during the Iron Age (Set 5-7).

### Admixture dating

To assess the time of formation of admixed populations detected through the mixture modelling, DATES^58^ was run using default settings (Supplementary Note S5.5). A mean generation time of 25 years, and the midpoint of the mean radiocarbon ages of all dated individuals in the target population were used.

### Y-chromosome and mitochondrial genomes analyses

Genetic sex and the presence of karyotypes 47,XXY and 47,XYY were determined from the ratios of reads mapping to the autosomes, X, and Y chromosomes (Supplementary Note S5.4.3). For individuals with an XY karyotype, we used bcftools v1.17 mpileup to call genotypes for positions located in the single-copy short-read callable 10Mb region of the Y-chromosome, excluding indels, triallelic locus and variants with less than 95% frequency in the population of non-clonal reads in a locus. Haplogroup paths were called conservatively by matching ancestral and derived calls from ISOGG 2019-2020 in a root-to-tip direction (Supplementary Note S5.4.2). For reads that mapped to rCRS mitochondrial reference genome, a consensus was called and the haplogroups classified using Haplogrep 2.0^100^ for all samples with over 5X mitochondrial coverage. Sequences were then aligned with mafft, and a phylogenetic tree was generated using raxML (Supplementary Note S5.4.1).

### Dendrochronology

We used dendrochronological data based on tree-ring measurements from 654 wood samples of *Quercus* sp. from 42 sample sites in Denmark covering the years between AD 300 and AD 800 (Supplementary Note S6.2). From the dendrochronological data we constructed an average growth-curve for Denmark within this time interval by use of TSAP-Win^TM^ (version 4.82b2).

### Pollen data in southern Sweden

We used all available pollen data from the province of Scania to investigate the anthropogenic vegetation changes before the Viking age (Supplementary Note S6.2). The pollen records were grouped into four characteristic biogeographical regions, standardised to include the 87 most common pollen/plant taxa and the period 4000–900 BP. We applied a simplified indicator-species approach using eight pollen taxa from which changes through time are expected to be mainly due to land-use change. Pollen proportions for these pollen indicators were then summarised in 100-year intervals.

## Data availability

Sequence data were deposited in the ENA under accession: xxxxxxxx

## Methods References

References 89 - 100

## Acknowledgements

The Lundbeck Foundation GeoGenetics Centre is supported by grants from the Lundbeck Foundation (R302-2018-2155, R155-2013-16338), the Novo Nordisk Foundation (NNF18SA0035006), the Wellcome Trust (WT214300), Carlsberg Foundation (CF18-0024), the Danish National Research Foundation (DNRF94, DNRF174), the University of Copenhagen (KU2016 programme) and Ferring Pharmaceuticals A/S to EW.. GK was supported by Maritime encounters, Riksbankens Jubileumsfond, grant M21-0018. JVMM was supported by European Research Council (101078151) and VILLUM FONDEN (VIL53099). GS and SS were supported by Riksbankens Jubileumsfond, grant P22-0641 Italy before Rome. MDC, SH, and LK were supported by ‘Constructing the Limes: Employing citizen science to understand borders and border systems from the Roman period until today’ (NWA, 2021-2026, project number: NWA.1292.19.364). KM was supported by ERC 864358 (AMI). AW was supported by Emil Aaltonen Foundation. TW was supported by Lundbeck Foundation. KKr was supported by Riksbankens Jubileumsfond M16-0455:1, Rise II.

## Author contributions

EW initiated the study. HMC, EW led the study. HMC, GK, TW, RN, LH, KKr, MS, EW conceptualised the study. LV, TW, LO, LH, KKr, MS, EW supervised the research. GS, PBD, SS, AS, CPEZ, TW, RN, EW acquired funding for research. HMC, GS, PE, FD, MLSJ, SB, PBD, MEA, KGS, AF, TA, IAk, SLA, BA, PB, NC, PC, EC, TTC, ACp, SDD, SD, JD, KMF, OG, UBG, ZTG, JH, SH, MHe, VH, MHø, MHo, RJ, MDJ, JWJ, TJ, OTK, AK, RK, KKj, LK, KK, ACL, TL, ML, NL, YM, VYM, KM, VMa, ALM, IM, VMo, EMo, NM, EMu, BHN, DPK, GP, MSPdL, HR, WR, AS, BS, VSl, VSa, LS, OCU, HV, JV, DV, AVo, SW, HWe, AW, HWi, KW, JZ, BC, JPD, LO, RN were involved in sample collection. HMC, MLSJ, PBD, AF, EVB, PC, EC, RJ, TJ, RK, LK, KK, NL, VMa, VMo, EMo, EMu, BHN, DPK, GP, MSPdL, WR, VSa, AVo, SW, HWi, KW, JZ curated bioarchaeological data. HMC, FVS, GS, PBD, MEA, LV, CG, JC, JaS, JeS, FEY, ML, SBM, BSP, MSPdL, SW, TSK, LO were involved in data generation. HMC, GK, JVMM, FVS, TP, TV, SMS, PBD, ARa, IAl, RAH, EKIP, WB, AI, ARo, AVa, ML, CPEZ, TSK, BC, JPD, RN, MS were involved in data analysis. HMC, GK, PE, LH, KKr, EW drafted the main text. HMC, JVMM, FVS, TP, TV, SMS, PE, RF, MJG, HMEL, MFM, MLSJ, SB, ARa, KGS, SS, AF, IAk, SLA, EVB, EC, MDC, UBG, JZG, ZTG, SH, MHe, MHø, RJ, MDJ, JWJ, TJ, OTK, KKj, VK, LK, KK, ACL, ML, VYM, KM, VMa, ALM, VMo, EMo, NM, EMu, DPK, HR, WR, AS, BS, VSa, LS, GT, HV, JV, AVo, SW, HWe, AW, HWi, KW, JZ, TSK, LH drafted suppl notes and materials. HMC, GK, JVMM, FVS, GS, TP, TV, SMS, PE, RF, MJG, HMEL, MFM, FD, MLSJ, SB, MEA, LV, CG, ARa, IAl, RAH, EKIP, KGS, SS, AF, WB, AI, ARo, JC, FEY, SLA, MGB, JD, JZG, ZTG, VH, MHø, MHo, RJ, HJ, MDJ, JWJ, OTK, KKj, VK, LK, ML, NL, KM, ALM, NM, EMu, BHN, DPK, HR, AS, VSa, LS, GT, HV, SW, AW, KW, JZ, CPEZ, TSK, TW, LO, RN, LH, KKr, MS, EW involved in reviewing and editing drafts.

## Ethics declarations

The authors declare no competing interests

## Additional Information

Supplementary Information is available for this paper.

Reprints and permissions information is available at www.nature.com/reprints.

## Extended Data Figures

**Extended Data Fig. 1.**
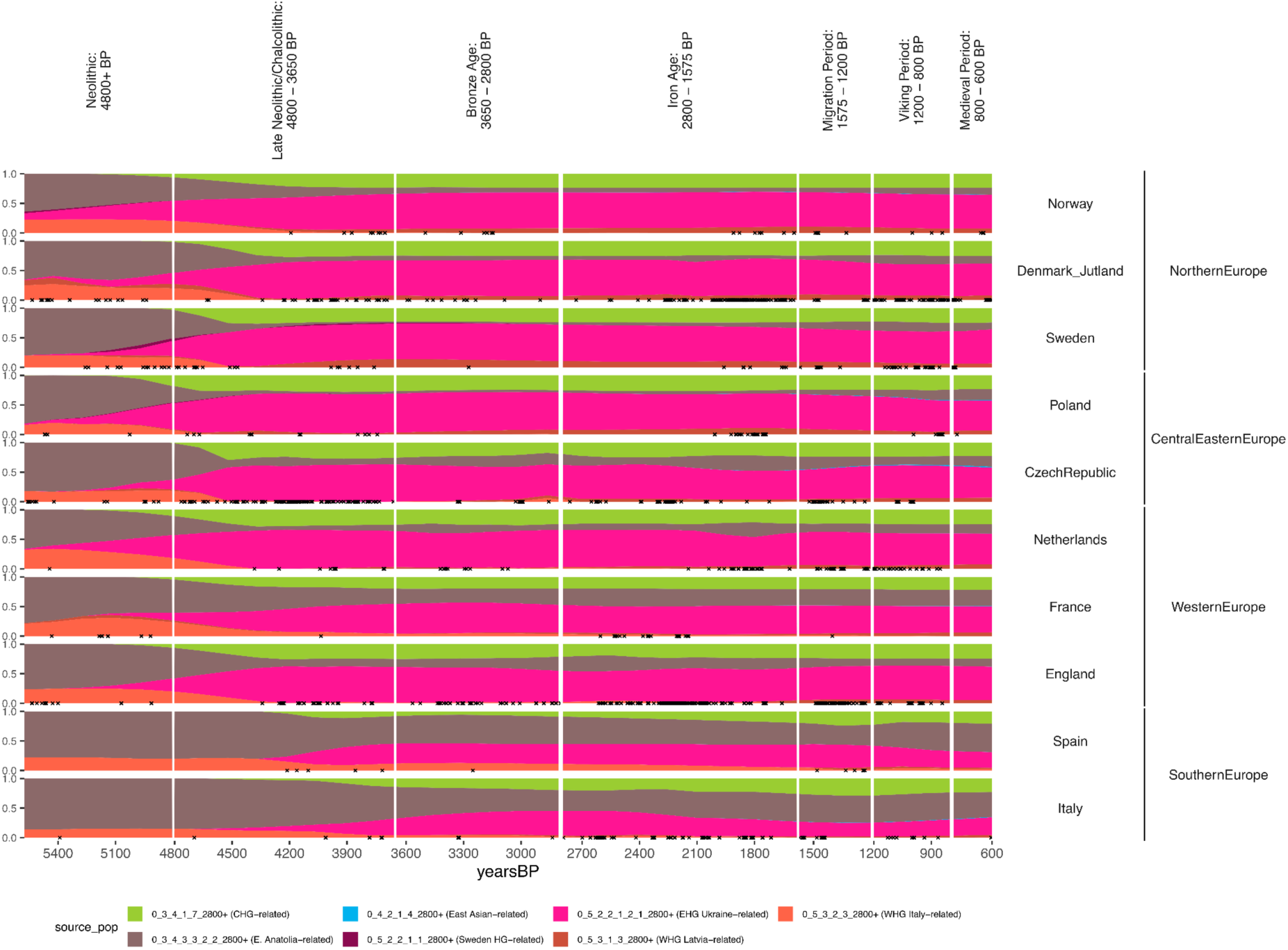
Spatiotemporal Kriging results showing the general trends of the proportion of ancestries using Hunter Gatherer and Early Neolithic Farming Sources (Set 1). The arrival of Steppe Ancestry, modelled as EHG (pink) and CHG (light green), onto a backdrop of Farming related ancestry, modelled as Early Anatolian Farming (brown) and Western Hunter-Gatherer (orange) is apparent. The ‘Europe-wide’ kriging parameters are applied here. Individuals within 250 km of the point used for each country (Supplementary Fig. S5.134) are indicated with an ‘x’. While these results represent the general trends, they do not capture the heterogeneity present in many regions, and hence should be interpreted together with the full mixture modelling results (Supplementary Fig. S5.18).

**Extended Data Fig. 2.**
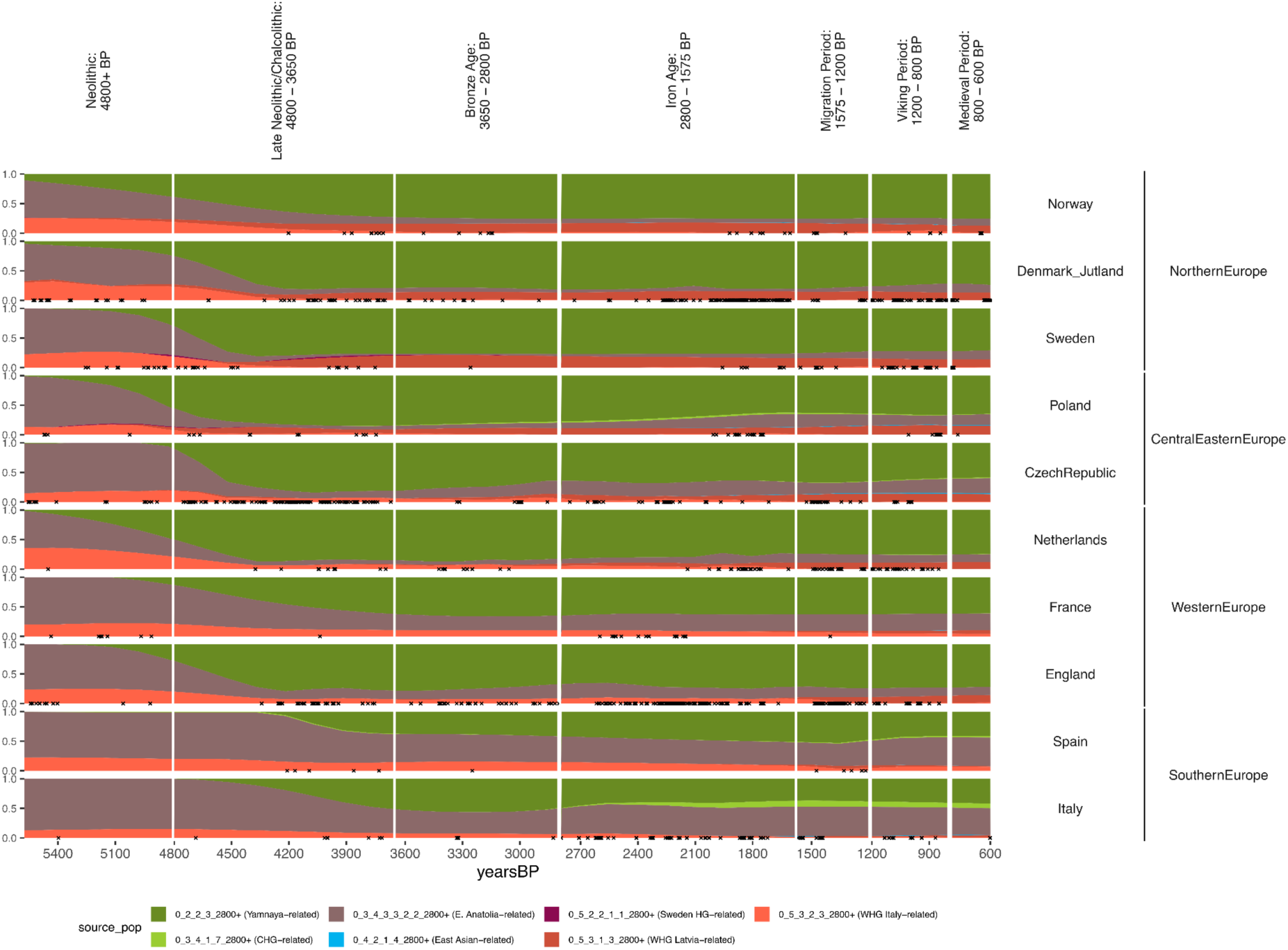
Spatiotemporal Kriging results showing the general trends of the proportion of ancestries when adding a ‘Yamnaya-related’ source (Set 2). Here, Steppe Ancestry that was previously modelled as EHG (pink) and CHG (light green), is now modelled directly (darker green). The ‘Europe-wide’ kriging parameters are applied here. Individuals within 250 km of the point used for each country (Supplementary Fig. S5.134) are indicated with an ‘x’. While these results represent the general trends, they do not capture the heterogeneity present in many regions, and hence should be interpreted together with the full mixture modelling results (Supplementary Fig. S5.18).

**Extended Data Fig. 3.**
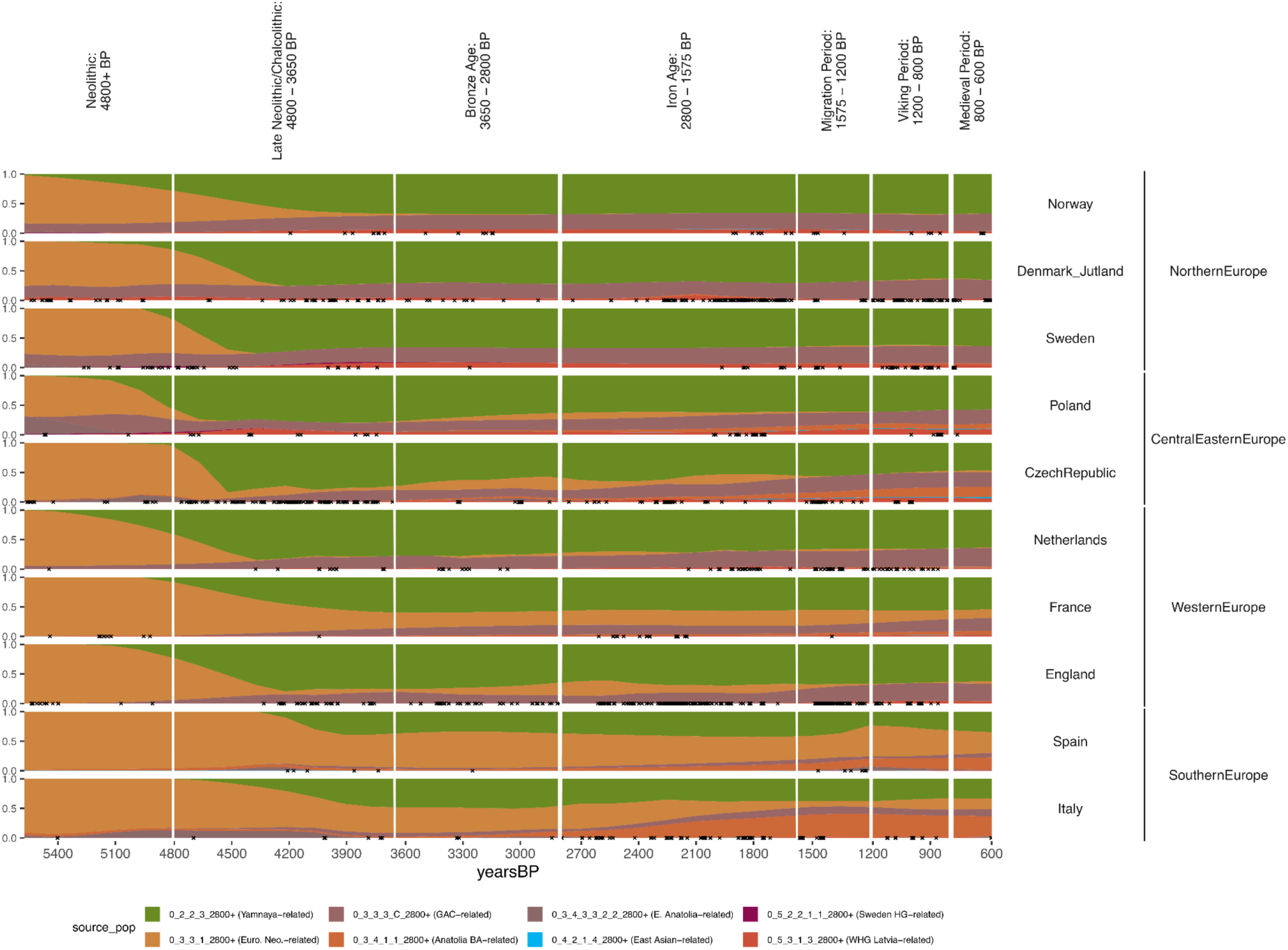
Spatiotemporal Kriging results showing the general trends of the proportion of ancestries of when including additional Farming-related sources (Set 3). To the north, Bronze Age populations are modelled only as Yamnaya and Globular Amphora, whereas in the south, some European Farming ancestry persists. The ‘Europe-wide’ kriging parameters are applied here. Individuals within 250 km of the point used for each country (Supplementary Fig. S5.134) are indicated with an ‘x’. While these results represent the general trends, they do not capture the heterogeneity present in many regions, and hence should be interpreted together with the full mixture modelling results (Supplementary Fig. S5.18).

**Extended Data Fig. 4.**
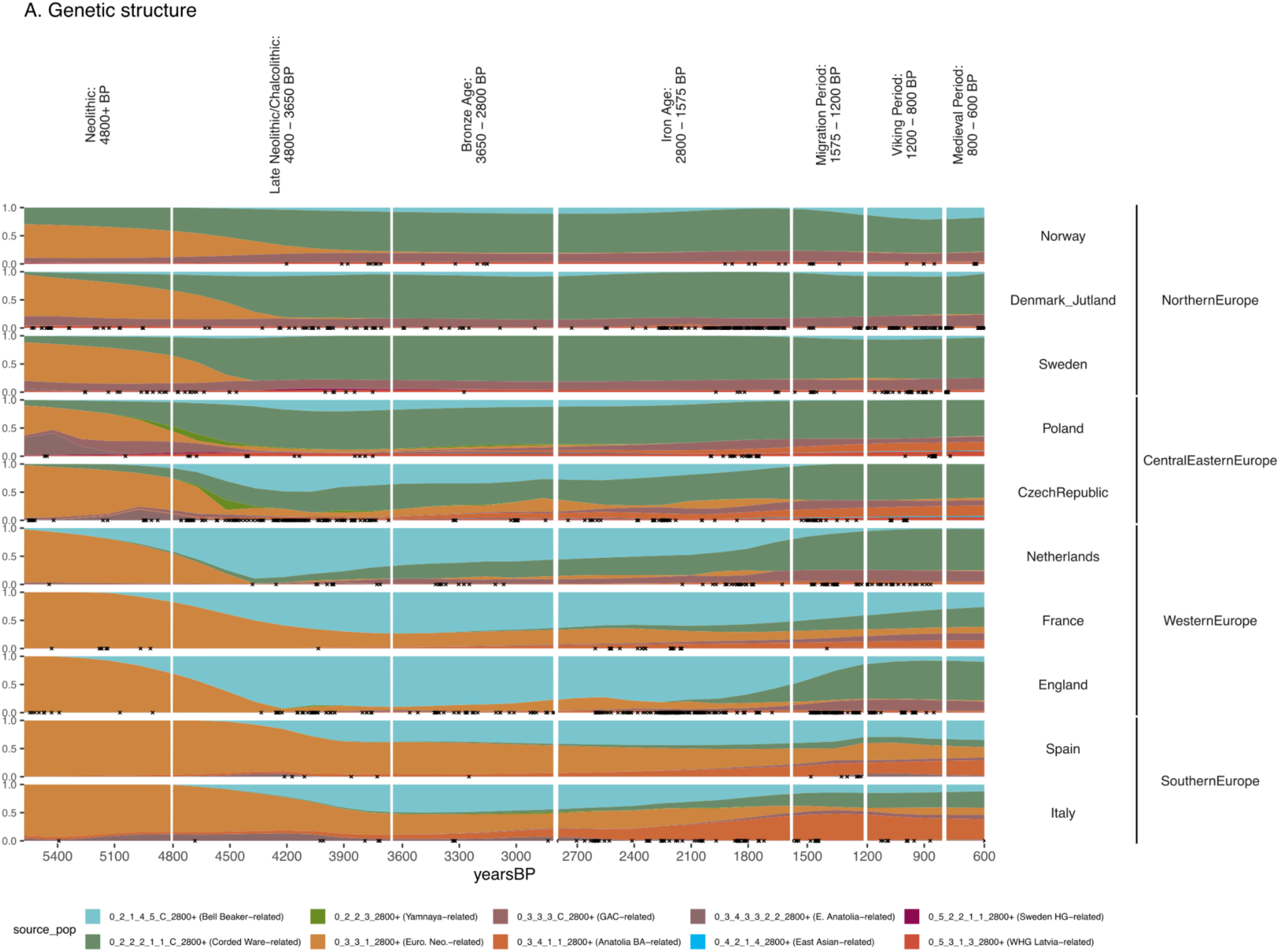
Spatiotemporal Kriging results showing the general trends of the proportion of ancestries when including additional Corded Ware and Bell Beaker-related sources (Set 4). Steppe Ancestry from the Bronze Age populations are modelled only as Corded Ware-related to the North, and Bell Beaker-related in the south. The expansion of Bronze Age Anatolian-related ancestry in Italy during the Iron Age, and to lesser degrees Czech Republic, France and England is apparent. The ‘Europe-wide’ kriging parameters are applied here. Individuals within 250 km of the point used for each country (Supplementary Fig. S5.134) are indicated with an ‘x’. While these results represent the general trends, they do not capture the heterogeneity present in many regions, and hence should be interpreted together with the full mixture modelling results (Supplementary Fig. S5.18).

**Extended Data Fig. 5.**
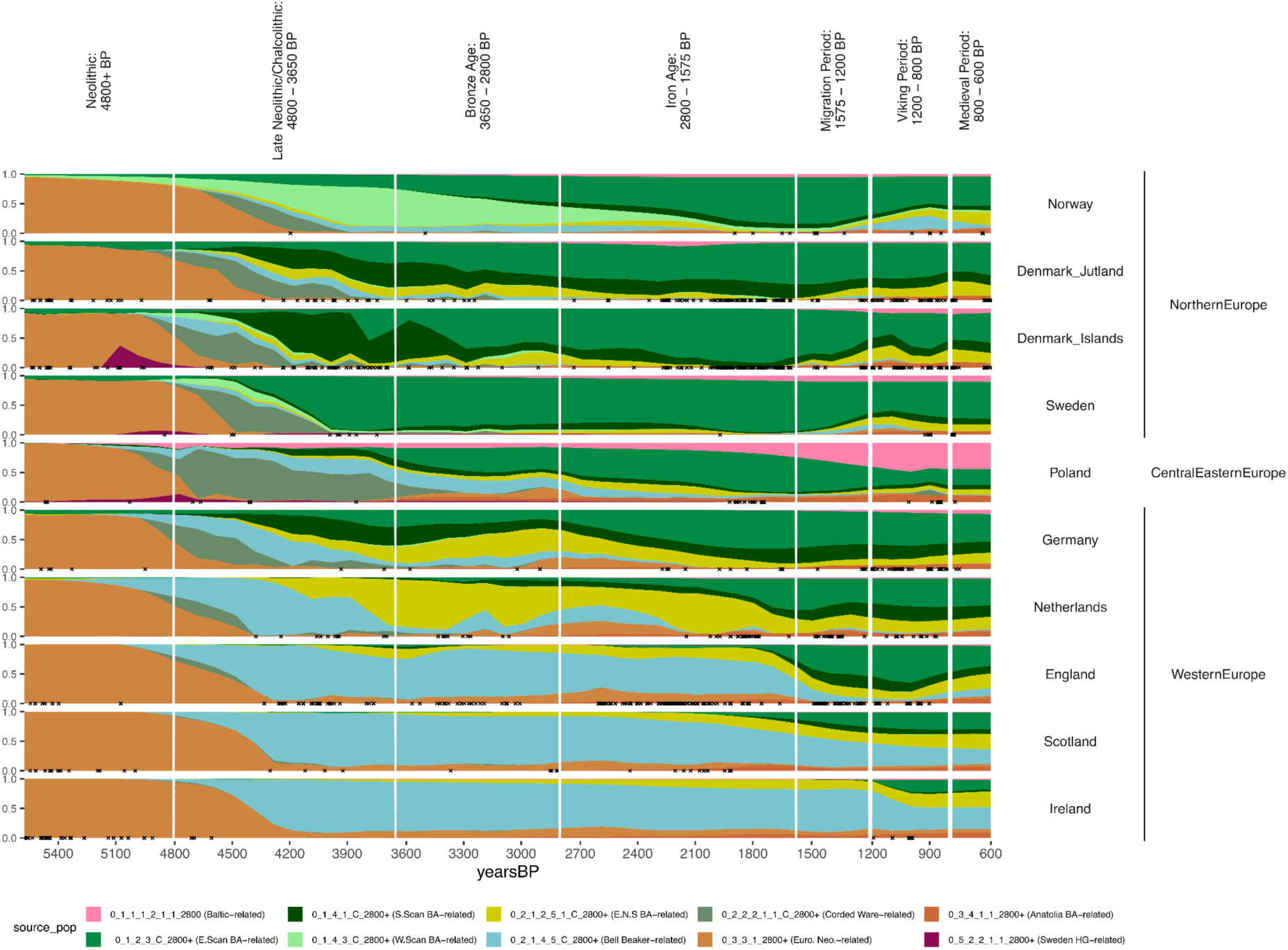
Spatiotemporal Kriging results showing the general trends of the proportion of ancestries when including additional Bronze Age Scandinavian and Baltic-related sources (Set 5). In the Late Neolithic / Chalcolithic, the regions of Northern Europe are distinct and modelled by various sources. Throughout the Bronze Age and Iron Age, Eastern Scandinavian ancestry becomes widespread, despite the persistence of some local Bronze Age ancestry. During the Migration Period, individuals from England and the Netherland experience an increase in Scandinavian ancestry reflecting Iron Age Jutland. Individuals from Denmark and Sweden experience and increase in East North Sea ancestry. Norway experiences an increase in Bell Beaker-related ancestry during the Viking Period. The transition in Poland from Iron Age (East Germanic) to Medieval (Slavic) coincides with the arrival of ‘Baltic BA’-related ancestry. The ‘North / West Europe’ kriging parameters are applied here. Individuals within 200 km of the point used for each country (Supplementary Fig. S5.134) are indicated with an ‘x’. While these results represent the general trends, they do not capture the heterogeneity present in many regions, and hence should be interpreted together with the full mixture modelling results (Supplementary Fig. S5.18).

**Extended Data Fig. 6.**
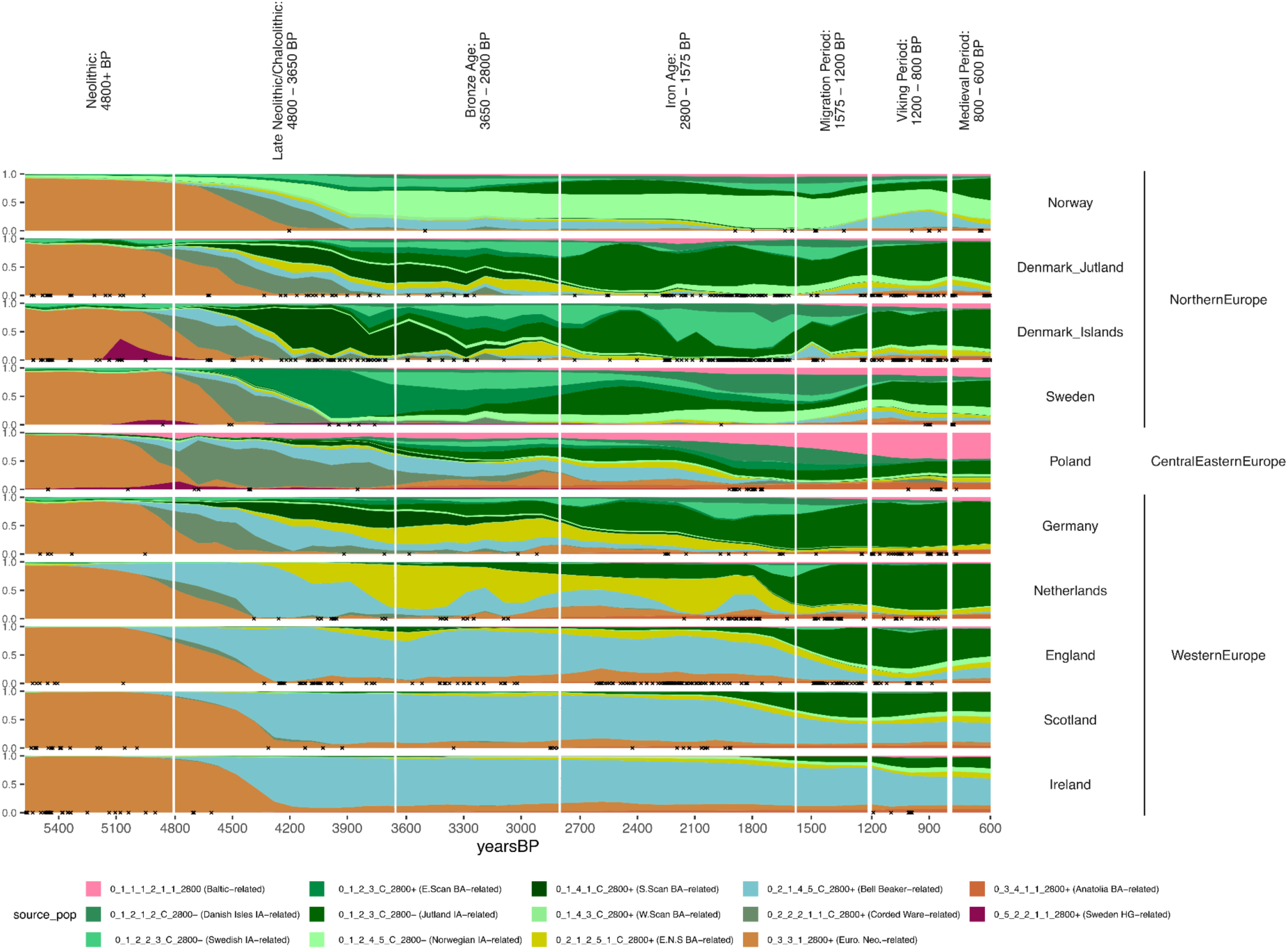
Spatiotemporal Kriging results showing the general trends of the proportion of ancestries when including Iron Age sources from Norway, Sweden, the Danish Isles and Jutland (Set 6). Here, the migration period shows an increase of ‘Jutland IA’-related ancestry in England, the Netherlands, the Danish Isles. The ‘North / West Europe’ kriging parameters are applied here. Individuals within 200 km of the point used for each country (Supplementary Fig. S5.134) are indicated with an ‘x’. While these results represent the general trends, they do not capture the heterogeneity present in many regions, and hence should be interpreted together with the full mixture modelling results (Supplementary Fig. S5.18).

**Extended Data Fig. 7.**
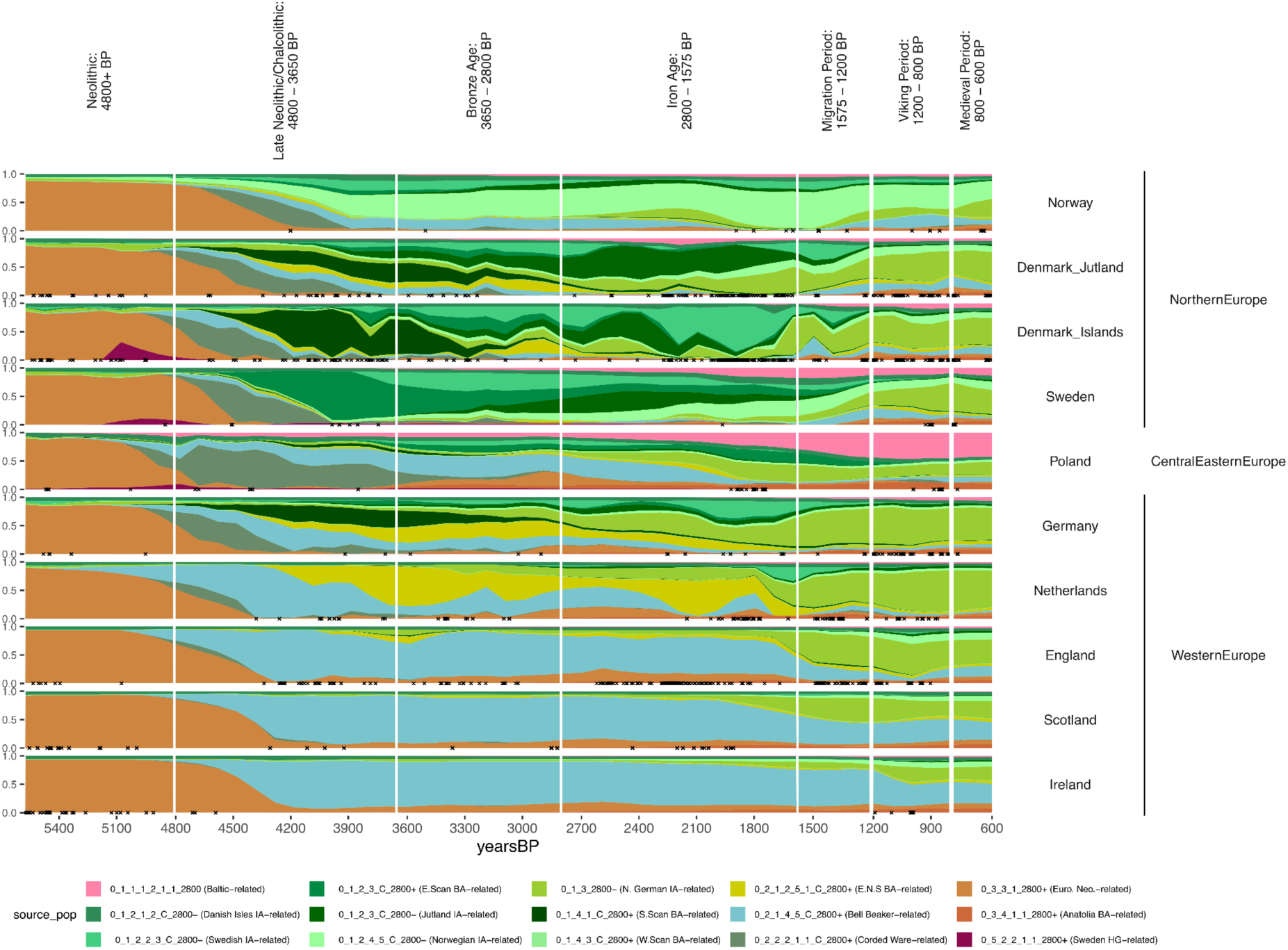
Spatiotemporal Kriging results showing the general trends of the proportion of ancestries when including an additional Iron Age source from Northern Jutland (Set 7). Here, we show migration period migrations transitions mentioned in Set 6 are better modelled by the Northern/Germany Southern Jutland source. In addition, Jutland sees a transition from local to Northern Germany/Southern Jutland ancestry. Individuals within 200 km of the point used for each country (Supplementary Fig. S5.134) are indicated with an ‘x’. While these results represent the general trends, they do not capture the heterogeneity present in many regions, and hence should be interpreted together with the full mixture modelling results (Supplementary Fig. S5.18).

**Extended Data Fig.-8.**
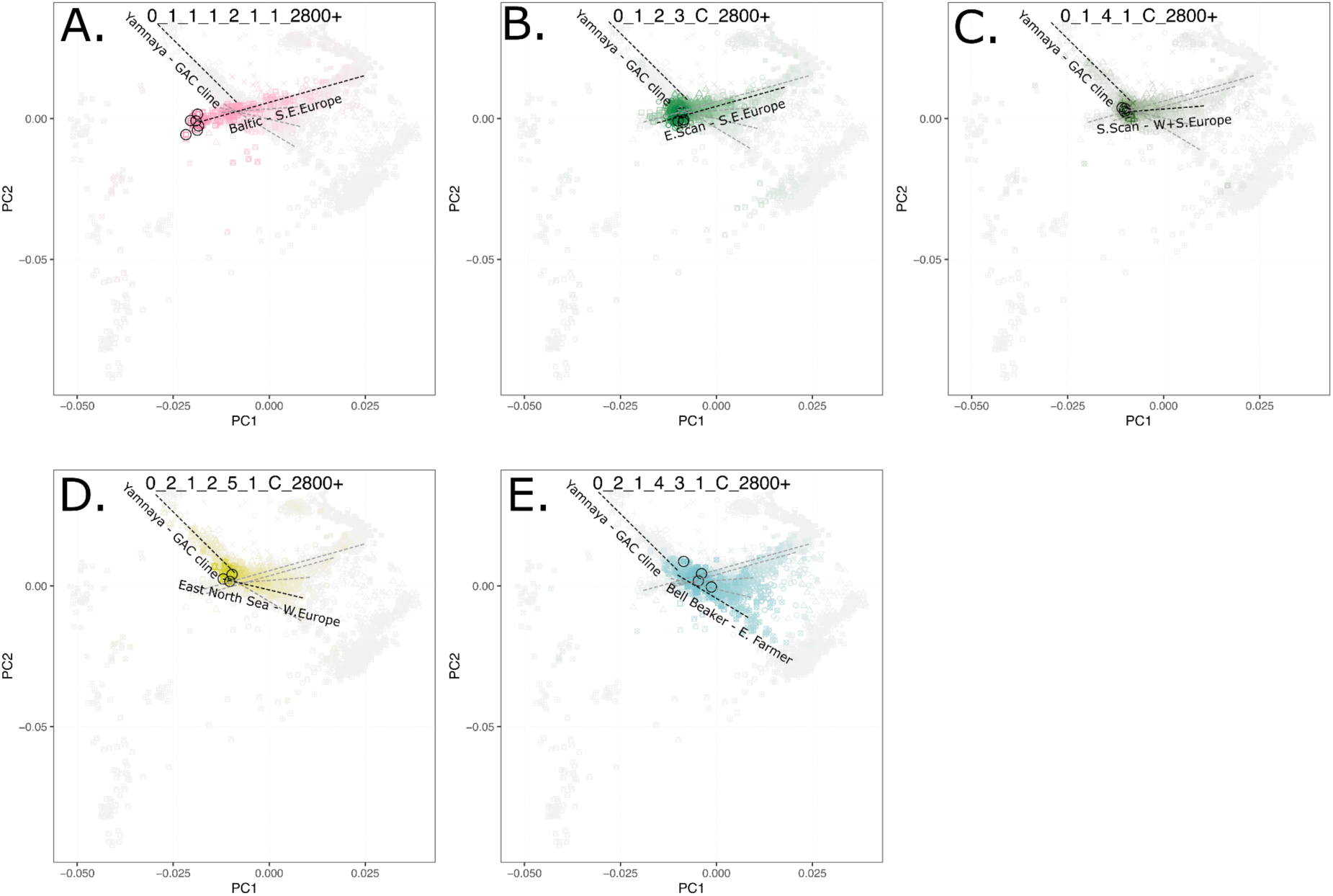
A subset of mixture modelling results from Set 5 displayed on the western Eurasian PCA, (Supplementary Note S5.3.1, Supplementary Fig. S5.20), showing mixture modelling proportions for a series of subclusters from the Corded Ware (North) (A, B, C) and Bell Beaker (D, E) clusters, revealing clines admixing with groups of varying European Farmer- and Bronze Age western Mediterranean-related ancestry. Source individuals are circled, and admixture proportions follow a cline from full colour (100%) to grey (0%).

**Extended Data Fig.-9.**
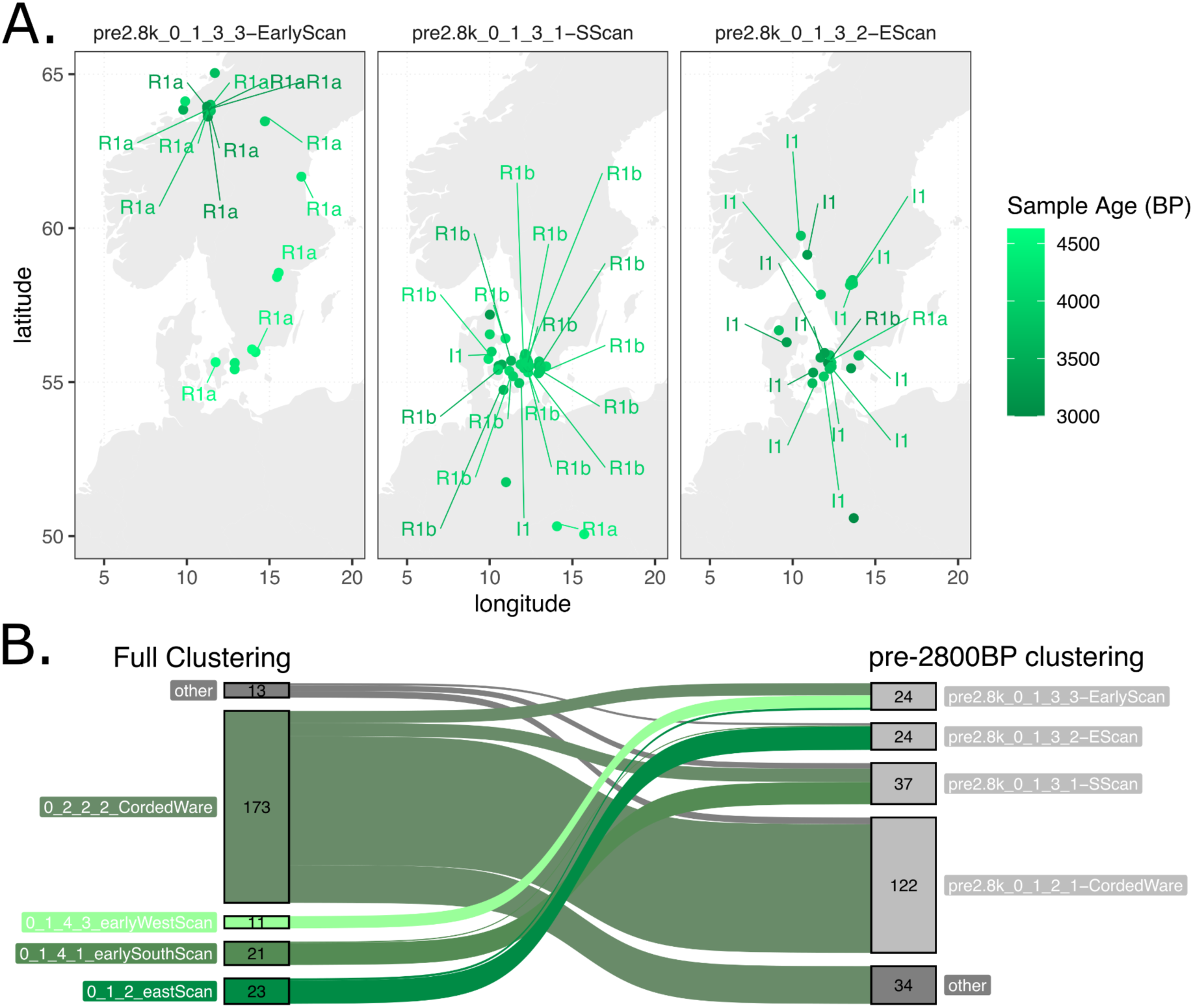
Scandinavian IBD clusters from pre-2800BP clustering. (A) Geographical distribution of individuals within the Scandinavian sub-clusters from the pre-2800 BP re-clustering. For males with sufficient coverage, major Y-haplogroups are noted. (B) Sankey diagram showing the correspondence between the three main Scandinavian clusters and the Eastern Corded Ware clusters in the Full and pre-2800 BP clustering.

**Extended Data Fig.-10.**
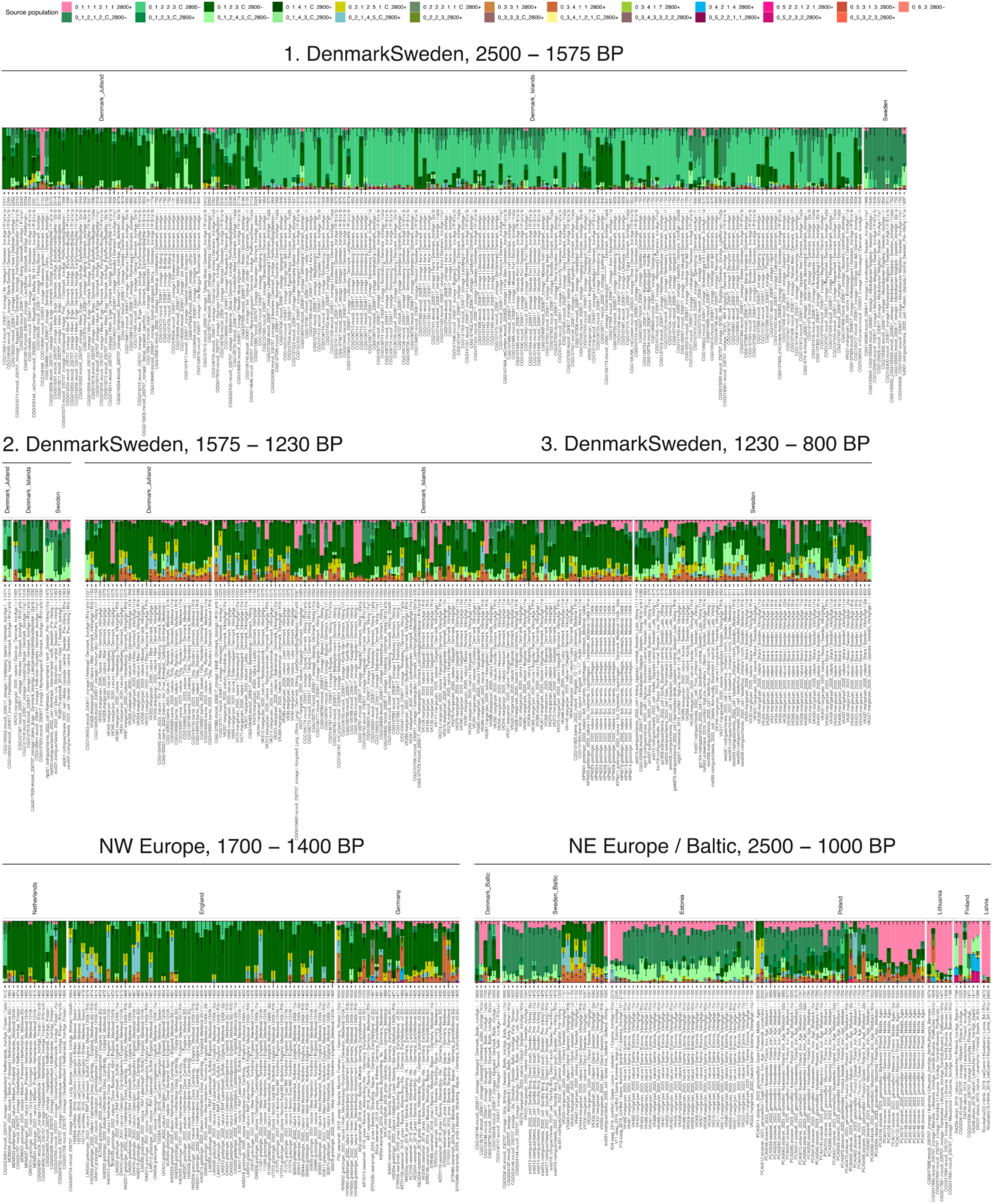
A northern European subset of IBD mixture modelling results for Iron Age sources (Set 6). Row 1 shows variation from Denmark_Jutland to the Islands of Denmark, to southern Sweden. Row 2 shows Denmark and Sweden during the Migration Period (1575–1200 BP, left) to the Viking Age (1200–800 BP, right). Row 3 shows the surrounding regions to the west (left) and east (right). Subset from Supplementary Note S5.3, Supplementary Fig. S5.18.

**Extended Data Fig.-11.**
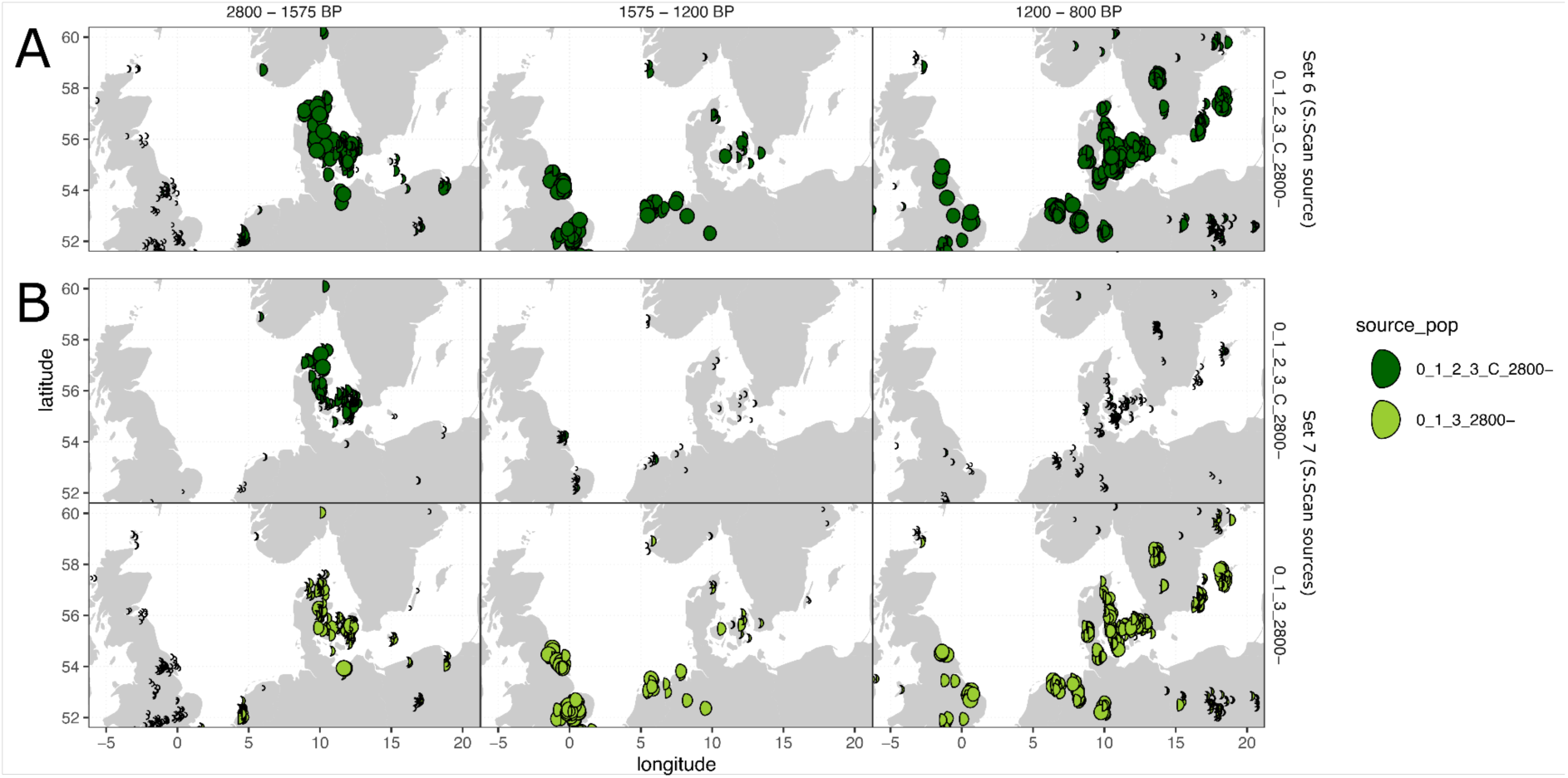
A subset of IBD mixture modelling results showing the proportion of Southern Scandinavian Iron Age ancestry for northern Europe with varying source sets. A) Set 6, which contains a single Southern Scandinavian Iron Age source (0_1_2_3_C_2800-, northern Jutland), in comparison to B) Set 7, with two Southern Scandinavian Iron Age sources (0_1_2_3_C_2800-, northern Jutland and 0_1_3_2800-Mecklenburg (northern Germany). The proportion of ancestry modelled is indicated by the proportion filled and the size of each circle. Full mixture modelling results for northern Europe are shown in Supplementary Fig. S5.25 and Supplementary Fig. S5.26

**Extended Data Fig.-12.**
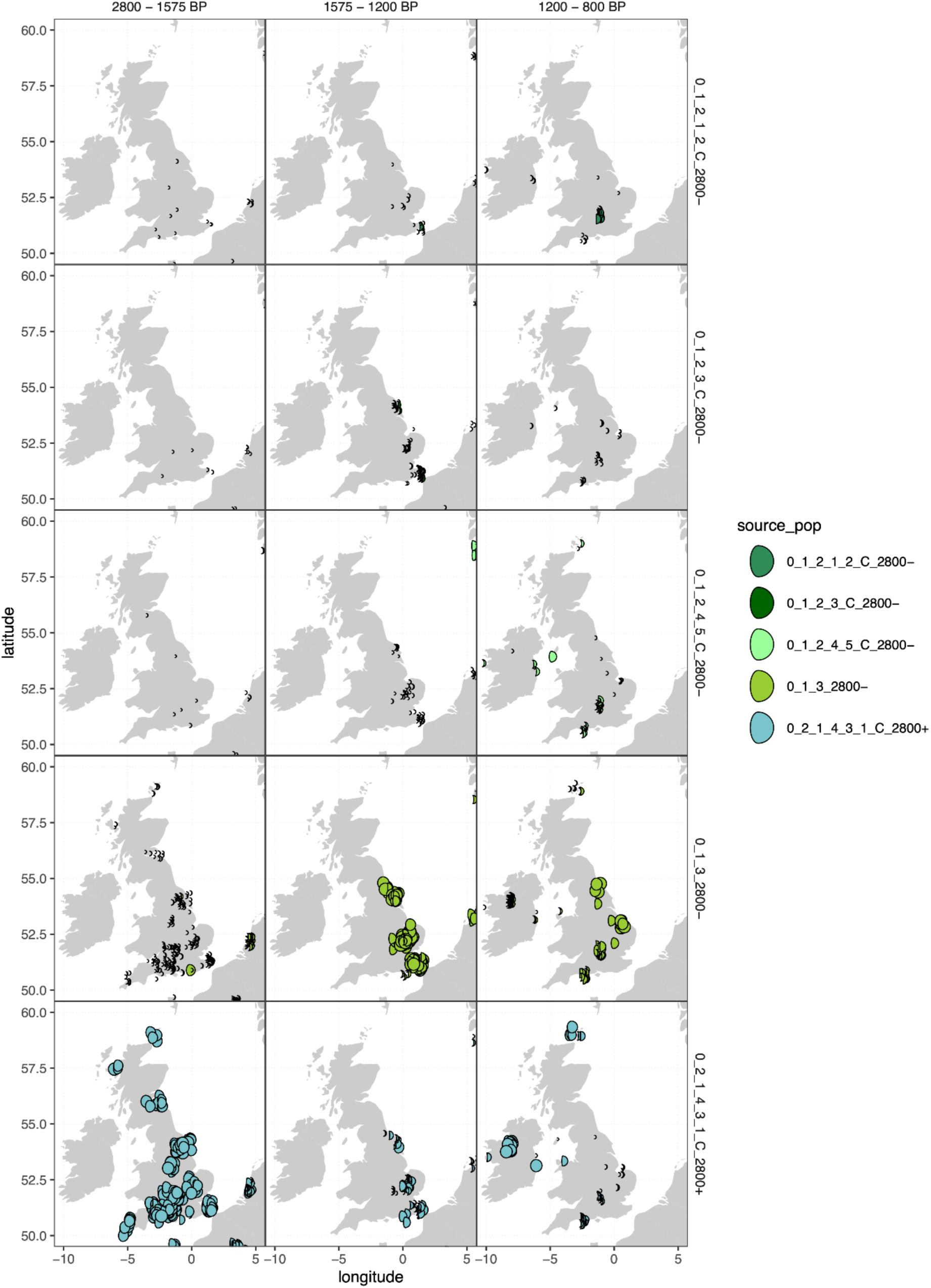
A subset of IBD mixture modelling results showing the proportion of ancestry in Britain and Ireland (Set 7). In column 1 (2800–1575 BP), the dominant ancestry modelled is 0_2_1_1_2 Celtic Bronze Age. In column 2 (1575–1200 BP) during the Anglo-Saxon period, a transition causing individuals to be modelled primarily as 0_1_3_2800-Southern Scandinavian Iron Age (Mecklenburg, northern Germany) has occurred, with small proportions of 0_1_2_3_C_2800-Southern Scandinavian Iron Age (northern Jutland, Denmark). In column 3 (1200–800 BP), the appearance of other Scandinavian ancestries (cluster 0_1_3_2_2_2 Eastern Scandinavian Iron Age (Sweden) and cluster 0_1_6_2 Western Scandinavian Iron Age (Norway)) is apparent during the Viking Age. Full mixture modelling results for northern Europe are shown in Fig. S5.26.

**Extended Data Fig.-13.**
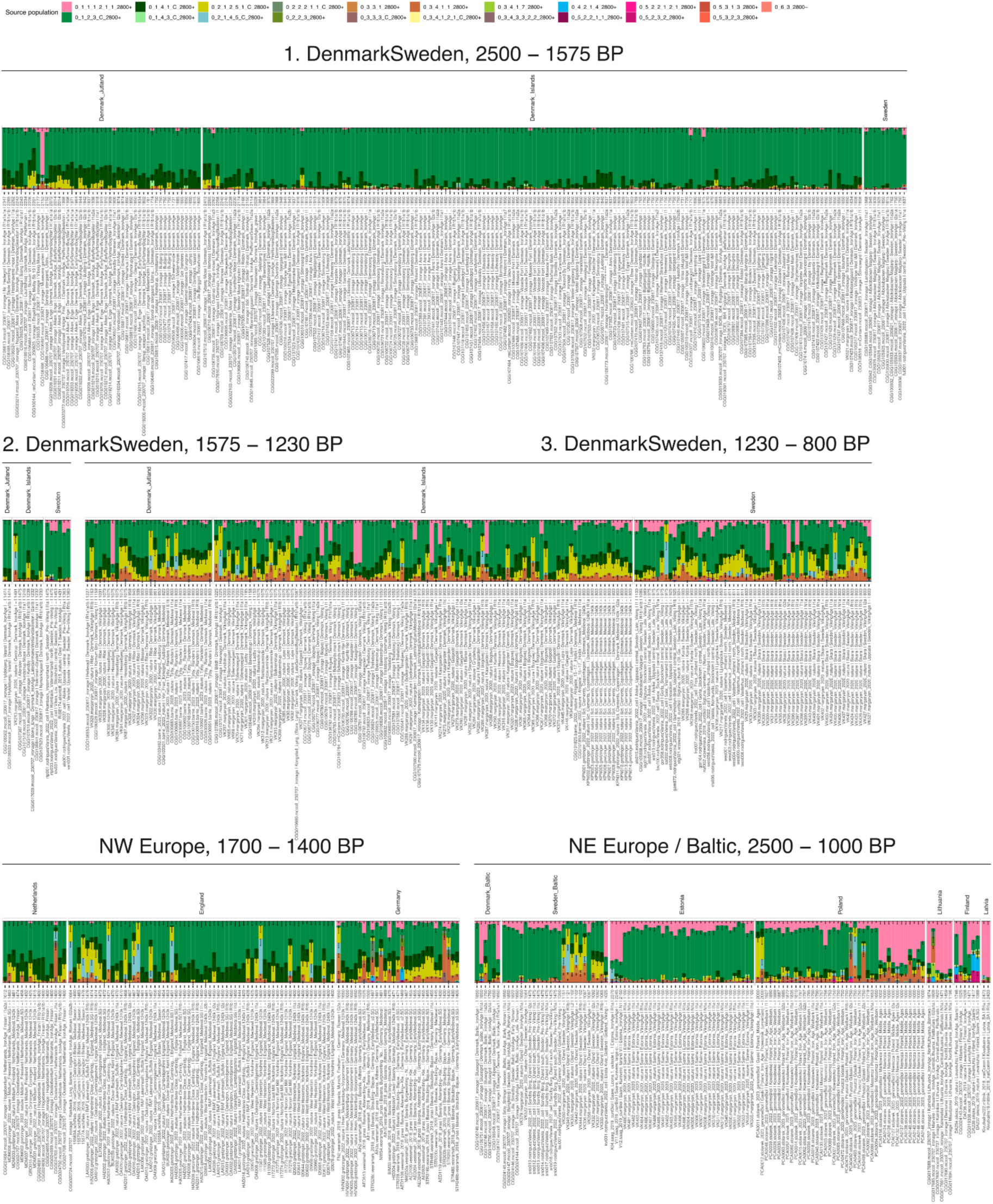
A northern European subset of IBD mixture modelling results for Bronze Age sources (Set 5). Row 1 shows the decreasing proportion of Southern Scandinavian ancestry from Denmark_Jutland to the Islands of Denmark, to southern Sweden. Row 2 shows Denmark and Sweden from the Migration Period (1575–1200 BP, left) to the Viking Age (1200–800 BP, right). Row 3 shows the surrounding regions to the west (left) and east (right). Subset from Supplementary Note S5.3, Fig. S5.18

**Extended Data Fig.-14.**
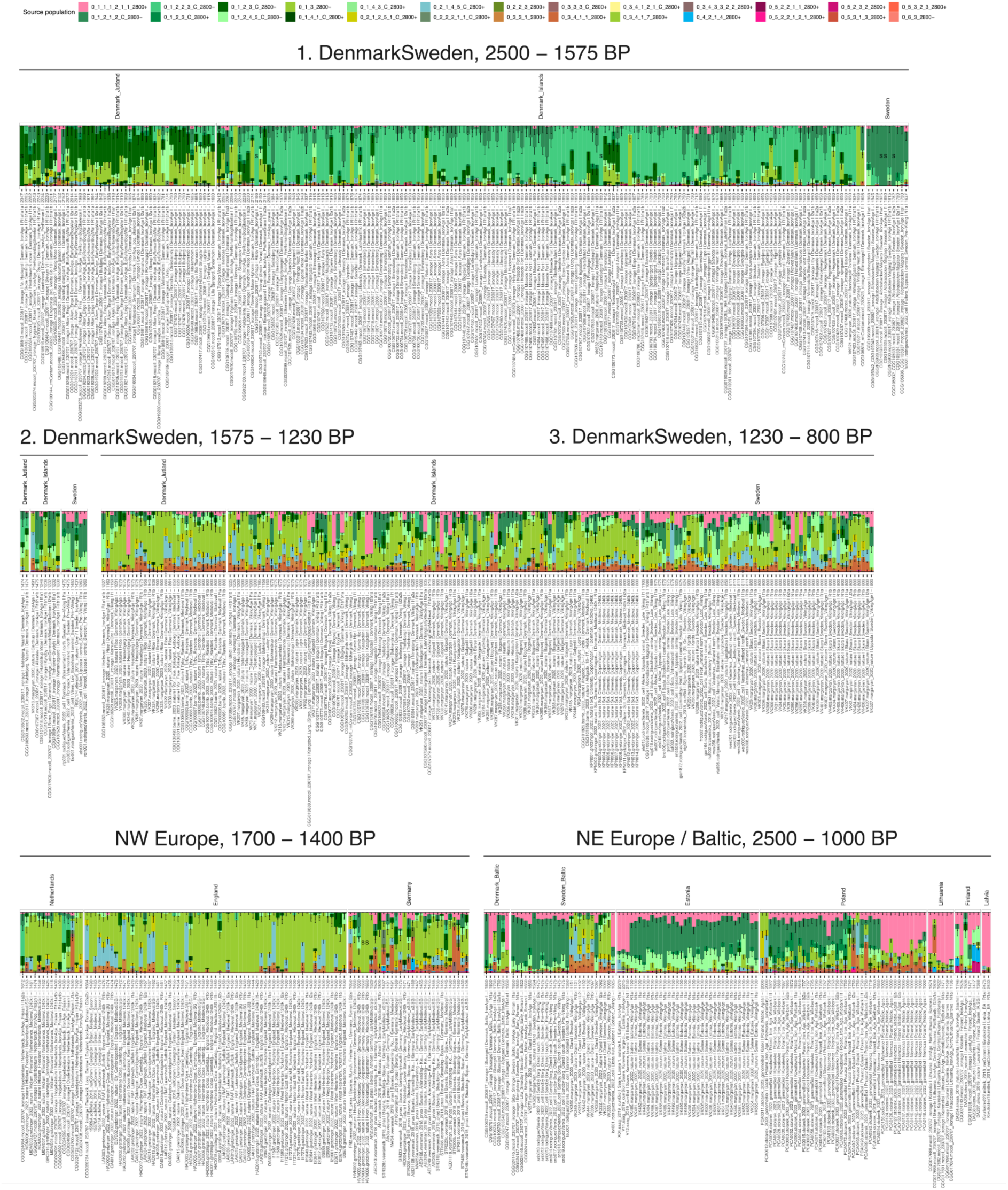
A northern European subset of IBD mixture modelling results for Iron Age sources, when including two Southern Scandinavian Iron Age sources (Set 7). Row 1 shows variation from Denmark_Jutland to the Islands of Denmark, to southern Sweden. Row 2 shows Denmark and Sweden during the Migration Period (1575–1200 BP, left) to the Viking Age (1200–800 BP, right). Row 3 shows the surrounding regions to the west (left) and east (right). Subset from Supplementary Note S5.3, Fig. S5.18.

**Extended Data Fig.-15.**
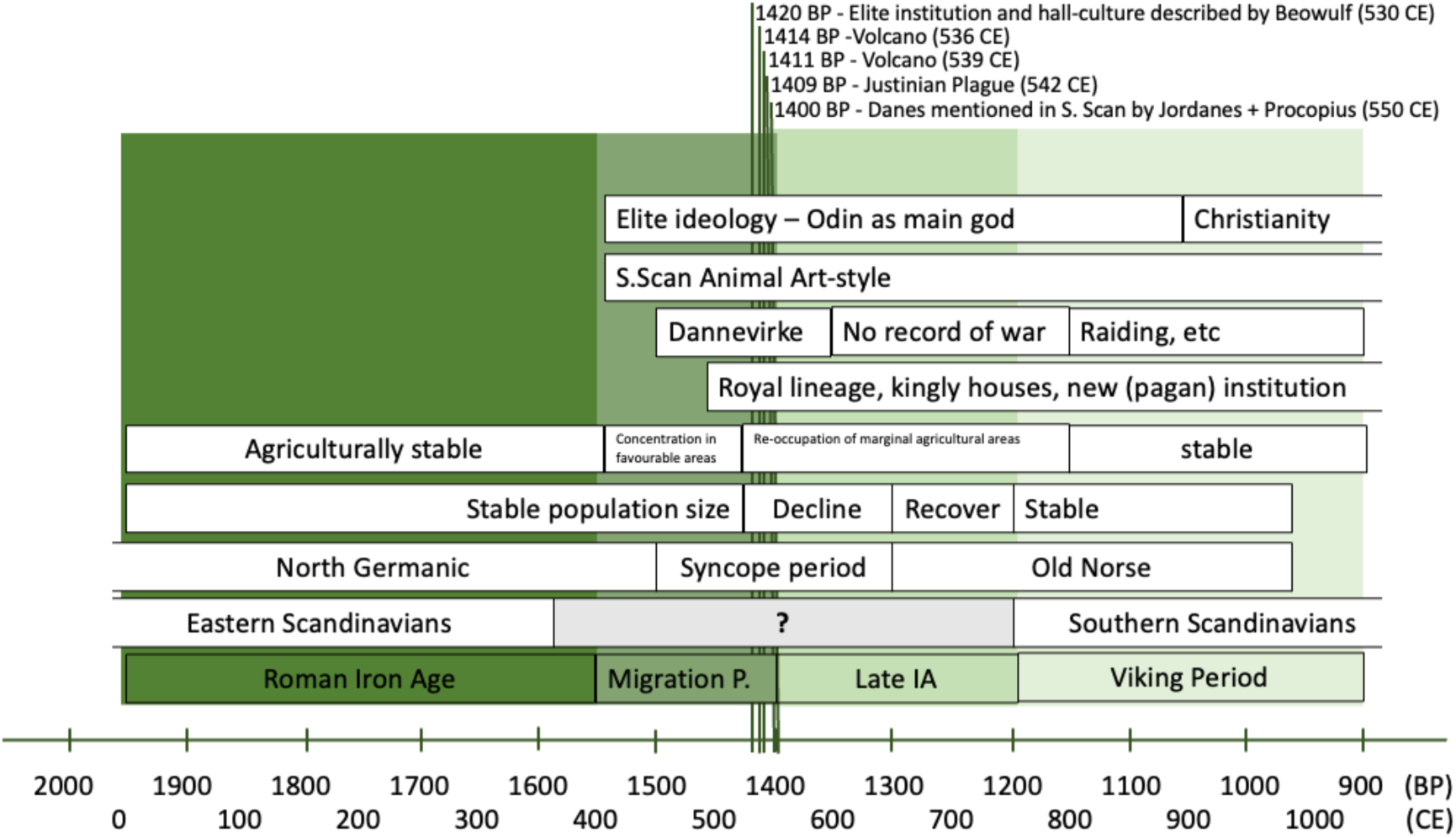
Timeline showing climatic events, and social, demographic, linguistic and genetic shifts in the Danish Isles and southern Sweden from the Migration Period to the Viking Age.

