## Supplementary Notes S7 for "Steppe Ancestry in Western Eurasia and the Spread of the Germanic Languages"

#### Archaeological supplementary material


S7.1. Archaeological and linguistic background to Scandinavian prehistory 2850 BC–750 AD. 2

S7.1.1. Introduction 2

S7.1.2. Middle and Late Neolithic 3

S7.1.3. Bronze Age 5

S7.1.4. Iron Age 8

S7.1.5. Glossary and Triangulation Table 13

S7.2. Site descriptions for Northern Europe 19

Denmark 19

Norway 161

Sweden 207

S7.3. Site Descriptions for Western Eurasia (excluding Northern Europe) 217

Austria 218

Bulgaria 221

Crimea 221

Czech Republic 225

Finland 228

France 230

Germany 245

Georgia 246

Hungary 247

Italy 249

Lithuania 250

The Netherlands 253

Portugal 262

Russia 264

Slovakia 274

Spain 277

UK 284
References 289

### S7.1. Archaeological and linguistic background to Scandinavian prehistory 2850 BC–750 AD.

Kristian Kristiansen, Lotte Hedeager, Guus Kroonen, Søren M. Sindbæk, Per Ethelberg, Sophie Bergerbrant, Karl-Goran Sjogren

#### S7.1.1. Introduction

We present a brief summary of the main archaeological and linguistic trends in Scandinavia from around 2850 BCE until the beginning of the Viking period 800 CE. In doing so we apply the following terminology and chronology as outlined in Table S7.1.1.

**Table S7.1.1:** Abbreviations of the periods used in the text. BP (before present) indicates years before 1950 CE as per convention in radiocarbon chronology.

| **Archaeological Period** | **Year (BCE / CE)** | **Years (BP)** |
|---|---|---|
| Early Neolithic | 4000 BCE - 3300 BCE | 5950 – 5250 BP |
| Middle Neolithic A | 3300 BCE - 2800 BCE | 5250 - 4750 BP |
| Middle Neolithic B | 2800 BCE - 2300 BCE | 4750 - 4250 BP |
| Late Neolithic | 2300 BCE - 1700 BCE | 4250 - 3650 B |
| Early Bronze Age | 1700 BCE - 1100 BCE | 3650 - 3050 BP |
| Late Bronze Age | 1100 BCE - 500 BCE | 3050 - 2450 BP |

| | | |
|---|---|---|
| Pre-Roman Iron Age | 500 BCE - 1 CE | 2450 - 1950 BP |
| Roman Iron Age | 1 - 375/400 CE | 1950-1575/1550 BP |
| Migration Period | 375/400 AD – 575 CE | 1575/1550 - 1375 BP |
| Germanic / Vendel / Merovinger Iron Age | 575 CE - 750/800 CE | 1375 - 1200/1150 BP |
| Viking Age | 750/800 CE – 1050 CE | 1200/1150 - 900 BP |

#### S7.1.2. Middle and Late Neolithic

Steppe-related migrations into northern Europe and Scandinavia had different origins, and thus also brought along different cultural and perhaps linguistic traditions. In Jutland and NW Germany/Holland, we find the Single Grave Culture (SGC) arriving 2850 BCE and quickly transforming landscapes into open grazing land and heathland[259]. Once they had consolidated, they started to expand towards North and East Jutland and the Danish islands 2600–2300 BCE. Here, they adopted cultural and religious influences from the surviving Neolithic Funnel Beaker Culture (FBC) population, creating a hybrid culture[260]. Contrarily, the Battle Axe Culture (BAC) in Sweden had a mixed background with an important Corded Ware (CW) contribution from Finland, and possibly from the SE Baltic[261,262]. In Finland, the CW arrives already by 2900 BCE[263]. In Sweden, there are only scattered finds related to BAC from before 2600 BC, whereas the period from 2600 to 2200 BCE has abundant finds. Thus, Swedish-Norwegian BAC emerged in the late Middle Neolithic B (MNB2). Based on radiocarbon dating of burials, BAC begins 2600 BCE and ends c. 2200 BCE[264]. This represents essentially a new beginning, bringing together elements from FBC, Finnish CW, and Central European Bell Beaker Culture (BBC) groups. This syncretic BAC product remained uniform throughout the period. In

Denmark, the period from around 2600–2500 BCE was a breakpoint too, with the momentous expansion of the SGC into northern Jutland and moderately so into the Danish islands[260,265]. The Steppe ancestry haplogroup (R1a) for the first time emerges manifestly in the genetic data[266].

Thus, we observe a related pattern of rapid primary migrations around 2850 BCE into Jutland in the west, and into Finland to the east. After consolidation and population build-up, a new wave of secondary migrations from 2600 BCE to 2300/2200 BC colonised the remaining parts of South Scandinavia, creating a hybrid material culture.

These different cultural and genetic traditions lived on, even if the Late Neolithic (LN) period saw a more unifying material culture linked to the mass production and distribution of high-quality flint daggers, imitating metal prototypes[267]. The LN was a period of intense demographic and agricultural expansion, witnessed in pollen diagrams as well as high ratios of non-local strontium values, especially in Sweden[268], less so in Denmark[269]. For Sweden, Blank et al[268] observed that 'Radiogenic ratios representing eastern and northeastern regions also appear on Late Neolithic Gotland. This might reflect a general trend of more northern and eastern influences in southern Sweden during this time. Based on pottery decorations and temper, intensified connections between middle Sweden and more eastern regions, such as Finland, has been proposed already in the last part of Middle Neolithic[261].

After 2000 BCE, South Scandinavia enters a more developed metal economy, even if according to current terminology the Bronze Age only started after 1700 BCE. In the period 2000–1700 BCE, we can observe several cultural innovations: from Sweden the introduction of gallery graves into the Danish islands, and parts of East Jutland. Also, Swedish house types, inspired by Únětice prototypes, were introduced[270]. Additionally, metalwork of Únětice origin or local imitations were produced and traded within the same broad region of East Scandinavia, while metalwork of West European, primarily English origin dominated in West Scandinavia[271,272].

#### S7.1.3. Bronze Age

During the first few centuries of the Scandinavian Bronze Age (1700–500 BCE), the old cultural divisions between East and West Scandinavia lived on, until the onset of the homogenising Nordic Bronze Age Culture from 1500 BCE[273] (chapter 5:3). In the period 1700–1500 BCE, leading up to the more unified Nordic Bronze Age, East Scandinavia was linked to trade and interaction with the Carpathian tell culture[274], while West Scandinavia had connections with NW Europe and the emerging Tumulus culture in South Germany [275] .

The Nordic Bronze Age period represented a climax of cultural innovation, wealth, and interaction both inside Scandinavia and with the outer world[275,276]. It is reflected in more diverse and frequent non-local strontium signatures in Denmark[277] and in Sweden[278], and extensive copper import from various areas; especially Wales and the Italian Alps during the Early Bronze Age, and Iberia and the Austrian Alps during the Late Bronze Age[279] [280]. It was also a period of continued northward expansion along the Norwegian coast, and the coasts on both sides of the Bothnian Bay in Sweden and Finland, resulting in the construction of thousands of cairns. They represent a new maritime economy[281,282]. The Late Bronze Age witnessed Nordic colonisation of the coastal region of Mecklenburg and Pomerania until river Vistula, resulting in a hybrid Nordic/local elite culture[276,283].

The cultural homogeneity represented by the thousand-year-long Nordic Bronze Age culture was a vehicle for the development of shared values and identities, most probably also a shared language, i.e Palaeo-Germanic. The presence of this language in the given archaeological setting is suggested by (1) loanwords from Palaeo-Germanic into Finnic and (possibly from there) into Saamic[284,285], and (2) loanwords from Celtic into Palaeo-Germanic[284,285]. The transmission of the first category of loans, which includes e.g. Fi. *hakea* 'to seek' from Palaeo-Germanic **sāgeja-*, predates the Germanic sounds shifts of the mid-first millennium BCE. It postdates the arrival of Finnic and Saamic in the East Baltic, which recent studies have correlated with the introduction of Siberian ancestry after the mid-second millennium BCE and no later than the mid-first millennium BCE[286]. To the south, the second category of loans, from (North) Celtic into Palaeo-Germanic, could have likewise been transmitted during the Late Bronze Age. These include **rīg-* 'ruler', **drūto-* 'jester', and **lēgjo-* 'healer, leech' from Celtic **rīg-*, **drūto-* and **le.a/egi-*, suggesting perhaps the adoption of certain societal innovations. The location of contact cannot easily be delimited, and might have occurred over land or over sea[287], [284].

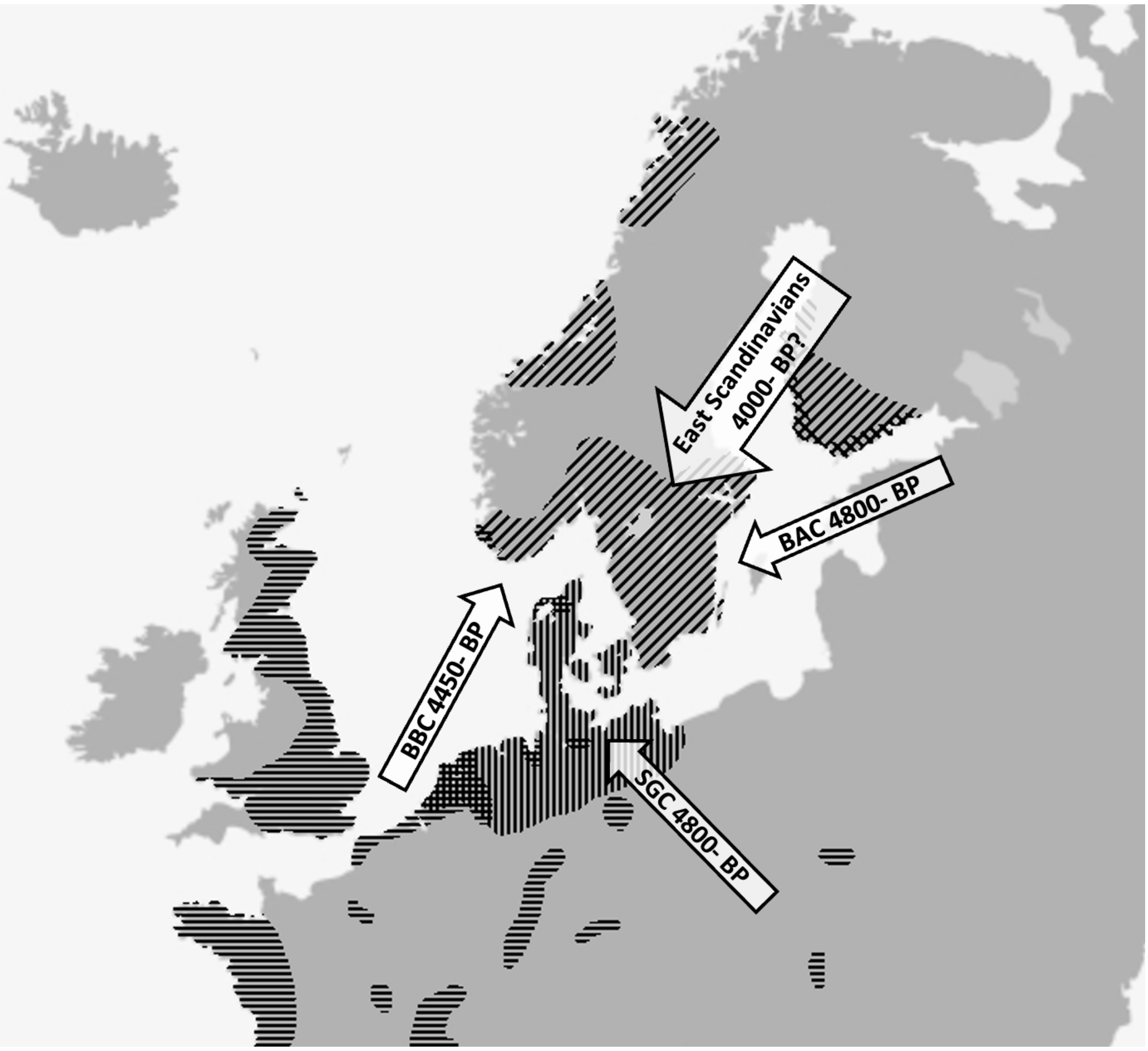


**Figure S.7.1.1. The four potential vectors for the arrival of Germanic to Scandinavia, Bell Beaker (BBC - horizontal lines), Single Grave Culture (SGC - vertical lines), Battle Axe Culture (BAC - diagonal lines rising), and 'East Scandinavians', potentially associated with the Finnish Kiukainen Culture (diagonal lines falling).**

**Table S7.1.2. A Table summarising the sources for Germanic described in the literature and the new 'East Scandinavian' source.**

| Suggested Sources | References | Timing | Associated Genetic cluster | Genetic support |
|---|---|---|---|---|

| | | | | |
|---|---|---|---|---|
| Bell Beaker | Lomborg 1973[289]; Pietro-Martinez 2008 [290]; Precott 2017[291] | By ~4600 BP | 'Bell Beaker' cluster | Very poor - Palaeo-Germanic and Proto-Germanic exist within the genetic 'Corded Ware' region |
| Corded Ware (Battle Axe) | Forssander 1933[292]; Hinsch 1956[293]; Price 2015[294] | By ~4800 BP | 'Western/Early Scandinavian' cluster | Poor - W.Scan ancestry decreases in prevalence around 3500 BP. Does not border Celtic region (loan words) |
| Corded Ware (Single Grave Culture) | Childe 1934[295]; Globe 1944[296]; Kristiansen 1984[297], Iversen & Kroonen 2017[298] | By ~4800 BP | 'Southern Scandinavian' cluster | Poor - S.Scan ancestry decreases in prevalence around 3500 BP. Does not border Finno-Saamic region (loan words) |
| New 'East Scandinavian' | | First seen genetically ~4000 BP, likely associated with start of the Late Neolithic (4200 BP) | 'East Scandinavian' cluster | Best - Distribution across Scandinavia at time and location of loan words |

#### S7.1.4. Iron Age

The northern European Early Iron Age (500 BCE–375 CE) and the Migration Period (375–575 CE) are characterised by a series of revolutionary transitions: the 'democratisation' of metallurgy through ease of access to iron[276], the rise and fall of the Western Roman Empire[299] and the subsequent 'Barbaric' invasions into and within Europe[300–302]. Linguistically, these periods coincide with the formation and spread of the Germanic languages. These languages, which include German, Dutch, Frisian, English as well as the Nordic languages, diverged from the common linguistic ancestor, Proto-Germanic, which is commonly assumed to have existed in Southern Scandinavia and Northern Germany throughout the Pre-Roman Iron Age[303,304]. From the end of the Pre-Roman Iron Age, Proto-Germanic disintegrated into East, North and West Germanic through a complex series of consecutive demographic processes, spreading across much of Europe[304–307].

When the East Germanic-speaking Goths emerged in Late Antique sources, they were settled in South East Europe and at the Black Sea during the 3rd century CE, becoming a major threat to the Romans[308,309]. However, they had previously been settled at the lower Vistula in the 1st century CE and then gradually expanded towards the southeast. Their own oral history records an exodus from Scandinavia to Poland across the Baltic Sea[310–312]. This scenario, which traditionally takes the East European Wielbark culture as a vector, is heavily disputed[313], even if there is agreement about close cultural connections between Eastern Sweden/Gotland and the Vistula region[314,315]. Following the Hunnic invasion c. 375 CE, some Pontic Goths entered the territories of the Roman Empire, contributing to its fall, and established a kingdom in Italy, and another in France and Iberia. In Italy, the Goths were succeeded by a West Germanic group, the Langobards, who similarly traced their origins back to Southern Denmark or Northern Germany[316,317]. Roman author Tacitus places them around the lower Elbe in the 1st century CE, spread south through Czechia and Hungary, and eventually established a kingdom in Italy from 568 to 774 CE.

During the Roman period, we can identify ongoing political/military competition between Jutes and Angles in South Jutland/North Schleswig in the form of territorial expansion and regression[318,319]. Both groups are characterised by specific settlement types/farms[320–323], and the territorial conflicts between them resulted in the construction of several defence walls, such as Olgerdike[324,325], and massive weapon sacrifices after battles during the period[326], such as Thorsbjerg[327–329], Nydam[330,331], Ejsbøl[332,333] and Illerup[334–336]. Conflicts came to a partial standstill after c. 400 CE with massive migrations to England. It left room for new political constellations to expand, such as the Danes[321], whose archaeological origin remains unclear. *Dania* was first mentioned around 490–500 CE as established north of the Frisians and the Saxons[337,338]. However, we observe a major change of material culture after 400 CE with the introduction of a new animal art style of Gothic/Hunnic inspiration[339] as well as other forms of elite culture such as gold bracteates with inspiration from Roman medallions and with occasional reference in runic writing to the main Norse god Odin[340]. This cosmological universe continued to the end of the Viking period[341]. A massive influx of gold accompanied these changes, most probably shared booty from participation in Gothic/Hunnic wars with the Roman Empire. However, the role of Hunnic influence on these major changes in South Scandinavia remains debated[312,342].

The Migration Period was a time of great upheaval in Western and Northern Europe. Germanic groups entered the formerly Roman provinces from across the Rhine and formed the Frankish, Vandal and other kingdoms[302]. The period also saw volcanic activity resulting in decades of globally decreased temperatures[343] and reduced plant growth in Scandinavia, a situation which may have contributed to the spread of the so-called Justinian plague and associated population collapse and recovery (Supplementary Note S7.3, S7.4). In parts of Britain, immigrating West Germanic Anglo-Saxons replaced local Romance and Celtic-speaking populations. In Scandinavia, runic inscriptions document the first use of Northwest Germanic dialects[344], which through a series of defining changes, evolved into Old Norse, the language spoken by Viking-Age Scandinavians from ~ 750 to 1150 CE[345].

Whereas the Migration Period was traditionally defined as a period of ‘folk migrations’ of Germanic and other tribes, recent scholarship is highly divided over the scale of these population movements as well as the authenticity of the origin stories of ‘Barbarian’ peoples [299,312,346–349]. While previous studies have shown that some Goths, Langobards and Anglo-Saxons carried northern European ancestry, confirming the specific origin within northern Europe has not been possible[350–354]. Thus, the specific northern European origins, as well as the potential genetic impact of these peoples on their regions of settlement, remain heavily disputed.

By the later Migration Period, a new group of large princely halls was introduced across Scandinavia[355,356]. Finally, we see possible evidence of a political shift in the construction of Danevirke in Southern Jutland, a south-facing moat and rampart earth stretching more than 5 km across the peninsular near Schleswig, whose second phase dates to around 500 CE[357]. Thus, the period between 400 and 550 CE in Scandinavia covers several potentially major population dynamics.

Within this period of transition, volcanic activity (536 CE) and the Justinian Plague (540 CE) meant that Scandinavia saw a population decline that did not fully recover until around 650 CE (Supplementary Note S7)[358]. Although the migrations and plague may have caused abandonment or depopulation of marginal subsistence areas[359], there was continuity to some degree in more fertile and southern areas[360], also related to intensified food production[361]. This is shown in the pollen data from southernmost Sweden, where woodland regeneration occurs in uplands, with continuity of agricultural production in the most favourable areas (Supplementary Note S7.4). In southern Norway, variation between different climatic zones is noted, with different societal impact from place to place which does not directly correspond with the climatic data[343]. A decline in population size at this time in Southern Norway has been attributed to the results of the volcanic events[362], however for those who survived, the subsequent improving conditions and relative abundance of resources due to a lower population size would have created the opportunity for rapid expansion, as attested to in historical sources in other areas. Despite these dramatic events, there is cultural and linguistic continuity from the beginning of the Migration Period until the end of the Viking period.

While some early medieval groups had clearly defined origin myths, there are conflicting texts and myths about the origin of the Danes. Writing in or shortly after 551 AD, the Roman author Jordanes states in his history of the Goths, *Getica*, (ch. 1.23)[362] that the *Dani* in *Scandza* were of the same stock as the *Suetidi* ('Swedes'). Another Roman author, Procopius (*De Bello* VI.15.1-4; 27-29)[363] claims that the *Heruli* "passed" the *Dani* on their way back to *Thule*. This could refer to almost anywhere between Albania and Scandinavia. The 12$^{th}$ century *Chronicon Lethrense*[364]invokes the Swedish legendary king Ypper, who ruled from Uppsala "in the time of King David". Ypper's sons Dan, Nor and Østen, it claims, became the rulers of Denmark, Norway and Sweden, which recalls the spread of BA East Scandinavians from Sweden to Norway and Denmark. According to Saxo Grammaticus' *Gesta Danorum*[365], completed around 1200, the Danes traced their origin from king Dan, while "the English race" originated from his brother, Angul. Meanwhile, the Icelandic Heimskringla by Snorri Sturlasson[366] holds that the *Aesir* gods travelled from the Black Sea through *Saxlandt* and from there to Funen and Sealand, perhaps recalling yet another origin myth. There are, then, rather conflicting memories, and no genuine migration myth as many other nations. The reason may be that the Danes – like the Franks – had formed as an agglomeration with no "core" ethnic group.

During the Viking period, the Old Norse language, as well as the linguistic area in which it was spoken, including Norway, Iceland and Sweden, was referred to as *dǫnsk tunga*, i.e. the Danish tongue[367]. The origin of this linguistic label is debated and it has been argued to reflect either the Pre-Viking Age political dominance of Denmark or the central position of Denmark as a gateway to Scandinavia[368],[369]. Both explanations find support in the genomic expansion of Southern Scandinavians to the Danish Isles and southern Sweden just prior to the Viking Age. From a cultural perspective, the label has been interpreted as expressing a sense of ethnolinguistic cohesion[370]. At any rate, it seems reasonable to assume that this identity arose in the south and became current across Scandinavia during the period leading up to the Viking Age. The rapid and relatively uniform transition from Proto-Norse to Old Norse across Scandinavia, marked by a set of defining sound-changes, may further support a model in which Old Norse spread as an elite dialect[371], driven by political dominance of its speakers in the south and cultural adoption in the north of Scandinavia. Of note, being part of the Old Norse language community resulted in privileged legal status with tangible implications in matters of justice and inheritance[372]. Intriguingly, the linguistic precursor of Old Norse ethnonym *danir*, often narrowly translated as 'Danes', was borrowed into Saami, cf. South Saami *daatje*, Lule Saami *dádtja* 'Norwegian or Swede', North Saami *dáža* ~ *dáčča*, Inari Saami *taažâ*, Skolt Saami *taaʒ̂ʒ̂* 'Norwegian'[373], the meanings of which further suggest that the term was in use as a pan-Scandinavian autonym, i.e. among Old Norse speakers living both in the north and the south, already before the Viking Age.

#### S7.1.5. Glossary and Triangulation Table

**IBD Clusters**

Clusters formed using the total shared length of IBD segments between all pairs, using a hierarchical clustering method. The deepest branches of the cluster are named 0_1_x, 0_2_x, 0_3_x etc. Subclusters within the first cluster (0_1_x) are named as 0_1_1_x, 0_1_2_x etc. Clusters are split into 2800+ and 2800- depending on the age of the individuals.

**IBD Sources**

A subset of individuals from specific subclusters. For example, the Yamnaya IBD Source individuals are 5 individuals associated with the Yamnaya culture, of the 12 individuals from within the 0_2_2_3_2800+ cluster. All individuals for each source come exclusively from one cluster, however not all individuals are included as sources.

**IBD Mixture Modelling**

A supervised ancestry modelling method in which target individuals are modelled from source individuals using non-negative least squares (NNLS). For each individual in the dataset, a palette is formed representing the proportion of IBD shared with all 386 clusters in the dataset. The palette of each target individual is then modelled as a linear combination of the palettes of the source individuals using NNLS: the result must be positive, and sum up to one. As a result, the shared IBD segments with all clusters is informative in the modelling, not just the segments shared between the target and source individuals. Mixture modelling using NNLS based method has previously been applied on 'allele-matching profiles'[374] and chromopainter chunks[375].

**Migrations (as inferred by IBD Mixture Modelling)**

In the genetic data we detect regional changes in ancestry based on changing patterns of IBD sharing reflected by the palettes. We infer that these changes represent the expansion of ancestry, which we refer to as migrations. In instances in which sampling over the transition period is sparse, like the Migration Period in Southern Scandinavia, it remains unclear whether the transition resulted from migrations of a large number of people over a short period of time, or gradual influx over hundreds of years.

**Yamnaya-related Cluster (0_2_2_3_x)**

An IBD sub-cluster containing 52 individuals, including 13/15 individuals archaeologically assigned to the Yamnaya culture. The 5 individuals used as the 'Yamnaya' source for IBD modelling are included in the 0_2_2_3_2800+ subcluster.

**Corded Ware (East)-related Cluster (0_2_2_2_x)**

An IBD sub-cluster containing 177 individuals, including 63/64 individuals archaeologically assigned to the Corded Ware (and Battle Axe) cultures. Few Bell-Beaker assigned individuals (n = 8) exist in this cluster. The 2 individuals used as the Corded Ware (East) source for IBD modelling are included in the 0_2_2_2_1_1_C_2800+ subcluster.

**Corded Ware (North)-related Cluster (0_1_x)**

An IBD subcluster containing 1297 individuals, found primarily across Northern Europe. The Steppe ancestry in these individuals is modelled primarily by the Corded Ware source. This cluster contains the Bronze Age sub-clusters (Baltic - 0_1_1), East Scandinavian (0_1_2), South Scandinavian (0_1_4_1_x) and West Scandinavian (0_1_4_3_x) and the clusters containing the Iron Age sources (see below)

**Bell Beaker Cluster-related Cluster (0_2_1_x)**

An IBD sub-cluster containing 737 individuals, including 81/107 individuals archaeologically assigned to the Bell Beaker. No Corded Ware assigned individuals exist in this cluster. The 6 individuals used as the 'Bell Beaker' source for IBD modelling are included in the 0_2_1_4_5_C_2800+ subcluster.

**West Scandinavian Bronze Age Cluster (0_1_4_3_2800+)**

An IBD cluster of individuals found exclusively within the bounds of Norway. Associated closely with the R1a y-haplogroup. The 5 individuals used as the West Scandinavian BA source for IBD modelling are included in this cluster.

**East Scandinavian Bronze Age Cluster (0_1_2_3_2800+)**

A IBD cluster of individuals first found in Sweden around 4,000 BP. Associated with the I1 y-haplogroup. The 4 individuals used as the East Scandinavian BA' source for IBD modelling are included in this cluster. Individuals within this cluster are modelled primarily with Steppe Ancestry, but are modelled with additional Latvian/Lithuanian HG ancestry, relative to the West and South Scandinavians.

**South Scandinavian Bronze Age Cluster (0_1_4_1_2800+)**

An IBD cluster found primarily in Denmark and the southern tip of Sweden. Associated with the R1b y-haplogroup. The 4 individuals used as the 'South Scandinavian BA' source for IBD modelling are included in this cluster.

**Norwegian Iron Age IBD Source**

Individuals from an IBD cluster (0_1_2_4_5_C_2800-) found primarily in Norway. Individuals within this cluster carry small proportions of Western Scandinavian BA ancestry, together with varying proportions of Eastern Scandinavian BA. The admixture event between the Western BA and Eastern Scandinavian BA ancestry likely occurred 3500-3000 BP.

**Swedish Iron Age IBD Source**

Individuals from an IBD cluster (0_1_2_1_2_C_2800-) found primarily in Sweden. Individuals within this cluster tend to be modelled exclusively as Eastern Scandinavian BA ancestry.

**Danish Isles Iron Age IBD Source**

Individuals from an IBD cluster (0_1_2_2_3_C_2800-) found primarily on the Danish Isles. Individuals within this cluster are modelled mainly by Eastern Scandinavian BA ancestry, with small amounts of local South Scandinavian BA ancestry. The admixture event between the local BA and Eastern Scandinavian ancestry likely occurred 3500-3000 BP.

**Jutlandic Iron Age IBD Source**

Individuals from an IBD cluster (0_1_2_3_C_2800-) found primarily on the Danish Isles. Individuals within this cluster are modelled similarly to the Danish Isles cluster individuals, but with a higher proportion of Southern Scandinavian BA ancestry. The admixture event between the local BA and Eastern Scandinavian ancestry likely occurred 3500-3000 BP.

**North Germany Iron Age IBD Source**

Individuals from an IBD cluster (0_1_2_3_C_2800-) found in Northern Germany. Individuals within this cluster are modelled similarly to the Jutland IA source individuals. The admixture event between the local BA and Eastern Scandinavian ancestry likely occurred 3500-3000 BP.

**Table S7.1.3 Triangulation Table**

| **Time Period** | **Archaeological** | **Genetic** | **Linguistic** |
|---|---|---|---|
| 5250 – 4750 BP<br>3300 - 2800 BCE | Middle Neolithic A<br>Funnel Beaker Culture | Presence of Farming-related ancestry | Pre-Indo European Languages |
| 4750 – 4250 BP<br>2800 - 2300 BCE | Middle Neolithic B | Arrival of Steppe (Corded Ware-related) ancestry | Arrival of Indo-European languages |

| | | | |
|---|---|---|---|
| 4250 – 3650 BP | Late Neolithic<br>Cultural Borders between Jutland/NW Germany and East Denmark/Sweden. | Genetic structure present (West Scandinavian in Norway, East Scandinavian in Sweden, East and South Scandinavian in Denmark | |
| 3650 – 2540 BP | Nordic Bronze Age<br>Borders dissolve, homogenisation of cultures across Scandinavia | East Scandinavian ancestry now found widespread across Scandinavia, varying degrees of admixture | Insight into geographical distribution based on Celtic loan words incorporated into Palaeo-Germanic, Palaeo-Germanic loan words incorporated into Finno-Saamic. These loans predating the so-called Germanic sound shifts, which had happened by the beginning of the Iron Age |
| 2450-1950 BP | Pre-Roman Iron Age. Regionalization and cultural borders return | Genetic borders persist between Sweden (E.Scan) and Denmark (E.Scan+S.Scan) and Norway (W.Scan+E.Scan) | Proto-Germanic likely in Southern Scandinavia |
| 1950-1575/1550 BP | Roman Iron Age. Strong trade with Roman Empire luxury products, and social/political interaction | Genetic borders persist between Sweden (E.Scan) and Denmark (E.Scan+S.Scan) and Norway (W.Scan+E.Scan) | Runes found in Denmark, Norway and Sweden (crossing genetic borders).<br>Proto-Germanic gradually splits into East, North and West Germanic. |
| 1575/1550-1375 BP | Migration Period.End of Roman influence. Major cultural changes, and formation of new elite animal style | Limited sampling in Scandinavia.<br>Transition from Bell-Beaker-related to S. Scan in Britain | Transition from Proto- to Old Norse begins.<br>Arrival of West Germanic to Britain. |

| | | | |
|---|---|---|---|
| 1375-1200/1150 BP | Germ./Vendel/Mer.Period. Consolidation of changes introduced during the Migration Period. First royal centres | Limited genomes due to cremation | Transition from Proto- to Old Norse complete |
| 1200/1150-900 BP | Viking Period. Continuation and expansion of previous period, external colonisation | Genetic border between Denmark and Sweden dissolved. Norwegian border persists. | Period of Old Norse as a common language of Viking Age Scandinavians. |
| 900- | Middle Ages. Christianity, formation of nation states | | Rise of East and West Norse dialectal features, start of the fragmentation of Old Norse into its descendant languages. |

### S7.2. Site descriptions for Northern Europe

The following sections contain the archaeological descriptions of the sites in Northern Europe for the new samples in this study. Each site description is followed by the inventory number of the samples concerned (CGG inventory number (GeoGenetics inventory number), Museum inventory number). When available, the $^{14}$C dating, isotope data and the sex identification of the samples are given.

#### Denmark

#### Denmark, Bornholm, Bornholms Sønder, Pedersker parish, Slusegård

Coordinates: 54.9997, 14.9985

Sofie Laurine Albris

Slusegård is the largest Early Iron Age burial site on the Island of Bornholm in the Baltic Sea. It is situated near the mouth of the stream Øle Å, stretching along each side of an old hollow road. The cemetery lies close to the south coast of Bornholm[376–381].

On the surface, some burials have been marked by boulders, stone settings and trenches. The burial site was subject to smaller investigations in the 19th century, while large-scale excavations of over 6000m$^2$ were carried out by Ole Klindt-Jensen during the years 1958 to 1964[381]. Over some parts of the site a find-rich burnt layer was observed, and all was covered by sand drift.

Slusegård has around 1400 excavated graves. There are both wealthy graves, some with Roman imports and burials with no grave goods. Most graves were cremations (928), while there were also some inhumation graves (467), placed both in supine and crouched positions [376,380,381]. At least 43 of the inhumations were boat-graves, where boats and boat parts were used as coffins[376]. The boat graves contained both females and males, where the latter were the wealthiest, some containing weapons. Finds of boat graves indicate a network of connections from northern Poland to Jutland in Denmark with Bornholm in the middle[378].

At Slusegård around 110 graves from the 2nd century showed signs of ritual plundering[377]. Many skeletons from Slusegård show indications of violence, with decapitations or broken limbs. One of the graves sampled in this project had an iron point stuck in the skull. In all, five graves, two from the Early Roman Iron, one from the Late Roman Iron Age and two just dated to the Roman Iron Age were sampled and yielded results within this project. The samples come from both rich graves and sparsely equipped graves.

The burial site is dated to a 500-year period from around 50 BC into the 5th century CE[379]. The inhumation graves mainly date to the period between the middle of the 1st to the late 3rd century. The oldest boat graves were from the 2nd century, and boat burials were reaching into the 3rd century CE (B2-C2). The samples included in this study are listed in Table S7.2.1.

| **CGG No** | **Grave** | **Sample description** | **Grave description** | **Period** |
|---|---|---|---|---|
| CGG_2_106751 | Garve 129 | Male adult, placed in supine position, head in N | Iron knife, and two pots plus and iron point embedded in the cranium | Early Roman Iron Age |
| CGG_2_106752 | Grav 2 | Male? Adult placed in half a boat | Iron brooch, iron knife, bone com, four pots, textile and hide fragments | Late Roman Iron Age |
| CGG_2_106747 | Grave 198 | Female mature, crouched on the right side, head in N | Bone comb, iron knife, remain of a wooden vessel and five clay vessels | Late Roman Iron Age |
| CGG_2_106749 | Grave 1144 | An individual aged 16–17, crouched on the right side, head inN | Two bronze brooches,S-shaped bronze hook, 3 glass beads, spindle whorls of bone, iron knife with partly preved wooden handle, iron strap buckle,bronze sewing needle, bronze terminal strap and one c´lay vessel, | Early Roman Iron Age |
| CGG_2_106748 | Grave 1241 | Male adult, supine position, head in NNE | Iron arm-ring, iron knife, iron rivets, two iron iron rings, iron strap buckle and iron awl | Roman Iron Age |
| CGG_2_106750 | Grave 951 | Female 16–17, placed in supine position | No artefacts found | Roman Iron Age |

**Table S7.2.1:** List of the samples studied from Denmark, Bornholm, Bornholms Sønder, Pedersker parish, Slusegård and summary of the corresponding anthropological and archaeological studies.

#### Denmark, Falster, Falsters Nørre District, Idestrup Parish, Sønder Tåstrup

Coordinates: 54.724635, 11.974114

Sophie Bergerbrant and Marie Louise Jørkov

Human bones including three human crania (IX, X and XI) were given to the Zoological museum in 1944 by L. Kring from Nykøbing Falster. Thenon-cranial human bones are assumed to belong to cranium XI. Bones from cattle, sheep and horse were also given to the museum at this time. All the bones come from a wetland in Sønder-Tostrup, Idestrup parish. The two sampled crania: cranium IX (CGG_2_106739) and cranium XI (CGG_2_106740) are dated to the Early Neolithic and the Pre-Roman Iron Age, respectively. Showing that the bones ended up in the wetland at two very distinctly different times. If the interpretation that the non-cranial bones belong to cranium XI then the Pre-Roman Iron Age person was probably deposited in a more or less complete state. Both individuals are males aged 25–35 years at the time of death.

CGG_2_106739 (ZMK81/1944, AS136/2007, IX) an adult male; UCIAMS-243272, 4795±15 BP, 3640–3520 BCE >30kDa collagen yield (%) 7.9, d15N (‰) 9.8. d13C (‰) -20.1. C/N (atomic) 3.2.

CGG_2_106740 (ZMK81/1944, AS136/2007, XI) an adult male; UCIAMS-243273, 2135 ±15 BP, 350-50 BCE >30kDa collagen yield (%) 8.7, d15N (‰) 10.4. d13C (‰) -20.5, C/N (atomic) 3.2.

#### Denmark, Falster, Falsters Nørre, Eskilstrup, Borremose (070102-38)

Coordinates: 54.844, 11.914 (coordinates not exact findspot)

Sophie Bergerbrant and Marie Louise Jørkov

Skeletal remains of two individuals were recovered during peat digging in 1948 in Johs. Krabbes Mose. In addition a cranium of a cattle and an oak stump were also found. The objects were sent to the museum by L. Kring from Nykøbing Falster. The remains were found at c. 3 m depth. The remains are from one adult (individual I) and one juvenile (individual II).

CGG_2_106742; NM424/48, I has been dated to the Early Neolithic, 3700–3630 BCE (UCIAMS-243275 4870±15 BP, >30kDa collagen yield (%) 9, d15N (‰) 10.4. d13C (‰) -19.9. C/N (atomic) 3.3).

#### Denmark, Falster, Falsters Nørre, Eskilstrup, Sørup Mose and Tåderup mose

Coordinates: 54.844881, 11.91337

Sophie Bergerbrant

A number of artefacts from the Neolithic to the Roman Iron Age have been found at different parts of the wetland area Sørup Mose. The exact location of the human bones has not been possible to relocate. At least three human skeletal elements, but possible more have been found in the wetland bog. The finds have mainly been done within the time period 1934–1945 during peat digging. In this project human remains have been sampled from different parts of the wetland.

CGG_2_106741 sample is one of two human remains found in one area of the wetland area (ZMK82/1944) in 1944. CGG_2_106741 dates to the Early Neolithic 3710–3630 BCE.

The second sample CGG_2_106735 (NM397/45) was found at 1.2 metres depth 20 metres from dry land while peat digging in 1945. Though not preserved it should have been found together with a skull. The remains are of a well preserved male that originally was determined to be c. 1.7 metres tall, animal bones were submitted to the museum at the same time, but there is no information of the human and animal bones found together in the wetland. The radiocarbon date dates this find to the Pre-Roman Iron Age, 400–230 BCE.

CGG_2_106741 (ZMK82/1944, Ind.2) adult female, BCE (UCIAMS-243274, 2270±15 BP, >30kDa collagen yield (%) 4.7, d15N (‰) 9. d13C (‰) -20.1, C/N (atomic) 3.2)

CGG_2_106735 (NM397/45) adult male, 400–230 BCE (UCIAMS-243264, 2270±15 BP, >30kDa collagen yield (%) 8.8, d15N (‰) 11.1. d13C (‰) -19.7, C/N (atomic) 3.2).

#### Denmark, Falster, Falsters Nørre, Nørre Vedby, Nørre Vedby

Coordinates: 54.917, 11.837 (coordinates not exact findspot)

Sophie Bergerbrant and Marie Louise Jørkov

Human remains were recovered during peat digging in the wetlands close to Nørre Vedby Church. The human remains consisted of fragments from two skulls (a calvarium and a fragment of a calva). The human remains were found together with mandibles of wild boars.

CGG_2_106743 (ZMK 25/1942) has been dated to the Early Neolithic 3630-3380 BCE (UCIAMS-243276 4740±15 BP, >30kDa collagen yield (%) 13.9, d15N (‰) 10.4. d13C (‰) -19.1. C/N (atomic) 3.3).

#### Denmark, Falster, Idestrup, Elkenøre (07.02.05-91)

Coordinates: 54.73, 11.98

Anders Fischer

Pre-Roman Iron Age – Roman Iron Age (100 BCE–64CE)

During peat digging in 1942 a number of human skeletal remains were salvaged. Among these are cranial parts of two individuals, characterised by a pale and a dark patina, respectively. Via AMS dating, the pale one turned out to be from the Iron Age: 2029±26 BP (UB-37909; $\delta^{13}C$ -20.5, $\delta^{15}N$ 10.8). The dark cranium dates to the Early Neolithic. Based on parallels to better documented sites, the bone assemblage is inferred to be the result of sacrificial depositions.

CGG_2_022099 (AS_73/2007,_ZMK_232/1942,_37:12;_pale_individual, NEO889).

#### Denmark, Funen, Kalvehavegaard, Kølstrup sogn, Bjerge Herred (080106-150)

Coordinates: 55.4391, 10.6329

Malene R. Beck

In 2016 and 2017, Museums of Eastern Funen excavated a ploughed-down Bronze Age round barrow at Kalvehavegård, near the southern shore of Kerteminde Fjord on Funen, Denmark[382].

The barrow covered a central grave containing the cremated bones of a man and a woman in an oak-log coffin, accompanied by an extraordinary set of grave goods. With a date of around 1400 BC, this is one of the earliest examples of a cremation grave in Denmark. A flat-grave cemetery, originally containing at least 11 inhumation graves, was discovered on the northern periphery of the barrow. $^{14}C$ dates reveal that most of these graves are from Nordic Bronze Age Period I (1700–1500 BCE), but at least one is from Late Neolithic I (2250–1950 BCE). The flat graves appear to have numerous maritime associations, with contents including various mollusc shells and perhaps seaweed. The dead were buried in a fashion indistinguishable from the LN tradition. The NBA II barrow and the central grave appear to represent a break or change in tradition and ideas but the use of an existing burial ground indicates the continuation of a family tradition. Four samples from this site are included in the study (Table S7.2.2).

| CGG No | Grave id | Poz No | $^{14}C$ dating BP |
|---|---|---|---|
| CGG_2_100212 | Grave A12 | Poz-103940 | 3240±35 |
| CGG_2_100215 | Grave A79 | Poz-103942 | 3285±35 |
| CGG_2_100220 | Grave A77 | Poz-103945 | 3315±35 |
| CGG_2_100222 | Grave A78 | Poz-103946 | 3715±35 |

**Table S7.2.2:** List of samples from Denmark, Funen, Kalvehavegaard, Kølstrup sogn, Bjerge Herred and corresponding $^{14}C$ dating.

#### Denmark, Funen, Odense county, Bjerge hundred, Kølstrup parish, Nymarksgaard (080106-28)

Coordinates: 55.43973, 10.6324

Johan Zakarias Gårdsvoll & Mads Dengsø Jessen

In 1937 (j.nr. NM1 89/42) a mound with a stone built double grave was excavated by Helweg Mikkelsen at Nymarksgaard not far from the Ladby ship burial. The skeletal remains were sent to the National Museum of Denmark in 1939, but no grave goods were allegedly found. In 1940 K. Thorvildsen from the National Museum of Denmark also inspected the site without finding any bones or grave goods. In 1981 a left humerus presumably of a man aged 35–55 was carbon-14 dated to 1290±70 BP but it is uncertain whether the bone stems from this site or not.

CGG_2_107529 (NM1 89/42).

#### Denmark, Funen, Odense county, Åsum hundred, Fraugde parish, Fraugde/Fravde (080803-7)

Coordinates: 55.3563933, 10.4951952

Johan Zakarias Gårdsvoll & Mads Dengsø Jessen

In 1890 in J. E. Boesen inspected and wrote about a grave and a bronze fibula which led to the 1895 excavation by Carl Peter Neergaard who excavated more than 80 graves – 9 inhumation and the rest urns and cremations. The two sampled inhumations contained several types of grave goods, such as (Grave 4) pearls of glass and amber, bronze rings and two bronze fibulas, and (Grave 22) pottery, a bone comb and 4 glass pearls. These two graves can be dated to the Late Roman Iron Age.

CGG_2_107398 (NM1 C 8376-88, Grave 4).

CGG_2_107399 (NM1 C 8448-50, Grave 22).

#### Denmark, Funen, Odense, Bjerge, Rynkeby, Illemose (080111)

Coordinates: 55.390942, 10.589975

Sophie Bergerbrant

The site is a small so-called weapon-sacrifice place and the weapons date the site to the 3rd century CE[383]. The mire Illemose contains c. 40 artefacts of metal and wood as well as animal and human bones[384]. Løvschal et al.[383] compares the site with Alken Enge, based on its probable long time use, the composition of the deposition. Though at the time of writing that they did not have access to any radiocarbon dates of the humans.

At Illemose minimum eleven individuals have been found (Fund og Fortidsminder 080111-22[384], though only two complete craniums were found. They have been interpreted as being the remains from adult men[384]. Some of the human bones have been lost but some of the materials are still in collection at Antrologisk samling. Two individuals were sampled CGG_2_106709 (NMC7180-7203, C7202, II) and CGG_2_106710 (NMC7180-7203, C7202, I). Both individuals have been interpreted as male.

Both individuals has been radiocarbon dated to the Pre-Roman Iron Age (CGG_2_106709, UCIAMS-272879, 2200±15 BP, >30kDa collagen yield (%) 0.8, d15N (‰) 10.4. d13C (‰) -

20.3. C/N (atomic) 3.2 / UCIAMS-243259, 2265±15 BP>30kDa collagen yield (%) 1.2, d15N (‰) 11. d13C (‰) -20.4. C/N (atomic) 3.2, giving a combined date of 2233±11 BP 380–200 BCE and CGG_2_106710, UCIAMS-272880, 2245±20 BP, >30kDa collagen yield (%) 3.6, d15N (‰) 10.3. d13C (‰) -19.8. C/N (atomic) 3.1 / UCIAMS-243260, 2300±15 BP >30kDa collagen yield (%) 2.2, d15N (‰) 10.3. d13C (‰) -20.4. C/N (atomic) 3.2) giving a combined date of 2265±13 BP 400–230 BCE).

The dates show that they either were placed in the wetland at the same time or in close proximity to each other. The calibrated time span is 380–200 BCE and 400–220 BCE. Combined all four dates, if they were deposited at the same time, indicate a time period around 390–230 BCE (2254±8 BP).

As Løvschal et al.[383] argues the site resembles Alken Enge, though the radiocarbon dates indicate that the human remains were deposited at least 180 years earlier than the human remains from Alken Enge.

#### Denmark, Funen, Odense, Båg, Skydebjerg parish, Neverkær Mose Høgholm (080215–18)

Coordinates: 55.3978, 10.0852

Sophie Bergerbrant

During dredging work in 1974 in a mine were bones and a layer of hassle sticks foundT. The bones came both from a human and animals. The human remains were found c. 43 cm under the surface. The find is dated to the Pre-Roman Iron Age 350–100 BCE, UCIAMS-275217

2145±15 BP (>30kDa collagen yield (%) 5.9, d15N (‰)19.8. d13C (‰) -20.0. C/N (atomic) 3.1).

CGG_2_106804 (OBM1303).

#### Denmark, Funen, Odense, Højstrupvej (080407-31)

Coordinates: 55.39891, 10.34511

Sophie Bergerbrant

A small cemetery partly excavated by E. Albrectsen 1950, one inhumation and cremation grave have been excavated at the site (Albrectsen 1968:36).

CGG_2_106835 (FSM 8869-76, grave 1) is an individual grave oriented west-southwest-east-northeast. The individual was placed in a right side hocker position with the head in the west. The grave contained an adult female who was buried with a bronze ankle ring, pieces of bronze mounted with a silver sheet disk, a bronze strap buckle, a bronze ring, an iron ring, iron mount, 29 glass-, three amber and one clay beads and a clay pot[380,385]. The grave dates to the early parts of the Late Roman Iron Age[385].

#### Denmark, Funen, Odense, Skovby, Bogense parish, Lillestrand (080601-21)

Coordinates: 55.555.399, 10.062.143

Sophie Bergerbrant

A human cranium and other skeletal elements were found in Fogense mose in the late 1970's (Fund og Fortidsminder, 080601-21). The landowner was digging for a water hole and a cranium was found. Later the museum excavated the heaps of soil from the maschine digging the water hole and found additional human bones from the post cranium. The bones and the cranium are assumed to come from the same individual.

CGG_2_100145 (OBM6140) is an undated remains of a male.

#### Denmark, Funen, Odense, Skovby, Melby parish, Engeldrup Bro (080605-19)

Coordinates: 55.5407, 10.2292

Sophie Bergerbrant

In the middle of 1960's in a by cattle created water hole while drinking in the stream a group of hunters found human remains (Fund og fortidsminder, 080605-19). A well preserved human skeleton was found during a later excavation by Anders Jæger, Bogense Museum. The only bones that were missing were from the knee and below, the latter is thought to have been destroyed by the cattle. Only the cranium was brought into the museum, the remaining remains were reburied at the site. The remains were found at a depth of c. 50 cm.

CGG_2_100144 (OBM6139).

#### Denmark, Funen, Odense, Vends, Tværvejen (080701-56)

Coordinates: 55.476.544, 9.896.608

Sophie Bergerbrant

Human remains placed in hocker position were found 2008 during an excavation by Odense Bys Museer[386]. In what probably had been a smaller bog area. The individual was placed in a possible prehistoric pit from a possible prehistoric peat digging. The remains was of a male aged c. 40 years of age. Under the human remains were a few horse bones found. In the vicinity of the human remains was a shard of pottery found but no other objects (Bonde 2011a&b).

CGG_2_100143 (OBM 4907) has been dated to the Pre-Roman Iron Age/ Early Roman Iron Age, 200 BCE - 60 CE (AAR-13785, 2068±41BP; Bonde 2011a).

#### Denmark, Funen, Odense, Vends, Udby parish, Rolund No 18

Coordinates: 55.4462, 9.8420

Sophie Bergerbrant

A single grave excavated by E. Albrectsen 1951. The grave (CGG_2_106837) contained the remains of an adult/mature male placed right side hocker in a west-east oriented grave. There were traces of a wooden coffin, the objects found in the grave was an iron knife and one ceramic vessel. The grave is dated to the Early Roman Iron Age[380,385].

CGG_2_106837 (Kranie, kæbe).

#### Denmark, Funen, Odense, Åsum, Fraugde, Nørrebjerg (080803-202)

Coordinates: 55.3626, 10.4798

The excavated area contains remains from many prehistoric periods. The excavated area contained hundreds of house remains and some burials. The sampled material comes from one inhumation grave excavated at the site. Probably dating to the Viking Age.

CGG_2_105541 (OBM4937X1242).

#### Denmark, Funen, Svendborg, Salling, Hillerslev, Skregeskov

Coordinates: 55.2341, 10.3628

Human remains found while draining in 1975, during excavation a human cranium and skeletal remains from the upper body were found together with cattle bones. The remains are from an adult female dating to the Pre-Roman Iron Age.

CGG_2_106806 (FSM6863, Grav 2, x2-7) has been dated to 2230±15 BP, 380–200 BCE (UCIAMS-275218, >30kDa collagen yield (%) 2.6, d15N (‰) 9.6. d13C (‰) -21.3. C/N (atomic) 3.1)).

#### Denmark, Funen, Svendborg, Vindinge, Frørup, Sludegaard Sømose (090606-34 A)

Coordinates: 55.2528, 10.7450

Sophie Bergerbrant

Human remains found during peat digging in 1946. In the wetland area there are a number of objects from ceramic vessels, flint tools from the Neolithic, a neck ring from the Late Bronze Age and animal bones have been found (fund og fortidsminder 0906066-34 A).

CGG_2_105358 (NM467/46) is a male over 40 years old.

#### Denmark, Funen, Svendborg, Vindinge, Avnslev, Regstrup

Coordinates: 57.0068, 9.7710

In 1895 were a number of artefacts given to the museum that were expected to come from a burial. Later in 1980 a number of Early Roman Iron Age burials were excavated at the site.The burial is dated to the Early Roman Iron Age.

CGG_2_106836 (FSM2680, Grav J), aDNA XX.

#### Denmark, Funen, Svendborg, Vindinge, Flødstrup parish, Kavslunde Eng (090605-41)

Coordinates: 55.4191, 12.0304

Sophie Bergerbrant

CGG_2_106805 (OBM6964) has been dated to the Early Bronze Age, period II 1500–1410 BCE. UCIAMS-282664, 3170±20 BP, >30kDa collagen yield (%) 12.0, d15N (‰) 9.4. d13C (‰) -20.7, C/N (atomic) 3.2.

#### Denmark, Jutland, Aalborg county, Hornum hundred, Hasseris parish, Gammel Hasseris Grusgrav (120506-18)

Coordinates: 57.057, 9.893

Johan Zakarias Gårdsvoll & Mads Dengsø Jessen

In 1961 (NM1 759/61 and 759/62 and 759/63) Oscar Marseen excavated two inhumation graves with conspicuous amounts of grave goods. The sampled grave (Grave 1) contained a wooden bucket, a bronze pin, wooden boxes with bones, 16 pearls of bone, glass and amber, and a rosetta fibula. The finds place the grave in the Late Roman Iron Age.

CGG_2_107417 (NM1 759/61, Grave 1).

#### Denmark, Jutland, Aalborg, Fleskum, Romdrupgård

Coordinates: 57.0028, 10.0339

An inhumation grave was excavated by O. Marseen in 1958. The grave was oriented west-east and contained one individual in a hocker position on the right side wuíth head in the west. The individual is determined as being an Infant II that was buried with four vessels. The grave has been dated to the Early Roman Iron Age [380].

CGG_2_106816 (ÅHM4578–81) genetically determined to XX.

#### Denmark, Jutland, Aalborg, Fleskum, Sønder Tranders, Sdr. Tranders

Coordinates: 57.0508, 9.9063

Lars Egholm Nielsen

The excavation of the area excavated most of part of a large settlement with cultural layers from the Early Iron Age. Many of the pre-Roman Iron Age and Early Roman Iron Age phases are remains of a row-village. The village changed character in the transition to Late Roman

Iron Age, to dispersed settlements. The area stopped being used at the beginning of the younger Germanic Iron Age. Totally 191 house construction of which ¾ of the house constructions dates to the Early Iron Age has been excavated. At the site a cemetery from late Roman Iron Age and early Germanic Iron Age containing 22 inhumations and 12 cremations have also been excavated. Two inhumations from the Early Pre-Roman Iron Age have also been found at the site. One of these burials appears to be a male that was executed.

Both the settlement and the cemetery are part of a larger complex, west of Sdr. Tranders that have been continuously excavated during the last 20 years. All in all around 420 buildings, of which c. half dates to Early Iron Age, and around 80 burials from both Early and Late Iron Age have been excavated. The settlement area is almost totally excavated, while the central part of the cemetery has not been examined.

CGG_2_106040 (AMH 6784 x 3160, grave A6651) Early Pre-Roman Iron Age Period I, inhumation of an executed male oriented SW–NE. The deceased had been placed with head and torse facing the ground and with his hands bound together. The individual is determined to be a male aged 28–40 years old c. 168 cm tall. There are indications that the individual had been killed by a sword or a knife.

CGG_2_106041 (ÅMH 6784X720, grave A625) A burial oriented SW–NE, dating to Early Pre-Roman Iron Age Period I. The individual that was placed in extreme hocker is determined to be a male aged 30–45 years, c. 169 cm tall. In the foot end of the grave were an iron pin with coiled head found. SampleID 40764 AAR-33313 1884±25 BP, AMH6784X3163; SampleID40765 AAR-33314 1996+-26 BP

#### Denmark, Jutland, Aalborg, Gislum, Fovlum, Hestbek III

Coordinates: 56.717, 9.227

Sophie Bergerbrant

Human remains were found during drainage of a wet peat area 1949. The remains belonged to a 25–35 year old female and are dated to the later parts of the Pre-Roman Iron Age.

CGG_2_106492 (VMÅ 933) a female dated 25–35 years old of local origin SrI 0.71016. The sample is dated to the Pre-Roman Iron Age 200–40 BCE (AAR 16688, 2119±30 BP, Collagen %7.9, C:N ratio 3.2, δ13C VPDB ‰ -19.6, δ15N AIR ‰ 10.6)[387].

#### Denmark, Jutland, Aalborg, Gislum, Gislum, Stistrup Kær VII (120205-67)

Coordinates: 56.7499, 9.2818

Sophie Bergerbrant

The cranium of a female aged 25–35 was found while peat digging in a dried out inlet. The inner side of the frontal bone showed traces of a possible infection (Nielsen et al. 2020).

CGG_2_106494 (VMÅ 895; ) has been dated to the Middle Neolithic, 3370–3040 BCE (AAR-16685 4510±40 BP; $\delta^{13}C$ -20.6, $\delta^{15}N$ 9.0). The individual was probably of local origin with a SrI value of 0.71011[387].

#### Denmark, Jutland, Aalborg, Hellum, Bælum, Smidie

Coordinates: 56.8612, 10.1386

An inhumation grave, found by chance and partly excavated 1942[380]. An east-west orientated burial measuring c.1x2 m. The grave contained three individuals (Sellevold et al. 1984:115), here individual 1 has been analysed, a young individual 15–17 years old. The grave contained a number of pots and an iron knife (Sellevold et al. 1984:115).

CGG_2_106817 (ÅHM no museum number) has been dated to Early Roman Iron Age 25–130 CE (UCIAMS-287279 1935±15 BP, δ13C -20.5, δ15N 12.2) and determined via aDNA to be a female (XX).

#### Denmark, Jutland, Ålborg, Hindsted, Visborg parish, Glerup (120413-66)

Coordinates: 56.2314, 10.1904

Sophie Bergerbrant

A small cemetery containing two graves and four individuals. The cemetery was found during agricultural work 1935. The skeletal material cannot be assigned to the different graves. Grave 1 included a single edge-sword, an iron scissor, an iron knife and at least five pots. Grave 2 contained three ceramic vessels. The artefacts date the graves to Early Roman Iron Age[380].

CGG_2_107480 (NM C23335-46, Skeleton 3c), a mature make, sampled material a C.

CGG_2_107479 (NM C23335-46, Skeleton 4d), an adult male, sampled material an M2.

#### Denmark, Jutland, Aalborg, Hindsted, Oue parish, Stubberup

Coordinates: 56.7141, 9.9415

Bjarne Henning Nielsen

Sample: VMÅ 755 Stubberup, Oue sogn

Central coordinates, a number of finds have been found while draining the mire south of Stubberup stream 1937. The finds consist of part of a human skeleton, a child c. 12 years old, bones from horse, cattle sheep and dog and two clay pots, shards, and six bone awls.
The finds have been interpreted as four depositions close to each other. Within a 8m2 metre area were human bones from the same individual found together with one bone awl, some animal bones and some shards. 55 metres from the area with human bones were two bone deposits found, seven metres from each other. One contained cattle bones. the mandible of a dog and shards. The second included animal bones, many teeth, some shards and five bone awls. 80 metre NE of these were one more animal bone deposition found.
All the depositions can be dated to the Early Iron Age.

CGG_2_106490 (VMÅ 755) has been dated to Pre-Roman Iron Age, 2159±50 BP (UBA-31576 (d13C-21.1, d15N 10.9, C:N 3.2 collagen yield (%) 12.7, dated material bone from a femur[388]).

#### Denmark, Jutland, Aalborg, Hornum, Hasseris Parish, Hamahus

Coordinates: 55.6702, 10.3333

An east-west stone cist grave, with internal dimensions 2x1.5 m, excavated 1963. The cist contained the remains of ten individuals and one iron knife, six ceramic vessels and shereds from a number of more vessels. The burial are dated to the Early Roman Iron Age[380].

CGG_2_106811 ÅHM4753-45 (NM I 313/63) Grav B, ll b, aDNA XY.

#### Denmark, Jutland, Aalborg, Hornum, Hasseris, Scheelsminde

Coordinates: 57.0190, 9.9094

CGG_2_106508 (ÅHM3844 X2888 (see comments). The correct number for this individual is ÅHM0075_grav_III).

#### Denmark, Jutland, Aalborg, Slet, Vilsted parish, Vilsted IX (120714-56)

Coordinates: 56.901, 9.283 (not exact find spot)

Sophie Bergerbrant

Very little information is known about this find. It was found in a bog in Vilsted parish in 1951.

CGG_2_106495, VMÅ 897) is a male 25–35 years old with pathologies on the lower limb that probably affected his mobility. IT has been dated to the Pre-Roman Iron Age 200 BCE–30 CE, 2065±40 BP (AAR-16686, C:N ratio 3.2, δ13C VPDB ‰ -19.9, δ15N AIR ‰ 11.7). He is probably of local origin based on the SrI results 0.70955[387] (Nielsen et al. 2020).

#### Denmark, Jutland, Aalborg, Aars, Hyllebjerg parish, Hyllebjerg X

Coordinates: 56.822, 9.28 (not exact coordinates)

Sophie Bergerbrant

A cranium was found in the wetland 1946 and three years later was the skeletal remains of a torso found. The human remains (VMÅ 389; CGG_2_106493) are one of a female aged 35–45 of a probable height of c. 153 cm with calcified cartilage which could be due to trauma or an infection. The individual had a SrI value that indicates that it was of local origin (0.70930)[387]. In the mire a flint axe, a flint dagger, bone tools and Iron Age pottery have been found. Though none in association with the human remains[389].

CGG_2_106493 (VMÅ 389) has been dated to Late Neolithic 1890–1690 BCE (AAR 16681 3480±27 BP, collagen % 9.8, C:N ratio 3.2, δ13C VPDB ‰ -19.7; δ15N AIR ‰ 9.1[387].

#### Denmark, Jutland, Aalborg, Aars, Museums garden

Coordinates: 56.812, 9.501

Sophie Bergerbrant & Bjarne Henning Nielsen

Human remains, two craniums and other bones from two individuals, that have no clear find circumstances but most likely come from a wetland in the area, both have the Museum number VMÅ2600. A ribbon from one of the skeletons was submitted for dating by M. Gregersen, Retsmedicinsk Institut, Aarhus Universitet, as the police was called for on the suspicion that the skeletons were modern and perhaps victims of a crime.

CGG_2_106486 & CGG_2_106491 are taken from one individual that is a female and has been dated to the Pre-Roman Iron Age 360–50 BCE (AAR-11902 2147±49 BP).

#### Denmark, Jutland, Aalborg, Aars, Skivum, Skivum VI

Coordinates: 56.868, 9.585 (Not exact coordinates)

Sophie Bergerbrant

During peat digging in 1941 west of Skivum was a cranium found, no further information about the find circumstances exist (Fund og Fortidsminder 120811-80). Only the skull is preserved.

CGG_2_106484 (VMÅ 894) has been dated to the Pre-Roman Iron Age 380–270 BCR (AAR 16684 2210±28 BP, C:N ratio 3.2, δ13C VPDB ‰ -19.7 and δ15N AIR ‰ 11.5). The individual is a male adult of 35+ years old. The strontium isotope ratio (0.70914 and 0.770996) for the individual indicates that the person was local[387].

#### Denmark, Jutland, Aalborg, Aars, Sønderup, Sønderup XI

Coordinates: 56.807, 9.628

Sophie Bergerrant

During peat digging in 1952 were the human remains of a woman aged 18–22 dating to the Pre-Roman Iron Age found at a depth of c. 1m. The remains consisted of a cranium without its mandible and some small bones, despite looking for the remaining bones none were found within the area of 1 metres from the cranium. The remains show traces of three healed injuries on the occipitaö bone and the left and right parietal bones posterior[387].

CGG_2_106496 (VMÅ 898) has been dated to the Pre-Roman Iron Age 400–200 BCE (AAR 16686, 2278±35 BP, C:N ratio 3.2, δ13C VPDB ‰ -19.9, δ15N AIR ‰ 11.1). The strontium isotopes values 0.71054 indicates that the female was local[387].

#### Denmark, Jutland, Aalborg, Aars, Ulstrup, Sønderup Østergaard

Coordinates: 56.865, 9.415

Sophie Bergerbrant

Two human remains probably found during peat digging, the remains were found together with pottery (VMÅ972). The remains was of one male (individual #1) and one female (#2) (van der Sluis 2017).

CGG_2_106488 is a female individual that has been dated to the Pre-Roman Iron Age, 360–60 BCE (UCIAMS-275210 2140± 15 BP (>30kDa collagen 11.8, d15N 11.8, d13C -20.7 C/N (atomic) 3.1)[388]. This individual has earlier been dated to 400–190 BCE (UBA-31275 2257± 49 BP, van der Sluis 2017). Giving a combined date of 350–100 BCE.

CGG_2_106489 is a male individual that has been dated to the Roman Iron Age 120–220 CE (UCIAMS-272875 1885±15 BP, >30kDa collagen yield (%) 9.5, d15N (‰) 11.1. d13C (‰) -20.8, C/N (atomic) 3.2) so the wetland was used at different times to deposit human remains within.

#### Denmark, Jutland, Aalborg, Aars, Aars, Skolesamling

Coordinates: 56.796, 9.397

Sophie Bergerbrant

This is a cranium that has no exact findspot related to it, it is part of the museum teaching kit but is most likely a cranium found in wetlands in the area for the museum (Aars).

CGG_2_106485 (VMÅ 2170, II; ) has been dated to the Pre-Roman Iron Age, 400–230 BCE (UCIAMS-272874, 2285±15 BP, >30kDa collagen yield (%) 4.5, d15N (‰) 10.2. d13C (‰) -19.2. C/N (atomic) 3.2).

#### Denmark, Jutland, Århus, Vester-Lisbjerg, Lisbjerg parish, Bulbjerg (150604-16)

Coordinates: 57.1586, 9.0266

A cemetery that was excavated 1900–1906[380,390]. The cemetery contained 89 inhumations and 18 cremations, 44 of the inhumations contained preserved skeletal remains[380]. The cemetery contains both inhumations and cremation graves. The samples investigated in this study are listed in Table S7.2.3. Graves, object poor and object rich graves[390]. Grave 94 is dated to the transition period Pre-Roman Iron Age to Early Roman Iron Age (60 BCE–70 CE).

| **CGG No** | **Museum id** | **Sample description** | **Grave description** | **Period** |
| --- | --- | --- | --- | --- |

| | | | | |
|---|---|---|---|---|
| CGG_2_107471 | NM C11211, Grave 39 | East-west oriented grave, individual placed in hocker on the right side with head in west. Adult | Iron razor, iron knife, seven ceramic vessels, sherds and animal bones | Early Roman Iron Age |
| CGG_2_107472 | NM C111262, Grave 46 | East-west oriented grave, individual placed in hocker on the right side with head in west Maturus | Bronze finger-ring, two iron knives, iron awl, three bone pins, five ceramic vessels and animal bones | Early Roman Iron Age |
| CGG_2_107474 | NM C11502, Grave 72 | A west-northwest –east-southeast oriented grave, individual placed in hocker on the right side with head in west. Adult/mature female | Iron brooch, two knives, iron awl, bone pin, seven ceramic vessels and animal bones | Early Roman Iron Age |
| CGG_2_107476 | NM C12547, Grave 94 | A west-northwest –east-southeast oriented grave, individual placed crouched on the right side with head in west. Adult | Bronze ring, iron pin, iron razor, iron knife and five ceramic vessels and parts of a sixth vessel.. | 60 BCE–70 CE (UCIAMS-275221 20055±15 BP, >30kDa collagen yield (%) 16.2, d15N (‰) 9.0. d13C (‰) -19.1. |

| | | | | |
|---|---|---|---|---|
| | | | | C/N (atomic) 3.1) |
| CGG_2_107477 | NM 11383-91, Grave 57 | A west-northwest –east-southeast oriented grave, individual placed in hocker on the right side with head in west. Adult/mature female | Six ceramic vessels and animal bones | Early Roman Iron Age |
| CGG_2_107478 | NM C12427, Grave 87 | A west-northwest –east-southeast oriented grave, individual placed in hocker on the right side with head in west. Mature female | Two bronze brooches, six bronze pins, silver berloque, S-shaped silver hook, bronzering, two beads of silver wire, one amber bead, one goldfoil bead, one bone bead, iron pin, iron awl, two iron knives, iron ring, eight clay pots, mussel shell and animal bones | Early Roman Iron Age |

**Table S7.2.3:** List of samples studied from Denmark, Jutland, Århus, Vester-Lisbjerg, Lisbjerg parish, Bulbjerg, context description and their dating.

#### Denmark Jutland, Aalborg, Hyldebjerg, Vaarst, (120104-195)

Coordinates: 56.9462, 10.0553

Torben Trier Christiansen

A small cemetery that was partially excavated in rescue excavation in 2019. The size of the cemetery is unknown, however, it is connected to a ridge that is found a few hundred metres west of the Grønhøjgård cemetery. Only two graves are excavated, one heavily disturbed by ploughing. The disturbed grave contained a female grave with a rich and unusual burial equipment dating to the Early Germanic Iron Age. Among the objects in the grave was an a-brakteat made of silver.

CGG_2_106502 (ÅHM7081 X1), Grave 1 an almost complete female skeleton, 32–40 years of age and 166 cm tall. The grave contained an almost complete fibula, a bone comb, part of a possible claff button, a bone spindle whorl and a small ceramic vessel. The sample is dated to the period 410–540 CE (UCIAMS-286999, 1610 ±15BP, N15 12.2, C13 -20.3) aDNA result XX.

CGG_2_106503 (ÅHM7081 X26) UCIAMS-287000, has been dated to 1610 ±15 BP (N15 12, C13 -20.5, 410-540 CE, aDNA result XY).

#### Denmark, Jutland, Aalborg, Lille Bjerggård, Sønderholm, (120511-102)

Coordinates: 57.0107, 9.7506

Torben Trier Christiansen

In connection to agricultural work two graves were found on a ridge in 1972[391]. The bones were well preserved due the chalky soil. One of the graves contained a richly furnished female, dating to the transition between Late Roman Iron Age and -Early Germanic Iron Age (c. 400 CE). South of the female burial were the remains of a partly disturbed grave containing human bones from an individual estimated to have been 190 cm tall. Though all upper body skeletal parts were gone due to the earlier disturbance. The second burial contained a fragmented iron knife and could not be more specifically dated.

CGG_2_106510 (ÅHM0977 A1, aDNA result XX).

#### Denmark, Jutland, Aalborg, Mellemholm, Nørholm, Hornum, (120508-55)

Coordinates: 57.02053, 9.712001

Torben Trier Christiansen

A partly excavated cemetery, placed on a small plateau close to the south facing a larger ridge on the southern Limfjorden coast. Due to chalk in the ground the human remains were unusually well preserved in some of the graves. So far 19 inhumation burials have been excavated over a number of excavation seasons. The graves date to the Early Roman Iron Age to the Early Germanic Iron Age. Both male and females interred at the cemetery, most of them were adults.Some of the graves contain no or just a few objects, however, some graves held a more comprehensive burial equipment. A couple of the male graves contained weapons and one female burial contained a rosette fibula.

CGG_2_106499 (ÅHM5963 X69 (same individual as X129)).
CGG_2_106500 (ÅHM5963 X129 (same individual as X69), aDNA result XX).
CGG_2_106810 (ÅHM5963, A15, x54) aDNA result XY, is an individual found in a east-west oriented graved with among other things an iron knife, a belt buckle, and an awl.The artefacts dates the burial to Late Roman Iron Age.The cranium were found deeper than the rest of the skeletal remains.

#### Denmark, Jutland, Aalborg, Sdr. Tranders, Sdr. Tranders by, Sønder Tranders parish, (120113-51)

Coordinates: 57.0088, 9.9721

Torben Trier Christiansen

Part of a larger cemetery excavated in 2005[392]. The cemetery is placed along a larger elevation in the landscape. The cemetery contained 28 burials of which 23 were inhumations. The cemetery is believed to date to the Late Roman Iron Age– Early Germanic Period. Four of the five cremations graves were very similar to the inhumations as the remains were placed in rectangular graves in the same size as the inhumations. Four of the inhumations contained unusual well-preserved skeletal material. This was three adult women and one adult male. The cemetery's full size is not known, later excavations indicate that it was larger than believed in 2005.

CGG_2_106506 (ÅHM5401 X97, Inhumation grave A97)
CGG_2_106507 (ÅHM5401 X126, Inhumation grave A95) has been dated to 1830 ±20 BP, (UCIAMS-282660 (>30kDa collagen yield (%) 2.4, d15N (‰)11.2. d13C (‰) -20.0. C/N (atomic) 3.2)).

CGG_2_106509 (ÅHM5401 X113. Inhumation grave A98).

#### Denmark, Jutland, Dover and Skanderup parishes, Alken Enge

Coordinates: 56.04, 9.84

Pre Roman Iron Age–Roman Iron Age (193 BCE–206 CE)

From the Alken Enge site a large number of human remains have been recovered. At the site ceramic, animal bones, weapons and wooden objects have been found during the excavations. The ceramic, animal bones, weapons and wooden objects stem for a long period of time. The human remains, however, seem to be from one occasion that fall within the time period 2 BCE to 54 CE (Holst et al 2018).

The human remains have been found in several parts of the 40 ha large wetland area. The samples analysed here come from a concentrated deposition excavated 2009–2014. Animal gnawmarks indicate the human remains were deposited sometime after death. Remains from at least 82 individuals were identified based on the excavated left femurs. The sex specific traits indicate an exclusive or near-exclusive male population. The age span is from subadults to mature, with a predominance of individuals in the age span of 20 to 40 years of age. Antemortem trauma occurs to a limited extent while perimortem trauma in the form of sharp force trauma has been recorded on numerous human remains[383,393].

The human remains have been interpreted as deceased from a larger combat that was deposited in calm water sometime after death[383,393]. Fourteen individuals from the site have been analysed for this article, see Table S7.2.4 for details about the analysed material.

| **Sample id** | **Museum id** | **Lab nr** | **$^{14}$C Dating** | **±** | **date** | **Collagen yield(%)** | **C:N** | **Marin input** |
|---|---|---|---|---|---|---|---|---|
| CGG_2_019201 | SBM1028 2013, X2244 | AAR-24489 | 2051 | 21 | 149 BCE–CE 20 | 6.0 | 3.2 | - |

| | | | | | | | | |
|---|---|---|---|---|---|---|---|---|
| CGG_2_019202 | SBM1028 2013, X2854 | | | | | | | |
| CGG_2_019203 | SBM1028 2014, X4052 | AAR-24495 | 2035 | 21 | 98 BCE–CE 57 | 3.7 | 3.2 | - |
| CGG_2_019204 | SBM1028 2014, X3305 | AAR-24494 | 2042 | 22 | 106 BCE – CE 36 | 7.2 | 3.2 | 0% |
| CGG_2_019205 | SBM 1028 X991 | AAR-18054 | 1987 | 25 | 44 BCE–CE 115 | 8.8 | 3.2 | 0% |
| CGG_2_019206 | SBM 1028 X955 | AAR-18053 | 2022 | 25 | 94 BCE – CE 65 | 8.4 | 3.2 | 0% |
| CGG_2_019207 | FHM 1880 QKM | AAR-24496 | 1926 | 25 | CE 29-205 | 7.8 | 3.2 | - |
| CGG_2_019208 | SBM1028 X2381 | AAR-24490 | 2113 | 18 | 194 – 51 BCE | 7.3 | 3.2 | 0% |
| CGG_2_019209 | SBM1028 X1611 | AAR-18060 | 2009 | 25 | 51 BCE – CE 77 | 4.8 | 3.2 | 0% |
| CGG_2_019210 | SBM1028 X912 | AAR-18050 | 1994 | 25 | | 10.2 | 3.1 | 3% |
| CGG_2_019211 | SBM1028 X409 | AAR-18045 | 2084 | 25 | 171 BCE–CE 1 | 10.7 | 3.2 | 1% |

| | | | | | | | | |
|---|---|---|---|---|---|---|---|---|
| CGG_2_019212 | SBM1028 X952 | AAR-18052 | 1976 | 30 | 41 BCE – CE 121 | 9.5 | 3.1 | 0% |
| CGG_2_019213 | FHM 1880 OOA | AAR-10.5 | 2477 | 30 | 770 – 425 BCE | 10.5 | 3.2 | - |
| CGG_2_019214 | SBM1028 X1000 | AAR-18055 | 1972 | 25 | 38 BCE – CE 120 | 8.6 | 3.1 | 1% |
| CGG_2_019215 | SBM1028 X1406 | AAR-18059 | 1923 | 25 | CE 28-206 | 7.7 | 3.2 | 0% |
| CGG_2_019216 | SBM1028 X2048 | AAR-24488 | 2005 | 31 | CE 28-206 | 4.3 | 3.3 | 0% |

**Table S7.2.4:** List of samples from Denmark, Jutland, Dover and Skanderup parishes, Alken Enge and corresponding $^{14}C$ dating.

#### Denmark, Jutland, Fleskum, Romdrup, Christianshøj

Coordinates: 57.0016, 10.0422

A stone-framed inhumation grave containing remains of three humans excavated in 1951. In the grave two ceramic vessels were recovered, dating the burial to the Early Roman Iron Age [380].

CGG_2_106815 (ÅHM414, Skelet 1b) aDNA determination XX, earlier osteological estimated an adult male.

#### Denmark, Jutland, Grønhøjgård, Gunderup, Fleskum, Vaarst, Aalborg (120104-192)

Coordinates: 56.9444, 10.0481

Torben Trier Christiansen

An unusually large and rich cemetery from Late Roman Iron Age to Early Germanic Iron Age, containing primarily inhumation graves[394]. A substantial part, over 100 graves, of the cemetery were excavated in 2018–2022. The cemetery covered an area of almost 5000 square metres. A number of small metal artefacts and jewellery found by metal detectors indicates that many graves have been destroyed over the years. The burials are placed in long rows along the top of a ridge. In the north part of the cemetery the post holes of a contemporary probable three-aisled house was found. The structure was surrounded by the largest graves at the site. The excavated burials showed a large variation both in shape and content. Both large chamber tombs containing multiple individuals and more ordinary inhumations were among the excavated graves. Even if a large part of the graves have been reopened during the Iron Age, and objects clearly were removed, did still some burials contain an unusual rich and varied burial equipment. Human remains were registered in a number of the graves.

The cemetery is a high-status burial ground of a size and character that is without regional parallels, and the Grønhøjgård cemetery is clearly just one part of a larger cemetery complex. Another two cemeteries from the time period are located on small ridges a couple of hundred metres northwest and northeast (Hyldebjerg cemetery). A few metres north of the site are traces from a larger settlement that probably could be connected to the cemeteries[394].

CGG_2_106504 (ÅHM7032 X122, grave A43) has been dated to 1700BP±15 (UCIAMS-275211, >30kDa collagen yield (%) 14.1, d15N (‰) 10.7. d13C (‰) -19.6. C/N (atomic) 3.1) (260–420 CE).

#### Denmark, Jutland, Hjørring, Børglum hundred, Svennum Offermose, Jerslev Parish (100106-254)

Coordinates: 57.403, 9.88

Sidsel Wåhlin

Roman Iron Age (25–130 CE)

VHM00500x00022 is one of six human skulls deposited in a narrow topographical bog[395]. The bog was excavated in 2013 and 2015 by the Vendsyssel Historiske Museum. The site is a complex ritual deposit site where offerings were deposited in small oval peat cuts in a very structured manner during the Pre Roman and Roman Iron Age. According to its $^{14}C$ date the skull is older than the context it was deposited in, as it belongs to the final set of deposits the six human skulls and some animal bones. Some of these are dated to the 5th century CE. The ceramics connected with the skulls are dated to the early Roman Iron Age (first-second century CE). Five of the skulls were sampled but only one yielded sufficient DNA to be admitted to this study.

CGG_2_016034 (VHM00500x00022) UCIAMS-233750, 1935±15 BP, >30kDa collagen yield (%) 4.7, d15N (‰) 10.9. d13C (‰) -20.3, C/N (atomic) 3.1.

#### Denmark, Jutland, Hjørring, Børglum, Brønderslev parish, Lygtebæk (100101-191)

Coordinates: 57.2577, 9.982

Sophie Bergerbrant, Sidsel Wåhlin

In 2021 a human cranium and a mandible was brought to the museum for examination. The human remains have so-called bog patina and most likely come from a wetland. The material is given to the museum. The cranium and the mandible do not match. Possibly the mandible was found together with ceramics in 1947 during peat digging. From this find there is information that the pots were found with a human mandible but the mandible never reached the museum. The finds are found in a bog on Gl. Burholdts Jorde.

CGG_2_106526 (VHM 01192x000002) only gave limited genetic information but is the remains of a female (XX).

#### Denmark, Jutland, Hjørring, Børglum, Jerslev parish, Aalstrup, Mellergaard (100106-259)

Coordinates: 57.2789, 10.0142 (not exact coordinates)

Sidsel Wåhlin

In 1937 the landowner of Mellergård, Ålstrup found a human cranium and some stone c. 60 cm beneath the current surface of the bog. The finds were given to the museum.

CGG_2_106530 (VHM 22600, VHM 077/1937) according to the genetic analysis this is the remains of a female (XX).

#### Denmark, Jutland, Hjørring, Børglum, Sejlstrup parish, Kornum Bro (100109-11)

Coordinates: 57.32, 9.86

Sophie Bergerbrant, Sidsel Wåhlin

A human cranium without the mandible was found 1958 close to Kornum bridge during dredging as part of the preparation for work on the bridge. Archaeologists from the Vendsyssel Historiske Museum were sent to the site. They found among other things a flint axe, an antler axe and flint. The objects date to both the Neolithic and the Bronze Age .It should be noted that the finds were scattered and not observed in situ.

CGG_2_106529 (VHM 1965/0131, VHM 046/1922) has been dated to 1230±15 BP (UCIAMS-275213, (>30kDa collagen yield (%) 13.9, d15N (‰) 10.5. d13C (‰) -20.0. C/N (atomic) 3.1)).

#### Denmark, Jutland, Hjørring, Børglum, Stenum parish, Stenum (100111-21)

Coordinates: 57.32, 9.86 (coordinates not exact findspot)

Sidsel Wåhlin

During peat digging in 1922 a cranium without its mandible was found. It is the remains of a younger adult. No more information than that was given to the museum

CGG_2_106522 (VHM 14759, VHM 046/1922) are genetically determined to be a male (XY).

#### Denmark, Jutland, Hjørring, Børglum, Tolstrup parish, Vibsig Mose (100113-192)

Coordinates: 57.3, 9.91

Sophie Bergerbrant Sidsel Wåhlin

Over an area of 40x40 metres were three cranium, antler, animal bones, stones and shards (VHM 1948/0145, VHM 1947/0535, VHM 1947/0520, VHM 016/1947) found during peat digging 1947 and 1948. One of the crania VHM 1947/0520s was an infant I/II.

CGG_2_106535 (VHM 1948/0145) is an adult and has been dated to the Pre-Roman Iron Age (UCIAMS-275214 2150BP±15 BP (>30kDa collagen yield (%) 13.0, d15N (‰) 10.6. d13C (‰) -20.7. C/N (atomic) 3.3)). The sample is genetically determined to be a female (XX).

#### Denmark, Jutland, Hjørring, Børglum, Tårs parish, Sæsing (100114-302)

Coordinates: 57.45, 10.08

Sophie Bergerbrant, Sidsel Wåhlin

The human cranium (CGG_2_106525, VHM 1945/0038, VHM 052/1945) without its mandible was found while peat digging in 1945. The skull has been radiocarbon dated to the Pre-Roman

Iron Age period, UCIAMS-275212 2235±20 BP (>30kDa collagen yield (%) 11.8, d15N (‰) 10.7. d13C (‰) -20.5. C/N (atomic) 3.1).

CGG_2_106525 (VHM 1945/0038, VHM 052/1945).

#### Denmark, Jutland, Hjørring, Børglum, Vrejlev parish Tollestrup Mose (100116-195, 196, 213 & 218)

Coordinates: 57.3461, 10.037 (general coordinate for the mire, however the individual finds are located to specific plots in the mire)

Sidsel Wåhlin, Anders Fischer and Sophie Bergerbrant

In Tollestrup Mose at least the remains of six human and some artefacts have been found during peat digging, mainly in the period between 1938 to 1943, in different parts of the wetland.

CGG_2_106527 (VHM 22943, VHM 039/1938) a cranium without a mandible was found in 1938, the cranium shows traces of damage due the spade on the nose ridge. The remains have genetically been determined as female (XX).

CGG_2_106528 (VHM 1940/0043, VHM 029/1938) Is a well preserved complete skeleton placed on its back and held down by a birch branch in the bog. The human remains have previously been labelled as a well preserved female almost complete, though the genetic analysis determined it as male (XY).

CGG_2_023270 (VHM 041/1941) human remains and a small piece of rope were found at a depth of c. one metre in an ancient peat pit. The human bones were not placed in an anatomically correct position. A stone was found in the mouth of the skull. Our sampling is based on a -4 tooth. It is dated to the Late Bronze Age or possibly early Pre-Roman Iron Age (780–420 BCE) 2491±38 BP (UB-39167; $\delta^{13}C$ -20.0, $\delta^{15}N$ 11.4). VHM 41/1941, 1941/10. NEO948.

CGG_2_023271 (VHM 48/1941, 1941/26) was found together with another skull and other human bones in 1941 (however the second skull and the mentioned bones are not in the Museums collection). The sampling is a +7 tooth. It is dated to the Late Bronze Age or Pre-Roman Iron Age (730–380 BCE) 2373±38 BP (UB-39166; $\delta^{13}C$ -20.1, $\delta^{15}N$ 11.2). VHM 48/1941, 1941/26. NEO949.

#### Denmark, Jutland, Hjørring, Dronninglund, Hellevad parish, Hellevad (100205-184)

Coordinates: 57.215, 10.157 not precise coordinates, only located to Parish

Sophie Bergerbrant, Sidsel Wåhlin

A cranium (CGG_2_106553, VHM 1944/0062) damaged by the spade found while peat digging in the parish Hellevad. The exact find spot for this find is unknown, the coordinate relates to the parish it is found within. The radiocarbon date shows that this is dated to late Germanic Iron Age 670–770 CE, UCIAMS-275215 1275±15 BP (>30kDa collagen yield (%) 9.1, d15N (‰) 9.8. d13C (‰) -20.1. C/N (atomic) 3.1).

CGG_2_106553 (VHM 1944/0062).

#### Denmark, Jutland, Hjørring, Horns, Ellinge parish, Strandby (100302-6)

Coordinates: 57.4933, 10.4989

Sophie Bergerbrant, Sidsel Wåhlin

A small cemetery with six graves from the Roman Iron Age was excavated by dentists Valdemar Friis for the Vendsyssel Historiske Museum. The bones were not separated by grave but found in a mixed box without individual numbering in the Museum in 2019. Only one skull with teeth was preserved and has been analysed for this project.

CGG_2_106531 (VHM 01200x000002) is radiocarbon dated to the Early Roman Iron Age 1-120 CE (UCIAMS-272878, 1970 ±15 BP, d13C -19.1, d15N 13.2, C/N 3.2).

#### Denmark, Jutland, Hjørring, Vennebjerg, Sankt Olai parish, Hestehavens mose (100612-91)

Coordinates: 57.4219, 10.0055

Sophie Bergerbrant, Sidsel Wåhlin

A human cranium was found while peat digging in 1994. The remains were found in Hestehavens mose c. 1200 metres south from Fulsig and 15 metres from the stream. The cranium was found at a depth of 75 cm. The skull was given to the museum as a gift.The remains are one of a probable adult male.

CGG_2_106524 (VHM 1942/0085, VHM 057/1942) has been dated to the Pre-Roman Iron Age 400–220 BCE (UCIAMS-282661, 2275±20 BP, >30kDa collagen yield (%) 2.0, d15N (‰) 11.6. d13C (‰) -20.6, C/N (atomic) 3.2).

#### Denmark, Jutland, Hørmested, Nygård Mose (100307-94)

Coordinates: 57.478, 10.216

Sidsel Wåhlin, Anders Fischer and Sophie Bergerbrant

Pre-Roman Iron Age (390–197 BCE)

In a bog close to Uggerby Stream one robust human skull without mandible, animal bones and ceramics were found during peat digging in 1944. Our DNA sampling is based on a tooth 5- and has been dated on cranial bone to 2229±35 BP (UB-39161; $\delta^{13}C$ -20.6, $\delta^{15}N$ 11.3).

CGG_2_023274 (VHM_083/1944,_genstand_1944/0083a, NEO952).

#### Denmark, Jutland, Lauritshøj (14.11.08-104)

Coordinates: 56.26, 10.31

Iron Age (367 BCE – 201 BCE)

Human bones have been found in a former bog during a large-scale excavation in 2014[396]. In the southern part of the bog remains of a young woman have been retrieved. The bones of the woman were not found in anatomically correct order. The cranium and the left arm bones were missing. Though teeth have been found during sieving. In this part of the bog there were also over 30 so-called bog pots registered.

CGG_2_018916 (Lauritshøj II FHM 5478) has been dated to 2220±15 BP (UCIAMS-232723 ,>30kDa collagen yield (%) 6.8, d15N (‰) 9.7. d13C (‰) -20.1, C/N (atomic) 3.1).

#### Denmark, Jutland, Illerup, Skanderup (16.02.08-53)

Coordinates: 56.05, 9.93

Roman Iron Age (AD 64–253)

Weapon sacrifice sites

Skeletal remains from humans were found near the Roman Iron Age weapon deposited in Illerup Ådal[397]. Two almost complete human remains were found as well as remains from more individuals[398]. The place is best known for its large weapon deposits of equipment from vanquished armies originally deposited in an Iron Age small lake between 200 to 500 CE[399]. The human remains are found up-stream to Alken Enge[383]. Three individuals were sampled for this study for details see Table S7.2.5.

| **CGG No** | **Museum id** | **Lab nr** | **BP** | **±** | **date** | **Collagen yield(%)** | **C:N** |
|---|---|---|---|---|---|---|---|
| CGG_2_019200 | FHM1880QEV | AAR-24500 | 1892 | 35 | 64–235CE | 7.9 | 3.2 |
| CGG_2_019207 | FHM1880QKM | AAR-24496 | 1926 | 25 | 29–205 CE | 7.8 | 3.2 |
| CGG_2_019213 | FHM1880OOA | AAR-10.5 | 2477 | 30 | 770 –425 BCE | 10.5 | 3.2 |

**Table S7.2.5:** List of samples from Denmark, Jutland, Illerup, Skanderup and corresponding $C^{14}$ dating.

CGG_2_019200 (Illerup_FHM_1880_QEVAAR-24500) had been dated to 1892±35 BP (AAR-24500, Collagen yield % 7.9, C% 42.8, N% 15.7, C:N 3.2, $\delta^{13}C$ -19.0, $\delta^{15}N$ 17.31, FHM 1880 QEV[397]).

#### Denmark, Jutland, Sejlstrup, Nedergaard (Smidstrup) (100109-13)

Coordinates: 57.39, 9.88

Sidsel Wåhlin, Anders Fischer and Sophie Bergerbrant

Late Bronze Age or Pre-Roman Iron Age (725 BCE–392 BCE)

Human mandible found in 1943 during peat digging in a bog owned by the person who gave the specimen to the museum. The site is located close to Liver Å. Our DNA sampling is based on CGG_2_023281 ( VHM 65/1943, VHM 1945/20. NEO953).

CGG_2_023274 (VHM_083/1944,_genstand_1944/0083a) has been dated to 2383±32 BP (UBA-39163; $\delta^{13}C$ -20.1, $\delta^{15}N$ 10.1).

CGG_2_023281 (VHM 065/1943, 1945/0020, NEO953).

#### Denmark, Jutland, Thisted, Mors

Coordinates: 56.84, 8.72

Sophie Bergerbrant

The remains of a subadult (a mandible) found in wetland om Mors was given to the museum in 1905 by the dentist Teilman-Friis from Nykøbing Mors. No other find circumstances are known.

The sample CGG_2_106711 (NM23/190) has been dated to the Late Middle Age-Early Modern time (1450–1620 CE) (UCIAMS-243261, 385±15 BP (>30kDa collagen yield (%) 3.2, d15N (‰) 14.2. d13C (‰) -19.7, C/N (atomic) 3.2).

#### Denmark, Jutland, Vendsyssel

Coordinates: unspecified spot in Vendsyssel

Sophie Bergerbrant Sidsel Wåhlin

The sample CGG_2_106523 (VHM 01197x00001, VHM 01197) comes from a cranium without its mandible. It has a bog patina and the only known find circumstances is that it comes from Vendsyssel. It is the remains from an adult male seen both with the osteological analysis and the genetic ones.

CGG_2_106523 (VHM 01197x00001, VHM 01197).

#### Denmark, Jutland, Vendsyssel (Holger Friis' kaelder)

Coordinates: 57.22, 9.75

Anders Fischer and Sophie Bergerbrant Sidsel Wåhlin

Pre-Roman Iron Age to Roman Iron Age (170 BCE–43 CE)

A ‘bog patinated’ cranium with mandible without specific find circumstances but most likely from Vendsyssel. Our DNA sampling is based on tooth +6. It has been dated on cranial bone to 2043±43 BP (UBA-39165; $\delta^{13}C$ -20.6, $\delta^{15}N$ 11.0).

CGG_2_023272 (VHM_00807x1, NEO950).

#### Denmark, Jutland, Vennebjerg, Mygdal, Vester Klodske

Coordinates: 57.5065, 10.11347

In 1989 two stone-built burials were excavated in Vester Klodske. Both graves date to the Early Roman Iron Age. Here grave 1 was sampled,it contained the remains of an individual placed on the right side hocker and buried with a small bronze ring, an iron knife and four ceramic vessels.

CGG_2_106803 (VHM1991/9, Grave A1), a DNA XY.

#### Denmark, København, Smørum, Smørum, Alpegard

Coordinates: 55.7497, 12.2894

A large excavation during 2020 found remains of a large number of long houses (over 70) and other settlement remains, including over 1100 pits. The dates of the long houses are to the Germanic Iron Age and the larger pits mainly to the Late Bronze Age. In 2021 was one female inhumation grave dating to the Late Roman Iron Age excavated.

CGG_2_106548 (Tak 1597x4588).

#### Denmark, Langeland, Dagsmose, Fodslette (09.03.01-21)

Coordinates: 54.82, 10.73

Anders Fischer and Sophie Bergerbrant

Pre-Roman Iron Age (550–200 BCE)

During peet extraction under as much as 3 metres of sediment several prehistoric artefacts were found together with a number of bones of wild and domestic animals. The assemblage also consists of the skulls of three humans[400]. Based on biological anthropology, the one in question here represents a 17–18 years-old individual (probably female). At the instance of sampling, a complete skull with three teeth and both *pars petrosae* was available. The sampling is based on tooth 4+. Observation on tooth +6 of caries and calculus.

CGG_2_021419 (AS_26/78,_LMR_A_6342,_07:10, NEO883), has been dated to 2326±47 BP (UBA-39138; $\delta^{13}C$ -20.2, $\delta^{15}N$ 11.3).

#### Denmark, Langeland, Langelands Sdr. district, Tryggelev, Rørsløkke Mose (090307-26)

Coordinates: 54.8023, 10.7017

Sophie Bergerbrant

In the mire have a number of artefacts, human and animal bones from different periods been found during peat digging.. On the south side of the mire has flint axes and TRB ceramics been found. Though the bog is mainly known for its Bronze and Iron Age deposits (Fund og Fortidsminder 090307-26)[401]. The mire is probably most well-known for the find of two Late Bronze Age lurs found by peat diggers in 1886 (Broholm 1946:174).Though in the mire has also a large ceramic vessel from the latter part of the Early Iron Age been found as well as undated animal bones (horse, cattle and dog) as well as human remains. The human remains are dating to the Bronze Age period I and hence forgoes the famous lur find.

CGG_2_106744 (NM22/1901) is a male tha thas been dated to Early Bronze Age 1730–1520 BCE, i.e. PI (UCIAMS-243277, 3350±15 BP (>30kDa collagen yield (%) 5.1, d15N (‰) 10.3. d13C (‰) -20.5, C/N (atomic) 3.2).

#### Denmark, Langeland, Langelands Sønder, Humble parish, Humble (090303-113)

Coordinates: 54.8296, 10.6990

In 1922 an inhumation containing a skeleton and ceramic vessel was found. The grave was oriented N–S. Sampled remains were deposited at the anthropological collection in 1990 and labelled grave 2. The sampled material is dated to the Viking Age (770–890 CE) and is a female with mtHG U2e1a.

CGG_2_106772 (LMR12608, Grave 2) has been dated to 1210±20 BP (UCIAMS-282663, >30kDa collagen yield (%) 1.2, d15N (‰) 12.4. d13C (‰) -19.8, C/N (atomic) 3.2).

#### Denmark, Lolland, Maribo county, Lollands Sønder hundred, Gloslunde parish, Hoby (070504-1)

Coordinates: 54.838, 11.142

Johan Zakarias Gårdsvoll & Mads Dengsø Jessen

In 1920 (j.nr. NM1 78/20) a grave was disturbed in a former gravel pit at Hoby. The find was reported to the National Museum of Denmark and Hans Christian Broholm was sent to inspect and excavate the site[402]. The grave was a flat inhumation grave with a NE-SW orientation, containing an incomplete human skeleton (CGG_2_107481) lacking parts of the lower extremities. Parts of the deceased's legs still lay in situ when excavated thus, indicating that the deceased had been lying with his head at the northeast end of the grave. Beneath the legs, a wooden layer was observed which was probably the remains of the base of a wooden coffin. A fragmented mandible was found with some teeth still present in situ and it was later determined that the individual was possibly an adult male aged between 25–35. The grave contained numerous grave goods of extraordinary kind, such as a large bronze basin (Eggers type E97), a bronze saucepan (E140), a bronze pitcher with a trefoil mouth (E126), a bronze tray (E115), a bronze situla (E24) and two silver cups of Roman make (E168). In addition, the grave contained seven metal fibulae of the 'Rollenkappen' type and several high-status goods such as gold rings, drinking horns (only mountings preserved), and many other objects. In total, the Hoby grave ranks among one the richly furnished graves from the Early Iron Age and has therefore also been denoted a princely grave[380,403,404]. The strontium isotope signature from the remains of the mandible suggests a local origin of the entered man[403]. The area near the grave contained a larger settlement, and local historical notes indicate that there might have been several additional burials in the vicinity.

CGG_2_107481 (NM1 C 17946-64, Skeleton 1, Kiste B).

#### Denmark, Lolland, Maribo Lollands Nørre, Birket parish, Uglemose (070401-110)

Coordinates: 54.8862, 11.31484

Sophie Bergerbrant

During peat digging in 1943 a human cranium was found in Uglemose, Birket parish (Fund og Fortidsminder, 070401-110). In 1948 the museum visits the bog/mire in which it has been reported to have been found three or four wooden helmets, a boat and a wooden club, as well as pottery from the Neolithic and Roman Iron Age, a stone axe, and an Early Iron Age bone spear head and cranium from two children. The wetland area has clearly been used over a long period of time (Fund og Fortidsminder, 070401-110). The human remains sampled for this project (CGG_2_106712) dates to the Late Neolithic/ Early Bronze Age 1870–1630 BCE.

CGG_2_106712 (NM634/43, ?) are the remains from an adult male (UCIAMS-243262, 3420±15 BP (>30kDa collagen yield (%) 6.2, d15N (‰) 10.1. d13C (‰) -19.3, C/N (atomic) 3.2)).

#### Denmark, Lolland, Maribo, Lollands Nørre, Birket parish, Hejringe (070401-112)

Coordinates: 54.881623, 11.344825

Sophie Bergerbrant

During peat digging in 1947 the skeletal remains from an adult male was found in Hejringe Enge. The individual is osteological determined as an adult male and is dated to the late Bronze Age Period IV (1020–900 BCE).

CGG_2_106737 (NM827/47) UCIAMS-2432710, 2810±20 BP (>30kDa collagen yield (%) 6.1, d15N (‰) 8.1. d13C (‰) -19.4, C/N (atomic) 3.2).

#### Denmark, Lolland, Maribo, Lollands Nørre, Halsted parish, Juellinge (070403-16)

Coordinates: 55.3075, 12.25271

Sophie Bergerbrant

A small cemetery with four inhumations, grave 4 was found during work in 1908 and the other three graves were excavated by g: Rosenberg in 1909[380]. The graves were placed in a row all oriented N–S.

CGG_2_106771 (NMI C13854-72, Gr. 2, A, RPP). The individual was placed on the right side in a hocker position with the head in the north. The grave contained one bronze kettle, a bronze ladle and strainer set, two drinking horns, a silver pin, three silver brooches, a s-shaped gold hook, a gold berloque, two small gold beads, a wooden casket with bronze mounts, a bone comb, a bronze scissor, a bronze knife, a bronze pin, two ceramic vessels and animal bones. The deceased was determined to be an old female dating to the Early Roman Iron Age, B2[380].

#### Denmark, Lolland, Vigsnæs (Frk. Brandts Mose), Vigsnæs (07.06.22)

Coordinates: 54.89, 11.65

Anders Fischer

Pre-Roman Iron Age (405–208 BCE)

Bog patinated human remains for which there is (currently) no provenance data except a label saying “Vignsæs VIII, Saxkøbing (Frk. Brandts Mose)”. Tooth +5 was sampled for the present project.

CGG_2_017610 (Vigsnæs,_Saxkøbing_(Frk_Brandts_Mose), ZMK 80/1944. NEO126) has been dated to 2289±33 BP (UB-38224, no collagen was available for measuring dietary isotopes).

#### Denmark, Møn, Borre Enge, Borre (05.05.02-96)

Coordinates: 55.01, 12.43

Anders Fischer and Sophie Bergerbrant

Germanic Iron Age (660–780 CE), uncertainty due to unresolved reservoir effect

Remains of a human found in a bog. Submitted to the Zoological Museum of Copenhagen University 1944 by a school teacher. Later on the find was transferred to the Anthropological Laboratory, where at the time of our sampling it had yet not got an inventory number. We sampled a *pars petrosa*, that has been dated to 1298±26 BP (AAR-25172; $\delta^{13}C$ -19.25, $\delta^{15}N$ 16.06).

The combination of topography, date and dietary isotopic values of this skeleton is unusual in a Danish context. The nitrogen value implies a considerable consumption of protein from high trophic level (freshwater) fish. This may indicate a provenance of the individual out-side Denmark. The combined carbon and nitrogen values most probably imply a considerable reservoir effect, which may result in an age off-set of several hundred years.

CGG_2_017609 (Borre_Enge,_Møn, NEO125).

#### Denmark, Møn, Præstø county, Mønbo hundred, Keldby parish, Landledgård (050506-26)

Coordinates: 54.9886848, 12.3730929

Johan Zakarias Gårdsvoll & Mads Dengsø Jessen

In 1952 (j.nr. NM1 380/52) Ole Klindt-Jensen excavated 4 inhumation graves and based their dating to the Early Roman Iron Age on the ceramic grave goods and an iron knife.

CGG_2_107450 (NM1 380/52, C26762, Grave 3).
CGG_2_107449 (NM1 380/52, C26763, Grave 2).
CGG_2_107451 (NM1 380/52, C26764, Grave 4).

#### Denmark, Møn, Præstø county, Mønbo hundred, Magleby parish, Maglebjerghøj (050507-237)

Coordinates: 55.0062, 12.4818

Johan Zakarias Gårdsvoll & Mads Dengsø Jessen

In 1965 (j.nr. NM1 466/72) during ploughing in a field at Maglebjerghøj several graves containing skeletal remains were disturbed. In 1971 Birgitte B. Henriksen inspected the site and excavated the remains of at least 5 highly disturbed inhumation graves with 4 adults and one child. For some of the graves the position in which they had been buried could be observed as hocker. One of the graves contained undefined iron objects and an amber pearl. The positioning of the bodies in the graves could indicate a dating to the Late Roman Iron Age, though a carbon-14 dating could also suggest that the graves might be dated to the Early Germanic Iron Age.

CGG_2_107522 (NM1 573/71, Grave E).

#### Denmark, Nr. Hedegård, Sønder Tranders, Fleskum, Ålborg, Jutland (120113-24)

Coordinates: 57.0195, 9.96671

Excavations at the site have shown that there was a settlement from Bronze Age Period V/VI to the Early Roman Iron Age. A cemetery has been excavated 30 to 40 m north of the village enclosure fence. It contains both cremation and inhumations. Due to a later settlement and modern agricultural practices many of the graves at the settlement have been destroyed, despite this there are about 200 graves preserved. The cremation dates to Pre-Roman Iron Age period IIIB to Early Roman Iron Age phase B1, while the inhumations dates to Roman Iron Age phase

B2. Though the inhumation grave we sampled are radiocarbon dated to the Pre-Roman Iron Age.

CGG_2_106813 (ÅHM3844, Grave A238, XY) has been dated to 2335±20 BP, N15 10.3, C13 -20.8 420–380 BCE (UCIAMS-287001).

#### Denmark, Svendborg, Langelands Sønder, Lindelse, Bogøvej

Coordinates: 54.8711, 10.703

Excavated in the late 1980’s by Langelands Museum. The cemetery dates to the Viking Age and is located on a small drumlins of mixed moraine material with a high chalk content. The preservation conditions for bone material were thus varying, but often good. It is located c. 500 m south of Lidelse Nor on a small hill located on a small hill 500 m south of Lidelse Nor in an (an elongated area measuring 70 m x 30 m in a northeast-southwest orientation). The cemetery was discovered in 1920, when four inhumations were excavated. Over time a total of 49 graves have been excavated, three of which were double graves.

The majority of the deceased had been placed in a supine position, though a few in hocker. Some Individuals had been buried with grave goods, e.g. knives, whetstones, buckles, and/or glass beads.

Two individuals had been buried in carriages, minus wheels (e.g. Grave BA).

One male burial contained an Arabic silver coin, an axe dated to the late Viking Age and a strike-a-light, and . One skeleton appears to have been decapitated (Grave T). Most of the graves were oriented east-west. Of the sexed individuals, 15 were male and 22 female. Only a few were subadults. The graves are generally dated to the 10th century CE, with a possibility of a few graves dated to the early 11th century CE.

During the excavations in 1987 and 1988 were activities dated to the late Bronze Age, including 17 pits (some contained flint tools, large amounts of pottery, small fragments of bronze, and some antler tools) discovered.

A number of burials were sampled and published in Margaryan et al. 2020[405] these were VK286 Denmark_Bogovej Grav BJ, VK288 Denmark_Bogovej Grav BA, VK292 Denmark_Bogovej Grav A.D., VK320 Denmark_Bogovej Grav S, VK338 Denmark_Bogovej Grav BV, VK361 Denmark_Bogovej BX, VK362 Denmark_Bogovej LMR 12077, VK363 Denmark_Bogovej BT, VK364 Denmark_Bogovej BN, VK365 Denmark_Bogovej BS, VK366 Denmark_Bogovej BY, VK367 Denmark_Bogovej D, VK368 Denmark_Bogovej T of which some has been resampled for this project. The additional samples analysed in this study are listed in Table S7.2.6.

| **CGG No** | **Grave** | **Relations to** | **Genetic sex** |
|---|---|---|---|
| CGG_2_106774 | Grav AD (AP?) | VK292_FS | XY |
| CGG_2_106775 | Grav AU | - | XY |
| CGG_2_106776 | Grav BE, younger | - | XY |
| CGG_2_106777 | Grav BE, older | - | XY |
| CGG_2_106778 | Grav BB | - | XY |
| CGG_2_106779 | Grav BU | - | XX |
| CGG_2_106780 | Grav BZ | - | XX |
| CGG_2_106781 | Grav O | | XY |

| | | | |
|---|---|---|---|
| CGG_2_106782 | Grav AO3 | Same as VK292_FS | XY |

**Table S7.2.6:** List of the samples studied from Denmark, Svendborg, Langelands Sønder, Lindelse, Bogøvej, their relation to each other and their sex.

#### Denmark, Svendborg, Langelands Sønder, Lindelse, Kumle Høje

Coordinates: 54.8943, 10.7370

The Viking Age cemetery containing a minimum of 11 individuals was excavated by Langelands Museum in 1998, it dates to 10th century CE. Both male and female adults were buried at the site. One of the graves was a double burial with two men buried on top of each other, one placed on its back and the other facing down. A male cranium in another grave had a healed cranial trauma.One female was buried with a 17 cm long iron knife. One male grave had bones showing that he had broken his legs (perimortem). One female burial was covered with large stone slabs. The burial with must equipment was a female in supine position, she was buried with with iron artefacts, amber, and part of a bronze belt buckle. Several of the male graves not mentioned earlier showed signs of trauma.

One burial, grave O has previously been analysed for aDNA[405].

CGG_2_106785 (LMR12845, teeth set 4).

CGG_2_106786 (LMR12845, Grave E).

#### Denmark, Svendborg, Langelands Sønder, Lindelse, Kaagården

Coordinates: 54.8495, 10.7660

LMR11563

The site was excavated by Langelands Museum in the time period 1984–1987[406]. Cemetery contained 95 graves and was placed on a small hill. It was used from the Late Germanic Iron Age to the Late Viking Age, however most graves dated to the Late Viking Age.

The graves with must burial equipment were found on the top of the hill. The deceased were both adults and subadults. The subadults were buried with iron knives. The cremations were probably done close to a beach as seen by the inclusion of sand and the shells of marine mollusks in some of the graves.

Six graves from the site has earlier been analysed for aDNA (VK274 Denmark_Kaargarden 391, VK275 Denmark_Kaargarden 217, VK276 Denmark_Kaargarden BH, VK285 Denmark_Kaargarden Grav BZ, VK287 Denmark_Kaargarden Grav BS, VK317 Denmark_Kaargarden Grav BF99)[405]. THe additional samples analysed in this study are listed in Table S7.2.7.

| CGG No | Grave | Genetic relation | Sample description |
|---|---|---|---|
| CGG_2_106787 | Grav AK, x27 | 2nd related to CGG_2_106790 | Placed in a light hocker position, determined as male(?) aged 35–55 and buried with an iron knife, aDNA XY |
| CGG_2_106788 | Grav AY, x51 | | The deceased was placed in supine position, and determined as female 25–35, aDNA XX |
| CGG_2_106789 | Grav BR, x106 | | Partly destroyed by the creation of burial BZ, placed in supine position osteological female 18–20, aDNA XX |
| CGG_2_106790 | Grav CA, x139 | 2nd related to CGG_2_106787 | Male aged 45–55 years old, placed in supine position, buried with an iron knife and a dog. A a lump of ocher and a grinding stone probably belongs to this grave, aDNA XY |

| | | | |
|---|---|---|---|
| CGG_2_106791 | Grav DO, x220 | | Placed in supine position, no objects was found in the grave, osteological a mae 45–60 years old, aDNA XY |
| CGG_2_106792 | Grav DS, x297 | | The grave contained a partly preserved child aged 5–6 years of age, aDNA XY |

**Table S7.2.7:** List of the samples studied from Denmark, Svendborg, Langelands Sønder, Lindelse, Kaagården, their relation to each other and summary of the anthropological and archaeological studies.

#### Denmark, Svendborg, Langelands Sønder, Magleby, Nordenbrogård "Hesselbjerg"

Coordinates: 54.7813, 10.7356

A small cemetery containing 5 burials. Grave 3 to 5 was excavated by J. Winter. The sampled burial is grave 5 and is the remains of a young male (age 18–19) buried in hocker position. No artefacts were found in the grave, based on the surrounding graves it probably dates to the Early Roman Iron Age[380].

CGG_2_106773 (LMR C605-21, Grave 5), aDNA XY.

#### Denmark, Svendborg, Langelands Sønder, Tryggelev, Bødgårds mark/ Bødkergårds

Coordinates: 54.8043, 0.6878

The cemetery was excavated by Langelands Museum in 1997. It is a Viking Age cemetery placed on a small hill, and it contains five preserved inhumations. Three of the graves were oriented east-west and these had an iron knife as burial equipment..The two neighbouring burials were in a north-south direction, one of these contained a small ceramic vessel. Some Early Iron Age settlement remains were also excavated.Based on the typology of the artefacts the graves are broadly dated to the 9th century CE. Based on stratigraphy is grave D younger than grave C

The burial D, an adult male, was oriented southwest-northeast in supine position. The cranium was separate from postcranium and found adjacent to the lower part of the torso. The mandible was found upside down in the"original head end". A knife was found underneath the skull. This grave has earlier been analysed in Margaryan et al 2020 (VK291)[405].

Grave H, an(adult male placed on a right side hocker position in a north-south direction with head in north, both arms and legs were bent. A small ceramic vessel (14 cm dm) was placed next to the torso. Burned animal bones and charcoal were found in the fill.This grave has earlier been analysed in Margaryan et al 2020 (VK289)[405].

Samples from the following graves were analysed:

CGG_2_106783 (LMR13372, Grave C), aDNA analysis determined the deceased as XX.

CGG_2_106784 (LMR13372, Grave K, skelton 1), aDNA analysis determined the individual as XX.

#### Denmark, Zealand, Alsted, Simonsborg

Coordinates: 55.4031, 11.6601

Sophie Bergerbrant and Marie Louise Jørkov

A cemetery that was excavated in the 1960 until no more graves could be found. The First excavator was H. Thrane and the second D. Liversage[380,407]. All the graves sampled here have the general museum number NM586/66. The cemetery covered c. 25x32 m, it was placed on a long boulding clay sloop but in an area with sand. The area had been disturbed by sand digging, the people digging for the sand avoided the areas where they thought they would find human bones[407]. Hence the cemetery probably originally contained more burials then are known today. Sand digging is believed to occur from the Middle Ages to modern times and the excavated 37 graves was by the Liversage[407] to be a fragment of the possible hundred of original graves. The graves are dated to the Early Roman period[380,407]. Though the grave dated for this study (grave L24) gives a radiocarbon date to the Viking Age, 770–890 CE. Ten graves from this cemetery were sampled by Melchior et al 2010 for mitochondrial haplogroups, of those only six yielded any genetic results and 22 were sampled by Jørkov (2007; Jørkov et al. 2010) for stable isotope analysis[408], [409]. The graves that have been sampled twice are shown in Table S7.2.8. The other sampled graves were grave 5 (female 35–55) no genetic results and grave 6 (male 25–40) MtHG H[410].

| CGG No | Grave | Sample description | Burial | Grave description | Mito haplotype |
|---|---|---|---|---|---|
| CGG_2_106713 | Grave 17 | Female aged 18–25 | crouched on left side head | one clay pot | MtHG U4a1a |

| | | | | | |
|---|---|---|---|---|---|
| CGG_2_106714 | Grave 21 | Female aged 35–55 | crouched on the right side head N | bronze sewing needle, iron knife, clay spindle whorl | MtHG H (Melchior et al 2010)<br>MtHG H1g1 |
| CGG_2_106715 | Grave 26 | adult/mature | crouched on the right side, head in the N | No grave goods | MtHG H2a2b |
| CGG_2_106716 | Grave 27 | Female mature | crouched on the right side head N | iron rod , 1 clay pot and unburnt clay | MtHg I2'3 |
| CGG_2_106717 | Grave 29 | Child aged 8 | crouched on the right side head N | Part of bronze pin | MtHG H1 |
| CGG_2_106718 | Grave 30 | Male adult/mature | crouched on the right side head N | sherds from clay pot | MtHG U2e2a1 htY I1a~ |
| CGG_2_106719 | Grave 31 | Female adult/mature | lying with flexed legs and head N | bronze sewing needle, possible traces of leather purse | MtHG U4a1a |
| CGG_2_106720 | Grave 34 | Child aged 7 | crouched on right side Head N | No grave goods | MtHG J1c8a htY R1b1a1b1a1a1b |
| CGG_2_106721 | Grave 36 | Male mature | crouched right side head N | eight small stones | MtHG T2b5 htY I1a2a1a |
| CGG_2_106722 | Grave 10 | Female mature/senil s<br>Genetic male | Crouched on the right side head N | Iron knife , iron awl, spindle whorl, two clay pots and unburnt clay | no result in Melchior et al 2010<br>MtHG H1ak hgY R1b1a1b1a1a1b |
| CGG_2_106723 | Grave 13 'løsfund' | Female adult | Disturbed grave | | MtHG U5a1c2a1 |

| | | | | | |
|---|---|---|---|---|---|
| CGG_2_106724 | Grave 19 | Male mature | Disturbed grave - collection of bones - done in historic time | | MtHG J1c4<br>hgY R1b1a1b1a1a |
| CGG_2_106725 | Grave L24 | Female mature | Crouched on the right side head S | Bronze sewing needle UCIAMS-282662, 1940±20 BP, >30kDa collagen yield (%) 1.6, d15N (‰) 10.2. d13C (‰) -20.4, C/N (atomic) 3.2 | MtHG HV9a1 |
| CGG_2_106726 | Grave 32 | Male juvenilis 14–15 | Crouched on the right side head N | Sherds from clay pot | MtHg U2e2a1<br>hgY I2a1b1a2b1a~ |
| CGG_2_106727 | Grave 3 | Female? 16–18 | crouched on the left side | No grave goods | MtHG T2b5 |
| CGG_2_106728 | Grave 2 | Male 35–55 | extended supine position with flexed legs and arms together over the chest | No grave goods | MtHH H1e1a<br>htY R1b1a1b1a1a1b1a |
| CGG_2_106731 | Grave 11 | Osteological Male 30–40<br>Genetical XX | Supine position | band of iron bent in an angle | no result in Melchior et al 2010<br>MtHH H10e<br>Related with CGG_2_106730 |

| | | | | | |
|---|---|---|---|---|---|
| CGG_2_106732 | Grave 14 | Adult female and infant (our sample adult) | The adult held the infant in its arms | No grave goods | MtHG U5b (Melchior et al 2010)<br>HtMG U5b3e |
| CGG_2_106733 | Grave 18 | Female 35–55 | crouched right side head N | No grave goods | MtHG T2b (Melchior et al 2010)<br>MtHG T2b5 |
| CGG_2_106729 | Grave 7 | Female adult | Crouched on the left side head N | Iron knife, iron awl and clay pot | MtHG I (Melchior et al 2010)<br>HtMG I3b |
| CGG_2_106730 | Grave 9 | Male adult | Partly disturbed grave crouched on the right side head Nc | One clay pot | MtHG H (Melchior et al 2010)<br>MtHG H10e, hgY R1b<br>Related with CGG_2_106731 |

**Table S7.2.8:** List of the samples studied from Denmark, Zealand, Alsted, Simonsborg and summary of the corresponding anthropological and archaeological studies.

#### Denmark, Zealand, Bjørnstrupgård, Røsnæs (03.01.05-128)

Coordinates: 55.74, 10.94

Anders Fischer

Pre-Roman Iron Age (200 BCE –10 CE)

Human remains found in a stone set grave (AS_4/68,_NM1_551/68). The grave was excavated in 1968. It contained no objects, only the remains of an individual were found.

CGG_2_019645 (AS_4/68,_NM1_551/68) is a male and has been dated to 2086±39 BP (200 BCE to CE 10) (UBA-35720; $\delta^{13}C$ -19.8, $\delta^{15}N$ 10.7; petrous bone).

#### Denmark, Zealand, Engbjerg, Høje Tåstrup (02.02.07-84)

Coordinates: 55.65, 12.24

Late Roman Iron Age (200–375 CE)

SØL 964 Engbjerg

The burial site was excavated in 1998 by Søllerød Museum[411] and dates to Late Roman Iron Age (200–375CE) and Early Germanic Iron Age. The excavating archaeologist was Linda Boye. The cemetery contained 24 burials of which many are children's burials. The burials with most artefacts all contain females. See Table S7.2.9 for information on the samples included in this study.

| **CGG No** | **Grave** | **Haplotype** | **Grave** | **Sex/Age** |
|---|---|---|---|---|

| | | | | |
|---|---|---|---|---|
| CGG_2_019088 | Grave 3 | C1b2 | Inhumation grave, coffin 2.2x0.6 m, remains of the coffin show that it was made of oak, the deceased had been placed in a supine position. In the area of the head were animal bones, the head had possibly been placed on a pillow, which was indicated by a light colouration in the ground below the head and that the head had been separated from the body during decomposition. A ceramic vessel had been placed by the head and another one by the feet. Outside the coffin in the south side a bone comb had been placed. The comb is the base for the dating. | Sex unknown. Subadult, 16–17 years Related to CGG_2_019091 |
| CGG_2_019090 CGG_2_106821 | Grave 18 | C1b2-C2 | Remains of a oak-coffin, 2.2x0.75 m., contained an inhumation grave. The deceased was placed in a hocker position with the face turned towards east. A hair-pin was found next to the head, on the neck and the torso four fibulas were placed, two amber beads, a glass bead and a gold foil bead. A spindle whorl was placed in front of the face. On the south side of the grave a bone comb and animal bones were found. In the mouth of the deceased had a piece of glass been placed. In the northern end of the grave a ceramic vessel and sheep bones were found. | Female, 25–35 years |
| CGG_2_019091 | Grave 20 | C2 | In the traces of a 2.5x0.72 m coffin an inhumation was found. The deceased had been placed in a supine position. In the neck area a | Male, 20–25 years |

| | | | | |
|---|---|---|---|---|
| | | | blue glass bead was found, by the shin a bone a comb had been placed. By the head aceramic vessel been placed and another one by the feet. | Related to CGG_2_019088 |

**Table S7.2.9:** List of samples studied from Denmark, Zealand, Engbjerg, Høje Tåstrup and context description.

#### Denmark, Zealand, Frederiksborg county, Horns hundred, Kyndby parish, Kyndby (010205-93)

Coordinates: 55.854, 12.129

Johan Zakarias Gårdsvoll & Mads Dengsø Jessen

In 1952 (j.nr. NM1 523/52) and 1953 (j.nr. NM1 470/53) Mogens Ørsnes-Christensen excavated a cemetery with 9 inhumation graves[412], occasionally covered with oval stone pavements. Accompanying dog skeletons in the graves is a recurring phenomenon. One of the graves (not part of this analysis) contained the famous Kyndby sword dated to around 1250 BP. Grave IVa (CGG_2_107519) also had a dog skeleton and a stone cover and was interred with another individual (not sampled) which might have been interred at an earlier point in time. All the graves must be dated to the Late Germanic Iron Age. In 2003 (j.nr. 199/03) the site was revisited by Museum Færgegården (L. Dokkedal og S. A. Sørensen) and there appears to have been a series of fire and cooking pits with burnt animal bones related to the cemetery, perhaps as part of some kind of feasting.

CGG_2_107519 (NM1 523/52, Grave IV A) UCIAMS-282676 1335 BP ±20 (>30kDa collagen yield (%) 2.4, d15N (‰) 10.8. d13C (‰) -19.4. C/N (atomic) 3.2)

#### Denmark, Zealand, Frederiksborg county, Lynge-Frederiksborg hundred, Frederikssund parish, Bilidt (010302-2)

Coordinates: 55.847, 12.062

Johan Zakarias Gårdsvoll & Mads Dengsø Jessen

In 1887–1888 during digging for gravel in a sandy hill at Bilidt several inhumation graves with preserved skeletal remains were disturbed. The finds were reported to the National Museum of Denmark and the site was inspected and a grave (Grave A) was excavated by Andreas Peter Madsen and later in 1888 Vilhelm Christian Boye was also sent to inspect the site after reports of new findings. The skeletal remains of at least 5 individuals in 4 graves have been registered in the sandy hill. One of them (Grave A) found in 1887 was surrounded by smaller stones and contained extraordinarily rich grave goods such as a single-edged sword laying on the chest of the body, sword sheath fittings of tinned bronze and 19 game pieces made from bone. In total more than 20 graves have been found over the years in the sandy hill and in the adjacent area that faces straight out towards Roskilde Fjord. The sword and its sheath's fittings (Grave A) can be dated to around 1320–1350 BP and thereby date the cemetery to the early parts of the Late Germanic Iron Age[412,413].

CGG_2_107386 (NM1 C 6026-28, C 6092-93, Grave A).

#### Denmark, Zealand, Frederiksborg county, Lynge-Frederiksborg hundred, Uggeløse parish, Kirkebakkegård (010313-41)

Coordinates: 55.824, 12.313

Johan Zakarias Gårdsvoll & Mads Dengsø Jessen

In 1965 (j.nr. NM1 522/66), Henrik Thrane excavated a richly furnished inhumation grave. The grave contained gold rings, a golden disc with a face, fibulas, a bronze bucket, glass object, a ladle and sieve of Roman make and pottery, as well as several animal bones from for example sheep and pig. The grave can be dated to the Late Roman Iron Age.

CGG_2_107422 (NM1 522/66).

#### Denmark, Zealand, Frederiksborg county, Lynge-Frederiksborg hundred, Jørlunde parish, Sperrestrup (010306-30)

Coordinates: 55.8103, 12.2749

Johan Zakarias Gårdsvoll & Mads Dengsø Jessen

In 1890, Vilhelm Christian Boye registered some of the finds from 1858–1864 that had been made in a marl pit at Sperrestrup. The finds consisted of a glass drinking vessel and a gold ring, further to the south of these a comb made of bone and a wooden bucket which had been placed beside a human skeleton were found. The finds can be dated to the Late Roman Iron Age.

CGG_2_107410 (NM1 19043).

#### Denmark, Zealand, Frederiksborg, Lynge-Frederiksborg, Frederiksborg Slots, Gadevang Mose (010301-200)

Coordinates: 55.9619, 12.2874

In 1940 during peat digging human remains were found. The police examined the find place and could see that the remains were placed in supine position c. 1.5m under the surface. On the left side of the cranium a round hole can be seen. It was interpreted as a healed trepanation.

CGG_2_107531 (NM 455/40).

#### Denmark, Zealand, Frederiksborg, Ølstykke, Egedal Mose

Coordinates: 55.8042, 12.2081

Rise earlier sample are found close by (RISE274, NM643/44 OxA-30483 2068±25BP, d13C (‰) -19.85) (Allentoft et al. 2015) with a SrI result (0.71143) indicating a local origin (Frei et al 2019).

CGG_2_107534 (NM 771/41) is an older male are dated to the Pre-Roman Iron Age/Roman Iron Age 50 BCE– 60 CE (UCIAMS-272885, 2010±15 BP, >30kDa collagen yield (%) 1.5, d15N (‰) 11.0. d13C (‰) -20.4. C/N (atomic) 3.2).

#### Denmark, Zealand, Frederiksborg, Ølstykke, Veksø parish, Værebro Å, Snydebro

Coordinates: 55.7556, 12.2140

Sophie Bergerbrant

Human remains of at least nine individuals have been found in the vicinity of Snydebro. Remains of at least five humans have been collected by K. Nilesent along Værebro Å, most close to Snydebro, two children and three adults. Remains from another four individuals were found during clearing Værebro Å by Snydebro and given to the museum by E Andersen. The samples individuals sampled for this project dates from the Neolithic. Close to Snydebro has two Late Neolithic axes have been found (1968 & 1969).

CGG_2_105364 (VÆM385, cranium 385) has been dated to the Late Neolithic 2130–1940 BCE (UCIAMS-272871, 3645±15 BP, 30kDa collagen 4.5, d15N 10.2, d13C -20.2, C/N 3.2).

CGG_2_105362 (VÆM559, cranium 559) has been dated to the Middle Neolithic 2560–2340 BCE (UCIAMS-272870, 3935±20 BP, >30kDa collagen 8.2, d15N 10.4, d13C -19.9, C/N 3.2).

#### Denmark, Zealand, Fredriksborg, Lynge-Fredriksborg, Oppe Sundby parish, Sanddal (010310-63)

Coordinates: 55.8130, 12.0810

Katrine Ipsen Kjær

Due to road construction work related to the new Roskilde Fjord-connection, south of the town Frederikssund, an Iron Age burial site was located and excavated in 2015. A total of 61 inhumations were examined, where human bones were preserved in 16 graves. Four of these contained children between the age of 0–6 years old. Both the male and female deceased were placed in the hocker-position in the graves.

The grave goods were relatively poor and consisted of ceramic vessels, a few small iron nails, a small amount of bronze needles, and food offerings[414].

The $^{14}$C-results dates the burial site in the late Early Iron Age to the early Late Iron Age (B2/C1).

Grave A456 (skeleton ROM3306_X186) was placed in a North-South direction with the head placed in the north and had a grave marker. The skeleton was almost fully preserved, it was placed in a hocker position on the right side. Behind the cranium were two ceramic vessels placed. The deceased has been determined as male c. 180 cm tall, aged 35–45 years old[414]. Bone dated by the excavator dates the grave to the Late Roman Iron Age 86–242 CE (Kjær 2019). However, the tooth dated within this project dates the grave to the Early Roman Iron Age (1940±15 BP, UCI-232729). The samples included in this project are listed in Table S7.2.10.

| **CGG No** | **Museum id** | **Sample information** |
|---|---|---|
| CGG_2_019437 | ROM 3306 A237X119 | AAR-25274, 1844±27 BP |
| CGG_2_019438 | ROM 3306 A240X82 | AAR-25275, 1785±29 BP |
| CGG_2_019439 | ROM 3306 A257X77 | AAR-25276, 1939±27 BP |
| CGG_2_019442 | ROM 3306 X186 | UCIAMS-232729 1940±15 BP >30kDa collagen yield (%) 9.9, d15N (‰) 10.9. d13C (‰) -20.0, C/N (atomic) 3.2 andAAR-25279, 1840±29BP. Combined X-Test fails at 5%, X2-Tes: df=1, T=9.3(5% 3.8) 1919±1BP (70–210 CE) |

**Table S7.2.10:** List of samples from Denmark, Zealand, Fredriksborg, Lynge-Fredriksborg, Oppe Sundby parish, Sanddal and their dating.

#### Denmark, Zealand, Fyrendal parish, Skafterup Mark (040503-13?)[407]

Coordinates: 55.2234871, 11.5115596

Johan Zakarias Gårdsvoll & Mads Dengsø Jessen

CGG_2_107501 (NM 20457, Skafterup I).

#### Denmark, Zealand, Gerlev parish, Mærkebækgård Syd (010203-159)

Coordinates: 55.803288, 12.004319

Ole Thirup Kastholm

Due to road construction work related to the new Roskilde Fjord-connection an Iron Age burial place was located and excavated in 2016–2017[414]. A total of nine grave pits holding in all 10 inhumations were examined. The burials covered an area of c 500 m2, and according to the trial excavation results the burial place seems to have been fully examined. The semi-circular layout of the burial place indicates that the burials might have been placed around a burial mound. Though historical maps reveal a small mound or hill at this exact place, any archaeological trace of it was not to be recovered. The site is situated on a small headland in the landscape c. 12 m.a.s.l., stretching south and delimited by the watercourse of Mademose Stream.

A group of five graves had a N-S orientation, and according to pottery finds in one of these this group should be dated to the Roman Iron Age. The rest of the graves were orientated E-W, and these were C14-dated to the Viking Age, more precisely from the late 9th to the early 11th century CE. One of the Viking Age burials (A4417) held two individuals interred at two different events. The secondary individual (grave A14417a) had a small whetstone, an iron knife and beads, while the primary individual (grave A4417b) had an iron knife. The DNA sample is from the latter, which is C14-dated to (2 $\sigma$) 780–980 CE (CGG_02_105350, UCIAMS-233740, 1140±15 BP, >30kDa collagen 6.4, d15N 11, d13C -19.8 C:N 3.2).

CGG_2_105329 (ROM3204, A4417b/x5721) and CGG_02_105350 (A4417b x5721).

#### Denmark, Zealand, Gerlev Strandvej 10 (010203-87)

Coordinates: 55.820214, 12.032846

Jens Winther Johannsen

The two samples come from the southernmost of two burial mounds called Iversbjærgene, which were previously found close to the western shore of Roskilde Fiord. The remains of the mound were excavated after a so-called fishtail flint dagger (of Lomborg's type IV) was found on the site in the late 1990s. The dagger indicates that the burial mound belonged to the last phase of the Late Neolithic. The excavation revealed the last remains of the mound, which seems to have been surrounded by a stone frame. The mound contained an east-west oriented gallery grave covered by a stone package including several shells. The top stones of the cist were missing, but the side stones consisted of slender split stone slabs. Remains of a white limestone pavement were preserved in the floor. The internal dimensions of the cist were 250 x 100 centimetres. The unburned bones of at least three individuals were found scattered in the filling. Fragments of burnt bone, also found in the filling, together with a fragment of a serrated bronze sickle, may represent the remains of a Late Bronze Age cremation.

The bones of individuals 1 and 2 were selected for study:

Individual 1 was identified as a male between 25–35 years old (closer to 25 than 35). The bones were strong and large, and the presence of prominent muscle attachment markings on the knee and heel bones suggest a muscular physique. Mild dental calculus was observed. There was a lack of wisdom tooth development (agenesis) in the preserved mandible.

Individual 2 was probably a male over 40 years old. His bones were not as robust as those of Individual 1, but still somewhat stronger than the unexamined Individual 3. Significant tooth loss (with signs of healing during the individual's lifetime) was observed in the preserved left half of the jaw. There was also evidence of osteoarthritis in the lumbar vertebrae and a finger joint.

The third individual, who was not sampled, had only a fully preserved right humerus, a left kneecap (patella), a left talus (ankle) and a right navicular (foot). These bones were more delicate and smaller than those identified and sorted as belonging to Individuals 1 and 2, suggesting that they belonged to a third, possibly female, individual. The individual was adult, i.e. over 18 years of age, but was probably older (at least over 30 years of age) as the joint surfaces, where they could be observed, were not as sharp as in younger individuals. Based on the humerus, the stature is estimated to have been between 154 and 157 centimetres.

A small stone cist was found just outside the stone frame of the mound. Nothing was found in it. and function remains uncertain.

CGG_2_106513 (Indiv 1 MFG 114/97) has been dated to 3560±20 BP (UCIAMS-272876, >30kDa collagen yield (%) 5.1, d15N (‰) 9.9. d13C (‰) -20.0, C/N (atomic) 3.2 (2010–1770 BCE)).

CGG_2_106514 (Indiv 2 MFG 114/97).

#### Denmark, Zealand, Holbæk county, Ars hundred, Store Fuglede parish, Kalmergården (030106-137)

Coordinates: 55.56924, 11.2511

Johan Zakarias Gårdsvoll & Mads Dengsø Jessen

The initial grave, the west-grave, is dug into a shallow, natural depression about 45m north of the old crossing of the Halleby Stream. The bottom of the grave is level and rests on the sterile, clayish subsoil. The interred is placed in supine position, on his back with straight legs – the right might have been slightly bent – and the arms are situated along the body. Between the legs the decapitated head could be found with the face towards the crotch. Consequently, the placing of the person seems quite orderly and the grave must have been backfilled just after the decapitation. Soon after, a large stone was erected on top of the grave. There can hardly be any doubt that the stone and the content underneath it was known to the wider public, when the second burial took place (see below). The entire skeleton in the west-grave had male attributes, he was between 35–50 years of age at the time of death, and approx. 171 cm in height. He had only moderate tooth-wear and was in general good health with almost no osteoarthritis, and had an unusually large skull of approx. 1.700 $cm^3$. He did however have enamel hypoplasia, indicating that he at a young age had suffered from malnutrition or some kind of severe illness. He had a distinct underhung jaw which would have made him very recognisable. The cut to his neck hit right on his second cervical vertebra and just glanced the lower part of his jawbone. It cut the head clean off.

The second grave, the east-grave, clearly had a more careless character to it. The grave pit itself is slanting toward the large stone and the body is almost entirely placed in a natural depression, wherefore only the upper part of the corpse rests in clayish subsoil. As if almost thrown into the pit, the body is placed on its side with the arms flung across the breast and the legs, especially the right leg, bent towards the torso. The head was severely damaged by the decapitation and the cranium rested by the knees while the jaw, which had been separated by the killing blow, was placed by the neck. Presumably as a result of it still being attached to the facial skin. Seemingly, the grave-pit was dug close to the large stone, almost under it, and appears very narrow, with hardly any space to contain the body. Placed on the side the skeleton appears as would a body falling down a hillside indicating a casual and hasty handling of the decapitation as well as the interment. The skeleton has been determined as a male that at the time of death was between 21–35 years old, and approx. 173 cm in height. All teeth were present, but the inside of the upper incisors had a visible flat wear, which could indicate them being used for a special purpose and as a ’tool’. The bones appeared normal except for a fusion of the right hip bone and the sacrum resulting in a deterioration of the entire hip and femur and a completely stiff right leg. The leg would have pointed outwards in an almost 90-degree angle. Consequently, the person was severely impaired and would have had an acute limp if he could even walk at all. The top of the skull was cut off from behind in a line from the topmost vertebra and out basically in the middle of the nose. Of course the blow would have killed instantly, but the execution was either less professional or the man would have pulled his chin back at the last second. As a consequence, the jaw was left hanging by the skin of the chin, and would presumably have been a rather macabre sight to behold.

During the examination of the skeletal remains it was confirmed that both individuals were beheaded and as both cut surfaces had very sharp edges the decapitation must have been carried out with a very sharp tool. In either case, the decapitation was pre-mortem and most likely the cause of death as no other trauma was detected. Furthermore, both cuts were singular and effective thus indicating orchestrated and controlled beheadings[415–417].

CGG_2_107579 (NM1 2654/79, East, beheaded half).

CGG_2_107580 (NM1 2654/79, West).

#### Denmark, Zealand, Holbæk county, Ars hundred, Ubby parish, Udby (030109-62)

Coordinates: 55.615936, 11.2187132

Johan Zakarias Gårdsvoll & Mads Dengsø Jessen

In 1905 (j.nr. NM1 195/05) in a sandpit with disturbed graves, H. Kjær, The National Museum, inspected and excavated the remains of at least 5 inhumation graves on the northern side of a small hill, of which 3 contained grave goods. The pottery could be dated to the Early Roman Iron Age. Earlier a cranium (Cranium C) had been found at the same site.

CGG_2_107505 (NM1 C 12894, Cranium C).

#### Denmark, Zealand, Holbæk county, Merløse hundred, Grandløse parish, Store Grandløse (030302-40)

Coordinates: 55.6819295, 11.7297839

Johan Zakarias Gårdsvoll & Mads Dengsø Jessen

In 1903 (j.nr. NM1 158/03), H. Kjær, The National Museum, excavated 3 inhumation graves containing 3 individuals, with several different types of grave goods. Grave C (CGG_2_107503) of the present study contained a pottery vessel from the Early Roman Iron Age.

CGG_2_107503 (NM1 C 11874, Grave C).

#### Denmark, Zealand, Holbæk county, Merløse hundred, Holbæk parish, Tveje Merløse (030303-28)

Coordinates: 55.656, 11.631

Johan Zakarias Gårdsvoll & Mads Dengsø Jessen

In 1951 (j.nr. NM1 330/51) during gravel digging in a pit several graves were disturbed. The find was reported to The National Museum of Denmark and Hans Norling-Christensen inspected and excavated 6 inhumation graves forming a small cemetery, and again in 1961 (j.nr. NM1 723/61) Mogens Ørsnes-Christensen inspected a single grave there. The total number of graves is 6–7. Grave 4, probably containing an adult male, has been carbon dated to 1710±70 BP (240 e.Kr. C-14; 275 e.Kr. Cal. Sample nr.: Tveje Merløse, Gr. 4). Grave 5 has been sampled for the present study.

CGG_2_107431 (NM1 330/51, Grave 5).

#### Denmark, Zealand, Holbæk county, Merløse hundred, Stenlille parish, Stenlille (030309-7A)

Coordinates: 55.539, 11.586

Johan Zakarias Gårdsvoll & Mads Dengsø Jessen

In 1914 (j.nr. NM1 43/14) T. Thomsen inspected a gravel pit at Stenlille in which 2 inhumation graves with Roman imports and preserved textiles and skeletal material had been disturbed. Later in 1952 (j.nr. NM1 710/52) Hans Norling-Christensen conducted an excavation at the site. Among the finds were an 'Østlandskedel' dating the grave material to the later parts of the Early Roman Iron Age.

CGG_2_107502 (NM1 C 16396-400).

#### Denmark, Zealand, Holbæk county, Ods hundred, Fårevejle parish, Fårevejlegård (030403-504)

Coordinates: 55.8851, 11.4383

Johan Zakarias Gårdsvoll & Mads Dengsø Jessen

In 1977 and 1980 (j.nr. NM1 2080/77) Per Poulsen, the National Museum, excavated 6 inhumation graves of 4 adults and presumably 2 children. The graves were oval and the interred were all placed in hocker position. The grave form dates the graves to the Roman Iron Age.

CGG_2_107533 (NM1 2080/77, Grave E).

CGG_2_107532 (NM1 2080/77, Grave F).

#### Denmark, Zealand, Holbæk county, Tuse hundred, Hørby parish, Baunehøj (030705-29)

Coordinates: 55.710, 11.672

Johan Zakarias Gårdsvoll & Mads Dengsø Jessen

In 1938 (j.nr. NM1 523/38) in a sandy hill near the top of Baunehøj at the headland of Tuse an inhumation grave under a flat field with a preserved cranium was disturbed. The find was reported to the National Museum of Denmark and Carl Johan Becker was sent to inspect the site. Next to the cranium a ceramic vessel was found, and it is possible that both stemmed from the same disturbed grave. The typology of the ceramic vessel and the context in which the cranium was situated suggest that the date of burial occurred in the Late Roman Iron Age.

CGG_2_107468 (NM1 523/38).

#### Denmark, Zealand, Holbæk county, Tuse hundred, Hørby parish, Hørby Skov (030705-41)

Coordinates: 55.7381625, 11.6452876

Johan Zakarias Gårdsvoll & Mads Dengsø Jessen

In 1898 (j.nr. NM1 639/98) in a sand pit 2 inhumation graves were excavated by C. Fredstrup. The pottery grave goods can be dated to the Early Roman Iron Age.

CGG_2_107484 (NM1 C 9289-90).

#### Denmark, Zealand, Holbæk county, Tuse hundred, Kundby parish, Gedebjerg (030707-223)

Coordinates: 55.70024, 11.50478

Johan Zakarias Gårdsvoll & Mads Dengsø Jessen

In 1960 (j.nr. NM1 473/62), Hans Norling-Christensen excavated a single inhumation grave in a gravel pit. The grave was 1.8 metres deep and contained a single individual with a single, ornamented vessel. The vessel can be dated to the Early Roman Iron Age.

CGG_2_107447 (NM 473/60, C27446, Skeleton 1).

#### Denmark, Zealand, Holbæk, Løve, Bakkendrup, Bakkendrup

Coordinates: 55.563, 11.249

A cemetery partly placed under an old road has been excavated two times. The cemetery was damaged and stray human bones discovered at the surface. The inhumations date to the Late Iron Age and the Viking Age. During the first excavation 1979 ten burials and bones from another minimum seven individuals were recovered[380]. In 1983 another five burials were excavated at the site. Three samples from this site have earlier been analysed for their DNA, two of the surface found bones and grave 16[405]. The samples analysed in this study are listed in Table S7.2.11.

| **CGG No** | **Grave id** | **Sample description** | **Additional information** | **Time period** |
|---|---|---|---|---|

| | | | | |
|---|---|---|---|---|
| CGG_2_106483 | FFF F54-586 AS 59/79, Lavo H, Grave 9 | Oriented NW-SE, the deceased was placed in hocker with head in NW Female adult Genetically XX | Bronze disk, 41 glass beads, mone amber bead, four rock crystal beads and two iron knives with wooden handles | Viking Age |
| CGG_2_106826 | Grave 10 (F54-586) | NW-SE oriented grave with the individuals placed in supine position, male maturus Genetically XY | Bronze buckle, iron knife with wooden handle and unidentified iron object | Viking Age |
| CGG_2_106827 | Grave 16 (RAS 45/83) | Genetically XY Earlier analysed (VK315) | | |

**Table S7.2.11:** List of the samples studied from Denmark, Zealand, Holbæk, Løve, Bakkendrup, Bakkendrup and summary of the corresponding anthropological and archaeological studies.

#### Denmark, Zealand, Holbæk, Løve, Reerslev parish, Almager (030210-47)

Coordinates: 55.79016, 12.29043

Sophie Bergerbrant

A small cemetery containing four probable inhumation graves of which one (grave B) was found by workers while digging for sand and one (grave A) was excavated by H. Kjær in 1928 (Liversage 1980: 16).

Grave A (CGG_2_107467 (NM C17361) is dated to the Early Roman Iron Age based on the related graves as it contains no objects[407]. The grave was oriented southwest-northeast with the head in the south. The individual buried in the grave is determined to be an adult male[380].

#### Denmark, Zealand, Holbæk, Løve, Ruds Vedby, Lyngmose (030211-27)

Coordinates: 55.5334, 11.3455

In Lyngmose were human and animal bones found while peat digging 1933–1934 (NM 503/34).

CGG_2_107526 (Skeleton I) is a male of 25–30 years old and has been dated to 3770–3640 BCE, i.e. the Early Neolithic, (UCIAMS-275223, 4930±20 BP (>30kDa collagen 3.2, d15N 7.2, d13C -21.0 C/N (atomic) 3.2)).

#### Denmark, Zealand, Holbæk, Merløse, Stenlille, Sandlyngmose (030309-106)

Coordinates: 55.57171, 11.55200

Anders Fischer

In 1949 the remains of a human were found while peat digging.

CGG_2_107524 (NM 512/49) is a female of 35-40 years old and has been dated to the Germanic Iron Age 400–550 CE (AAR-11351, 1620±29 BP; femur sin.).

#### Denmark, Zealand, Holbæk, Ods, Asnæs parish, Asnæs (030401-242) and Asnæs Gymnasiums Sportsplads (030401-406)

Coordinates: 55.8150, 11.5029 (coordinates for Asnæs)

Two cemeteries from the Roman Iron Age found in Asnæs found c. 600 metres from each other. Sb No 242 (NM231/74) was excavated by S. Thorsen in 1974–1975, skeletal remains have been found in 19 graves [380]. The samples examined in this study are listed in Table S7.2.12. The cemetery was disturbed by ploughing and has probably been fully excavated. The cemetery contains both inhumation graves and cremation graves (Fund og fortidsminder 030401–242). Sb 406 (NM 3313/80) was excavated by L Jørgensen in 1980 and held 13 graves[380]. Both cemeteries date to the Early Roman Iron Age.

| **CGG No** | **Museum id** | **Sample description** | **Grave finds** |
|---|---|---|---|
| CGG_2_107388 | NM 3313/80, Gave 10 | Possible female aged 16–25, crouched right side, head in S | Two bronze broaches,a bronze pin,one clay pot and animal bones |
| CGG_2_107390 | NM 3313/80, Gave 18 | Young individual 16–19, crouched right side head in N | No artefacts found |
| CGG_2_107393 | NM 3313/80, Gave 46 | A possible male aged 45+, crouched right side, head in S | No artefacts found |

| | | | |
|---|---|---|---|
| CGG_2_107389 | NM 3313/80, Grave 17 | An individual aged 35?, crouched right side, head in S | No artefacts found |
| CGG_2_107391 | NM 3313/80, Grave 22 | Female aged 45+, crouched right side, head in S | No artefacts found |
| CGG_2_107392 | NM 3313/80, Grave 31 | Male? 20–30, crouched right side, head in S | No artefacts found |
| CGG_2_107441 | NM P231/74, Grave O | Male? Aged 20–30, crouched left side head n S | No artefacts found |
| CGG_2_107442 | NM P231/74, Grave P | Female? 40+, crouched left side head n S | One clay pot and some smaller stones |
| CGG_2_107443 | NM P231/74, Grave R | Female? 20–30, crouched on the left side, head in S | No artefacts found |
| CGG_2_107444 | NM P231/74, Grave S | Female 18–25, disturbed grave with head in S | One clay pot and some smaller stones |
| CGG_2_107445 | NM P231/74, Grave T | Female 45+, crouched left side head n S | One iron knife, one iron pin, thunder stone and a few smaller stones |

**Table S7.2.12:** List of samples from Denmark, Zealand, Holbæk, Ods, Asnæs parish, Asnæs (030401-242) and Asnæs Gymnasiums Sportsplads and context description.

#### Denmark, Zealand, Holbæk, Ods, Fårevejle parish, Bøgebjerg

Coordinates: 55.8057, 11.4311

An inhumation grave that was excavated 1928 by G. Raklev, the inhumation was found on the top of a rise[407].

CGG_2_107446 (NM C20369-71) is a mature male placed in a southeast.northwest oriented grave. The skull was placed in the south and the body was placed in a crouched position on the left side[380]. The grave contained an iron knife, a polishing stone, one clay pot and hind bones from a lamb. The grave is dated to the Early Roman Iron Age[407].

#### Denmark, Zealand, Holbæk, Ods, Højby parish, Trundholm Mose (030405-331)

Coordinates: 55.8837, 11.5800

Sophie Bergerbrant

During work at the former bog/mire area Trundholm Mose in 1951 human remains were found. The find is c. 500 metres south east of the famous find of the Trundholm wagon. In a different part of the wetland c. 3.5 km northwest from the sampled find was 1940 human remains found while ploughing. On of those skeletons was analysed within the Rise project (RISE276) which was most likely a male of local origin (SrI 0.710053) that had MtDNA haplogroup T2 b and is was dated to Late Bronze Age, OxA-30485, 2525±25BP (Frei et al 2019) d13C (‰) -18.86.

CGG_2_106734 (784/51 skelet I) is a male that is dated to the Pre-Roman Iron Age 370-190 BCE (UCIAMS-243263, 2210±15 BP, d15N(‰) 11.1, d13C(‰) -20.3).

#### Denmark, Zealand, Kelleklinte Mose, Ubby (03.01.09-263)

Coordinates: 55.62, 11.26

Anders Fischer

Pre-Roman Iron Age (380–197 BCE)

In a bog, measuring c. 450 by 300 m, a local farmer found part of a human skull during peat digging in 1944. The material salvaged from the bog also included bones of swan and (pond) tortoise. With the assistance of a local doctor of medicine, these finds were soon given to the Zoological Museum at the University of Copenhagen. A *pars petrosa, dex.* from the human skull was sampled.

CGG_2_022103 (ZMK 8.2.1944, AL 37:23) has been dated 2215±26 BP (UBA-37911).

#### Denmark, Zealand, Kongsted Lyng (Store Lyng), Kongsted (05.03.05-35)

Coordinates: 55.23, 12.06

Anders Fischer

Viking Age (895–1024 CE)

During peat extraction in a bog, c. 1000 by 350 m, items of presumably prehistoric date were revealed on several occasions: a trephinated human skull (Fig. S7.1), a cranium of a 7–8 years-old child, a ceramic vessel of Neolithic type, a buckle of bronze of Iron Age type, a wooden wheel and a wooden shaft, the latter two possibly also dating to the Iron Age[418]. Tooth 8+ of the first mentioned skull was sampled for the present project.

CGG_2_019689 (NM 547/42, NEO787) has been dated twice: 981±37 BP (UBA-37894; $\delta^{13}C$ -19.7, $\delta^{15}N$ 12.4) and 1127±29BP (UBA-38237; dentine, insufficient for measuring $\delta^{13}C$ and $\delta^{15}N$) $^{14}C$ years BP. R_Combine 1072±23 BP.

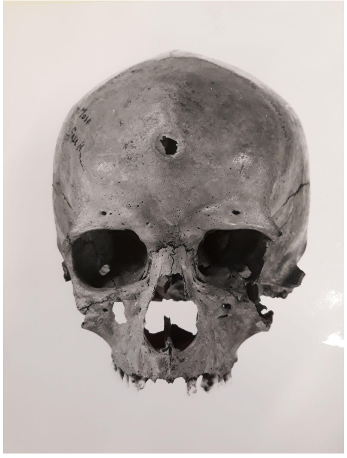

**Fig. S7.1:** Trephinated skull, revealed by peat extraction in Kongsted Lyng 1942. Photo: Laboratory of Biological Anthropology, UCPH.

#### Denmark, Zealand, København, Lejre

Coordinates: 55.6170, 11.9694

A cemetery containing 49 graves dating to the Viking Age[380]. The site was excavated by the National Museum 1953 to 1968. Four graves have been sampled for aDNA for an earlier study; VK90 Denmark_Lejre Grav 902, VK92 Denmark_Lejre Grav 935, VK247 Denmark_Lejre Grav 804, VK385 Denmark_Lejre Grav 321[405].

CGG_2_106830 (Grave 281) the deceased were placed in supine position. In the burial was also an iron knife found. Osteological the grave was determined to be an adult/maturus probable female, the aDNA result was XX.

#### Denmark, Zealand, København, Smørum, Sengeløse parish, Lille Vasby (020211-50)

Coordinates: 55.6691, 12.2092

A small fully excavated cemetery. The burial ground was excavated in 1964 by D. Liversage[407]. The burial ground was placed above what has been interpreted as a Late Neolithic ritual place, containing scattered human bones[407]. The cemetery contains 19 inhumation graves of which six are dated to the Early Roman Iron Age; the remains are either undated or belong to the Late Neolithic[380]. Grave 16, skeleton 72 is radiocarbon dated to the Early Roman Iron Age (1–120 CE). See Table S7.2.13 for a list of samples investigated in our study.

| **CGG No** | **Museum id** | **Sex/Age** | **Sample description** | **Period** |
|---|---|---|---|---|
| CGG_2_107464 | NM 460-64, 2 | Na/30–40 | UCIAMS-282674 3710±20 BP (>30kDa collagen yield (%) 1.3, d15N (‰) 9.5. d13C (‰) -20.5. C/N (atomic) 3.2) 2200–2030 BCE | Late Neolithic I |
| CGG_2_107466 | NM 460-64, 35 | Na/16–20 | | Late Neolithic ? |
| CGG_2_107465 | NM 460-64, 49 | Na/20–30 | UCIAMS-282675 3700±20 BP (>30kDa collagen yield (%) 2.3, d15N (‰) 8.3. d13C (‰) -20.7. C/N (atomic) 3.2) 2200– | Late Neolithic I |

| | | | | |
|---|---|---|---|---|
| | | | 2030 BCE | |
| CGG_2_107463 | NM 460-64, 8 | Female/ 20–30 | | Late Neolithic? |
| CGG_2_107453 | NM 460-64, Grave 1, Skeleton 7 | Male/40+ | A grave that probably was disturbed in the Iron Age, interpreted by the excavator as as remains of LN ritual activity | Late Neolithic? |
| CGG_2_107456 | NM 460-64, Grave 12, Skeleton 61 | Female/ 40 | No artefacts in the grave | Early Roman Iron Age |
| CGG_2_107457 | NM 460-64, Grave 13, Skeleton 59 | Female/45, | crouched on left side spindle whorls (clay),bronze sewing needle,1 clay pot | Early Roman Iron Age |
| CGG_2_107458 | NM 460-64, Grave 15, Skeleton 67 | 13–16 | No artefacts in the grave | Early Roman Iron Age |
| CGG_2_107459 | NM 460-64, Grave 16, Skeleton 72 | Male/40–50 | Crouched on the right side. One clay pot | Early Roman Iron Age (1–120 CE) UCIAMS-275220 19655±15 BP (>30kDa collagen yield (%) 2.6, d15N (‰) 10.1. d13C (‰) -19.6. C/N (atomic) 3.2) |
| CGG_2_107460 | NM 460-64, Grave 17, Skeleton 19 | Female/ 35–45 | No artefacts in the grave | Early Roman Iron Age |
| CGG_2_107461 | NM 460-64, Grave 19, Skeleton 92 | Female?/25–40 | Crouched on the side. No artefacts in the grave | Early Roman Iron Age |

| | | | | |
|---|---|---|---|---|
| CGG_2_107462 | NM 460-64, Grave 21, Skeleton 90 | | Crouched on the left side oriented NE-SW. Bronze sewing needle, 1 clay pot | Early Roman Iron Age |
| CGG_2_107454 | NM 460-64, Grave 4, Skeleton 26 | Male/ 55+ | Crouched on the right side. Sherds from clay pot | Early Roman Iron Age |
| CGG_2_107455 | NM 460-64, Grave 9, Skeleton ? | Male?/25–40 | No grave No 9 exists so this is probably from grave 10 or 11 which dates to the LN | Late Neolithic? |

**Table S7.2.13:** List of samples studied from Denmark, Zealand, København, Smørum, Sengeløse parish, Lille Vasby, their context description and dating.

#### Denmark, Zealand, København, Smørum, Værløse parish, Værløse (Alugod) (020216-34)

Coordinates: 55.76537, 12.32188

Sophie Bergerbrant

A small cemetery consisting of three inhumation graves. The graves were found and inspected in 1944[380]. For this project the only grave with preserved human bones (grave I) was sampled. Grave one orientated east-west and contained a bronze ladle, and strainer set, two silver brooches, silver pin, two glass beads, shards of two ceramic vessels, and a now lost bronze vessel. The grave continued the remains of an adult female [380](139).

CGG_2_105397 (NM260/44) has been dated to the Roman Iron Age 120–310 CE, 1830±20 BP, (UCIAMS-273013, >30kDa collagen yield (%) 1.0, d15N (‰) 10.6. d13C (‰) -20.5, C/N (atomic) 3.2).

#### Denmark, Zealand, København, Sokkelund, København, Kongens Nytorv

Coordinates: 55.6804, 12.5858

Marie Louise Jørkov

Two well preserved skulls with associated mandibles and a loose mandible were found in the early 1940's during the construction of bomb shelters at Kongens Nytorv in front of Magasin du Nord. The skulls were found by digger Eigil Nielsen and handed over to Eigil Knuth. They were later transferred to Copenhagen Museum in 1992 and ended up at the Anthropological Collection at Copenhagen University shortly after. The date of the skulls are unknown but could possibly be from the Medieval Period. Individual 1 which was included in this study was genetically determined to be female (XX) and osteologically aged to be ca. 30–45 years old.

CGG_2_107536 (AS11/92, Skeleton I).

#### Denmark, Zealand, København, Tune, Karlstrup, Karlslunde (020504-4)

Coordinates: 55.5528, 12.2088

Sophie Bergerbrant

A mound excavated by Christian Leif Vedbæk 1965-66[419]. Four gallery graves and twelve inhumation graves were found during the excavation; at least 31 individuals had been buried in the mound[419–421]. The skeletal material was sent to the *Antropologisk Samling* before the final numbering of the graves were decided. Therefore, the grave numbers in AS's catalogue and Vedbæk's report are not correlated. A key to the renumbering in tAS archive was used to ensure that the right sample is correlated to the right archaeological context. The grave numbers for each grave are also different in the catalogue of Aner and Kersten (1973) than in Vedbæk's report. In Aner and Kersten (1973) all graves are numbered from A and onwards, while Vedbæk's report uses Roman numerals.

The mound hence contains remains from the Late Neolithic to Early Bronze Age (PII). One grave has previously been sampled for both Strontium Isotope[269] and for genetic data[422].

Nine individuals were sampled for this project, however, only six yielded enough genetic data to be included in this study (Table S7.2.14). Remains from two Gallery graves (Nos II and III, Aner & Kersten B & C)) and from the double grave XVI (Aner & Kersten grave Q) were sampled (see 20) table).

CGG_2_106702, CGG_2_106704, CGG_2_106705, CGG_2_106706, CGG_2_106707 and CGG_2_106708 are included within this study.

The radiocarbon dates (c.2040 to 1880 BCE) from Gallery Grave II indicate that it was used during Late Neolithic I, Based on the radiocarbon dates (2140–1760 BCE) Gallery Grave III were used during a longer time span. Gallery Grave II contained no artefacts, while Gallery Grave III held a flint dagger (Lomborg type VIB), two amber beads and two bone awls[419,421] so no archaeological comparison between the content of the gallery graves can be done.

Gallery grave II contained the remains of four individuals: two children, one c. 8 years old and one c. 10 years old, a person aged 14–15 years and an older adult. Only a few fragments of the older adult were present[421,423]. Today there are five individuals related to this grave in AS. Two of those are adults and the remaining are subadults of which have been determined to be around 5–6 years old, 7–8 years old and just as a subadult. No objects were found in the gallery grave[419,424].

Gallery grave III contained remains of five individuals (Vedbæk 1987: 23; Ebbesen 2007:44). According to Balslev Jørgensen (1965) they were children aged 2–3, a 12-year old, a person aged 14–15, an 18-year-old and an older adult. The 18-year-old was the last person placed in the grave. Balslev Jørgensen (1965) argues that this person had an old break in the back, giving the person a humpback. The latter has not been observed in later osteological examinations[406,425]. The grave also contained a flint dagger (Lomborg type VIB), two amber beads and two bone awls [419,421,424]. Today there are five skulls present in AS and some other human bones.

A number of inhumations probably from the Early Bronze Age period I and II are included in the mound. However, the bone material from these were preserved in a way to be included in this study. Some of these graves contained no artefacts but some indicate that wealthy males were interred. The final grave in the mound is the double grave XVI. It contained the remains of two individuals: Skeleton A (RISE 20) and Skeleton B (RISE 21). Skeleton A was found with the head in SSE, connected to the remains were a sword, two fibulas, three double buttons (one with an inlaid amber disc), a razor, a bronze knife, two bronze fish hooks (including one with remains of rope), a tweezer, a small gold piece and a flint-strike-a-light small balls of yarn or string and wooden sticks[419,420]. Skeleton B was found with the head in NNV with a razor, an awl, a small piece of bronze, a flint-strike-a-light, a piece of pyrite and some small pieces of worked wood associated with the deceased[419,420]. A previous aDNA analysis determined the skeleton (RISE 21) to be that of a male, with a: HG N1a1a1a2. Today in AS there is also a mandible (RISE 19) of a child connected to this grave. There is no information of where this was found. Therefore, it is difficult to say if this is part of the grave attire or just part of the fill. A radiocarbon date demonstrates that the mandible from the child predates the two adult burials (1510–1420 cal BC). Skeleton A dates to 1510–1290 cal BC and skeleton B to 1430–1280 cal BC, indicating that the double burial probably occurred around between 1400 –1300 cal BC.

Strontium isotope values show that all three individuals analysed within the RISE suggest nonlocal origin, but they seem to come from different areas. The individual with the highest 87Sr/86Sr value, RISE 20, had an unusually low δ18O value (-7.37 ‰) for analysed Danish samples (Frei et al 2019). The other two individuals had δ18O values between -3 to -5 ‰ (carbonate values VPDB) which strengthens the hypothesis that they come from different areas (Frei et al 2019).

The objects in the grave are all of Nordic style, possibly indicating that the deceased is from another area of the Nordic region. There is one similar bronze double button in Billegravsgård, Bornholm [420], which could indicate a connection to Bornholm, although there are many other possibilities.

| Analysed skeleton | Lab nr C14 | $^{14}$C | ± | δ$^{13}$C | d$^{15}$N | C% | C: N | Sampled material | Osteological age and sex | CGG No |
|---|---|---|---|---|---|---|---|---|---|---|
| Gallery grave II | | | | | | | | | | |
| Cranium A | UCIAMS -243256 | 3635 | 20 | -19.817 | 10.6 | 42.0 | 3.2 | Tooth 7+ | Adult, unknown | CGG_2_106705 |
| Cranium B | UCIAMS -243257 | 3620 | 15 | -20.352 | 10.9 | 43.7 | 3.3 | RPP | Adult, unknown | CGG_2_106706 |
| Cranium C | Not collagen enough for C14 | - | - | - | - | - | - | RPP | Subadult | CGG_2_106707 |
| Cranium possibly C | UCIAMS -243258 | 3645 | 15 | -20.4 | 10.9 | 43.7 | 3.3 | RPP | Subadult 8–10 | CGG_2_106708 |
| Gallery grave III | | | | | | | | | | |
| Cranium C | UCIAMS -243253 | 3520 | 15 | -20.094 | 9.8 | 41.9 | 3.2 | LPP | Subadult | CGG_2_106702 |
| Cranium D | UCIAMS -243254 | 3665 | 15 | -20.279 | 11.2 | 44.6 | 3.3 | LPP | Adult, male | CGG_2_106703 |
| Cranium E | UCIAMS -243255 | 3640 | 15 | -19.756 | 10.6 | 43.5 | 3.2 | Tooth -7 | Adult, unknown | CGG_2_106704, CGG_2_107471 |
| Grave XVI | | | | | | | | | | |

| | | | | | | | | | | |
|---|---|---|---|---|---|---|---|---|---|---|
| Skeleton A | OxA-28045 &<br>OxA-28046<br>Com.d | 3162<br>&<br>3113<br>3138<br>701- | 27<br>&<br>27<br>20 | -19.6<br>&<br>-19.5<br>X2-Test: df=1 T=1.6( 5% 3.8) | | 42.7<br>&<br>42.8 | 3.1<br>&<br>3.1 | Tooth -7 | Adult, male | CGG_2_010393 (RISE<br>&<br>CGG_2_106701 |
| Skeleton B | OxA-28047 | 3092 | 29 | -19.8 | | 46.5 | 3.1 | Tooth | Adult, male | CGG-2 010394 (RISE 21)<br>Mt HG N1a1a1a2 |
| Just mandible | OxA-28160 | 3196 | 25 | -19.3 | | 41.9 | 3.4 | tooth | Subadult c. 12 | CGG-2 010392<br>(RISE 19) |

**Table S7.2.14:** List of samples studied from Denmark, Zealand, København, Tune, Karlstrup, Karlslunde, isotopic analysis, $^{14}$C dating and summary of the anthropological studies.

#### Denmark, Zealand, København, Volborg, Allerslev parish, Allerslev (020601-80)

Coordinates: 55.6044, 11.9796

The grave was found by workers while digging for gravel. The grave was investigated by P.V. Glon 1939. The grave was oriented north-south and the remains (CGG_2_107387, NM 551/39) was found with a silver brooch and a gold finger ring. The remains were determined to be an adult/mature male[380].

#### Denmark, Zealand, København, Voldborg, Gevninge Parish, Gevninge Overdrev (020603-59)

Coordinates: 55.626, 11.939

An inhumation grave excavated in 1954, found in a sandpit. The grave contained no preserved objects.

The sample CGG_2_107535 (NM 349/54) is a male aged of 25–35 years old. The grave is radiocarbon dated to Pre-Roman Iron Age/ Early Roman Iron Age 100 BCE–60 CE, UCIAMS-282677 2030±20 BP (>30kDa collagen yield (%) 2.6, d15N (‰) 10.8. d13C (‰) -20.3. C/N (atomic) 3.1).

#### Denmark, Zealand, Københavns county, Ramsø hundred, Højelse parish, Ølby (020105)

Coordinates: 55.47386, 12.16255

Johan Zakarias Gårdsvoll & Mads Dengsø Jessen

A single skull found with a comb and a ceramic vessel handed over by a private visitor to the National Museum of Denmark. Information is limited, but the finds date the skull to the Roman Iron Age.

CGG_2_107415 (NM2 14295).

#### Denmark, Zealand, Københavns county, Sokkelund hundred, Gladsakse parish, Højgård (020303-30)

Coordinates: 55.7336, 12.4591

Johan Zakarias Gårdsvoll & Mads Dengsø Jessen

In 1905 (j.nr. NM1 19/05), a single inhumation grave was excavated by local administrator T. Thomsen. The grave contained several grave goods, mostly pottery, but also a Swastika-shaped fibula. The grave is from the Late Roman Iron Age.

CGG_2_107401 (NM1 C 12920-22).

#### Denmark, Zealand, Københavns county, Sokkelund hundred, Hvidovre parish, Hvidovre (020305-3)

Coordinates: 55.650, 12.467

Johan Zakarias Gårdsvoll & Mads Dengsø Jessen

In 1933 (j.nr. NM1 334/33), Mogens Bellmann Mackeprang excavated 8 inhumation graves in a farm field. Grave A (CGG_2_107483) contained a bronze fibula and silver pin, and other silver fragments. The fibula (Almgrens group V) places the graves in the Early Roman Iron Age.

CGG_2_107483 (NM1 334/33, Grave A).

CGG_2_107482 (NM1 334/33, Grave H).

#### Denmark, Zealand, Københavns county, Sokkelund hundred, Hvidovre parish, Landlystvej 54-56 (020305-8)

Coordinates: 55.6602166, 12.4743842

Johan Zakarias Gårdsvoll & Mads Dengsø Jessen

In 1940 (j.nr. NM1 723/52) H. N. Christensen and H. N. Kunwald excavated 8 inhumation graves with several bronze fibulas and a gold ring, as well as some pottery. The graves are dated to the Early Roman Iron Age by the grave goods.

CGG_2_107452 (NM1 723/52, 641/40, C 24174-76, Grave 4).

#### Denmark, Zealand, Københavns county, Sømme hundred, Himmelev parish, Kildehus (020405-44)

Coordinates: 55.6187, 12.0193

Johan Zakarias Gårdsvoll & Mads Dengsø Jessen

In 1947 (j.nr. NM1 976/47) a group of local workers from a gravel pit brought in skeletal material from 4 inhumation graves (a minimum of 3 adult and 1 child) and the pottery grave goods. The pottery can be dated to the Early Roman Iron Age. An additional 12 skeletons presumably have been excavated in the same gravel pit.

CGG_2_107485 (NM1 976/47, Skeleton 1).

#### Denmark, Zealand, Københavns county, Tune hundred, Karlslunde parish, Mosede Fort (020504-14)

Coordinates: 55.5644395, 12.2745516

Johan Zakarias Gårdsvoll & Mads Dengsø Jessen

In 1913 (j.nr. NM1 277/13) a team from the National Museum (Kjær, Rosenberg, Neergaard, Raklev and Blinkenberg) excavated a cemetery from the Early Roman Iron Age. The pottery grave goods (NM1 C1595 to C16024) have dated the skeletal material. Table S7.2.15 provides a sample list for this study.

| **CGG No** | **Museum id, Grave** |
|---|---|
| CGG_2_107492 | NM1 C 15922-5, Grave TT |
| CGG_2_107493 | NM1 C 15961-63, Grave J taken out due to low coverage |
| CGG_2_107497 | NM1 C 15981, Grave JJ |
| CGG_2_107490 | NM1 C 15985, Grave MM |
| CGG_2_107494 | NM C15987, Grave OO |
| CGG_2_107489 | NM C15991, Grave SS |
| CGG_2_107496 | NM C16001-002, Grave YY |
| CGG_2_107488 | NM C16004, Grave Bb |
| CGG_2_107487 | NM C16013, Grave Ii |

| | |
|---|---|
| CGG_2_107500 | NM C16014, Grave Kk taken out due to low coverage |
| CGG_2_107499 | NM C16015, Grave Ll |
| CGG_2_107498 | NM C16019, Grave Nn |
| CGG_2_107495 | NM C16021, Grave Pp |
| CGG_2_107486 | NM 15968a, Grave Z |

**Table S7.2.15:** List of samples studied from Denmark, Zealand, Københavns county, Tune hundred, Karlslunde parish, Mosede Fort.

#### Denmark, Zealand, Kaarebjerg Agre, Tørslev Hage, Gerlev Sogn

Coordinates: 55.8085, 12.050

Jens Winther Johannsen

In the spring of 2020, ROMU[426] investigated the remains of a grave on the easternmost part of the sandy peninsula Tørslev Hage on the west side of Roskilde Fjord. The grave was discovered during construction work and initially investigated by the police. It was therefore severely disturbed when ROMU became involved. Although some of the bones were scattered as a result of the disturbance, they were generally well preserved. This was due to the fact that the grave was filled with shells, possibly from a kitchen midden, which had prevented the calcium from leaching out of the bones. An examination of the bones showed that the grave must have contained two adult males, one about 25–30 years old and about 175 centimetres tall (X11) and the other 40–45 years old and about 182 centimetres tall (X12). Their stature indicated good nutrition and no signs of deficiency diseases were observed on the bones. Both had dental calculus, and the teeth of the oldest individual were in rather poor condition: several had been lost during life, there was significant tooth wear, signs of periodontal disease, an abscessed tooth, and several cavities. The latter suggests that the older man's diet for a large part consisted of carbohydrates. The two individuals appear to have been buried in two stone-lined coffins with different orientations but in the same plot. Stratigraphic observations suggest that the older individual (X11) was buried first and that some of the stones from his coffin were reused when the younger individual was buried. A bone dress pin was found on the upper left arm of the younger individual. A bifacial flint dagger of Lomborg type IB found during the police investigation shows that the grave belongs to the first half of the Late Neolithic (2350–1950 BC).

CGG_2_106770 (ROM3667, Skeleton x11).

CGG_2_106838 (ROM3667, Skeleton x12).

#### Denmark, Zealand, Lundbjerggård (020512-37)

Coordinates: 55.6272, 12.1338

Katrine Ipsen Kjær

Due to residential development of the minor village Vindinge, southeast of Roskilde, a burial site was located and excavated in 2017–2018. A total of 13 inhumation graves were examined. The samples analysed in this study are listed in Table S7.2.16. The burial site was situated at the highest area and was also placed exactly in the middle between the contemporaneous burial sites Stålmosegård and Vindinge Vest.

The burials were mostly oriented N-S, except for two, which were orientated NE-SW.

In all the graves human bones were preserved, which showed that men, women and children were present. All of the deceased were buried in separate graves.

All the graves contained grave goods dominated by personal equipment. Monstrous fibulas, as well as simple fibulas, beads from glass, amber and metal, hair needles, golden finger rings, combs, knives, ceramic vessels and food offerings, were present. A single glass beaker was present in two of the graves, both as imports from the Roman Empire.

According to the different kinds of fibulas and the glass beakers, the site is dated to the Late Roman Iron Age, primarily C1b-C2.

| **CGG No** | **Museum id** |
|---|---|
| CGG_2_105323 | ROM3469, A1/x9 |
| CGG_2_105327 | ROM3469, A598/x336 |
| CGG_2_105325 | ROM3469, A652/x127 |
| CGG_2_105322 | ROM3469, A653/x130 |
| CGG_2_105326 | ROM3469, A655/x496 |

**Table S7.2.16:** List of samples studied from Denmark, Zealand, Lundbjerggård.

#### Denmark, Zealand, Maderne, Store Fuglede parish (03.01.06-97)

Coordinates: 55.57 11.25

Anders Fischer

Late Bronze Age (900–673 BCE)

Bog area with sacrificial depositions spanning Neolithic to Iron Age periods. At the site in question a human skull was found during repair of a drainage pipe in 1981. Archaeological inspection revealed a flint axe of Late Neolithic type, pottery of Pre-Roman Iron Age character as well as fragmented bones of several animals and humans. Previous radiocarbon dates on human remains (K-4088) and a horse bone (K-4089) from the site span the period Iron Age to Viking Age. For the present study, a right human *pars petrosa* was sampled.

CGG_2_021421(AS 9/85, NM1 3841/8, NEO885) has been dated to 2624±40 $^{14}$C years BP (UBA-39140).

#### Denmark, Zealand, Mandemarke Bakker (040411-232)

Coordinates: 54.9647177, 12.5006051

An inhumation in a hocker position was excavated in 1955. The buried individual was a female aged c. 50 years. The grave is radiocarbon dated to the transition from Pre-Roman Iron Age to Roman Iron Age, 50 BCE–70 CE.

CGG_2_107523 (NM 465/55) has been dated to 1995±15 BP (UCIAMS-272884 ,>30kDa collagen yield (%) 1.9, d15N (‰) 10.1. d13C (‰) -20.3, C/N (atomic) 3.2).

#### Denmark, Zealand, Præstø county, Bjæverskov hundred, Vråby parish, Læbrogård (050113-6)

Coordinates: 55.3503662, 12.1533117

Johan Zakarias Gårdsvoll & Mads Dengsø Jessen

In 1929 (j.nr. NM1 4/29) Carl Peter Neergaard excavated two inhumation graves in a gravel pit, dating them via grave goods to the Late Roman Iron Age.

CGG_2_107403 (NM1 C 21853-63, Grave 1).

#### Denmark, Zealand, Præstø county, Bårse hundred, Beldringe parish, Hastrup (050202-37)

Coordinates: 55.269, 12.020

Johan Zakarias Gårdsvoll & Mads Dengsø Jessen

In 1945 (j.nr. NM1 583/46), Carl Johan Becker excavated 2 inhumation graves in a gravel pit, and another one in 1946. In particular the last excavated grave contained several grave goods, mainly pottery, and can be dated to the Late Roman Iron Age.

CGG_2_107419 (NM1 583/46, Grave II).

#### Denmark, Zealand, Præstø county, Bårse hundred, Bårse parish, Broskov (050203-50)

Coordinates: 55.1457358, 11.9982349

Johan Zakarias Gårdsvoll & Mads Dengsø Jessen

In 1927 (j.nr. NM1 440/27) Hans Christian Broholm excavated 5 inhumation graves (including Grave A and D) in a gravel pit at Broskov. As early as 1876 there had been reports of finds of many skeletal remains of humans and even horses. The sampled graves contained several types of grave goods, such as (Grave A) bronze sieve and ladle, a possible bronze vessel, and (Grave D) a silver fibula (Mackeprang type X), pottery, a bronze bucket and a bone comb. The graves can thereby be placed in the Late Roman Iron Age.

CGG_2_107394 (NM1 C 18915-17, Grave A).

CGG_2_107395 (NM1 C 18934-38, Grave D).

#### Denmark, Zealand, Præstø county, Bårse hundred, Skibinge parish, Endegaarde (050210-58)

Coordinates: 55.1080068, 12.0638179

Johan Zakarias Gårdsvoll & Mads Dengsø Jessen

In 1946 (j.nr. NM1 340/46 and 57/52), Hans Norling-Christensen excavated two inhumation graves. The graves contained 3 ornamented vessels and a three-layered bone comb placing them in the Late Roman Iron Age.

CGG_2_107416 (NM1 340/46).

#### Denmark, Zealand, Præstø county, Bårse hundred, Udby parish, Egebjerg (050212-18)

Coordinates: 55.0654, 11.9584

Johan Zakarias Gårdsvoll & Mads Dengsø Jessen

In 1895 (j.nr. NM1 254/95) Georg Frederik Ludvig Sarauw excavated 6 inhumation and 3 cremation graves, perhaps in the vicinity of an older mound. The sampled graves contained several types of grave goods such as (Grave 1) a comb, a pearl, a fibula and pottery, and (Grave 5) several pearls of glass and amber, a bronze ladle and sieve, several pottery vessels and 5 fibulas (one of these a rosetta fibula) and a gold ring. These finds place both graves in the Late Roman Iron Age.

CGG_2_107397 (NM1 C 7888-92, Grave 1).

CGG_2_107396 (NM1 C 7901-22, Grave 5).

#### Denmark, Zealand, Præstø county, Bårse hundred, Ørslev parish, Kirkebjerggaard (050215-24)

Coordinates: 55.346, 11.964

Johan Zakarias Gårdsvoll & Mads Dengsø Jessen

In 1945 (j.nr. NM1 777/45) during digging for sand in a field at Kirkebjerggaard an inhumation grave under a flat field with preserved skeletal remains was disturbed. The find was reported to the National Museum of Denmark and Peter Vilhelm Glob was sent to lead an inspection and an excavation of the site. Next to the cranium lay the remains of a wooden bucket with a handle and fittings made of bronze and a small ceramic cup. Next to the feet lay a large and a small ceramic vessel, the large one contained a little piglet. Besides the remains in the ceramic vessel the grave also contained remains of a sheep, a cow, a pig and a dog or a fox. Earlier reports of finds of skeletal remains in the same field could indicate that the grave is just one out of many other graves from a flat field burial ground. The typology of the ceramic vessels and the positioning of the body in the grave suggest that the date of burial occurred in the Late Roman Iron Age.

CGG_2_107423 (NM1 777/45).

#### Denmark, Zealand, Præstø county, Bårse hundred, Øster Egesborg parish, Lekkende (050216-27A)

Coordinates: 55.0657215, 12.0131568

Johan Zakarias Gårdsvoll & Mads Dengsø Jessen

In 1934 (J.nr. 365/34) Mogens Bellmann Mackeprang excavated two inhumation graves located in a gravel pit. They both contained dateable grave goods, such as silver and bronze pearls, and pottery, placing them in the Late Roman Iron Age.

CGG_2_107402 (NM1 C 23061-63, Grave 1).

#### Denmark, Zealand, Præstø county, Fakse hundred, Hylleholt parish, Borritshoved (050303-1A)

Coordinates: 55.2924, 12.1001

Johan Zakarias Gårdsvoll & Mads Dengsø Jessen

In 1937 (j.nr. NM1 354/37), Hans Christian Broholm excavated 8 inhumation graves, where the two sampled internments were equipped with the most grave goods, such as (Grave 1) a small knife, and (Grave 4) a similar knife, pottery, and iron spike, and an ornamented spindle whorl. The graves can be dated to the Early Iron Age.

CGG_2_107470 (NM1 354/37, Grave 1).

CGG_2_107469 (NM1 354/37, Grave 4).

##### Denmark, Zealand, Præstø county, Fakse hundred, Hylleholt parish, Vemmetofte Skovridergård (050303-15)

Coordinates: 55.2541586, 12.2237027

Johan Zakarias Gårdsvoll & Mads Dengsø Jessen

In 1926 (j.nr. NM1 468/26) K. Klem inspected and excavated 6 inhumation graves in a gravel pit. In 1970 (j.nr. NM1 377/70), T. Ramskou excavated a quite rich female grave with a well-preserved skeleton, pottery, glass beads and a silver fibula, and an arm ring from the Early Roman Iron Age. In 1987 (j.nr. NM1 6630/87), a metal detectorist found a bronze saucepan of Roman origin in the same area.

CGG_2_107414 (NM1 468/26, Grave 1).

#### Denmark, Zealand, Præstø county, Hammer hundred, Hammer parish, Hammer Strandhuse (050401-13)

Coordinates: 55.1462609, 11.8701669

Johan Zakarias Gårdsvoll & Mads Dengsø Jessen

In 1953, (j.nr. NM1 366/53) a private Degenkolw excavated 3 inhumation graves with two adults and a child. The child was equipped with a three-layered comb, and the pottery grave goods date the graves to the Late Roman Iron Age.

CGG_2_107418 (NM1 366/53, Skeleton 2a).

#### Denmark, Zealand, Præstø county, Hammer hundred, Kastrup parish, Neder Vindinge (050403-12)

Coordinates: 55.0312651, 11.8865266

Johan Zakarias Gårdsvoll & Mads Dengsø Jessen

In 1925 (j.nr. NM 159/24), Johannes Balthasar Brøndsted excavated an inhumation grave with different types of pottery as grave goods. These date the grave to the Roman Iron Age.

CGG_2_107406 (NM1 C 18998-99, Grave A2).

#### Denmark, Zealand, Præstø county, Hammer hundred, Køng parish, Øbjerggaard (050404-24)

Coordinates: 55.1162152, 11.829094

Johan Zakarias Gårdsvoll & Mads Dengsø Jessen

In 1955 (j.nr. NM1 539/56), C. L. Vebæk inspected and excavated 8 graves discovered by a local farmer in a sandpit. Relatively well-preserved inhumations but most of them lacking grave goods. A few indeterminate pieces of iron were found.

CGG_2_107433 (NM1 539/56, Skeleton I).
CGG_2_107434 (NM1 539/56, Skeleton 2).
CGG_2_107436 (NM1 539/56, C 27384, Skeleton 5).
CGG_2_107435 (NM1 539/56, C 27383, Skeleton 7).

CGG_2_107436 has been dated to the Pre-Roman Iron Age- Roman Iron Age transition (60 BCE–60 CE) UCIAMS-275219 2025±15 BP (>30kDa collagen yield (%) 2.6, d15N (‰) 10.1. d13C (‰) -19.6. C/N (atomic) 3.2)).

#### Denmark, Zealand, Præstø county, Hammer hundred, Toksværd parish, Toksværd (050410-22)

Coordinates: 55.2526298, 11.8796892

Johan Zakarias Gårdsvoll & Mads Dengsø Jessen

In 1921 (j.nr. NM1 224/21) while digging for marvel in a hill an inhumation grave under a flat field was disturbed. The site was visited and excavated by K. Friis Johansen, The National Museum. It contained a skeleton and a well-preserved foot-vessel from the Early Roman Iron Age.

CGG_2_107504 (NM1 C 18895).

#### Denmark, Zealand, Præstø county, Hammer hundred, Vester Egesborg parish, Vester Egesborg (050412-5)

Coordinates: 55.155344, 11.8190732

Johan Zakarias Gårdsvoll & Mads Dengsø Jessen

In 1855 a private excavation revealed 7 inhumations graves in a gravel pit, and they were handed over to the National Museum in 1872. Based on the ceramic grave goods found with one of the individuals the cemetery can be dated to the Early Roman Iron Age.

CGG_2_107506 (Skeleton 2).

CGG_2_107507 (Skeleton 3).

#### Denmark, Zealand, Præstø county, Stevns hundred, Hellested parish, Barup Sandgrav (050603-20)

Coordinates: 55.332, 12.231

Johan Zakarias Gårdsvoll & Mads Dengsø Jessen

In 1955–1959 and 1961 (j.nr. NM1 658/61) multiple inspections and excavations were conducted by Olfert Voss, Christen Leif Pagh Vebæk, Mogens Ørsnes-Christensen, Ole Klindt-Jensen, Henrik Thrane and Bjørn G. Stürup in a sandy hill at Barup occasioned by several instances of disturbances of graves in the sand pit and subsequent reports to the National Museum of Denmark. In total the traces of 10 graves have been found, many of them containing grave goods. From the grave goods and/or from the positioning of the bodies in the graves one grave could be dated to the Viking Age and the rest (including Grave 7) to the Late Roman Iron Age.

CGG_2_107385 (NM1 658/61, Grave 7).

#### Denmark, Zealand, Præstø county, Tybjerg hundred, Næstved parish, Næstved Mark (050707-1)

Coordinates: 55.22959, 11.77806

Johan Zakarias Gårdsvoll & Mads Dengsø Jessen

In 1901 (j.nr. NM1 676/01) T. Thomsen excavated a series of inhumation graves in a hill. The cemetery contained a grave with several grave goods such as pottery, a comb and silver pearls dating the cemetery to the Late Roman Iron Age.

CGG_2_107407 (NM1 C 11362, Grave 3).

CGG_2_107408 (NM1 C 11363, Grave 5).

#### Denmark, Zealand, Præstø county, Tybjerg hundred, Næstved parish, Skyttemarksvej 59-61 (050707-0A)

Coordinates: 55.237, 11.755

Johan Zakarias Gårdsvoll & Mads Dengsø Jessen

In 1932 (j.nr. NM1 779/32), Hans Christian Broholm registered a privately excavated richly furnished female inhumation grave that was partly covered with stones. Among these there was a swastika-shaped fibula, other silver jewellery and several pearls of glass and bronze, not to mention pottery. These finds place the grave in the early parts of the Late Roman Iron Age[427].

CGG_2_107409 (NM1 C 20509-21).

#### Denmark, Zealand, Præstø county, Tybjerg hundred, Rislev parish, Rislev (050708-7?)

Coordinates: 55.2815323, 11.7331049

Johan Zakarias Gårdsvoll & Mads Dengsø Jessen

Table S7.2.17 lists the samples provided from this site.

| **CGG No** | **Museum id** |
|---|---|
| CGG_2_107510 | NM A3979, Skeleton 1 |
| CGG_2_107509 | NM A3979, Skeleton 2 |

**Table S7.2.17:** List of samples studied from Denmark, Zealand, Præstø county, Tybjerg hundred, Rislev parish, Rislev.

#### Denmark, Zealand, Præstø county, Tybjerg hundred, Skelby parish, Øsbergårds Mark (050710-50)

Coordinates: 55.3131, 12.0087

Johan Zakarias Gårdsvoll & Mads Dengsø Jessen

In 1934 (j.nr. NM1 260/34) Mogens Bellmann Mackeprang, The National Museum, excavated 3 disturbed graves with 1 skeleton in each. All three were inhumations and with relatively sparse amounts of grave goods. The ceramics can be dated to the Early Roman Iron Age.

CGG_2_107439 (NM1 C 23058, Grave 1).

CGG_2_107438 (NM1 C23059, Grave 2).

CGG_2_107437 (NM1 C23060, Grave 3).

#### Denmark, Zealand, Præstø, Bårse, Udby parish, Skovgårde (050212-27)

Coordinates: 55,0792, 11,9579

A cemetery whose first archaeological traces were found in 1943[428]. Eighteen north-south originated inhumation graves have been excavated between 1943 and 1988 The cemetery's layout was divided into four groups and one horse . All graves date to Late Roman Iron Age C1b and C2 [428]. Fourteen individuals have earlier been sampled for genetic analysis, eleven of these yielded information about MtDNA haplogroups [410]. These were the graves 4, 5, 7–9, 202, 204, 205, 207–210, 400 and 1943. Five belonged to haplogroup H (4, 7, 208, 210 and 1943), two to haplogroup J (204 and 207), two to haplogroup K (5 and 205), one to haplogroup V (209), one to haplogroup U3a (8) and three gave no results (9, 202 and 400) [410]. In this study grave 202 has yielded genetic information we also sampled and gained information from grave 1. Melchior et al. (2010)[429] argues that in difference from Bøgebjerggård the haplogroups seen here are the ones we see in modern Scandinaivans. Samples included in this study are listed in Table S7.2.18.

| CGG No | Grave | Sample description | Grave description |
|---|---|---|---|
| CGG_2_106797 | Grave 1, x240 | child | A grave containing a number of object from silver fibulas to textile fragments |
| CGG_2_106798 | Grave 205, x309 | Female? c. 25 | The grave contain among other things glass beads, a comb and ceramic vessels |
| CGG_2_106801 | Grave 9, x31 | Female aged 55–58 | A grave containing a number of objects including fibulas, beads and a comb |
| CGG_2_106802 | Grave 202, x294 | Female 18–20 | The grave contain among other things a silver fibula, pendants and spiral silver beads |

**Table S7.2.18:** List of the samples studied from Denmark, Zealand, Præstø, Bårse, Udby parish, Skovgårde and summary of the corresponding anthropological and archaeological studies.

#### Denmark, Zealand, Præstø, Bårse, Vordingborg parish, Eksercérpladsen, (050213–148)

Coordinates: 55,0151, 11,9204

Sophie Bergerbrant

A small inhumation cemetery that was excavated in 1989 (Fund og Fortidsminder, 05213-148; accessed 2023-11-01). The cemetery was structured in two groups, one northern containing three graves and eastern with eight graves. The graves have few grave goods but based on the ones found the cemetery dates to the Roman Iron Age. In the vicinity of the graves were traces of settlement activities in the form of different types of pits.

CGG_2_106796 (SMV7122, grave 58).

#### Denmark, Zealand, Præstø, Bårse, Vordingborg parish, Græsbjerg (050213-129)

Coordinates: 55,0124, 11,9558

Sophie Bergerbrant

A cemetery containing 21 inhumation graves[380]. The cemetery was excavated 1978–79. The sample in this study comes from grav A (CGG_2_106800, SMV20/79). A grave that contained two individuals, the sampled individual is skeleton 1. The grave was orientated N–S and dates to the Roman Iron Age. The grave contained a clay button and fragments of either an iron pin or iron brooch. Skeleton 1 are the remains of an adult female and skeleton 2 of an adult/mature probable female[380].

CGG_2_106800 (SMV20/79, grave A).

#### Denmark, Zealand, Præstø, Bårse, Øster Egesborg parish, Bøgebjerggård, Røstofte (050216-132)

Coordinates: 55.0340, 11.9935

Six graves containing seven individuals were excavated in 1992, Earlier, in 1983, had three human skeletons been found while ploughing, one of these graves were excavated. The area contained preserved graves and spread human remains. In 2000 one more grave was excavated at the site and during excavation 2020 human remains from at least two more graves were found. The graves date to the Early Roman Iron Age, though grave 7 (CGG_2_106795) has been radiocarbon dated to the transition Pre-Roman Iron Age – Roman Iron Age.

Eight individuals have previously been sampled for aDNA[410,429], Seven of these have published mtDNA (Melchior et al 2010). The graves earlier analysed are the graves A, B, E, 4, 4A, ,6, 7 , 8 and marts 2000. Two of these belonged to haplogroup I (B and E), two to haplogroup H (6 and 7), one to haplogroup R0a (4), one to haplogroup U2e (6) and to haplogroup U5b (marts 2000). Grave 4A did not yield any genetic result[410]. According to Melchior et al (2010)[429] the haplogroups R0a, U2 and I are rare in present day Scandinavia. They argue that the presence of R0a suggests that the male buried in grave 4 was a slave or a soldier with connection to regions further south. Samples included in this study are listed in Table S7.2.19.

| Sample id | Museum id | Gender | Age | Description |
|---|---|---|---|---|
| CGG_2_106793 | SMV7273, Grave March 2000 | Female | 35–45 | Deceased placed in hocker. The grave contained six glass- and amber beads and one ceramic vessel |
| CGG_2_106794 | SMV7273, Grave B | Female | 25–30 | |
| CGG_2_106795 | Grave 7 | Male | 25–30 | Pre-Roman Iron Age- Roman Iron Age 50 BCE to 60 CE, UCIAMS-275216 2020 ±15 BP (>30kDa collagen yield (%) 12, d15N (‰)13.2. d13C (‰) -20.3. C/N (atomic) 3.1) |

**Table S7.2.19:** List of the samples studied from Denmark, Zealand, Præstø, Bårse, Øster Egesborg parish, Bøgebjerggård and summary of the corresponding anthropological, archaeological and isotopic studies.

#### Denmark, Zealand, Præstø, Hammer, Hammer, Hundstrup Mose (050401-12)

Coordinates: 55.1526, 11.9176

During peat digging in 1947 were human and animal bones found. It was the remains of two adults, two children and three infants[430]. The two children were covered by branches and placed next to each other and with them was an iron knife found. About five metres from the two children were the adults and the infants found. The remains were well preserved but not completely preserved. The adults were determined to be male aged 25–35 and 30–50. The two children were aged 9–11 and 11–13. The infants were all newborn. None of the individuals showed any traces of violence[332].

Strontium isotope analysis of the two adults and the older child indicates that only the younger male (VK296) was local[332]. See Table S7.2.20 for additional information.

The radiocarbon dates indicate that the individuals had not been placed in the bog at the same time. The older child is radiocarbon dated to 500–610 CE and the younger male and child dates to the period 660–780 and the older male 670–830 CE. Indicating that the people ended up in the wetland at atleast two occasions.

CGG_2_107518 (individual 1) has been dated to 1345±15BP (650–760 CE) (UCIAMS-272883, >30kDa collagen yield (%) 8.7, d15N (‰) 11.3. d13C (‰) -19.3. C/N (atomic) 3.2)). The two adults have previously been analysed for genetic information[405].

| **Sample id** | **Museum id** | **Sex/Age** | **Period** | **Haplotyåe** |
|---|---|---|---|---|
| CGG_2_107518 | NM 673/47, Skeleton 1 | UCIAMS-272883 | 1345±15BP (650–760 CE) | Same as VK296 |
| VK296 | Hundrup Mose sk 1 | Younger male, 25–35 | 660–780 CE | mtDNA haplogroup HV6, Y_haplogroup I1a1b1a1 |
| VK297 | Hundrup Mose sk 2 | Older male, 45+ | 670–830 CE | mtDNA haplogroup J1c2h, Y_haplogroup I1a1b1 |

**Table S7.2.20:** List of samples studied from Denmark, Zealand, Præstø, Hammer, Hammer, Hundstrup Mose, their dating and genetic analysis results.

#### Denmark, Zealand, Præstø, Hammer, Sværdborg Parish, Over Vindinge

Coordinates: 55.070, 11.890

Helle Vandkilde & Anna Tornberg

The site of Over-Vindinge boasts two large gallery graves built within 300 m of one another and excavated in the early 20th century. The cists are 3 m long and about 1 m wide while narrowest at their rear end and at the entrance. The shape of the burial chamber is rectangular-roundish and oriented W-E with the long sides in N and S respectively with a cover of large stone slabs. The last internment was placed supine on the floor while previous interments were moved to the entrance part where kept in a certain order. Both cists counted several internments, which took place over a long period of time, c. 2200–1700 BCE, corresponding to the LN II and NBA IA periods according to the radiocarbon dates. The burial goods are few and inconspicuous albeit confirming the date to the final Neolithic. Along with several other gallery graves the Over-Vindinge cists were built 2200–2100 BC at the time when metallurgy became fully established. Little if any activity took place in the burial space after c. 1800–1700 BC. Gallery Grave I is the better known of the two cists due to the pelvis bone with the spearhead tip still sitting in situ[431]). Gabs between the stones were sometimes filled in by pressing bones into them including the pelvis bone with the spearhead tip, according to Kjær[431]. Identifying and classifying the individuals at Over-Vindinge is quite a puzzle and forms part of an ongoing-trauma-pathological study[432]. Preliminarily: Gallery Grave I comprised 6 individuals while Gallery Grave II held 4 individuals (MNI). They were all males, some of them quite old with pathological issues. Six were sampled for DNA, three had sufficient DNA for further sequencing, all from Gallery Grave I (Table S7.2.21).

| CGG No | Inventory NM - Panum | BP | cist | DNA | Sex |
|---|---|---|---|---|---|
| CGG_2_105335 | NM36/11, A 26881-86 PMD 124 | 3505±24 | 1 | low | XY |
| CGG_2_105334 | NM36/11, A 26881-86 PMD 125 | 3480±23 | 1 | sufficient | XY |
| CGG_2_105337 | NM36/11, A 27518 PMD 123 Individ G | 3659±24 | 2 | low | XY |
| CGG_2_105332 | NM36/11, A2688-86 n. 7 PMD 127B | 3465±35 | 1 | sufficient | XY |
| CGG_2_105333 | NM36/11, A26881-86 n. 9 PMD 126 "V" | 3549±26 | 1 | sufficient | XY |
| CGG_2_105336 | NM36/11, A 26881-86 PMD 122 | 3566±25 | 2 | low | XY |

**Table S7.2.21:** List of samples studied from Denmark, Zealand, Præstø, Hammer, Sværdborg Parish, Over Vindinge, dating and DNA content and sex identification.

#### Denmark, Zealand, Præstø, Tyberg, Tyberg, Tybjerg Mose (050711-18)

Coordinates: 55.337, 11.764

During peat digging in 1945 was a pot from the iron age, a wooden club, human and animal bones found. The animal bones were placed in two piles within an area that had been fenced off by wooden stakes. C.J. Becker investigated the site after it had been discovered. It was the human remains from two individuals. They were found c. 3m from the animal bone piles. The wooden club and the potsherds were found next to the humans.

CGG_2_107512 (NM 377/45, Skeleton 1) was according to the information disturbed but was found upright in the mire. It is the remains of an adult male and is dated to the Late Bronze Age/Pre-Roman Iron Age 750–170 BCE (K-3607, 2320±70).

Skeleton II was found scattered around individual I and is incomplete, possibly a female[380].

#### Denmark, Zealand, Rosenåen, Brøndbyvester parish (02.02.03-48)

Coordinates: 55.63, 12.41

Pre-Roman Roman Iron Age (160–1 BCE)

A small Early Roman Iron Age cemetery containing five or possibly six inhumation graves and one cremation grave. All the graves are orientated east-west. Samples were taken from grave 31 (TAK1594_X31) and grave 42 (TAK1594_X42) but only grave 31 yielded enough data to be included in this study. Grave 31 was placed in a hocker position with the head in the west facing south, the body had fallen forward. The deceased had a S-shaped knife and an awl placed in the area of the left upper arm. The grave also contained organic remains, possibly leather and iron residue from a possible needle. The human remains were determined to belong to a woman that was aged 45+.

CGG_2_011889 (MAp1162, Rosenåen X31, prev.), UCIAMS-233748, 2075±15 BP, >30kDa collagen yield (%) 10.6, d15N (‰) 11.6. d13C (‰) -19.9, C/N (atomic) 3.1 (160-1 BCE).

#### Denmark, Zealand, Roskilde, Fløng Parish, Trekroner-Grydehøj (020401-17)

Coordinates: 55.6572, 12.1479

Ole Thirup Kastholm

Grydehøj is the name of a small hill in the now developed area of Trekroner some 4 kilometres east of the town centre of Roskilde. Beginning in the late 1990s, more than 70 hectares of farming land were archaeologically surveyed by trial excavations before residential development. During this, a burial place on the isolated hill of Grydehøj was located in 2005 and excavated in 2007[433–435].

The excavation revealed graves and burial monuments from the Late Stone Age, the Early Bronze Age, the Late Bronze Age/Early Iron Age, and the Viking Age. During the Late Neolithic Period (c. 2000–1700 BCE) three stone cists covered with one or more mounds were situated on the hilltop. In the Early Bronze Age (c. 1700–1100 BCE) a mound was built neighbouring the stone cists to the west. Later the mound was expanded to facilitate a new Early Bronze Age grave, and a ditch was dug around the eastward foot of the mound, probably to make the mound appear larger compared to the Neolithic mounds. From the very Late Bronze Age or the beginning of the Pre-Roman Iron Age (c. 600–400 BCE) are three urn graves, buried at the western foot of the hill. During the late eight or early ninth century CE, a Viking Age burial ground was established with the old cluster of mounds as a point of departure. The 27 inhumations were primarily found on the western slope of the hill. Altogether 23 skeletons were registered. Two single graves contained additional burned human skeletal material, while five graves held two or more persons. The graves with two or more individuals could be divided in two groups: double burials, where the bodies have been interred at the same time in the same pit, and primary/secondary burials characterised by a primary grave with one individual and a secondary grave dug within the boundary of the first one.

The sample is from individual 3 in the grave A505, a Viking Age inhumation burial belonging to the latter group of graves, with primary/secondary burials. A505 held the remains of three individuals: Two women, a half man, a stallion, a dog cut in two halves, a sheep foetus and a small menhir constituting a complicated sequence of events sealed with a stone covering. The lower female (individual 3) was seemingly the burial's primary interred person. She has been interpreted as a sorceress due to a number of extraordinary elements in the layout and inventory of the burial.

CGG_2_106841 (ROM 2285, Grave A505: individual 3: right petrous bone) has been dated to , 1300±20 BP (UCIAMS-282665, >30kDa collagen yield (%) 2.2, d15N (‰) 10.2. d13C (‰) -19.4, C/N (atomic) 3.2).

#### Denmark, Zealand, Roskilde, Vindinge Parish, Vindinge Vest I-II (020512-36)

Coordinates: 55.62415, 12.13301

Nadja M.K. Mortensen

Due to residential development, a large settlement from the Roman Iron Age with an adjacent burial site from the same period was uncovered and excavated in 2018. A total of 79 inhumation graves were examined[436]. The majority of the graves (70) were uncovered at the burial site while nine graves were located among the settlement features. Six of these graves were placed together in a cluster just south of the burial site while the remaining three graves were found alone. The samples investigated in this study are listed in Table S7.2.22.

Of the 79 examined graves 66 contained grave goods. Both children's graves as well as graves with adults contained grave goods. The amount and the worth of the objects in the graves varied. The grave goods consisted of simple as well as monstrous fibulas, beads from glass, amber and metal, finger rings made from gold, silver and bronze, a belt bracket, needles, weights and weight hooks, combs, knives, ceramics, wooden dishes and food offerings.

Both settlement and burials indicate a long settlement continuity throughout the Roman Iron Age and the findings reflect a society of increasing wealth; the amount of objects in the graves increased and the objects became more exclusive.

A total of 41,800 $m^2$ were excavated of which the burial site covered 5,400 $m^2$. The site is considered fully uncovered to the West but continues under the existing village to the East.

| CGG No | Museum id |
|---|---|
| CGG_2_105317 | ROM3385, A4009/x5283 |
| CGG_2_105313 | ROM3385, A4010/x5490 |
| CGG_2_105319 | ROM3385, A4038/x4694 |
| CGG_2_105318 | ROM3385, A4039/x6426 |
| CGG_2_105316 | ROM3385, A4047/x5582 |

**Table S7.2.22:** List of samples studied from Denmark, Zealand, Roskilde, Vindinge Parish, Vindinge Vest I-II.

#### Denmark, Zealand, Sejerø parish, Havnegård (030607-162)

Coordinates: 55.884, 11.153

Johan Zakarias Gårdsvoll & Mads Dengsø Jessen

In 1969 (j.nr. NM1 545/69) David Liversage excavated a single, stone-covered inhumation grave with two skeletons in a farm field. The lowest and oldest (Grave II B) is sampled here

(CGG_2_107517). The grave contained 3 knives and a few fragmented pieces of pottery. Based on the knives the grave must presumably be dated to the Late Germanic Iron Age.

CGG_2_107517 (NM1 545/69, Grave II B).

#### Denmark, Zealand, Smørum, Bredebjerggård 8

Coordinates: 55.6404, 12.2571

At the site both remains of settlements and one burial have been excavated. The burial is an inhumation dated based on the ceramic in the burial to the Late Roman Iron Age. The deceased was placed in supine position[437].

CGG_2_106822 (TAK1180, Grave 477) aDNA XY.

#### Denmark, Zealand, Smørum, Sejlsbjerg

Coordinates: 55.64423, 12.2073

Among other things did Kroppendal Museum excavate a small Early Iron Age cemetery in 2017–2019[438]. The cemetery contained four inhumations burials. That dates to the Roman Iron Age close by where a grave from the Pre-Roman Iron Age was found. In the excavated area was another Roman Iron Age burial found. Sampled here are grave 549, an inhumation where the deceased was found with stomach facing down, though this were interpreted as a post-burial taphonomy. The grave is dated to Early Roman Iron Age. The skeleton was well preserved and was determined to be the remains of a female 146 cm tall aged 20–23.

CGG_2_106820 (TAK1621, Grave 549) aDNA XX.

#### Denmark, Zealand, Sokkelund, Assistens Kirkegård

Coordinates: 55.6912, 12.5501

Marie Louise Jørkov

The Assistens cemetery, section G dates from the late 18th to mid 20th century. It was excavated by Copenhagen Museum in 2009–2011, where more than 850 graves were recovered. Individual SB15088 probably dates to the late half of the 19th Century and/Early 20th Century AD. She was ca. 5 months pregnant at the time of death. Individual SB15138 was a 33 year-old female who died in the last quarter of the 19th century. The remains were well preserved.

CGG_2_105338 (KBM3830, SB15088).

CGG_2_105340 (KBM3830, SB15138).

#### Denmark, Zealand, Sorø county, Alsted hundred, Alsted parish, Alsted (040101-40)

Coordinates: 55.3792, 11.6773

Johan Zakarias Gårdsvoll & Mads Dengsø Jessen

In 1978 (NM1 2541/78) in a steep moraine at Alsted an inhumation grave with preserved skeletal remains was disturbed. The find was reported and Svend Nielsen was sent to inspect and excavate the site. In the grave that had been heavily disturbed by ploughing remains of a ceramic vessel was found. The body was positioned on its right side and in hocker position.

According to the landowner he had often come across bones in this area of his land. The excavated grave therefore presumably represents one out of many from a flat field burial ground. The typology of the ceramic vessel and the positioning of the body in the grave suggest that the date of burial occurred in the Late Roman Iron Age.

CGG_2_107384 (NM1 2541/78).

#### Denmark, Zealand, Sorø county, Ringsted hundred, Haraldsted parish, Haraldsted (040206-52)

Coordinates: 55.467, 11.811

Johan Zakarias Gårdsvoll & Mads Dengsø Jessen

In 1953 (j.nr. NM1 442/53) Georg Kunwald excavated 3 inhumation graves in a gravel ridge. Presumably a man, female and a child (Grave 2). The presumed male was equipped with pottery grave goods that can be dated to the Roman Iron Age.

CGG_2_107508 (NM1 442/53, Grave 2).

#### Denmark, Zealand, Sorø county, Slagelse hundred, Boeslunde parish, Espe (040301-75)

Coordinates: 55.4006, 11.3225

Johan Zakarias Gårdsvoll & Mads Dengsø Jessen

In 1947 (j.nr. NM1 611/47), Peter Vilhelm Glob excavated 8 inhumation graves, where only one contained grave goods, a rectangular, ornamented bronze fibula (Ørsnes K3), stirrups and a few iron fragments placing the grave in the later parts of the Late Germanic Iron Age. The two sampled graves (Grave 4 and 5) are dated to the same period on the basis of the rich grave.

CGG_2_107516 (NM1 611/47, Grave 4, Espe II).

CGG_2_107515 (NM1 611/47, Grave 5).

#### Denmark, Zealand, Sorø county, Slagelse hundred, Hejninge parish, Trelleborg

Coordinates: 55.389, 11.308

Johan Zakarias Gårdsvoll & Mads Dengsø Jessen

The circular Viking Age fortress of Trelleborg on the island of Zealand in Denmark is well known as a military fortress built by Harold Bluetooth or by his son Svend in AD 980/981. The site is crucial for the understanding of the beginning of the Danish state formation and has therefore been intensively investigated. Trelleborg is situated at a promontory bordered by the rivers Tude Å and Vårby Å in southeastern Zealand with a view towards the Great Belt. It was excavated in the years 1934–42 by the National Museum[380,439–441].

Believed to belong to the decades prior to the construction of the fortress the excavator found several structures, and among these were two wells number 47 and 121. Well 47 was located within house 4 in the southeastern building complex of the fortress. It was oval and measured 1.8 × 2.35m at the surface and narrowed down to 0.90 × 1.20 m at the bottom. It was 2.13 m deep. Among the artefacts in the fill was a tenth century tortoise brooch, bronze bowl, an arrowhead, and pottery, and different kinds of household tools. In this well, skeletal remains of two children were found (CGG_2_107581 and CGG_2_107582). Both skeletons were fairly complete and well preserved, though fragmented.

Well 121 was located in the northeastern building complex. It was circular with a diameter of 3–3.75 m and reached a depth of c.3m below the surface of the subsoil. In the upper fill layers, fragments of pottery, semi-finished antler for combs, whetstone, a bone skate and iron fragments were found. The lower layers provided three belt buckles, a knife fragment and various iron objects. However, none of the finds seem to be associated with the primary function as a sacrificial well as they derive from the secondary fill material. Both wells and the adjoining trench are cut by the building wall and thus from the period before the fortress. Human skeletal remains of two children were found in the well (CGG_2_106832 and CGG_2_106833). Both skeletons were well preserved though incomplete and fragmented. Skull fragments from both children were present and some few postcranial parts, and the excavator has described both children as if they were complete when thrown in the well. The strontium isotope values of all four children indicate local origin.

Both wells had several bones from different kinds of animals such as horse, pig, cow, dogs and others. Some of these are interpreted as thrown in the wells as complete animals, while others most have been dismembered.

The burial ground at Trelleborg was excavated between 1938 and 1940 and altogether 133 graves containing 157 individuals were unearthed. The majority of graves held a single individual, while three of the graves must be interpreted as mass burials and there are also two double burials. All burials were inhumations in rather shallow graves, originally marked above ground, probably by a small mound. The graves are oriented east-west, with the dead generally facing east. The burial ground is situated outside the eastern entrance of the fortress, with burials scattered on both sides of the main east-west axis that leads to the complex and the cemetery is assumed to be the burial place for the deceased occupants of the fortress complex. Both graves 38 (CGG_2_106824) and 54 (CGG_2_106823) are believed to be males between 35–55 years old, and none of them had any grave goods. They belong to the south-eastern part of the cemetery and grave 38 to the perimeter. The strontium isotope value of grave 54 indicates that he is a non-local. Grave 1 (CGG_2_071583), AS21/45, Q1440 is believed to be a female between 35–55 years old. The grave goods contained; A single-edged knife (Q 1442), length 8.9 cm. The knife is broken off at both ends; Four-sided tapered iron fragment (Q 1443). Length 4.1 cm; Red, barrel-shaped glass bead (Q 1441). Height 0.8 cm, diameter 1.1 cm; Five Iron Nails (Q 1444–1447). The female grave belongs in the most south-western part of the cemetery at Trelleborg, orientated ESE-WNW and with her head in WNW. Table S7.2.23 provides a list of the samples under investigation in this study.

| **CGG No** | **Grave id** |
|---|---|
| CGG_2_107581 | Well 47a |
| CGG_2_107582 | Well 47b |
| CGG_2_071583 | AS21/45, Q1440, Grave 1 |
| CGG_2_106824 | Grave 38 |
| CGG_2_106823 | Grave 54 |
| CGG_2_106832 | Pit/well 121 |
| CGG_2_106833 | Pit/Well 121 |

**Table S7.2.23:** List of samples studied from Denmark, Zealand, Sorø county, Slagelse hundred, Hejninge parish, Trelleborg and their context description.

#### Denmark, Zealand, Sorø county, Slagelse hundred, Hemmeshøj parish, Hemmeshøj (040305-3A)

Coordinates: 55.420, 11.301

Johan Zakarias Gårdsvoll & Mads Dengsø Jessen

In 1920 (j.nr. NM1 393/20), K. Friis Johansen excavated 4 inhumation graves in a sand pit at Hemmeshøj. As early as 1892 there had been reports of finds of skeletal remains in the sandy hill. Only one of the graves (Grave 1) from Johansen's excavation contained grave goods (2 pottery vessels, a spindle whorl and an awl) which dates the entire cemetery to the Late Roman Iron Age.

CGG_2_107420 (NM1 C 17994, Grave 4).

CGG_2_107400 (NM1 C 17995, Grave 5).

#### Denmark, Zealand, Sorø county, Slagelse hundred, Slots Bjergby parish, Blæsenborgvej (040315-48)

Coordinates: 55.3982, 11.3105

Johan Zakarias Gårdsvoll & Mads Dengsø Jessen

In 2001 (j.nr. 8189/00-14), P. O. Nielsen excavated a large, semi-disturbed pit with human remains of presumably two individuals and unornamented pottery. In the disturbed pit also neolithic pottery as well as a polished flint axe was found. The former pottery could suggest the skeletal material belongs to the Early Germanic Iron Age.

CGG_2_107383 (NM1 8189/00-14).

#### Denmark, Zealand, Sorø, Vemmelev Forlev "Forlev I"

Coordinates: 55.3769, 11.2573

A cemetery counting 15 inhumation and a large number of cremation graves. Excavated by G. Sarauw 1895[380].

Grave 7 A Viking Age inhumation grave containing an iron mount and lock probably for a wooden chest.

CGG_2_106828 (Grave 7), aDNA XY.

#### Denmark, Zealand, Strandlunden 11, Gerlev (MFG 191/02)

Coordinates: 55.820668, 12.032673

Jens Winther Johannsen

The sample comes from the northernmost of two burial mounds called Iversbjærgene, which were previously found close to the western shore of Roskilde Fiord. The remains of the burial mound were excavated prior to development. The mound was surrounded by stones and its diameter was estimated at 7–8 metres. At the time of the excavation, a layer about 20 centimetres thick was preserved. This included shells, possibly because the sod for the mound was collected from the nearby salt marsh. Under the central and southern parts of the mound two postholes and a circular pit, just under 1 metre in diameter, were found, all of unknown function. Near the centre of the mound an apparently intact part of a grave, less than 1 square metre in size, was found. The grave contained human bones - parts of the skull, left scapula, upper arm and forearm. The bones were identified as those of a male aged 25–35 years. The individual had strong and large bones and must have appeared robust. Moderate calculus was observed on almost all of the teeth (19 preserved). No other pathological changes were observed. The deceased had apparently been laid on a stone slab and covered with a layer of white (lime?) stone. Flint flakes and a transverse arrowhead were found in and around the grave. Burnt and unburnt bones, including finger bones, were found in a heavily disturbed area to the east of the grave.

CGG_2_106515 (Strandlunden II Gerlev MFG 191/02, stone cist) has been dated to 3070±15 BP (UCIAMS-272877, >30kDa collagen yield (%) 1.2, d15N (‰) 8.9. d13C (‰) -19.4, C/N (atomic) 3.2 (1410–1270 BCE, Montelius Period II)).

#### Denmark, Zealand, Holbæk county, Ods hundred, Asnæs parish, Aastofte (030401-142)

Coordinates: 55.842, 11.422

Johan Zakarias Gårdsvoll & Mads Dengsø Jessen

In 1945 (j.nr. NM1 773/45) during work in a sandy hill at Aastofte an inhumation grave (Grave 1) under a flat field with preserved skeletal remains was disturbed. The find was reported to the

National Museum of Denmark and Peter Vilhelm Glob was sent to inspect the site. Next to the first grave Glob found another grave (Grave 2) which had been heavily disturbed by digging for sand and ploughing. No grave goods were recorded in neither of the graves. The positioning of the bodies in the graves suggests that the date of burial occurred in the Early Roman Iron Age.

CGG_2_107440 (NM1 773/45, Grave 1).

#### Denmark, Zealand, Stålmosegård (020512-04)

Coordinates: 55.6301, 12.1373

Mette Høj

Stålmosegård is a large burial ground from the Roman and Early Germanic Iron Age. The excavations have spanned a 40-year period. They were begun by the National Museum in 1972 and completed by ROMU in 2013[442–444]. The burial site is located on a significant sand hill north of Vindinge village and stretches in a 135 m long and 30–40 m wide belt oriented northeast-southwest; an area of 4500 m2 in total. The inhumation burial custom was dominant in Zealand during this period. This also applies at Stålmosegård, where 112 inhumations have been excavated, almost all oriented N-S. The main part dates to the Late Roman Iron Age C2-C3 and the Early Germanic Iron Age – a period in which the burial ground shows continuity, which is a peculiar view on Zealand, where most burial grounds from the Late Roman Iron Age ended in C3. Cremation graves, on the other hand, are virtually absent on Zealand, so it is remarkable that such have been found from both C3 and D at Stålmosegård, and that these appear to be linked to a ditch system that ran through large parts of the burial site. Stratigraphic observations and excavation data indicate that the ditches are contemporaneous with the burial site's period of use in C3-D. It has been suggested that they may mark a division of the burial field into families/kindreds, but finds of cremations embedded in the ditches suggest that this is unlikely to be the full explanation. For now, the ditch system stands as a unique and somewhat enigmatic phenomenon.

As far as construction and inventory are concerned, the graves at Stålmosegård do not stand out from the other burial grounds of the period in Zealand: inhumations without significant elements of imported goods or local prestige items. There are both coffin burials and simple graves without traces of a coffin. At least 10 large graves are interpreted as chamber graves. 17 graves had been re-opened, which is interpreted as ritually conditioned interventions that may have included the removal of not only objects, but also bones or other parts of the buried person. Re-openings are known not only in Denmark, but also in large parts of Northern Europe.

Among the grave goods, locally produced objects such as ceramic vessels, combs, knives and spindle whorls dominate. It is the women who appear most visibly through rich sets of jewellery and costume accessories. The site's richest grave is grave CL (dated to C3), which, in addition to an elaborate breast ornament, contained imported items such as a Roman denarius, struck under Commodus 175–176, as well as a seed of the bladdernut (*Staphylae)*, found growing wild in the southeastern European area. The closest parallels are to be found in graves in Poland and southern Germany.

The bone material was poorly preserved in the sandy subsoil, and only four graves can be osteologically sex determined. In addition, another grave has been singled out as a boy's grave in the DNA analysis from the Center for Geogenetics at the University of Copenhagen (sample ID CGG_2_106516 to CGG_2_106519, Table S7.2.24).

| CGG No | Museum id |
|---|---|
| CGG_2_106516 | ROM 555 AUx34 |
| CGG_2_106517 | ROM 555 Ax1 |
| CGG_2_106518 | ROM 555 CÅx1 |
| CGG_2_106519 | ROM 555 DNx1 |

**Table S7.2.24:** List of samples studied from Denmark, Zealand, Stålmosegård.

#### Denmark, Zealand, Varpelev, Varpelev Vest (Lillevangsgården) sb No 6

Coordinates: 55.365, 12.188

Section Author 1, Section Author 2 & Section Author 3

Burial examined in 1861 by A. Stunk. The grave was found during gravel digging . A richly furnished burial dating to the Late Roman Iron Age[380]. The grave from this site has earlier been sampled for genetic information, grave 1, but it gave no information[410].

CGG_2_107413 (NM 19674-85) is a mature male buried with gold spiral finger-ring, a bronze bucket, a bronze ladle and strainer, three glass vessels, a bronze mount for a wooden bucket, 13 gaming pieces of glass, sherds from a ceramic vessel and animal bones.

#### Denmark, Zealand, Varpelev[380]

Coordinates: 55.365, 12.188

Grave 1 was sampled by Melchior et al. (2010)[410] but gave no genetic results.

CGG_2_107412 (NM C3059-97a, Varpelev IV, Grave n).

CGG_2_107411 (NM C3059-97a, Varpelev IX, Grave P).

#### Denmark, Zealand, Aamosen-Gammelvind

Coordinates: possibly in the vicinity of 55.583, 11.533

Anders Fischer

Germanic Iron Age (650–704 CE)

Fragment of a human skull, which was found within a portion of mercantile peat fuel during the 1940s or early 1950s. A previous owner of the specimen, amateur archaeologist Eigil Gammelvind Petersen, was informed that the fuel derived from Major Aamose, possibly the part of the bog that belongs to the neighbourhood of the town of Sorø (personal communication). A sample of the *maxilla* or *os zygomaticum* was sent to the isotope laboratory at Bradford University, which reported the following values: $\delta^{13}C$ -18.33, $\delta^{15}N$ 12.6, C:N 3.2, collagen yield 17.0%. Subsequently, a part of the unfiltered collagen extract was sent for AMS dating in Poznan.

CGG_2_017649 (Aamosen-Gammelvind), has been dated to 1345±30 BP (Poz-17028).

### Norway

#### Norway, Central Norway, Aunvollen, Steinkjer, Trøndelag (T20248)

Coordinates: 64.1641, 11.8140

Birgitte Skar

The Aunvollen grave was investigated through a rescue excavation in 1981 by Lars Stenvik. Originally the grave was probably a simple inhumation grave. The grave was relatively disturbed from ditching when it was found. Human skeletal remains from three individuals were found. Two of the individuals were considered to be women aged between 35–49 and 19–20 respectively, in addition, an infant app 1–1,5 years old was found. Of particular interest were the male grave gifts found in the grave, consisting of a sword, sickle, comb, playing dice, scissors, knife. Some recovered rivets may either be the remains of a boat or a wooden cist. $^{18}O$ analysis of the material indicates that both adults had spent some time during their life either in Europe or the British Isles[445]. Several gravemounds are known from the same area.

One of the adult individuals has previously been sequenced mtDNA H[446].

A7437/T20248 the infant, Modern (Tra – 14441).

A7437/ T20248 the young woman, 651–759 CE (Tra14912).

A7437/ T20248 the older woman, 649–673 CE (Tra15329).

CGG_2_105600 (A3729/T2408 b).

CGG_2_105601(A3729/T2408 c).

CGG_2_105602 (A3727/T2413).

CGG_2_105603 (A3727/T2413).

#### Norway, Central Norway, Eid, Åfjord (T14795h)

Coordinates: 64.0212, 9.9131

Merete Moe Henriksen

In 1933, 8 human teeth, animal bones, and seashells were found in the rock shelter Epskaret on the island of Lines, a small island situated 3 kilometres off the coast of Central Norway, north of the Trondheim fjord. Located 45 m above sea level the rock shelter faces the Atlantic Ocean in the west. The rock shelter measures 8 by 5 metres reaching a maximum height of 4–5 metres at the entrance. Its height gradually diminishes, leaving only 0,5 metres between the floor and the roof in the innermost part of the shelter. In front of the shelter is a small plateau measuring 3 by 8 metres (Henriksen 2014:206).

The excavation in 1933 uncovered a 30 cm thick occupation layer inside the shelter. A thin charcoal layer was also found on the plateau in front of the shelter, suggesting the plateau served as an additional living space. A vertical division was observed in the layer inside the shelter, presumably representing different activity zones. One part consisted of fatty soils mixed with fire-cracked stones, whereas the other part was seen to consist of fatty soils containing seashells and animal bones. In the latter, the 8 human teeth were found, alongside a flint drill, four small flint scrapers and five flint flakes, a disc-shaped pearl, four small needles made of bone, a fragment of a pendant of bone reused as a needle and six semicircular ornaments of antler and bone. The ornaments were found at the lower level of the occupation layer, as were the scrapers, flint flakes and flint drill. The semicircular ornaments of antler and bone were laid out in a circle measuring c. 50 cm in diameter and the human teeth were found inside the circle formed by the ornaments [430].

##### Anthropological study summary

A preliminary odontological examination of the human teeth was undertaken in 2009[430] and demonstrates that the teeth belong to at least three different individuals, all of them adults. Both premolars from the upper jaw and incisors from the lower and upper jaws are present. One of the teeth shows clear signs of rickets (vitamin D deficiency). The teeth are all heavily worn, and the wear is probably caused by sliding movements/grinding, indicating that they were used for functions other than just eating. No evidence of tooth decay/caries or tooth diseases could be identified, and the teeth appear healthy and sound.

A preliminary analysis of the animal bones was carried out by Gordon Turner-Walker in 2010 mainly to clarify whether there were more human remains amongst the material, but also to provide information on the use of the shelter. No human remains other than the teeth were identified. As fragile and well-preserved fish bones were found in the same layer as the human teeth, one would expect to find additional human remains if these had been present. This strongly suggests that the human teeth were deposited in the shelter separated from the skull. Among the animal remains, bones from fish, seabirds and seals were recorded, as well as bones from a small canid (probably a fox). Along with the large number of seashells, the animal bones in the cave link its use to the exploitation of marine resources.

CGG_2_105640T (Individual T 14795h, premolar, sex: XX, Mt DNA: U4c2a). CGG_2_105641 (individual T14795h) did not yield enough DNA.

One tooth from the rock shelter had been dated in 2009 to 2456–2202 BC (BP 3850 ± 35 BP, TUa-7563).

#### Norway, Central Norway, Følstad, Trøndelag, Inderøy gnr 151/35 (T16444)

Coordinates: 63.8409, 11.3867

#### Birgitte Skar

The human remains were found in a cairn located on a terrace 71 metres above sea level and 400 metres from the shore. From the site there are wide views towards the Trondheim fjord and parts of Verdal. The cairn along with the human remains were found during work in the limestone quarry "Hylla kalkverk" in 1946. What was left of the cairn was excavated by A. Eisassen in 1946[447]. The cairn had a diameter of 13 metres and a height of 1.30 metres. In the centre was a large oval boulder. The human remains were found underneath this boulder. No traces of a stone cist were present, and most likely the dead was placed directly on the underlying bedrock. From the positioning of the bones, the dead appears to have been laid out orientated east-west with the head towards the east. No burial goods were recorded. 3–4 metres from the centre was an inner kerb made up of large boulders. Fragments of a human skull were found 25 metres east of the cairn but was not handed in to the Museum. Following the excavation in 1946, K. E. Schreiner identified the remains as belonging to a male, aged c. 40 years. A more recent osteological analysis carried out by Hilde Fyllingen identified the bones as belonging to a female, aged 25–30 years although the post-cranial skeleton is robust[448,449]. DNA analysis now confirms that the individual was a male. The individual would have had poor dental health as an abscess was recorded on the lingual side of M2 and on the buccal of M1. Spondylosis was detected in a fragment of the vertebral body. Pronounced muscle attachments of the arms indicate hard physical labour[448].

CGG-2-105643P (T 16444, petrous bone, Mt DNA H5, sex XY) has been dated to 1985±15 BP (UCIAMS_272873 d13C -20.74, d15N 11.41, >30kDa collagen 2.1 C:N 3.1) .

CGG-2-105644 (tooth, sample M2, NA).

Norway, Central Norway, Trøndelag Inderøy, Sund Gnr 136/1 (T18863/A5186)

Coordinates: 63.8604, 11.3070

Birgitte Skar

The Sund site was found in the autumn 1967 and was first excavated by Sverre Marstrander, the year after more bone material was found and Oddmunn Farbregd excavated the rest of the site from 5-7th of August 1968.

Sund is situated at a small promontory on the inlet of the Borgenfjord, which is a branch of the Trondheim Fjord. The finds were discovered in a gravel pit from which masses were removed, thus the site may well have been larger originally[451]. Large amounts of human and animal bones were found seemingly directly under a 20-30 cm thick layer of topsoil, covering an area of app. 10-15 metres in length. The material was divided into 6 concentrations, of these concentrations, concentration 4-5 contained bones from 16 human individuals and some animal bones densely packed. In this concentration there were found 3-4 larger stones and at one end of this concentration a standing flagstone was documented. The colour of the soil in this concentration was reddish-yellow, thus differing from the other concentrations. A simple bone needle was also found here. It is suggested by Farbregd that this concentration may have been remnants of a cist. Farbregd also refers to a small pile of stones comprising one flagstone and several fragments, observed by the first excavator of Sund, Sverre Marstrander. Farbregd suggests that these stones are the remnants of cists that have been partially removed. Thus the human individuals may originally have been buried in cists, a confinement that also explains the bone concentrations. According to Farbregd the use of stone cists also explains the difference of 400 years between individuals in concentration 6 versus concentration 4–5, but in particular it clarifies the multiple individuals that in his mind may have been buried successively in concentration 4-5.

Sund is suggested by Fyllingen (2010) to constitute at least 22 individuals (according to our analysis more likely 31-32 individuals). Adults of both sexes, juveniles and children were present. The lack of clear anatomical connection between skeletal parts, Farbregd et al.[451] take to indicate that the skeletons had either been in a stage of heavy decomposition or that the skeletons had been cut up at the time of deposition. This has, however, not been confirmed in later analyses by neither Torgersen nor Fyllingen. Close to the skeletons was a charcoal- and soil-filled pit and around the circumference of the site was a semi-circular ditch. However, neither the pit nor the ditch could be directly stratigraphically connected to the constructions with skeletons. The pit has, however, been dated to be contemporaneous with the human remains in concentration 6. Farbregd suggests that the firing in the pit is related to the burial activity and potentially to a death cult. The only artefact found on the entire site was a bone needle of a type normally attributed to the Late Neolithic/EBA.

Within the human bone assemblage, bones of rodents, small ruminants, horses and mammals were found. While Marstrander suggests that the ditch could be remnants of a house, Farbregd summarises that 1) the skeletal remains and the coal pit are contemporaneous – dating the site to EBA. 2) The circular ditch is the remainder of a round grave monument, potentially a cairn that has later been removed. The cairn would thus have been 12 metres in diameter. 3) The piling of skeletons would be the results of use of cists. 4) Re-opening and burial would likely have happened successively, also causing some mixing of the context both by animals and humans. Fyllingen (2010) on the other hand states that on the whole the site can neither be interpreted as an ordinary burial site, nor as a settlement site. One would assume that more artefact material and structures were recovered if Sund had been either of the two. Fyllingen suggests that her analysis of the human remains rather indicate a massacre.

The following description is based on Fyllingen 2010 and the original osteological analysis by Johan Torgersen[451]. The Sund material is very fragmented, and fragments were not labelled and numbered individually at the time of recovery. It was, however, sorted according to individuals. Due to the decomposed character of the individuals and a situation where several individuals were deposited together and potentially even disintegrated before burial, it is assumed that such sorting would still imply a certain amount of interpretation. When the material was sampled for the present project, it was deposited in the Schreiner Collections. Here the material was packed in individual boxes and plastic bags according to provenience on the site. The material is thus estimated to comprise between 20-30 individuals, Fyllingen[449] uses 22 as a minimum number of individuals.

Sexing is only accurate for one female and 3 males. Age distribution is as follows: 14% were under 6 years, 32 % between 6-15, 19% between 15 and 25 years, 35% were adults. Fyllingen comments the high number of children/ juveniles as being unusual; while child mortality is expected to be high during the first 4 years, the reported 32% for juveniles represents a large proportion of the total number of individuals.

The material has been analysed with regard to pathology and trauma. Only 2 of the 22 analysed individuals showed neither pathology nor trauma. The results can be summarised as follows: 7 individuals all older than 25 showed signs of cuts ante-mortem, peri-mortem and post-mortem. 13 individuals showed pathology, 3 of these were children with signs of malnutrition and signs of infection, 9 adults suffered spondylosis, *osteochondritis dissecans* and various degenerative changes, enamel hypoplasia, signs of infection. The evidence of both old and new trauma of both defensive and fatal character was found in 54 % of the adult individuals at Sund. For more detailed information on the character of injuries, see Fyllingen[449]. None of the children had injuries indicative of trauma, this, however, does not exclude that injuries occurred. Only violent attacks with weapons would leave their mark on the very small bones of these skeletons.

The dental material reflected a coarse diet from an early age; even children's teeth were severely worn. Caries were only found in a couple of cases. This indicates a diet which is low on starchy foods that breaks down to sucrose or sugar. Secondary dentine was present in most of the adults, where molars were worn down evenly at the chewing surface. According to Fyllingen this could be a result of a diet based on cereal products that contains a great deal of minerogenic material from grinding.

Rickets disease was detected in three cases indicating that approximately 12 % of the population suffered from Vitamin D deficiency. This disease already starts in childhood and results in bending of the weight-bearing bones often followed by fracturing. This is particularly surprising as the site is very close to the sea with an abundance of marine food resources. The disease has, however, previously been found in Danish and Norwegian Neolithic individuals. The presence of Rickets, Harris lines, dental hypoplasia and *cribra orbitalia* is a sign of long-term diet insufficiency/ starvation that is found in 20% of the children's remains from Nordland in the north to Østfold in the south. Fyllingen takes this to be a sign of food restrictions and potentially that some groups did not eat seafood.

30 samples CGG_2_105604 - CGG_2_ 105633 from an estimate of 21 individuals from Sund were sampled for DNA sequencing and further analyses, these represented 14 adults and 7 juvenile/subadults. The children's teeth in the collection are without roots or very badly preserved. From the sampled individuals we have sampled petrous and mainly molars, but in a couple of cases incisors were the only opportunity to get material that was well enough preserved. This in effect means that from the present analysis, we will not be able to state the potential family relations for the children from Sund.

CGG_2_105604 (A5186/T18863, Sund box A cranium A) W3a1c

CGG_2_105605 (A5186/T18863, Sund box A cranium A) W3a1c

CGG_2_105606 (A5186/T18863, Sund box A cranium A) NA

CGG_2_105608 (A5186/T18863, Sund box A cranium E) NA

CGG_2_105609 (A5186/T18863, Sund box A cranium E) NA

CGG_2_105610 (A5186/T18863, Sund box A cranium B) J1c2h

CGG_2_105612 (A5186/T18863, Sund box A cranium C) U5a2c

CGG_2_105614 (A5186/T18863, Sund box A 4b) W3a1c

CGG_2_105615 (A5186/T18863, Sund box A 4b) W3a1c

CGG_2_105617 (A5186/T18863, Sund box A cranium F ( 24)) NA

CGG_2_105622 (A5186/T18863, Sund Box B Sund 3) U4b1a3a

CGG_2_105623 (A5186/T18863, Sund Box C) J2a1a1a

CGG_2_105626 (A5186/T18863, Sund box D Sund 4 nr 26) U5b2a2c

CGG_2_105628 (A5186/T18863, Sund box D Sund 4-5 nr 9, 18) T1a1

CGG_2_105629 (A5186/T18863, Sund box D Sund 4-5 nr 9, 18) NA

CGG_2_105631 A5186/T18863, Sund box E Sund 5 (1)) K2a5a

CGG_2_105632 (A5186/T18863, Sund box E Sund 5 (2)) J1c5a1

Further information on dating: The bone needle found at Sund dates the site typologically to the Late Neolithic/ Early Bronze Age.

Old Radiocarbon dates: Skull from concentration 4 – 5 ( Box A) has been dated to 3300± 100 BP. Femur from concentration 6 has been dated to 2900 ± 90 BP. Charcoal from the coalpit has been dated to 2950± 90 BP. Table S7.2.25 provides information on a selection of analysed specimens.

| CGG ID | Dating lab | individual | $^{14}$C dating | MtDNA |
|---|---|---|---|---|
| CGG_2_105612 | UCIAMS-275208 | box A cranium C | 3485±15 BP | U5a2c |
| CGG_2_105628 | UCIAMS-275209 | box D Sund 4–5 nr 9, 18, petrous | 3505±15 BP | T1a1 |
| CGG_2_105604 | UCIAMS-282658 | box A cranium A, petrous | 3445±20 BP | W3a1c |
| CGG_2_105608 | UCIAMS-282659 | box A cranium E, M1 | 3595±20 BP | X2b+226 |

**Table S7.2.25:** List of some of the samples studied from Norway, Central Norway, Trøndelag Inderøy, Sund Gnr 136/1, their dating and genetic analysis.

#### Norway, Central Norway, Trøndelag, Bergvin (T12775)

Coordinates: 63.74603, 11.2983

Merete Moe Henriksen

The human remains of two adult individuals (cranium and post-cranial bones) were found in a cairn located north of the hill “Bergvinshaugen” 1 km from the shore, on a large promontory with wide views towards the Trondheim fjord[452]. The cairn and most of the human remains were found during the extraction of gravel in 1922, and the cairn was excavated by Theodor Petersen the following year[453].

The cairn had a diameter of 13 metres and a height of 1.75 metres. The stone cist with the remains of two individuals was 2 metres long and had been constructed using a dry-stone technique, where smaller slabs had been placed on top of each other. The stone cist was orientated north-northwest – south/southeast. No artefacts were found. A human skull was discovered at the southern end of the cist, which led Petersen to conclude that the dead had been buried with the head towards the south. The lower jaw was found next to the skull between the skull and the eastern wall in the cist, with the chin bone facing away from the skull, indicating rodent activity. Post-cranial bones were found in the northern part of the cist, between the smaller slabs in the eastern wall, as part of the wall had collapsed. Large amounts of Littorina were found in the cist, especially in the southern end close to the skull.

An osteological analysis carried out by Hilde Fyllingen identified the bones as belonging to one male c. 35 years of age and that of a woman aged 25–35 years. Rickets (Vitamin D deficiency) were observed in the femur and ulna of the female. Animal bones of horse or cow were also present[448,449].

No radiocarbon dates exist from this burial. As no burial goods were found, a likely date to the Bronze Age is based on the cairn`s location in the landscape which is similar to most Bronze Age cairns in the area as well as the construction of the cist with walls made up of slabs. This detail can be observed in several cists at the Bronze Age burial site at Toldnes in Steinkjer and Skjeggesnes in Alstahaug (T 18383). The occurrence of Littorina in the burials is also mentioned in Karl Rygh`s excavation reports from Toldnes[454].

CGG_2_105634 (T12775, tooth M1).

CGG_2_105635T (T12775, tooth M2), Mt DNA: U2e2a1a2, sex: XX.

#### Norway, Central Norway, Trøndelag, Bøle, Flatanger M

Coordinates: 64.375798, 10.631408

Torkel Johansen

Cave find, Early Iron Age. The remains of (minimum) 3 individuals were found by a student in the cave "Bølekonta" in 1986. Parts of the skeletons were taken in during a subsequent police investigation. An archaeological inspection was carried out in 1987, as well as a subsequent archaeological investigation in 1988. The original report identified two individuals (Herje 1987); however, an anthropological assessment by Sellevold (1988) identified a third individual. A total of 4 hearths were found in the cave, some of which contained a lot of burnt bone material. A partially burnt log was also found in one of the hearths (unr. 33). SW in the cave was also a small collection of elbow shells (unr. 8). The skeletons, netting, shells, hearths and possibly also animal bones from the site indicate that the cave has been a place of residence for people over a period of time.

The human remains were located in an approx. 30 sq m. large room in the cave. The human remains were found partly disarticulated.

CGG_2_107036 (skeleton 1, T21061:2, A7271, tooth C) has been dated to 1570 ± 90 BP, cal. AD 333–649, d13C (‰) -18,8 (lab.ref. DF-1848, T-7760). This individual must have been lying between the hearth and the eastern wall of the cave, with its head to the SE. Other human remains were however scattered around the cave floor. The pathological damage to skeleton 1 (cut marks in the pelvis) indicates that this individual has been exposed to lethal violence.

CGG_2_107035 (skeleton 2, T21061:3-4) has been dated from differents elements to:

1860± 140 BP, cal. AD 174–442, d13C (‰) -19,5 (lab.ref. DF-1848, T-7354).

1570± 50 BP, cal. AD 412–598, d13C (‰) -20,7 (lab.ref. DF-3754, TUa-4820).

1630± 35 BP, cal. AD 364–545, d13C (‰) - 20,9 (lab.ref. DF-3754, TUa-4821).

The bones from Skeleton 2 in the report (T21061:3-4) were found scattered in the cave; the skull was wedged under a stone, while the femurs lay together in the NW, together with other bones and a stone sinker. The damage to skeleton 2 may have occurred postmortem. The remains were disarticulated in the cave.

The discrepancy in the dating results suggest that the disarticulated remains catalogued as Skeleton 2 represent multiple individuals.

In addition, three fireplaces in the cave have been dated:

Fireplace no. 1, charcoal (museum no. T21061:41/q): 1740± 70 BP, cal. 129–436 (lab.ref. DF1848, T-8314).

Fireplace no. 2, charcoal (museum no. T21061:42/g): 1460± 70 BP, cal. 428–674 AD (lab.ref. DF1848, T-7761).

Fireplace no. 4, charcoal (museum no. T21061:43/k): 1750± 70 BP, cal. 124–435 AD (lab.ref. DF1848, T-8312).

Calibration curve: OxCal v4.4.4 Bronk Ramsey (2021); r:5; Atmospheric data from Reimer et al (2020). Museum no. T21061:2, Anatomical no. A.7271.

#### Norway, Central Norway, Trøndelag, Bomo, Snåsa M (T17680:c/007)

Coordinates: 64.263118, 12.446387

Torkel Johansen

Chamber burial, 4th century AD. Found in a previously disturbed chamber measuring approx. 2,3 x 0,8 m, which had previously been covered by several slabs[455]. The assemblage contained a clay pot, a bronze needle, parts of a fragmented human cranium and long-bone fragments. Excavated by F. Gaustad.

CGG_2_107020 (T17680:c/007, left petrous).

#### Norway, Central Norway, Trøndelag, Hitra Island, Fauskland gnr 8/1 (T11792 c)

Coordinates: 63.6057, 14.8586

Birgitte Skar

The skeletal material and artefacts were found in 1917 while digging gravel in the tidal zone on the farm of Fauskland, in Fausklandsvågen in the innermost part of Barmfjorden on the island of Hitra[456,457]. The gravel was dug at low tide while the site would in 1917 have been submerged at high tide. The material was first discovered when the gravel was spread on a road leading to the farm, when human bones were discovered. In the same let of gravel a dagger was found. Further investigations in the gravel pit by the Fjord revealed several skeletal fragments and a wrist-guard made of bone at app. 30–40 cm depth. There is no doubt in the mind of the finder that the bones, the dagger and the wrist-guard belong to the same context. The location of the material is estimated to have been deposited about 3 metres below the contemporary shore-level; although mentioned as a grave[281] the find circumstances indicate a drowning and rapid encapsulation in marine sediments.

The human skeletal material represents 41 fragments of a skeleton including the top of the skull and a lower jaw with 6 molars as well as many postcranial elements.

The artefacts found in this context consist of a flint dagger of type I X and a wrist-guard made of bone. The artefacts are considered to belong to the individual.

CGG_2_105916, CGG_2_105917T and CGG_2_105918 representing M1, M2 and M3 respectively, Mt DNA: J2a1a1a, sex: XY.

Two samples from T11792c have been radiocarbon dated in 2022 to 3682+/-12 BP (Tra 17220) and 3682± 13 BP (TRa 17221) representing the skull and the jaw of the same individual respectively.

#### Norway, Central Norway, Trøndelag, Jøa, Namsos M

Coordinates: 64.6544, 11.2084

Torkel Johansen

Burial, Late Iron Age. Part of an unusual Late Iron Age cemetery where the individuals were buried in a sitting position in pits (hocker-type burials). Excavated by S. Marstrander in 1965–1966[455]. 23 burials were excavated in total. The burials had the form of circular cylindrical pits dug into a shell-sand bank, without visible markings. The pits were usually 1 to 1.5 m deep. The dead had been buried sitting at the bottom of the pits with their legs pulled up under them.

CGG_2_107026 (Jøa burial XV, Namsos M, T18653, A7444, right petrous) has been dated to 1575±20 BP ( UCIAMS-282670, >30kDa collagen yield (%) 7.4, d15N (‰) 17.0. d13C (‰) -15.3, C/N (atomic) 3.2). This burial contained the remains of a female individual, ca. 30 years old. The bones were deposited in a 68 cm deep pit, but occurred to have been disarticulated or disturbed after deposition. Preserved remains consisted of cranial fragments, 5 vertebrae, a femur and one unidentified long-bone. The burials also contained a small knife and nails. Osteological determination by Bernhard Getz. Marine diet.

CGG_2_107027 (Jøa burial XI, Namsos M (T18650, A7441, left petrous) has been dated to 1325±20 BP (UCIAMS-282671, , >30kDa collagen yield (%) 5.6, d15N (‰) 10.1. d13C (‰) -20.9, C/N (atomic) 3.2). This burial contained the remains of an male individual ca. 40 years of age, height approximately 163 cm. The inhumated remains were deposited in a 118 cm deep funnel-shaped pit, covered by ash-filled earth, fire-cracked stones and charcoal. A knife was deposited near the deceased's left hip. Osteological determination by Bernhard Getz. Terrestrial diet.

CGG_2_107028 (Jøa burial X, Namsos M (T18649, A7440, right petrous). This burial contained a male individual, ca. 165 cm of height according to the osteological analysis by B. Getz. The burial contained however a brooch, 2 knives, 4 beads and 2 combs - indicating a female assemblage according to the original catalogue. The burial was situated in a sitting position in a stone covered pit, 110 cm deep. The lower part of the body was in its original place; the upper part of the body had however been displaced by the weight of the earth-fill.

CGG_2_107032 (Jøa burial I, Namsos M (T28746, A4991, right petrous) is an adult female, ca. 45 years old. The burial had been dug out before the excavation to a depth of 110 cm. It is assumed that the disarticulated bones unearthed prior to the excavation come from this burial. There were no artefacts retrieved. Terrestrial diet. It has been dated to 1330±20 BP (UCIAMS-282672, , >30kDa collagen yield (%) 4.7, d15N (‰) 10.7. d13C (‰) -20.7, C/N (atomic) 3.2).

CGG_2_107033 (Jøa burial V, Namsos M (T18646, A4993, tooth, M3, maxilla). This burial contained an undisturbed skeleton of an adult female ca. 40 years old, 157 cm of height, buried in a hocker position. The burial pit was funnel shaped, and contained ash-filled earth and fire-cracked stones. A singular knife accompanied the deceased. Terrestrial diet. It has been dated to 1350±20 BP (UCIAMS-282673, >30kDa collagen yield (%) 4.4, d15N (‰) 11.4. d13C (‰) -20.5, C/N (atomic) 3.1).

CGG_2_107034 (Jøa burial VII, Namsos M (T18648, A4995b, A7439, right petrous) is an adult female, ca. 60 years old., ca. 160 cm of height. The skeleton appeared undisturbed, but had been partly displaced due to the pressure from the surrounding soil. The burial contained (apart from the human remains) an early oval brooch and a knife. The pit was refilled by ash-filled earth and fire-cracked stones.

#### Norway, Central Norway, Trøndelag, Leka, Solsem gnr 10/2 (T10834)

Coordinates: 65,0537, 11,5510

Birgitte Skar

The cave on the island of Leka in Trøndelag was found by some boys in spring 1912 and later excavated and documented by Theodor Pettersen in 1912-1913. Pettersen considered that the cave was untouched since prehistory due to a challenging access. The cave belongs to a group of caves in central and northern Norway containing prehistoric wall paintings. The documentation from the excavation is unfortunately less satisfactory by modern standards.

The cave is approximately 40 metres long and the archaeological remains can be described as three separate areas following the topography of the cave. The entrance is blocked by a large boulder. Area I consists of an extended area from the entrance and app 11 metres into the cave, this area is like the entrance very steep, going down from the cave entrance to an area app. 10 metres below - the cave floor (78 m.a.s.). The boulder and the steepness are of importance since this would have been a hindrance for both animals and humans to enter the cave thus the cave interior was at the time of discovery free from modern traces of use. Area II is narrower but widens further into the cave, this is where an excavated cultural layer (app 10–20 cms) covered by an app 50 cm deep top layer of accumulated dust and plant remains revealed 1534 bone fragments and artefact material. The cultural layer also comprised shells and charcoal. The stratigraphy in Area II was considered to be intact at the time of excavation The walls of Area II converge to end in a sharp angle towards Area III. Area III which lies in complete darkness has a floor that is covered in water-rolled stones and stones that have fallen from the roof. Here artefacts and bones were found between and under stones, there is no clear cultural layer in this area. 334 fragments of bones were according to Pettersen`s original report found scattered on the surface of Area II and Area III. Several fireplaces or traces of fire were established from between Area II and Area III, as well as in the darkest part of Area III. Two lines of stones demarcate an entrance to Area III and yet another entrance to the innermost part of the cave with wall-paintings.

The skeletal material was analysed in 1913 by Herluf Winge, the report is a handwritten letter to Theodor Pettersen. There were findings of human bones both in the central cultural layer (Area II) and in the innermost part of the cave (Area III) in relation to the cave paintings. The samples for DNA and radiocarbon dating were taken from Area III in the innermost part of the cave. The contexts included all together human skeletal remains from 3 – 4 adult humans and part of a jaw with three teeth from a baby (less than 6 months old). No distinct signs of violence, cutting or ocular signs of disease could be stated from the human bones. In addition, animal bones were found in the cave: fish and bird bones, mammal bones consisting of: Otter, harbour seal, harp seal, unspec. seal, goat, sheep, cattle, horse, water rat[458], thus a mixture of wild and domestic animals. Unfortunately, only a little of the animal osteological material has been retained in our collections. In Area II of the cave the assemblage includes an 8 cm long bodkin made from the radius of sheep or goat. A rectangular and worked piece of antler, two pieces of a quartzite grind-stone, while Area III contained a 9 cm long point made from antler, a 8,7 cm long carved toe-joint of harpseal with incisions making it resemble a rough sculpture of a man and suggested by Bjerck[459] to be a “brummer”, an 8,6 cm long point of violet slate, a 5cm long carved figure made from the limb of a bone of cattle depicting a sea-bird. This piece has been suggested to depict the great auk, a 9,7 cm long whistle-like object made from bone of a larger sea-bird. In addition some pieces of wood were found both in Area II and III.

Of particular interest are the cave paintings that consist of altogether 24 drawings in red paint (Norsted 2011; according to Sognnes 2002 only 21). The two main panels consist of an eastern wall comprising 14 human figures in different size, some of them distinctly drawn as men – this panel is extended with a simple but very large cross. While on the western cave wall the panel includes all together 10 figures, some of them less distinct due to preservation, but assumed originally to have been human figures and potentially part of a cross. For a more detailed description – see references below -the paintings have been the focus of the literature.

CGG-2-105642T (TRa-17217, tooth I, DNA: j1c2h sex: XX) has been dated to 3300 ± 15 BP (rounded date).

Further information regarding dating:

Two human samples from Area III of the inner cave were radiocarbon dated in 2022: TRa 17217 from the femur of a small child: 3302 +/13 BP and TRa 17218 from the pelvis of a grown up individual: 3433+/13 BP. TRa 17218 is potentially from the same individual that has provided the sequenced incisive ( CGG-2-105642). Earlier two radiocarbon dates have been made: one (T-4295) from shells (*patella vulgate*) found in the cultural layer giving 3310± 80 BP calibrated to 1715± 155 BC. One (T-4429) from a bone of sheep or goat (the only animal bones retained in our collections) these bones are incised by a sharp metal edge dated to 2120± 50 BP calibrated to 255± 135. The exact location of these earlier dated bones is unknown. The slate arrow-head indicates an earlier use and can be morphologically dated to the Late Neolithic, while the main use of the cave is estimated to be Bronze Age. Sognnes[458] writes that the fireplaces in Area III that included preserved wood and possibly the dated domestic bones must derive from a later Iron Age episode. We can conclude that the archaeological remains in the cave are accumulated over time, but are likely associated with episodic use of the cave very likely for ritual purposes.

#### Norway, Central Norway, Trøndelag, Lund, Namsos M (T21438)

Coordinates: 64.7690, 11.5964

Torkel Johansen

Human remains found in a ditch, approximately 100 m from the farmyard and Lund, and ca. 100 m from the shoreline, 5–10 m above sea level. The site lies in a cultivated area on a subsoil consisting of shell sand. The burial was discovered in the ditch profile, which had cut a ca. 1 m deep and 1 m wide pit in which the human remains were located. The pit was located on a flat area below a ridge where a small burial ground is located. The site was subject to a secondary archaeological inspection by I. Smestad and L. Stenvik, but no more human remains were found. The osteological assessment suggests that the remains belonged to a male, approximately 19–35 yrs (based on tooth-wear and the fact that the third molar had not erupted), estimated height ca. 160 cm. The burial form closely resembles similar “hocker type” burials from Jøa[455], suggesting a Late Iron Age provenance.

CGG_2_107024 (T21438, tooth, I) has been dated to 1240±20 BP (UCIAMS-282669, >30kDa collagen yield (%) 13.3, d15N (‰) 12.4. d13C (‰) -19.2, C/N (atomic) 3.1).

#### Norway, Central Norway, Trøndelag, Røyne, Ørlandet M (T16237:e/006, A4469)

Coordinates: 63.7158, 9.6422

Torkel Johansen

Early Iron Age inhumation burial, found in a gravel pit deposit at a depth of ca. 0,5 m below the surface. The burial comprised the remains of a female individual ca. 50 yrs of age, height approx. 155 cm (osteological assessment by K.E. Schreiner). The burial goods included a typical “set” of one bone comb, two flat bone needles (”hair-pins”), one round bone needle and a sickle. The assemblage is typical assemblage from phase C3 (Late Roman Period 310/20–400 AD, cfr. Slomann[460], representing a wider set of assemblages typically occurring in middle and northern Norway during this period.

CGG_2_107029 (T16237:e/006, A4469, tooth, M1).

#### Norway, Central Norway, Trøndelag, Storfosen, Ørland M (T15808/A4407)

Coordinates: 63.6586, 9.4218

Torkel Johansen

Female burial, Merovingian Period (late 6th / early 7th century). Found during digging for a foundation wall in the farmyard of the Storfosna estate[461,462]. A female cranium was discovered at a depth of 1,1 m below the surface, along with two small conical brooches, two polyedric bronze needles, 82 beads (blass, clay, amber and bronze), a knife and remnants of fittings for what was likely a small wooden box. The human remains belonged to a female individual ca. 40–50 years of age (osteological determination by K.E. Schreiner).

CGG_2_107038 (T15808 / A4407, tooth, M3, mandibula).

#### Norway, Central Norway, Trøndelag, Valstad, Verdal M (T2186/A0019)

Coordinates: 63.7691, 11.5218

Torkel Johansen

Female burial. Found in a stone chamber, 2 m long and 0,5 m wide, oriented east-west in a large burial mound, 15 m i diameter. Beyond the human remains (T2186), the assemblage comprised a glass beaker, 2 electrum finger rings, 3 metal needles, a rectangular belt-buckle, an iron key and 61 beads (1 silver, 1 amber and 59 glass beads). The combination of a glass beaker (Eggers type 212), a bronze needle of Beckmann form 124 and the finger ring of Beckmanns form 8b places the find in the Late Roman Period C2, 250/60–300 AD[463–465].

CGG_2_107031 (T2186 / A0019, tooth M3).

#### Norway, Central Norway, Trøndelag, Vik, Ørlandet M (T28126:11)

Coordinates: 63.6993, 9.6402

Torkel Johansen

Female burial, Migration period / 5th century AD. Found during professional excavation by G. Grønnesby / M. Aspvik in 2019. The female individual was buried sideways in a shallow pit with no visible markings on the surface. The subsoil in which the pit was dug consisted of marine sediments, sand, corals and shell fragments/shell sand with good conditions for bone preservation. The skeleton was nearly complete, buried with an assemblage consisting of a cruciform brooch, a bone bead, spindle-whorl, scissors, 2 keys, an iron fibula, an iron needle and nails from what was possibly a small box. A C14-sample from the skeleton provided the result 1742± 15 BP, cal. AD 242–345 D (95,4%) (Lab ref. TRa-145200017). However, the cruciform brooch in the assemblage is a diagnostic artifact from the Migration Period[460,466]. According to the excavation report, the early C14 result is possibly influenced by the marine reservoir effect; the burial is also located along the western coast of the Trøndelag region.

CGG_2_107025 (T28126:11, Left petrous bone) has been dated to 1742± 15 BP, cal. AD 242–345 D (95,4%) (TRa-145200017).

#### Norway, Central Norway,Trøndelag, Toldnes (T2408/A3729, T2413/A3727)

Coordinates: 63.9758, 11.3946

Merete Moe Henriksen

The project includes the analysis of five samples from four individuals (three adults and one child) from Cairn IX and Cairn X.

CGG_2_105599 (A3729/T, 2408a).
CGG_2_105600 (A3729/T, 2408b).
CGG_2_105601P (A3729/T, 2408c).
CGG_2_105602 (A3727/T, 2413).
CGG_2_105603T (A3727/T, 2413).

DNA was extracted from two individuals, one from each of the two cairns:

Cairn IX:_CGG_2_105601P (A3729/T 2408c, 1 petrous, adult, sex: XX, Mt DNA: J1b1a1).

Cairn X: T 2413 (burial 2): CGG_2_105603T (A3727/T 2413, tooth M2, adult, sex: XY, Mt DNA: I1a1) (Tua-2471) had been dated in a previous study in 2000 to 3105± 85 BP, cal. 1535–1121 BC (95,4 %).

Cairn II: T 1267 (TUa-2470) had been dated in a previous study in 2000 to 3040± 90 BP, cal. 1497–1019 BC (95,4 %), cal. 1461–1045 BC (92,8 %).

From 1879–1880 and in 1905–1906 Karl Rygh undertook excavations of several cairns on the farms Toldnes, Holan, and Langås, on a small peninsula in Beitstad Fjorden, the inner part of the Trondheim Fjord[463]. In the Bronze Age, the cairns would have been located on small islands and promontories facing the mouth of the narrow passage that previously existed between the island Inderøy and the mainland. In all, Rygh excavated 24 cairns on these three farms, and his excavations remain the most comprehensive investigations undertaken of an Early Bronze Age burial site in Norway.

Many more cairns have been registered in the area since Rygh's time[467]. 10 out of the 24 cairns contained burial goods, and 22 bronze objects were found. 47–48 burials were recorded in the cairns. They contained the unburnt remains of c. 36 individuals. 14 of the burials contained inhumation graves whereas 19 were cremation burials. For 15 of the burials, it is not known whether they represent inhumation or cremation burials[467].

The cairns at Toldnes have probably been used for burials over a considerable period. A c. 2-meter-long stone cist commonly occurred at the centre of the cairns with secondary burials surrounding these. The cremation burials were found in smaller cists. In some of the cists with inhumation burials, more than one individual had been interred whereas other cists were empty.

Animal bones were found in two cists. The stone cists had been constructed using several smaller slabs placed on top of each other in the side walls, or the walls in the cists were made up of standing stone slabs. Some of the cists were filled with shell/beach sand as well as seashells[467,468].

Description of burials with analysed osteological material:

##### Cairn IX (Toldnes – Lille Drakenhaugen)

Significantly destroyed cairn, 15 meters in diameter. Close to the centre of the cairn were the remains of a stone cist, measuring 2, 40 x 0,35/0,55 metres and orientated E-W. The walls of the stone cist were made up of flat slabs, with larger slabs placed at the two ends. The bottom of the cist was filled with gravel and sand as well as the broken fragments of the slabs that would have covered the cist. The unburnt remains of three [448] were found in the eastern part of the cist. The cist also contained fragments of burnt bones, but these could have belonged to a secondary burial that ended up in the cist when the cover slabs collapsed.

##### Cairn X (Toldnes – Limbuhaugen)

This was the largest cairn at Limbuhaugen measuring approximately 17 metres in diameter. Two well-preserved crania and other skeletal remains (T 2409, T 2410) are thought to originate from a cist located at the centre of the cairn (burial 1). To the southeast of this cist was a second cist, 2.2 metres long and orientated N-S (burial 2). It contained a cranium along with other skeletal remains (T 2413). A bronze tweezer with broad, thin pinchers slightly bent in (T2411) was found on the western side wall, towards the southern end (T 2411). Two fragments of a bronze razor were also recovered from the burial. Today only one fragment of the outer edge of the blade is preserved (T 2412). It is decorated with opposing triangular ornaments that form a zig-zag motif along the back of the razor. The other fragment that cannot be located today was sketched for Rygh`s publication of the burial site (Rygh 1906:11, fig. 8). As shown in this sketch, the knife would have had a horse-headed handle with ears forward and small protruding eyes. In the western part of the cairn was a small cist that contained burnt bones.

Anthropological study summary

During Karl Rygh’s excavations, the unburnt remains of c. 36 individuals were found in the cairns at Toldnes, Holan, and Langås. Only some of these bones were handed to the Museum (from Cairn II, IX, and X), while the rest were reburied in the cairns. Consequently, only seven individuals from the burial site have been the subject of osteological analysis. The first thorough examination of the human remains was carried out by Hilde Fyllingen in 2002[448,449].

Fyllingen`s results are as follows[448]:

Cairn II, T. 1267: Sex: Female? Pathology: Ulna is unusually curved. It is possibly a case of Rickets even though this is unusual in the upper extremities.

Cairn IX, T. 2408 (three individuals): Sex: Undetermined. Age: under 5; Sex: Female. Age: 25–35; Sex: Male. Age: 25–35. Stature: 172–173 cm. Pathology: The right Tibia is unusually curved. This is a sign of healed Rickets. The female and the child from the same grave show no obvious signs of pathology or trauma.

Cairn X, burial 1, T. 2409 & T 2410: Sex: Female. Age: 35+ (T 2409), Sex: Male. Age: 35+ (T. 2410)

Cairn X, burial 2, T. 2413: Sex: Male. Age: 25–35. Stature: 185 cm. Pathology: The radial tuberosity, of the right radius, is turned inwards. This is probably due to the tendon being severed (trauma) at an early stage, before the bone was properly developed. On the lower extremities, especially the femur, the blood vessels are abnormally marked on the bone. This could indicate high blood pressure or other cardiovascular conditions. Trauma: The left tibia has small lesions across the shaft. These occurred post-mortem. Appear to be either very narrow gnawing marks from an animal or shallow scraping marks (done by a human). The proximal shaft of the radius has shallow post-mortem marks that appear to be either gnawing or wear.

#### Norway, Northern Norway, Bø, Føre 1 and 2 (Ts 5337/A4732 and Ts 5338/A4729)

Coordinates: 68.6666, 14.4687

Zanette Tsigaridas Glørstad

The two burials are part of a burial site located at promontory facing the Arctic Ocean. Ten burials from the site were excavated in 1955 (Askeladden ID 8915), some buried deep in drift sand. The primary use of the site, as of other burial sites in the surroundings, is reflected by cairn-burials and cist graves underlying stone cairns, dating to the Roman and Migration period (AD 0–550). The use of the area is also reflected through boat houses and settlement remains, and boat burials dating to the Late Iron Age (AD 550–1050).

Føre 1 (Ts 5337/ A4732) was accidentally discovered in 1954, and the items sent to the museum. A well preserved, adult skeleton was placed in a large stone cist covered by slabs and a low cairn. The head was placed towards the south, resting on a stone slab where a clay vessel was also placed. The grave goods included numerous weapons and personal equipment, placed alongside the body. The grave also contained four bear claws and the split skeleton of a large dog. The front part of the dog lay below the deceased’s feet, while the hind part lay across the stone slab that served as a pillow. Beneath the skeleton was a layer of charcoal with large pieces of charcoal, as well as an organic layer assumed to be from a bearskin judging from the claws. Osteologically, this individual was probably a male, aged 20–25. Based on the weapons, the grave is type-dated to the $4^{th}$ century.

Føre 2 (Ts 5338/ A4729) Constructed much like grave 1, but the sides of this coffin were built of rounded stones instead of slabs, and the grave was intact when excavated. An osteologically determined female, aged 30–35[469], was lying on her back with the head towards the south and slightly flexed legs. Several bones, probably originating from a dog, were found by the individual's feet. Among the rich burial goods was a complete Roman glass beaker and a ceramic pot, two bronze hairpins, and two knives. On one of her feet lay a spinning-whorl, and her right hand seems to have held the beak of the seagull. As in Føre 1, the grave contained seven bear claws assumed to be part of an organic layer possibly representing a bear pelt, under the deceased. The dead and the associated grave goods are described as placed in an unusually symmetrical and meticulous order[469]. The grave is typologically dated to the 4th century[470].

CGG_2_107015 (Ts 5337/ A4732, Føre 1, tooth, M1), CGG_2_107014 (Føre 2, tooth, M2).

#### Norway, Northern Norway, Nord-Rollnes (Ts 10340/A5212)

Coordinates: 68.8703, 17.0454

Zanette Tsigaridas Glørstad

The burial was excavated in 1970, following the discovery of the grave during construction work". In addition to the remains of an adult skeleton, the burial included an axe found near the deceased's right hip, a scythe by the individual's left hand, and two knives both placed in a wooden sheath of which some fragments were preserved, placed parallel by the left arm. In addition, the grave contained several iron nails, a piece of flint, and unidentified fragments of iron as well as of animal bones, including the remains of what is assumed to be a cow’s skull. The grave is dated to AD 550–800, the Merovingian Period. The individual is a man of about 60 years old. Based on the layout of the remnants in the grave, the excavation report estimated the man to have been 164,8 cm tall. The teeth bore signs of heavy dental wear, in particular two of the back mandibles which were both asymmetric due to extensive wear.

The listing of CGG_2_107019 under Nord-Rollnes is based purely on the close genetic kinship with CGG107019, as the context and provenance for CGG_2_107019 is uncertain.

CGG_2_107017 (Ts 10340/ A5212, tooth, M2).

CGG_2_107019 (A3729/T2408, tooth M3)

#### Norway, Northern Norway, Nordland, Alstahaug, Skjeggesnes, (T18383/A4924)

Coordinates: 63.7460, 11.2983

Birgitte Skar

The Bronze Age cairn at Skjeggesnes was excavated in 1963 following an incident involving an illegal excavation by local inhabitants who stumbled upon a stone cist with human remains. The excavation was led by Harald Egenæs Lund and was undertaken in only a few days due to the lack of funding. The cairn at Skjeggesnes is situated on a promontory 20–30 meters above sea level and with wide views towards the sea. The excavated and now reconstructed cairn constitutes one of more than 16 cairns that cluster on the same promontory. At least some of these cairns are likely to be of Bronze Age origin[471]. At the time of the excavation in 1963 the cairn measured 20x23 m. On top of the cairn in the mid-section was a slightly elevated construction of stone in the shape of an overturned boat, measuring 12x6 meters and orientated N-S. The centrally placed burial was located just beneath this construction. The stone cist containing the remains of what was considered to be two dead individuals as well as grave goods, was 2 meters long and constructed by using a dry-stone technique, where smaller slabs had been placed on top of each other[472]. The cairn contained well-preserved human bones of what was considered to be two individuals. Following the excavation in 1963, the human remains were examined by Bernhard Getz at the Anatomical Institute at the University of Oslo. A more recent osteological analysis was carried out by Hilde Fyllingen[448]. Fyllingen identified the bones as belonging to a male 35–50 years of age and that of a woman aged 30–40 years. The male individual had degenerative changes to the spine of the vertebra located in the neck as well as degenerative changes at the wrist, due to trauma to the ligaments of the elbow joint. The female would have had poor dental health due to calcified plaque as well as caries[448]. See Tables S7.26 for samples included in this study.

Artefacts from the burial include a bronze razor with a horse headed handle, a bronze pin and the remains of an asbestos tempered ceramic pot. The burial has been typologically dated to Montelius Period III (1300–1100 BC). The following Table S7.2.26 provides the list of the samples studied.

| CGG No | sample | Supposed individual | Mt dna | sex |
|---|---|---|---|---|
| CGG-2-105636T | premolar | Individual 1 male | J1c8 | XY |
| CGG-2-105637T | M2 | Individual 1 male | J1c2 | XY |
| CGG-2-105638T | M1 | Individual 2 female | I1a1 | XX |
| CGG-2-105639 | M3 | Individual 2 female | NA | ? |

**Table S7.2.26:** List of samples studied from Norway, Northern Norway, Nordland, Alstahaug, Skjeggesnes, material and sex.

NB: The project includes sequencing of two teeth from each of the supposed individuals, an adult male (premolar and molar (M2)) and a female (M1, M2). The DNA analysis however indicates that the grave contained two males and one woman based on different Mt DNA.

CGG_2_105637 (AI 4923) is a male individual that had been dated in 2000 to 1535–1121 BCE, 3105±85 BP (TUa-2472).

#### Norway, Northern Norway, Nordland, Enge, Sømna M (T16392:c/003, A4502)

Coordinates: 65.2747, 12.1271

Torkel Johansen

Provenance: Burial consisting of a male adult skeleton, ca. 50 years of age, height ca. 171-172 cm[450]. The burial assemblage also comprised a sword and arrowhead from the Merovingian Period[473]. The burial was found due to gravel extraction from a mound by a local farmer, whereupon a further investigation was made by Einar Høvding. The body in the grave was oriented exactly N-S and lay on the original ground surface; the height from the grave to the top of the mound was 1.10 m. The head rested on a flat stone. Stones also appear to have been placed around the dead. The sword lay by the skeleton's left side. The location lies in a sheltered cove below Nordstrandura on the northern side. There were according to the original catalogue entry 6–7 burial mounds here, consisting of shell sand; as such is not found either in the small cove or on the property in general, the sand must have been brought from elsewhere to build the mounds. Museum no. T16392:c/003, Anatomical no. A.4502.

CGG_2_107037 (tooth, M1).

#### Norway, Northern Norway, Nordland, Holm, Bindal M (T16707:b, A4568)

Coordinates: 65.1713, 12.1152

Torkel Johansen

Provenance: Male burial with skeletal remains and a spearhead. Found during digging of a cellar in a mound of earth, obviously a burial mound, of which there are many on the farm[440]. The skeleton lay at a depth of approx. 1 m from the top of the mound, level with the ground surface and was, according to information from the finder, oriented with the head to the south, facing down. The legs were bent up under the body with the knees towards the west. If these observations are correct, the individual was laid to rest in a squatting position. The spearhead "stood vertically on the skeleton at the height of the shoulder blade and with the point facing down" according to the finder. Marks of the stake could not be seen in the pile fill, which was typical beach gravel. Museum no. T16707:b, Anatomical no. A.4568

CGG_2_107030 (right petrous bone).

#### Norway, Northern Norway, Nordland, Lund, Steigen (Ts 1168-1181/A1511)

Coordinates: 67.9853, 15.1812

Zanette Tsigaridas Glørstad

The archaeological remains were found in 1897 in a burial mound built of stone and earth, with a height of 90 cm and a "fairly wide" circumference. Its exact location is unknown, but the site lies in a small bay at the outer coast, facing the North Sea. Several burial mounds are recorded in the area. The burial was associated with a large variety of smithing tools as well as e.g., a horse bit, a knife with an ornamented bone handle, and 16 bone gaming pieces. The individual was a male adult, described in the museum records as a large built person. The cranium has traces of severe trauma, with a deep cut across the backside of the head, and another cut across the lower crania towards the neck, both cuts possibly from a sword or similar[450]. The finds can be dated no closer than to the Late Iron Age (ca. 550–1050), but most likely to the Viking Age (AD 800–1050)[474].

CGG_2_107010 (Ts 1168-1181/ A1511, tooth, M3).

#### Norway, Northern Norway, Nordland, Tomeide, Nesna M (T19564:l,m,o)

Coordinates: 66.2627, 12.7243

Torkel Johansen

Multiple burial with three individuals from the Early Iron Age, in addition to the disarticulated remains of a fourth. Find assemblage containing knife, comb, strike-a-light, arrowhead and unidentified iron rods. Found during construction work, followed by an excavation by A.S. Alsvik[475] and F. Gaustad. No mound was visible, but aspects of the surrounding layers suggested that a mound with an assumed diameter of 8–9 m had previously covered the burials. 3 skeletons laid side by side with their heads towards the NE. According to the museum catalogue, the skeletons were: A - man, approx. 60 years, B - female, approx. 20 years, C - probably boy, 8–9 years. Above the skeletons, a pit containing the disarticulated remains of a fourth individual, a child aged 12–13 years was also unearthed. The male individual (A) had the remains of a possible arrowhead in his spine. The placement of individuals A–C suggests they were buried at the same time according to the excavator. On the basis of the comb, the primary burials belong to the Late Roman Period (assumed 4th century AD in museum catalogue). The samples stem from individual A and B, as well as the disarticulated skeleton above the primary burials.

CGG_2_107021(Individual A, T19564:l, tooth M3 left).
CGG_2_107022 (Individual B, T19564:m, left petrous).
CGG_2_107023 (individual T19564:o, left petrous).

CGG_2_107022 (Individual B) has been dated to 130–240 CE (UCIAMS-282870, 1855±15 BP, >30kDa collagen yield (%) 3.5, d15N (‰) 11.1. d13C (‰) -20.0, C/N (atomic) 3.1).

CGG_2_107023 are disarticulated remains above the primary burials have been dated to 599–652 CE (UCIAMS-282668, 1430±20 BP,) >30kDa collagen yield (%) 10.2, d15N (‰) 14.5.

d13C (‰) -18.1, C/N (atomic) 3.2. Museum no. T19564 l, m, o Det kgl. norske videnskabers selskab. Museet. Antikvarisk avdelings tilvekst 1975, museum no. T19564.

#### Norway, Northern Norway, Troms, Balsnes (Ts 874-882/A4274)

Coordinates: 69.5407, 18.8530

Zanette Tsigaridas Glørstad

A burial was found "under stone and a thin layer of soil during ploughing" in 1889, in a gently inclined slope towards the Balsfjord in the NE, around 10 masl. A skeleton of an adult male was found together with a number of iron weapons and tools, including remains of a sword, two spearheads, forked arrowheads, horns and a sickle (Askeladden ID 57007). The finds can be dated no closer than to the Late Iron Age, i.e. the Merovingian Period (ca AD 550–800) – the Viking Age (AD 800–1050)[474,476].

CGG_2_107012 (Ts 874-882/ A4274, tooth, M2).

#### Norway, Northern Norway, Troms, Karlsøy, Engholmen 1 and 2, (Ts 299-318/ A2806 and Ts 715-722/A4275)

Coordinates: 70.1173, 19.0177

Zanette Tsigaridas Glørstad

Archaeological remains from Engholmen are derived from the eroded Iron Age cemetery at the southern end of Helgøya and the outlying skerries, consisting originally of up to 15 recorded burials, of which many are now eradicated (Askeladden ID 38503, ID 74787-1). Two burials, both with skeletal remains and grave goods, were collected from the site when it was taken into use as a churchyard in the late 19th century.

Engholmen 1 (Ts 299-318/ A2806) was found in 1877 in a low round mound of sand, possibly with remains of kerbstones. There were no signs of a burial chamber. The burial included a skeleton of an adult male[450], with a sword placed at the right hip, one bone comb by the head and another one by the feet, and with three bone harpoon heads next to the body. The grave goods also included a knife, a bone spoon, a whetstone and eight forked bone implements, the latter possibly for hunting fur animals[474]. The burial is dated to the 7th century CE.

Engholmen 2 (Ts 715-722/ A4275) was a pit-grave found in 1886, ca. 30 m. from Engholmen 1. The remains of an adult male[450], were found together with a spearhead, a bone comb, 8 iron arrowheads, a whetstone, and fragments of iron and bone implements of unknown use. The whetstone was placed under the deceased's head, the spearhead on the chest, and the comb by the head. It is dated to the 7th century CE.

Samples studied: CGG_2_107011 (Ts 299-318/ A2806, Engholmen 1, tooth M1). CGG_2_107013 (Ts 715-722/ A4275, Engholmen 2, tooth M3).

#### Norway, Northern Norway, Ytre Kvarøy (Ts 6657/A5119)

Coordinates: 66.4900, 12.9343

Zanette Tsigaridas Glørstad

The burial is part of a cemetery containing 13 burials excavated in 1968–1975 (Askeladden ID 63115). No discernible markers were found, and all graves are assumed to be pit-graves. An account of selected groups of jewellery are published in Vinsrygg 1979[477], while a majority of the burials are described in Klokkervold 2015[469], based on archival material. The cemetery is located on a cultivated shore terrace, ca 8 masl., overlooking the Kvarøy fjord. A layer of shell sand ca 20–30 cm below the surface contributed to good preservation conditions for osteological remains.

This grave (grave 13) was discovered during an inspection in 1974 and excavated the following year. The body was placed with the head in the northeast. Under the lower jaw was a conical brooch, and over the chest the remains of a necklace or similar ornament with 25 beads, a bronze spiral, a walrus tooth, as well as 5–6 cowrie shells. 3 beads were also found under the inner molars in the woman's mouth, possibly serving as a form of Charon’s coin. The grave also contained a spinning whorl, two knives, a comb and a vessel. The remains of the front section of a dog as well as cattle remains were found by the feet. The individual is a female, around 40 years old. Based on the associated jewellery, the grave is dated to the beginning of the 6th century.

CGG_2_107016 (Ts 6657/ A5119, tooth, M1).

#### Norway, Southern Norway, Buskerud, Tyrifjorden (A4177)

Coordinates: 60.0572, 10.2212

Zanette Tsigaridas Glørstad

The osteological remains were recovered in 1902 from Svartøyene, two small islets with a total of 8 registered burial mounds, largely damaged (Askeladden ID 9589-90,19424, 49009, 77186). The finds were collected by a Dr. Thaulow (as reported in a 1903 newspaper article), and transferred to the paleontological museum in Oslo in 1935. The remains include at least 7 crania, and additional skeletal remains. In 1894, a Bronze Age copper-alloy knife was recovered from an open cist in one of the cairns (Ringerike museum, RM05228). The burials can be dated to 1800 BC–200 AD[450].

CGG_2_107002 (box 1, tooth, M3)

CGG_2_107003 (box 4, tooth, M2), has been dated to 1955±20BP (UCIAMS-282666(10–130 CE) >30kDa collagen yield (%) 2.3, d15N (‰) 11.6. d13C (‰) -20.2, C/N (atomic) 3.1).

Another dating was obtained from an unspecified fragment for a previous study: 1980±55 BP, cal.35 BC–80 AD.

#### Norway, Southern Norway, Østfold, Kråkerøy

Coordinates: 59.1787, 10.9190

Lene Melheim

A male skeleton found c. 4 m deep in blue clay on an ancient seabed at Rødsmyr on Kråkerøy, Østfold, Fredrikstad, when a water reservoir was constructed in 1945. Cattle bones were found nearby (but not collected). The osteological analysis and the fact that the head was found at a distance from the rest of the body, led Per Holck[478] to argue that the man had been violently killed and dumped in the sea, hence possibly a sacrificed victim or a 'bog body'. During the Neolithic and Early Bronze Age Kråkerøy was an archipelago of small islands, with rich evidence of habitation.

The upper part of the head, lower jaw and most of the larger bones from the extremities are preserved (Schreiner collection anthr. no. 4482, lacks museum no.). Identified by Holck as a male individual, 172 cm tall, c. 60 years of age, and with signs of lethal violence, e.g. unhealed sword injuries to head and spine, stress, malnutrition, and scoliosis.

When teeth were sampled in 2017, Evangelia Daskalaki observed that the skeleton is well preserved, but not complete. The top of the cranium and a part of the left temporal bone are assembled with steel wire and glue. The lower jaw has 6 teeth attached (3 left and 3 right molars). There is also a substantial number of post-cranial parts.

The skeleton was dated to the transition EBA–LBA on the basis of a C14-date conducted by Holck[478]: 2860±90 BP. AMS dating initiated by Per Persson in 2016, gave an earlier date, with a tighter probability range: 3055±35 BP, or 1390–1265 cal. BCE (Rijksuniversiteit Groningen). 13C was measured at -20.58 and 15N at 10.68.

CGG_2_105645 (A4482, petrous bone).
CGG_2_105646 (A4482, tooth M1).

#### Norway, Southern Norway, Verket

Coordinates: 59.61397, 10.4185

Lene Melheim

Burial find from unconventional Late Neolithic gallery grave (stone cist), discovered in a sand quarry at Verket on the Svelvikryggen morraine, an isthmus in Drammensfjorden, Hurum, Akershus. There were two stone cists (Verket 1 and 2) lying c. 150 m apart at c. 20–21 m asl, close to the seashore and probably on a small island at the time of their construction[479].

Verket 1: Mus. no. C10476–C10480: 5 flint daggers of Lomborg's types I A/B (3), I C (1) and II A (1), human remains of 3–5 individuals (Schreiner collection anthr. no. 584, 584a, b, c). Verket 2: Mus no. C19886-87: Hagebyhöga axe, flint dagger type VA, human remains (unpreserved). Kulturminnesøk: Askeladden id 19898.

Verket 1 was an untypical stone-built chamber with multiple burials, situated in a southerly slope. The five 14.8–19 cm long flint daggers date to LN I. The human remains and the daggers were discovered c. 10 cm deep in fine sand, under a large boulder (4.3 x 3.7 x 2.5 m), when it was blown up. Dug into the ground beneath the natural boulder was a burial chamber of standing slabs, c. 2.2 x 1.6 m, oriented E-W. The southern end had an opening of c. 0.6 m.

One skull (584), postcranial remains originally identified as three individuals (584a, 584b and 584c). 584a is best preserved, 584b next best, 584c has the least number of bones. It is not clear whether the skull belongs to individual a or c (not b). Bones from two other, less well-preserved individuals (a child and an older person) were identified among the bones in 584b. Details and pictures from anthropological study by Per Holck are published by Einar Østmo (2011:95–103).

584a–c belong to at least five individuals, of which three are identified as male, c. 20–25 years of age and quite robust and tall (173 ± 4 cm, 166 ± 4 cm, 167 ± 4 cm) and with signs of violence and/or disease. Muscular attachments on humerus and femur are well developed on all three. A flattening of the femur (platymeri) was identified on two. 584a has an unhealed cut to the heel, a healed injury to the collarbone, and signs of arthrosis on several limbs. 584b has signs of malformation; possibly related to tuberculosis.

A new evaluation of the human remains was done by Evangelia Daskalaki in 2017, when the teeth were sampled, observing as follows: 584 is a well-preserved cranium, missing the right temple and jaw. The teeth preserved in the upper jaw are the right side main molars, and on the left side the first premolar and the first two molars (6 in total). The enamel shows little wear and there are no signs of caries. The skull most likely belongs to 584a,or possibly 584c, but not 584b since the articulated surfaces of the skull and the neck vertebrae do not match.

A conventional radiocarbon date was done by Rolf Scheen[457] on specimen 584a (3600±50 BP). The three male individuals were later AMS radiocarbon dated by Østmo (2011:92-93): 584a: 3630±40 BP, 584b: 3390±35 BP, 584c: 3610±35 BP. C14-dates are partly overlapping with but also somehow later than the typological date, indicating, according to Østmo, that the chamber had been used for multiple burials for a longer duration of time, c. 2100–1600 BC.

CGG_2_105650 (A584/C10476, tooth M3).

#### Norway, Southern Norway, Vestfold, Jarlsberg Herregaard (A1500/C20300/a-i)

Coordinates: 59.2908, 10.3867

Zanette Tsigaridas Glørstad

The burial was found under a low mound in 1901, when rocks were removed for reuse. The burial consists of a male adult skeleton, found with a spearhead, a shield boss, a gold fingering, remains of mountings for two drinking horn, a ceramic vessel and a large amount of shell fragments. Estimated dating is ca. the Roman Iron Age, ca AD 0–400[480]. The precise location of the mound is unknown, but it belonged to a small group of two-three mounds, possibly situated 150m SE of the farmhouse on the slope down towards the Aulie River (Askeladden ID22233). These mounds have now been removed by levelling and cultivation.

CGG_2_107001 (A1500/ C20300/a-i, tooth, M3).

#### Norway, Western Norway, Rjoa, Sola, Rogaland (S4115)

Coordinates: 58.9002, 5.6522

Håkon Reiersen & Sean Dexter Denham

The burial was found in a chamber in a damaged mound in 1920 (Askeladden ID 5537) at the farmstead Joa/Rjoa in the southern end of the Hafrsfjord. The burial is dated by its inventory of jewellery and ceramic vessels to the latest part of the Roman period (AD 325–400). As the jewellery composition is typical for female burials at the time, it is assumed to be a female burial. Several splendid archaeological finds indicate that elites dwelled at this farmstead at the time of the burial. Notable finds include a lost serpent-head finger ring from the late Roman period and a large gold hoard from the Migration period[481].

CGG_2_107044 (S4115, petrous bone).

#### Norway, Western Norway, Skeie, Klepp, Rogaland (S8510 Ia, S8616/A4766)

Coordinates: 58.6939, 5.5472

Håkon Reiersen & Sean Dexter Denham

The two burials stem from the northern and southern part (Askeladden ID 5335 and 65582) of a burial site situated in the beach zone facing the North Sea. They form a part of a larger stretch of ‘beach burial sites’ along the coast of the Jæren region[482,483]. The burials were all inhumations, and due to shell sand covering the burials, preservation conditions for skeletal remains were favourable. The burials were situated in chambers in stone cairns of either circular or rectangular shape. In total, 7 burials were excavated by O. Møllerop in 1958–60[484]. The burials are all dated by context to the latest part of the Roman period and the early Migration period (AD 300–500). A few other burials excavated in the late 19th and early 20th century stem from the same periods. The area provides one of few favourable natural ports along the Jæren coast, and is thus strategically situated in relation to maritime traffic. A small gold hoard from the same period indicates a farmstead of some importance, and it is situated between several centres of power from the period[481].

The skeleton from burial S8510 Ia has been osteologically determined to be a man of about 60 years old. It was one of three burials in a large rectangular cairn. Originally, the cairn was made for a woman c. 20 years old, buried in a rectangular chamber (grave Ib). Later, a second woman c. 20–30 years old was buried in another, similar chamber (grave II). In southwestern Norway, the main rule for excavated rectangular cairns is that they represent female burials from the Migration period[485]. However, at some point after female burial Ib, a male burial secondarily was put into the chamber. The remains of the woman Ib and her associated grave goods were pushed to the edges of the chamber, before the corpse of the man was put down. Due to the atypical placement of a man in a rectangular cairn, it is likely that the man and woman were in some way related. While few objects stem from the male burial, it is dated in relation to the female burial. The female burial Ia is dated by context to c. AD 400, and the male burial Ib presumably should be dated to the early Migration period (AD 400–450). The individuals Ia and Ib are on permanent display in the exhibitions at the Museum of Archaeology, University of Stavanger[486].

The skeleton from burial S8616 (Skeie, Northern field, Grave IV) has been osteologically determined to probably be a man, 30–35 years old. As the burial was situated in a circular cairn and the burial included few objects, contextually it is likely that the deceased was indeed male. Female burials in the burial field on the other hand were placed in rectangular cairns with extensive grave goods. The man's preserved grave goods was only a handled vessel, dating the burial to the Migration period (AD 400–550). After excavations, the skeletal remains were deposited in the Schreiner Collections, Oslo. The collection protocol claims that the skeletal remains from S8616 got the new id number A7087. However, comparisons with field photos suggests that there has been a mix up with another id number from Skeie, A4766, which is actually the correct assemblage for S8616.

CGG_2_107042 (S8510 Ia, tooth, M2), CGG_2_107043 (S8616/A4766, tooth, M3).

#### Norway, Western Norway, Sogn and Fjordane, Skongeneshulen

Coordinates: 62.0302, 5.1266

Zanette Tsigaridas Glørstad

The osteological remains were recovered in 1886 by the geologist Dr. Hans H. Reusch who carried out an excavation within the cave[487,488]. The cave is situated around 100 m from the shoreline and ca. 20 m above the sea, at the northern peak of Vågsøy, a small island at the outer west coast. The cave is spacious, about 20 m wide and 30 m high in the entrance.

Remains of at least six unburned individuals were found in a gravel layer, beneath ca. 0,7– 0.8 m deep layer of manure from sheep who used the cave for shelter. These are assumed to represent primary burials (Bergsvik 2007). A shard from a bucket shaped pot, typologically dating to around 500 AD, was found in close association with the bones (mus. No. C13178). Slightly deeper were two layers of kitchen midden, 5–10 cm. thick and with a mutual distance of 20–30 cm. They consisted of shell (patellas and a few specimens of buccinum and blue shell), coal, splinters, ash, rounded stones, bones of ruminant animals, fish and probably also of humans. A pointed bone tool was found in the deepest midden layer (mus. No. C13179).

It has previously been assumed that the human remains were contemporaneous with the pottery remains, dating them to around 500 AD, the Migration Period. The C14 analysis for the current study suggests the human remains are older, and the pottery reflect a later use of the cave.

CGG_2_107004 (Box 1, Ind 1, tooth M3).

CGG_2_107005 (Box 1, Ind 2, tooth, M2).

CGG_2_107006 (Box 2, Ind B, tooth P3).

CGG_2_107007 (Box 2, Ind A, tooth P3).

CGG_2_107008 (Box 2, Ind C, tooth P3).

CGG_2_107009 (Box 2, Ind D, tooth, C).

CGG_2_107004 (Box 1, Ind 1/ A1570) has been dated to 1955±20 BP (UCIAMS-282667, >30kDa collagen yield (%) 2.2, d15N (‰) 15.0. d13C (‰) -17.5, C/N (atomic) 3.1).

#### Norway, Western Norway, Vik, Klepp (S14383)

Coordinates: 58.7121, 5.5258

Håkon Reiersen & Sean Dexter Denham

The farmstead Vik is situated immediately north of the farmstead Skeie, with two individuals included in this study (see Site description). Similar to the situation at Skeie, there are several 'beach burial sites' here. In 1936, a collection of human bones from Vik was sent to Bergen for osteological analysis (Osteological ID while the bone collection was in Bergen: JS 249). Although it has not been possible to retrieve the original context information for these bones, the provenance to Vik is regarded as certain. Well-preserved unburnt bones are rare in the Rogaland region, and as the local shell sand preserved the bones of the neighbouring farmstead Skeie, similar preservation conditions are highly likely also at Vik. It is therefore assumed that the bones stem from one or several of the beach burial sites at Vik.

As part of a research project[489], an osteological analysis of the bone assemblage was carried out by S. D. Denham, identifying 5–8 individuals among the bones. Radiocarbon dates of five femurs from different individuals provided a rather uniform date range with median dates within the Merovingian period (AD 600–800), although a slightly later date is possible for two individuals (lab no. UBA-36742-46). Petrus bones from three individuals were chosen for sampling. A conical brooch from the same period has been found at Vik by metal detecting, indicating graves disturbed by ploughing. Burials from the Migration period are also known from the Vik burial fields. The Vik bone assemblage is especially interesting viewed in relation to the older Migration period burials from the neighbouring farmstead Skeie.

CGG_2_107039 (S14383.1, petrous bone).

CGG_2_107040 (S14383.2, petrous bone).

CGG_2_107041 (S14383.3, petrous bone).

### Sweden

#### Sweden, Öland, Bengtstorp,Lilla Smedby, Mörbylånga, Sörby Störlinge

Coordinates:

Bengtstorp: 56.51, 16.43

Lilla Smedby: 56.41,16.43

Mörbylånga: 56.50, 16.42

Sörby Störlinge: 56.80, 16.73

Helene Wilhelmson

Early to Late Roman Burials.

The samples were selected from a much larger number of burials (circa 20) but many did not yield enough material for DNA-analysis as only tooth roots were sampled. The human remains are curated at the Historical museum, Tumba, under the accession id per site as given in table 1. All samples are from teeth, details specified in Table S7.2.27.

| **SAMPLE ID** | **ID*** | **SITE** | **ACCESSION SITE ID** | **GRAVE** | **EXCAVATION YEAR** | **PERIOD** | **DATE** | **SAMPLED TOOTH** |
|---|---|---|---|---|---|---|---|---|
| **CGG_2_024144** | Oland 1099 | S Bårby (Mörbylånga) | 1785/67 | 5 | 1966–1967 | Late Roman Iron Age | 200–400 CE | FDI 35 |
| **CGG_2_024145** | Oland 1038 | Sörby Störlinge | 28364 | 6 | 1964–66 | Early Roman Iron Age | 0–200 CE | FDI 23 |

| | | | | | | | | |
|---|---|---|---|---|---|---|---|---|
| | | (Gärdslösa) | | | | | | |
| **CGG_2_024146** | Oland 1036 | Lilla Smedby (Smedby) | 23267 | 1 | 1944 | Early Roman Iron Age | 21 BCE–236 CE (1905+-45 uncal.BP) | FDI 42 |
| **CGG_2_024147** | Oland 1092 | S Bårby (Mörbylånga) | 1214243 | 9, upper | 1904 | Early Roman Iron Age | 0–200 CE | FDI 43 |
| **VK522** | Oland 1052 | N Kvinneby (Stenåsa) | 24846 | 2 (lower burial) | 1953 | Late Roman Iron Age | 386+–80 (2325 +/-45 BP) | FDI 13 |

**Table S7.2.27:** List of the samples studied from Sweden, Öland, Bengtstorp,Lilla Smedby, Mörbylånga, Sörby Störlinge, their site and dating.

The individuals sampled in this study have previously undergone a detailed bioarchaeological analysis including radiocarbon dating and isotope analysis for migration and diet[490–492]. The results are summarised in Table S7.2.28.

The sampled individuals are both males and females of varying age of death. They comprise both individuals presenting a local strontium and oxygen profile and clearly non-local individuals. During the Early Iron Age (500 BCE–400 CE) the rate of non native born individuals on Öland was 30%, compared to 68% in the Late Iron Age (400–1050 CE)[492]. The average of the $\delta^{13}C$ values is-19.8 and $\delta^{15}N$ 13.7 for the Roman Iron Age (ie. 0–400 CE)[490]. The sampled individuals had a diet corresponding to the average person of the period with the exception of 1052. This individual had comparably lower values i.e. a slightly more terrestrial diet with less protein.

The two non-local individuals (id 1038, 1036) both have a very high strontium value (above 0.73) and comparable to baseline data available from for example mid-northern Sweden and Finland[491].

| **SAMPLE ID** | **ID*** | **AGE, YEAR (Y)** | **SEX** | **D13 C /D15N** | **SR /O** | **WILHELMSON & PRICE 2017** | **STATURE** | **NOTE** |
|---|---|---|---|---|---|---|---|---|
| **024144/ VK579** | Oland 1099 | 18–19 y | Female | - 19.8 / 13.7 | 0.7131 / -6.1 | local | 154.0 +- 4,52 | Very short stature, outlier |
| **024145** | Oland 1038 | Mature, ca 40 y | Female | -19.9/ 13.2 | 0.7358 /-5.3 | non-local | NA | |
| **024146** | Oland 1036 | 20-30 y | Female? | -19.2 /13.9 | 0.7314 /-6.6 | non-local | NA | |
| **024147** | Oland 1092 | Mature-old, 40+ | Male | -19.5/ 14.6 | 0.7158 /-5.8 | local | NA | Perimortem SFT and BFT; skull. |
| **VK522** | Oland 1052 | 19 y | Male | -20.3 / 11.2 | 0.7125 / -6.0 | local | 163.2 | Short stature for a male |

**Table S7.2.28:** Summary of bioarchaeological data for the samples from Table S7.28: List of the samples studied from Sweden, Öland, Bengtstorp,Lilla Smedby, Mörbylånga, Sörby Störlinge and observations per sampled individual. NA=not available[490,492,493].

The individuals sampled in this study were all found buried in lime stone cist burials, oriented north-south. This is the most common inhumation burial orientation and a common grave architecture during the Early Iron Age in Öland (Wilhelmson 2017a). Cist burials are often reused and comprise more than just a single human individual. This was the case of burial 1092, which was the upper burial in a cist with two complete individuals layered

#### Sweden, Southern Sweden, Scania, Höja gård, Västra Skrävlinge 3

Coordinates: 55.58 3.06

Pre Roman or Early Roman Iron Age (150 BCE to CE 75)

This barrow was situated in SW Scania, in the eastern outskirts of Malmö. It was excavated in 1966 by Claes Wahlöö, Malmö Museum (Wahlöö 1966, Malmö Museum inv. MHM 1264). The excavation revealed the remains of a substantial round barrow, estimated to *c*. 30 m in diameter and at least 1.75 m high, although the original height would most likely have been larger.

In and below the mound, a sequence of 16 inhumation burials and three cremations were found, dating to the Late Neolithic, Early Bronze Age, and the Early Iron Age. It was also clear that the mound had been constructed in several phases[278].

In the first phase, four inhumation graves dating to the Late Neolithic were constructed, only covered by small low mounds (A1, A17–A19). The grave goods were sparse but flint daggers, amber beads and schist pendants. The skeletons were poorly preserved but the positions seem to be extended supine. Probably, the traces of ard ploughing found below the mound belong to this phase.

In the Early Bronze Age, period II, an inhumation containing a *c*. 2m tall male (A8) was constructed and covered by an oblong mound, probably *c*. 20x8 m large. The male was accompanied by a period II bronze dagger and a flint arrowhead. A further 6 burials were inserted into this mound in the EBA period II–III (A2, A15, A20–A23). Of these, A15 was a cremation.

An enlargement of the mound took place in the EBA period III, when it was extended to 20 m diameter. In the centre of this mound and on top of A8 was an inhumation grave (A2) with period III artefacts (bronze tweezers, flint strike-a-light). The skeleton was poorly preserved. Another inhumation (A10) was inserted into this mound. It contained a bronze tutulus dating to period III. A further enlargement of the mound to ca 30 m diameter took place after this, but the dating is unclear.

In the peripheral parts of the mound, two inhumations from the Early Iron Age were found, A3 and A16. A3 contained an iron sickle and an iron needle, but A16 was findless. The skeleton was lying in outstretched supine position, but no construction was visible.

Due to poor preservation of most of the skeletons, only grave A16 yielded sufficient DNA for analysis. It was dated to

RISE219 (MHM 1264, A23) has been dated to 3101±27 uncal BP (OxA-29661).

RISE220 (MHM 1264) has been dated to 3251±30 uncal BP (OxA-29738).

CGG_2_015491 (MHM 1264, NEO50).

CGG_2_015492 (MHM 6902, NEO046) has been dated to 2030±33 uncal BP (UBA-39936, δ13C -20.3, δ15N 10.4). This corresponds to 150 cal BC–cal AD 75 (95.4%); however, 93.8% of the probability is confined to the range 115 cal BC–cal AD 75.

#### Sweden, Southern Sweden, Scania, Maglarp parish, Albäcksbacken

Coordinates: 55.3777, 13.0950

Torbjörn Ahlström

The site Albäcksbacken (Maglarp 13) is situated west of the town Trelleborg in southwestern Scania, Sweden. A gravel ridge, oriented in an NNE-SSW direction, has been excavated intermittently between the years 1934 to 1977[494–497]. Several cremations and inhumations were unearthed, dating from the Late Neolithic to the Iron Age. An osteological analysis of the Iron Age interments was published by Arcini[498]. For the present analysis, we have sampled specimens from the 1977 excavation[497], curated at the Lund University Historical Museum. Grave A13a 1 & 2 is a double grave, dated to the Late Neolithic, with two interments lying on the side. This grave was disturbed when another grave was dug into A13a, referred to as A13b (CGG_2_105923). This disturbance caused an uncertainty as to which burial the artefacts belonged. Therefore, A13b was classified as possibly Late Neolithic. The present analysis has confirmed genetic association with A13a, thus rendering the question mark obsolete. Grave A26 (CGG_2_105941), identified as belonging to an adult female, was furnished with an amber button and a bronze ring, and initially dated to the Roman Iron Age. The present analysis demonstrates a genetic affinity with the Late Neolithic individuals sampled, thus changing the date of the grave to the Late Neolithic. Apart from the four Late Neolithic individuals sampled, we have sampled eleven Iron Age individuals, dated to the Pre-Roman Iron Age and Roman Iron Age (Table S7.22.9). Four graves have dateable finds, seven lack finds altogether. Arcini[498] provided three radiocarbon dates for Iron Age interments, where the date from grave

A15b is relevant here. Tornberg[499] provides radiocarbon dates for the two individuals in A13a. The list below is compiled from Jacobsson[497], Arcini [498] and Tornberg[499].

| CGG No | Individual id | Date | information |
|---|---|---|---|
| CGG_2_105922 | A13a Indiv 2 | LN, UBA-24005, 3604±30 uncal BP, d13C-20.6, d15N 10, C:N ratio 3.29 | An inhumation grave without any markings containing two adult females with the heads in different ends of the grave. They were placed on the side and were interred with a re-cut flint dagger and a flint flake. |
| CGG_2_105921 & CGG_2_105920 | A13a Indiv 1 | LN, UBA-24004, 3588±30 uncal BP, d13C-21, d15N 9.9, C:N ratio 3.23 | See above |
| CGG_2_105928 | A15b | Pre-Roman Iron Age–Early Roman Iron Age U-8155 2000±80 uncal BP 200 BCE–220 CE | Inhumation grave 2.3x0.72x0.22m, E-W orientation. A male placed in hocker position , aged 30–35, no artefacts in the burial. |
| CGG_2_105926 | A15a | Pre-Roman Iron Age–Early Roman Iron Age On stratigraphic evidence older than A15b | Inhumation grave, damaged, orientation E-W. Male placed in hocker position, no artefacts in the grave |
| CGG_2_105925 | A14 | Early Roman Iron Age | Inhumation grave 2.4x0.62x0.3 m, orientated E-W. A male 35–40 years old placed outstretched on his back. A clay pot was found in the grave |
| CGG_2_105930 | A16 | Early Roman Iron Age, B | Inhumation grave 2.25x0.8x0.25m, orientation W-E. Male aged 20–30 placed in supine position with a spear head, an iron knife and a bone pin. |

| | | | |
|---|---|---|---|
| CGG_2_105932 & CGG_2_105933 | A23 | Roman Iron Age | Inhumation grave with a larger stone in the lower part of the grave, orientated W-E. A male aged 25–30 in supine position with Iron knife and one iron pin or awl |
| CGG_2_105923 | A13b | Late Neolithic | Burial above 13a, some of the bones from grave 13a were placed central in the grave, flint flake and recut flint dagger |
| CGG_2_105934 | A16 F18 | Early Roman Iron Age B | Inhumation grave, oriented W-E, a male aged 25–30 placed in a supine position with slightly flexed arms buried with one iron knife and one bone comb |
| CGG_2_105935 | A11 | Roman Iron Age | Inhumation grave 2x0.76x0.32 m, orientated WNW-ESE, A male 35+ placed outstretched on his back. No artefacts were found in the grave. |
| CGG_2_105936 & CGG_2_105937 | A24 | Probably Roman Iron Age | Inhumation grave orientated NNE-SSW, A male aged 25–30 placed in supine position, no objects found in the burial |
| CGG_2_105938 | A19 | Probably Roman Iron Age | Damaged inhumation in WSW-ENE orientation. A male in supine position with whetstone pendant |
| CGG_2_105939 & CGG_2_105940 | A25 | Probably Roman Iron Age | Inhumation grave orientated N-S, Male aged 50–60 placed in hocker position with a larger stone at the back, no objects in the grave |
| CGG_2_105941 | A26 | Late Neolithic | Inhumation in SSW-NNE orientation. A female aged 30–40 in supine position. interred with one amber button and a ring (finger?) of bronze |

| | | | |
|---|---|---|---|
| CGG_2_105942 & CGG_2_105943 | A27 | Later Roman Iron Age– Migration period | Inhumation in E-W ordination. Female aged 40–50 placed in supine position, interred with iron sickle and an iron fragment |

**Table S7.2.29:** List of samples studied from Sweden, Southern Sweden, Scania, Maglarp parish, Albäcksbacken, their isotopic analysis and context description.

#### Sweden, Western Sweden, Bohuslän, Assleröd

Coordinates: 58.4172, 11.3251

Karl-Göran Sjögren

Late Neolithic gallery grave, excavated in 1863 by August Malm[500] and again in 1920 by Georg Sarauw. The chamber was damaged due to shell gravel extraction and only two stones remained of the chamber, but the find bearing area measured ca 3 by 1.8 m. The grave was located on a late glacial shell bank that contributed to the preservation of human bone. Remains of at least eight people were found, among which a child. Two crania were measured and published by Gustaf Retzius[501] (no 41 and 42). The grave also contained 3–4 flint daggers, 7 flint flakes, some pottery, and animal bones from horse, pig and sheep. No 14C datings have been done, but Sr isotopes were published by Sjögren, Price & Ahlström[502]. An M3 tooth from a 35–45 years old male was sampled from this grave.

CGG_2_106210 (GAM 2216, 84).

#### Sweden, Western Sweden, Bohuslän, Frölundaborg

Coordinates: 57.6779, 11.9356

Karl-Göran Sjögren

Late Neolithic gallery grave, excavated and removed in 1930 by Nils Niklasson. It was constructed of seven gneiss slabs, oriented north-south and partly damaged. The remaining length was ca 1.5 m and the width was ca 1 m. No roof blocks were preserved. The chamber was surrounded by a low mound. The grave was located on a late glacial shell bank that contributed to the preservation of human bone. Remains of three individuals were found: a male ca 20 years old (ind A), a female 20–25 years old (ind B) and a child c. 6 years old (ind C). The finds in the chamber consisted of a decorated bone needle and three flint flakes. Sr isotopes were published by Sjögren, Price & Ahlström[502]. Inv no GAM 45859-45863. No 14C datings have been made.

CGG_2_106210 (GAM 2216, 84).

#### Sweden, Western Sweden, Falbygden, Landbogården

Coordinates: 58.177602, 13.434700

Torbjörn Ahlström

The passage grave at Landbogården in Gökhem parish was excavated in 1987 by Lars Bägerfeldt [503] [504]. Osteological material from the tomb was analyzed by Ahlström [505], light isotopes by Lidén [506], and strontium isotopes by Sjögren et al.[502]. This tomb is one of the smallest known in the area, featuring a rectangular chamber made of limestone slabs measuring only 2.7 x 1 m, with a passage approximately 4 m long extending from the eastern side.

Although no roof blocks remained and the northernmost stone in the western wall was missing, the rest of the chamber did not exhibit any obvious signs of damage. Some of the passage's side stones were missing, while others were lying down. The structure was surrounded by the remains of a cairn or mound about 7 by 5 m, though the original size is unknown due to damage from ploughing.

A total of 857 human and 84 animal bone specimens were retrieved. Among these, at least 344 fragments belong to the articulated individuals (B-E) and 87 fragments to the bone package (F), with 426 fragments coming from disarticulated bones. Individual A , a complete skeleton, was dated to the Iron Age. With respect to the neolithic bone material, the minimum number of individuals is estimated to 16 (5 children and 11 adults).

In total, 13 radiocarbon datings were conducted on human bone samples from Landbogården. Initial datings in the 80's conducted by means of conventional methods without pretreatment of samples suggested early neolithic dates. However, renewed datings with accelerator propose a concentration of dates around 3000 BC (i.e. early middle neolithic).

CGG_2_105989 (SHM32201, box 8212, Ind F2) has been dated to 2030±20 BP (UCIAMS-275224).

### S7.3. Site Descriptions for Western Eurasia (excluding Northern Europe)

The following sections contain the archaeological descriptions of the sites from Western Eurasia for the new samples in this study. Each site description is followed by the inventory number of the samples concerned (CGG inventory number (GeoGenetics inventory number),

Museum inventory number). When available, the $^{14}C$ dating, isotope data and the sex identification of the samples are given.

### Austria

#### Austria, Dürrnberg, Hallein

Coordinates: 47.66 13.09

Karin Wiltschke-Schrotta & Holger Wendling

Dürrnberg near Hallein, in the northern part of the Austrian Alps, is one of the most important Iron Age sites in Central Europe[507–511]. The excellent geographical location of the Dürrnberg mountain zone at the northern exit of the Salzach valley into the Salzburg basin combines trade routes to the Alpine foothills and the inner Alpine valleys and a vital mineral resource: salt!

This outstanding economic position forms the basis for the rapid economic growth soon after the initial occupation c. 600 BC. The archaeological record includes mining sites, settlements and almost 400 tombs with 950 documented burials of the late Hallstatt and La Tène periods[510,511]. The finds from the graves contain exceptional precious artefacts which indicate the prominent position as an economic centre. As such, the Dürrnberg node linked the Iron Age communities of Central Europe through the valleys and passes of the Inner Alps with the Mediterranean cultures of northern Italy, the south-eastern Alps and the Balkan Adriatic[512,513]. The graves contained a variety of southern imports, northern Baltic amber objects, and ornamentation and style from the central West-Hallstatt communities from both sides of the upper Rhine. These indicate the role of Dürrnberg as a transfer and relay station on one of the major eastern Alpine communication routes. The imports have been adapted in processes of cultural appropriation and integrated into local cultural traditions[510]. Against this background, the innovative role of Dürrnberg is evident. It underlines its transformative role in the emergence of the artistic and cultural innovation of the La Tène period[511,514].

Based on the physical human remains, anthropological research addresses the question of how the Iron Age mining community managed subsistence, workload and physical impact of mining and living in an alpine region[515]. Individual and communal health status is interpreted by collecting data on the distribution of diseases, and patterns of trauma. Demographic data shows a rather even distribution of male and female individuals in different age categories. A detailed analysis concerning the different cultural epochs is still pending until the last two major burial sites will have been investigated.

CGG_2_101225 (NHMW_HD_288-2_Zahn_46).

CGG_2_101232 (NHMW_HD_228-4_Zahn_36).

#### Austria, Hallstatt, Gmunden

Coordinates: 47.56, 13.64

Doris Pany-Kucera, Anton Kern(†) & Georg Tiefengraber

The first evidence of human presence in the Hallstatt High Valley, where the cemetery and the salt mines are situated, dates to the Neolithic, maybe already c. 7000 years ago. First traces of systematic salt mining can only be documented for the Bronze Age[516]. However, one of the world's oldest continuously used industrial and cultural landscapes developed around the

Hallstatt salt mines. Salt determined life in this region for a long time and is still being mined today.

The world-famous cemetery in the high valley above the Hallstatt lake dates to the Early Iron Age period in Europe (8th–4th centuries BC), and is name-giving for the "Hallstatt period". The graveyard contains inhumations and cremations, only a few of them were urn burials. A large number of graves was excavated in the 19th century by a team around Bergmeister Johann Georg Ramsauer, whose documentation of the Hallstatt burial site is exceptional for that time. Until 2023, in total c. 1500 of nearly 2000 excavated graves were unearthed documented, although only a small part of the skeletons is preserved[517]. As the new excavations since 1993 have shown, the original number of graves must have been considerably higher and comprised around 4000–5000 burials. The cemetery was in use until the early La Tène period and revealed outstanding and precious items from throughout Europe and outside, reporting the extensive network of contacts of the Hallstatt people, based on the salt trade.

Some aspects on the skeletons have been investigated in detail, like entheseal changes, chronic sinusitis, and potential child labour, and are partly published[518–520]. A complete presentation of the different burials and a detailed chronological seriation is still pending.

CGG_2_101213 (NHMW_KE45_M3_FD_18).

CGG_2_101214 (NHMW_KE54_P4_FD_25).

CGG_2_101215 (NHMW_KE46_P4_FD_15).

CGG_2_101216 (NHMW_KE55_P4_FD_25).

CGG_2_101217 (NHMW_Inv.A.850_M3_FD_48).

CGG_2_101219 (NHMW_Inv.A.918_P4_FD_15).

CGG_2_101222 (NHMW_Mo_8_1938_P4_FD_45).

CGG_2_101223 (NHMW_KE23_P4_FD_45).

CGG_2_101225 (NHMW_HD_288-2_Zahn_46).

CGG_2_101232 (NHMW_HD_228-4_Zahn_36).

#### Bulgaria

##### Bulgaria, Varna county, Koriyata (at Suvorovo)

Coordinates: 43.353, 27.587

The archaeological site of Koriyata is located near the town of Suvorovo, in the Varna region. The site was a Middle Copper Age settlement (4750–4600 BC), with later occupation during the Early Mediaeval period (10th century AD). During the excavations in 2011 a single skeleton of the end of Late Antiquity period (end of 6th–beginning of 7th c. AD) was cleared to the north of one of the Chalcolithic buildings[521].

CGG_2_018087 (Koriyata_(Varna)_-_from_Inga_Merkyte).

#### Crimea

##### Crimea, Suvlu-Khaye

Coordinates: 44.74, 33.88

Alexei Voloshinov, Vyacheslav Masyakin

The Suvlu-Khaya necropolis is located in the southwestern part of the Crimean peninsula, in the eastern part of Bakhchisarai. 46 burials dated by the end of 3 –first part of 5 centuries AD were excavated by A.A.Voloshinov and V.V. Masyakin in 2009–2019[522,523] .

The samples for genetic studies were taken from the burials attributed to different chronologic periods.

The first group of burials dated to the 3rd century AD corresponds to the final stage of the Late Scythian archaeological culture. Archaeological data reveals significant Sarmatian cultural influence in Crimea in this period. The burials were made in the undercut graves. In most cases, the buried lay with their heads to the east.

CGG_2_017683 is a male of 35–45 years old. His skull was artificially deformed which is very characteristic for the Sarmatian society. On the skull in the region of the right frontal tubercle there is a rhomboid opening from an intravital penetrating injury with traces of healing[522]. The skeleton lay in a wooden coffin and accompanied by red-lacquered vessels, a fragment of a Roman bronze spoon, an animal bone and an iron knife. Dating - the first half of the 3rd century AD. Burials of the second cultural-chronological group were found in catacombs, which were used for multiple burials. The catacombs are oriented from west to east and southwest to northeast. In those cases when it was possible to fix the bones in situ, the buried were placed with their heads toward the entrance of the chamber. Such a burial tradition is presumably associated with the Alans. The burial inventory is represented by numerous stucco ceramics and pottery red-lacquered vessels, amphoras, ornaments and costume details, and weapons. Burials in the catacombs date back to the 4th to first half of the 5th centuries AD. This group includes burials in catacombs 7 (burial 3, CGG_2_017681) and 32 (CGG_2_017682).

It was suggested that together with obvious Alanian influence, the burial tradition of the second period shares some characteristics of Chernyakhov, Przeworsk, and Wielbark cultures which are related to the ancient Germanic tribes[523]. According to historical evidence, the so-called Dory State with a multicultural population was established by the Eastern Goth tribes in Crimea in the 3–5 centuries AD.

CGG_2_017681 (N1).
CGG_2_017682 (N2).
CGG_2_017683 (N3).

#### Crimea, Chersonesus City

Coordinates: 44.36, 33.29

Vladimir Kolosov

The necropolis is located on the Black sea shore in the northern part of the Greek colony settlement of Chersonesus in the outskirts of present-day Sevastopol on the Crimean Peninsula. The necropolis was discovered as a result of archaeological excavations by the archaeological team of the Chersonesus Museum under the leadership of D.G.Belov. The first two burials were discovered during excavations in 1935, and in the following 1936 91 more burials were discovered. In the course of the following years, a number of burials were also discovered in this sector of the settlement. As a result, more than 150 graves were identified and studied in total The most burials were found within the area bounded by two transverse streets of the ancient Cheronesus city - the eighth (VIII) from the east and the tenth (X) from the west. The northern border of the necropolis runs along the coastline, the southern border along the Chersonesos gully[524] (273). At present time the osteological materials are housed in the Peter the Great Museum of Anthropology and Ethnography (Kunstkamera) of the Russian Academy of Science (collection 5656).

The main bulk of the burials are inhumations made in the ground pits. The buried were stretched on their back (mainly with an eastern orientation) or crouched on the left or right side. A significant number of children's burials in amphorae were also excavated. The inventory is rather poor, but it allows the burial to be dated to the early stage of the Greek colonization of the Black sea territories within the 5th–4th centuries BC[524] ( 275–278).

The necropolis dates back to the early stages of the history of Chersonesus, which should be characterized by a high level of social, population and ethnic diversity. It is sometimes argued that the crouched burials are associated with the autochthonous Taurian population. An alternative assumption is that such burials reflect the social position of the buried (Ivanov, 2013, 13–17; Stoyanov 2004)[525,526].

All samples taken for the genetic study are from the burials made in simple graves pits without any inventory.

CGG_2_021478 (5656-21) is an adult laying stretched off the back with its head oriented to the East[527].

CGG_2_021475 (5656-24) is a poorly preserved skeleton of an adult destroyed by the later burials. Possibly laying crouched on the right side with its head to the east[527].

CGG_2_021474 (5656-259 is a child crouched on its left side with its head to the south-west[527].

CGG_2_021473 (5656-34) is an adult stretched on its back with its head to the east. The right hand lay on the pelvic bones, the left was bent upward at the elbow joint. Traces of ash and coal were found in the burial [527].

CGG_2_021476 (5656-35) is an adult lay stretched on its back, head to the east. The right arm was bent and lay on the stomach, the left - on the chest. Traces of ash and coal were found in the burial[527].

CGG_2_021472 (5656-45) is an adult stretched on its back with its head to the East-South-East[527].

### Czech Republic

#### Czech Republic, Central Bohemian Region, Holubice

Coordinates: 49.17, 16.81

Migration Period, Langobard

Holubice is located in the Vyškov district, 15 km east of Brno[528]. M. Čižmář. K. Gaislerová and I. Rakovský conducted excavations here in 1979–80. In 105 graves, 86 individuals were anthropologically determined: 22 men, 32 women and 34 children. According to M. Stloukal, the burial ground could last about 50–60 years and was used by a 40–60 member group. From a chronological point of view, J. Tejral included it in the Langobardic period (Middle Danube phase 4: 510 / 20–540 / 50) (2005 tab.3).

According to the cemetery plan, the graves appear to be divided into three groups, which could lead to the possibility of identifying ancestral structures in Langobar society[529]**.**

CGG_2_021948 (Holubice_H93).

CGG_2_021956 (Holubice_H32).

CGG_2_021963 (Holubice_H88).

CGG_2_021972 (Holubice_H41).

#### Czech Republic, South-Moravia Region, Lužice

Coordinates: 48.83, 17.06

Migration Period, Langobard

Lužice at Hodonín is the largest Langobardic burial ground from the 6th century in Moravia found so far, counting 120 skeletons[530,531]. (Head of archaeological research was Z. Klanica 1981– 1990). Average height of the bodies was 177 cm in males and 163 cm in females.

The osteological material from Lužice presents strongly pronounced relief of musculature attachments. Excessive loading of the respective groups of muscles stressed the bone edges on the fringe of the hip bone as well as those on the bones of the lower and upper extremities. It seems that this population was hard working.

CGG_2_021981 (Luzice_H110).

CGG_2_021991 (Luzice_H75).

CGG_2_021993 (Luzice_H50).

#### Czech Republic, Ústí Region, Radovesice II

Iron Age, La Tène

Coordinates: 50.40, 14.06

Václav Smrčka

The La Tène burial ground Radovesice II is located in the Teplice district near Razice. In 1974–77 J. Waldhauser[532,533] examined 37 skeletal and cremation graves. Some individuals were stored in coffins made of wooden logs.

CGG_2_021735 (Radovesice_II_Hv_7_16345).

CGG_2_021736 (Radovesice_II_Hv._19_16364).

CGG_2_021737 (Radovesice_II_Hv_15_16312).

CGG_2_021738 (Radovesice_II_Hv_20_16365).

CGG_2_021740 (Radovesice_II_Hv_11/81_16309).

CGG_2_021839 (Radovesice_II_Hr_5_(16373).

CGG_2_021846 (Radovesice_II_Hr_16_(17332).

#### Czech Republic, Ústí Region, Lovosice

Coordinates: 50.5099972, 14.05

Václav Smrčka, Marek Půlpán

The site was discovered by Marek Půlpán in 2002 and belongs to the Hallstat Bylany culture. The findings reflect a community engaged in agriculture, metalworking, and trade, with a complex social organization and rich cultural practices. These insights help paint a broader picture of the development and interactions of ancient European societies during this period. Grave 28A contains two burials A and B which contained respectively a cremated individual and a female skeleton of 24-35 years old. Artefacts such as a wire torc, rion pin with bronze patina and several fragmented poteries were found associated with the individuals in the burials.

CGG_2_021890 (Lovosice_II_obj_1_/_2002).

### Finland

#### Finland, Isokyrö, Levänluhta

Coordinates: 62.94, 22.41

Anna Wessman and Kristiina Mannermaa

Migration period-Merovingian period (400–700 CE)

The Levänluhta find material consist of commingled human remains from 98 individuals (MNI) buried along with artefacts and animal bones[534]. During the Iron Age this spring site was a pond or a small lake, and it has been archaeologically investigated since the 1800s[535,536]. Excavations have yielded an impressive range of finds, including precious copper alloy brooches, arm rings and other dress implements together with an imported copper alloy cauldron (Vestland type), suggesting that most of the buried individuals were women[535]. Osteological studies have confirmed that Levänluhta was a cemetery mainly for women and children[537,538], which is rather uncommon for Finland. Levänluhta and Käldamäki are unique sites in Finland.

The genomic data from Levänluhta was obtained from four mandibular molars which have the museum numbers KM 2:1:a2 (right lower first premolar+left lower third 44 molar), 2:1:a3 (right lower third molar) 2:1:a16 (left lower first molar) and 2:1:a29 (right lower first and third molar). Three of the samples (coded: DA234, DA238 and DA236) have been previously published[539]. All individuals are females based on the aDNA.

CGG_2_019338 (Sr2_Levanluhta_(48)).

#### Finland, Vöyri, Käldamäki

Coordinates: 63.15, 22.17

Anna Wessman and Kristiina Mannermaa

Migration period-Merovingian period (420–620 CE).

Found by accident in 1935 in connection to draining work, next to a large erratic stone boulder. Small-scale excavations took place there in 1936 and 1937. Six commingled individuals have been excavated together with cow bones, a wooden axe shaft and a bronze rod. The individuals are believed to have been deposited in shallow water[535]. Previous radiocarbon dates from 1988 (Ua 991: 1500±85 and Ua 992:1550±80 BP) suggest that the site is from the end of the Migration period to early Merovingian period[534].

CGG_2_024143 (Käldamäki_1) is a female individual.

### France

#### France, Aisne Haut de France, Bucy le Long

Coordinates: 49.39, 3.39

Jean-Paul Demoule

The Celtic cemetery of Bucy-le-Long (Aisne department, France) was excavated as a rescue excavation during the 1980–1990’. It provided about 200 well-preserved La Tène A–B graves (480–320 BC), with 4 aristocratic chariot burials, 6 main typological phases, that is probably about six generations. The cemetery is organised in several groups of graves, probably family groups. It is one of the best studied and published Celtic cemeteries all over France[540–542].

About 50 samples (petrous bones) were selected in 2018 by Fabrice Demeter and Jean-Paul Demoule; 38 have enough preserved aDNA; 19 of 50 are currently above 1X and will therefore be used in the more advanced analysis. About 18 C14 dates were made, mainly coherent with the relative typochronology (Table S7.2.30).

| **CGG No** | **N° grave** | **site** | **$^{14}$C cal BC** | **Typological phase** | **Groups of graves** |
|---|---|---|---|---|---|
| CGG 2 022418 | BLH 55 02/03/18 | Bucy le Long | 480–320 BC | IIA | 2 |
| CGG 2 022420 | BLH 213 02/03/18 | Bucy le Long | **485** | IIA | 4 |
| CGG 2 022421 | BLH 224 02/03/18 | Bucy le Long | 435 | IIIB–C | BFT |
| CGG 2 022424 | BLH 368 02/03/18 | Bucy le Long | *245* | IIIB–C | NW |
| CGG 2 022425 | BLH 17 02/03/18 | Bucy le Long | **473** | IIA | 2 |
| CGG 2 022426 | BLH 37 02/03/18 | Bucy le Long | 480–320 BC | IIA | 2 |
| CGG 2 022427 | BLH 38 02/03/18 | Bucy le Long | 480–320 BC | IIA | 2 |
| CGG 2 022428 | BLH 48 02/03/18 | Bucy le Long | 480–320 BC | IIA | 2 |
| CGG 2 022429 | BLH 57 02/03/18 | Bucy le Long | 480–320 BC | undated | 2 |
| CGG 2 022430 | BLH 59 02/03/18 | Bucy le Long | **435** | IIA | 2 |
| CGG 2 022431 | BLH 63 02/03/18 | Bucy le Long | *452* | IIIA | 2 |
| CGG 2 022432 | BLH 64 02/03/18 | Bucy le Long | 480–320 BC | IIB | 3 |
| CGG 2 022433 | BLH 67 02/03/18 | Bucy le Long | **490** | IIB | 3 |
| CGG 2 022434 | BLH 69 02/03/18 | Bucy le Long | 480–320 BC | IIA–C | 3 |
| CGG 2 022436 | BLH 369 Good 02/03/18 | Bucy le Long | 480–320 BC | IIC | 2 |
| CGG 2 022437 | BLC 67 02/03/18 | Bucy le Long | 480–320 BC | IIB | « BLC » ? |
| CGG 2 022438 | BLH 54 02/03/18 | Bucy le Long | 480–320 BC | undated | 2 |

| | | | | | |
|---|---|---|---|---|---|
| CGG 2 022440 | BFT 202 02/03/18 | Bucy le Long | **420** | IIC | BFT Est |
| CGG 2 022441 | BFT 223 02/03/18 | Bucy le Long | **370** | IIIA | BFT Est |
| CGG 2 022442 | BFT 226 02/03/18 | Bucy le Long | 480–320 BC | IIC | BFT Est |
| CGG 2 022443 | BFT 228 02/03/18 | Bucy le Long | **320** | IIIB–C | BFT Est |
| CGG 2 022444 | BLH 5 02/03/18 | Bucy le Long | 480–320 BC | undated | 2 |
| CGG 2 022445 | BLH 16 02/03/18 | Bucy le Long | **465** | IIB | 2 |
| CGG 2 022447 | BLH 27 02/03/18 | Bucy le Long | 480–320 BC | II A–C | 2 |
| CGG 2 022451 | BLH 53 02/03/18 | Bucy le Long | 480–320 BC | IIA–C | 2 |
| CGG 2 022452 | BLH 56 02/03/18 | Bucy le Long | *407* | IIA | 2 |
| CGG 2 022453 | BLH 58 02/03/18 | Bucy le Long | **470** | IIA–C | 2 |
| CGG 2 022454 | BLH 60 02/03/18 | Bucy le Long | **378** | IIA | 2 |
| CGG 2 022455 | BLH 66 02/03/18 | Bucy le Long | **379** | IIC | 2 |
| CGG 2 022456 | BLH 70 02/03/18 | Bucy le Long | 480–320 BC | undated | 3 |
| CGG 2 022457 | BLH 75 02/03/18 | Bucy le Long | 480–320 BC | undated | 3 |
| CGG 2 022458 | BLH 79 02/03/18 | Bucy le Long | 480–320 BC | IIIA | 2 |
| CGG 2 022459 | BLH 82 02/03/18 | Bucy le Long | *491* | IIIA | 2 |
| CGG 2 022460 | BLH 114 02/03/18 | Bucy le Long | 480–320 BC | | 5 |
| CGG 2 022461 | BLH 150 02/03/18 | Bucy le Long | 480–320 BC | undated | 3 |
| CGG 2 022463 | BLH 188 02/03/18 | Bucy le Long | **511** | IIA | 4 |
| CGG 2 022464 | BLH 198-1? 02/03/18 | Bucy le Long | *314* | IIA | 2 |

| | | | | | |
|---|---|---|---|---|---|
| CGG 2 022465 | BLH 369 Nord? 02/03/18 | Bucy le Long | 480–320 BC | IIC | 2 |

**Table S7.2.30:** List of samples studied from France, Aisne Haut de France, Bucy le Long and their dating. Caption: $^{14}C$ column : **bold : ± compatible dates ;** *italic : incompatible dates.* Absolute dates BC = BP - 1950.

#### France, The Jura Culture, Bourgogne Franche Comté

Bruno Chaume

The Tumulus of Moidons/Parançot belong to what Patrice Brun defined as the Culture from the Jura. These necropolises were set up on the French Jura Plateau; They form a cultural group, homogeneous all along the First Iron Age (Hallstatt) on their artefact production as well as on their funerary practices. One of the important points would be to check if these populations were linked genetically. A few burials from the Early and Late First Iron Age should put some light on the genetic traits of these populations. The hypothesis of sedentary clan families present since the Bronze Age is suggested by archaeologists to explain the high number of tumulus within a small space. It will be very informative to study kinship within a group of tumulus sharing the same territory[543–546].

On the Langres Plateau, at Vix and in the Châtillonnais, at the North-West of Burgundy, populations were occupying these regions as early as the beginning of the First Iron Age with a well-established network. The site of Vix was at the centre of the economic activity and the commercial trade over long distances at the end of the 6th and beginning of the 5th century B.C. These relatively intensive contacts between autochthon populations and Mediterranean groups, seem to have led to genetic exchanges. It will also be very interesting to check whether genetic changes occurred within local populations and when they occurred. Then the question will be to answer if the entire population was concerned or if it was only the elite.

At Vix particularly, it will be very interesting to compare the genome of the Vix Princess with this of the other human remains excavated in the region and that yielded DNA.

CGG_2_023660 (Les_Moidons_Tumulus_1_120).

CGG_2_023662 (Les_Moidons_Tumulus_4_96).

CGG_2_023663 (Les_Moidons_Tumulus_8_1922_35).

CGG_2_023664 (Les_Moidons_Tumulus_1_1925_99).

CGG_2_023668 (Les_Moidons_Tumulus_1_1925_100).

CGG_2_023669 (Les_Moidons_Tumulus_4).

CGG_2_023671 (Les_Moidons_Tumulus_sous_la_Rochette_37).

CGG_2_023681 (Les_Moidons_Tumulus_3_1918_19).

CGG_2_023687 (Les_Moidons_Tumulus?_1923_45).

CGG_2_023688 (Les_Moidons_Tumulus_4_1920_116).

CGG_2_023691 (Les_Moidons_Tumulus_?_1921_115).

CGG_2_023692 (Les_Moidons_Tumulus_?_1924_82).

CGG_2_023693 (Les_Moidons_Tumulus_8_1920_51).

CGG_2_023695 (Les_Moidons_Tumulus_?_1924_78).

CGG_2_023708 (Les_Moidons_Tumulus_4_30).

CGG_2_023709 (Les_Moidons_Tumulus_4_128).

CGG_2_023710 (Les_Moidons_21).

CGG_2_023714 (Les_Moidons_88).

CGG_2_023716 (Les_Moidons_Tumulus_4_125).

CGG_2_023717 (Les_Moidons_Tumulus_4_1920_118).

CGG_2_023718 (Les_Moidons_Tumulus_7_1920_sep._b_11).

CGG_2_023724 (Les_Moidons_Tumulus_10_46).

#### France, Camp du Château

Coordinates: 46.94, 5.85

The Camp du Château (Salins, Jura) was a hilltop settlement with a summit plateau of less than 1 ha. It was defended by natural cliffs and ramparts, most of which have been dated to the 1st Iron Age. Sporadic traces of occupation have been attested in the late Bronze age, beginning of Hallstatt (Ha C), but the imprint of the end of the First Iron Age was the most striking. Numerous protohistoric necropolises in the area have been excavated since the 19th century, notably those at Les Moidons.

CGG_2_023655 (Camps_du_Château_1910_Corps_49).

CGG_2_023656 (Camp_du_Château_1910_Corps_5).

CGG_2_023657 (Camp_du_Château_1910_Corps_6_B).

CGG_2_023722 (Camp_du_Château_Sépulture_1922).

CGG_2_023725 (Camps_du_Château_Sépulture_2).

#### France, Champ Peupin

Coordinates: 46.89, 5.84

Tumulus of “Champ Peupin” (ø 23 m) has been excavated in 1869 and by M. Piroutet in 1921. In the grave D, unfortunately looted, fragments of an iron wheel tyre were evidence of the presence of a wagon. 2 fibulae date the grave of the late Iron age (530/500 BC).

CGG_2_023690 (Tumulus_4_28).

#### France, Necropolis Les Moidons and Parançot

Coordinates: 46.86, 5.85 and 46.87, 5.83

The necropolis of Les Moidons (groups of Les Moidons and Parançot) are located in the french part of the Jura. Around 90 tumuli with 234 graves identified, the Moidons group is one of the most important necropolis of the eastern part of France[547]. In the object association matrix of all the tombs, 4 groups were identified for a period extending from the Neolithic to the Second Iron Age (La Tène). The majority of the graves belongs to the 1st Iron Age (800–450 BC).

CGG_2_023660 (Les_Moidons_Tumulus_1_120).
CGG_2_023662 (Les_Moidons_Tumulus_4_96).
CGG_2_023663 (Les_Moidons_Tumulus_8_1922_35).
CGG_2_023664 (Les_Moidons_Tumulus_1_1925_99).
CGG_2_023668 (Les_Moidons_Tumulus_1_1925_100).
CGG_2_023669 (Les_Moidons_Tumulus_4).
CGG_2_023671 (Les_Moidons_Tumulus_sous_la_Rochette_37).
CGG_2_023681 (Les_Moidons_Tumulus_3_1918_19).
CGG_2_023687 (Les_Moidons_Tumulus?_1923_45).
CGG_2_023688 (Les_Moidons_Tumulus_4_1920_116).

4624 CGG_2_023691 (Les_Moidons_Tumulus_?_1921_115).

4625 CGG_2_023692 (Les_Moidons_Tumulus_?_1924_82).

4626 CGG_2_023693 (Les_Moidons_Tumulus_8_1920_51).

4627 CGG_2_023695 (Les_Moidons_Tumulus_?_1924_78).

4628 CGG_2_023703 (Les_Moidons_Tumulus_4__27).

4629 CGG_2_023708 (Les_Moidons_Tumulus_4_30).

4630 CGG_2_023709 (Les_Moidons_Tumulus_4_128).

4631 CGG_2_023710 (Les_Moidons_21).

4632 CGG_2_023714 (Les_Moidons_88).

4633 CGG_2_023716 (Les_Moidons_Tumulus_4_125).

4634 CGG_2_023717 (Les_Moidons_Tumulus_4_1920_118).

4635 CGG_2_023718 (Les_Moidons_Tumulus_7_1920_sep._b_11).

4636 CGG_2_023724 Les_Moidons_Tumulus_10_46).

4637 CGG_2_023659 (Parançot_Tumulus_5_près_du_corps).

4638 CGG_2_023667 (Parançot_17).

4639 CGG_2_023672 (Parançot_Tumulus_10_Sépulture_centrale_95).

4640 CGG_2_023673 (Parançot_Tumulus_13_52).

4641 CGG_2_023674 (Parançot_Tumulus_4_53).

4642 CGG_2_023676 (Parançot_Tumulus_1_29).

4643 CGG_2_023677 (Parançot_Tumulus_1_24).

4644 CGG_2_023678 (Parançot_Tumulus_7_6).

4645 CGG_2_023679 (Parançot_Tumulus_7_Corps_3_108).

4646 CGG_2_023682 (Parançot_Tumulus_9_10).

4647 CGG_2_023685 (Parançot_Tumulus_13_Vieillard).

4648 CGG_2_023686 (Parançot_Tumulus_2_1923_89).

CGG_2_023698 (Parançot_85).

CGG_2_023699 (Parançot_Tumulus_13_Vieillard).

CGG_2_023700 (Parançot_Tumulus_1_1913_75).

CGG_2_023702 (Parançot_Tumulus_1_122).

CGG_2_023704 (Parançot_Tumulus_9_8).

CGG_2_023706 (Parançot_Tumulus_6_124).

CGG_2_023707 (Parançot_Tumulus_4_123).

CGG_2_023712 (Parançot_Tumulus_10_110).

CGG_2_023713 (Parançot_Tumulus_2).

CGG_2_023719 (Parançot_Tumulus_1_1900_16).

CGG_2_023723 (Parançot_Tumulus_1_1928_70).

#### France, Leuglay, tumulus des Montagnottes

Coordinates: 47.80, 4.83

Stone tumulus of 10 m in diameter, central grave of a warrior with a large iron sword and an S-shaped ceramic urn. Ha C dating (circa 750 BC).

CGG_2_023658 (Corps_9).

#### France, Maisey-le-Duc, tumulus de la Tête de Maisey S8

Coordinates: 47.85, 4.67

Three burial mounds measuring between 14 and 20 metres in diameter were excavated in the Tête de Maisey area at the end of the 19th century. The excavations were not very methodical, and only 5 burials were found in the mound 3. A bronze torc, bronze bracelets and anklets, date these graves to the end of the First Iron Age (Ha D2–D3).

CGG_2_023647 (Tumulus_de_la_Tête_de_Maisey_S3,_92.586.0).
CGG_2_023648 (Maisey-le-Duc,_tumulus_de_la_Tête_de_Maisey_S8).

#### France, Nod-sur-Seine, tumulus du Puits de Nod

Coordinates: 47.78, 4.62

This stone burial mound was around 10 m in diameter and 1m high. It was excavated in 1965 by René Joffroy and his team. The perimeter of the burial mound was defined by a well-constructed stone crown. A single burial, facing south-east/north-west, with the head to the south-east, occupied the centre. The body bore a bronze banded bracelet on the left arm and 2 bracelets on the right arm, one in schist, the other in bronze with oval decoration. The 3rd bronze bracelet could not be located in the tomb.

CGG_2_023651 (Tumulus_du_Puits_de_Nod_n°_2,_crâne_1).

#### France, Pothières, tumulus de Crevan

Coordinates: 47.90, 4.49

Four burial mounds were excavated by J. Lagorgette in the 1930s on the edge of the plateau facing Mont Lassois (Vix) on the west side. These stone mounds, each around fifteen metres in diameter, yielded 9 burials, including one of a warrior. In this case, the sword in an iron scabbard had been placed to the right of the body. The weapon dates from the early Second Iron Age Iron Age (LTA) like most of the other graves.

CGG_2_023635 (Vix,_tumulus_3_St_25).

#### France, Sainte Colombe-sur-Seine, Tumulus 3

Coordinates: 47.87, 4.52

This stone mound was excavated by R. Joffroy in 1973. It was located less than 1km from Mont Lassois (Vix). It was 18 m in diameter and 2 m high. The central tomb had been looted in the past, but some objects had remained in place or were located in the vertical cone of the clandestine excavation. Fragments of a bronze bowl, a painted truncated-cone ceramic plate and a duck-head fibula date this burial complex to the very early La Tène period (LTA).

CGG_2_023637 (Sainte_Colombe-sur-Seine,_Tumulus_3).

#### France, Tumulus de La Forêt de Châtillon S3

Coordinates: 47.82, 4.60

The tumulus of theChâtillon’s Forest, probably identified as the Val Thibaut tumulus, was explored at the end of the 19th century. 7 burials were identified. The bronze ornaments (torc, bracelets and anklets) date the graves to the end of the First Iron Age (Ha D2–D3) and the beginning of the Second Iron Age (LTA).

CGG_2_023644 (Tumulus_de_La_Forêt_de_Chatillon__1).

#### France, Vix mont Lassois

Coordinates: 47.90, 4.54

The hilltop settlement on the plateau of Saint-Marcel

Mont Lassois is a coherent and organised complex consisting of a quasi-urbanised space (a plateau on the summit comprising some 5 ha), a defensive circuit (on the edges of the plateau, its slopes and at the foot of the flanks of mont Saint-Marcel), burial grounds and open settlements in the Seine valley. Significant remains found on the plateau of Saint-Marcel and in the cemeteries at the foot of mont Lassois date to the Late Bronze Age (Final Bronze Age III)[544]. Thereafter, for a period of around two centuries, there was a hiatus in occupation until the Late Hallstatt period (Hallstatt D1).

The settlement on the upper plateau on mont Lassois is arranged on either side of a north-south axis. This "main street" structures the access to some fifteen enclosures delimited by palisaded ditches which contained the dwellings. In the south of the plateau, three buildings raised on piles represent three enormous collective granaries.

The settlement's regular layout suggests that it was planned from the outset and that this building work was controlled by the local power. Indications of a social hierarchy are discernible in the types of buildings erected and in the enclosed spaces: classic two-aisled houses are next to the monumental apsidal buildings.

Five large apsidal buildings stood at the heart of this organisation and in the centre of the largest enclosure. Two were extraordinarily large: Building 1 is 35 m long and 21 m wide and Building 2 is 25 m long and 11 m wide. In 2013, a sixth apsidal building of equally monumental size (30 m long and some 18 m wide) was found in the enclosure next to that containing the five other apsidal buildings. These buildings show to what degree the Hallstatt craftsmen mastered carpentry. For Building 1, they created an inner space of 500 $m^2$, spanned by a roof whose ridge towered at least 15 m above ground level.

The exact purpose of this building remains hypothetical: was it a public or private building, a seat or power, a religious or a domestic space? It is likely that it fulfilled all these functions since the political and religious spheres were not separate in complex chiefdoms.

Access to the plateau remains an open question, even though our Swiss colleagues from the University of Zurich have uncovered what is likely to be a monumental gate set in the western rampart at the site of Champ de Fossé. Another ascent to the plateau, via the eastern ramparts, may have existed, as suggested by the flattening of the ramparts' crests.

Monumentalising the defensive system

On the upper plateau, only the eastern edge appears to have been fortified by a rampart of *Pfostenschlitzmauer* type dated to the Late Hallstatt. Under this 9m-wide rampart, an earlier defence of the Late Bronze Age (Final Bronze Age III) had been built[546,548–550]. Excavations have shown that the Mont Lassois defences were monumental in character and went far beyond what was needed for purely defensive purposes. For example, the bank on which Rampart 3 was built (of which hardly anything remains) was 4 m high and 30 m wide at its base. The ditch that flanked it on the southern side was impressive, being 25 m wide and 10 m deep. The interior area protected by this complex defensive system (*intra muros* space) was some 40–45 ha in extent and we are only just beginning to understand how it was organised.

The tumulus of the Lady of Vix

The princely tomb of the Lady of Vix was discovered at the end of 1953 and excavated by R. Joffroy, M. Moisson and R. Paris in January/February 1954[544,546,551,552]. The tumulus, surrounded by a ring of stones, measured 38m in diameter and was on average 1m high. The 9m3 burial chamber housed the body of the deceased and her impressive viaticum. The "Lady of Vix" rested on the body of a chariot, the 4 wheels of which had been dismantled in antiquity and placed against the eastern wall of the burial chamber. The dead woman wore a hollow gold torque around her neck (Fig. S7.2). A second bronze torque, on which a leather strap had been wound, was placed on her stomach. On each arm were three lignite bracelets and a bronze bracelet with amber beads. The deceased was wearing a pair of bronze rings on her ankles. Eight fibulae were found on and beside the body, five of them decorated with amber, coral and gold leaf. The four ash wheels of the chariot were arranged along the eastern wall of the tomb. The body of the chariot was decorated with bronze plates and balusters. Typological parallels between the shell of the hubs and the balusters decorating the body's railing exist with the chariot discovered in a tomb in the Ca' Morta necropolis in Como (Italy), demonstrating the existence of links between these distant territories.

The most spectacular element of the viaticum is a large bronze crater, 1.64 m high and weighing 208,600 kg (Fig. S7.3). The vessel was cast in a cauldron; the handles, riveted to the mouldings on the rim, and the foot were cast using the lost-wax technique. The largest bronze vase known to us from antiquity was made around 530 BC by Greeks in a workshop in Magna Graecia near the towns of Taranto and Sybaris. A locally-made silver phiale and two ceramic bowls made in Athens, dated to 520/515 BC, had been placed on the rim of the crater lid. An oenochoe and three basins of Etruscan origin completed the bronze tableware set.

CGG_2_023310 (Vix,_tumulus_1,_Dame_de_Vix).
CGG_2_023626 (Vix,_Les_Tillies_Fait_29_Sp_04).
CGG_2_023627 (Vix,_Les_Tillies_Fait_14_Sp_06).
CGG_2_023628 (Vix,_Les_Tillies_Fait_14_Sp_07).
CGG_2_023630 (Vix_Mont_Saint_Marcel_Fait_1009).
CGG_2_023634 (Vix_les_Locheres).

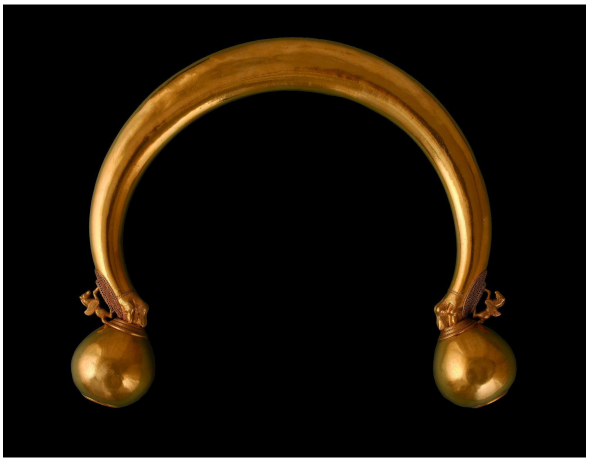

**Fig. S7.2** Gold torc of the Vix Princess. Photo Dominique Geoffroy.

Despite the presence in the burial of a number of imported objects that have been dated with some accuracy, it is not possible at present to date the burial with an accuracy of less than a quarter of a century, but it must be dated to the end of the first quarter of the 5th century BC. The DNA analysis presented in this article confirms that the woman was of indigenous origin and belonged to the social elite.

A new excavation of the tumulus was carried out in 2019 by a team from INRAP; although only a small half of the tumulus was excavated, a number of objects of finery were found, including fibulae, and the architecture of the monument was better understood.

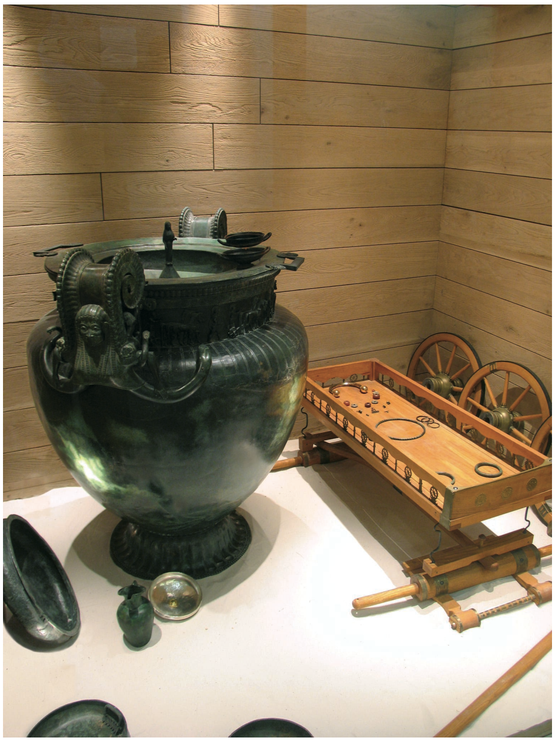

**Fig. S7.3:** Reconstitution of the Vix chamber Photo: Bruno Chaume (Musée du Pays du Châtillonnais).

### Germany

#### Germany, Saarland, Rubenheim

Coordinates: 49.17, 7.207

Walter Reinhard

Tumulus 17 (dm. 25.70/20.30; h. 0.99) lies to the west of the Rubenheim "Schornwald" necropolis, which comprises a total of 33 tumuli. It was the subject of a comprehensive study in 1984 by the Archaeological Society of the Saarpfalz District under the direction of the author. The burial mound, built for a Bronze Age D cremation grave, was clearly raised at the end of the Hallstatt period and surrounded by a stone crown. In addition to the richly endowed female grave 1 from the Late Hallstatt period, two unfurnished burials from the same period were discovered in the mass of the burial mound. Grave 2, oriented SE-NW, and Grave 3, oriented SW-NE, with a female aged between 50 and 60 years[553].

CGG_2_023734 (Rubenheim,_SPK._G._Hügel_17,_grab_3,_Nr._1).

CGG_2_023735 (Rubenheim, SPK. G. Hügel ?, Nr. 2).

### Georgia

#### Georgia, Digomi Valley, Treligorebiis

Coordinates: 41.767, 44.76

Irakli Akhvlediani

Treligorebiis situated on the natural hills of the Digomi Valley, a part of the Mtkvari (Kura) River valley between Mtskheta and Tbilisi, along the contemporary Georgian Military Road. The site was inhabited from the Chalcolithic Period until the Early Middle Ages. During the Late Bronze Age and Early Iron Age it was the largest settlement in the Caucasus, occupying five natural hills stretching about one mile along the main road connecting the city with other regions of Georgia. The favourable geographical location of the site, on the crossroads of the east-west and south-north trade routes, was the basis of its privileged state throughout the Caucasus.

The first clear evidence for sociopolitical complexity in southern Caucasia appears in the Late Bronze Age. The Late Bronze Age is marked most conspicuously by the reappearance of numerous permanent settlements in the form of variably sized stone-masonry fortresses built atop hills and outcrops. These fortified settlements are often associated with large cemeteries, such as Treligorebi located on the outskirts of modern Tbilisi, Georgia.

The multi-proxy study (grains, phytoliths, miliacin and isotopes) carried out on the Treli-Treligorebi site, during the transition from the Late Bronze Age to the Early Iron Age, demonstrated the presence of millet cultivation and consumption by domestic animals and humans, of up to 40%[554].

CGG_2_021779 (Digomi_Kat_N1092).

### Hungary

#### Hungary, Szeged-Kiskundorozsma-Nagyszék II. (site 26/72 No. 34, M5 motorway)

Coordinates: 46.27, 20.06

The archaeological site was excavated by Csaba Szalontai and Katalin Tóth during two seasons, in 1998 and 1999. The site is situated on the Banks of Maty Creek in Nagyszék near the western boundary of Szeged-Kiskundorozsma. A part of an extensive Sarmatian settlement (approximately 40.000 square metres of the known 72.000–108000 square metre settlement) and the remains of a small cemetery including 24 graves had been excavated [555–557]. The settlement seems to have been long-lived. Its history started in the second half of the 2nd century AD and lasted well into the 5th century AD, based on the excavated wheeled pottery, thinned with pebbles.

The majority of the graves had been looted by contemporary robbers. Based on the grave goods the cemetery spanned a short period (the late 2nd and the first half of the 3rd century AD).

CGG_2_021918 (Kiskundorozsma_1537).

CGG_2_021922 (Kiskundorozsma_1617).

CGG_2_021923 (Kiskundorozsma_1607).

CGG_2_021924 (Kiskundorozsma_1624).

#### Hungary, Madaras-Halmok, Madaras

Coordinates: 46.06, 19.27

The excavation of the Sarmatian period cemetery at Madaras-Halmok is attributed to Mihály Kőhegyi (Baja, Türr István Museum). The excavation began in 1963 and continued with several interruptions until 1975. The completely excavated burial site is Hungary's largest Sarmatian period cemetery, used from the late 2nd-century to the turn of the 4th–5th century. In the 145,000 square meter area, 666 graves were found, although not all of them were Sarmatian graves, as several Árpádian Age (11–14th c. CE) graves (45 graves) in the center of the cemetery and a larger Árpádian Age settlement were also excavated. Most of the graves in the cemetery were robbed. The skeletal material is very fragmentary. The results of the archaeological and anthropological analyses were summarized in [530,558].

CGG_2_021896 (Madaras_287).

CGG_2_021897 (Madaras_427).

CGG_2_021898 (Madaras_342).

CGG_2_021899 (Madaras_368_B).

CGG_2_021902 (Madaras_195).

CGG_2_021908 (Madaras_194).

CGG_2_021909 (Madaras_215/A).

CGG_2_021912 (Madaras_214).

CGG_2_021913 (Madaras_335).

CGG_2_021916 (Madaras_398).

CGG_2_021917 (Madaras_218).

### Italy

#### Italy, Marche, Ripa Bianca

Coordinates: 43.71, 13.09

Serena Sabatini

Ripa Bianca or Ripabianca lies in a sub-coastal area a few kilometres north of the modern city of Ancona in the Marche region. The region became part of the so-called Longobard/Lombard kingdom at the time of its maximum extension[559]; however, the site from which our sample was collected is not published and we are not able to provide detailed information about the context.

CGG_2_019380 (MYRAN_B).

### Lithuania

#### Lithuania, Kaunas, Marvelė

Coordinates: 54,89, 23,87

Rimantas Jankauskas

The Marvelė burial site is located in the western part of the Kaunas city, on the lower terrace of the southern (left) bank of Nemunas River. Archaeological excavations were conducted during 1991–2011 seasons. Total area estimated 30–40 thousand sq.m., containing no less than 1590 human graves (both inhumations and cremations)[560], over 100 graves of horses[561]. Burials should be attributed to the cultural group of the "Flat burials of the central Lithuania", and dating stretches from the 2nd to the 12th centuries, i.e. embracing Roman, Folks Wandering, Viking periods.

CGG_2_017685 (I001).

#### Lithuania, Kelmė, Maudžiorai

Coordinates: 55.85, 22.62

Rimantas Jankauskas

Excavated episodically from 1964 till 1984. Over 300 flat burials dated 8–9th c.c. Archaeologists attribute them to Samogitian entity, although there are artefacts typical to Curonians and Semigalians[562–565].

CGG_2_017694 (1460B).

#### Lithuania, Panevėžys, Berčiūnai

Coordinates: 55.74, 24.22

Rimantas Jankauskas

Barrows located west from Panevėžys town, on the right (western) bank of Nevėžis river. Around 40 semi destructed barrows, dated 3rd–7th c.c. Barrows, known since the beginning of the 20th c., are 7–12 m wide and up to 1,2 m high, some of them with stone circles.

CGG_2_017690 (I287C).

#### Lithuania, Radviliškis, Kairėnėliai

Coordinates: 55.51, 23.66

Rimantas Jankauskas

Partly deranged during recent earthworks and agricultural activities inhumations. Excavated in 1977–1978, 30 inhumations and one cremation found, archaeologically dated 5th–6th c.c. AD.

Burials are quite rich in artefacts (gender-specific jewellery, weapons, instruments and tools)[566].

CGG_2_017691 (729).

#### Lithuania, Raseiniai, Kalniškiai

Coordinates: 55.26, 23.44

Rimantas Jankauskas

Flat burials, known since 1935–37. Investigated from 1984 till 2004. In total, 277 burials were found, among them 259 inhumations dated 5–6$^{th}$ c.c. and 18 7–8$^{th}$ c.c. cremations. Skeletal materials are poorly preserved. Inhumations rich with artefacts (gender-specific jewellery, tools, weapons), some in wooden coffins, males oriented NW, females SE[567].

CGG_2_017692 (1500(18)).

#### Lithuania, Rokiškis, Vaineikiai

Coordinates: 55.72, 25.53

Rimantas Jankauskas

Vaineikiai barrows, lined with stones, are known from 19th c., their total estimated number - c.50. During excavations in 1977–1978, three barrows were investigated which contained 15 inhumations, mostly dated to the 3rd–4th c.c., some to the 7–8th c.c[568,569].

CGG_2_017687 (I348).

#### Lithuania, Šiauliai, Kalneliai

Coordinates: 55.98, 23.10

Rimantas Jankauskas

Partly destroyed barrows or burials lined with stone circles. Inhumations roughly N-S oriented. Dating according to artefacts – 2nd–4th c.c. AD[570].

CGG_2_017688 (1457).

### The Netherlands

#### The Netherlands, Hogebeintum

Coordinates: 53.33, 5.85

Stijn Heeren, Lisette Kootker

The cemetery of Hogebeintum was discovered and researched when the *wierde* (mound) was quarried for its fertile soil in 1904–1905. At least 142 burials were documented, but the cemetery must have been (much) larger. Judging from the grave goods and radiocarbon dates of cremated bone and charcoal, the cemetery was in use from the early 5th to the first half of the 8th century CE (approx. 400–730 CE; Dutch Early Medieval period). A few older dates were explained as resulting from the use of old wood for cremation[571]. The catalogue holds at least 94 cremation burials, several buried dogs, and at least 48 human inhumation burials. To date, 28 (partial) skeletons are still available for research[571]. Ten graves were selected for multi-dental elemental Sr-O-C sampling[572]. Five of these individuals (FM95, FM96, FM98, FM99, and FM101) were selected for $^{14}C$ and aDNA analysis, of which four are included in the present study (Table S7.2.31).

| | | | | | | | | | Costa | | |
|---|---|---|---|---|---|---|---|---|---|---|---|
| **CGG ID** | **Sample ID** | **Sex** | **$^{14}C$ code** | **$^{14}C$ date BP** | **Calibrated date** | **Element (FDI)** | **$^{87}Sr/^{86}Sr$** | **$\delta^{18}O_{PDB}$ (‰)** | **$\delta^{13}C$ (‰ vs PDB)** | **$\delta^{15}N$ (‰ vs AIR)** | **C:N** |
| CGG_2_024691 | FM95 – kist 249 | M | Ua–69910 | 1587 ± 29 | AD 419 – AD 547 | 36 | 0.710382 | –4.77 | –19.9 | 10.9 | 3.2 |
| | | | | | | 37 | 0.710669 | –6.15 | | | |
| | | | | | | 28 | 0.709513 | –5.79 | | | |
| CGG_2_024692 | FM96 – kist 249 | M | Ua–69911 | 1688 ± 28 | AD 258 – AD 419 | 26 | 0.710270 | –5.27 | –19.8 | 11.2 | 3.2 |
| | | | | | | 17 | 0.710618 | –5.57 | | | |

| | | | | | | | | | | | |
|---|---|---|---|---|---|---|---|---|---|---|---|
| | | | | | | 38 | 0.709540 | –5.6 | | | |
| CGG_2_024694 | FM99 – kist 249 II | M | Ua–69913 | 1684 ± 28 | AD 258 – AD 422 | 46 | 0.710224 | –6.53 | –20.3 | 11.2 | 3.2 |
| | | | | | | 47 | 0.710896 | –6.32 | | | |
| CGG_2_024695 | FM101 – 28–360 | F | Ua–69915 | 1677 ± 29 | AD 257 – AD 432 | 46 | 0.711104 | –4.78 | –19.8 | 10.4 | 3.2 |
| | | | | | | 37 | 0.711370 | –5.82 | | | |
| | | | | | | 48 | 0.710635 | –5.45 | | | |

**Table S7.2.31:** List of the samples studied from Hogebeintum, their biological sex (aDNA), $^{14}C$, and Sr-O-C-N isotope data. Sr-O-C isotope and $^{14}C$ data were previously published. All $^{87}Sr/^{86}Sr$ are incompatible with the expected local Sr signature and provide evidence for childhood mobility[572,573]. Key: FDI - World Dental Federation; PDB: Peedee Belemnite; AIR: Ambient Inhalable Reservoir. The 2SE internal error of the $^{87}Sr/^{86}Sr$ were <0.000010.

#### The Netherlands, Oosterbeintum

Coordinates: 53.33, 5.87

Stijn Heeren, Lisette Kootker

A small cemetery was excavated in 1987 on the southeastern part of the *wierde* of Oosterbeintum. The cemetery held between 33 to possibly 48 cremated burials and 46 inhumation graves. The cremation graves date between the early 5th century and the first half of the 8th century CE, while the inhumation graves are probably a bit younger, from the second half of the 5th century to the first half of the 8th century CE[574].

In a previous study, McManus et al, 2013[575] 24 dental elements of the Oosterbeintum population for strontium isotope analysis, restricted to a single dental element per individual[575]. In Kootker and Heeren, 2022[572], an additional sixteen samples from eleven individuals were selected, in order to obtain subsequent molar 1-2-3 data where available. Five individuals were included in this study (Table S7.2.32; S60, S335, S410, S487 (485B), and S570).

| | | | | | | | | | Costa | | |
|---|---|---|---|---|---|---|---|---|---|---|---|
| **CGG ID** | **Sample ID** | **Sex** | **$^{14}$C code** | **$^{14}$C date (BP)** | **Calibrated date** | **Element (FDI)** | **$^{87}Sr/^{86}Sr$** | **$\delta^{18}O_{PDB}$** | **$\delta^{13}$C ‰ (PDB)** | **$\delta^{15}$N ‰ (AIR)** | **C:N** |
| CGG_2_024699 | S60 | F | Ua–69995 | 1611 ± 27 | AD 416–AD 538 | 37 | 0.709822* | - | –19.8 | 13.7 | 3.2 |
| | | | | | | 38 | 0.709781 | –4.51 | | | |
| CGG_2_024707 | S487 (485B) | F | Ua–70003 | 1561 ± 27 | AD 430–AD 569 | 36 | 0.709322 | –4.59 | –20.3 | 11.7 | 3.2 |
| | | | | | | 44 | 0.712460* | - | | | |
| CGG_2_024708 | S570 | F | Ua–70004 | 1561 ± 27 | AD 430–AD 569 | 26 | 0.708884 | –4.67 | –20.2 | 9.1 | 3.2 |
| | | | | | | 27 | 0.708791* | - | | | |
| CGG_2_024702 | S410 | M | Ua–69998 | 1556 ± 28 | AD 430–AD 575 | 36 | 0.709133 | –4.00 | –19.4 | 9.1 | 3.2 |
| | | | | | | 35 | 0.708824* | - | | | |
| | | | | | | 38 | 0.709110 | –6.63 | | | |

| | | | | | | | | | | | |
|---|---|---|---|---|---|---|---|---|---|---|---|
| CGG_2_024700 | S335 | M | Ua–69996 | 1535 ± 28 | AD 435–AD 599 | 36 | 0.708874 | –5.89 | –19.5 | 11.6 | 3.2 |
| | | | | | | 27 | 0.709232* | - | | | |
| | | | | | | 48 | 0.708156 | –5.94 | | | |

**Table S7.2.32:** List of the samples studied from Oosterbeintum, their biological sex (aDNA), $^{14}$C, and Sr-O–C-N isotope data. All $^{87}$Sr/$^{86}$Sr are incompatible with the expected local Sr signature and provide evidence for childhood mobility[572,573,575]. Key: BP - Before Present; FDI - World Dental Federation; * - data generated by[575]; PDB: Peedee Belemnite; AIR: Ambient Inhalable Reservoir.

#### The Netherlands, Valkenburg (ZH)

Coordinates: 52.17, 4.43

Lisette Kootker, Maura De Coster

A few kilometres from where the Lower German *limes*, recently recognized as a World Heritage site by UNESCO, ends in the North Sea, the unique Roman cemetery of Valkenburg Marktveld has been excavated. Located circa 500 m. south of the auxiliary fort, the archaeological site plays a key role in understanding the population dynamics in a Roman borderscape and coastal landscape. This cemetery was used between 50 – 300 CE for the entire military community that consisted of men, women and children, who lived in the vicinity of the auxiliary fort. The remains of over 650 individuals were recovered, 134 of which are inhumations (47 adults, 87 children and infants)[576,577] an extraordinary number as cremation dominates the Roman burial record in the Netherlands.

Within the framework 'Constructing the Limes: Employing citizen science to understand borders and border systems from the Roman period until today' (C-Limes), funded by the Dutch Research Council (NWO) as part of the Dutch Research Agenda (NWA, 2021–2026, project number: NWA.1292.19.364), 28 adult individuals were available for Sr-O-C isotope research.

As part of the collaboration between C-LIMES and the GeoGenetics Centre at the University of Copenhagen, petrous parts and/or dental elements of 27 individuals were collected for aDNA analysis (Table S7.34). A more detailed paper integrating archaeological and historical data, $^{14}$C and Sr-O-C-N isotope data, and aDNA data (kinship, etc.) from Valkenburg Marktveld is forthcoming (De Coster et al., in prep). The following Table S7.2.33 gives the list of the samples studied from Valkenburg (ZH), dating, inventory number and type of material used for aDNA analysis.

| | | | | | **Old sample/individual/Find/Feature ID numbers** | | | | |
|---|---|---|---|---|---|---|---|---|---|
| **CGG ID** | **Sample ID** | **$^{14}$C code** | **$^{14}$C date (BP)** | **Calibrated date** | **ROB** | **Lonnée** | **Find** | **Feature** | **Sample type** |
| CGG_2_107754 | CL002 | DeA-36906 | 1956 ± 19 | AD 13 – AD 123 | I085 | 5 | 102 | | Bone |
| CGG_2_107733 | CL005 | - | - | - | I081 | 10 | 260 | | Tooth |
| CGG_2_107746 | CL005 | DeA-39646 | 1982 ± 19 | 39 BC – AD 114 | I081 | 10 | 260 | | Bone |

| | | | | | | | | | |
|---|---|---|---|---|---|---|---|---|---|
| CGG_2_107763 | CL006 | DeA-39647 | 1852 ± 19 | AD 125 – AD 233 | I079 | 15 | 265 | | Bone |
| CGG_2_107747 | CL007 | DeA-39648 | 1878 ± 19 | AD 120 – AD 225 | I002 | 22 | 38 | | Bone |
| CGG_2_107760 | CL008 | - | - | - | I007 | 23 | 170 | | Bone |
| CGG_2_107767 | CL010 | DeA-39649 | 1944 ± 18 | 98 BC – AD 25 | I028 | 30 | 223 | | Bone |
| CGG_2_107761 | CL013 | - | - | - | I037 | 39 | 313 | | Bone |
| CGG_2_107734 | CL013 | - | - | - | I037 | 39 | 313 | | Tooth |
| CGG_2_107749 | CL015 | DeA-39650 | 2129 ± 23 | 342 BC – 52 BC | I024 | 42 | 248 | | Bone |
| CGG_2_107735 | CL017 | DeA-39651 | 1910 ± 19 | AD 72 – AD 207 | I021 | 50 | 181 | | Tooth |
| CGG_2_107745 | CL017 | DeA-39651 | 1910 ± 19 | AD 72 – AD 207 | I021 | 50 | 181 | | Bone |
| CGG_2_107757 | CL018 | - | - | - | I043 | 52 | 237 | | Bone |
| CGG_2_107736 | CL018 | DeA-39652 | 1898 ± 19 | AD 80 – AD 213 | I043 | 52 | 237 | | Tooth |
| CGG_2_107752 | CL019 | DeA-39653 | 1936 ± 20 | AD 21 – AD 200 | I051 | 53 | 118 | | Bone |
| CGG_2_107737 | CL021 | - | - | - | I055 | | | | |
| CGG_2_107738 | CL023 | DeA-39654 | 1936 ± 20 | AD 21 – AD 200 | I041 | 62 | 334 | | Tooth |
| CGG_2_107750 | CL023 | DeA-39654 | 1936 ± 20 | AD 21 – AD 200 | I041 | 62 | 334 | | Bone |
| CGG_2_107768 | CL025 | - | - | - | - | 16 | 8 | 070.0016 | Bone |

| | | | | | | | | | |
|---|---|---|---|---|---|---|---|---|---|
| CGG_2_107739 | CL026 | DeA-36907 | 1912 ± 19 | AD 70 – AD 206 | I031 | 66 | 263 | | Tooth |
| CGG_2_107744 | CL026 | DeA-36907 | 1912 ± 19 | AD 70 – AD 206 | I031 | 66 | 263 | | Bone |
| CGG_2_107740 | CL027 | - | - | - | I042 | 68 | 337 | | |
| CGG_2_107751 | CL027 | DeA-36908 | 1841 ± 22 | AD 127 – AD 244 | I042 | 68 | 337 | | Bone |
| CGG_2_107748 | CL029 | DeA-39656 | 1950 ± 19 | AD 16 – AD 125 | I018 | 69 | 167 | | Bone |
| CGG_2_107741 | CL029 | DeA-39656 | 1950 ± 19 | AD 16 – AD 125 | I018 | 69 | 167 | | Tooth |
| CGG_2_107765 | CL030 | DeA-36909 | 1926 ± 19 | 91 BC – AD 60 | I114 | 72 | 242 | | Bone |
| CGG_2_107742 | CL031 | DeA-36910 | 1879 ± 20 | AD 120 – AD 224 | I095 | 76 | 31 | | Tooth |
| CGG_2_107753 | CL032 | - | - | - | I052 | 86 | 189 | | Bone |
| CGG_2_107755 | CL033 | - | - | - | I120 | 87 | 280 | | Bone |
| CGG_2_107766 | CL035 | DeA-39657 | 2038 ± 19 | AD 27 – AD 203 | I121 | 95 | 59 | | Bone |
| CGG_2_107756 | CL041 | DeA-39660 | 1992 ± 19 | 42 BC – AD 106 | I134 | 129 | 71 | | Bone |
| CGG_2_107743 | CL041 | - | - | - | I134 | 129 | 71 | | Tooth |
| CGG_2_107762 | CL043 | DeA-36912 | 1863 ± 21 | 50 BC – AD 250 | I074 | 137 | 182 | | Bone |
| CGG_2_107764 | CL046 | DeA-39661 | 2026 ± 21 | AD 129 – AD 236 | I092 | 141 | 160 | | Bone |

| | | | | | | | | | |
|---|---|---|---|---|---|---|---|---|---|
| CGG_2_107758 | CL048 | DeA-39662 | 1894 ± 18 | 50 BC – AD 250 | I001A | 19 | 155A | | Bone |
| CGG_2_107759 | CL049 | DeA-39663 | 1976 ± 20 | AD 83 – AD 212 | I001B | 19 | 155B | | Bone |

**Table S7.2.33:** List of the samples studied from Valkenburg (ZH), dating, inventory number and material used for aDNA analysis..

### Portugal

#### Portugal, Estremadura, Gruta do Caldeirão

Coordinates: 38.67, 9.16

The layer of provenience is a thick, badger-burrowed, and homogenised dark cave earth containing abundant ceramics, mostly of the Bronze Age, and other artefacts ranging from the Late Neolithic to the 17th century. Among the iron artefacts, two buckles (one of armour, another of bridle) were diagnostic of the Visigothic period[578] (118–119; Fig. 9.6, nos. 3–4). A human bone sample from this layer had already been dated to the same time range indicated by your new results.

That the three samples from layer ABC–D yielded post-Neolithic ages is hardly surprising, as the human remains therein are associated with artefacts ranging from the Late Neolithic to the early Middle ages and, previously, a rib from this layer had already been dated to the Early Medieval time range. Most of the ceramics are Bronze Age, yet all the human bone dates so far are Medieval, which would seem to suggest that the funerary activity of the Bronze Age consisted of the deposition of incinerations.

The full list of results for layer ABC–D is below (Table S7.2.34). Considering that P11sc375 and P13–32 are the same individual and that the two TO dates are on the same sample, their averages can be calculated and, after calibration, compared with the result for P11–73. The three ages, in the rows highlighted yellow, are statistically distinct and so the dated rib belongs to a third individual. This evidence shows occasional but recursive use of the cave as a burial site between the mid-6th and the mid-9th centuries AD.

| **CGG number** | **Sample type** | **Lab dating code** | **$^{14}C$** | **±** | **Date** |
|---|---|---|---|---|---|
| CGG_2_018558 (P11-sc375) | left temporal with petrous | UBA-40089 | 1213 | 27 | |
| CGG_2_018559 (P13-32) | right temporal with petrous | UBA-40090 | 1258 | 31 | |
| | Average | | 1232 | 20 | 691–877 AD, 2σ |
| P11-73 | left temporal with petrous | UBA-40091 | 1313 | 26 | 657–767 AD, 2σ |
| P11-112 | rib fragment | TO-351 | 1420 | 50 | |
| replicate | rib fragment | TO-351a | 1490 | 70 | |

| CGG number | Sample type | Lab dating code | $^{14}C$ | ± | Date |
|---|---|---|---|---|---|
| | Average | | 1444 | 41 | 548–658 AD, 2σ |

**Table S7.2.34:** List of $^{14}C$ dating results for the samples from Estremadura, Gruta do Caldeirão, Portugal, layers ABC–D.

### Russia

Mazuninskaya culture

Elizaveta Chernykh

The Mazuninskaya culture was described by V.Gening (1968). Named after the Mazuniskiy necropolis, the culture was distributed in 3–5 centuries AD in the middle Kama River. The culture developed on the basis of the local P'yanoborye culture. The morphological studies revealed complex structure of the Mazunino populations which resulted from the admixture of different groups of European and Uralian groups. It is argued that Mazunino populations have the closest biological affinities with modern Uralic-speaking Mari and Moksha people but not Udmurts[579,580].

#### Russia, Boyarskiy

Coordinates: 56.05, 54.02

The Boyarskiy (Arai) burial ground attributed to the Mazuninskaya culture (the end of 3–4 centuries AD) is located on the narrow terrace of the right bank of the Kama River, Karakulinskiy district, the Udmurtia Republic, Russia. In 2002–2009 183 burials consisting 199 skeletons were excavated by E.M.Chernykh.

The burials were made in rectangular ground pits with a depth of 20 to 130 cm. The most skeletons lay with their heads oriented to the SW and WSW. Remains of wooden burial structures (similar to boxes) were found in 1/3 of the burials. In few cases plank log decking, burials in the decks, wrapping in birch bark, and swaddling were detected. There were cases of tying the legs of the deceased, the crooked position of buried on the side, burials without heads, and violations of anatomical order. The frequency of child burials is about 25%. About 42% of all burials have no inventory, a half of them are child ones. The number male and female non-inventory burials are equal. The specific male and female inventory kits can be detected. The Boyarsky burial ground share a specific feature of the Mazunino necropolises - the placement of “female” inventory kits not in the place where they were worn, but in special “donation (sacrificial)” places. While in other Mazunino burial grounds such places were usually located at the head of the buried in the Boyarskiy cemetery the feet area was used.

According to the cranial studies the population which left Boyarskiy cemetery resulted from the admixture of two groups. The first one is possibly related to steppe populations alike Sarmatians while second is close to historical Finnish-speaking populations from the Volga region.

CGG_2_021459 (Boyasky_III-V_AD_8).

CGG_2_021460 (Boyasky_III-V_AD_7).

CGG_2_021461 (Boyasky_III-V_AD_6).

CGG_2_021463 (Boyasky_III-V_AD_3).

#### Russia, Dybrovskiy

Coordinates: 53.63, 33.37

The Dubrovskiy burial ground (IV – early V centuries AD) is located on a high steep slope of the left bank of the Shekhostanka River (third-order tributary of the Kama River), Kiyasovsky district, the Udmurt Republic, Russia.

In 2009–2017 Perevozchikova and V.Bertz excavated 204 burials. Most burials are single made in narrow rectangular pits of about 0,6 m depth. The burials were arranged in rows. The buried lay stretched with their heads to the South down of the river flow. Several specific features of the burial rituals which are not characteristic for the local cemeteries of the time were noted. These are: coating the walls with clay, traces of firing, adding chalk to the bottom, backfilling with stones. While the male military inventory kit included swords, combat knives, spears, arrows, "pink salmon braids", and horse bits, the Female kit consisted of headdresses decorated with metal clips, plaques, beads and beads, temporal pendants, chains.

CGG_2_021462 (Dubrovsky_III-V_4).

CGG_2_021464 (Dubrovsky_III-V_2).

#### Russia, Ingushetia, Ekazhevo 1

Coordinates: 43.208907, 44.820441

V.Yu. Malashev, U.B. Gadiev, E.V. Balanovskaya

The Ekazhevo I burial ground (“Ekazhevo 1st mounds”) is located on the southern and southwestern outskirts of the village of Ekazhevo (Ingushetia, Russia) and was the necropolis of the Achim-Boarz settlement. As a result of excavations carried out by an expedition of the Institute of Archeology of the Russian Academy of Sciences (Moscow) in 1996, 2016 and 2018 together with the State Public Institution “Archaeological Center named after. E.I. Krupnov" (Nazran), 6 burials under the burial mounds in the catacombs, characteristic of the early stage of the Alan culture and dating from the middle of the 3rd to the beginning of the 4th centuries, were studied (mounds 6 and 7 – mid-3rd century AD, mounds 2 and 3 – second half of the 3rd century AD, mounds 1 and 4 – beginning of the 4th century AD)[581,582].

According to anthropological definitions and the military composition of the grave goods, the burials belonged to men. The size of the burial structures (the depth of the catacombs is 7–8 m), the funeral equipment (preserved after grave robberies in ancient times - bladed weapons, harness sets, high quality silver belt sets, gold plaques) indicate a high-status social group. The territory of the burial ground, where the burials of the military elite took place, was isolated from the main part of the necropolis.

CGG_2_021451 (Ekazhevo_2).

#### Russia, Kabardino-Balkaria, Zaragizh

Coordinates: 43.33, 43.71

B. Kh. Atabiev

Zaragizh is a complex of archaeological sites from the 3rd century - the middle of the 1st millennium AD on the border of the plain and foothill zones of the Caucasus, on the Cherek River near the village of Zaragizh (Chereksky district of Kabardino-Balkaria, Russia) [583]. Includes a vast settlement with a complex system of fortifications (the oldest - Koban cultures), burial mounds and soil burials located on the site of an earlier irrigation system. The settlement was discovered by V. M. Batchaev in 1986, the burial grounds by B. Kh. oval (early) and rectangular (late) catacombs with a stone foundation and, as a rule, a perpendicular dromos; the evolution of the design of the vaults of the chambers from shed to arched and lancet can be traced. about finished and extensive catacombs up to 7 meters deep. In the catacombs - from 2 to 21 stucco and circular bowls, mugs, jugs, weapons (including those with wooden and gold details, decorated in the cloisonne style), belt set, brooches and other decorations (including with the use of gold, various inserts), amulets, toiletries, Roman coins, indications, and others. Zaragizh is considered as the center of one of the Sarmatian-Alanian groups in the Caucasus.

CGG_2_021996 (Katak_N_30).

CGG_2_022000 (kat_N_291).

CGG_2_022002 (Kat_N_175).

CGG_2_022003 (Kat_N_36).2

CGG_2_022004 (Kat_N_333).

CGG_2_022005 (Kat_N_324).

#### Russia, Tuva, Aymyrlyg

Coordinates: 51.88, 95.62

Eileen Murphy, Vyacheslav Moiseyev

The cemetery complex of Aymyrlyg is located in the Ulug-Khemski region of Tuva, deep in southern Siberia and near the border with Mongolia. It spanned a vast distance of some 10 km and was located alongside a tributary of the mighty Yenisei river. It was excavated over a 16 year period between the 1960s and 1980s by Dr A.M. Mandelshtam and Dr E.U. Stambulnik of the Sayano-Tuvinsky expedition team from the Institute of the History of Material Culture in St. Petersburg because the entire zone was scheduled to be flooded as part of the workings of a hydro-electric power station. The human skeletal remains were brought to the Department of Physical Anthropology of the Kunstkamera Museum. A detailed osteoarchaeological and palaeopathological analysis was undertaken of the population for PhD research[584] and this study has formed the basis for additional and ongoing projects. Approximately 800 individuals were recovered from the burial ground – most of the earlier 607 interments dated to between the 5th and 2nd centuries BC and were attributed to the Uyuk culture of the later Scythian period[585]. The remainder of the population was derived from the Shurmak culture of the early Hunno-Sarmatian period and was associated with the expansion of the Xiongnu empire into Tuva. They are largely thought to date to between the 3rd century BC and the 2nd century AD[586,587].

Mandelshtam and Stambulnik (1992)[588] were of the opinion that the change from the Uyuk Culture to the Shurmak Culture involved the movement of new groups of people into Tuva. A detailed craniometric and osteometric analysis was undertaken on a number of individuals from Aymyrlyg[589]. The findings indicated that the Uyuk Culture individuals at Aymyrlyg displayed more European characteristics than Shurmak Culture individuals who displayed more pronounced Asian features. In addition, differences in the degree of Asian and European admixture were evident between Shurmak Culture individuals buried in ground pits and stone cists. This finding was interpreted as evidence that the population burying their dead at Aymyrlyg during the Hunno-Sarmatian period was heterogeneous, comprising both individuals of the preceding Uyuk Culture and those of the more recently arrived Shurmak Culture[589].

In a previous study, aDNA of mycobacterium bovis was identified in five individuals from the burial ground, none of which are included in the current study[590,591]. The date range of these individuals extended from the 4th century BC to the 4th century AD. A weak positive for brucellosis was identified in one individual but this could not be replicated at a later date.

CGG_2_021493 (6862-363, *Burial XX.10. Sk. 1,* Scythian). The individual was recovered from a rectangular stone-cist along with three other individuals including:

CGG_2_021495 (6862-364) The skeleton was near-complete but had been disturbed. It was situated near the southwestern wall of the tomb and the skull lay near a stone slab upon which was a clay pot. The skeleton was that of a 17–25-year-old female, with an estimated stature of 161.1 cm. Osteoarthritis was evident on several rib heads, while osteophytosis and Schmorl's nodes were present on thoracic and lumbar vertebrae. Reactive new bone formation, suggestive of a pulmonary infection, was observed on the visceral surfaces of three right ribs and on the dorsal and visceral surfaces of a left rib. Periodontal disease of considerable severity and a large dental abscess were visible, and the individual had lost one tooth ante-mortem.

CGG_2_021494 (6862-138, *Burial VI.10. Sk. 1,* Scythian). The individual was recovered from within a log house tomb along with the remains of three disturbed skeletons. The skeleton was near complete. The individual lay on the left side with the head to the west. The legs were flexed and the arms were extended with the hands in front of the pelvis. The skull lay on a stone slab. The individual was associated with a notable array of grave goods – fragments of an iron pin, covered with gold foil, were located near the slab associated with the skull. A further 11 objects appear to have been directly connected to Sk. 1, including a bronze mirror within a leather case. In addition, to these the distal ends of the lower arm bones, the left ilium and the midshaft of the left femur displayed green discoloration suggestive of original association with bronze objects. The skeleton was that of an adult female, aged 25–35 years, with an estimated stature of 156.5 cm. The bones were generally gracile and developmental dysplasia of the left hip was evident. The left leg was atrophied but osteoarthritis was visible on and on the bones of the right leg which suggested it had borne weight. In addition, pronounced Schmorl's nodes were evident on the lower thoracic and upper lumbar vertebrae. The morphology of the right arm bones was suggestive that she had used crutches that placed particularly strain on the right side of the body. Linear dental enamel hypoplasia is indicative of childhood physiological stress. Slight periodontal disease and one caries were visible.

CGG_2_021495 (6862-364, *Burial XX.10. Sk. 2,* Scythian). The individual was recovered from a rectangular stone-cist along with three other individuals including:

CGG_2_021493 (6862-363). The skeleton was practically complete. The individual lay on their right side, and had flexed arms and legs. The head was orientated towards the NW and was situated at the edge of a stone slab. A clay pot was located near the stone slab. An area of green discoloration on the left iliac crest is suggestive that the individual had been buried with a bronze object. The skeleton was that of an adult male of 25–35 years, with an estimated stature of 170.8 cm. A well-healed fracture was visible on the right clavicle. Vertebral osteophytosis was evident on cervical, thoracic and lumbar vertebrae, while Schmorl's nodes were evident on many thoracic and lumbar vertebrae. Osteoarthritis was visible at the distal end of the left first metacarpal. The individual was robust with pronounced entheses. Reactive new bone formation, suggestive of pulmonary infection, was evident on the visceral surfaces of five right ribs. Considerable periodontal disease, two caries and five dental abscesses were observed, and six teeth had been lost ante-mortem.

CGG_2_021492 (6862-359, *Burial XX.9. Sk. 1*, Scythian). The individual was recovered from a log house tomb along with the remains of a further eight individuals. The skeleton was practically complete. The individual lay on their left side, and had flexed arms and legs. The head was oriented to the west and the skull lay on a stone slab. A clay pot was positioned beside the stone slab at the head. Fragments of an iron object were located on top of the right side of the pelvis. A bronze container associated with a belt fragment and two pieces of an iron object were recovered adjacent to the left pelvis. A bronze buckle and fragment of cowrie shell were located beside the left femur, while the remains of an iron knife were retrieved from beneath the right femur. Three arrowheads lay near the individual's feet. The skeleton was that of a young adult male, with an estimated stature of 165.9 cm. Reactive new bone formation was present on the left humerus, left tibia and both fibulae, as well as on right rib 9 and left rib 7 and the anterior surfaces of lumbar vertebrae 1–3 and 5. These lesions are indicative of systemic infection but the new bone formation on the lumbar vertebrae are also suggestive of brucellosis or tuberculosis. Schmorl's nodes were visible on most thoracic and lumbar vertebrae. Periodontal disease of slight severity was observed.

CGG_2_021497 (6862-381, *Burial XXIII.10. Sk. 2*, Scythian). The individual was recovered from within a log house tomb but the archive report did not include details of the context. Based on the context numbers of skeletons it is evident that at least two other individuals were recovered from the tomb. Only the skull was available for analysis. The skeleton was that of a probable adult female aged 25–35 years. The left orbit had an abnormal enlarged appearance suggestive of the presence of a soft tissue mass within the orbit. The right orbit was also slightly enlarged and asymmetry of the nasal bones was evident. It was considered that the lesions may be compatible with the genetic condition neurofibromatosis[592,593].

CGG_2_021499 (7256-85 *Burial XXXI.177*, Hunno-Sarmatian (Xiongnu)). The archive report for the year of excavation (1982 or 1983) was unavailable but the context number suggests an individual burial, typical for the Hunno-Sarmatian phase of burial. The skeleton was that of a young adult female of 17–25 years, with an estimated stature of 145.8 cm. Lytic lesions, suggestive of a non-specific infection (or perhaps tuberculosis or brucellosis although not typical lesions) were evident in the right mandible and superior surface of the sixth lumbar vertebra, with a large sub-circular lesion measuring almost 30 mm present at the proximal metaphysis of the left tibia. Reactive new bone formation was visible at the midshaft of the right tibia. Slight periodontal disease and two caries were evident.

CGG_2_021498 (7256-98, Burial XXXI.205, Hunno-Sarmatian (Xiongnu). The archive report for the year of excavation (1983) was unavailable but the context number suggests an individual burial, typical for the Hunno-Sarmatian phase of burial. The skeleton was that of an adult female of 25–35 years, with an estimated stature of 149.7 cm. Plaques of reactive new bone formation were visible at the heads and necks of the left ribs 4–9 and right ribs 7. 8 and 11. The lesions are suggestive of a pulmonary infection. Schmorl's nodes were present. Slight periodontal disease, two caries and two abscesses were present. Five teeth had been lost ante-mortem.

#### Russia, Zaborinskiy

Coordinates: 56.23, 53.64

The Zaborinskiy burial ground (4 century AD) is located in the Sarapulskiy district, the Udmurt Republic, Russia on the right bank of the Kama River tributary Kyrykmas. The burial ground was excavated in 1983 and 2002 by N.L. Reshetnikov and V.A.Bernts. In total 30 burials arranged in clear rows were studied. The buried lay in 30–90 cm deep rectangular pits with their heads to North or North-East. In a half of burials funeral offering kits were found. No morphological studies have been done so far.
Both Dubrovskiy and Zaborinskiy burial grounds are attributed to the Mazunino culture (3–5 centuries AD) of the Pyanoborsk cultural community[579,580,594,595].

CGG_2_021458 (Zaboryinsky_IV-V_AD_1).

### Slovakia

#### Slovakia, Gerulata

Coordinates: 48.05, 17.14

The exact position of Gerulata is in Rusovce, a present- day suburb of Bratislava on the right bank of the Danube. The prosperity of Gerulata peaked in Traian- Hadrian´s period. The military detachment at Gerulata was formed by a cavalry troop of auxiliaries[596].

Archaeologists uncovered four burial grounds which date to the 1st–2nd and 3rd–4th centuries.

It appeared that the method of burial in these burial grounds was biritual i.e., skeletal and cremations[597].

CGG_2_021931 (Gerulata_Grave_58).

CGG_2_021934 (Gerulata_Grave_51).

CGG_2_021935 (Gerulata_Grave_21).

#### Slovakia, Trnava, Cifer-Pac

Coordinates: 48.30, 17.51

Cífer-Pác (South-western Slovakia) is an archaeological site that provides evidence of the coexistence of Avars and Slavs. Excavation in this area during the years 1971 to 1983 revealed a relatively large avar-slavic burial site dated to the 8th to 9th century[598]. The burial site consists of 119 skeletal graves with varying degrees of preservation. Based on the accompanying inventory of the graves, graves 1–38ab are dated to the 8th century and graves 39–119 from the 8th to the first half of the 9th century. No skeletal remains were preserved in 25 graves due to completely decomposed bones. The remaining 94 graves contained skeletal remains of 101 individuals, including 61 adults (20 males, 29 females and 12 individuals with undeterminable sex) and 40 juveniles[599]. No cremation burials occurred on this burial site and the equipment of graves was relatively diverse. Most of the graves contained only parts of ceramics and the objects of everyday use like needles, spindle whorls (female graves), iron knives and buckles (male graves), just a few individuals were buried with more valuable things or weapons: gilded bronze forging (CP33), necklace with metal and glass beads (CP22), two rings and lance (CP28), axe and lance (CP85), casted belt decorations with animal or plant ornaments (CP15, CP24, CP28, CP33). Furthermore, ten horse riders' graves (CP9, CP12, CP15, CP16, CP17, CP24, CP28, CP33, CP34 and CP109) were excavated. The walls of these graves were hardened with wooden posts and they contained skeletal remains of riders and probably their horses. In addition, the skull analysis of 22 individuals from this burial site revealed 6 cases of individuals with typical mongoloid cranial features (CP2, CP3, CP61, CP63, CP72, CP105) and 9 cases of mixomorphic individuals with both mongoloid and europoid cranial features including 3 individuals with higher percentage of mongoloid features (CP55, CP66, CP69).

In a previous study, we focused on the determination of mitochondrial haplogroups in this small Avar-Slavic population which lived in the 8th to 9th century, in order to determine the increased presence of possible "Avar" mitochondrial haplogroups (typical for the region of western and central Asia) in this mixed population, because the archaeological and anthropological research showed the incidence of Avar burial rites and also mongoloid cranial features of some individuals from this burial site[600].

Additionally a case report was published – dealing with a severe case of skeletal TBC found in one of the individuals from this burial site[601].

CGG_2_018917 (2/71).

CGG_2_018918 (3/71).

CGG_2_018920 (8/75).

CGG_2_018923 (40/79).

CGG_2_018924 (42/79).

CGG_2_018926 (102/83).

CGG_2_018927 (108/83).

### Spain

#### Spain, Madrid, Camino de las Yeseras

Coordinates: 40.40, -3.50

Jorge Vega

The site called “Camino de las Yeseras” is located on a raised terrace of the Henares River that shares its flow with the Jarama River, located near the confluence of both canals, in the Madrid district of San Fernando de Henares. The necropolis is located on the western edge of such a terrace.

It is a great archaeological site, with more than 22 hectares and more than 10.000 archaeological structures well identified and documented. Despite the site being occupied during the late Neolithic, the Campaniform period of the site needs to be also highlighted, with impressive internments, several defending pits, huts, different activities areas and great archaeological ground holes with symbolic material associated.

In 2006 within the western escarpment of the terrace which belongs to the floodplain of the Jarama river, a huge amount of visigothic tombs were detected and well identified. Emergency archaeological works were undertaken over 22 tombs that were going to be damaged as part of a building project. These graves are mainly simple pits with great rock slabs as the cover. Two of them present pit walls strengthened with great rock blocks creating a cist shape.

The tombs have been reused since some of them contain remains of several individuals placed surrounding the remains of the main individual. At least 52 individuals belonging to these 22 funerary structures have been well identified, with an axis of East-West. The good graves found are scarce, although they seem to provide a dating for 6th and 7th centuries A.D.

In conclusion after the anthropological analysis the current asserts are the following: the sex of the individuals has been well identified in 27 of the 52 cases, being 12 males and 15 females. Regarding the group of age, the majority of the individuals belong to the age-frame between 21–40 years old, which is the 49%, followed by the infancy group deceased at the age of 6 years old, which is the 21 % and followed by the infancy II group, with 7–12 years old, being the 13%.

It can be observed that the probability of death is rather high (19%) during the child age, it decreases during the youth and increases extremely during the adult age up to 81%. Only 12% of this population reached the mature age, keeping the probability of death as its previous age group. According to this data, only 2% of the population would exceed the age of 60 years old.

CGG_2_022031 (A118 E1 UE 1) is a child male.
CGG_2_022028 (A119 E3 UE 2) is a female adult.

#### Spain, Madrid, Estevillas Virgen de la Torre

Coordinates: 40.38, -3.56

Jorge Vega

The hispanic-visigothic necropolis of "Estevillas-Virgen de la Torre" was excavated between the years 2010–2011. His area was about 9.067 m² and 824 archaeological structures were excavated and well documented, of which 694 are tombs. Within them remains of up to 894 individuals have been excavated and studied. It is the biggest visigothic necropolis excavated in Spain.

The whole site spans over a little hill and its surroundings, with a central area where there are hardly any funerary pits due to the presence of the modern mining which most likely destroyed the structures that were located in such a zone. More specifically, these are two zones that rather lack structures related to the necropolis. Nearby to the south, on the west bank of the ancient greek called "Arroyo de los Migueles" we have excavated a settlement with 166 huts and about 2000 associated structures (kilns, pits, wine presses, cisterns…) between 2017 and 2023. Within the necropolis, the majority of the tombs are sorted in several streets very irregularly organized. There is more density of tombs in some specific locations, thus most of these have barely 1,5 metres of distance between them.

It is highly unlikely to find archaeological remains on the surface, although in very few cases small bones, high-fired indeterminate pottery or steel nails perhaps belonging to the coffins can be found. The majority of the tombs (70%) are E-W orientated, being NE-SW the axis of the rest. There are several types of tombs: simple pits, a pit with another previous pit and rock cists. Almost half of the interments have a lid or cover made of big rock slabs. Usually, the inside contains a single body, however some of them (33%) includes remains of several individuals, a clear sign of later reuse of the tombs. We have been able to document offerings and objects of personal good graves of the 20% of the tombs, composed mainly by belt buckles of different typology: with a scutiform base, lyriform etc. fibulae with an arch of hinge, trilaminar, some hoop-type earrings, necklaces of vitreous paste or amber beads, circular and rectangular section rings, some knives and many iron nails. Also, a small set of jars with trilobed rims and with a handle and several small bottles with one or two handles.

All these remains and archaeological material are pointing out that the necropolis was inhabited between the end of the 5th century A.D., with the arrival of the Visigoth as allies of the roman government, and the early 8th century with the beginning of the Muslim conquest.
The anthropological analysis shows a rather high child mortality (the 20% of this part of the populations) between 4 and 12 years old and a barely percentage of individuals framed in the mature age between 40 and 60 years old and old age from 60 years old on, being higher the percentage of men within the same age group. The causes for the high mortality of adult young women would be among the issues related to childbirth and the different diseases after it. The health state and the nutrition of these populations shows a high presence of injuries associated with an excess of certain muscle groups and tendons, almost always related to the action of certain physical activities, inherent to the development of work in the field, such as carrying heavy loads. There is also a high presence of spinal cord injuries, a homogeneous appearance in both men and women and is directly related to the mechanical overload of the spine when performing certain physical activities.

Traumatic injuries show an average rate of representativeness of the whole population. These are mainly fractures in long bones such as extremities and their distribution is greater within the adult male population.

Nutritional markers studied indicate a high incidence of caries, an average incidence of tartar, high dental wear, especially in molars, and an average rate of enamel hypoplasia that show specific cases of physiological stress during the life of an individual, which may be due to a period of starving during the childhood, or to some type of infectious disease.

CGG_2_022051 (EstevillasVirgen_de_la_Torre;_elem_124;_UE_03).
CGG_2_022052 (EstevillasVirgen_de_la_Torre;_elem_785;_UE_03).
CGG_2_022053 (EstevillasVirgen_de_la_Torre;_elem_785;_UE_01).

#### Spain, Madrid, Los Berrocales

Coordinates: 40.374901, -3.579209

Jorge Vega

This necropolis is located within the county of Madrid, 500 metres near the great necropolis of Estevillas, on the opposite bank of the creek called "Los Migueles". It is interesting that both necropolises are used at the same time during a span of time.
Within this area there was a Roman occupation from the middle of 1$^{st}$ century to the beginning of the 3$^{rd}$ century A.D., afterwards it was abandoned and then it was occupied again in the 4$^{th}$ century. Belonging to this latter period, has been excavated a necropolis with a total number of 42 human interments with a N-S orientated axis and they are arranged irregularly on disorganized streets.

There are 8 tombs without human remains, although there are 11 graves in which more than one single individual have been documented, most of the cases having two individuals. Human remains of up to six individuals have been excavated and documented in one tomb. The maximum number of individuals found is 55. The chronological frame for these interments is between $4^{th}$-$5^{th}$ centuries A.D.

Another necropolis from a later period (end of $5^{th}$ century – beginning of $6^{th}$ century) has been excavated. This latter shares its funerary space with the one mentioned before and it get to change even some of the other tombs' space context. This second necropolis has a total of 64 funerary pits, with an orientation axis W-E in the majority of the cases and an inner arrangement even more chaotic. The population buried seems to be of 82 individuals: 21 kids (up to 12 years old), 14 males and 13 females, being the rest unidentified. The maximum life expectancy age for this site is between 21 and 40 years old.

Scarcely remains of grave goods have been found and documented in the interments. Some metallic ornaments such as belt buckles, earrings, rings, keys or brooches have been found. Moreover, several unidentified fragments of iron and nails have been documented, which shows the existence of wooden coffins.

Close to these two necropolises and on the same river bank of the creek has been excavated another necropolis with 118 Muslim tombs in it, chronologically framed between the $8^{th}$ and the $10^{th}$ centuries. This cemetery is the elongation of the interments of the Estevillas necropolis, which was abandoned during the arrival of the Muslim population to the Iberian Peninsula since 711 A.D.

#### Spain, Toledo, Monterrey-Estiviel

Coordinates: 37.60, -0.97

Jorge Vega

The necropolis was identified during the building works for the new Wastewater Treatment Plant in Toledo, which was excavated in 2012. The site is located about 800 metres from the current course of the Tagus River on its right bank and 8 km downstream from the city of Toledo, the capital of the Visigothic kingdom since the middle of the 6th century.

The complex comprises 154 tombs being all inhumation burials, of which 137 contain human remains, 17 are looted (with remains of construction materials and even highly fragmented and deteriorated remains of individuals). Archaeological data points to the use of the site for a time span of c. four centuries (4st - 7th. centuries CE). The oldest tombs date from the 4th century BC. C., 104 of them contain Late Roman materials, present a N-S orientation and they would have been in use during the 5th century AD.

In the second half of the 6th century, coinciding with the establishment of Toledo as the capital of the Visigothic kingdom, a small necropolis with 33 E-W oriented tombs was established within this same space, sometimes even breaking previous tombs. We have identified the presence of 44 individuals and it seems to be occupied for a large part of the 7th century. In general, they are graves with a single individual buried in it, although with the presence of reuses with up to four individuals in the same grave. It is a cemetery associated with a rural establishment of an agricultural nature located nearby. Anthropological data show that the individuals buried are men, women and children. We have identified the remains of ten infantile individuals, highlighting a greater female presence (ten individuals of which half correspond to young women). The reasons for this high mortality in young women should be sought in the unequal incidence of diseases, work risks and, above all, infections related to childbirth.

CGG_2_022041 (Monterrey-Estiviel_El_93,_UE_02).

# UK

#### UK, Gloucestershire, Randwick Long Barrow

Coordinates: 51.76, -2.25

Alan K. Outram

Randwick Long Barrow lies atop Randwick Hill (Witts 1883) to the NW of Randwick village in Gloucestershire, England. The monument has not been investigated in detail in the modern era, but is a Cotswold-Severn type Neolithic long barrow displaying apparent Roman re-use[602]. The site was excavated in 1883 by Witts[603], whose limited account describes the finding of the remains of several human skeletons both outside the southern walls and within the chambers, alongside some animal bones. The chamber also yielded both flint flakes and Roman pottery[603], evidencing both prehistoric and Roman era activity. Teeth from two human mandibles were analysed in this study, from old collections originating from Gloucester Museum, however, the precise location of these within the barrow, in relation to these Victorian excavations, is not known. From a contextual standpoint the remains could have been either Neolithic or Roman, but the radiocarbon dates undertaken for this study concord with the latter.

CGG_2_020723 (Randwick_Long_Barrow,_Individual_1,_A3081).

CGG_2_020724 (Randwick_Long_Barrow,_Individual_2,_A3081).

#### UK, Orkney, Mine Howe

Coordinates: 58.73, -2.93

Nick Card and Jane Downes

Archaeological investigation at Mine Howe first took place in 1946, when a remarkable stone-built underground 'staircase' was discovered during local exploration of the site's central (natural) mound. In 1999, geophysical survey across the Mine Howe mound and its surrounds, followed by four seasons of targeted excavation, revealed a striking ritual complex dating to the mid to late Iron Age. The Mine Howe excavation established two key loci to the complex[288,604]:

1. a roughly circular enclosure surrounding the mound and its central subterranean staircase structure, the creation of which substantially modified the appearance of the mound. The enclosure featured a monumentalised revetted entrance on its western side, and excavation revealed a complex history of infilling, recutting and remodelling of the enclosure ditch throughout its history. Both the composition (artefacts, and extensive depositions of animal bone indicative of feasting) and dating (indicating deposition/re-deposition of relict materials) of the midden infills of the ditch point to ritual activity and a performative focus to the enclosure's use and the ditch infill events. The enclosure ditch was constructed in the second century BC at the earliest, but possibly as late as 50 cal BC, and final infill activity took place within the third to fourth centuries AD.

2. Approximately 8 metres west of the southern (north-facing) enclosure entrance terminal, a roughly circular stone-built structure c.4.5–4.75m in diameter was identified, in addition to several earlier, more ephemeral structures. Internal features and associated material identified

this structure as primarily a workshop for the production of both ferrous and non-ferrous metalwork. Construction and early use of the workshop structure took place 60 cal BC – cal AD5, followed by an established phase of intensive use 40 cal BC– cal AD 25 and 20 cal BC – cal AD50, a phase of decommissioning and closure, cessation of of metalworking activity and the associated burial of two individuals (CGG_2_018914 and CGG_2_018915, see below) dated to cal AD20–100, a reactivation of workshop activity after a short hiatus (cal AD 55–130) and a final accumulation of midden-enriched soils, accompanied by sporadic activity, from cal AD 70–140.

CGG_2_018914: A sub-oval, N-W aligned grave cut containing the extended, supine inhumation of a young woman was inserted against the north-west workshop wall, inside the structure, during the period when the formal use of the workshop came to an end (Burial C827, within grave cut context C897). It is possible that this may represent a specific decommissioning event related to this transition in use. The grave was capped with flagstones. The body was placed with head to the south, with arms beside the body and legs straight. Grave goods included two spiral toe rings and an unusual worked antler object on her chest. One ring had been placed on the middle toe of the left foot and another on the second toe of the right, perhaps simply ornamental, perhaps forming part of sandal-type footwear. Some of the bones – particularly the skull – were slightly crushed by the stone capping, but the inhumation was complete and fully-articulated. The grave had two backfilled deposits, the lower deposit of which contained potsherds, worked stones, whelk shells, crucible fragments, copper alloy sheet fragments, and large quantities of charred plant remains, including several hundred carbonised barley seeds, weed seeds, burnt peat and industrial waste. This atypical grave fill signature may represent smashing and deposition of pots by mourners during burial, with shells and grain representing food offerings. Crucibles and industrial waste may allude to the metal-working activities undertaken when the workshop was active. Osteological analysis established that the woman was 15/16–18/20 when she died and c. 5 foot 2 inches tall (possibly not fully grown). Schmorl's nodes noted on several vertebrae indicated significant physical activity including the carrying of heavy loads. Other pathologies included the presence of spina bifida, but only in its minor form which would not have affected her physically. There was also evidence that she suffered from anaemia, perhaps via poor diet or parasitic infection, genetic predisposition, or a combination of these. Severe iron deficiency may have been a contributing factor in her death at such a young age. Separate radiocarbon dates from two ribs (GU-15640 and GU-15641) returned consistent determinations which indicate that she died between cal AD15 and cal AD100 (Table S7.36).

CGG_2_018915: A second grave (Context C1863) was located 4.6m to the north-west of the workshop. A fairly shallow, irregular pit (1.45m x 0.54m by 0.27m deep) contained the flexed skeleton of an aged adult male placed on his left side, aligned NE – SW with head to the south-west, and with legs and feet together (Burial C1861). The body was carefully arranged, but not fitted within the pit - the head on the slope of the cut, the legs bent perpendicular to the body, and the toes of the left foot bent to fit within the pit but still protruding. Almost too small for the individual within it, the pit appears hastily dug, the grave filled seemingly midden-derived, and there were no grave goods. Several large flagstones capped the grave, and had badly crushed the skull – it is possible that they were rapidly flung rather than carefully placed. Initially placed at 25–35 years, this individual was later re-aged to late 40s or perhaps even early 50s, due to the presence of ossified thyroid cartilage, and degenerative joint disease. The teeth were not significantly worn, suggesting that he may have enjoyed a relatively non-abrasive, perhaps even high-status diet. He was c. 5 foot 5 inches tall and probably right-handed. Like the young female buried in the workshop, a very physical lifestyle was indicated with well-developed upper arm muscles and several joint/bone diseases including spinal osteoarthritis, Schmorl's nodes and spondyloarthropy. He also likely suffered from Paget's disease, a thickening but weakening of the bone. Together this suggests significant mechanical stress, a result of constant bending, lifting and carrying heavy weights on his back.

Several traumatic unhealed wounds suggested that this man met a violent end: puncture wounds to the scapula, cut marks on the lower jaw, scapula, ribs and left metacarpal were likely caused by two or more different weapons: a projectile and a slashing blade. Chop marks also imply use of an axe. The penetrating wounds were inflicted to the back while the chopping/slashing wounds were to both, around the neck, predominantly to the left side of the body, but also more widely across vertebrae, clavicle, sternum and mandible. It seems likely that he was fired at from behind and, once down, stabbed and slashed repeatedly while on the ground - probably still moving, as injuries were inflicted to both back and front. This violent death may simply indicate a frenzied attack, but the subsequent careful burial, facing away from the workshop, may also suggest a ritualised killing, and it may be significant that the individual was possibly marked out both by social status and by pathological conditions that could have been seen as ‘marks’ with supernatural significance. Two modelled radiocarbon dates taken from the rib bones (GU-15638, GU-15639) were contemporary with those of burial C897, suggesting that this man also died between *cal AD20* and *cal AD90* (Table S7.36).

Isotopic analysis for both individuals comprised bone (carbon and nitrogen) and tooth enamel (strontium, lead and oxygen). Results were remarkably similar in all five isotope systems, suggesting similar geographic origins and childhood diets. C and N values suggest a diet rich in animal protein from terrestrial and a moderate marine input, together with C3 plant consumption. There was no evidence from lead, strontium or oxygen to indicate that either individual originated from somewhere other than Orkney. The following Table S7.2.35: provides the list of samples studied from Orkney, Mine Howe, UK and their dating.

| CGG No | Site context | Sample code (GU) | Calibrated age range (95% confidence) | Modelled age range (95% confidence) |
|---|---|---|---|---|
| CGG_2_018914 | 897 | 15640 | 60BC–AD120 | AD15–90 |
| | | 15641 | 40BC–AD130 | AD20–100 |
| CGG_2_018915 | 1861 | 15639 | 50BC–AD120 | AD20–95 |
| | | 15638 | 50BC–AD130 | AD20–95 |

**Table S7.2.35:** List of samples studied from Orkney, Mine Howe, UK and their dating.

285. Aikio, A. An essay on Saami ethnolinguistic prehistory. in *A Linguistic Map of Prehistoric Northern Europe* (ed. Grünthal R & Kallio) vol. 226 63–117 (Société Finno-Ougrienne, Helsinki:

Suomalais-Ugrilainen Seura, 2012).

286. Saag, L. *et al.* The arrival of Siberian ancestry connecting the Eastern Baltic to Uralic speakers further east. *Curr. Biol.* **29**, 1701–1711.e16 (2019).

287. Koch, J. T. & Ling, J. ‘from the ends of the earth’: A cross-disciplinary approach to long-distance contact in bronze age Atlantic Europe. in *The Indo-European Puzzle Revisited* 157–171 (Cambridge University Press, 2023). doi:10.1017/9781009261753.016.

288. Patterson, N. *et al.* Large-scale migration into Britain during the Middle to Late Bronze Age. *Nature* **601**, 588–594 (2022).

289. Lomborg, E. Die Flintdolche Dänemarks. *Köpenhamn* (1973).

290. Prieto-MartÍnez, M. P. Bell beaker communities in thy: The first bronze age society in Denmark. *Nor. Archaeol. Rev.* **41**, 115–158 (2008).

291. Prescott, C. Language and Prehistory of the Indo- European Peoples A Cross-Disciplinary Perspective. Museum Tusculanums Forlag. 229–249 (2017).

292. Forssander, J.-E. *Die Schwedische Bootaxtkultur Und Ihre Kontinentaleuropäischen Voraussetzungen*. (Lund, 1933).

293. Hinsch, E. *Yngre Steinalders Stridsøkskulturer I Norge*. (John Grieg, Bergen, 1956).

294. Price, D. *Ancient Scandinavia: An Archaeological History from the First Humans to the Vikings.* (Oxford University Press, 2015).

295. Childe, V. G. *The Aryans: A Study of Indo-European Origins. (K. Paul, Trench, Trubner & Company, Limited, 1926). 8. Güntert, H. Der Ursprung Der Germanen*. (C. Winter, Heidelberg, 1934).

296. Glob, P. *Studier over Den Jyske Enkeltgravskultur (Aarbøger for Nordisk Oldkyndighed Og Historie)*. (København, 1944).

297. Kristiansen, K. Prehistoric migrations - the case of the Single Grave Culture and Corded Ware Culture. *Journal of Danish Archaeology* **8**, 211–225 (1989).

298. Iversen, R. & Kroonen, G. Talking neolithic: Linguistic and archaeological perspectives on how Indo-European was implemented in southern Scandinavia. *Am. J. Archaeol.* **121**, 511–525 (2017).

299. Heather, P. J. *Empires and Barbarians*. (Macmillan, 2009).

300. Drinkwater, J. F. *The Alamanni and Rome 213-496: (Caracalla to Clovis)*. (OUP Oxford, 2007).

301. Kelly, C. *The End of Empire: Attila the Hun & the Fall of Rome*. (W. W. Norton, 2009).

302. Wickham, C. *The Inheritance of Rome: Illuminating the Dark Ages 400-1000*. (Penguin, 2009).

303. Nielsen, H. F. *The Germanic Languages: Origins and Early Dialectal Interrelations*. (University of Alabama Press, 1989).

304. Ringe, D. A. *From Proto-Indo-European to Proto-Germanic*. (Oxford University Press, 2017).

305. Salmons, J. The evolution of Germanic. in *Handbook of Comparative and Historical Indo-European Linguistics.* vol. 2 1002.1027 (DE Gruyter Mouton, 2017).

306. Agee, J. R. Using Historical Glottometry to Subgroup the Early Germanic Languages. *Journal of Germanic Linguistics* **33**, 319–357 (2021).

307. Hansen, B. S. S., Kroonen, G. J. & Olander, T. Germanic. in *The Indo-European Language Family* 152–172 (Cambridge University Press, 2022). doi:10.1017/9781108758666.010.

308. Heather, P. & Matthews, J. The Goths in the fourth century. (1991) doi:10.3828/978-0-85323-426-5.

309. Wolfram, H. *History of the Goths*. (University of California Press, 1988).

310. Jordanes & Mierow, C. C. *The Origin and Deeds of the Goths: In English Version*. (Princeton University Press, 1908).

311. Svennung, J. *Jordanes und Scandia: Kritisch-exegetische Studien*. (Almqvist & Wiksell, 1967).

312. Hedeager, L. *Iron Age Myth and Materiality: An Archaeology of Scandinavia AD 400-1000*. (Routledge, 2011).

313. Steuer, H. *„Germanen“ aus Sicht der Archäologie: Neue Thesen zu einem alten Thema*. (Walter de Gruyter GmbH & Co KG, 2021). doi:10.1515/9783110702675.

314. Cieslinski, A. Grabhügel der Wielbark-Kultur – Gemeinschafts-, Familien- und Elitengrabmäler. Einführung in die Problematik. Archäologie in Schleswig. Sonderband. in *Internationale Sachsensymposion 2010* 171–187 (Haderslev, Danmark: Neumünster, 2011).

315. Wołągiewicz, R. Die Goten im Bereich der Wielbark-Kultur. In: J. Kmieciński (Hrsg.). in *Peregrinatio Gothica* (ed. Kmieciński, J.) vol. 7 63–98 (Katedra Archeologii Uniwersytetu

Łódzkiego, Łódź, 1986).

316. Foulke, W. D., Paul, W. D. & Edel, A. *History of the Langobards*. (Creative Media Partners, LLC, 2018).

317. Menghin, W. *Die Langobarden: Archäologie und Geschichte*. (Panorama Wsb, 1985).

324. Neumann, H. *Olgerdiget: et bidrag til Danmarks tidligste historie*. (Haderslev Museum, Haderslev, 1982).

325. Christensen, L. Nye undersøgelser af Olgerdiget–en langvold fra jernalderen. *Sønderjysk Månedsskrift* (2020).

326. Ilkjær, J. Danish War Booty Sacrifices. in *The Spoils of Victory: The North in the Shadow of the Roman Empire* (eds. Jørgensen, L., Storgaard, B. & Thomsen, L. G.) 44–65 (Danish National Museum, Copenhagen, 2003).

327. Engelhardt, C. *Thorsbjerg mosefund: beskrivelse af de oldsager, som i aarene 1858-61 ere udgravede af Thorsbjerg mose ved Sønder-Brarup i Angel; et samlet fund, henhørende til den*

*aeldre jernalder og bevaret i den kongelige samling af nordiske oldsager i Flensborg*. (Kinsey und Thiele, 1863).

328. Raddatz, K. *Der Thorsberger Moorfund. Gürtelteile Und Körperschmuck*. vol. 13 (Offa-Bücher, Neumünster, 1957).
329. Blankenfeldt, R. *Das Thorsberger Moor: Die Persönlichen Ausrüstungen*. vols VI-XI (Verein zur Förderung des Archäologischen Landesmuseums, 2014).
330. Rau, A. *Nydam Mose: Die personengebundenen Gegenstände : Grabungen 1989 - 1999*. vol. 72 (Jysk Arkæologisk Selskab, Højbjerg, 2010).
331. Engelhardt, C. *Nydam mosefund, 1859-1863*. (G.E.C. Gad, 1865).
332. Jørgensen, A. N., Andersen, H. C. H. & Andersen, H. C. H. *Ejsbøl Mose: die Kriegsbeuteopfer im Moor von Ejsbøl aus dem späten 1. Jh. v. Chr. bis zum frühen 5. Jh. n. Chr*. (Jysk Arkaeologisk Selskab, Højbjerg, 2014).
333. Ørsnes, M. *Ejsbøl I. Waffenopferfunde Des 4.-5. Jahrh. Nach Chr*. vol. 11 (Det Kongelige Nordiske Oldskriftselskab. København, 1988).
334. Ilkjær, J. *Illerup Ådal 1. Die Lanzen Und Speere*. vol. XXV:1 (Jysk Arkæologisk Selskab, Højbjerg, 1990).
335. Ilkjær, J. *Illerup Ådal 3. Die Gürtel. Bestandteile Und Zubehör*. vol. XXV: 3 (Jysk Arkæologisk Selskab, Højbjerg, 1993).
336. Ilkjær, J. *Illerup Ådal 2. Die Lanzen Und Speere*. vol. XXV: 2 (Jysk Arkæologisk Selskab, Højbjerg, 1990).
337. Andersen, K. H. Da danerne blev danske. (Aarhus University, 2016).
338. Parthey, G, Pinder, M. *Ravennatis Anonymi Cosmographia et Guidonis Geographica*. (Fb&c Limited, 2017).
339. Sahlin, B. *Die Altgermanische Thierornamentik*. (Stockholm and Berlin: Asher, 1904).
340. Imer, L. M. & Vasshus, K. S. K. Lost in transition. The runic bracteate from the Vindelev hoard. *NOWELE* **6**, 60–99 (2023).
341. Hedeager, L. Poetry and picturing in deep historical time. in *Re-imagining Periphery* (eds. Hillerdal, C. & Ilves, K.) 107–116 (Oxbow Books, 2020).

342. Fabech C, N. U. *The Sösdala Horsemen and the Equestrian Elite of Fifth Century Europé*. (Aarhus University Press, Aarhus, 2017).

343. van Dijk, E. *et al.* Climatic and societal impacts in Scandinavia following the 536 and 540 CE volcanic double event. *Clim. Past* **19**, 357–398 (2023).

344. Looijenga, T. *Texts and Contexts of the Oldest Runic Inscriptions*. (Brill, 2003).

345. Birkmann, T. A survey of ancient Nordic sources. in *The Nordic Languages, Part 1* (De Gruyter, Berlin, Boston, 2017).

346. Amory, P. *People and Identity in Ostrogothic Italy, 489-554*. (Cambridge University Press, 2003).

347. Pohl, W. Memory, Identity and Power in Lombard Italy. in *The Uses of the Past in the Early Middle Ages* (ed. Hen Y Innes) (Cambridge University Press, 2000).

359. Price, N. Climate change and the Fimbulwinter. in *Oxford Handbook of Scandinavian Archaeology* (Oxfor University Press).

360. Holst, M. Farmsteads and communities. in *Oxford Handbook of Scandinavian Archaeology* (Oxford University Press, Oxford, in press).

361. McNeill, W. *Plagues and Peoples*. (Knopf Doubleday Publishing Group, 1998).

362. Loftsgarden, K. & Iversen, F. The impact of the 6th century crisis – exploring burials as a proxy for population dynamics in iron age Scandinavia. *Nor. Archaeol. Rev.* 1–21 (2024) doi:10.1080/00293652.2024.2412018.

363. De Bellis, P. Book I-VIII. in *Procopius. De Bellis* (Harvard University Press, 1918).

364. Gertz, M. C. I. *Scriptores Minores Historiæ Danicæ Medii ævi Ex Codicibus Denuo*. (København, I. Kommission hos G. E. C. Gad, 1917).

365. Saxo, G. *The History of the Dane*. (D.S.Brewer, Cambridge, 1998).

366. Storlason, S. *Heimskringla*. (Dover Publications, Inc., New York, 1990).

367. Ottosson, K. 90. Old Nordic: A definition and delimitation of the period. in *The Nordic Languages, Part 1* (De Gruyter, Berlin, Boston, 2017). doi:10.1515/9783110197051-090.

368. Venås, K. 'Dansk tunge' og Håkon Melberg. *Syn Segn* **106**, 66–79 (2000).

369. Schulte, M. & Bandle, O. *The Various Old Nordic Dialects, Their Systems and Their Typology: A General Survey of the Classical Old Nordic of the High Middle Ages. in The Nordic Languages. An International Handbook of the History of the North Germanic Languages*. vol. 100 871–882 (De Gruyter Mouton, 2008).

370. Jesch, J. Speaking like a Viking: Language and cultural interaction in the Irish Sea region. in *Search of Vikings: Interdisciplinary Approaches to the Scandinavian Heritage of* (eds. Harding, S. E., Griffiths, D. & Royles, E.) 50–61 (CRC Press, 2015).
371. Dahl, Ö. The origin of the Scandinavian languages. in *Studies in Language Companion Series* ccxv–ccxxxv (John Benjamins Publishing Company, Amsterdam, 2001). doi:10.1075/slcs.54.11dah.
372. Riisøy, I. A. Danish legal procedure and a common Scandinavian law Riisøy. in *Viking Encounters* 220–231 (2020).
373. Aikio, A. *The Saami Loanwords in Finnish and Karelian*. (2009).
374. Hofmanová, Z. *et al.* Early farmers from across Europe directly descended from Neolithic Aegeans. *Proc. Natl. Acad. Sci. U. S. A.* **113**, 6886–6891 (2016).
375. Lawson, D. J., Hellenthal, G., Myers, S. & Falush, D. Inference of population structure using dense haplotype data. *PLoS Genet.* **8**, e1002453 (2012).
376. Crumlin-Pedersen, O. Bådgrave og gravbåde. in *Slusegårdgravpladsen III. Gravformer og gravskikke. Bådgravene* (eds. Andersen, S. H., Lind, B. & Crumlin-Pedersen, O.) vol. 14 93–263 (Jysk Arkæologisk Selskab, Højbjerg, 1991).
377. Lind, B. Gravformer og gravskikke. in *Slusegårdgravpladsen III. Gravforme og gravskikke. Bådgravene* (eds. Andersen, S. H., Lind, B. & Crumlin-Pedersen, O.) vol. 14 13–91 (Jysk Arkæologisk Selskab, Højbjerg, 1991).
378. Rein Seehusen, C. Baltic connections. Some remarks about studies of boatgraves from Roman Iron Age. Finds from the Slusegård and Weklice cemeteries. *Nordiske Fortidsminder* **C**, 287–313 (2010).
379. Rasmussen, B. *Slusegårdgravpladsen: Bornholm Fra 1. årh: F. Til 5. år H. E. v. T.* (Jysk Arkæologisk Selskab, Højbjerg, 2010).
380. Sellevold, B. Hansen, U.L., & Jørgensen, J.B. *Iron Age Man in Denmark.Prehistoric Man in Denmark*. vol. III (København: Nationalmuseet, 1984).
381. Klindt-Jensen, O. *Slusegårdgravpladsen I: Bornholm Fra 1. Årh. F. Til 5. årh. E.v.t.* vol. 14 (Jysk Arkæologisk Selskab, Højbjerg, 1978).

382. Beck, M. R. Old traditions meeting new ideas. in *Life and afterlife in the Nordic Bronze Age.* (ed. A., Svensson, A. & Apel, J.) 141–157 (Lund University, 2019).

383. Løvschal, M., Iversen, R. B. & Holst, M. K. De dræbte krigere i Alken Enge: Efterkrigsritualer i ældre jernalder. (2019).

384. Stenbak, M. K. Illemose. in *Fynske Minder. Årbog for Odense Bys Museer* (ed. Ganshorn, J.) 149–180 (Odense Bys Museer, Odense, 1994).

385. Albrectsen, E. Fynske jernaldergrave. 3, Yngre romersk jernalder. (1968).

386. Bonde, J. Et arkæologisk gensyn med Hedegaardsvej. *Vends* **2011**, 21–28 (2011).

387. Nielsen, B. H., Christensen, T. & Frei, K. M. New insights from forgotten bog bodies: The potential of bog skeletons for investigating the phenomenon of deposition of human remains in bogs during prehistory. *J. Archaeol. Sci.* **120**, 105166 (2020).

388. Van der Sluis, L. G. Investigating Paleodietary Changes from the Mesolithic to the Viking Age in the Limfjord Area in Northern Denmark. (Queen's University Belfast. Faculty of Engineering and Physical Sciences, 2017).

389. van Beek, R., Quik, C., Bergerbrant, S., Huisman, F. & Kama, P. Bogs, bones and bodies: the deposition of human remains in northern European mires (9000 BC–AD 1900). *Antiquity* **97**, 120–140 (2023).

390. Norling-Christensen, H. *Katalog over Ældre Romersk Jærnalders Grave I Århus*. (Amt. Nordiske Fortidsminder IV. Bind 2 Hefte.København: Det KGL. Nordiske Oldskriftselskab, 1954).

391. Brinch Madsen, H. En Nordjysk kvindegrav fra omkring 400 e.Kr. *Hikuin* **2**, 123–130 (1975).

392. Christiansen, T. T. Sdr. Tranders – en gravplads fra ældre germansk jernalder. in 14–17 (Årsberetning for Nordjyllands Historiske Museum 2005, 2006).

393. Holst, M. K. *et al.* Direct evidence of a large Northern European Roman period martial event and postbattle corpse manipulation. *Proc. Natl. Acad. Sci. U. S. A.* **115**, 5920–5925 (2018).

394. Christiansen, T. T. "Gudernes hjem" – helligområde og gravplads med udsigt. *SKALK* **1**, 3–9 (2021).

395. Wåhlin, S. *Six Human Skulls in a Bog, Svennum – a 1st Century AD Sacrificial Bog" in In the Darkest of Days. Exploring Human Sacrifice and Value in Southern Scandinavian Prehistory.*

(Oxbow books, 2024).

396. Mandrup, P., Lundby, P. & Laursen, K. Udgravningsberetning FHM 5478, Lauritshøj II, Skødstrup sogn, Øster Lisbjerg Herred, Århus kommune, Randers Amt Stednr. 14.11.08, sbnr. 104 Matrikel nr. 1ak, Skødstrup By, Skødstrup,. *KUAS journal, 2014-7.24.02.02/FHM-0018.Moesgaard Museum* (2017).

397. {Kanstrup, M., Heinemeier, J. & Olsen, J. Kulstof 14-resulateter. in *De dræbte krigere i Alken Enge. Efterkrigsritualer i ældre jernalder* (eds. {Løvschal, M., Iversen, R. B. & Kähler Holst, M.) vol. 108 193–208 (Jysk Arkæologisk Selskab, Højbjerg, 2019).

398. Møllerup, L. Det humane skeletmateriale. in *De dræbte krigere i Alken Enge. Efterkrigsritualer i ældre jernalder* (eds. {Løvschal, M., Iversen, R. B. & Kähler Holst, M.) vol. 108 157–184 (Jysk Arkæologisk Selskab, Højbjerg, 2019).

399. Ilkjær, J. Danish war booty sacrifices. in *The Spoils of Victory. The North in the shadow of the Roman Empire* (eds. Jørgensen, L., Storgaard, B., Thomsen, G. & L) 44–65 (National Museum, Copenhagen, 2003).

400. Skaarup, J., Nyegaard, G. & Aaris-Sørensen, K. *Yngre Stenalder På øerne Syd for Fyn*. (Langelands Museum, Rudkøbing Denmark, 1985).

401. Broholm, H.-C. *Danmarks Bronzealder*. vol. III (Arnold Busch: Copenhagen. Fund og Fortidsminder, 090307-26, 1946).

402. Eggers, H. J. *Der römische Import im freien Germanien*. (Hamburgisches Museum für Völkerkunde und Vorgeschichte, 1951).

403. Frei, K. M. & Klingenberg, S. Revisiting the Roman Iron Age Hoby chieftain burial after 100 years of its discovery - adding the strontium isotopic perspective. *Dan. J. Archaeol.* **10**, (2021).

404. Nordman, C. A. *et al. Nordiske Fortidsminder*. (Kgl. nordiske Oldskriftselskab, København, 1923).

405. Margaryan, A. *et al.* Population genomics of the Viking world. *Nature* **585**, 390–396 (2020).

406. Grøn, O., Hedeager, K. A. & Bennike, P. *Vikingetidsgravpladser På Langeland*. (Langelands Museum, Rudkøbing, Denmark, 1984).

407. Liversage, D. *Material and Interpretation: The Archaeology of Sjælland in the Early Roman Iron*

*Age*. (National Museum of Denmark, 1980).
408. Jørkov, M. *Drinking with the Rich and Dining with the Poor in Roman Iron Age Denmark. A Methodological Study Based on Stable Isotope Analysis*. (2007).
409. Jørkov, M. L. S., Jørgensen, L. & Lynnerup, N. Uniform diet in a diverse society. Revealing new dietary evidence of the Danish Roman Iron Age based on stable isotope analysis. *Am. J. Phys. Anthropol.* **143**, 523–533 (2010).
410. Melchior, L., Lynnerup, N., Siegismund, H. R., Kivisild, T. & Dissing, J. Genetic diversity among ancient Nordic populations. *PLoS One* **5**, e11898 (2010).
411. Boye, L. *Excavation Report SØL 964 Engbjerg, Høje Tåstrup Sogn, Smørum Herred, Københavns Amt., Stednr. 020207-84*. (1988).
412. Ørsnes-Christensen, M. Kyndbygravpladsen: Rige fund fra en fundfattig tid. *Fra Nationalmuseets Arbejdsmark 1955* 74–81 (1955).
413. Ravn, L. G. Strandgrave ved Bilidt. in *Simblegård-Trelleborg. Danske gravfund fra førromersk jernalder til vikingetid* (ed. Jørgensen, L.) (Arkæologiske Skrifter 3, København, 1989).
414. Kjær, K. I. Gravpladserne langs Fjordlandsvej. in *Kronprinsesse Marys Bro og arkæologien* (ed. Kastholm, O. T.) 104–125 (Roskilde, 2022).
415. Bennike, P. *Palaeopathology of Danish Skeletons*. (Akademisk Forlag, København, 1981).
416. Jessen & Olsen, J. Deviant Vikings – ritual executions at lake Tissø. in *In the Darkest of Days: Exploring human sacrifice and value in Southern Scandinavian prehistory* 101–108 (Oxford: Oxbow Books, 2023).
417. Lynnerup, N. & Skaarup. Den halshuggede fra Bakkendrup: Hvordan så han ud? *Årets gang 2007 – Kalundborg Museum* 42–43 (2007).
418. Koch, E. *Neolithic Bog Pots from Zealand, Møn, Lolland and Falster*. (Det Kongelige Nordiske Oldskriftselskab, 1998).
419. Vedbæk, C. L. *Fundberetning 4319, Karlstrup*. (1987).
420. Aner, E. & Kersten, K. *Die Funde Der älteren Bronzeziet Des Nordischen Kreises in Dänemark, Schleswig-Holstein Und Niedersachsen: Bornholms, Maribo, Odense Und Svendborg Amter*. (Neumünster: Karl Wachholz Verlag, 1977).

421. Ebbesen, K. Späteneolithische Schmuckmode. *Acta Archäologica* **66**, 219–279 (1995).
422. Allentoft, M. E. *et al.* Population genomics of Bronze Age Eurasia. *Nature* **522**, 167–172 (2015).
423. Balslev Jørgensen, J. *Rapport over Osteological Analysers Karlstrup*. (1965).
424. Ebbesen, K. Danske hellekister fra stenalderen. *Aarbøger for Nordisk Oldkyndighed og Historie.* 7–62 (2007).
425. Bennike, P. Facts or myths? A re-evaluation of cases of diagnosed tuberculosis in the past in Denmark. in *Tuberculosis. Past and present* (ed. G. Palfi, O. Dutour, J. Deák & I. Hutas) 511–518 (Golden Book Publ. TB Foundation, Hungary, 1999).
426. Johannsen, J. W. En senneolitisk grav på Tørslev Hage. in *Kulturen på spil. Årsskrift for museumsorganisationen ROMU 2020* (eds. Nielsen, J. & Struve, E. W.) 38–53 (Museumsforlaget ROMU, Roskilde, 2021).
427. Hansen, B. B. Kvinden på Maglebjerg: En rig grav fra yngre romersk jernalder ved Næstved. in *Aarbøger for nordisk Oldkyndighed og Historie 2008* 123–194 (Det Kongelige Nordiske Oldskriftselskab, København, 2016).
428. Ethelberg, P. *Skovgårde: Ein Bestattungsplatz Mit Reichen Frauengräber Des 3.Jhs. N. Chr. Auf Seeland.* (Det Kongelige Nordiske Oldskriftselskab: Copenhagen, 2000).
429. Melchior, L., Gilbert, M. T. P., Kivisild, T., Lynnerup, N. & Dissing, J. Rare mtDNA haplogroups and genetic differences in rich and poor Danish Iron-Age villages. *Am. J. Phys. Anthropol.* **135**, 206–215 (2008).
430. Jørgensen, L. *et al.* Førkristne kultpladser – ritualer og tro i yngre jernalder og vikingetid. in *Nationalmuseets Arbejdsmark 2014* (eds. Madsen, P. K. & Wass, I.) 186–199 (Nationalmuseet, København, 2014).
431. Kjær, H. Et mærkeligt arkæologisk-antropologisk fund fra stenalderen. *Aarbøger for Nordisk Oldkyndighed og Historie:* 58–72 (1912).
432. Helle, T. A. V. Violence and warfare before the Nordic Bronze Age. in *Oxford Handbook of Scandinavian Archaeology. Age farmers, herders, hunters and gatherers: The Funnel Beaker Culture, the Single Grave/Battle Axe Culture and the Pitted Ware Culture (4000–2400 BC)* (eds. Hedeager, L., Prescott, C. & Kristiansen, K.) (Oxford University Press, 2023).

433. Kastholm, O. T. & Ulriksen, J. Burials of Ritual Specialists? Case studies of the Graves from Trekroner-Grydehøj and Gerdrup, Sjælland. in *The Norse Sorceress. Mind and Materiality in the Viking World* (ed. L. Gardeła, S. B. &. P. P.) 381–400 (Oxford, 2023).

434. Ulriksen, J. A Völva's Grave at Roskilde, Denmark? *Offa* **71/72**, 229–240 (2018).

435. Ulriksen, J. Vikingetidens gravskik i Danmark. Spor af begravelsesritualer i jordfæstegrave. *Kuml* 161–245 (2011).

436. Kastholm, O.T., K.I. Kjær, N.M.K. Mortensen & P.Ø. Sørensen. Døden i Vindinge – et smugkig på to nye gravpladser. in *Årsskrift for Roskilde Museum 2018* 166–183 (Roskildle Museum, Roskilde, 2018).

437. Bönnelycke, A. *Rapport Om de Arkæologiske Undersøgelser Ved Bredebjerggård 8,TAK 1180.* (2008).

438. Jacobsen, M. & Pallesen. *TAK1621 Sejlsbjerg, Høje Tåstrup Sogn, Smørum Herred, Tidl. København Amt. Sted Nr. 02.02.07. Sb.nr. 145*. (2019).

439. Gotfredsen, A. B., Primeau, C., Frei, K. M. & Jørgensen, L. A ritual site with sacrificial wells from the Viking Age at Trelleborg, Denmark. *Dan. J. Archaeol.* **3**, 145–163 (2014).

440. Nørlund, P. Trelleborg, Nordiske Fortidsminder IV (1),. *Det Kongelige Nordiske Oldskriftselskab, Nordisk Forlag, Copenhagen* (1948).

447. Petersen, T. *Undersøkelse Av Gravfunn På Følstad, Røra S, Inderøy Pgd. Innberetning I Topografisk Arkiv,*. (1946).

448. Fyllingen, H. The Use of Human Osteology in the Analysis of Ritual and Violence During the early Bronze Age. A Case Study from Nord-Trøndelag. (2002).

449. Fyllingen, H. Society and violence in the early bronze age: An analysis of human skeletons from Nord-Trøndelag, Norway. *Nor. Archaeol. Rev.* **36**, 27–43 (2003).

450. Schreiner, K. E. Crania Norvegica II. *Instituttet for sammenlignende kulturforskning.Skrifter. Aschehoug,* 5 (1946).

451. Farbregd, O., Marstrander, S. and Torgersen, J. *Bronsealders Skjelettfunn På Sund, Inderøy, Nord-Trøndelag På Bakgrunn Av Andre Bronsealders Funn*. (Norske Vitenskapers selskap Mus. Rapp. A 1974-3, 1974).

452. Lund, H. E. En «tom» eldre bronsealders røisgrav på Skånes i Frol, Nord-Trøndelag. *DKNVS Forhandlinger,Det Kongelige Norske Videnskabers Selskab, Trondheim* **Bd. X, nr. 33**, 121–124 (1937).

453. Petersen, T. *Innberetning Om Undersøkelse Av Gravrøys På Bergvin. Topografisk Arkiv*. (1923).

454. Grønnesby, G. En reise gjennom liv og død, ild og vann. En bronsealders gravruin i Steinkjer. in *Det 10. Nordiske bronsealdersymposium. Trondheim 5.–8.okt. 2006* (ed. Brattli, T.) 66-–78 (Vitark 8. Akademika forlag, 2009).

455. Marstrander, S. Antikvarisk Avdelings Tilvekst 1965. in *Det Kongelige Norske Videnskabers Selskab Museet Årbok Trondheim* 141–147 (Det Kongelige Norske Videnskabers Selskab, Trondheim, 1966).

456. Binns, K. S. Jorda tas i bruk. in *Fosens historie. Fra istid til 1730* (ed. Aune, K.) 73–136 (Fosen historielag, Orkanger, 2005).

457. Scheen, R. De norske flintdolkene: en typologisk-kronologisk analyse. (Mag. Art Thesis, University of Oslo, 1979).

458. Sognnes, K. Prehistoric cave paintings in Norway. *Acta Archaeologica* 101–118 (1982).

459. Bjerck, H. B. On the Outer Fringe of the Human World: Phenomenological Perspectives on Anthropomorphic Cave Paintings in Norway. in *Caves in Context – the cultural significance of Caves and Rockshelters in Europe* 48 –65 (Oxbow Books, 2012).
460. Slomann, W. Der übergang zwischen der späten Kaiserzeit und der frühen Völkerwanderungszeit in Norwegen. in *Archäologische Beiträge zur Chronologie der Völkerwanderungszeit* (eds. Kossack, G. & Reichstein, J.) 61–64 (Dr. Rudolf Habelt, Bonn, 1977).
461. Petersen, T. *Et Gravfunn Fra Merovingertid På Storfosna*. vol. 27 96–99 (Det Kongelige Norske Videnskabers Selskab, 1939).
462. Petersen, T. *Det Kongelige Norske Videnskabers Selskab. Museet. Oldsaksamlingens Tilvekst 1939*. (1939).
463. Rygh, K. *Foreningen Til Norske Fortidsmindesmerkers Bevaring. Aarsberetning for 1879. Fortegnelse over de Til Det Throndhjemske Videnskaberselskabs Samling I 1879 Indkomne Sager, ældre End Reformationen*. (Af K. Rygh, 1879).
464. Marstrander, L. *Inntrøndelag I Romertid - Gravfunn Og Bosetning*. (Gunneria 43.: Det kgl. Norske Videnskabers Selskap-Museet, 1983).
465. Lund Hansen, U. *Römischer Import Im Norden. Warenaustausch Zwischen Dem Römischen Reich Und Dem Freien Germanien*. vol. 10 (Det Kongelige Nordiske Oldskriftselskab. København, 1987).
466. Straume, E. *Gläser Mit Facettenschliff Aus Skandinavischen Gräbern Des 4. Und 5. Jahrhunderts N. Chr*. vol. B (Universitetsforlaget, Oslo, 1987).
467. Grønnesby, G. Bronsealdergravfeltet på Toldnes og Holan i Sparbu. in *Nord-Trøndelag historielag Årbok for 1998* (ed. Bjørgum, A. A.) 148–163 (Nord-Trøndelag historielag, Steinkjer, 1998).
468. Grønnesby, G. En reise gjennom liv og død, ild og vann. En bronsealders gravruin i Steinkjer. in *Det 10 nordiske bronsealdersymposium. Trondheim 5–8 okt. 2006* (eds. Grønnesby, G. & Henriksen, M. M.) 66–78 (2009).
469. Klokkervold, A. Dyrenes rolle i germansk gravskikk. En komparativ analyse av dyrenedleggelser i nordnorske jernaldergraver. (2015).

470. Sjøvold, T. *The Iron Age Settlement of Arctic Norway. Vol. I: Early Iron Age*. (Norwegian University Press, Tromsø/Oslo, 1962).

471. Kaul, F. Fund, helleristninger og landskaber, Nord-Norge. in *Agrarsamfundenes ekspansion i nord* (ed. Sørensen, I. F. K. &.) (Nordlige Verdener, Nationalmuseet, København, 2012).

472. Lund, H. E. *Gravfunn Fra Bronsealderen (skjelettdeler Og Daterende Oldsaker). Skjeggesnes, Alstahaug Pgd, SV. Alstenøya*. (1963).

473. Det kongelige norske videnskabers Selskab, M. O. T. *Det Kongelige Norske Videnskabers Selskab, Museet. Oldsaksamlingens Tilvekst*. (1946).

474. Sjøvold, T. *The Iron Age Settlement of Arctic Norway. Late Iron Age. A Study in the Expansion of European Iron Age Culture Within the Arctic Circle*. vol. 10 (Hassell Street Press, 1974).

475. Alsvik, A. S. *Skjelettgrav Fra 4. årh. e.Kr. På Tommeide, Tomma, Nordland*. (Universitetet i Trondheim, Trondheim, 1978).

476. Brøgger, A. W. *Nord-Norges Bosetningshistorie. En Oversikt*. (Instituttet for sammenlignende kulturforskning, Oslo, 1931).

477. Vinsrygg, S. *Merovingartid I Nord-Noreg. Arkeologiske Avhandlinger Fra Historisk Museum*. (1979).

478. Holck, P. Kråkerøymannen – et gammelt kriminalmysterium? *Borgarsyssel Museum Årbok* 31–42. (1984).

479. Østmo, E. *Krigergraver. En Dokumentarisk Studie Av Senneolittiske Hellekister I Norge*. vol. XXVI (Kulturhistorisk museum, Universitetet i Oslo.Norske Oldfunn, 2011).

480. Resi, H. G. *Gravplassen Hunn I Østfold*. vol. 12 (Norske Oldfunn,O slo, 1986).

481. Reiersen, H. Elite milieus and centres in western Norway 200–550 AD. (University of Bergen, 2017).

482. Lillehammer, G. *Gravskikk På Kvassheimfeltet, Hå I Rogaland, SV Norge*. (AmS-Skrifter 13. Stavanger., 1996).

483. Hauken, Å. D. *Burials on the Beach The Iron Age Cemetery at the Hå Old Parsonage in Rogaland, SW Norway*. vol. 13 (Stavanger, 2023).

484. Møllerop, O. J. Hågravningene. in *Arkeologi, vern og museum: Odmund Møllerop 70 år 7* 99–

106 (AmS-Varia 19.Stavanger, 1992).

485. Lillehammer, G. Kvinnegraver og langhauger. in *Den lengste historia. Rogaland 10 000 f.Kr.-1350 e.Kr* (ed. Obrestad, T.) (Wigestrand, Stavanger, 2004).

486. Kristoffersen, S. *Møt Mennesket. Til Jord Skal vi Bli? Skjeletters Liv Etter Døden*. (AmS-Småtrykk 87.Stavanger, 2016).

493. Wilhelmson, H. *Perspectives from a Human-Centred Archaeology: Iron Age People and Society on Öland*. (Lund University, 2017).

494. Hansen, F. Järnåldersgravar vid Albäcksåns utlopp nära Trelleborg. 44–66 (1945).

495. Stjernquist, B. *Om Järnåldersgravfältet Vid Albäcksån I Maglarp*. vol. 3 (1953).

496. Jacobsson, B. Det forntida gravfältet vid Albäcksån. *Det gamla Trelleborg årsskrift 1978.Trelleborg* (1978).

497. Jacobsson, B. *Albäcksbacken I Maglarp: Gravar Från Stenålder, Bronsålder Och Järnålder, Fornlämning 7 Och 13, Maglarp 23:1, Maglarp Socken, Skåne*. (1984).

498. Arcini, C. Ståtliga var järnåldersmännen från Albäcksbacken. in *Carpe Scaniam. Axplock ur Skånes förflutna.* (ed. Karsten, P.) vol. 22 (Stockholm: Riksantikvarieämbetet. Arkeologiska Undersökningar Skrifter, 1997).

499. Tornberg, A. Health, cattle and ploughs. Bioarchaeological consequences of the Secondary Products Revolution in southern Sweden, 2300–1100 BCE. *Acta Archaeologica Lundensia* **8**, (2018).

500. Malm., A. W. Om några fynd i en forngrift vid Assleröd i Tossene socken, Bohuslän, samt om fornfolkets användning af stenredskap. in *Forhandlingerne ved de skandinaviske Naturforskeres 11th Möde i Kjöbenhavn 1873.* 619—624 (1874).

501. Retzius, G. *Crania Suecica Antiqua*. (1899).

502. Sjögren, K.-G., Price, T. D. & Ahlström, T. Megaliths and mobility in south-western Sweden. Investigating relationships between a local society and its neighbours using strontium isotopes. *Journal of Anthropological Archaeology* **28**, 85–101 (2009).

503. Bägerfeldt, L. & Blomqvist, Å. Fynd och förvåning. *Falbygden* **1987**, 27–39 (1987).

504. Arkæologisk Selskab, J. Anmeldelser 2002. *Kuml* **51**, 297–364 (2002).

505. Ahlström, T. *Underjordiska dödsriken: humanosteologiska studier av neolitiska kollektivgravar.* (Institutionen för arkeologi, Göteborgs universitet, 2009).

506. Lidén, K. *Prehistoric Diet Transitions: An Archaeological Perspective*. (Archaeological Research Laboratory, 1995).

507. Dobiat, C., Sievers, S. & Stöllner, T. *Dürrnberg und Manching: Wirtschaftsarchäologie im ostkeltischen Raum : Akten des internationalen Kolloquiums in Hallein/Bad Dürrnberg vom 7. bis 11. Oktober 1998*. (Römisch-Germanische Kommission. Deutschen Archäologischen Instituts Bonn,., 2002).

508. Maier, F. Gedanken zur Entstehung der industriellen Großsiedlung der Hallstatt- und Latènezeit auf dem Dürrnberg bei Hallein. *Germania* **52**, 326–374 (1974).

509. Stöllner, T. *et al.* The Economy of Dürrnberg-Bei-Hallein: An Iron Age Salt-mining Centre in the Austrian Alps. *The Antiquaries Journal* **83**, 123–194 (2003).

510. Wendling, H. Iron Age Ancestral Bonds: Consecutive Burials and Manipulated Graves at the

Dürrnberg Cemeteries (Austria). in *Grave Disturbances. Studies in Funerary Archaeology* (eds. E. Aspöck, A. K. & Müller-Scheeßel, N.) 157–174 (Oxford, oxbow, 2020).

511. Wendling, H. The Dürrnberg salt metropolis – Catalyst of communication and complexity in La Tène Central Europe. in *Crossing the Alps. Early urbanism between northern Italy and central Europe (900–400 BC)* (eds. Zamboni, L., Fernández-Götz, M. & Metzner-Nebelsick, C.) 393–414 (Sidestone, Leiden, 2020).

512. Wendling, H. *Ursprünge | Zeitsprünge – Reise in Die Urgeschichte Salzburgs. Handbuch Der Urgeschichtlichen Archäologie Einer Alpenregion*. (Salzburg Museum, Salzburg, 2018).

513. Wendling, H. Vom Caput Adriae gen Norden – Transalpine Kontakte in der Späthallstatt- und Frühlatènezeit des Salzburger Raumes, in: W. David and M. Guštin (eds.) The Clash of Cultures? The Celts and the Macedonian World. in 7–17 (Manching, kelten römer museum., 2018).

514. Egg, M., Goedecker-Ciolek, R., Schönfelder, M., Zeller, K. W. †. Ein eisenzeitlicher Prunkschild vom Dürrnberg bei Hallein, Land Salzburg. *Jahrbuch des Römisch-Germanischen Zentralmuseums Mainz* **56**, 81–103 (2009).

515. Zeller, K. W. *Der Dürrnberg Bei Hallein - Ein Zentrum Keltischer Kultur Am Nordrand Der Alpen*. (Keltenmuseum Hallein, 2001).

516. Brandtner, D., Kowarik, K., Reschreiter, H., Rudorfer, J. & Tiefengraber, G. 7000 Jahre Salz. in *Erforschung und Vermittlung des UNESCO-Welterbes Hallstatt-Dachstein/Salzkammergut* (eds. Grömer, K. & Mückler, H.) vol. 151 (Anthropologischen Gesellschaft in Wien, Wien, 2022).

517. Kern, A. The discovery of the cemetery. in *Kingdom of Salt. Monography* (eds. Kern, K., Rausch & Reschreiter) (Natural History Museum Vienna, 2009).

518. Pany-Kucera, D., Reschreiter, H. & Kern, A. Auf den Kopf gestellt? – Überlegungen zu Kinderarbeit und Transport im prähistorischen Salzbergwerk Hallstatt. *Mitt. Anthr. Ges. Wien* **140**, 39–68 (2010).

519. Pany-Kucera, D., Kern, A. & Reschreiter, H. Children in the mines? Tracing potential childhood labour in salt mines from the Early Iron Age in Hallstatt, Austria. *Childhood in the Past* **12**, 67–80 (2019).

520. Pany-Kucera, D., Berner, M., Reschreiter, Kern A., H. & Kowarik, K. Chronische Entzündungen

der Nasennebenhöhlen als Hinweis auf die Umweltbedingungen im eisenzeitlichen Hallstatt. in *Lebenswelten zwischen Archäologie und Geschichte. commemorative publication.* (ed. Drauschke, J., Kislinger, E., Kühtreiber, K., Kühtreiber, T., Scharrer-Liska, G., Vida, T.) vol. 150 985–995 (Schnell & Steiner, 2018).

521. Slavchev, V. A Mid-5th Millennium Settlement near Suvorovo, Varna District, Bulgaria. in *International Conference Where Are the Sites? Research, Protection and Management of Cultural Heritage, 5–8 December 2013* (ed. H. Angelova, M. Ö.) 29–34 (Ahtopol, p., 2013).

522. Kazarnitski, A. A. The anthropological description of the skeletons from the Late Antic burial ground of the Suvlu-Khaya. Preprint at (2016).

523. Voloshinov, A. A. & Masyakin, V. V. The German elements from the Suvlu-Khaya necropolis. Preprint at (2016).

524. Belov, G. D. The necropolis of Chersonesos of the classical era. *Soviet archeology* 272–284 (1950).

525. Stoyanov, R. V. Necropolis of Tauric Chersonesos V–I centuries BC. (Historical sciences. St. Petersburg, 2004).

526. Ivanov, A. V. Population of ancient Tauric Chersonesos - Byzantine Kherson according to anthropological data. Preprint at (2016).

527. Belov, G. D. Report on excavations in Chersonesos for 1935–1936. Preprint at (1938).

528. Tejral, J. Zur Unterscheidung des vorlangobardsichen und elbgermanischlangobardischen Nachlasses. in *Die Langobarden. Herrschaft und Identität* (eds. Pohl, W. & Erhart, P.) (Austrian Academy of Sciences press, Wien, 2005).

529. Droberjar, E. *Věk Barbarů. České Země a Stěhování Národů Z Pohledu Archeologie.* (Praha – Litomyšl, 2005).

530. Smrčka, V, Marcsik, A, Svenssonova, M. The dietary trends and social relations in the migration period. *Acta Universitatis Carolinae* 73–90 (2009).

531. Svenssonova, M. Population settled north of the middle Danube during Merovingian time( in Czech). (Masaryk University Brno, 2002).

532. Waldhauser, J. *Jak Se Kopou Keltské hroby.Laténská Pohřebiště Ze 4.-3- Století v Čechách*

*Nakladatelství Lidové Noviny 340*. (1999).

533. Waldhauser, J. Keltische Gräberfelder in Böhmen. *Bericht der Römisch-germanischen Kommission* **68**, 25–179 (1987).
534. Formisto, T. *An Osteological Analysis of Human and Animal Bones from Levänluhta.* (Stockholms universitet, 1993).
535. Wessman, A. Levänluhta: – a place punishment, sacrifice or just a common cemetery? *Fennoscandia Archaeologica* **26**, 47–71 (2009).
536. Wessman, A. *et al.* Hidden and Remote: New Perspectives on the People in the Levänluhta Water Burial, Western Finland (c. ad 300–800). *Eur. J. Archaeol.* **21**, 431–454 (2017).
537. Niskanen, M. Stature of the Merovingian-period inhabitants from levänluhta, Finland. *Fennoscandia Archaeologica* **23**, 24–36 (2010).
538. Maijanen, H., Junno, J.-A., Mannermaa, K., Niskanen, M. & Wessman, A. Re-analysis of the Levänluhta skeletal material: Sex and stature estimation of individuals in an Iron Age water burial in Finland. *Int. J. Osteoarchaeol.* **31**, 347–357 (2021).
539. Sikora, M. *et al.* The population history of northeastern Siberia since the Pleistocene. *Nature* **570**, 182–188 (2019).
540. Desenne, S., Pommepuy, C. & & Demoule, J.-P. Bucy-le-Long, Aisne, une nécropole de La Tène ancienne (Ve-IVe siècle avant notre ère). *Rev. Med. Picardie* **26**, (2009).
541. Demoule, J.-P. Chronologie et société dans les necropolis celtiques de la culture Aisne-Marne, du VIème au IIIème siècle avant notre ère. *Rev. Med. Picardie* **406**, (2005).
542. Pommepuy, C., Auxiette, G. & Desenne, S. Rupture et continuité dans les pratiques funéraires de La Tène ancienne et moyenne/finale à Bucy-le-Long (Aisne). *Rev. archéol. Picardie* **1**, 85–98 (1998).
543. Brun, P. Les ‘ Résidences princières ’ comme centres territoriaux : éléments de vérification. in *Les Princes celtes et la Méditerranée, Paris novembre 1987* 128–143 (Paris : La Documentation française, 1988).
544. Chaume, B. *Vix et son territoire à l'âge du fer: fouilles du mont Lassois et environnement du site princier*. (M. Mergoil, 2001).

545. Brun, P. & Chaume, B. Une éphémère tentative d'urbanisation en Europe centre-occidentale aux VIe et Ve s. av. J.-C. ? *Bulletin de la Société préhistorique française* **110**, 319–349 (2013).
546. Chaume, B. *et al.* Entre l'État et la chefferie simple : le complexe aristocratique de Vix/le mont Lassois. in *Vix et le phénomène princier.* (ed. Actes du colloque international de Châtillon-sur-Seine, 26–28 octobre) (Bordeaux : DAN@, 2021, 2021).
547. Piningre, J. F. & Gannard, V. Les nécropoles protohistoriques des Moidons et le site princier du Camp du Château à Salins (Jura). *Documents* **17**, (2004).
548. Brun, P. & Chaume, B. An unfinished trend : towards urbanization in Celtic regions north of Alps (575–450 BCA). in *Vix et le phénomène princier. Actes du colloque international de Châtillon-sur-Seine, 26–28 octobre 2016* 367–389 (Bordeaux : DAN@, 2021).
549. Chaume, B. Vix et l'émergence des principautés celtiques : l'hypothèse portuaire et le concept de port of trade. in *Les ports romains dans les Trois Gaules : entre Atlantique et eaux intérieures. Actes du colloque international d'archéologie portuaire, Nantes 21–22 juin 2018* (eds. Mouchard, J. & Guitton, D.) 435–452 (Gallia, 2020).
550. Chaume, B. Vix: the Temptation of the City. in *Crossing the Alps. Early Urbanism between Northern Italy and Central Europe (900-400 BC)* 349–360 (Sidestone Press, 2019).
551. Rolley, C. *La Tombe Princière de Vix*. vol. 1 (Picard, 2003).
552. Chaume, B. Tracking down the story of the discovery of the Vix princely burial : new results. *Germania* **96**, 93–138 (2018).
553. Reinhard, W. *Studien Zur Hallstatt- Und Frühlatènezeit Im Südöstlichen Saarland.* (Bliesbruck-Reinheim, 2003).
554. Herrscher, E, Bedianashvili, G, Chkadua, M, Vanishvili, N, Messager, E, Martin, L. Millet crop and consumption during Late Bronze Age in Georgia: First human and animal isotopic evidence. *American Journal of Physical Anthropology* 141 (2014).
555. Paja, L. Anthropological study of four Sarmatian osteological series (Röszke (sites 48/60 and 48/75), Kiskundorozsma (sites 26/60 and 26/72)). in *Museological research along the alignment of the M5 motorway.* (ed. Szalontai, C. S.) 165–168 (University of Szeged, 2003).
556. Paja, L, Marcsik, A. Paleopathological and paleodemographical analysis of Sarmatian

osteological series originating from southern Hungary. *Acta Univ. Carol. Med. Monogr.* 57–64 (2009).

557. Szalontai, CS, Tóth, K. Szeged-Kiskundorozsma-Nagyszék II. (site 26/72 No. 34, M5 motorway). in *Museological research along the alignment of the M5 motorway* (ed. Szalontai, C. S.) 69–81 (University of Szeged, 2003).
558. Kőhegyi, M, Vörös, G. *Madaras-Halmok. Kr. U. 2–5. Századi Szarmata Temető. Monográfiák a Szegedi Tudományegyetem Régészeti Tanszékéről 1. Ed*. (Szegedi Tudományegyetem, 2011).
559. Gasparri, S. *Italia Longobarda: Il Regno, I Franchi, Il Papato*. (Laterza, 2012).
560. Bertašius, M. *Marvelė. Ein Gräberfeld Mittel-Litauens. Vidurio Lietuvos Aukštaičių II–XII A. Kapinynas*. vol. 1 (Kaunas, 2005).
561. Bertašius, M. *Marvelė. Ein Bestattungsplatz Mit Mittel-Litauischer Pferdegräber. Marvelės žirgų Kapiny-Nas*. vol. 2 (Kaunas, 2009).
562. Valatka, V. Maudžiorų (Kelmės raj.) kapinyno tyrinėjimai 1964 m. *Archeologiniai tyrinėjimai Lietuvoje* 10–11 (1965).
563. Valatkienė, L. Maudžiorų plokštinis kapinynas. *Archeologiniai tyrinėjimai Lietuvoje* 78–79 (1985).
564. Valatkienė, L. Maudžiorų plokštinio kapinyno kasinėjimai. *Archeologiniai tyrinėjimai Lietuvoje* 104–106 (1983).
565. Valatkienė, L. Maudžiorų senkapio (Kelmės raj.) tyrinėjimai 1978 ir 1979 metais. *Archeologiniai tyrinėjimai Lietuvoje* 89–92 (1979).
566. Stankus, J. Kairėnėlių plokštinis kapinynas. *Lietuvos archeologija. Vilnius* 63–79 (1984).
567. Kazakevičius, V. Aukštaičių kapinynas žemaičių etnokultūrinėje periferijoje. *Lietuvos archeologija, Vilnius* 57–67 (1999).
568. Simniškytė, A. Vaineikių pilkapyno tyrinėjimai 1998 metais. *Archeologiniai tyrimai Lietuvoje* 201–202 (1999).
569. Simniškytė, A. Vaineikių pilkapyno tyrinėjimai 1997 metais. *Archeologiniai tyrimai Lietuvoje* 155–156 (1997).
570. Salatkienė, B. Kalnelių pilkapiai. *Archeologiniai tyrimai Lietuvoje* 60–64. (1983).

576. Smits, E. Leven en sterven langs de Limes : het fysisch-antropologisch onderzoek van vier grafveldpopulaties uit de noordelijke grenszone van Germania Inferior in de Vroeg- en Midden-Romeinse tijd. Preprint at (2006).

577. Lonnée, H. A. & Maat, G. J. R. *Inhumations in a Roman Cemetery at Valkenburg-Marktveld (Zuid-Holland) in The Netherlands*. vol. 3 (Barge's Anthropologica, Leiden, 1998).

578. Zilhão, J. *Gruta Do Caldeirão. O Neolítico Antigo*. (Instituto português do património arquitectónico e arqueológico, 1992).

579. Shirobokov I.G., Chernykh E.M., Nechvaloda A.I. The anthropological characteristics of the skeletons from the Dubrovskiy burial ground. *Bulletin of Udmurt University. History and Philology Series.* 500–512 (2018).

580. Shirobokov, I. G., Chernykh, E M. The origin of the Kama populations of the middle of the first millennium AD (based on Boyarsky 'Arai' burial ground). *Bulletin of Udmurt University. History and Philology Series* 25–34 (2016).

581. Malashev V.Yu., Gadiev U.B., Valchak S.B., Friesen S.Yu., Shaushev K.B. Research of the

burial ground “Ekazhevsky 1st mounds. *Bulletin of the Archaeological Center* **VI**, 222–233 (2018).

582. Voronin, K. V. & Malashev, V. Y. Funerary monuments of the Bronze Age and Early Iron Age of the plain zone of the Republic of Ingushetia. Preprint at (2006).

583. Atabiev, B. K. Excavations near the village of Zaragizh of the Kabardino-Balkarian Republic, Archaeological discoveries 1995, L’or des princes barbares. *Archaeologia* **371**, (2001).

584. Murphy, E. M. An Osteological and Palaeopathological study of the Scythian and Hunno-Sarmatian period populations from the cemetery complex of Aymyrlyg, Tuva, South Siberia. (1998).

585. Mandelshtam, A. M. Ranniye kochevniki Skifskova perioda na territorii Tuvi,. in *Stepnaya Polosa Aziatskoi Chasti SSSR v Skifo-Sarmatskoye Vremya, Archeologiya SSSR* (ed. Moshkova, M. G.) 178–196 (Nauka, Moskva, 1992).

586. Stambulnik, E. U. Noviye Pamyatniki Gunno-Sarmatskova Vremeni B Tyvye: Nekotoriye Itogi Rabot,. in *Drevniye Kulturi Euraziiskih Stepei* 34–41 (Institute for the History of Material Culture. Leningrad, 1983).

587. Murphy, E. M. A Bioarchaeological Study of Xiongnu Expansion in Iron Age Tuva, South Siberia. in *Regimes and Revolutions: Power, Violence, and Labor in Eurasia from the Ancient to the Modern* (eds. Hartley, C. W., Yazicioğlu, G. B. & Smith, A. T.) (Cambridge University Press, 2012).

588. Mandelshtam A. M., S. E. U. Gunno-Sarmatskii period na territorii Tuvi,. in *Stepnaya Polosa Aziatskoi Chasti SSSR v Skifo-Sarmatskoye Vremya, Archeologiya SSSR* (ed. Moshkova, M. G.) 196–205 (Nauka, Moskva, 1992).

589. Bogdanova, V.I., Radzjun, A.B. Paleoantropologicheskiye Materiali Gunno-Sarmatskova vremeni iz Tsentralnoy Tuvi, Noviye Kollektsii I Issledovaniya Po Antropologii I Arheologii. *Antropologii I Etnografii* 55–74 (1991).

590. Taylor, G. M., Murphy, E., Hopkins, R., Rutland, P. & Chistov, Y. First report of Mycobacterium bovis DNA in human remains from the Iron Age. *Microbiology* **153**, 1243–1249 (2007).

591. Murphy, E., Chistov, Y., Hopkins, R., Rutland, P. & Taylor, G. Tuberculosis among Iron Age

individuals from Tyva, South Siberia: palaeopathological and biomolecular findings. *J. Archaeol. Sci.* **36**, 2029–2038 (2009).

592. Murphy, E., Donnelly, U. M. & Rose, G. Possible neurofibromatosis in a Scythian period individual from the cemetery of Aymyrlyg, Tuva, South Siberia. *International Journal of Osteoarchaeology* **8**, 424–430 (1998).

593. Murphy, E. M. Developmental defects and disability: the evidence from the Iron Age semi-nomadic peoples of Aymyrlyg, south Siberia. in *Madness, Disability and Social Exclusion (One World Archaeology 40)* (ed. Hubert, J.) vol. 1 60–80 (Routledge, 2010).

594. Goldina, R. D. Drevnyaya i srednevekovaya istoria udmurtskogo naroda (Ancient and Medieval history of the Udmurt people). Preprint at (2004).

595. Ostanina, T. I. Naselenie Srednego Prikam'ya v 3-5 vv. (The population of the Middle Kama in the 3–5 centuries AD). Preprint at (1997).

596. Pichlerová, M. Gerulata und seine Rolle im Bratislavaer Tor. *Archeologické rozhledy* **XXXVIII**, 435–445 (1986).

597. Smrčka V., Jambor J., Salaš M. Diet in the 1st–2nd centuries along northern border of the Roman Empire. *Anthropologie* **XXVI**, 39–54 (1988).

598. Kolník, T. Cífer-Pác, stanica z mladšej doby rímskej. Preprint at (1975).

599. Baldovič, M. Anthropological analysis of skeletal burial place Cífer-Pác (dist. Trnava) from 8th – 9th century AD. (2003).

600. Šebest, L. *et al.* Detection of mitochondrial haplogroups in a small avar-slavic population from the eigth-ninth century AD. *Am. J. Phys. Anthropol.* **165**, 536–553 (2018).

601. Kyselicová, K. *et al.* Molecular detection of Mycobacterium tuberculosis complex in the 8th century skeletal remains from the territory of Slovakia. **71**, 613–619 (2016).

602. Hutton, R. Romano-British reuse of prehistoric ritual sites. *Britannia* **42**, 1–22 (2011).

603. Witts, G. B. Randwick long barrow. *Proceedings of the Cotteswold Naturalists' Field Club* **8**, 156–160 (1883).

604. Card, N. & Downes, J. Mine Howe – the significance of space and place in the Iron Age. in *Sea Change: Orkney and Northern Europe in the later Iron Age AD 300-800* (eds. Downes, J. &

Ritchie, A.) 11–19 (Pinkfoot, 2003).
