## Supplementary figures and images for "Steppe Ancestry in Western Eurasia and the Spread of the Germanic Languages"

### Fig. S5.18

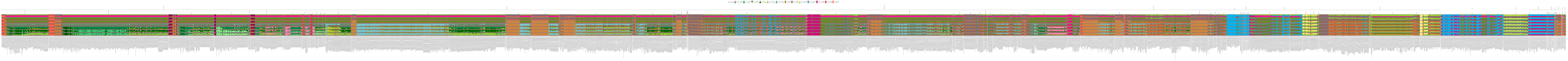

### Fig. S5.19

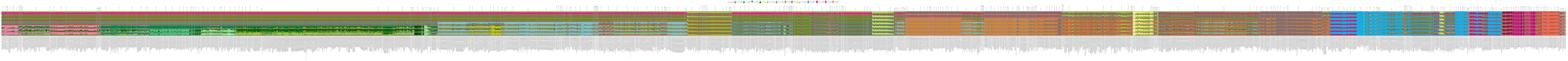

### Fig. S5.20

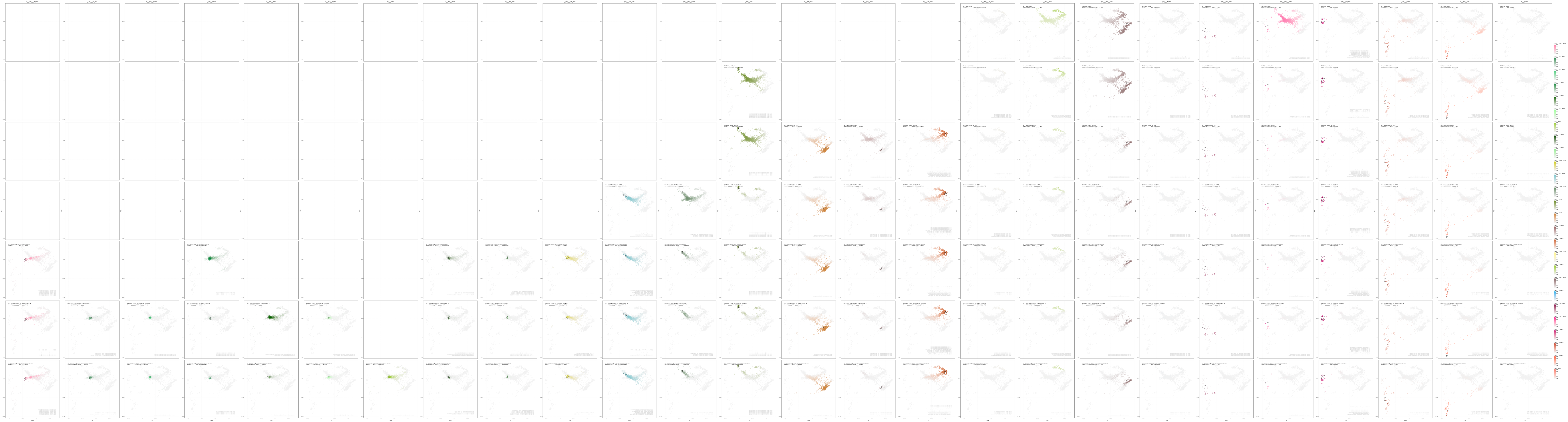

### Fig. S5.21

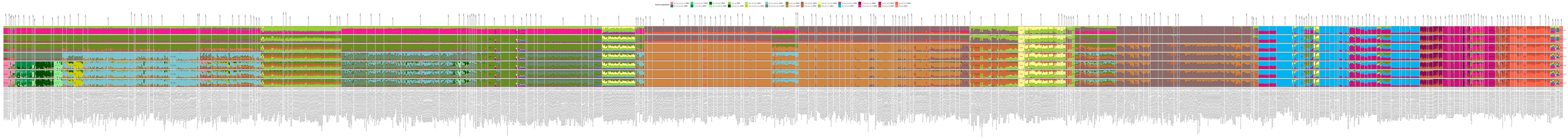

### Fig. S5.25

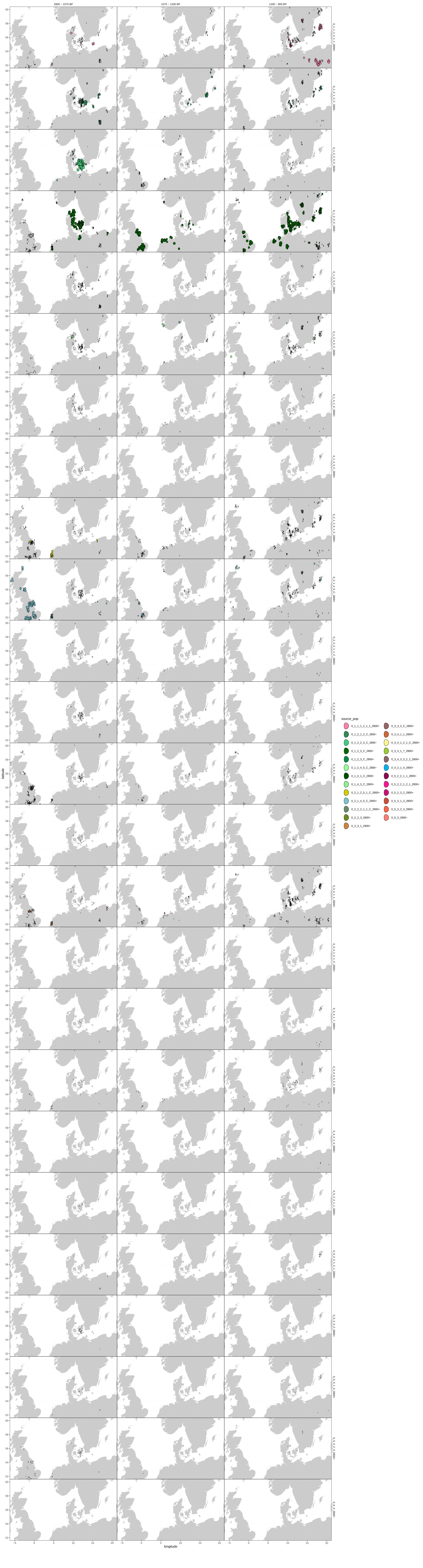

### Fig. S5.26

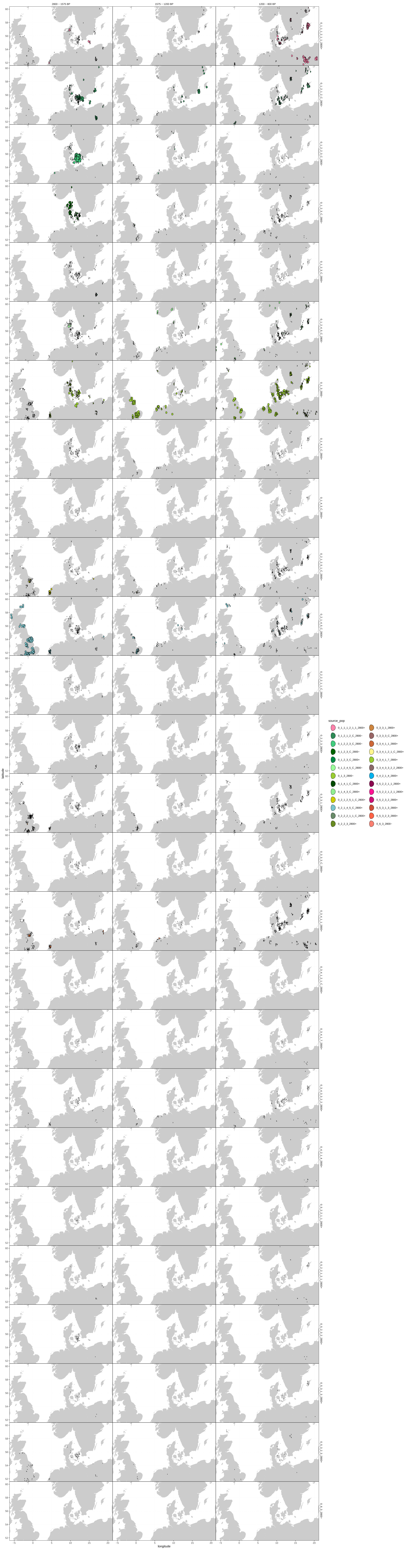

### Fig. S5.29

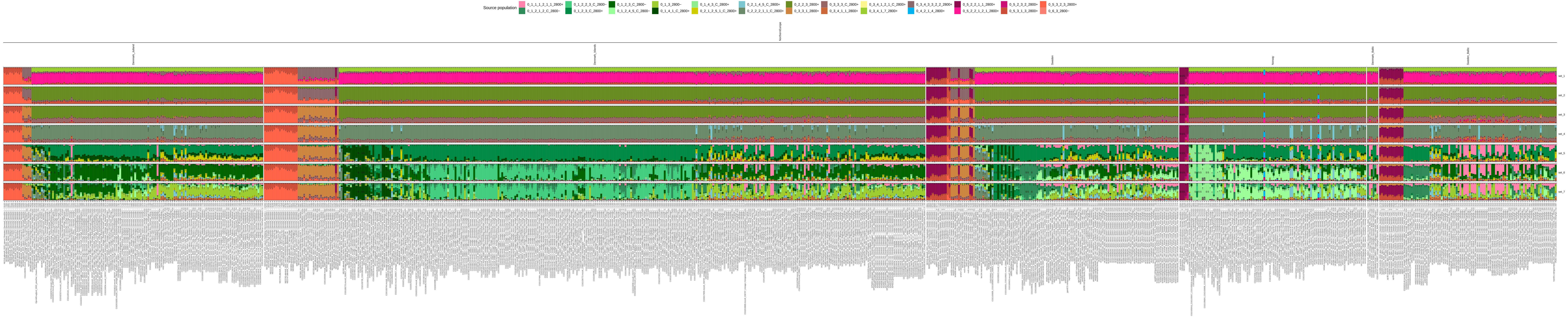

### Fig. S5.30

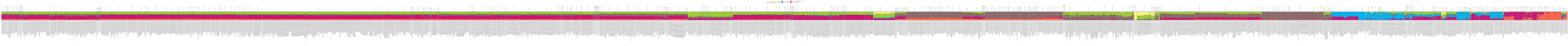

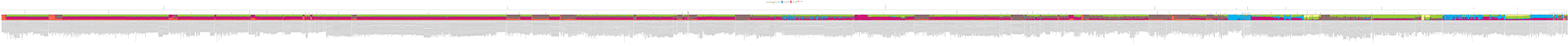

### Fig. S5.31

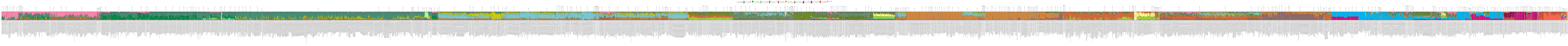

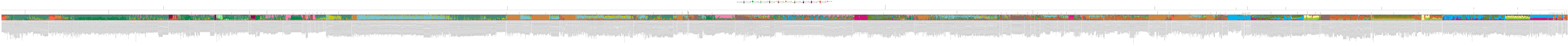

### Fig. S5.32

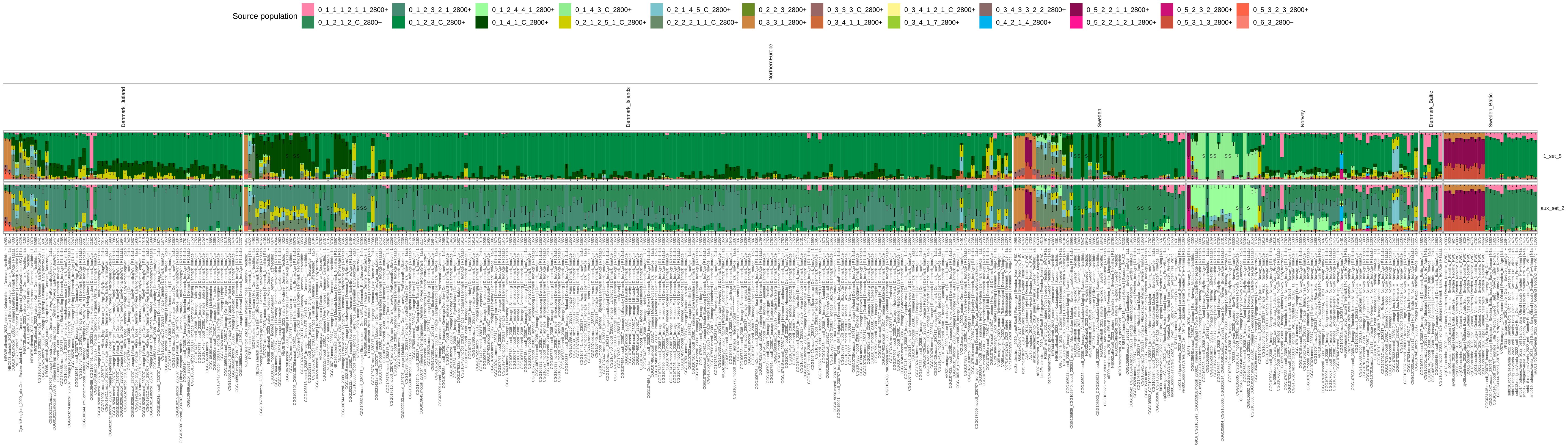

### Fig. S5.141

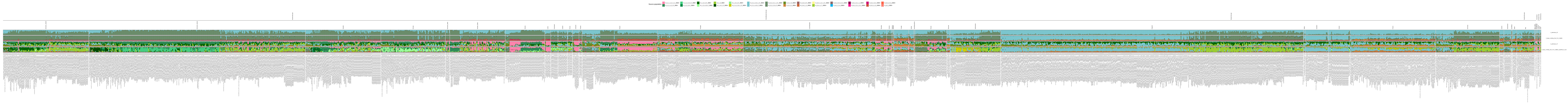

### Fig. S5.142

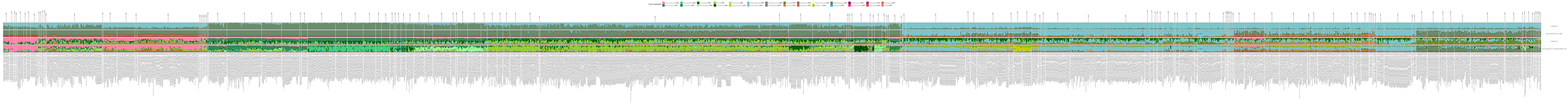
