## Supplementary Notes S1-6 for "Steppe Ancestry in Western Eurasia and the Spread of the Germanic Languages"

#### Genetics Supplementary Material

### S1. Data Generation

*Charleen Gaunitz, Lasse Vinner*

All aDNA laboratory procedures on non-amplified DNA were conducted in dedicated aDNA clean lab facilities at the Lundbeck Centre for GeoGenetics, University of Copenhagen, according to strict guidelines[101,102]. Post-amplification procedures were conducted in post-PCR laboratories physically separated from the clean lab facility.

During the project, a semi-automated data generation pipeline was introduced over a two-year period, hence the samples included in the present study were processed in an increasingly automated manner as procedures were automated for extraction, library preparation, PCR setup, post-amplification purification and library pooling.

#### Curation of included samples

In order to improve data quality and minimise destruction of unproductive samples, we implemented a thorough quality control check to make sure that all the samples analysed, teeth and petrous bones, were of suitable quality.

Macroscopic study of the surface of the teeth using a magnifier and an indirect source of light was performed. We only considered teeth that were fully formed, erupted and showing complete roots with the apex intact. We only retained teeth with roots of yellow to brownish colour. Teeth with roots of chalky or white colours were excluded.

The petrous bones were considered when the portion containing the cochlea was present. The petrous bones that were cut off too short from the temporal bone were excluded, as well as those that were showing a too chalky colour aspect and a very light weight.

#### Drilling

Drilling was performed manually. If possible, right or left petrous bone, one tooth and associated calculus were subsampled from one individual. Up to 150 mg of crushed subsample were stored in a 2 ml LVL barcoded tube (XLX2000, DSC-X20-BL-NS-SLP-S). While sampling for DNA analysis, a small piece of bone (up to 1 gram) was collected for radiocarbon dating. Sample status was documented photographically before and after the destructive sampling. Subsamples were stored at -20°C until demineralisation.

#### Demineralisation

Sample demineralisation was carried out in two steps. First, an initial short incubation step was performed at 37°C for 30 mins to increase the recovery rate for endogenous DNA, using 1.0ml of incomplete digestion buffer consisting of 0.5M EDTA and 30% N-Lauroylsarcosin[103]. The buffer was replaced with 1.8ml of freshly prepared digestion buffer (0.5M EDTA, Proteinase K and 30% N-Lauroylsarcosin) and incubated for 48 to 72 hours at 37°C on a rocking table. Demineralised samples (lysates) were stored at 4°C for no longer than 14 days, otherwise at -20°C for long-term storage.

#### Ancient DNA extraction

All aDNA extractions for this project were carried out according to Rohland et al.[104] using a modified version of the Qiagen PB buffer (Qiagen, cat: 19066). In total, two extractions were performed per lysate for initial library sequencing. One extraction per lysate underwent treatment with USER enzyme mix (NEB, cat: M5505L), the other remained untreated.

#### Manual extraction setup

For manual extractions, 1.8 ml of lysate was combined with 18 ml of binding buffer (modified Qiagen PB binding buffer: 500 ml Qiagen PB, 15 ml Sodium acetate 3 M, 1.25 ml 5 M NaCl, phenol red, adjusted to pH=5). The mix was passed through a large-volume silica column (Roche, cat: 05114403001) by centrifugation for 4 mins at 1500 rpm. The flow-through was discarded and the step above repeated 2 times for the full volume. The column was washed twice using 750 µL 80% ethanol + 20% 10 mM Tris-HCl (Qiagen, cat: 19065) and a centrifugation step of 30 seconds at 6-8 K·g, followed by a dry spin at full speed, according to kit protocol. The column was then transferred to a fresh 1.5 ml LoBind DNA Eppendorf tube (Eppendorf, cat: 0030108051) before final elution with 65 µl of 10 mM Tris-HCl + 0.05% Tween-20 incubated for 5 mins at room temperature, followed by a centrifugation step for 1 min at 11.5 K·g. Extractions were stored at 4°C until library preparation, no more than 2 days, or longer at -20°C.

#### Automated aDNA extraction setup

The aDNA extraction procedure was automated on the Biomek i5 Automated Workstation (Beckman Coulter) with the following adaptations: For each subsample, 150 µl of demineralised lysate were combined with 1560 µl of binding buffer and 10 µl of magnetic silica beads (G-Bioscience, cat: 786-915). The mixture was incubated for 15 mins and tip-mixed every 5 mins. Pelleted beads were washed twice in 450µl and 100 µl of 80% ethanol + 20% 10 mM Tris-HCl, respectively (Qiagen PE, cat: 19065). The final product was eluted in 35 µl of 10 mM Tris-HCl + 0.05% Tween-20. Extracted DNA was stored in non-skirted 96-well plates (VWR) at 4°C until library preparation for no more than 2 days, otherwise at -20°C. All extraction runs included a positive control sample and ≥1 negative control (buffer only). Robot deck specifications, method (.bmf format) and plastic/labware are provided on request.

To verify successful DNA extraction, 2 µl of the positive and negative control extracts were measured on QuBit (dsDNA HS Assay Kit, cat: Q32851). The QuBit threshold value for the positive extraction control was >0.1ng/µl for approval, whereas the negative was below the lower detection limit.

#### USER treatment

For the USER treatment, 10 µl of USER enzyme and 35 µl of DNA extract were mixed and incubated for 3 h at 37°C in a thermal cycler (SimpliAmp™ ThermoFischer). The treatment was performed in a 96-well format. After performing tests with reduced USER enzyme volume, the reaction volume of the USER enzyme was reduced to 2.5 µl. From May 2022, the USER treatment protocol reads: 2.5 µl USER enzyme + 7.5 µl H2O + 35 µl aDNA, incubated for 3 h at 37°C.

#### NGS library preparation

##### Manual preparation

Double-stranded DNA libraries were prepared according to Meyer & Kircher[89], Kapp et al.[90] or Gansauge et al.[91] from the USER- and one non-USER-treated extracts, using input template volumes of 42.5 µl (final reaction volume 50 µl) and 21.25 µl (final reaction volume 2 µl), respectively. Clean-up steps were performed after the end-repair and adapter-ligation step, using 10 volumes of modified Qiagen binding buffer (as above) and MinElute columns (Qiagen, cat: 28004) otherwise according to the MinElute kit protocol.

#### Automated NGS library preparation

The NGS library preparation for USER-treated and non-USER-treated libraries was also adapted to the Biomek i5 Automated Workstation (Beckman Coulter). Two different methods were written to accommodate the different input volumes of USER and non-USER extracts. Clean-up steps were performed after the end-repair and adapter-ligation step, using 10 volumes of modified Qiagen binding buffer (as above) and 10 µl of magnetic silica beads (G-bioscience, cat: 786-915). Pelleted beads were washed twice in 450 µl and 100 µl of 80% ethanol + 20% 10 mM Tris-HCl, respectively (Qiagen PE buffer). The purified products were eluted in 10 mM Tris-HCl + 0.05% Tween-20. Robot deck specifications, method (.bmf format) and plastic/labware are provided on request.

### Real-time PCR

We used real-time PCR for determining the number of PCR cycles required in the indexing PCR, for each batch of libraries. Excluding the control libraries, $c_t$-values for the sample libraries were converted to a consensus amplification cycle number, considering amplification curves, $c_t$-values and melting curves. The qPCR with a final reaction volume of 20µl was set up manually in a 96-well format (Roche, cat: 5102413001), using 1 µl of input material, 7 µl of $H_2O$, 500 nM forward- and 500 nM reverse primer (10 mM) and 10 µl of 2x Lightcycler 480 SYBR green I MasterMix (Roche, cat: 4707516001). The qPCR amplification was run in the POST PCR lab facilities using the Roche LightCycler 480 real-time PCR system (Roche).

**Table S1.2. Primer sequences**

| Primer ID | Primer Sequence (5'-3') |
|---|---|

| | |
|---|---|
| qPCR CH_P5_ | CTACTGACTTTCAGTGAGTGCAACCCACGACGCTCTTCCGATCT |
| qPCR CH_P7_ | CTCTCACATTGAATCCGACTAGGATACGTGTGCTCTTCCGATCT |

**Table S1.3. qPCR conditions**

| Temp | Time | Cycles |
|---|---|---|
| 95°C | 10 min. | 1 |
| 95°C<br>55°C<br>72°C | 30 sec.<br>30 sec.<br>30 sec. | 30<br>Detection |
| 95°C<br>55°C<br>95°C Slow ramp | 30 sec.<br>30 sec.<br>30 sec. | 1<br>Detection |

### Indexing PCR

The indexing PCR was set up either in a small PCR reaction volume of 50 µl for non-USER-treated libraries or in a big PCR reaction volume of 100 µl for USER-treated libraries. For the small PCR reaction setup, 25 µl of KAPA HiFi HotStart Uracil+ ReadyMix (Roche, cat: 07959079001) was mixed with 4 µl of UDI (8-bp index) or UDP (10-bp index) Illumina primer pairs (400 nM final conc. each) and 21 µl of template DNA. For the large PCR reaction setup, 50 µl of KAPA HiFi HotStart Uracil+ ReadyMix (Roche, cat: 07959079001) was mixed with 8 µl of UDI or UDP Illumina primer pairs (400 nM final conc. each) and 42 µl of DNA.

**Table S1.4. PCR amplification conditions.**

<table>
<tr><th>Temp.</th><th>Time</th><th>Cycles</th></tr>
<tr><td>98</td><td>45 sec.</td><td>1</td></tr>
<tr><td>98</td><td>15 sec.</td><td rowspan="3">12-18</td></tr>
<tr><td>65</td><td>30 sec.</td></tr>
<tr><td>72</td><td>30 sec.</td></tr>
<tr><td>72</td><td>min.</td><td>1</td></tr>
</table>

PCR amplification was performed in the post-PCR lab facilities. PCR amplification cycles were determined as described above. Indexing PCR setup was semi-automated on the Biomeki5 Automated Workstation (Beckman Coulter). The KAPA HiFi HotStart Uracil+ ReadyMix (Roche, cat: 07959079001) was dispensed manually to a non-skirted 96-well PCR plate (VWR) using a multi-dispenser pipette and then placed on the 96-well cooling element on the Biomek i5. Index primer pairs and template DNA were then transferred by the multichannel robot head to the plate. Robot deck specifications, method (.bmf format) and plastic/labware are provided on request.

#### Purification of amplified libraries

##### Manual procedure

For non-USER- and USER-treated libraries, respectively, 80 µl or 160 µl of MagBio High-Prep PCR beads per library were transferred into a 96-deep well plate (e.g., ThermoFisher cat: 267245). Subsequently, 50 µl or 100 µl, respectively, of amplified library were added to the beads, thoroughly mixed and incubated for 5 mins at room temperature. The plate was placed onto a magnetic plate and incubated until the supernatant was clear. While still on the magnet, the beads were washed twice with 200 µl of 80% freshly made ethanol. After discarding the last washing buffer, the plate was left to dry for 2-5 mins. The plate was removed from the magnet. The beads were resuspended in 35 µl of 10 mM tris-HCl pH= 8.5 (EB Buffer, Qiagen) and incubated for 2 mins, before clearing the supernatant on the magnet (Alpaqua Magnum FLX). Once clear, the supernatant was transferred to a storage tube (LVL technologies, 2DSC-X03-BL-NS-SLC-S).

##### Automated purification

The purification was set up in an automated 96-well format using the CyBioFelix liquid handler (Analytik Jena), following the conditions described above. Two protocols are currently available. One using the 96R Head, allowing the purification of 96 samples at once with a run time of 47 mins, and one using the Choice Head (8-Channel). For the latter, the number of samples to purify can be chosen. Minimum is 8, maximum is 96. The final purified DNA library (35µl) is stored in SX 300 1D and 2D barcoded tubes (LVL technologies, 2DSC-X03-BL-NS-SLC-S) at -20°C. The rack is scanned prior to purification using the Ziath express scanner. Labware and method are available upon request.

##### Quality control (Fragment Analyzer)

Library concentration and fragment length distribution was determined by using the 5300 Fragment Analyzer System (Agilent) with the HS NGS Fragment Kit (1-6000 bp), 500 (cat: DNF-474-0500, Agilent). The standard protocol provided by Agilent is followed to run the HS NGS Fragment Kit, using 2 µl of library input with 22 µl Diluent Marker (total 24 µl) in a semi-skirted Eppendorf twin.tec PCR Plate 96. For data analysis, the ProSize (Agilent) software was used.

#### Pooling of libraries

##### Manual procedure

Each sequencing pool consisted of a maximum of 96 libraries, with each library distinguishable by a unique index pair (Truseq UDP or UDI, Illumina). For initial sequencing, the libraries were pooled equimolarly.

##### Automated procedure

The preparation of sequencing pools was automated on the CyBioFelix liquid handler (Analytik Jena), with the possibility to pool up to 192 uniquely indexed libraries in one run. Input pooling volumes, rack position and optional dilution requirement were loaded via a .csv file.

The pooling method itself is divided into 2 parts, which can be run separately. In the first step, a dilution of all libraries is prepared. The dilution factor is the same for all samples in the plate (typically 5 or 10-fold) and created to ensure no pipetting of ≤ 1µl. In the second part of the method, the pooling of the diluted and undiluted libraries was carried out. Samples are transferred into a 1.5 ml Lobind Eppendorf tube, starting from the largest volume to the smallest volume. The final sequencing pool was purified manually using 1.6x volumes of paramagnetic beads (Magbio High prep Beads, cat: AC-60500) and washed with 80% freshly prepared ethanol (500µl), removing residual adapters and indexing primer. To compensate for possible loss during purification, pools were eluted in 2/3 of their original volumes. CyBioFelix method and labware files are available on request.

#### Sequencing

The molar concentration of sequencing pools was determined using qPCR (KAPA Library Quantification Kit, KAPA Biosystems, KR0405) according to kit protocol. All sequencing was done on the Illumina Hiseq 4000, as described in Allentoft et al.[28], or the NovaSeq6000 platform at the GeoGenetics Sequencing Core. Sequencing chemistry version 1.0 was used in the beginning of the project, but was later switched to the latest version 1.5 in mid-2021 due to updated kit version. In-house comparison between kit versions revealed no biases in subsequent analysis pipelines. For the USER-treated libraries, pools were sequenced on the Illumina S4 flowcell, 200 cycles, generating ≥10 G paired-end reads. For the non-USER-treated libraries, pools were sequenced on one lane of an Illumina S4 flow cell, 2x100 cycles, using Illumina's XP kit and workflow, generating 2.5 G sequencing reads of 100 bp paired-end. Loading concentrations for ancient DNA libraries deviate from Illumina recommendations for loading concentrations and differ between the two types of runs, i.e., full flow cells and XP workflows. For a full flow cell run, aDNA library loading concentration is optimal at 700 pM, and for the XP runs, loading concentration seems optimal at 550-600 pM. Each pool is quality-checked before sequencing, using Fragment Analyzer or Bioanalyzer to determine the average bp size and profile of the library, and a qPCR to determine molarity of the pool. The yields from full flow cells seem more stable >10 G reads than output from XP workflows ≥2.5 G reads. Demultiplexing of sequencing output was performed on BaseSpace Sequence Hub (Illumina).

### S2. Bioinformatics preprocessing pipeline

*Abigail Ramsøe, Isin Altinkaya, Thorfinn Sand Korneliussen*

#### Demultiplexing

The sequencing data was demultiplexed on the BaseSpace Sequence Hub using BCL Convert (*v4.0.3* BCL Convert Support, Illumina Inc.), allowing for one mismatch in the index. As this strategy allows the inclusion of reads with a single mismatch in the index sequence, it avoids the needless exclusion of reads. Furthermore, as the libraries were sequenced using Unique Dual Indexing (UDI), and the Illumina index sets are designed with a Hamming distance of four, the chances of a sequenced read being misassigned is nil. This generates two fastq files (read 1 and read 2) per library per lane – in total two fastq files per library for XP runs, and eight for full flow cells.

#### Trimming

AdapterRemoval --threads {threads} --file1 {input.R1} --file2 {input.R2} --minlength 30 --adapter1 {a1} --adapter2 {a2} --collapse-conservatively --basename {params.out} --gzip

As we sequence 100 bp reads, but, due to the characteristic short fragments of ancient DNA, we expect to sequence into the adapter sequence at the 3' end of the reads. As such, it is imperative to remove these technical sequences from the reads before downstream mapping and analysis. We use AdapterRemoval (2.3.2)[105] to remove adapter sequences from the 3' end of the sequenced paired-end reads. We specify the adapter sequences to be the ends of the Illumina TruSeq adapters, before the index sequence, as shown in Table S2.1. This is to allow variations in index sequence lengths.

**Table S2.1 Adapter sequences used by AdapterRemoval**

| | **Adapter 1** | **Adapter 2** |
|---|---|---|
| **Double-stranded libraries (and Santa Cruz Reaction)** | AGATCGGAAGAGCACACGTCTGAACTCCAGTCA | AGATCGGAAGAGCGTCGTGTAGGGAAAGAGTGT |
| **Single-stranded (ss2.0) libraries** | AGATCGGAAGAGCACACGTCTGAACTCCAGTCA | GGAAGAGCGTCGTGTAGGGAAAGAGTGT |

As reads below 30 base pairs are very problematic to align, they are discarded at this stage using the –min-length parameter. Here, paired-end reads with an overlap of 11 base pairs or greater were collapsed into longer consensus sequences using the –collapse-conservatively parameter, which achieves an increase in the length and the quality of the insert sequences. This generates three fastq files per library per lane: pair 1, pair 2 and collapsed reads.

### Mapping

```
bwa aln -l 512 -t {threads} {REF} {input.fq} > {output.sai}
```

We then map each of the generated fastq files against the human reference genome build 37 (hs37d5, md5sum 12a0bed94078e2d9e8c00da793bbc84e) using bwa aln (*0*.7.17)[94]. Here, we disable seeding (-l 512) and instead perform end-to-end alignment, as suggested for ancient samples[106] in order to maximise the number of reads mapping to the human genome. As each bwa aln step is an independent process, it happens in parallel for all trimmed fastqs in a given sequencing run.

```
bwa samse -r \"{params}\" {REF} {input.sai1} {input.fq1} > {output.sam}
```

```
bwa sampe -r \"{params}\" {REF} {input.sai1} {input.sai2} {input.fq1} {input.fq2} > {output.sam}
```

Next, when all alignments are completed, we use bwa sampe and samse (0.7.17)[94] on the paired- and single-end fastq and sai files respectively. At this step, we set the RG tags of the resulting sam files. The ID tag is set to the fastq name plus whether the reads were paired or collapsed during adapter trimming, the SM tag is a unique sample identifier, and the LB tag is the library barcode. This allows accurate marking of duplicates during downstream analyses.

```
samtools sort {input.cram} -@{threads} -m4G > {output.bam}
```

This produces sam files, which are then sorted and converted into bam files using samtools sort (*v1.10*)[94]. Lastly, paired- and single-end reads for each library and lane are merged together, such that there is one resulting bam file per library.

```
samtools merge {output.merged} {input.bams} -@{threads}
```

#### Duplicate marking

```
java -Djava.io.tmpdir={params.tmpdir} -XX:ParallelGCThreads={threads} -Xmx2g -jar {picard.jar} MarkDuplicates OPTICAL_DUPLICATE_PIXEL_DISTANCE=12000 I={input} o={output} REMOVE_DUPLICATES=false METRICS_FILE={params.metrics} TAGGING_POLICY=All VALIDATION_STRINGENCY=LENIENT
```

The library construction and sequencing processes create duplications of reads in libraries. Such duplicate reads add no extra information to the downstream analyses, and as such should be discarded. Duplicates were marked using Picard MarkDuplicates (2.25.0) with a pixel distance of 12000. For each duplicate read, this updates the flag to include the bitwise flag 1024, indicating that the read is a duplicate. These reads are not removed at this stage, but setting this flag prompts downstream software to disregard them.

#### Library complexity

```
decluster/decluster {input} -w -p 12000 -@{threads} -o {params.out}
```

For all screening libraries we extrapolated the possible yield from additional sequencing experiments. The approach is using a mark-and-recapture technique originally implemented in preseq[107] and works by downsampling the histogram of the PCR clonality multiplicities. The NovaSeq 6000 platform is, however, using a patterned flow cell technology, and depending on library load it will generate a non-negligible amount of cluster duplicates. The preseq software is therefore not directly applicable to our sequencing data. Existing tools for removing these cluster duplicates from single-end data are kmer based[108] and very memory-intensive. Therefore, we developed a program, decluster, that constructs the PCR duplication table from the mapped data (https://github.com/ANGSD/decluster). Using the output of decluster, and information about how many reads were sequenced, we project how many more reads would be needed to reach a given level of coverage and use this information to inform further sequencing.

366

367

### S3. DNA authentication

*Abigail Ramsøe, Thorfinn Sand Korneliussen*

In order to ensure the authenticity of the sequenced data, we employed three approaches: mapDamage to quantify the extent of deamination, and two contamination estimates. These contamination estimates were first run on a library-level basis, and any contaminated libraries were excluded. When the different libraries for a sample were merged, the authentication methods were then rerun to ensure that there was still no contamination flagged. This strategy ensures the high-confidence detection of both ancient and modern human contamination.

All analysis was performed on filtered files with reads with a base quality of at least 20, and a mapping quality of at least 30. For ANGSD, we removed the pseudoautosomal regions on the X chromosome.

#### mapDamage

```
mapDamage -i {input.bam} -r {REF}
```

In order to determine the authenticity of the ancient reads, the extent of the characteristic C->T deamination in libraries was quantified using mapDamage2.0[92]. Libraries were flagged for manual interpretation if the C->T extent at the 5' end, and/or G->A' extent at the 3' end did not follow expected parameters given the library type. mapDamage was also run on USER treated libraries, where we do not expect to see the ancient damage signal. Here, libraries were flagged if they exhibited the elevated C->T, as this could indicate a sample swap, or contamination from an ancient source.

### ContamMix

```
Rscript contamix/estimate.R --samFn ${base}_ra.final.bam --malnFn ${base}_312.aligned.fasta --figure ${base}_contam.pdf --trimBases 7 > ${base}.summary.txt
```

Firstly, we applied ContamMix in order to quantify the fraction of exogenous content in the set of reads mapping to the mitochondrial genome by comparing the mtDNA consensus genome to 311 possible contaminant genomes[109]. We only considered sites with at least 5x coverage and 70% base concordance (Supplementary Table S1.1).

### S4. Imputation

We generated genomes greater than 0.01X for 729 pieces of skeletal material. In 31 instances, skeletal material presumed to be from different individuals were genetically identical to another individual (Supplementary Note S5.6). After merging data for samples that were genetically identical, we were left with 14 unique individuals. In Supplementary Table S1.1, we report statistics for the all 729 pieces of skeletal material as well as the 14 merged sets. Merged data is indicated by an underscore joining the individual sample IDs (for example, the three samples CGG105916, CGG105917 and CGG105918 were merged as a single individual: CGG105916_CGG105917_CGG105918). There are therefore 4 lines for this one individual, so the Museum IDs for each can be traced. From the 729 pieces of skeletal material, the number of unique individuals that we have data for is 712.

Imputation was carried out on 4,009 already published ancient genomes[17,23,26,28,31,32,32,33,42–45,58,78,79,95,110–189] along with the newly generated genomes using GLIMPSE (v1.1.1)[96], as described in Allentoft et al.[28] and Sousa da Mota, B. et al.[37]. As a reference panel, we used the 30X coverage version of the 1000 Genomes (1000G) v5 phase 3 dataset[190] that was phased using the TOPMed panel[191], before being lifted over to hg19.

Whole-genome shotgun-sequenced genomes and genome-wide targeted capture (‘1240k’) enriched for 1.24 million sites[192] were included in this study. Studies have shown for this capture panel that off-target sites impute poorly even at higher coverages[193], so all off-target sites were excluded for all samples. For the 1240k on-target sites, imputation performs similarly for genomes of 0.1X-0.5X for shotgun and 1X for capture. Previous studies have demonstrated that shotgun samples at 0.1X are suitable for the same methods of demographic inference, even when applied on (Mesolithic and Neolithic) populations that are much more distantly related to the 1000G reference panel than the more recent samples that are the focus of this study[28,29]. As such, coverage cut-offs were set at 0.1X for shotgun-sequenced, and 1X for captured genomes. After this initial filtering, for each sample, the average genotype probability was calculated for all captured sites, and samples with a low average genotype probability (<0.90) were excluded (Fig. S4.1).

Of the new 712 unique individuals, 134 were not included in the clustering, due to low coverage or low gpAverage (n=93), being the lower coverage of a pair of close relatives in the dataset (n=35) or being contaminated (n=6) (See 'flags' column in Supplementary Table S1.1). These 134 individuals are included in the PCA projections in Supplementary Note S5.1.2., kinship information is provided in Supplementary Note S5.6., and contamination and coverage statistics can be found for merged and unmerged data in Supplementary Table S1.1. The remaining 578 new individuals were included in the final dataset of 4,587 samples for all IBD analyses.

The intersection of the 1.24 million capture sites and imputed 1000G sites left 1,085,103 SNPs. From there we kept sites with an imputation info score >0.5 and only considered the highly mappable regions in the genomes defined by the 1000G strict mappability mask, leaving the final number of SNPS at 690,211.

In comparison, a study with a similar processing pipeline that only included shotgun-sequenced ancient genomes resulted in a panel with over 10 times as many SNPs: 7,321,965[28]. However, for many regions and time periods of importance to this study, restricting to just capture samples would result in entire regions in time (e.g. Western Europe during the Bronze Age) and countries in general (e.g England and the Czech Republic) being under-represented (Fig. S4.2). To address the research questions of the study, it was therefore necessary to restrict to the smaller SNP set.

Metadata for the samples was curated from the respective papers and from Mallick et al.[194].

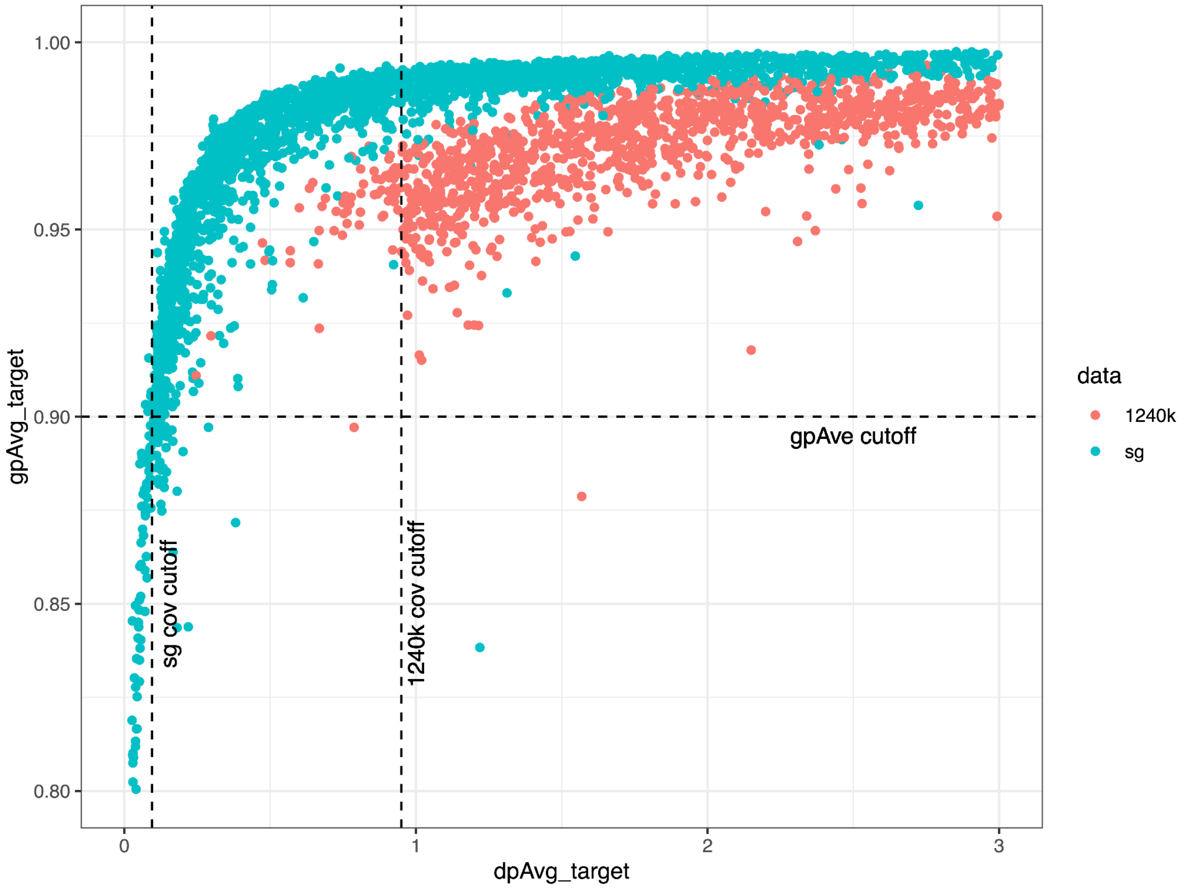

**Fig. S4.1. Relationship between the average genotype probability (gpAvg_target) and sequencing depth (dpAvg_target) for captured (1240k) and shotgun (sg) genomes, when restricting to target sites.**

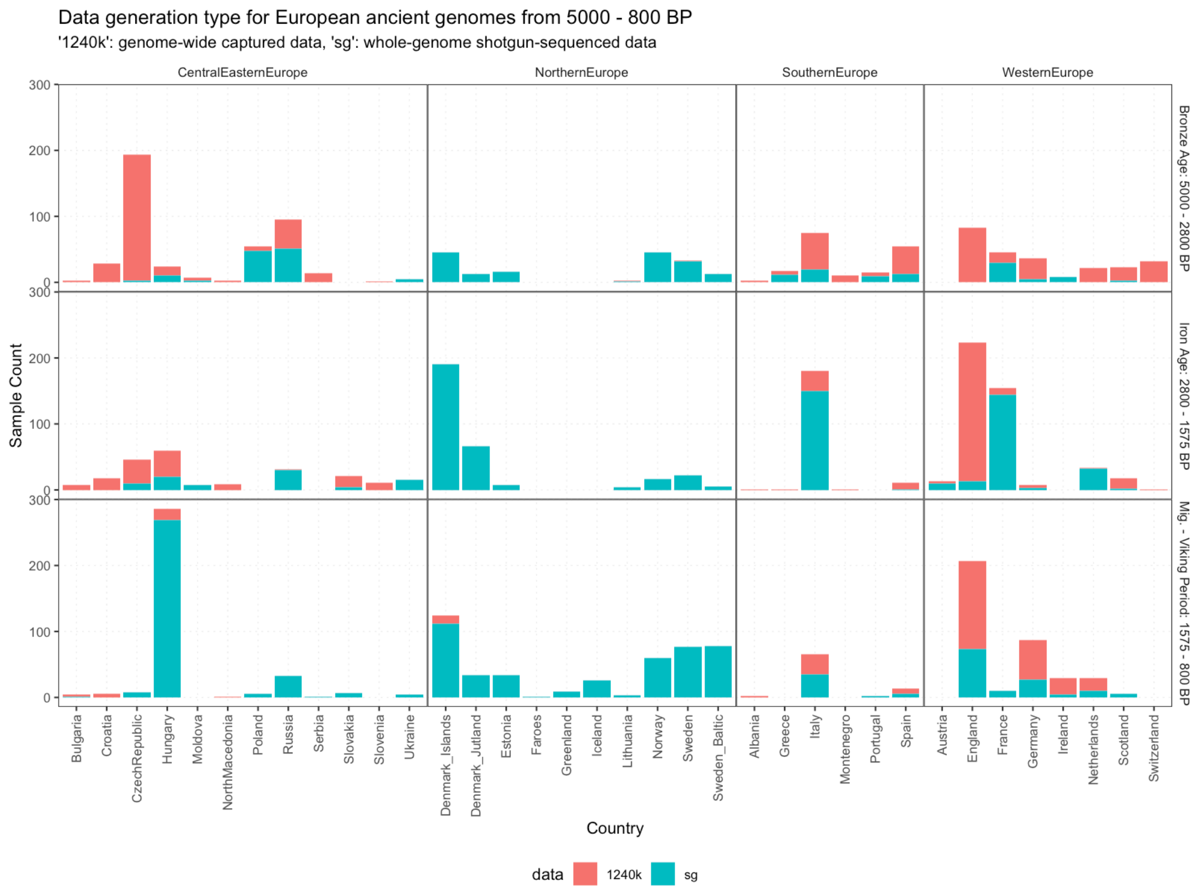

**Fig. S4.2. Geographical and temporal distributions of ancient genomes in the final imputed dataset for each data generation method (1240k capture and shotgun-sequencing).**

### S5. Population Genetics

*Hugh McColl, Frederik Valeur Seersholm, Thomaz Pinotti, Tharsika Vimala, J. Víctor Moreno-Mayar, Martin Sikora*

#### S5.1. PCA

##### S5.1.1. PCAs of clustered individuals

To infer basic demographic structure in our data, we carried out a principal component analysis (PCA) on the imputed dataset of 4587 individuals covering 690211 SNPs (see Supplementary Note S4) using GCTA (v1.94.1). To better visualise IBD clusters and finer-scale patterns in the data, we plotted two panels of decreasing size and diversity, the first containing all 'out-of-Africa' populations, (n=4,495) and the second focussing on Western Eurasia (n=3,870). As depicted in Fig. S5.1, the vast majority of samples from this study (highlighted with a black circle) fall within the European Bronze Age diversity, while a small number of samples display varying levels of Asian admixture.

Colouring the individuals in the PCA by their IBD clusters (Supplementary Note 6.4) shows a correlation between established clines in PCA space and the IBD clustering (Fig. S5.1). For each of the major four clusters discussed in the main text (Yamnaya, Corded Ware (East), Corded Ware (North), Bell Beaker), a PCA highlighting their subclusters has been plotted (Fig. S5.2- S5.5)

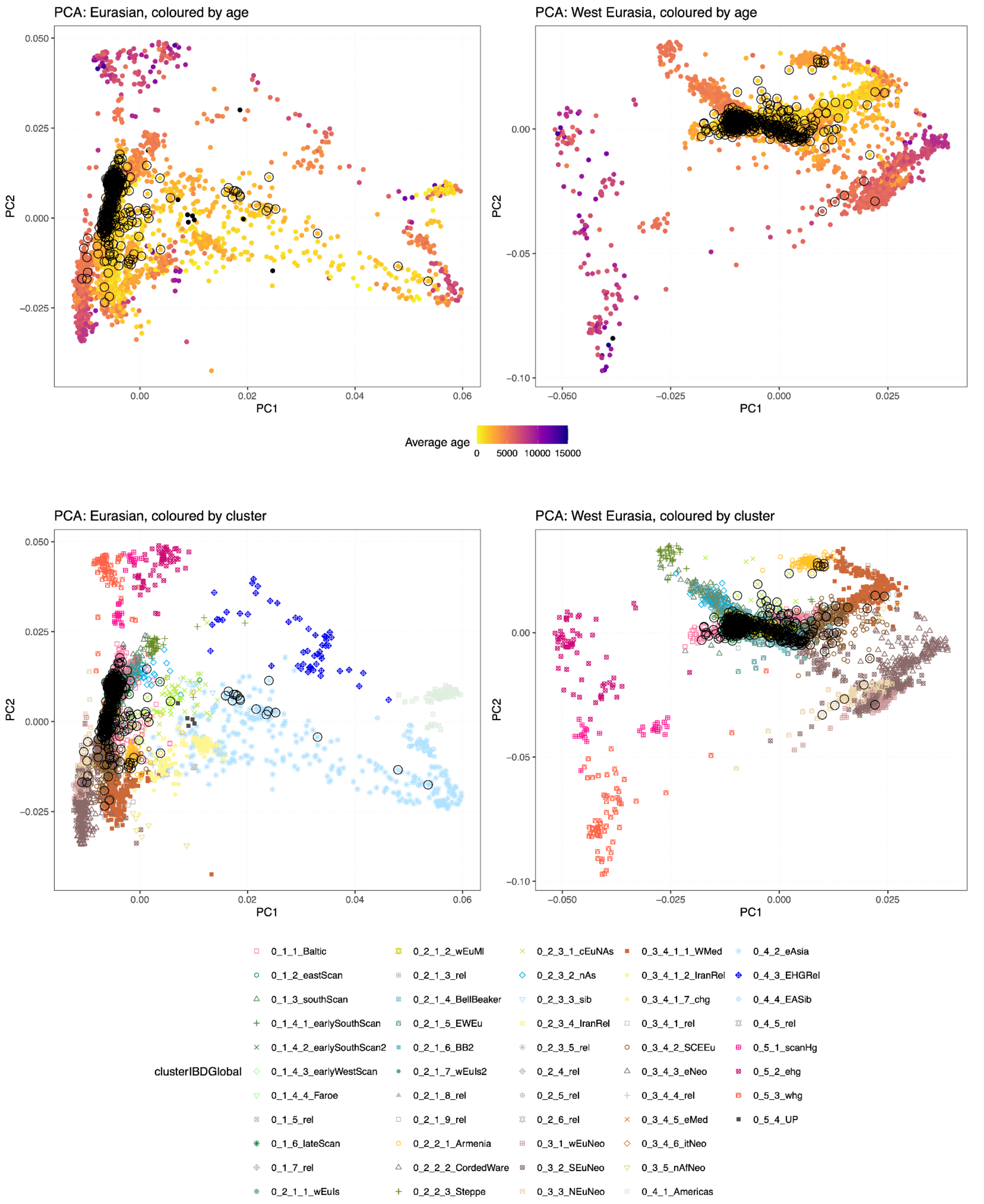

**Fig. S5.1. PCA of Eurasian and Western Eurasian ancient populations, coloured by age and by IBD cluster.** Circled individuals are newly generated in this project.

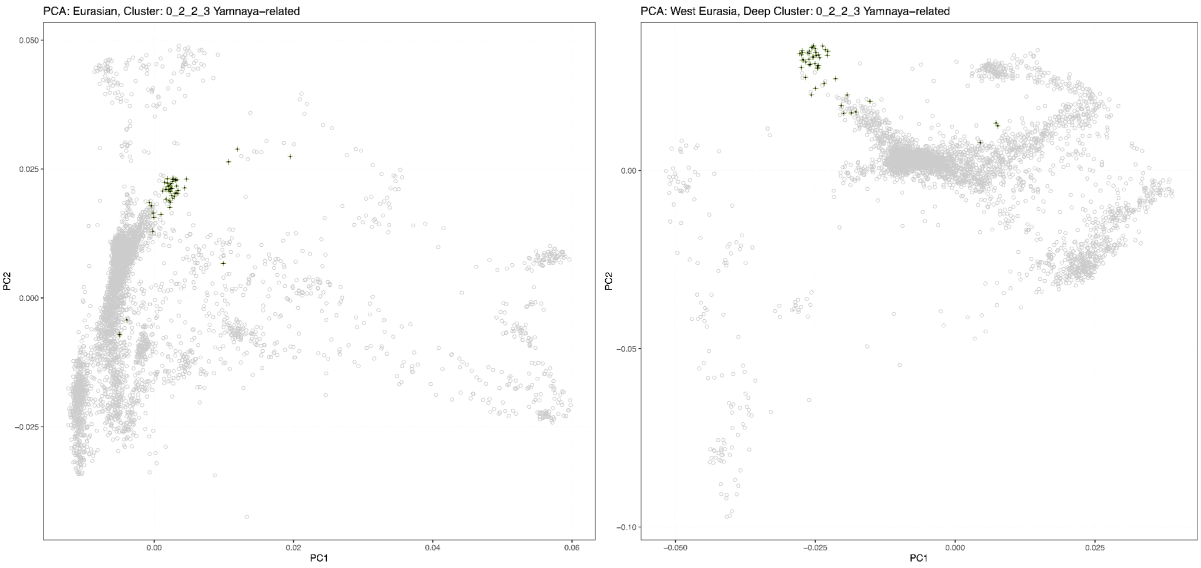

**Fig. S5.2. PCA of Eurasian and Western Eurasian ancient populations highlighting the Yamnaya-related cluster.** Individuals older than 2800 BP are indicated with a '.'

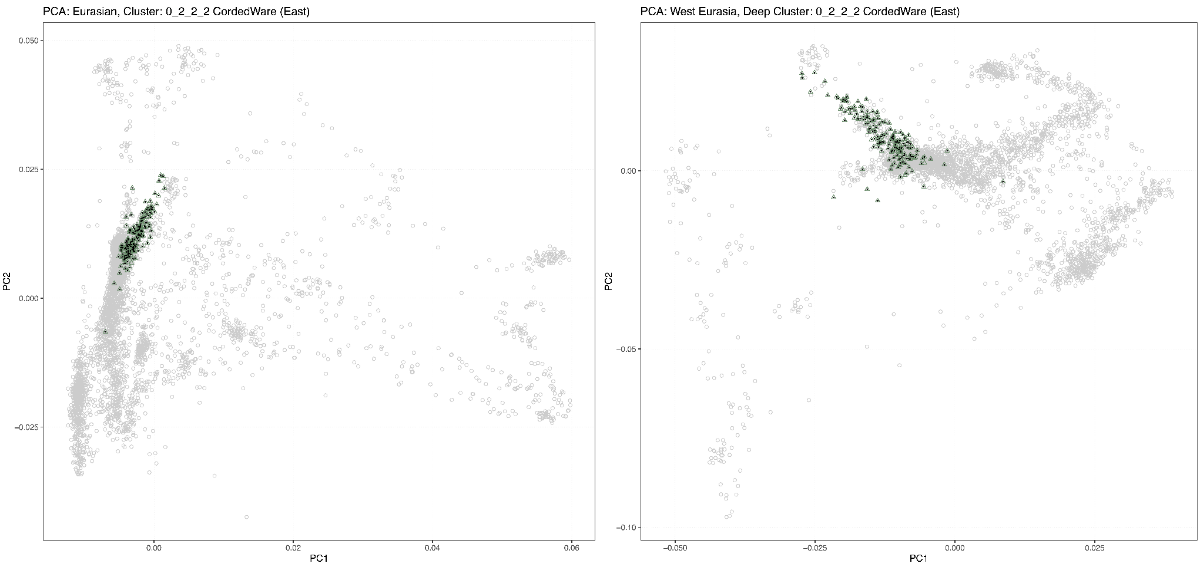

**Fig. S5.3. PCA of Eurasian and Western Eurasian ancient populations highlighting the Corded Ware (East)-related ancestry.** Individuals older than 2800 BP are indicated with a '.'

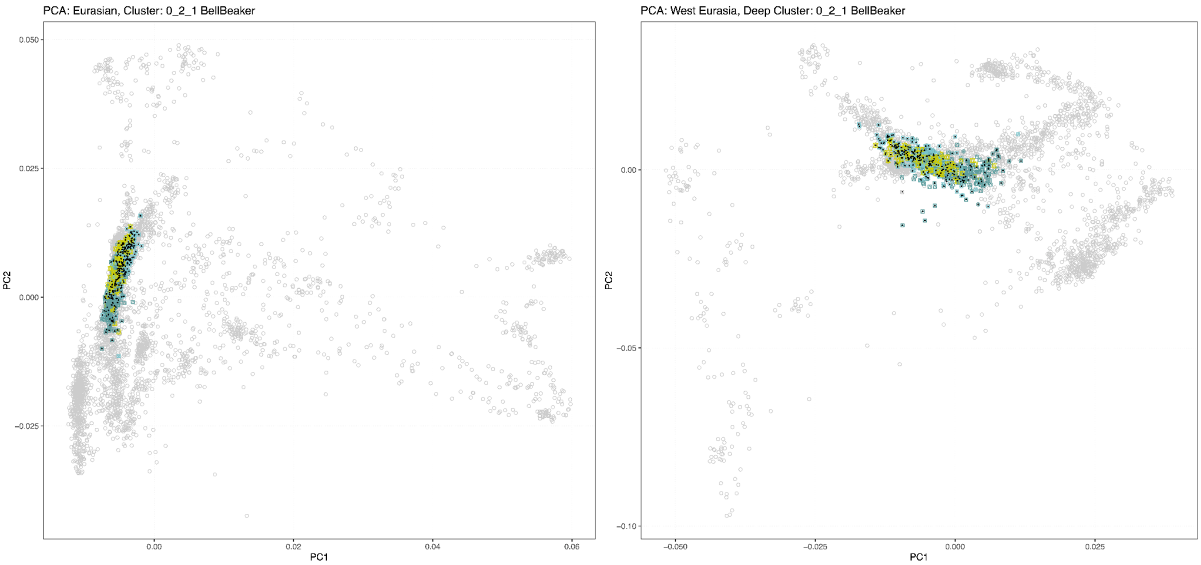

**Fig. S5.4. PCA of Eurasian and Western Eurasian ancient populations highlighting the Bell Beaker-related ancestry.** Individuals older than 2800 BP are indicated with a '.'

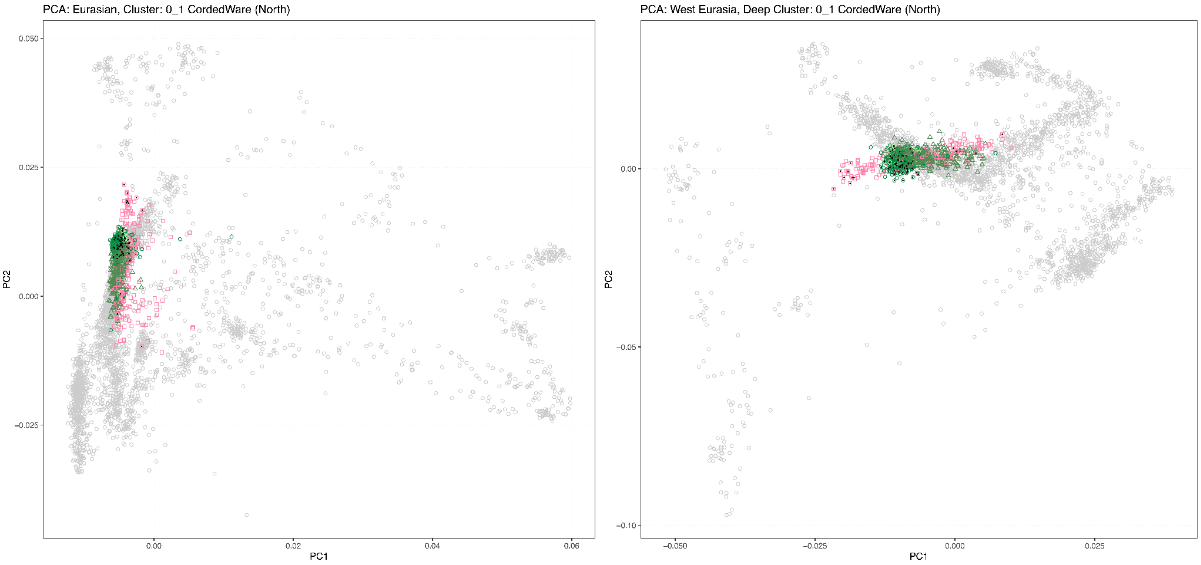

**Fig. S5.5. PCA of Eurasian and Western Eurasian ancient populations highlighting the Corded Ware (North)-related ancestry.** Individuals older than 2800 BP are indicated with a '.'

#### S5.1.2. PCA projection of contaminated, related and low gpAverage/depth individuals

Of the 751 genomes above 0.01X originally presumed to be from unique individuals, 174 were not included in the clustering. This was a result of contamination, low average GP values/low coverage, or high IBD sharing as a result of duplicate samples or closely related individuals. To recover additional information on these non-clustered samples, we projected pseudo-haploid data for these individuals on subsets of the clustered individuals using smartpca[195]. The same set of 690211 SNPs as described in S5.1.1 was used, using the default settings and lsqproject set to YES.

The first PCA (Fig S5.6) includes 'Out-of-Africa' populations, allowing individuals with East Asian ancestry to be detected. The second PCA (Fig S5.7) shows diversity within Western Eurasia, allowing individuals of farming ancestry to be distinguished from later populations. The third PCA (Fig S5.8) allows for Bell Beaker-related populations to be distinguished from Corded Ware-related populations. The fourth PCA (Fig S5.9) allows for Baltic-related populations to be distinguished from Scandinavian populations. The fifth PCA (Fig S5.10) allows for Western, Southern and Eastern Scandinavian populations to be distinguished from each other.

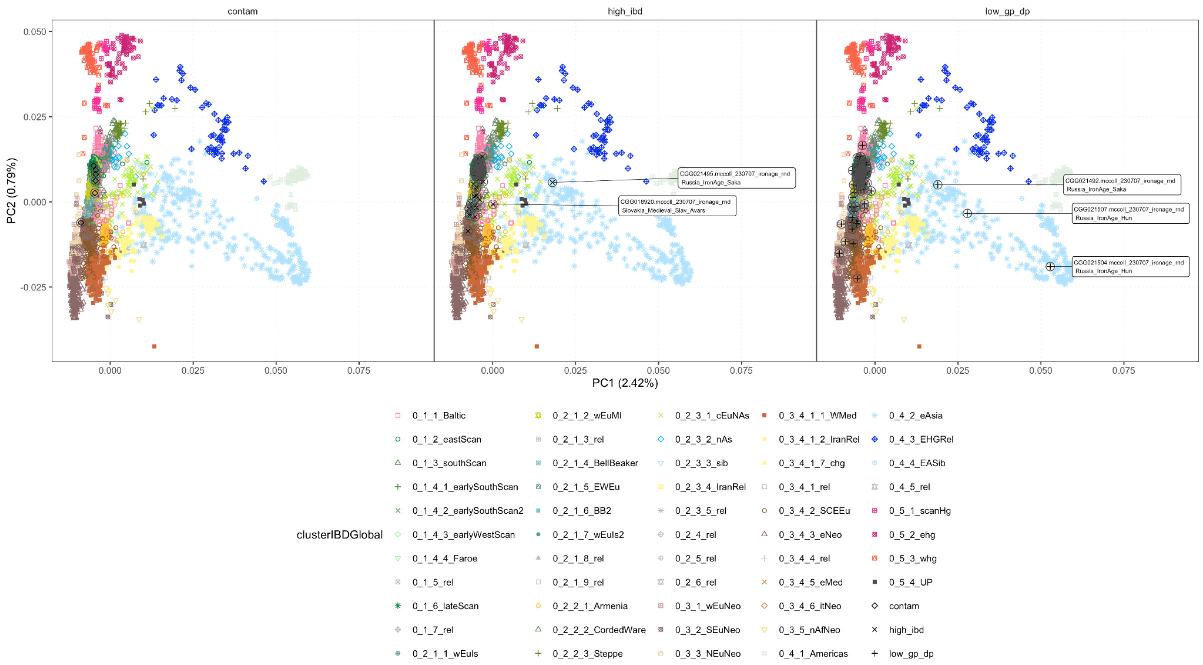

**Fig. S5.6. PCA projections of contaminated, high IBD sharing and low gpAverage or sequencing depths individuals on PCAs of ‘Out-of-Africa’ clustered ancient individuals**

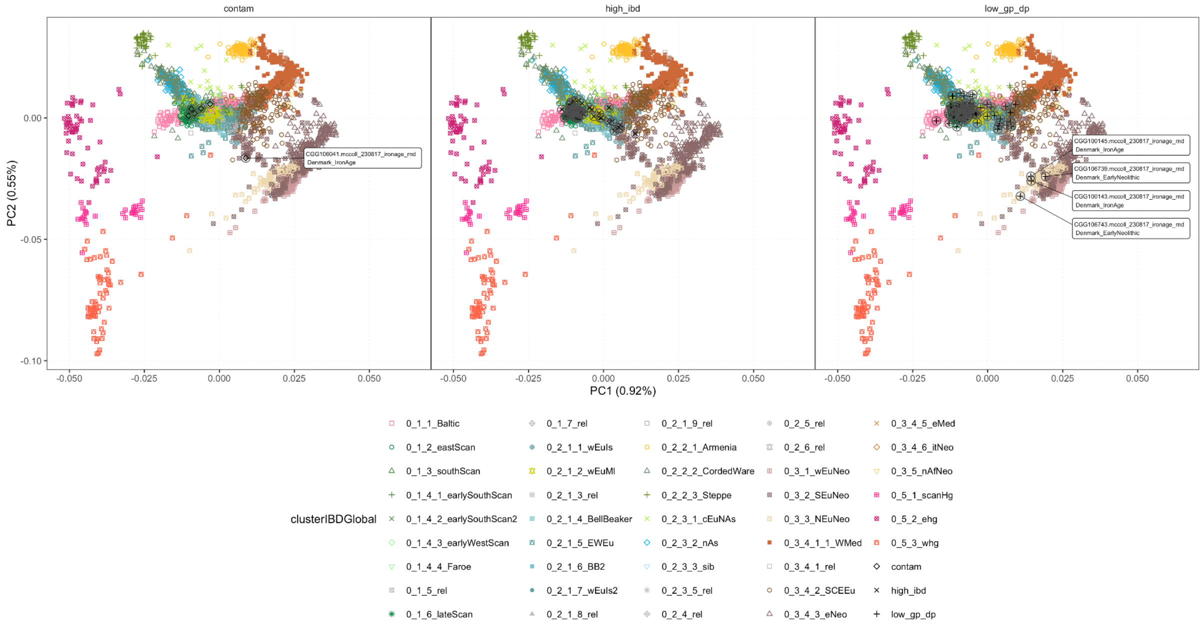

**Fig S5.7. PCA projections of contaminated, high IBD sharing and low gpAverage or sequencing depths individuals on PCAs of clustered ancient individuals from Western Eurasia.** All projected samples are circled, labelled samples are removed from subsequent projections.

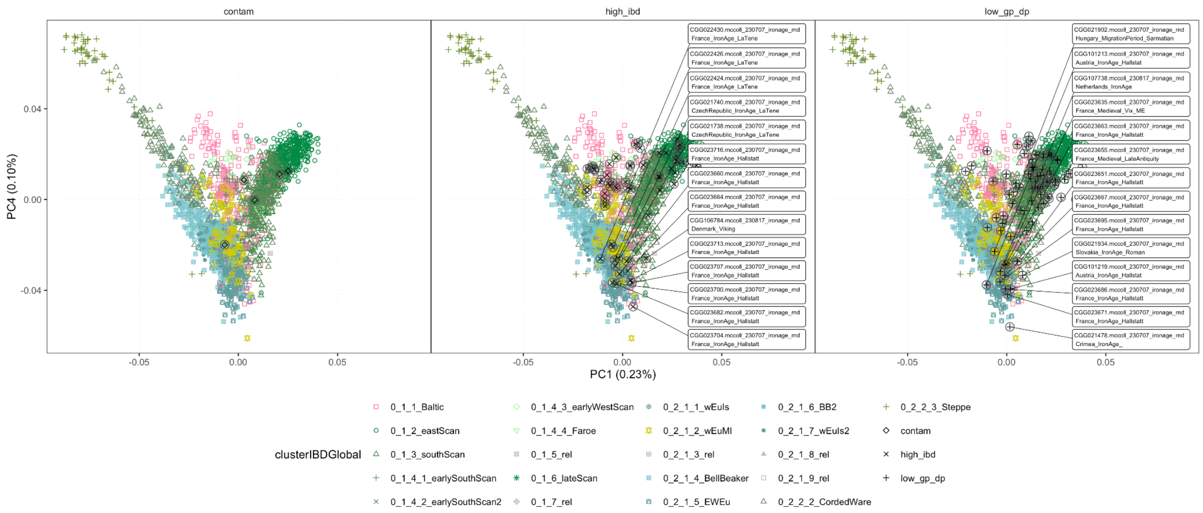

**Fig S5.8. PCA projections of contaminated, high IBD sharing and low gpAverage or sequencing depths individuals on PCAs of clustered ancient individuals from the Steppe and Corded Ware (North) and Bell Beaker clusters.** All projected samples are circled, labelled samples are removed from subsequent projections.

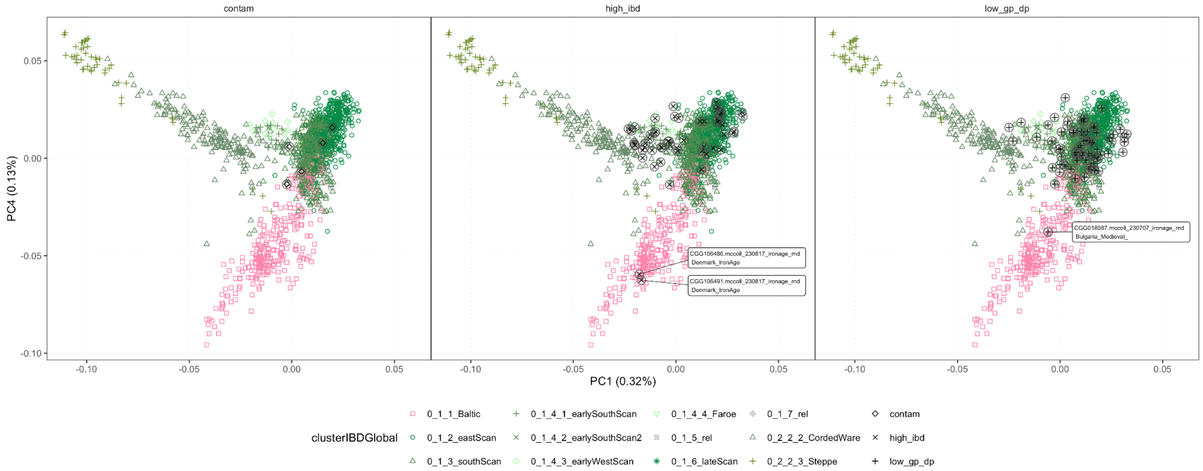

**Fig S5.9. PCA projections of contaminated, high IBD sharing and low gpAverage or sequencing depths individuals on PCAs of clustered ancient individuals from the Steppe and Corded Ware (North) clusters.** All projected samples are circled, labelled samples are removed from subsequent projections.

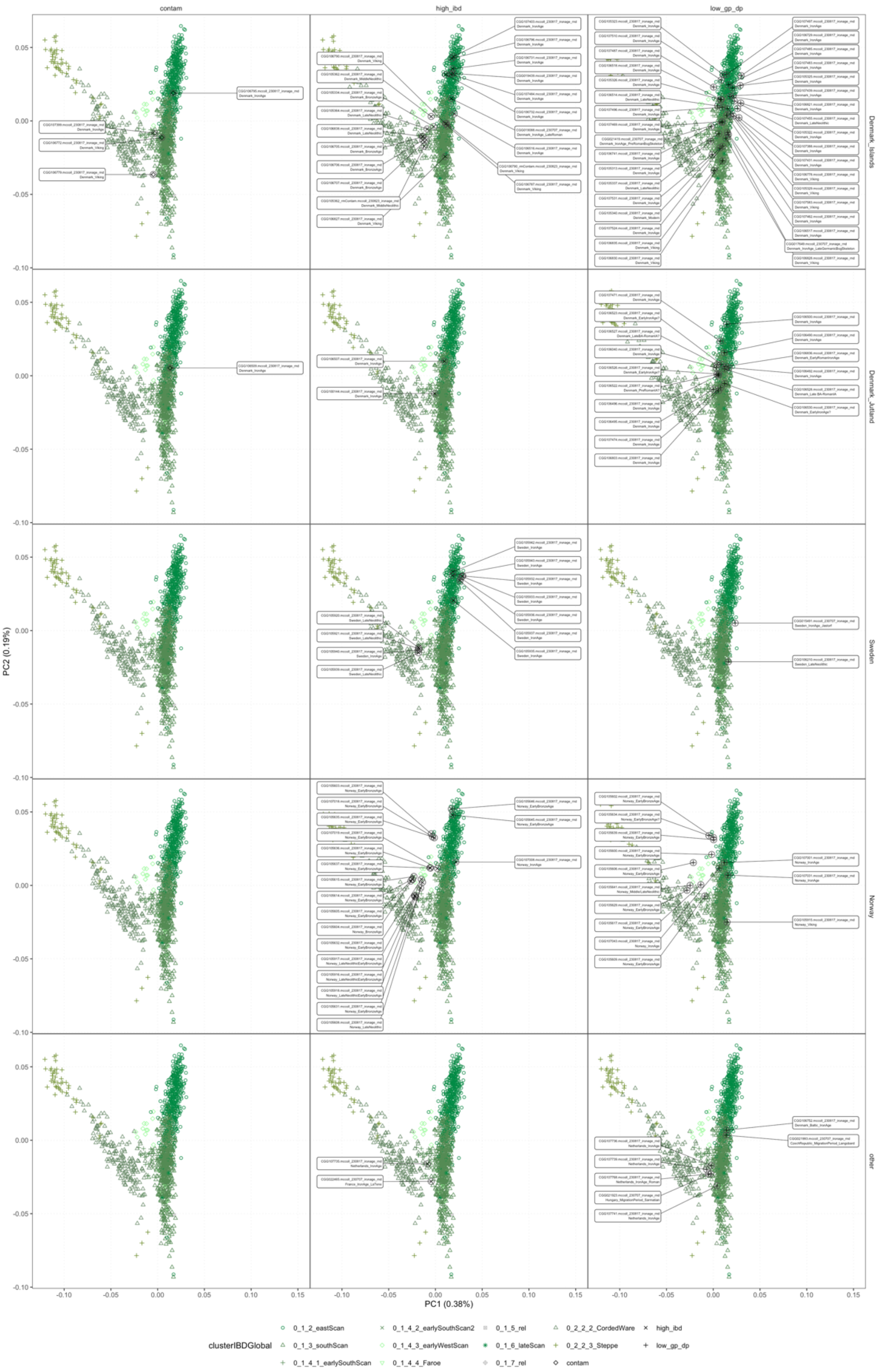

**Fig S5.10. PCA projections of contaminated, high IBD sharing and low gpAverage or sequencing depths individuals on PCAs of clustered ancient individuals the Steppe and Scandinavian Clusters.** All projected samples are circled.

#### S5.2. IBD clustering

##### S5.2.1. Whole dataset clustering

To identify IBD segments, we ran IBDseq[98] on the imputed panel. We removed any segments less than 2 cM, with a LOD score less than 3, and any hotspot regions with excess sharing across all individuals, as described in Allentoft et al.[28]. For all pairs, the total shared IBD length was summed and the number of IBD segments counted. We excluded any pairs sharing less than a total of 5 cM, and the lowest coverage of any pair sharing more than 1500 cM and had less than 150 segments, to avoid the formation of small clusters with only 1st and 2nd degree relatives. We then performed a network-based hierarchical clustering based on total shared IBD lengths[196], as described in Allentoft et al.[28].

By plotting sankey diagrams showing the relation between clusters and regions (Fig. S5.11, S5.12, S5.13), a correspondence between region and clusters is apparent. We note that almost all (81/107) individuals archaeologically associated with the Bell Beaker Complex fall within the 0_2_1_x clusters. Similarly 63/64 individuals associated with the Corded Ware Complex fall within the 0_2_2_2_x clusters and 13/15 individuals associated with the Yamnaya Culture fall within the 0_2_2_3_x clusters (see groupLabel and clusterIBDDeep columns in Supplementary Table S5.1). A fourth cluster, 0_1_x contains no individuals labeled in association with the Yamnaya, Corded Ware or Bell Beaker Cultures, however the Steppe ancestry of all Bronze Age (2800 BP and older) individuals is modelled by Corded Ware-related ancestry, rather than Bell Beaker-related (see 0_1_x clusters in Fig. S5.21, set 4).

We therefore refer to the 0_2_2_3_x clusters as Yamnaya-related, the 0_2_1_x clusters as Bell Beaker-related, 0_2_2_2_x as Corded Ware (East)-related (123/174 individuals are found in Central Eastern Europe), and 0_1_x as Corded Ware (North) related (66/72 individuals are found in Northern Europe).

Supplementary Table S5.1 contains the relevant archaeological (‘archGroup’ column), clustering (clusterIBDDeep column) and regional (‘region’ column) information for each ancient sample (sampleId column).

From the mixture modelling results discussed in Supplementary Note S5.3.1, a number of trends are apparent (Fig. S5.16, Fig. S5.23, Fig. S5.24), detailed below.

- Cluster 0_1 and subclusters (indicated by 0_1_…)
    - high Steppe ancestry and some Globular Amphora Culture (GAC) ancestry
    - found in Northern Europe
    - Steppe ancestry modelled by Corded Ware source, rather than Bell Beaker
- Cluster 0_2_1 and subclusters
    - high Steppe ancestry and some GAC ancestry and varying amounts of European Farmer ancestry
    - found in Western Europe
    - associated with Bell Beaker contexts.
- Cluster 0_2_2_2 and subclusters
    - high Steppe ancestry,
    - associated with Corded Ware contexts
- Cluster 0_2_2_3 and subclusters
    - highest Steppe ancestry, including Yamnaya, found in Western Eurasia
- Cluster 0_3 and subclusters

    - high Neolithic Farming ancestry
- Cluster 0_4 and subclusters
    - from across Asia and the Americas
- Cluster 0_5 and subclusters
    - Eastern and Western Hunter-Gatherer ancestry

From 5000 to 1575 BP, the Corded Ware (North) cluster was restricted primarily to Northern Europe, and the Bell Beaker cluster to Western Europe (Fig. S5.11). From 1575 BP onwards, the Northern Corded Ware cluster was more widespread. By using more fine-scale clusters (Fig. S5.12), it is apparent this spread is primarily by the Southern Scandinavian cluster. When replacing regions with countries (Fig. S5.13), the widespread appearance of Southern Scandinavian in England, Germany and the Netherlands is apparent. From 5000 to 2800 BP, individuals from these countries were primarily from Bell Beaker-related clusters.

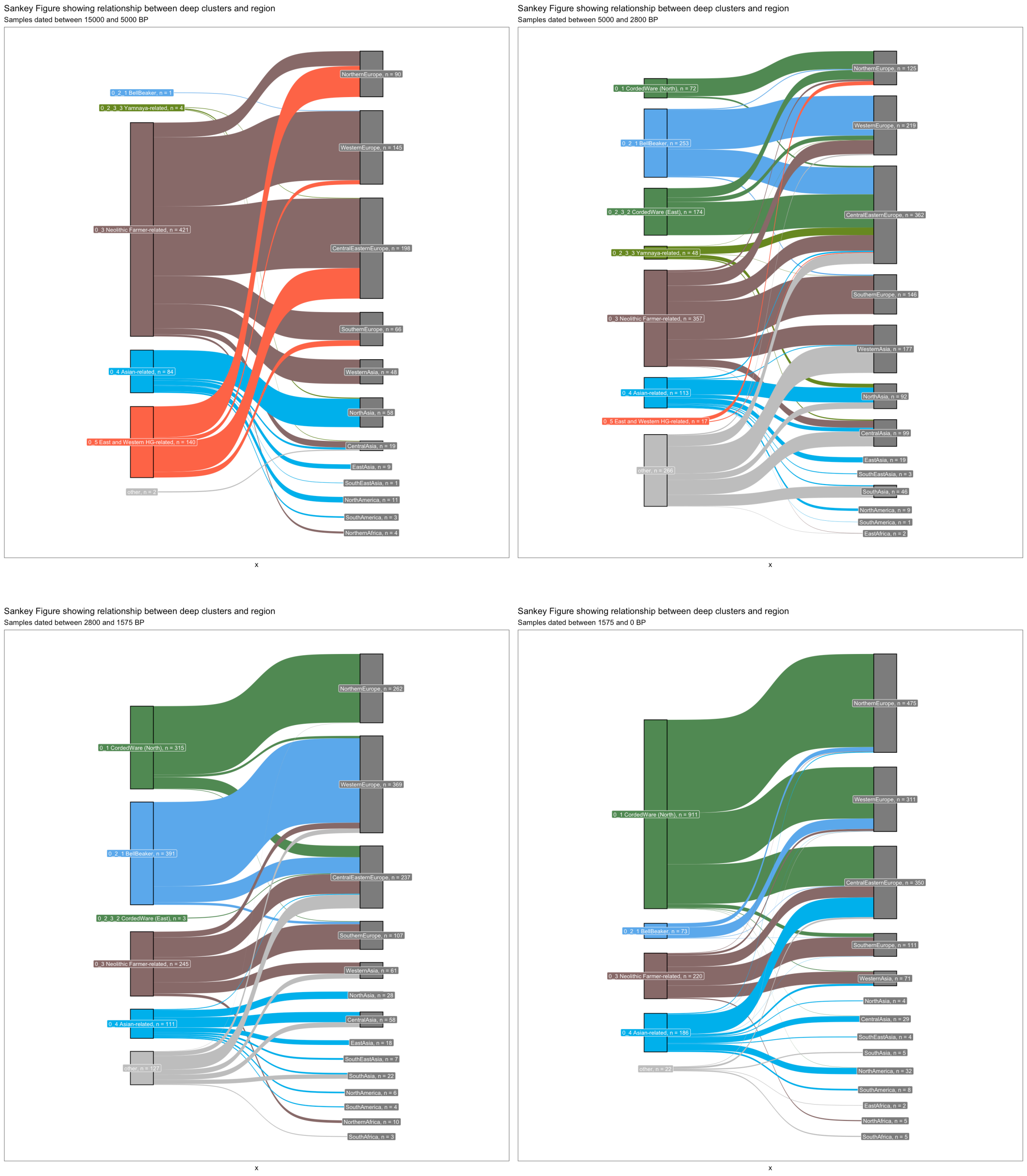

**Fig. S5.11. Sankey diagram showing the relationship between the deepest IBD clusters and geographical regions around the world, in different time bins (15000–5000 BP, 5000–2800 BP, 2800–1575 BP, 1575–0 BP).**

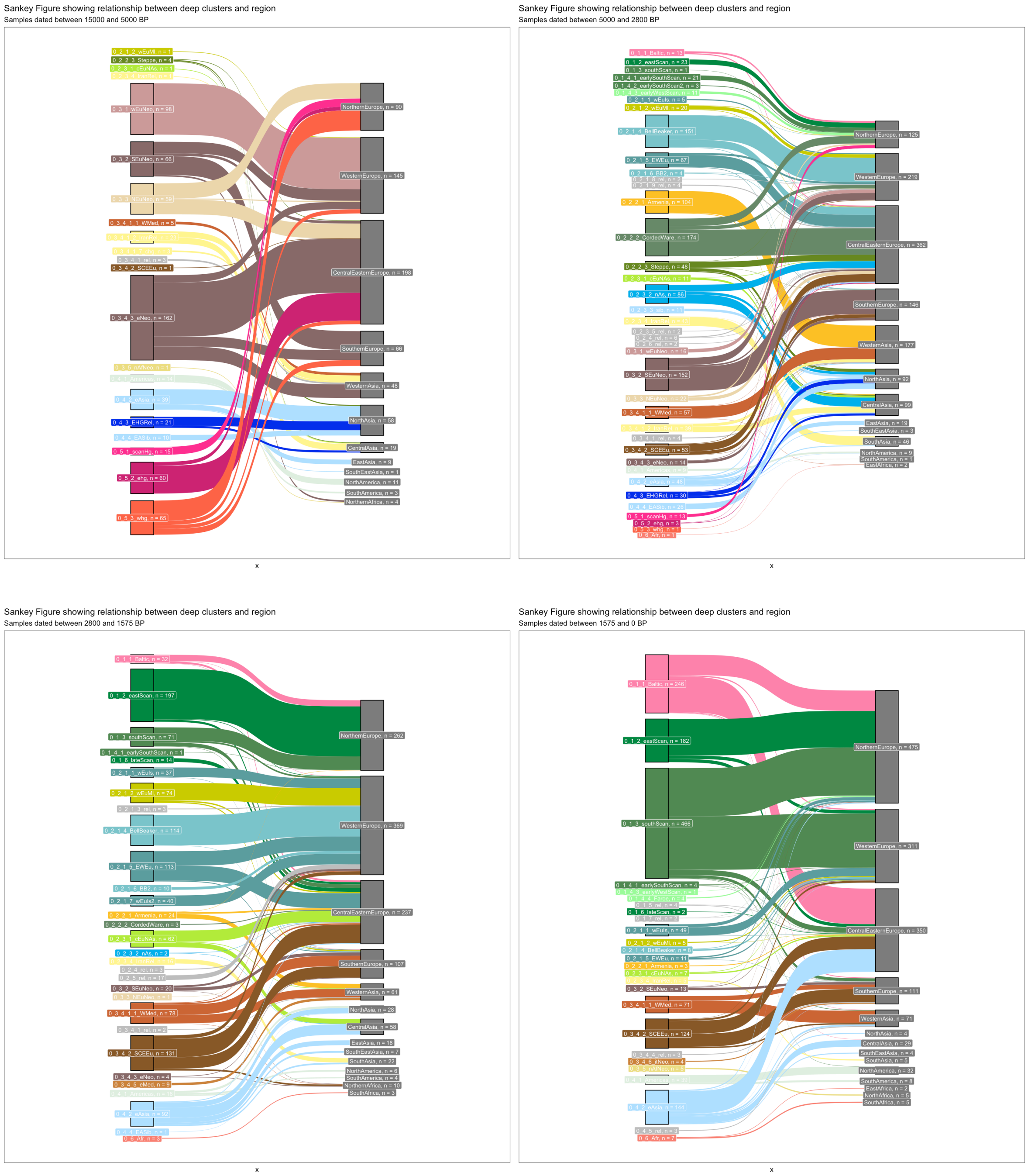

**Fig. S5.12. Sankey diagram showing the relationship between the deep IBD clusters and geographical regions around the world, in different time bins (15000–5000 BP, 5000–2800 BP, 2800–1575 BP, 1575–0 BP)**

Sankey Figure showing relationship between deep clusters and country
Samples dated between 15000 and 5000 BP

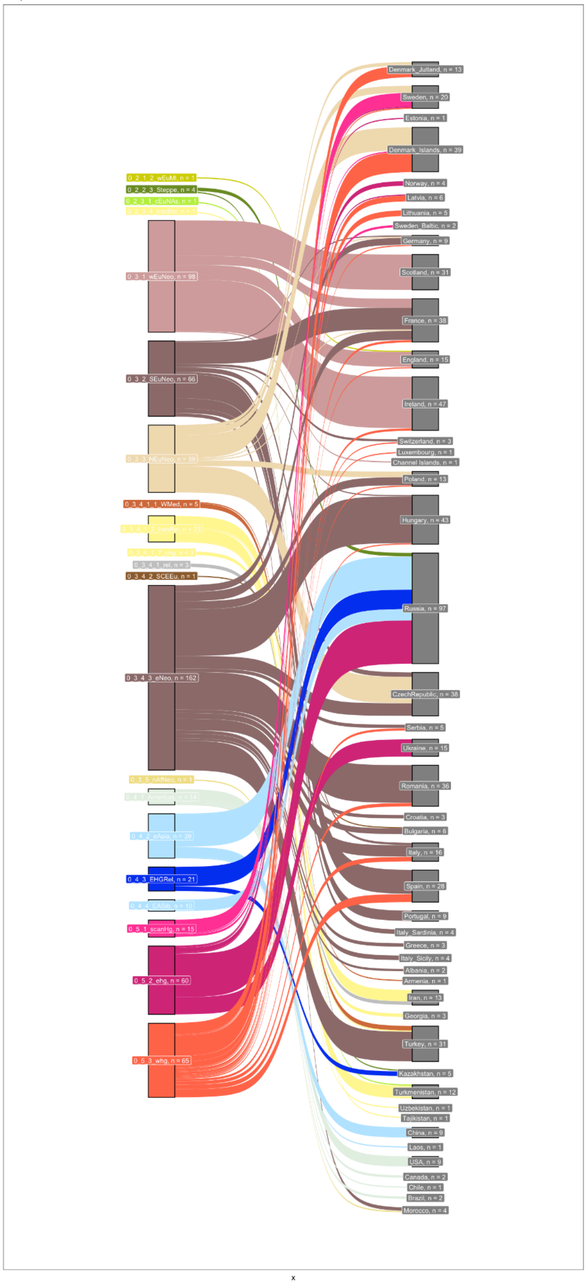

Sankey Figure showing relationship between deep clusters and country
Samples dated between 5000 and 2800 BP

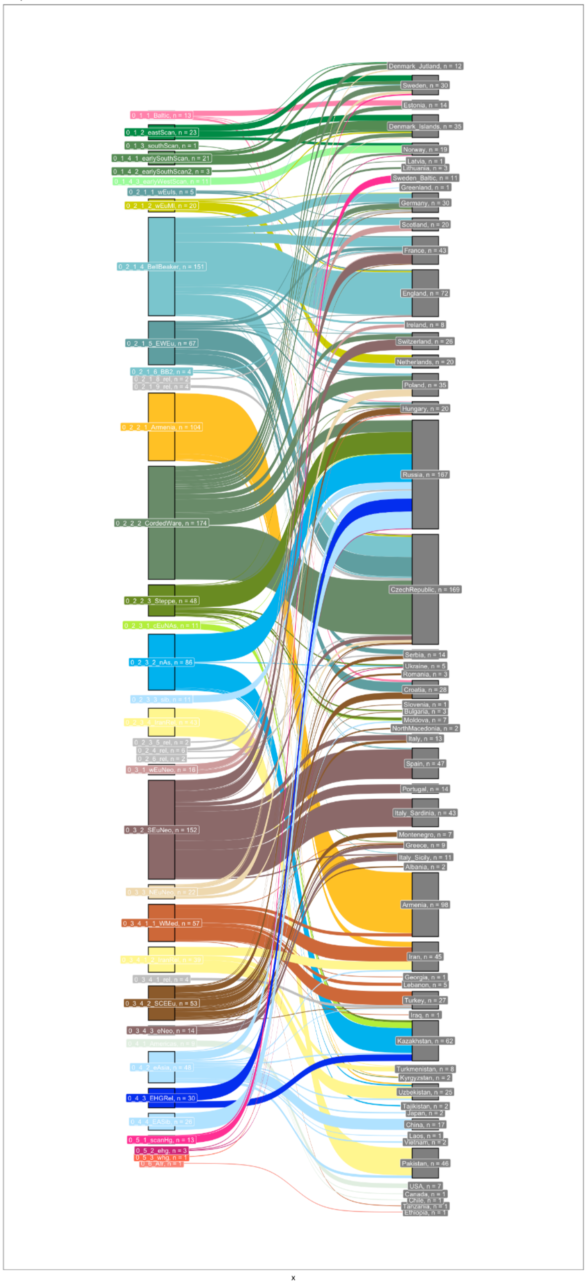

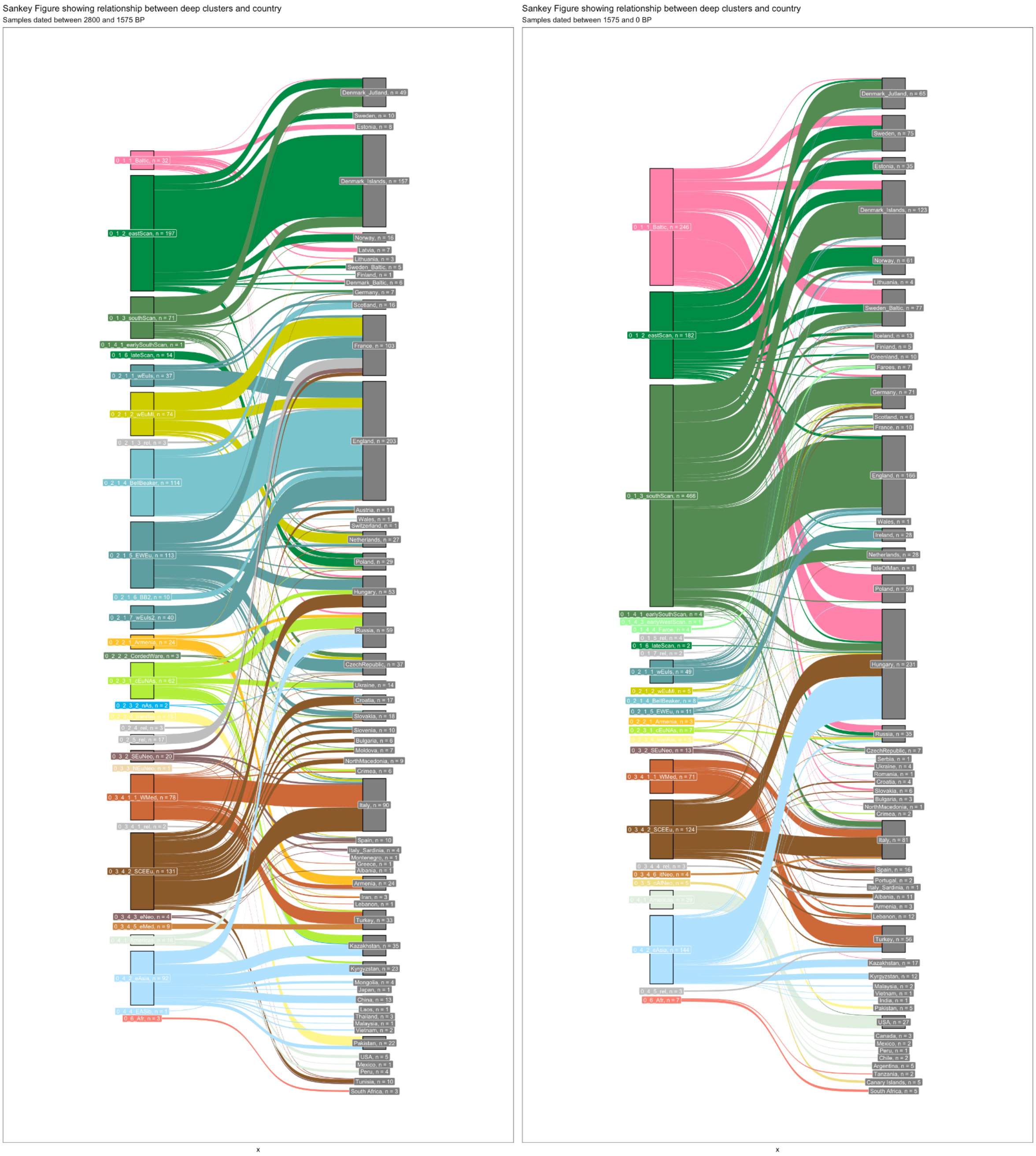

**Fig. S5.13. Sankey diagram showing the relationship between the deepest IBD clusters and countries around the world, in different time bins (15000–5000 BP, 5000–2800 BP, 2800–1575 BP, 1575–0 BP)**

#### S5.2.2. 2800BP+ Dataset clustering

**Rationale**

To get an understanding of the Bronze Age structure without having more recent admixed individuals influencing the clustering, we excluded all individuals dated to less than 2800 BP (pre2.8k) and reclustered the dataset as in S5.2.1, (Table S5.2). As the reclustering included samples from across all of Europe, we used a more general date of 2800 BP for the transition from the Bronze Age to the Iron Age in Europe, rather than the usual date of 2450 BP for Northern Europe.

**Results**

A clear difference between the two clustering runs is apparent in the reduction from 173 to 122 individuals in the Corded Ware (East) cluster in the pre-2800 BP clustering (Fig. S5.14). Many of these individuals are instead now found in clusters from Northern Europe in the pre-2800 BP clustering. The pre-2800 BP Early Scandinavian cluster contains 24 individuals rather than 11 in the corresponding Western Scandinavian cluster in the full clustering. Similarly, the pre-2800 BP Southern Scandinavian cluster sees 37 individuals rather than 21. From mixture modelling results (Fig. S5.15), we see that the samples that moved from the Corded Ware (East) cluster in the original modelling into the Corded Ware (North) clusters are modelled with only small amounts of smallest amounts of the Eastern, Western and Southern Scandinavian ancestry that is widespread from the late Bronze Age onwards. Instead, these early individuals are modelled with a large amount of Corded Ware Ancestry. However, from around 4000 BP, almost all Scandinavians are well-modelled as combinations of Eastern, Western and Southern Bronze Age ancestries. Combined, the results suggest a structured population in Scandinavia present from ~4600–4000 BP.

Of particular note, we find in the pre2.8k the Bronze Age Norwegians cluster together with the earliest Swedish and Danish individuals (Extended Data Fig. 9: pre2.8k_0_1_3_3-EarlyScan cluster), many of which previously fell in the Corded Ware (East) subcluster, and who are predominantly carriers of the R1a Y-haplogroup.

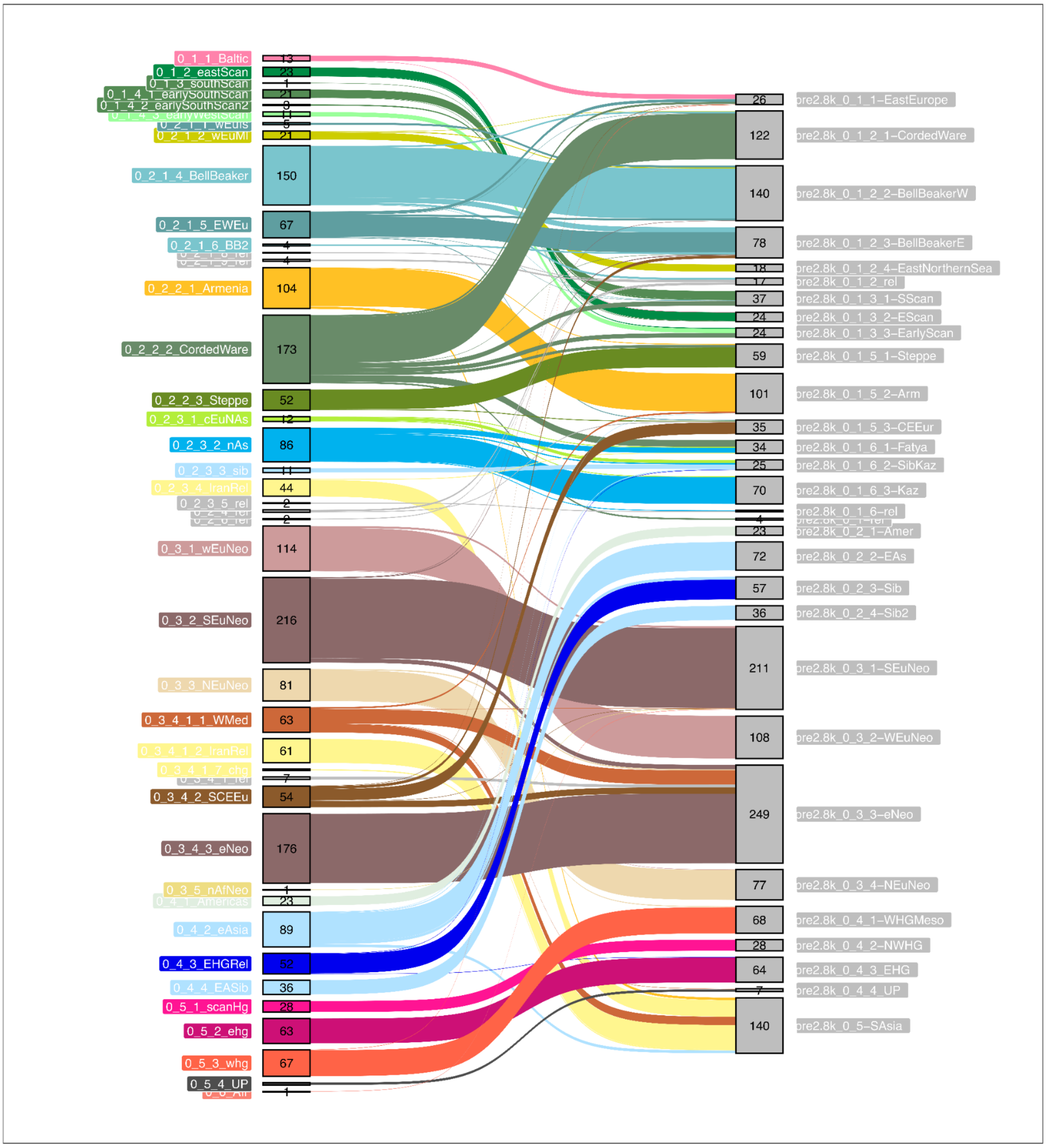

**Fig. S5.14. Sankey diagram showing relationship between full clustering (left) and pre2.8k clustering (right).**

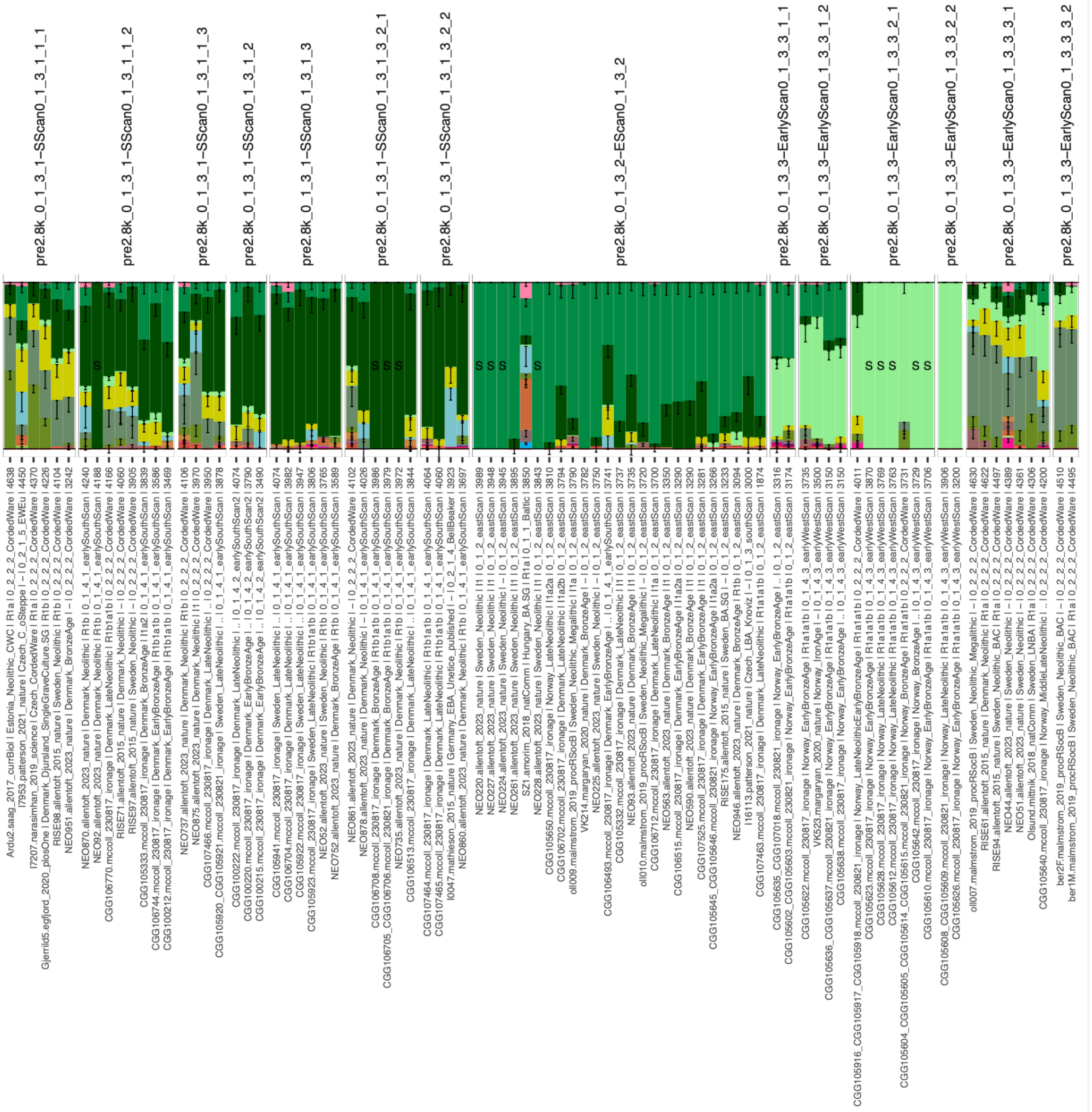

**Fig. S5.15. Mixture modelling results for Set 5 (upper panel, includes Bronze Age Scandinavian sources).** Samples are faceted on pre-2800 BP clusters (top of panel), and information on the original clustering is presented in the sample label below the panel. Samples that were originally in the Corded Ware cluster are modelled with high proportions of Corded Ware, Bell Beaker and Eastern North Sea Ancestry, and only small proportions of the local (Western Scandinavian, Southern Scandinavian) ancestries. This is particularly apparent in the pre2.8k_0_1_3_3-EarlyScan0_1_3_3_3_x clusters.

##### S5.2.3. Steppe ancestry within Farmer clusters

Many of the Bronze Age and later Southern European individuals only carry small amounts of Steppe ancestry, and hence cluster with Neolithic Farmers (Fig. S5.16, Fig. S5.30). By applying IBD mixture modelling with the representatives of the Yamnaya, Corded Ware and Bell Beaker clusters, we find the Steppe ancestry of the majority of these more southern individuals to be modelled as Bell Beaker-related (Fig. 3). By the Late Bronze Age onwards, irrespective of clusters, the Steppe ancestry in almost all Europeans can be well-modelled by Corded Ware (East) or the Bell Beaker sources (Fig. 3, Fig S6.17).

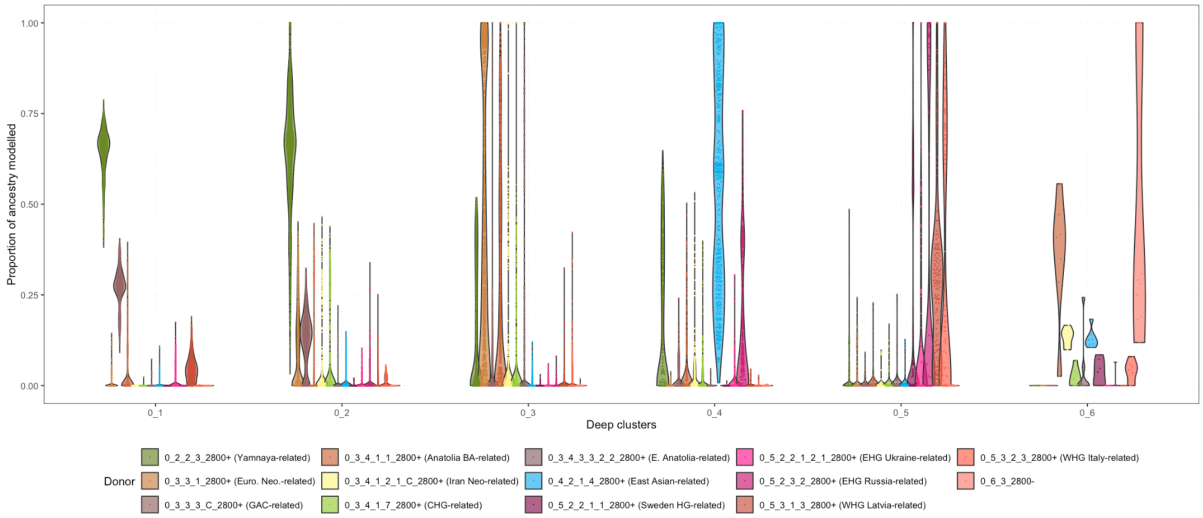

**Fig. S5.16. Violin plots showing the proportion of deep ancestries (Set Aux 1, Fig. S5.30) in the deepest levels of clustering for the entire dataset.** The main four subclusters of this study fall are the 0_1 cluster (Corded Ware (North)) and subclusters of the 0_2 cluster (Corded Ware (East), Bell Beaker, and Yamnaya), both of which are primarily modelled as the ‘dark green’ Yamnaya source (0_2_2_3_2800+). However, many individuals from the 0_3 cluster also carry some Steppe ancestry, despite being primarily modelled as the ‘brown’ Neolithic Farmer ancestry. Cluster 0_4 is modelled primarily with the ‘blue’ East Asian

Source, and cluster 0_5 by the 'orange' Western Hunter-Gatherer and 'purple' Eastern Hunter-Gatherer source.

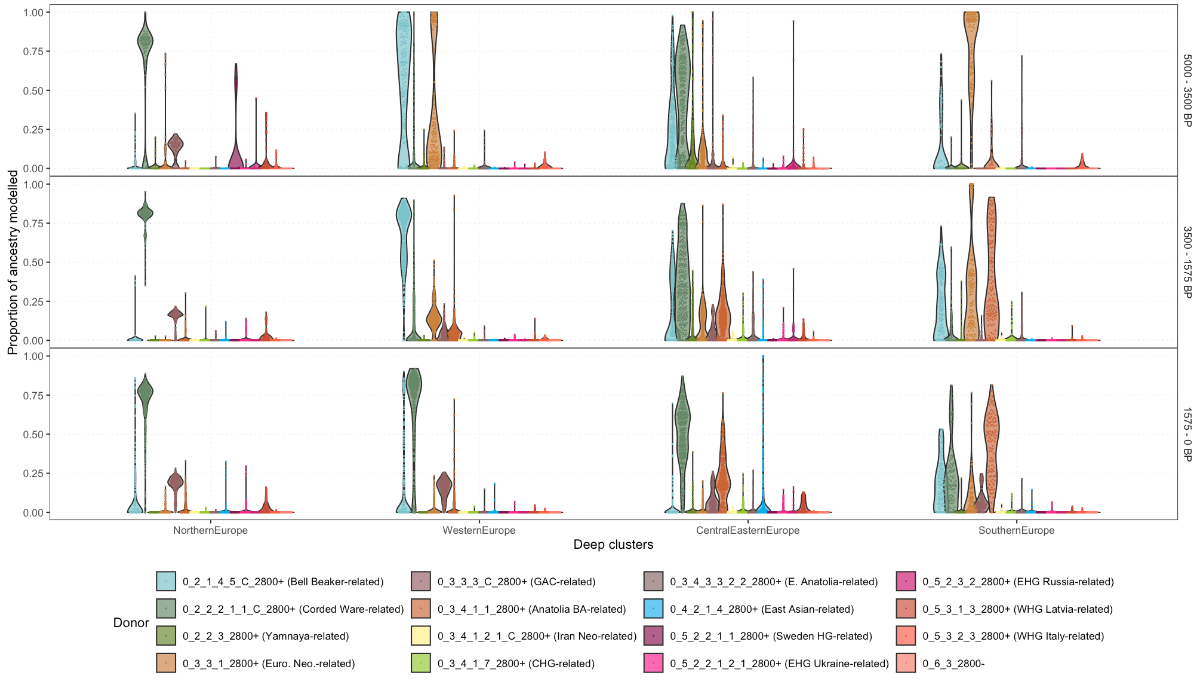

**Fig. S5.17 Violin plots showing mixture modelling results for Europe (Set 5), split into time bins.** The sources here include the three Steppe clusters: Bell Beaker (light blue), Corded Ware East (grey-green) and Yamnaya (dark green). In the earliest period (5000–3500 BP), a higher proportion of the Yamnaya source is present, compared to the later periods where for most individuals, Steppe ancestry is modelled entirely by the Bell Beaker and Corded Ware sources.

### S5.3. IBD mixture modelling

#### S5.3.1. Main mixture modelling sets

For IBD mixture modelling, we calculated total shared IBD lengths as in S5.2, but with a shorter minimal cut-off length of 1 cM per segment (based on the results from S5.3.9) and no lower limit on the total shared length . We undertook mixture modelling[135] on the total shared IBD lengths as described in Allentoft et al.[28], starting with the well-established diverse ancestries across Eurasia and adding more proximal sources (Table S5.3). These results were plotted as 'ADMIXTURE' style bar charts faceted by regions and country (Fig. S5.18), by IBD cluster (Fig. S5.19) and on the Western Eurasian PCA (Fig. S5.20).

When using a set of distal sources relevant to Europe during the Holocene (Western Hunter-Gatherers, Eastern Hunter-Gatherers, Caucasus Hunter-Gatherers, early Anatolian Farmers) and a series of out-groups (Table S5.3: Set 1), we find admixture proportions for Bronze Age Europeans consistent with the expectations; for Bronze Age Europe, individuals are modelled in primarily by the source populations for Yamnaya (Caucasus Hunter-Gatherer and Eastern Hunter-Gatherer) and Anatolian Farmers. By including the more proximal source, Yamnaya, the ancestry previously modelled as Caucasus Hunter-Gatherer and Eastern Hunter-Gatherer is now modelled by Yamnaya, despite all sources still being present (Fig. S5.20, Set 2, 3). We see similar patterns when including proximal admixed European Farmers to a more basal set with the distal Anatolian Farmers and Western Hunter-Gatherer source (Fig. S5.20, Set 3, 4), allowing us to progressively add more source clusters. When two source clusters used are too similar, large error bars appear and we reject the model. For each set, increasingly proximal sources are added. The full list of source individuals can be found in Table S5.3 and Fig. S5.22. and are described below.

Set 1: base_LvNoHg

The initial set included representatives of the four major ancestries present in Bronze Age Europe: Early Anatolian Farmers, Caucasus Hunter-Gatherer, Western Hunter-Gatherer from Southern Europe and Ukrainian Eastern Hunter-Gatherer. In addition, we included more distant source clusters, who are known to impact Western Eurasia from the Iron Age onwards: Norwegian Mesolithic, Latvia/Lithuania Mesolithic, Russian EHG, East Asian, Iran Neolithic and South Africans.

Set 2: base_LvNoHg_Yam

In addition to the clusters from Set 1, we included Yamanya as a source, who were previously modelled as Caucasus Hunter-Gatherer and Eastern Hunter-Gatherer.

Set 3: base_LvNoHg_Yam_Frm

Set 3 is identical to Set 2, but includes three clusters with Farming-related ancestry, representing known admixture events in Europe. The first cluster contains early European Farmers, here modelled as a mixture of early Anatolian Farmers and Western Hunter-Gatherer ancestry. The second cluster contains Globular Amphora Culture (GAC) individuals, who carry some Eastern Hunter-Gatherer ancestry similar to that of Yamnaya. A third cluster contains Bronze Age Anatolians, distinct from the early Anatolians in their additional Caucasus Hunter-Gatherer ancestry. Almost all Bronze Age and later Europeans carry GAC ancestry, whereas European Farmer ancestry is widespread in Southern and Western Europe, but not in the North.

Set 4_base_LvNoHg_Yam_Frm_CwBb5

Set 4 is identical to Set 3, but contains two additional clusters representing the major two archaeological cultures of the Bronze Age: Corded Ware ancestry and Bell Beaker ancestry.

Set 5: base_LvNoHg_Yam_Frm_CwBb5_scanENS

Set 5 is identical to Set 4, but contains 5 clusters related to various Western and Northern European Bronze Age clusters. These are a Bronze Age Southern Scandinavian cluster (0_1_4_1_earlySouthScan), a Bronze Age Eastern Scandinavian cluster (0_1_2_eastScan), a Bronze Age Western Scandinavia cluster (0_1_4_3_earlyWestScan), a Bronze Age Baltic cluster (0_1_1_Baltic) and a Bronze Age East North Sea cluster (0_2_1_2_wEuMl).

Set 6: base_LvNoHg_Yam_Frm_CwBb5_scanENS_IA

Set 6 is similar to Set 5, but now contains Iron Age Southern Scandinavians from Jutland (0_1_2_3_C_2800-), Iron Age Eastern Scandinavians from Danish Isles (0_1_2_2_3_C_2800-), Iron Age Eastern Scandinavians from Southern Sweden (0_1_2_1_2_C_2800-), Iron Age Western Scandinavians from Norway (0_1_2_4_5_C_2800-).

Set 7: base_LvNoHg_Yam_Frm_CwBb5_scanENS_IA_Gm

Set 7 is similar to Set 6, but includes a second more southern Iron Age Southern Scandinavian cluster from Northern Germany (0_1_3_2800-).

Mixture modelling results for all ancient individuals are plotted faceted by country (Fig. S5.18) and cluster (Fig. S5.19).

Due to the large size of the plots, a series of subsets from Fig. S5.18 and Fig. S5.25 are provided:

1. For reference, a plot including only the source clusters is shown in Fig. S5.22.
2. To visualise the Bronze Age diversity, individuals older than 2800 BP are shown in Fig. S5.21, faceted by cluster.
3. A subset of Fig. S5.25 showing only Danish, Swedish and Norwegian individuals is shown in Fig. S5.29.

We also note that the sources chosen are relevant to the specific research questions addressed in this study. For completeness, the plots include the results for all ancient individuals from the mixture modelling runs, but interpreting these results for other questions should be done with caution.

---

FIGURE INCLUDED SEPARATELY

---

**Fig. S5.18. Mixture modelling results for all ancient individuals. Results are faceted by region and country.** Within each country grouping, ancient individuals are ordered oldest to youngest, left to right. The 7 rows correspond to the 7 mixture modelling sets. For each set, sources are indicated by an 'S'.

---

FIGURE INCLUDED SEPARATELY

---

**Fig. S5.19. Mixture modelling results for all ancient individuals. Results are faceted by cluster**. Within each cluster, ancient individuals are ordered oldest to youngest, left to right. The 7 rows correspond to the 7 mixture modelling sets. For each set, sources are indicated by an 'S'.

---

FIGURE INCLUDED SEPARATELY

---

**Fig. S5.20. Mixture modelling results displayed on the Western Eurasian PCA.** Source individuals are circled, and admixture proportions follow a cline from full colour (100%) to grey (0%). The 7 rows correspond to the 7 mixture modelling sets.

---

FIGURE INCLUDED SEPARATELY

---

**Fig. S5.21. Subset of the cluster-faceted Fig. S5.25, plotting only samples older than 2800 BP.**

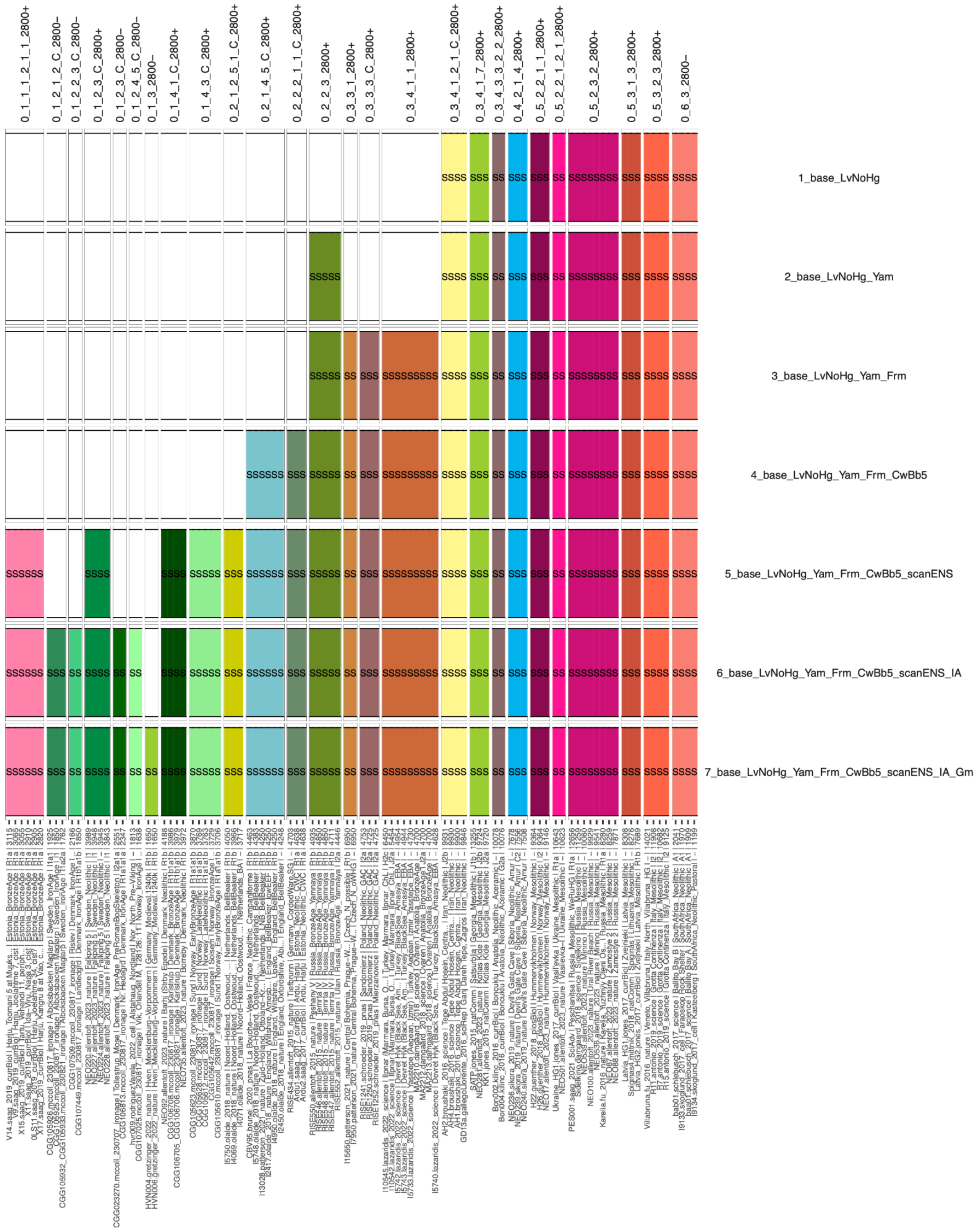

**Fig. S5.22. Mixture modelling results subset showing just source individuals used for Sets 1-8.**

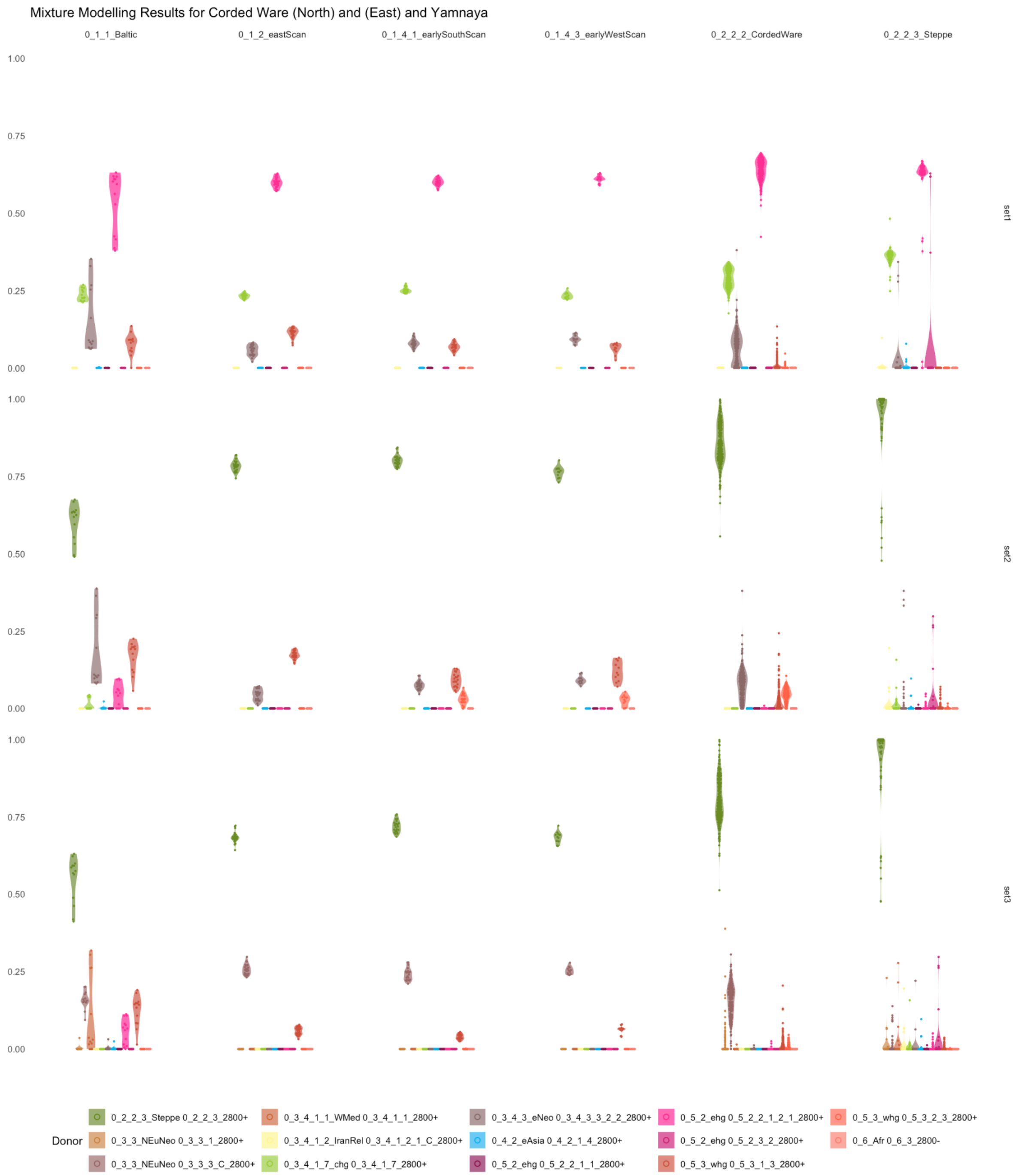

**Fig. S5.23. Violin plots showing subsets of results from Fig. S5.21, relevant for Corded Ware (East) and (North) clusters, for samples over 2800 BP.** Sets 1 and 2 show the variation in ancestry of Western Hunter-Gatherers (Southern Europe - 0_5_3_2_3_2800+) compared to Northern Western Hunter-Gatherers (Latvia, Lithuania - 0_5_3_1_3_2800+) for Southern, East and Western Scandinavians. Set 2 and 3 show the high Eastern Hunter-

Gatherer from Ukraine present in the Baltic Cluster. Set 2 shows the Yamnaya to Farmer ancestry cline present in the Corded Ware (East) cluster (0_2_2_3). Set 3 shows this ancestry modelled as GAC (0_3_3_3_C_2800+). Information on each set can be found in S5.3.1.

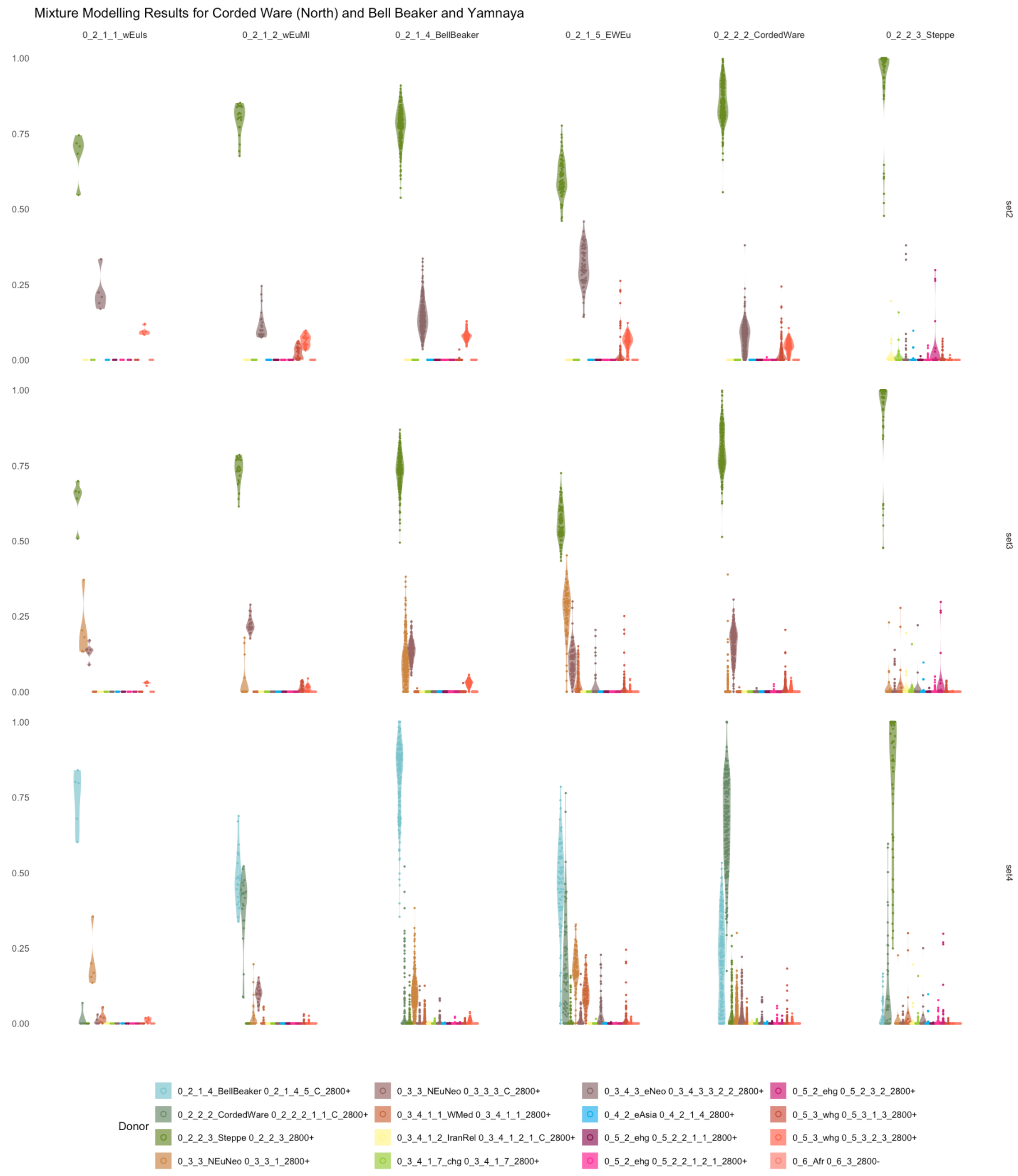

**Fig. S5.24. Violin plots showing subsets of Fig. S5.21, relevant for Corded Ware (East) and Bell Beaker clusters, for samples over 2800 BP.** Set 3 shows the European Farmer (0_3_3_1_2800+) present in most Bell Beakers, in addition to the GAC (0_3_3_3_C_2800+) ancestry found in the Corded Ware (East) individuals. Set 4 shows the high proportion of Bell Beaker ancestry relative to Corded Ware ancestry in most individuals. The East North Sea (0_2_1_2_WEuMl) cluster is unique, in the high proportion of Northern Western Hunter-Gatherer (Latvia, Lithuania - 0_5_1_3_3_2800+) ancestry (Set 2), low European Farmer (0_4_4_1_1_1_2800+) ancestry (Set 3) and equal proportion of Bell Beaker and Corded Ware ancestry (Set 4), reflecting both its geographical position between the two cultures and its position in the Western Eurasian PCA space (Fig. S5.13).

---

FIGURE INCLUDED SEPARATELY

---

**Fig. S5.25. Mixture modelling results overlaid on a map of Northern Europe for Set 6, including one Southern Scandinavian IA source.**

---

FIGURE INCLUDED SEPARATELY

---

**Fig. S5.26. Mixture modelling results overlaid on a map of Northern Europe for Set 7, including two Southern Scandinavian IA sources.**

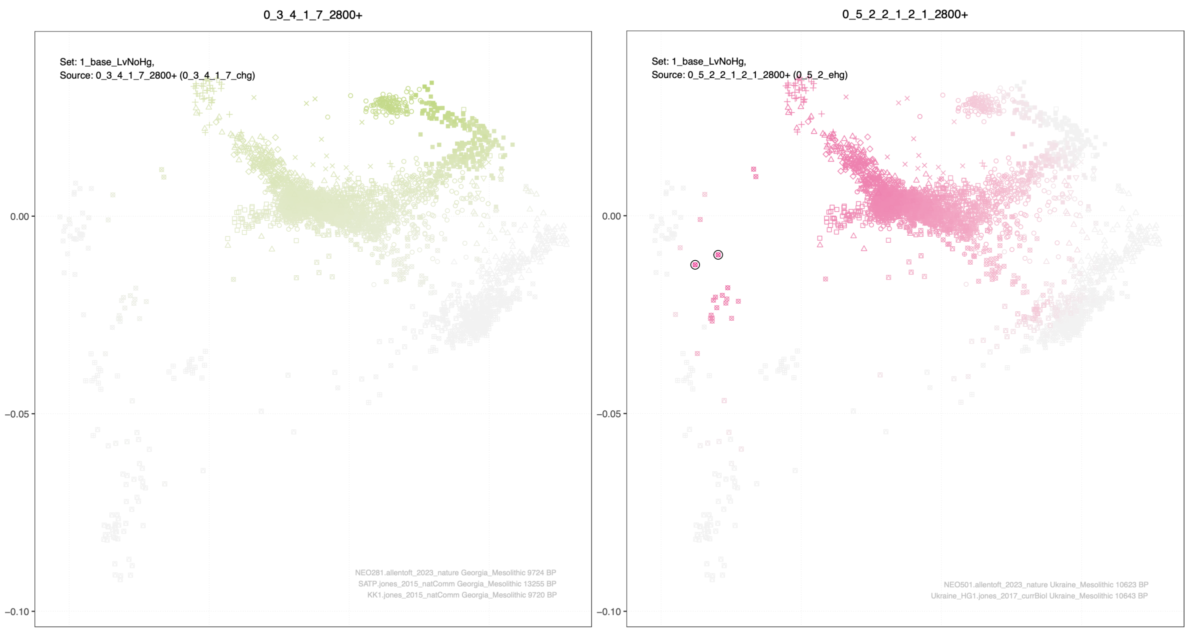

**Fig. S5.27. PCA results coloured by mixture modelling proportions when including more proximal European Steppe-related sources: Eastern Hunter-Gatherer ( pink), Caucasus Hunter-Gatherer (light green) source clusters (Set 1)**. Source individuals are circled. Subset of Fig. S5.20

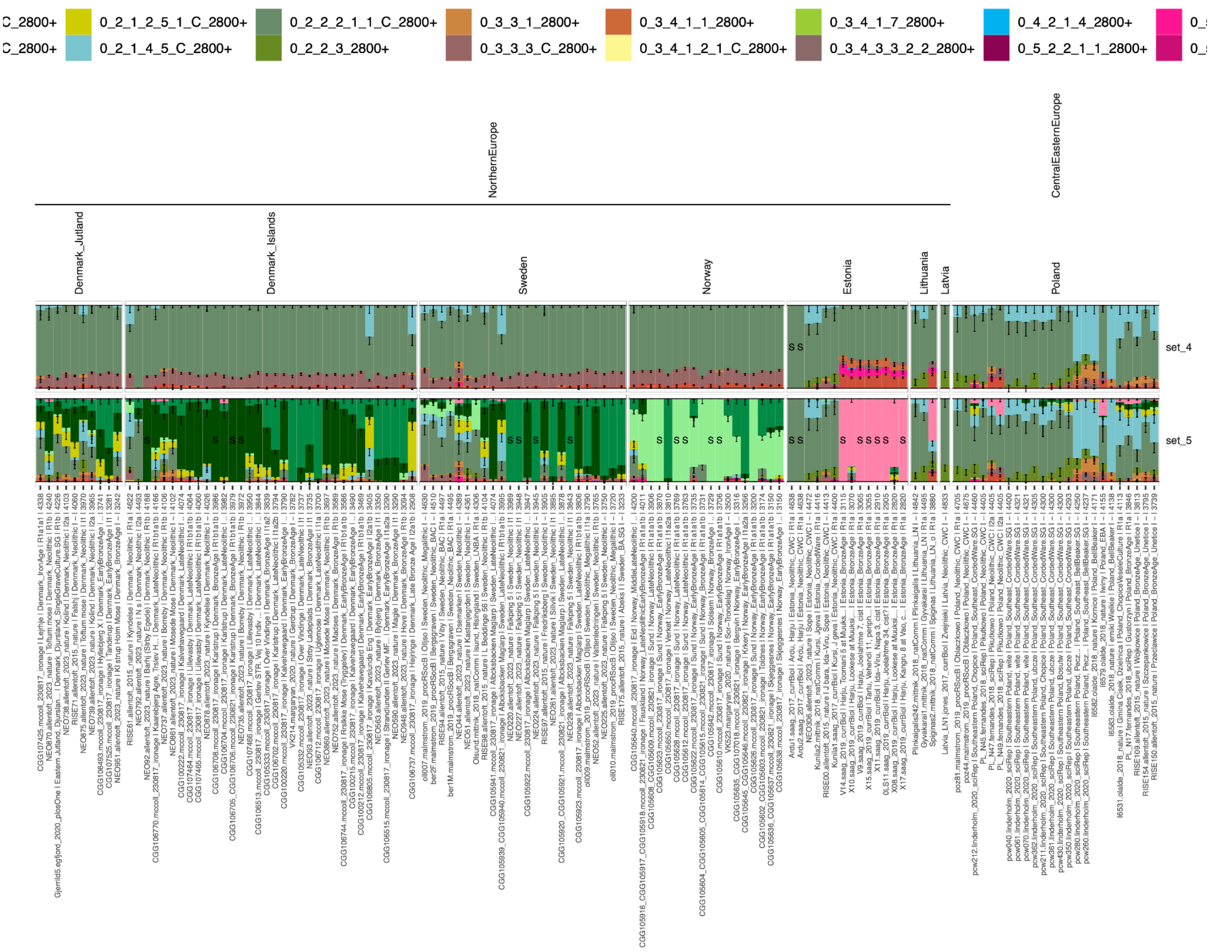

**Fig. S5.28. Subset of mixture modelling results from around the Baltic region, from 4800–2800 BP**. Both sides of the Baltic are modelled primarily as Corded Ware (East) ancestry (grey-green in Set 5), rather than Bell Beaker (blue). However, when including the Scandinavian sources (Eastern Scandinavian: light green, Western Scandinavian: bright green, Southern Scandinavian: dark green), the Estonian, Latvian, Lithuanian, and Polish individuals continue to be modelled by the more distal Corded Ware (East) source.

---

FIGURE INCLUDED SEPARATELY

---

**Fig. S5.29. A subset of Fig. S5.19, showing just Danish, Norwegian and Swedish samples.**

##### S5.3.2. Auxiliary mixture modelling sets

The ‘main’ mixture modelling sets were generated with more proximate source clusters added in each set. We also generated a set of auxiliary sets for specific research questions, as well as sets for the IBD mixture modelling tests (Supplementary Note S5.3.6). For these sets, the individuals used are detailed in Supplementary Table S5.4.

Auxiliary Set 1. Similar to main Set 3, except only a single Eastern Hunter-Gatherer and Western Hunter-Gatherer source cluster are included to demonstrate the Eastern–Western Hunter-Gatherer cline (Fig. 4). All main sets have additional sources from the Eastern–Western Hunter-Gatherer cline. Results can be seen in Fig. S5.30.

Auxiliary Set 2. Similar to main Set 6, but with Bronze Age early admixed Eastern and local (Western and Southern) Scandinavians as sources for later Bronze Age individuals. Results in Fig. S5.18. A comparison to the earlier, unadmixed Bronze Age individuals for Scandinavia is shown in Fig. S5.32.

Sets for testing the impact of temporal distance between source and target cluster (Supplementary Note S5.3.6):

2_base_DkHg, 3_base_ltAna, 4_base_euNeo

Sets for testing the impact of number of source individuals (Supplementary Note S5.3.6):

2.1sScan.1wScan.1eScan, 3.1sScan.5wScan.4eScan, 4.2sScan.5wScan.4eScan, 5.3sScan.5wScan.4eScan, 6.4sScan.1wScan.4eScan, 7.4sScan.2wScan.4eScan, 8.4sScan.3wScan.4eScan, 9.4sScan.4wScan.4eScan, 10.4sScan.5wScan.1eScan, 11.4sScan.5wScan.2eScan, 12.4sScan.5wScan.3eScan

---

FIGURE INCLUDED SEPARATELY

---

**Fig. S5.30. Mixture modelling results for Auxiliary Set 1**

---

FIGURE INCLUDED SEPARATELY

---

**Fig. S5.31. Mixture modelling results for Auxiliary Set 2**

---

FIGURE INCLUDED SEPARATELY

---

**Fig. S5.32. A subset of mixture modelling results for the Bronze Age Scandinavian sources (Set 5) and the admixed late Bronze Age sources (Auxiliary Set 2).** This subset includes only Danish, Norwegian and Swedish individuals between 5000 and 1200 BP.

##### S5.3.3. Admixture clines within Europe

We plotted the admixture proportions inferred by the IBD mixture modelling (Supplementary Note S5.2, Fig. 4 and Fig. S5.20) on top of the standard Western Eurasian PCA to explore the geographic apportionment of each genomic ancestry. By using the well-established distal sources for Europe, we first replicated previous established interactions between Mesolithic Hunter-Gatherer, Neolithic Farmer and Bronze Age Steppe populations[17,18], representing the major migrations into Europe. We see the relative proportions of Steppe- and Farmer-related ancestry along the Yamnaya-Neolithic Farmer cline in Fig. 4A and Fig. 4B, of Farming- and Western Hunter-Gatherer-related ancestry along the respective clines in Fig. 4B and Fig. 4C, and Western Hunter-Gatherer and Eastern Hunter-Gatherer along the respective clines in Fig. 4C and Fig. 4D.

This representation of our results with more proximal sources revealed a series of novel genetic clines and provides additional resolution to previously found clines within the densely overlapping Bronze Age PCA space. We modelled all individuals in the dataset using three representatives of Steppe ancestry (the Yamnaya-, Corded Ware (East)- and Bell Beaker-related clusters) as sources to explore the relationship between the early individuals from each cluster and later populations in time (Set 4, relevant subset in Fig. 4, full set in Fig. S5.20). To explore interactions with the Farming populations present in Europe during this time, we also included representatives of three Farming-related clusters; the Globular Amphora Culture (GAC) of North East Europe (0_3_3_3_C_2800+), European Farmers (0_3_3_1_2800+), and Levant/Bronze Age Anatolians (0_3_4_1_1_2800+). Despite all three being modelled primarily with Neolithic Farming ancestry, they are modelled with small proportions of North East European Hunter-Gatherer (Latvian/Lithuanian), Western Hunter-Gatherer (Italy) and Caucasus Hunter-Gatherer ancestry respectively (Fig. S5.21).

We find that the estimated Yamnaya admixture proportions previously shown to decrease along the cline connecting Yamnaya individuals and the dense clustering of BA diversity in Auxiliary Set 1 (Fig. 4A) to now be modelled by Corded Ware source (Fig. 4F). The decrease corresponds with increasing Farming-related ancestry, which is here modelled as GAC (Fig. 4H, Fig. S5.23.). We interpret this cline to correspond to the admixture with GAC previously documented to occur prior to the arrival of Steppe ancestry in Europe[28]. This cline corresponds to one of the four previously mentioned clusters, the 'Corded Ware (East)' cluster (Fig. 2).

Overlapping with the admixed European tip of this Corded Ware (East) cline, we find a series of additional clines, representing additional admixture between early European Steppe people already carrying GAC ancestry and additional Farming-related groups in Europe. The first cline within this diversity extends from this point to the European Farmer cline. The Steppe ancestry in this cline is modelled by the Bell Beaker-related source (Fig. 4G), and the additional Farming ancestry by the European Farmers (Fig. 4I). We interpret this cline to represent additional admixture with Farming sources within Europe who themselves carry some Western Hunter-Gatherer ancestry (Fig. 4C). This cline corresponds to another of the four previously mentioned clusters, the Bell Beaker cluster (Fig. 2). Very few individuals within Europe are modelled as Yamnaya when the Corded Ware and Bell Beaker source clusters are included (Fig. 4E).

From within the Corded Ware (East) and Bell Beaker diversity, we find an additional two clines extending to the Levant/Bronze Age Anatolians cluster (Fig. 4J). The Steppe ancestry of the first cline is modelled as Bell Beaker-related and corresponds to the 0_2_1_5_x subcluster (Fig. S5.12) within the main Bell Beaker cluster and contains many Hallstatt and La Tene individuals (Table S5.1), which we interpret as admixture within the range of these cultures, from France to the Black Sea. However, no suitable Bronze Age source cluster could be identified, suggesting a higher degree of continuity and complexity within the Bell Beaker-related populations of this region, consistent with previous studies[186]. The Steppe ancestry of the second cline is modelled as Corded Ware (East) ancestry (Fig. 4F) and corresponds to the Corded Ware (North) cline (Fig. 2), which we interpret as admixture between Northeast and Southeast Europe. For both clines, the additional Farmer ancestry is modelled as Levant/Bronze Age Anatolian (Fig. 4J).

Individuals with varying proportions of Corded Ware, Bell Beaker and Eastern Mediterranean Bronze Age ancestry are present as a cloud between the two clines. Entering this cloud at various angles we find three Corded Ware (North) subclusters (Eastern Scandinavian: Extended Data Fig.-8B, Southern Scandinavian: Extended Data Fig.-8C, Baltic: Extended Data Fig.-8A) and one Bell Beaker subcluster (Eastern North Sea: Extended Data Fig.-8D), which we interpret as admixture into different regions of Europe with varying proportions of the Farming-related sources (Fig. 4H, Fig. 4I, Fig. 4J).

We find an additional cline at the other end of the Corded Ware (North) cluster, extending towards the Eastern Hunter-Gatherers relative to the cline of Corded Ware (East) and GAC ancestry. At the end of this cline and corresponding to their position in the PCA, we find Estonian Bronze Age individuals of the Baltic subcluster modelled with additional Ukrainian Eastern Hunter-Gatherer ancestry, in addition to the Lithuanian/Latvian Hunter-Gatherer ancestry modelled in the other Corded Ware (North) clusters (Fig. S5.23). Notably, the presence of this ancestry makes the Baltic subcluster distinct from the other Corded Ware

(North) subclusters. Even the Bronze Age individuals with the highest Farmer ancestry from this cluster have higher Eastern Hunter-Gatherer ancestry than any Bronze Age individual from the other Corded Ware (Northern) subclusters, despite its southern location in Croatia (Fig. S5.21).

##### S5.3.4. Variation in Hunter-Gatherer ancestry in Steppe-related populations

We detect differences in structure within the Corded Ware (North) subclusters (Eastern Scandinavian, Western Scandinavian, Southern Scandinavian, Baltic), the Corded Ware (East) cluster and the Steppe cluster (including Yamnaya) by first modelling with the deeper ancestry sources (Fig. S5.23). When using the Set 2 mixture modelling sources, we find all clusters have relatively high Eastern Hunter-Gatherer (Fig. S5.27, Ukraine Eastern Hunter-Gatherer: 0_5_2_2_1_2_1_2800+, Russia Eastern Hunter-Gatherer: 0_5_2_3_2_2800+) and Caucuses Hunter-Gatherer ancestry, the expected sources for Steppe ancestry[43]. However, the Eastern Scandinavian cluster here is modelled with an additional small proportion of Lithuanian/Latvian Hunter-Gatherer ancestry (0_5_3_1_3_2800+).

By including Yamnaya (0_2_2_3_2800+) as a source (Fig. S5.23: Set 3), the Caucasus Hunter-Gatherer ancestry and much of the Eastern Hunter-Gatherer ancestry is now modelled as Yamnaya, revealing the additional Hunter-Gatherer ancestry present in the various groups is present in more detail. Here, we reveal small proportions of the Lithuanian/Latvian Hunter-Gatherer ancestry (0_5_3_1_3_2800+) in all Corded Ware (North) clusters, although the proportion in the Eastern Scandinavians remains highest. An additional distinction is that the Southern and Western Scandinavian clusters are modelled with some Italian Western Hunter-Gatherer ancestry (0_5_3_2_3_2800+), unlike the Eastern Scandinavians. Finally, we see the

Baltic cluster is modelled with additional Ukrainian Eastern Hunter-Gatherer ancestry, unlike the Scandinavian clusters.

By plotting the admixture proportions on the standard Western Eurasian PCA, a broader pattern is visible (Fig. S5.33). In the region of the Corded Ware (North) cluster (Fig. 2), the Latvian/Lithuanian Hunter-Gatherer ancestry is widespread (Fig. S5.33B). In the region of the Bell Beaker cluster (Fig. 2), Western Hunter-Gatherer ancestry is widespread (Fig. S5.33C). Ukrainian Eastern Hunter-Gatherer ancestry is present along the Baltic cluster (Fig. S5.33A).

However, individuals within the Bell Beaker and Corded Ware (North) clusters are also modelled with European Farmer and GAC-related Farming ancestry (Supplementary Note 5.3.3). European Farmers are modelled with some Western Hunter-Gatherer and Latvian Lithuanian Hunter-Gatherer ancestry, and the GAC individuals are modelled with Latvian Lithuanian Hunter-Gatherer ancestry. By including these Farming clusters as sources (Set 3), the excess Latvian/Lithuanian (Fig. S5.33E) and Western Hunter-Gatherer (Fig. S5.33F) can be visualised.

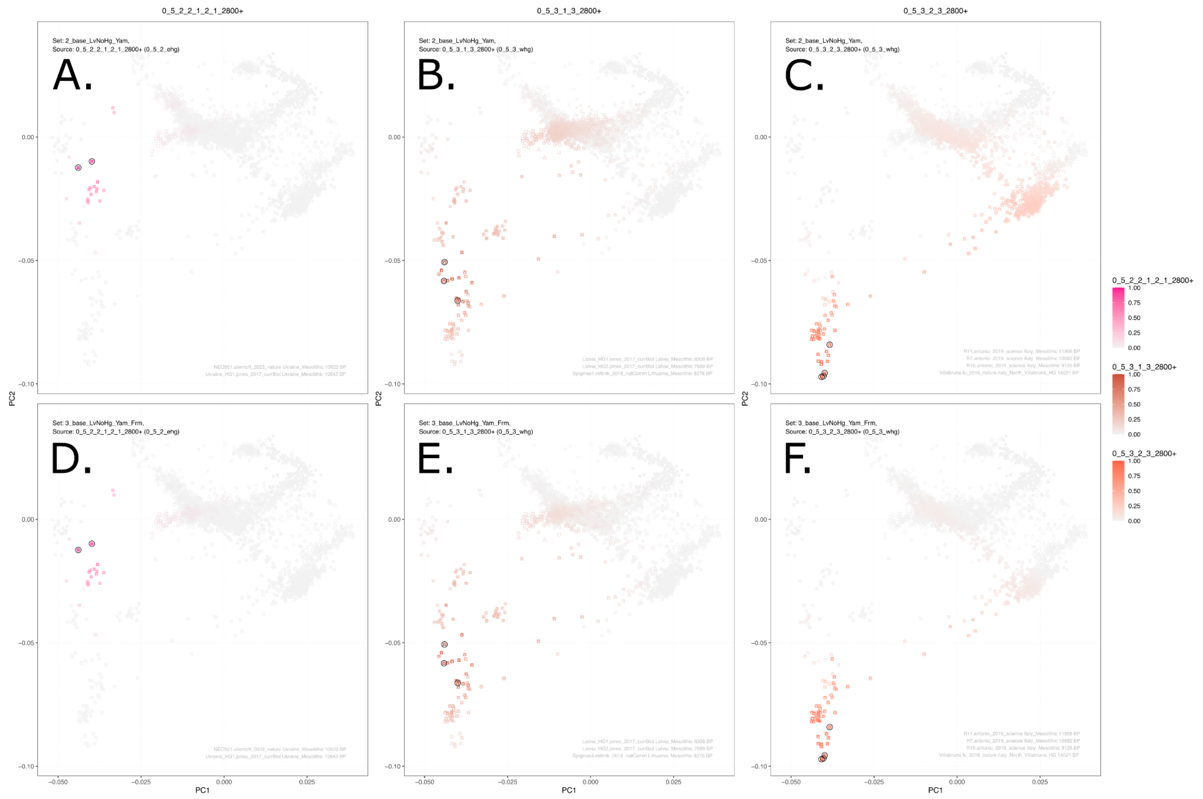

**Fig. S5.33. A subset of the mixture modelling results for Set 3 (A, B, C) and Set 4 (D, E, F).** The first column represents the Eastern Hunter-Gatherer source, the middle column represents the Latvian/Lithuanian Hunter-Gatherer source, and the right column indicates the Western Hunter-Gatherer source. Set 4 is similar to Set 3, but includes additional Farming related sources. The variation seen between row 3 and 4 results from the Hunter-Gatherer ancestry present in the additional Farming sources.

To validate the results using more traditional methods, we first confirmed a higher proportion of Hunter Gatherer ancestry in the Eastern Scandinavians using D-statistics in admixtools2 [197] on the 1240K sites as filtered for imputation. For individuals from the three main Scandinavian clusters, as well as some early Corded Ware (East) individuals, we calculated D(Outgroup, Scandinavian_test, Western Hunter-Gatherer, Eastern Hunter-Gatherer). We see a subtle trend with the highest proportion of WHG ancestry in Eastern Scandinavians, then

Western Scandinavians, Southern Scandinavians and finally the early Corded Ware individuals (**Fig. S5.34**). We coloured each point by the proportion of Latvian/Lithuanian HG ancestry from the IBD mixture modelling results (Set 3).

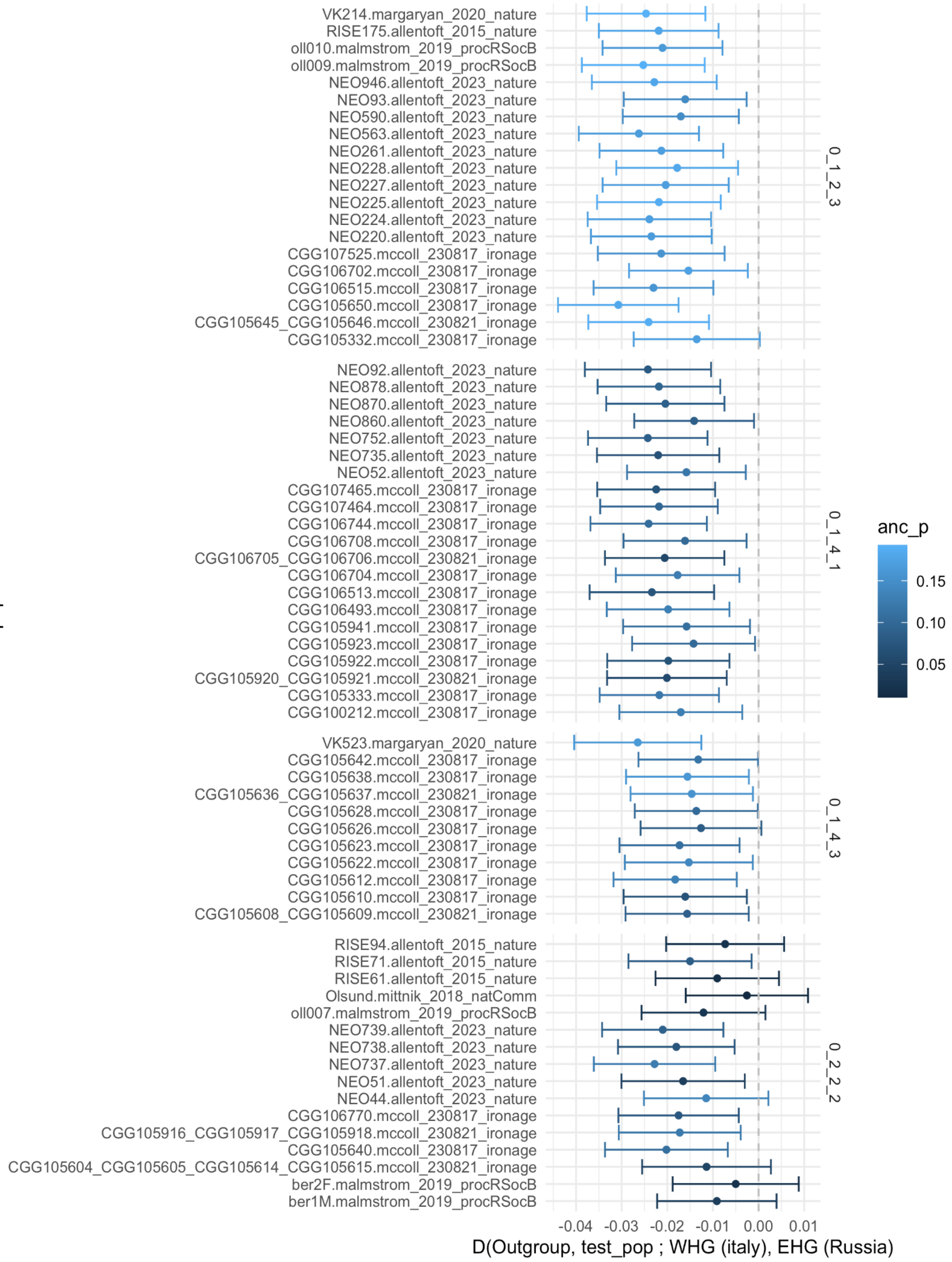

**Fig. 5.34. Statistic showing the variation in Western Hunter-Gather and Eastern Hunter-Gatherer in individuals from the Eastern Scandinavian BA (0_1_2_3), Southern Scandinavian BA (0_1_4_1), Western Scandinavian (0_1_4_3) and the Corded Ware (East) clusters.** The colour represents the proportion of ancestry modelled as 'Lithuanian/Latvian HG' ancestry in Set 2 of the IBD Mixture Modelling.

To identify the specific hunter-gatherer source, we ran qpadm using admixtools2 [197] on the same set of filtered 690,211 sites used for IBDseq based analyses. As targets we used individuals from a Baltic Bronze Age cluster (0_1_1_1_2_1_1_2800+) who are modelled with some Eastern Hunter Gatherer), an Eastern Scandinavians Bronze Age cluster (0_1_2_3_C_2800+) who are modelled with some Latvian Lithuanian HG ancestry), West and Southern Scandinavian Bronze Age clusters (0_1_4_3_C_2800+ and 0_1_4_1_C_2800+) who are modelled with extra Western HG), and an east Corded Ware cluster (0_2_2_2_1_1_C_2800+), who are not modelled with additional HG ancestry.

For all qpadm runs, we included individuals from the following clusters as outgroups in in the right group: an Iran Neolithic clusters (0_3_4_1_2_1_C_2800+), an East Asian cluster (0_4_2_1_4_2800+), an African cluster (0_6_3_2800-), the Caucasus HG cluster (0_3_4_1_7_2800+), and an early Anatolian Farmer cluster (0_3_4_3_3_2_2_2800+).

In the left group, we fixed Yamnaya (0_2_2_3_2800+) and Globular Amphora (0_3_3_3_C_2800+), based on modelling suggesting Bronze Age Northern Europeans carry GAC, rather than European Farmer ancestry (Supplementary Figure S5.23: set 3).

We then selected a series of HG individuals from across Europe to 'rotate' through the left and right groups, including known Hunter Gatherer genetic structure from the region[44–46,113]: These included individuals form a Pitted Ware cluster (0_5_1_1_1_1_2800+), an early EHG related Sweden Mesolithic cluster (0_5_1_2_2_2800+), a Norway Mesolithic EHG related cluster (0_5_2_2_1_1_2800+), a Lithuanian/Latvian Mesolithic cluster (0_5_3_1_3_2800+), a Central European WHG related Mesolithic cluster, including individuals from France, Britain and Switzerland (0_5_3_2_1_1_2800+), an early Italian WHG Mesolithic cluster (0_5_3_2_3_2800+), a Danish WHG-related Mesolithic related cluster (0_5_3_3_1_2800+), and a Late Swedish individual, who clustered with Danish Mesolithic individuals but is modelled similarly to the Latvian and Lithuanian HGs (0_5_3_3_3_2800+).

For an initial 2 population model, the above hunter-gatherers were included only as right groups. For the 3 population model, one of the hunter-gatherers were included in the left group, along with the Yamnaya and Globular Amphora groups, and the rest were included as in the right group. For the 4 population model, two hunter gatherer groups were included in the left group, along with the Yamnaya and Globular Amphora groups, and the rest were included as in the right group.

Information on the specific individuals used in each group and the results from the qpAdm analyses can be found in Table S5.5.

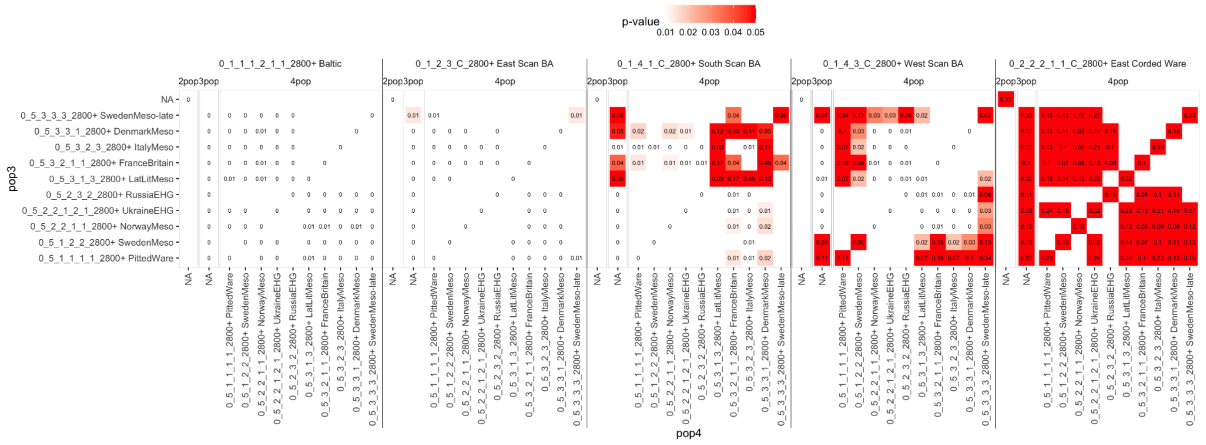

**Fig. 5.35. qpAdm P-values for two population, three population and four population models.** Yamnaya and Globular Amphora sources are fixed for all combinations, and Hunter-Gatherer sources are rotated through in the three and four population models. Values for infeasible results are not shown.

In the initial 2 population model, with only Yamnaya and GAC in the left group, and all outgroups and HGs in the right, we found only the East Corded Ware cluster (0_2_2_2_1_1_C_2800+) can be feasibly modelled with a significant p-value (0.12). The results can be seen in Fig. S5.35. The raw output can be found in Supplementary Table S5.5.

Under a 3 population model, we find good fits for the Southern Scan (0_1_4_1_C_2800+), with an additional HG source from Sweden, Denmark, France/Britain/Switzerland or Latvia/Lithuania. For Western Scan (0_1_4_3_C_2800+), we find fits with Swedish Meso-late and individuals from the EHG-related early Swedish Mesolithic or Pitted Ware culture. For both targets, the distinction between hunter-gatherer ancestries reflects the region in which they are found.

The Eastern Scandinavians cannot be modelled under the three population model. Here, the most significant result (p-value of 0.014) is found when using the late Swedish Mesolithic source. Under a 4 pop model, the Eastern Scandinavian individuals still do not fit. Combined, these results suggest that the specific source for the Hunter-Gatherers is not part of the established Mesolithic populations structure within Scandinavia, and does not exist in our dataset, but it point to a region along the coastline of Scandinavia or the Baltic region, consistent with the IBD mixture modelling results.

##### S5.3.5 Comparisons with ancIBD

A previous study applying the IBD clustering and mixture modelling methods[28] similarly used IBDseq to detect IBD segments in whole genome data. The recent release of ancIBD[36], a method tailored specifically for the ancient genomic data and the 1240K capture sites, is of potential relevance here. While this method has been shown to accurately call longer segments of IBD (> 8 cM), further investigation into the signal-to-noise ratio was required for the shorter segments (< 1cM), which are more informative of deeper relations and were included in mixture modelling by Allentoft et al[28].

We ran ancIBD (v0.6), as described in the online documentation (https://ancibd.readthedocs.io/en/latest/index.html). Allele frequencies were calculated from the samples themselves. We applied the —mask function using the provided mask file, as recommended for segments less than 8cM. We also included only segments that contained at least 220 SNPs per cM. To test whether more stringent filtering improved the signal to noise ratio, we created two additional subsets of the data retaining only segments with at least 330 and at least 440 SNPs per cM.

To visualise the difference between the IBDseq and ancIBD results at the various lengths fit with expectations, we separated the segments into 1-2cM, 2-4cM, 4-8cM, 8-12cM, and 12cM+ bins, and then summed the total length of the segments for each pair of individuals, plotting the results as a heat map. As the clustering results were broadly consistent with the Western Eurasian PCA and well established expectations from the human population history, we ordered the samples within the plot by the broader clusters.

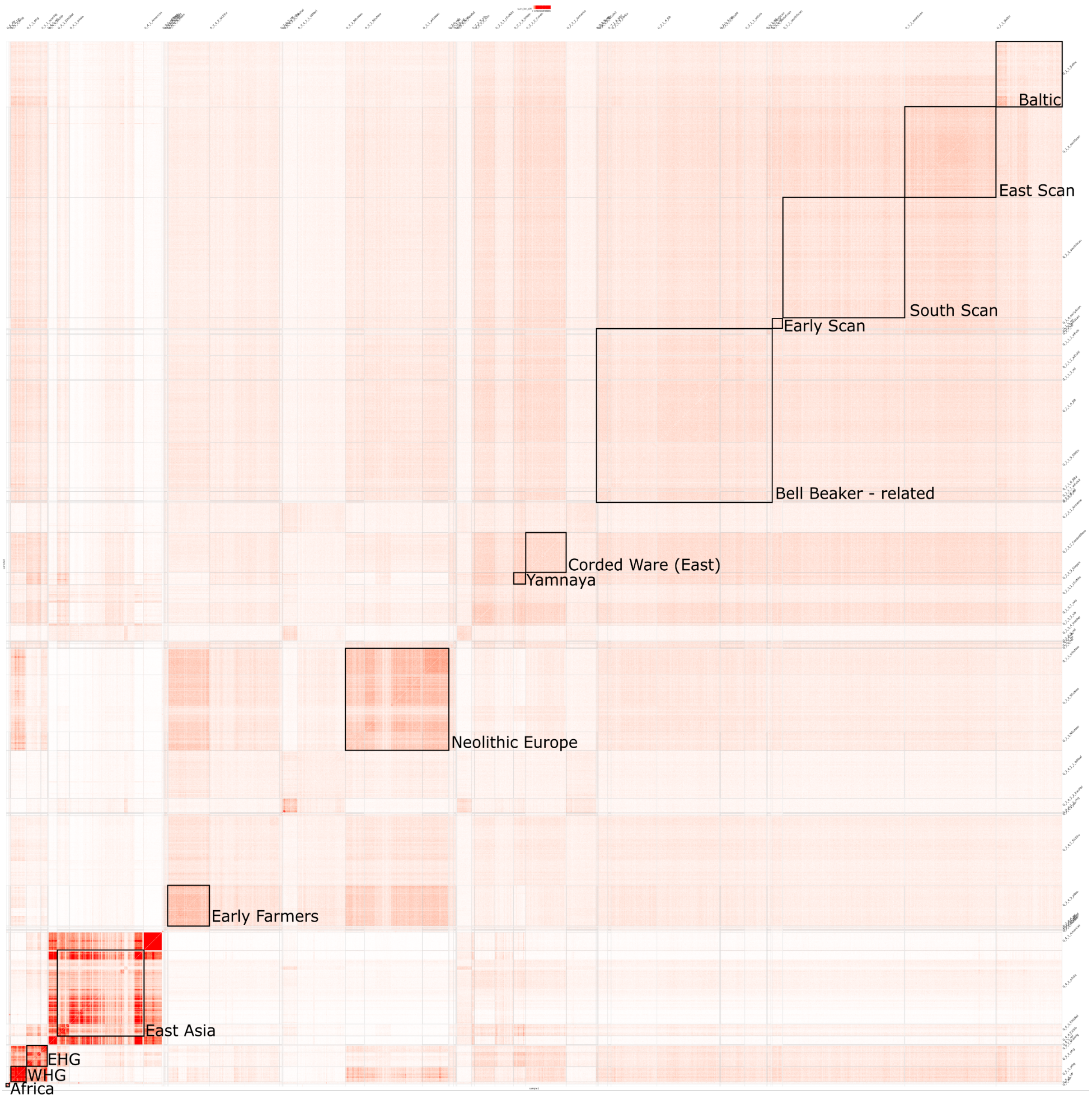

**Fig. S5.36. Annotated heatmap showing the sum of IBD segments between 1-2cM for all pairs of ancient individuals, called by IBDseq, ordered by cluster.** Clusters of interest have been highlighted.

For the subset of the data from IBDseq, in the bin with the shortest segments (1-2cM) we see the clusters consistent with expectations, as annotated in **Fig. S5.36**. For example, Hunter-Gatherer, Neolithic and Steppe related clusters have the highest sharing within their own broad clusters. In addition, we see evidence of known admixture events; the Early Farming cluster shares considerably less with WHG than the admixed European Farmer cluster, and Steppe-related clusters from within Europe show evidence of admixture with Neolithic Farmers, however the Yamnaya-cluster does not.

For IBDseq results, these patterns clearly are visible in the 1-2cM (**Fig. S5.37A)** and 2-4cM (**Fig. S5.38A)**, and to a lesser degree the 4-8cM bins **(Fig. S5.39A)**. In the 8-12cM (**Fig. S5.40A)** and 12+cM (**Fig. S5.41A)** only within-cluster sharing is apparent.

In contrast, these patterns are not visible in the ancIBD results for 1-2cM (**Fig. S5.37B,C,E)** or 2-4cM (**Fig. S5.38B,C,E)**. Here, more sharing that does not fit with expectations is present; East Asians and Bronze Age Europeans, and Africans with many out-of-Africa populations. In the 4-8 (**Fig. S5.39B,C,E)** and 8-12cM (**Fig. S5.40B,C,E)**, distinction between the clusters is more apparent, but more noise is still remains present.

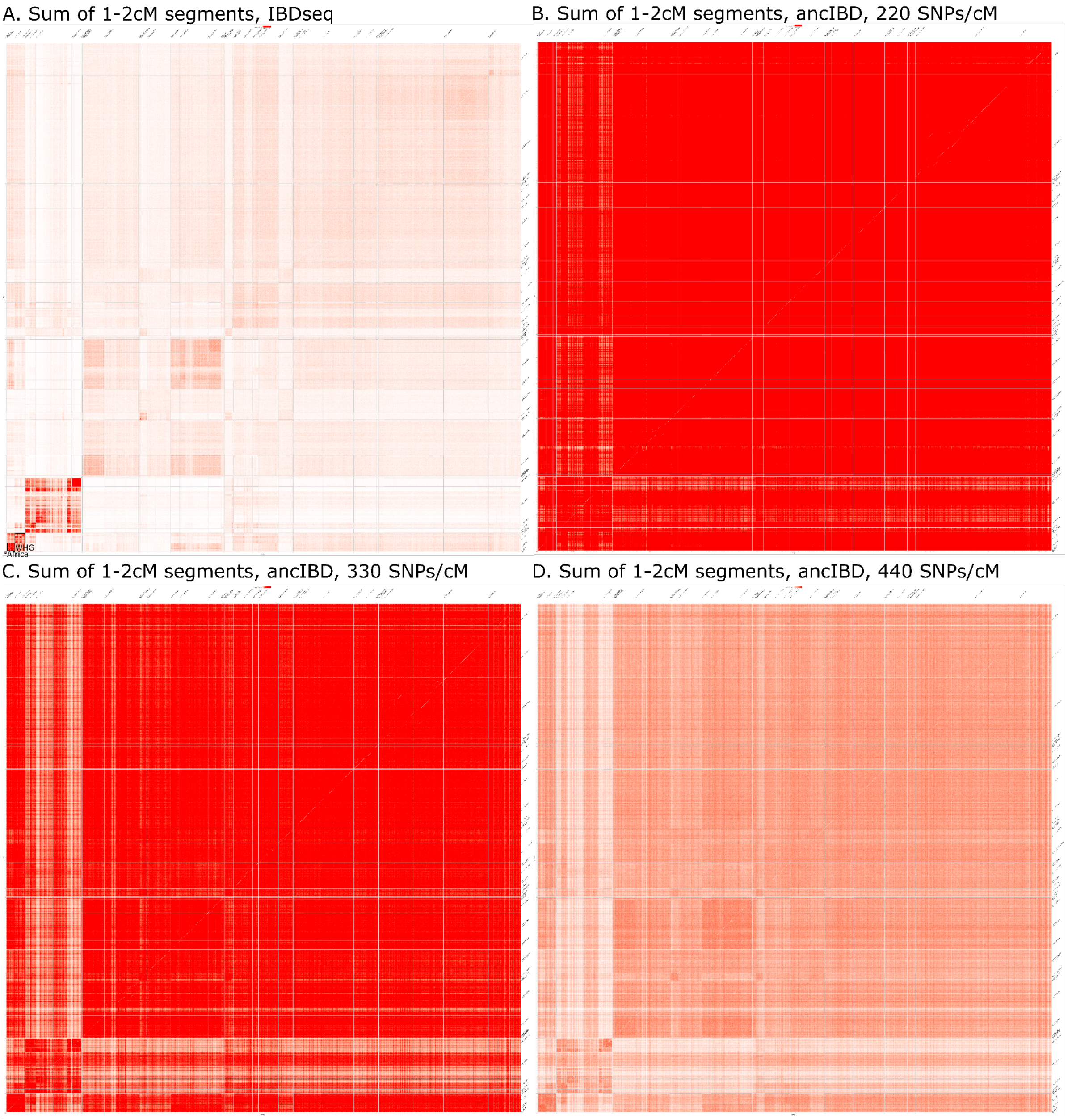

**Fig. S5.37. Heatmap showing the sum of IBD segments between 1-2cM for all pairs of ancient individuals, called by (A) IBDseq, (B) ancIBD at 220 SNP per cM, (C) ancIBD at 330 SNP per cM and (D) ancIBD at 440 SNP per cM .** Results from 0-100 cM are shown from white to red.

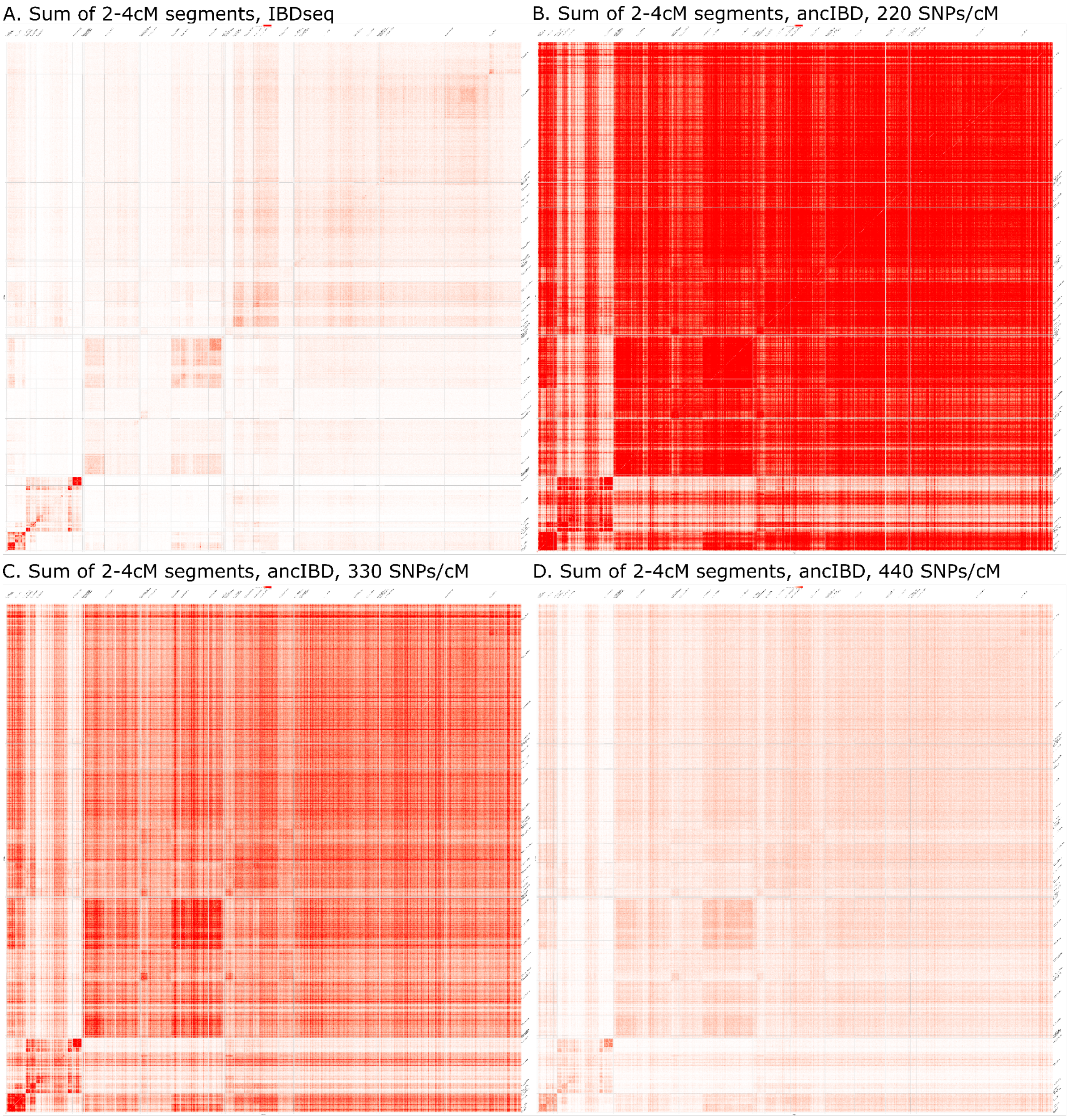

**Fig. S5.38. Heatmap showing the sum of IBD segments between 2-4cM for all pairs of ancient individuals, called by (A) IBDseq, (B) ancIBD at 220 SNP per cM, (C) ancIBD at 330 SNP per cM and (D) ancIBD at 440 SNP per cM .** Results from 0-100 cM are shown from white to red.

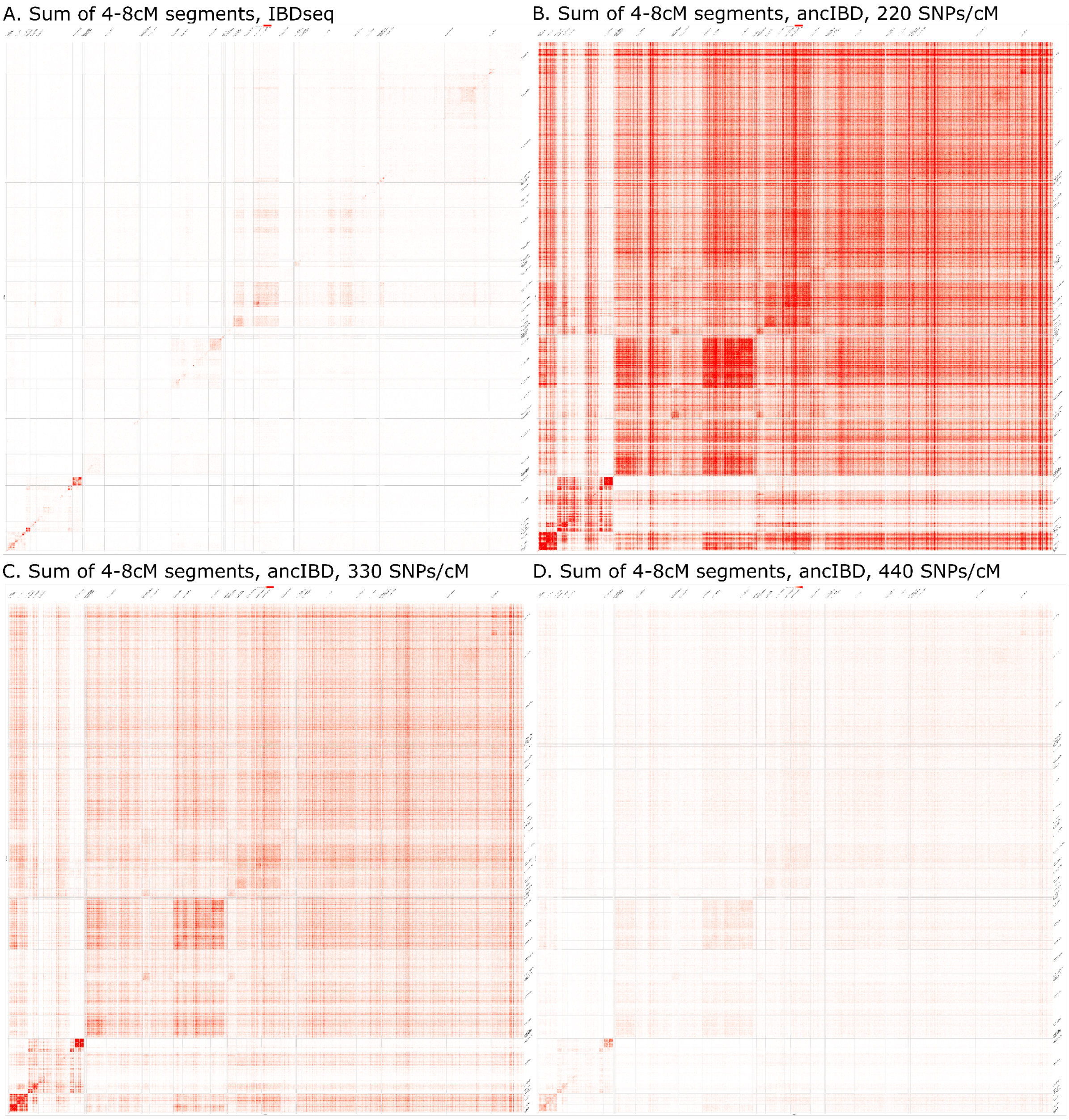

**Fig. S5.39. Heatmap showing the sum of IBD segments between 4-8cM for all pairs of ancient individuals, called by (A) IBDseq, (B) ancIBD at 220 SNP per cM, (C) ancIBD at 330 SNP per cM and (D) ancIBD at 440 SNP per cM .** Results from 0-100 cM are shown from white to red.

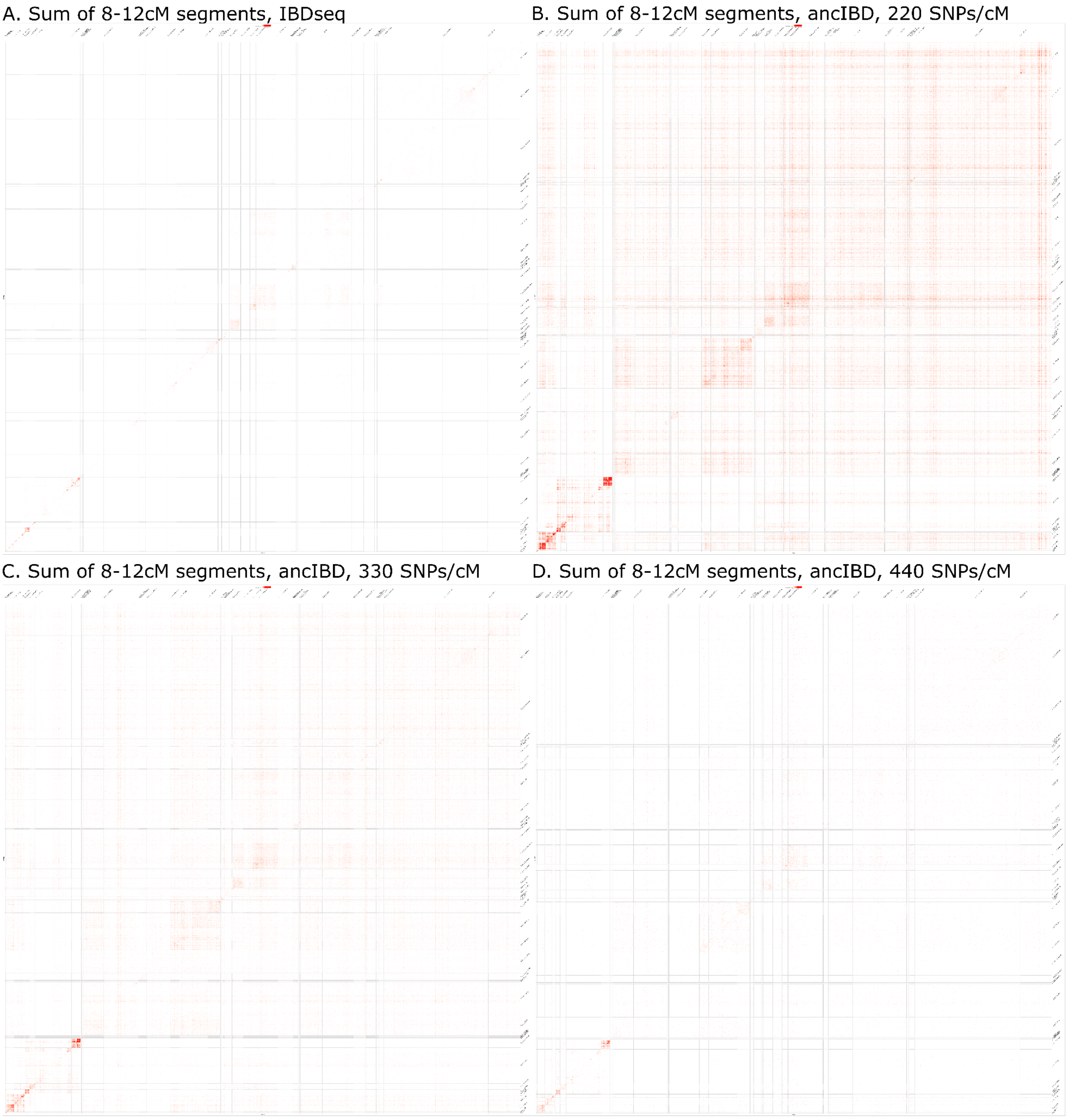

**Fig. S5.40 Heatmap showing the sum of IBD segments between 8-12cM for all pairs of ancient individuals, called by (A) IBDseq, (B) ancIBD at 220 SNP per cM, (C) ancIBD at 330 SNP per cM and (D) ancIBD at 440 SNP per cM .** Results from 0-100 cM are shown from white to red.

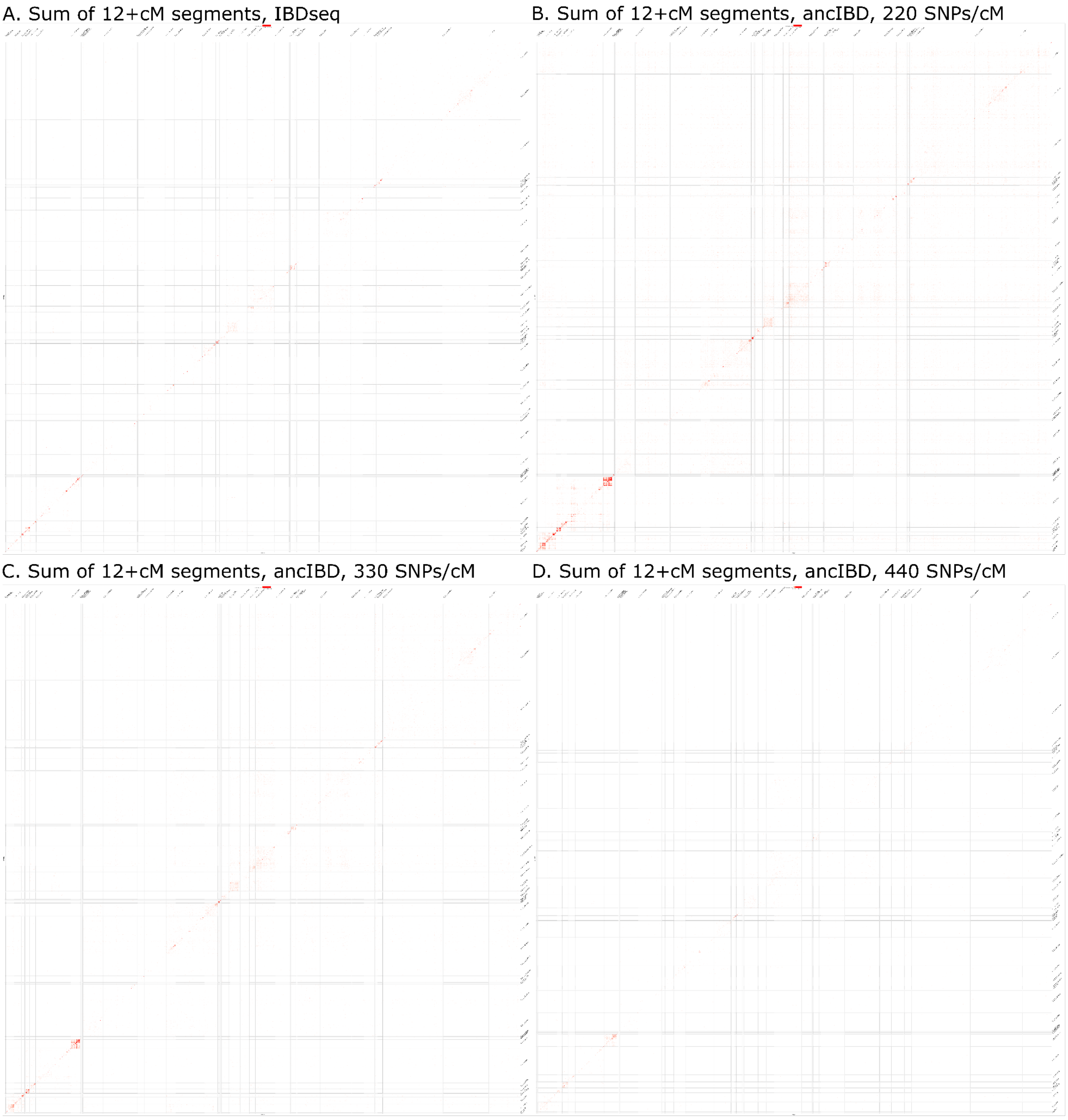

**Fig. S5.41. Heatmap showing the sum of IBD segments above 12cM for all pairs of ancient individuals, called by (A) IBDseq, (B) ancIBD at 220 SNP per cM, (C) ancIBD at 330 SNP per cM and (D) ancIBD at 440 SNP per cM .** Results from 0-100 cM are shown from white to red.

**Mixture Modelling Tests.**

In addition, to identify how the mixture modelling results varied by segment length, we aggregated and performed mixture modelling on subsets of the data with lower limits of 1cM, 2cM, 4cM, 8 cM and 12cM for individual segments. The mixture modelling failed when limited to segments above 8 or 12 cM, due to the small number of long, shared fragments between clusters.

When attempting to replicate the transition in England related to Anglo-Saxon migrations using Bell Beaker and Corded Ware related sources, transition is apparent in all IBDseq runs using the shorter cut offs (1cM+ and 2cM+) but not the longer (4cM+) (Fig. S5.42.). For all ancIBD runs with these sources, the transition is not apparent.

When including Iron Age sources from Scandinavia (Set 6), a transition around 1600 BP is apparent for all sets and cut-offs (Fig. S5.43). However, for all ancIBD runs and the >4cM IBDseq run, there are large amounts of noise, with large error bars and the presence of unexpected sources. Furthermore, the ancIBD runs fail to distinguish between the Southern Scandinavian ancestry arriving with the Saxons from and the Eastern and Western Scandinavian ancestry arriving with the Vikings[78].

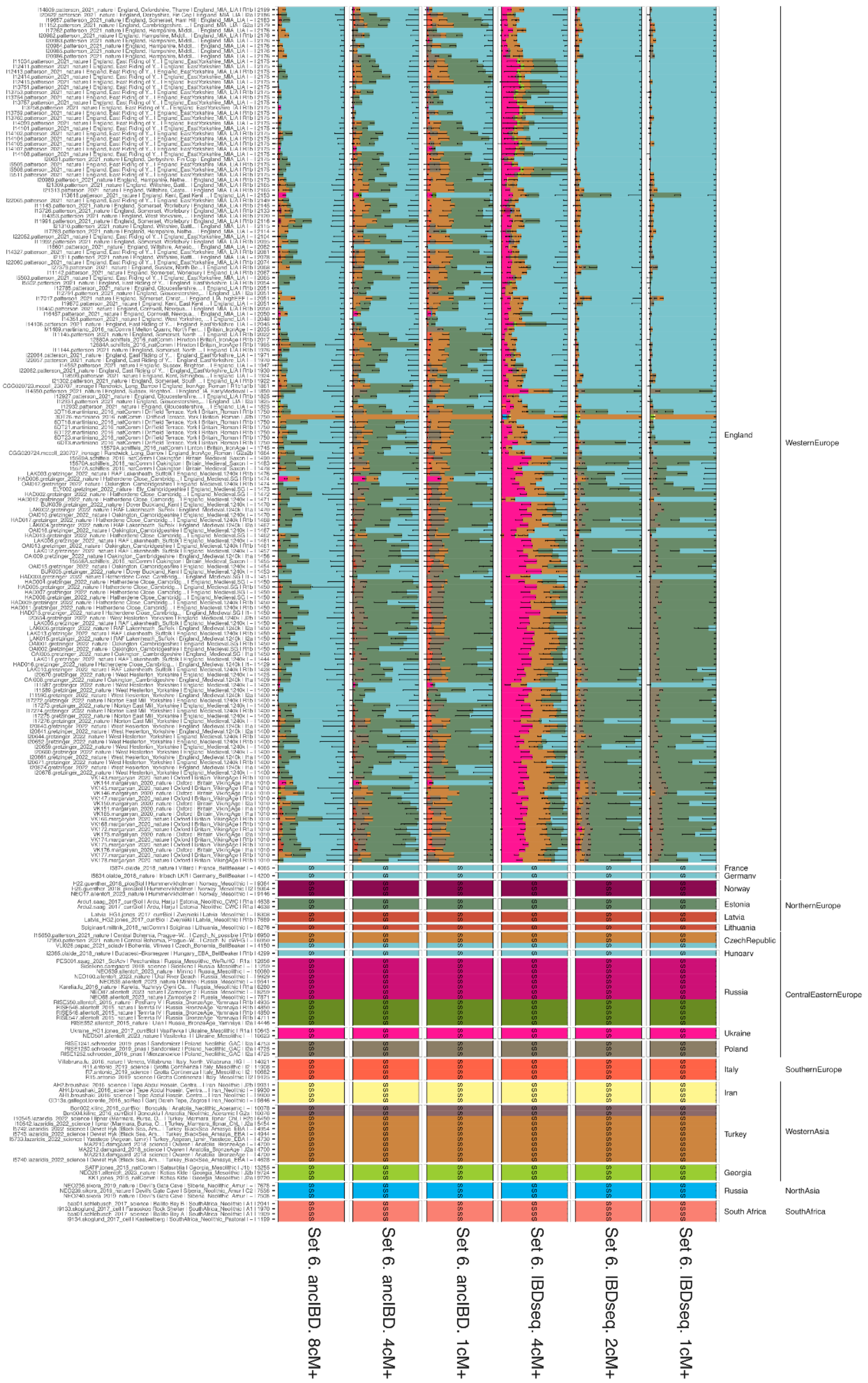

**Fig. S5.42. A subset of the mixture modelling results (set 4) highlighting Bell Beaker-related (blue) and Corded-Ware-related (grey green), for ancIBD >4cM, ancIBD >2cM, ancIBD >1cM, and IBDseq >4cM, IBDseq >2cM, IBDseq >1cM (left to right), showing samples from England between 2200 BP and 1400 BP, and Vikings from 1010 BP.**

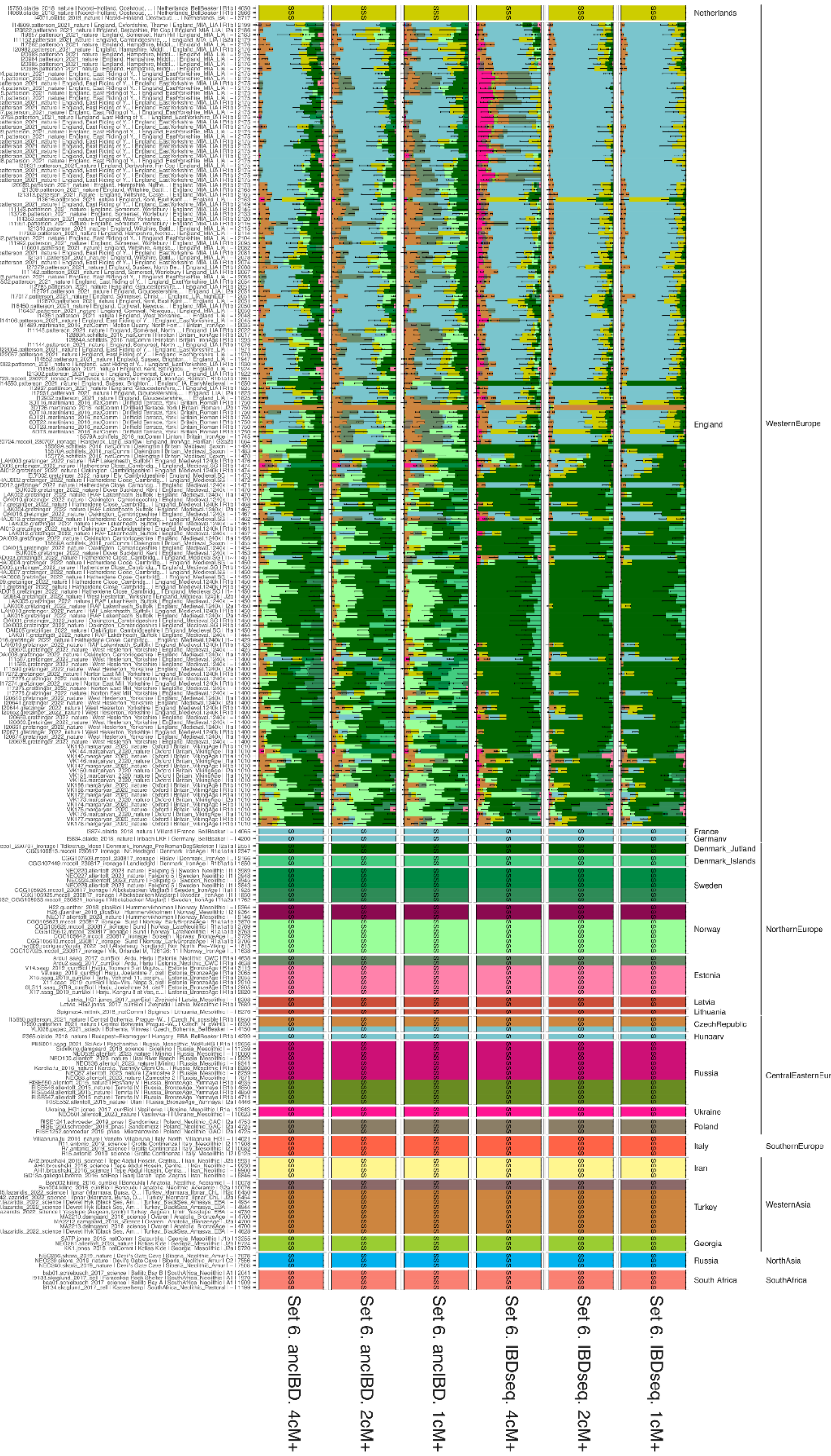

**Fig. S5.43. A subset of the mixture modelling results (set 4) highlighting Bell Beaker-related (blue), Jutland Iron Age (dark green) and Norwegian Iron Age (bright green),**

**for ancIBD >4cM, ancIBD >2cM, ancIBD >1cM, and IBDseq >4cM, IBDseq >2cM, IBDseq >1cM (left to right), showing samples from England between 2200 BP and 1400 BP, and Vikings from 1010 BP.**

#### S5.3.6 Impact of temporal differences and cluster size.

As IBD decays in length over time, the more temporally distant one source population is to a target, relative to another source population, potentially may bias results. To investigate this, we modelled the Neolithic Danish individuals (5700 - 4900 BP) with various sources, who have previously been identified as carrying both Western Hunter Gatherers and Neolithic Farming ancestry[29].

We used four source sets with varying temporal and geographical distance from the targets (**Fig S5.44**):

- a 'base' set, with early Western Hunter-Gatherers from Italy (14,000 - 9,000 BP) early Anatolian Farmers (10,000 BP)
- a 'base_DkHG' set, with Danish Hunter-gatherers (6500 - 5700 BP) in the place of the early Anatolian farmers
- a 'base_ltAna' set, with later Anatolian Hunter-gatherers (8300 - 7900 BP)
- A 'base_euNeo' set, with late European Farmers (6300 - 6000 BP) in the place of the early Anatolian farmers

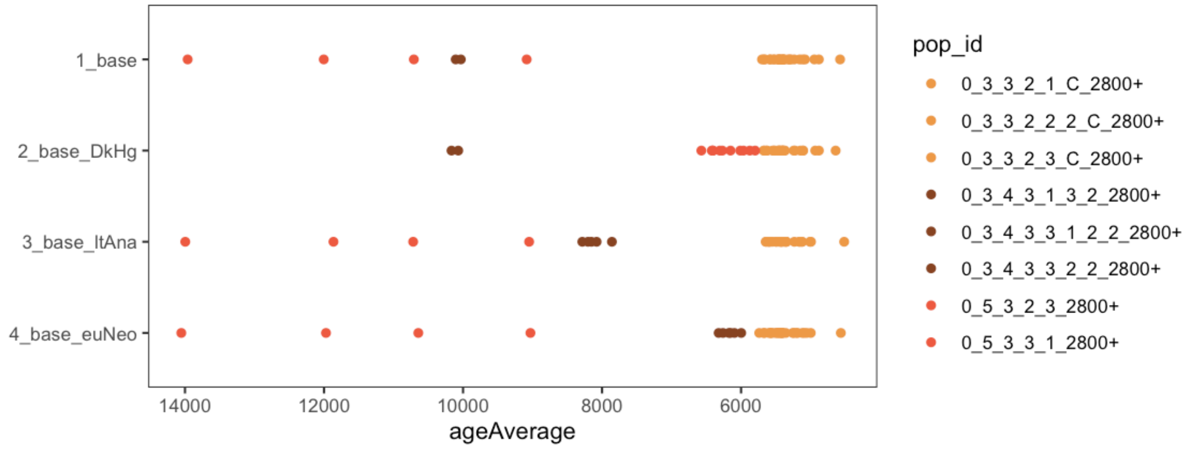

**Fig S5.44. Timeline showing the different source clusters for Hunter-Gatherers (red) and Farmers (dark brown), and the Danish Neolithic targets (light brown).**

For the four sets, we modelled the target individuals (**Fig S5.45**) and found subtle variations between the various sets. When using the closer Danish Hunter Gatherers, we see slightly less Farming ancestry. When using the later Anatolian farmers, we see slightly more farming ancestry. For the local farming ancestry, we see the highest proportion of Farming ancestry modelled. However, this source cluster itself is modelled with a mean of 10.1% Hunter-Gatherer ancestry, and as a result, when used as a source, it reduces the proportion of Hunter-Gatherer ancestry modelled in the Danish Neolithic individuals. When correcting for this, we find the error bars for all source sets to overlap. This indicates that at this density and time span, IBD mixture modelling can effectively model individuals thousands of years apart, by leveraging intermediate clusters within the dataset.

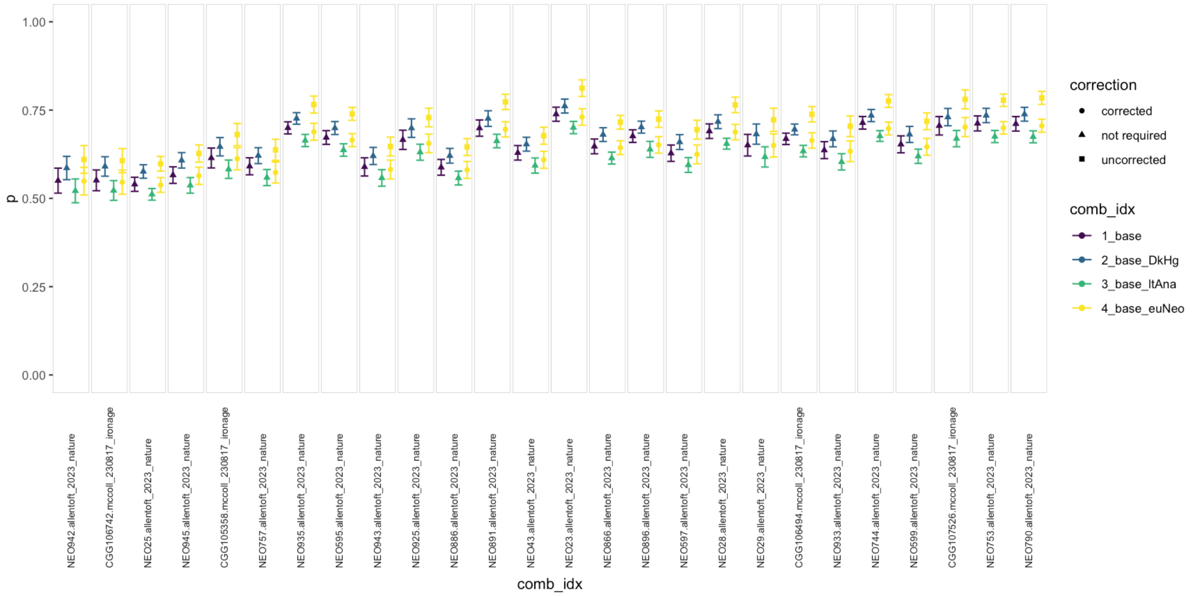

**Fig S5.45. Mixture Modelling results showing the proportion of Farming Ancestry in Danish Neolithic target individuals, when using the four sets of source clusters.** To account for the ~10% HG ancestry present in the Farming sources in the 4_base_euNoe set, a corrected value accounting for this is provided (circles).

In addition to potential bias as a result of temporal distance, we also address potential impacts of varied source group sizes. Although the proportion of IBD each individual shares with each cluster is normalised by the number of individuals in the cluster, using less individuals as sources may potentially create bias.

To investigate the impact of the size of source clusters on the modelling results, we remodelled all samples with varying numbers of source individuals in the Bronze Age sources for East (0_1_2_3_C_2800+, n= 4), West (0_1_4_3_2800+, n = 5) and South (0_1_4_1_C_2800+, n = 4) Scandinavia (Table S5.6), the clusters most significant to the results of this study.

**Table S5.6. Number of source individuals in the various sets for source size comparisons**

| Set | South Scan BA (0_1_4_1_C_2800+ | West Scan BA (0_1_4_3_2800+ | East Scan BA (0_1_2_3_C_2800+) |
|---|---|---|---|
| main_set_5 | 4 | 5 | 4 |
| 2 | 1 | 1 | 1 |
| 3-5 | 1-3 | 5 | 4 |
| 6-9 | 4 | 1-4 | 4 |
| 10–12 | 4 | 5 | 1-3 |

When modelling individuals associated with the Iron Age in Western Europe (0_2_1_4_2800-), Norway (0_1_2_4_5_C_2800-), Sweden (0_1_2_1_2_C_2800-), Denmark_Isles (0_1_2_2_3_C_2800-), and Denmark_Jutland (0_1_3_3_C_2800-), we find varying the number of individuals in the source cluster has no apparent impact (Fig. S5.46).

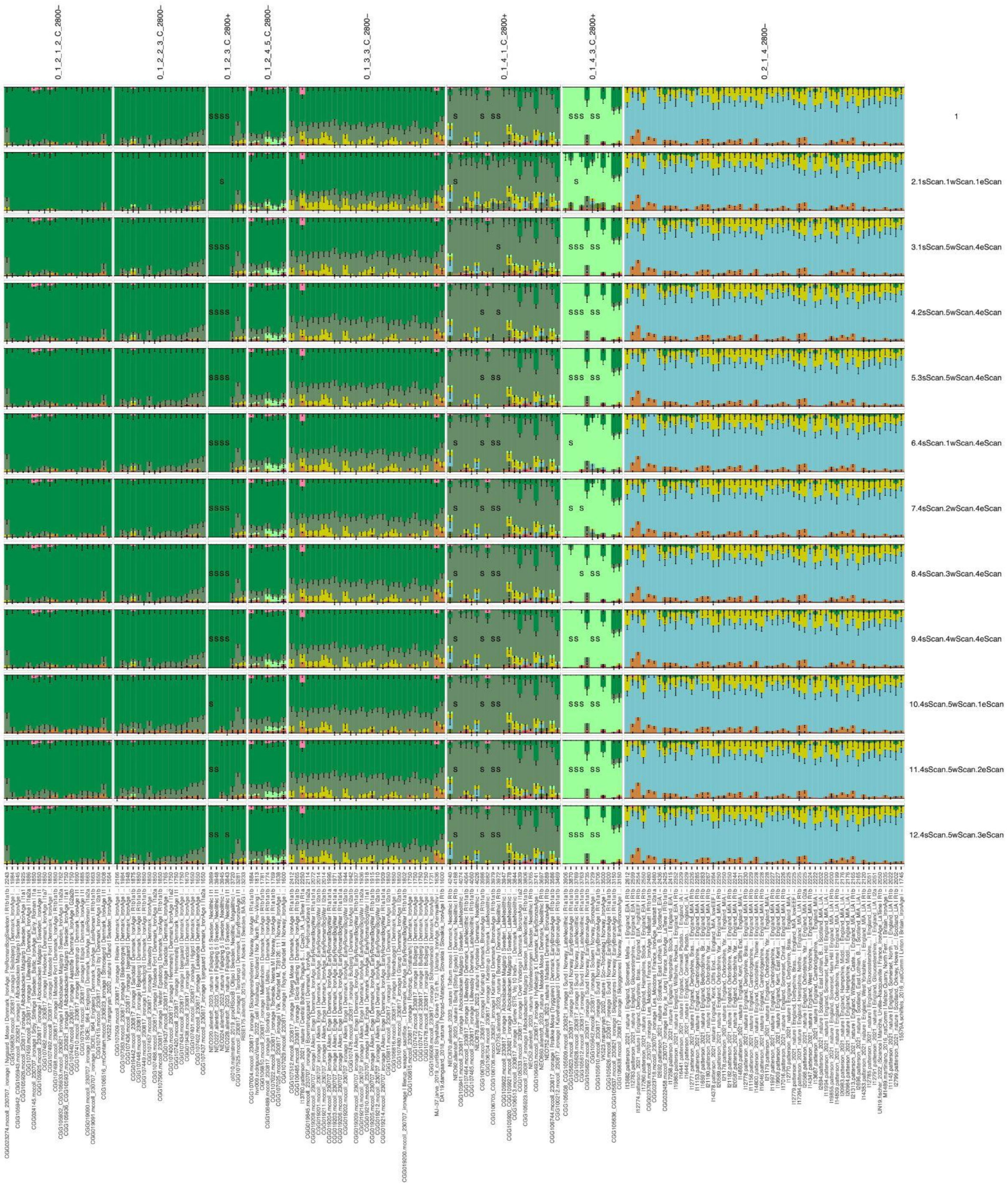

**Fig S5.46. Mixture Modelling results showing variation in proportions of Eastern Scandinavian (0_1_2_3_C_2800+), Southern Scandinavian (0_1_4_1_C_2800+) and Western Scandinavian (0_1_4_3_C_2800+) for various combinations of the number of source individuals.** The top row corresponds to the main mixture modelling set 5. The number of individuals in each set is indicated on the right, and the specific source individuals are indicated with an 'S'.

##### S5.3.7 Correlation between Genetics, Archaeology and Linguistic Groupings

To assess whether there was any statistical significance genetic results and individuals believed to be linked with specific cultural or linguistic groups, we assigned individuals to each of these groups where possible. We assigned individuals to 'Yamnaya', 'Corded Ware' and 'Bell Beaker' groups based on the information present in the metadata table from published studies (Supplementary Table S5.7). For the linguistic groups, we assigned individuals to 'Celtic' from England, Scotland and France, dated to the pre-Roman Iron Age (2800 - 2000 BP). For Germanic, we included individuals from early Iron Age Denmark (2800 -1575 BP), and Sweden and Norway from the approximate time of the appearance of Runes (2200 - 1575 BP), ending at the Migration Period. The samples and archaeological or linguistic assignment can be found in Supplementary Table S5.7.

For each of the three sources, we calculated the variance of the modelling results between the groups of individuals, based on the archaeological and linguistic assignments, using *aov* (ANOVA) in R. We found highly significant ($p < 0.001$) differences for groups within each of the source sets (Supplementary Table S5.7). We then compared the means of the significant ANOVA results by applying a Tukey HSD (Honest Significant Difference) test to the ANOVA results using *TukeyHSD* in R.

With regards to the archaeological groupings, we found for the 0_2_1_4_3_1_C_2800+ source, the group with the highest average proportion modelled by this source was the individuals assigned to the Bell Beaker group, and the mean was most significantly different from the other two Archaeological groupings (Fig. S3.5.7.1, Supplementary Table S5.7). The same distinction was present for 0_2_2_2_3_C_2800+ and the Corded Ware group, and for 0_2_2_3_2800+ and the Yamnaya group.

For the linguistic groupings, we found the largest difference in the means between the comparisons of Celtic and Germanic (Fig. S3.5.7.2, Supplementary Table S5.7). We found that for 0_2_1_4_3_1_C_2800+, the group with the highest average was the various groups assigned as Celtic, with the largest differences to the Germanic groups. Similarly, for 0_2_2_2_3_C_2800+ as a source, we found the highest average was the various groups assigned as Germanic, with the largest differences to the Celtic groups.

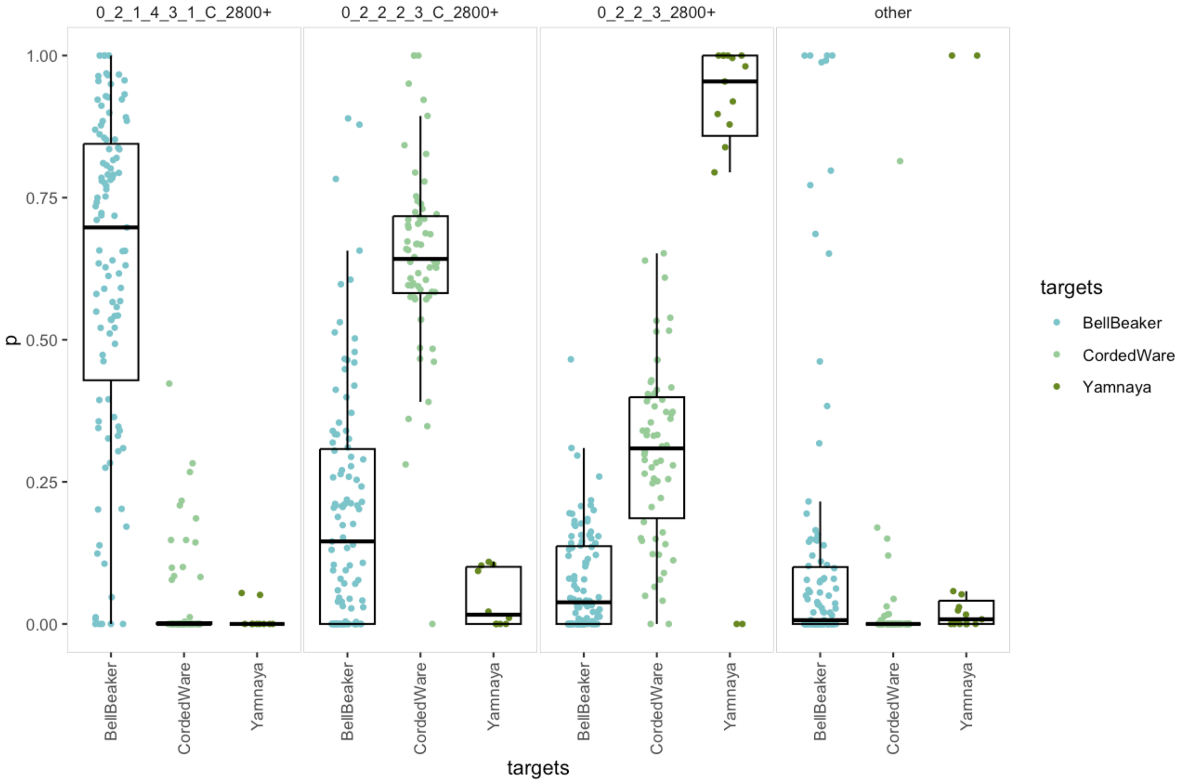

**Fig. S3.47. Proportion of the Steppe related ancestry in ancient individuals archaeologically assigned to the Bell Beaker, Corded Ware or Yamnaya Cultures.**

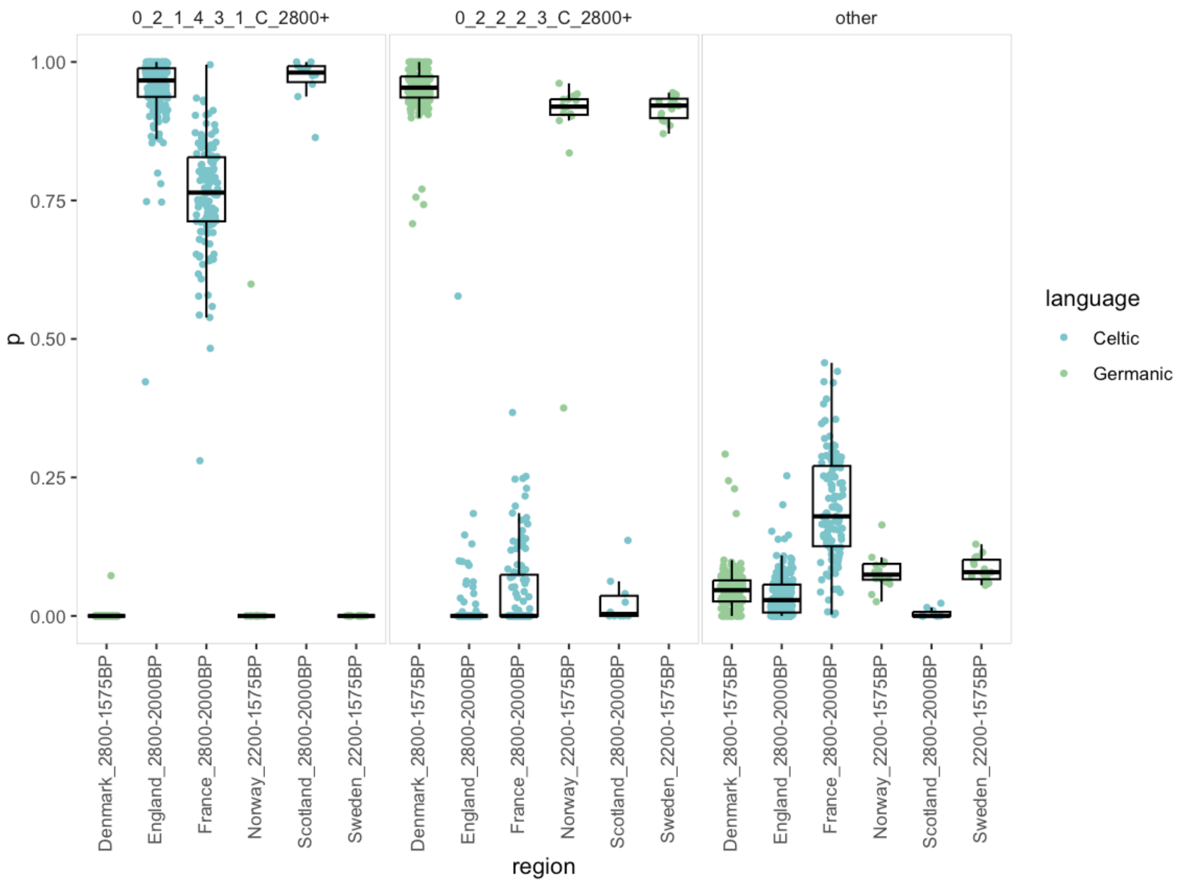

**Fig. S3.48. Proportion of the Steppe related ancestry in Iron Age individuals originating in regions assigned linguistically as Celtic or Germanic.**

#### S5.3.8. Spatiotemporal kriging

To understand the general trends of changes in ancestry that occurred throughout time, we performed spatiotemporal ordinary kriging as described by Racimo et al[39]. By using observed mixture modelling results and C14 or context dates for all ancient individuals, we are able to infer ancestries at unsampled locations and times.

The parameters used matched those of Racimo et al[39] with a few changes, to account for the differences in the sampling and focus of the studies (Table S5.8.). Resolution was increased from 5,000 points to 12100 points, as a result of the increased number of samples and shorter time range of interest (5000 years rather than 10,800 years).

For Mixture Modelling Sets 1 - 5, in which the patterns are relevant across Europe, the 'range' parameter of the vgmST function matched that of Racimo et al[39]: 5000. For Sets 5-7, in which patterns are relevant on a more local scale, the range parameter was reduced to 2000, to allow for a distinction between Jutland, the Danish Isles and Sweden to be visible. Similarly, the number of time slots and lags varied correspondingly for the sets, as described in (Table S5.8.).

Table S5.8. Variation in to parameters applied in Racimo et al[39] and the current study

| Parameter | Racimo et al 2023 | Europe Wide | North/West Europe |
|---|---|---|---|
| Time range | 10,800 - 0 BP | 5600 - 600 BP | 5600 - 600 BP |
| Number of time slots | 19 | 33 | 51 |
| Lags | 50 | 50 | 10 |
| Resolution | 5000 | 12100 | 12100 |
| Range (vgmST) | 5e3 | 5e3 | 2e3 |
| Minimum latitude | 35 | 35 | 50 |
| Max latitude | 72 | 70 | 70 |
| Minimum longitude | -20 | -10 | -10 |
| Maximum longitude | 80 | 30 | 20 |
| Admixture proportions | ADMIXTURE results for for HG (Hunter Gatherers), | IBD Mixture Modelling results for Sets 1-5 | IBD Mixture Modelling results from Set 5-7 |

| | | | |
|---|---|---|---|
| | NEOL (Neolithic Farmers) and YAM (Yamnaya) | | |
| Number of ancient individuals | 1546 | 3143 | 1825 |

**Results**.

Spatio-temporal kriging was able to be fit for the ancestries indicated in Table S5.3.9. The results are plotted in Figs S5.49 to S5.133. Timelines for specific points across Europe (see Fig S5.134) for mixture modelling sets 1-4 using the 'Europe Wide' parameters are plotted in Extended Data Figs 1-4, for mixture set 5-7 using the 'North/West' parameters are plotted in Extended Data Figs 5-7. The results for set 5, using the 'Europe Wide' parameters are plotted in Supplementary Fig. S5.174

**Table S5.9. Source sets that spatio-temporal kriging was performed on.** ^ and # indicate that 'Europe Wide' and 'North / West Europe' parameters were used respectively. * indicates sets where sampling was less dense and lag of 50 was applied.

| Source | Set 1^ | Set 2^ | Set 3^ | Set 4^ | Set 5^ | Set 5# | Set 6# | Set 7# |
|---|---|---|---|---|---|---|---|---|
| 0_1_1_1_2_1_1_2800 (Baltic-related) | | | | | X | X | X | X |
| 0_1_2_1_2_C_2800- (Swedish IA-related) | | | | | | | X | X |
| 0_1_2_2_3_C_2800- (Danish Isles IA-related) | | | | | | | X | X |
| 0_1_2_3_C_2800+ (E.Scan BA-related) | | | | | X | X | X | X |
| 0_1_2_3_C_2800- (Jutland IA-related) | | | | | | | X | X |

| | | | | | | | | |
|---|---|---|---|---|---|---|---|---|
| 0_1_2_4_5_C_2800- (Norwegian IA-related) | | | | | | | X | X |
| 0_1_3_2800- (N. German IA-related) | | | | | | | | X |
| 0_1_4_1_C_2800+ (S.Scan BA-related) | | | | | X | X | X | X |
| 0_1_4_3_C_2800+ (W.Scan BA-related) | | | | | X | X* | X* | X* |
| 0_2_1_2_5_1_C_2800+ (E.N.S BA-related) | | | | | X | X | X | X |
| 0_2_1_4_5_C_2800+ (Bell Beaker-related) | | | | X | X | X | X | X |
| 0_2_2_2_1_1_C_2800+ (Corded Ware-related) | | | | X | X | X | X | X |
| 0_2_2_3_2800+ (Yamnaya-related) | | X | X | X | X | | | |
| 0_3_3_1_2800+ (Euro. Neo.-related) | | | X | X | X | X | X | X |
| 0_3_3_3_C_2800+ (GAC-related) | | | X | X | X | | | |
| 0_3_4_1_1_2800+ (Anatolia BA-related) | | | X | X | X | X | X | X |
| 0_3_4_1_2_1_C_2800+ (Iran Neo-related) | | | | | | | | |
| 0_3_4_1_7_2800+ (CHG-related) | X | X | | | | | | |
| 0_3_4_3_3_2_2_2800+ (E. Anatolia-related) | X | X | X | X | X | | | |
| 0_4_2_1_4_2800+ (East Asian-related) | X | X | X | X | X | | | |
| 0_5_2_2_1_1_2800+ (Sweden HG-related) | X | X | X | X | X | X | X | X |
| 0_5_2_2_1_2_1_2800+ (EHG Ukraine-related) | X | | | | | | | |
| 0_5_2_3_2_2800+ (EHG Russia-related) | | | | | | | | |
| 0_5_3_1_3_2800+ (WHG Latvia-related) | X | X | X | X | | | | |
| 0_5_3_2_3_2800+ (WHG Italy-related) | X | X | | | | | | |
| 0_6_3_2800- (African-related) | | | | | | | | |

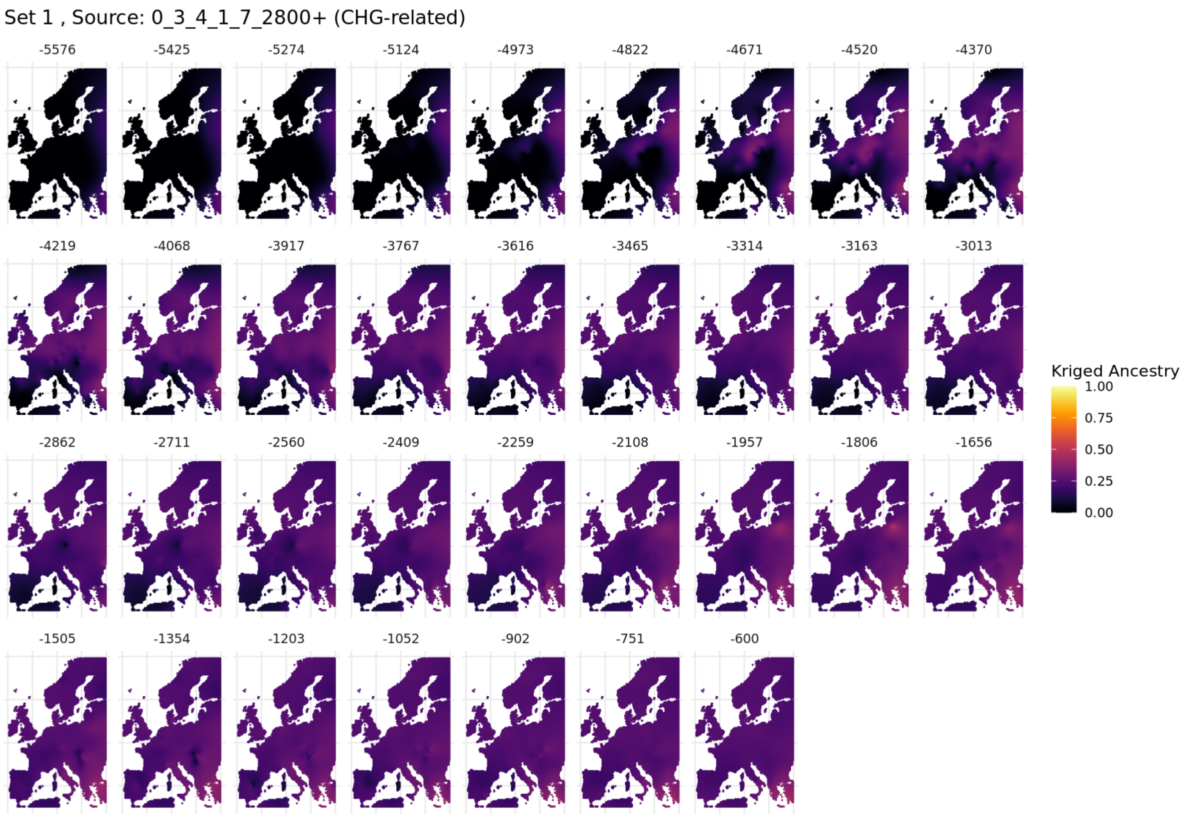

**Fig. S5.49. Spatiotemporal kriging results for Set 1 using 'Europe-Wide' parameters.**

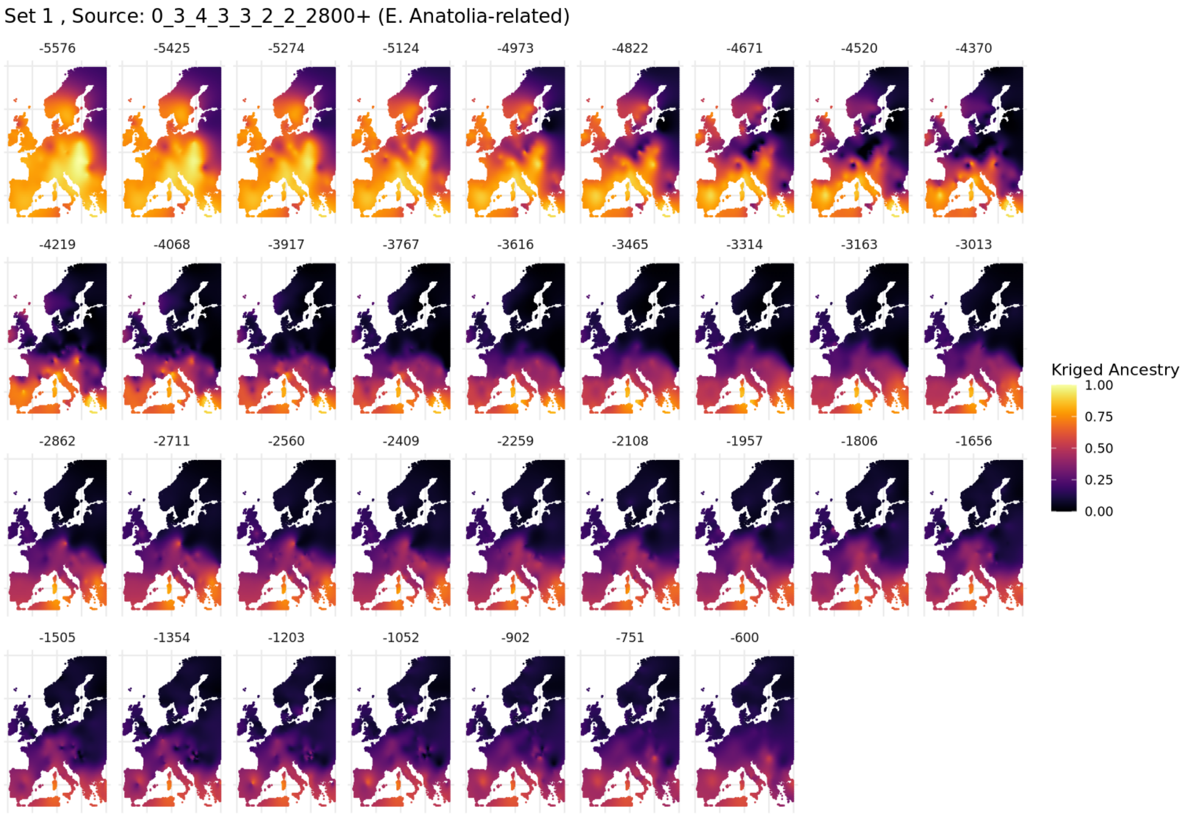

**Fig. S5.50. Spatiotemporal kriging results for Set 1 using 'Europe-Wide' parameters.**

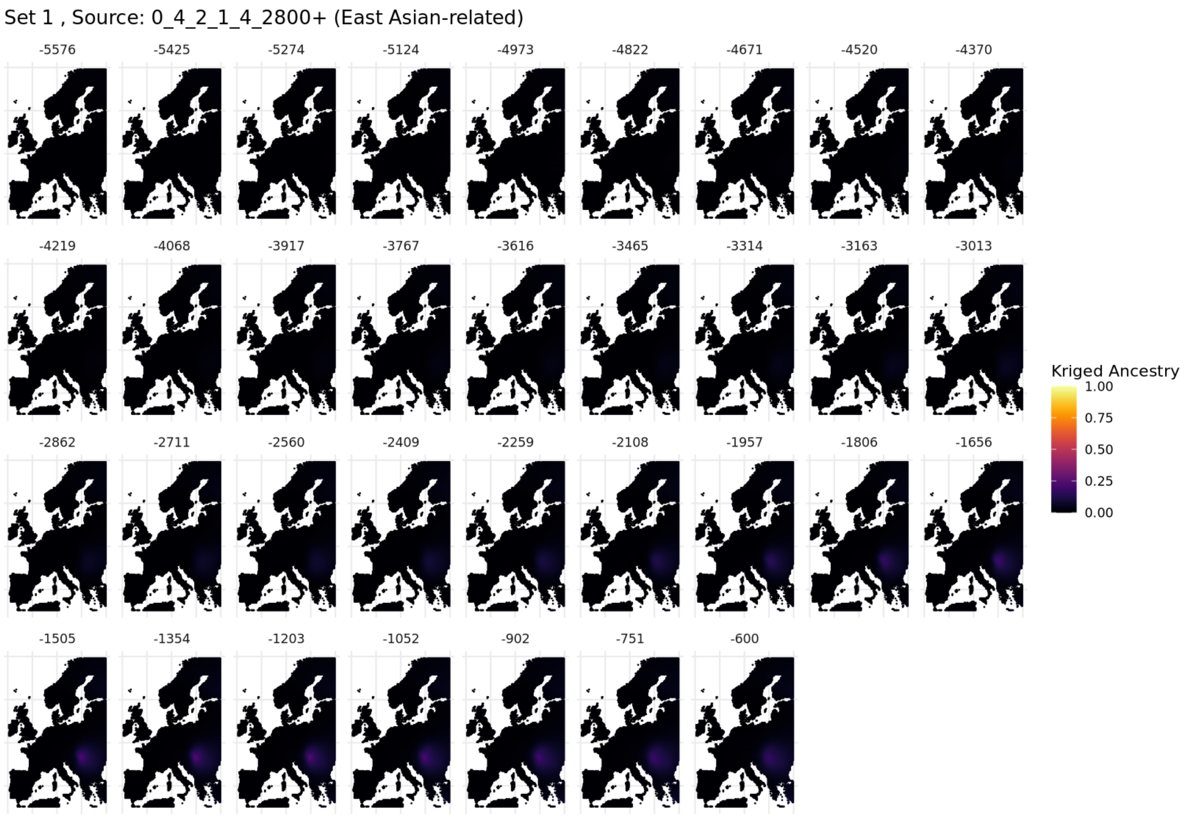

**Fig. S5.51. Spatiotemporal kriging results for Set 1 using 'Europe-Wide' parameters.**

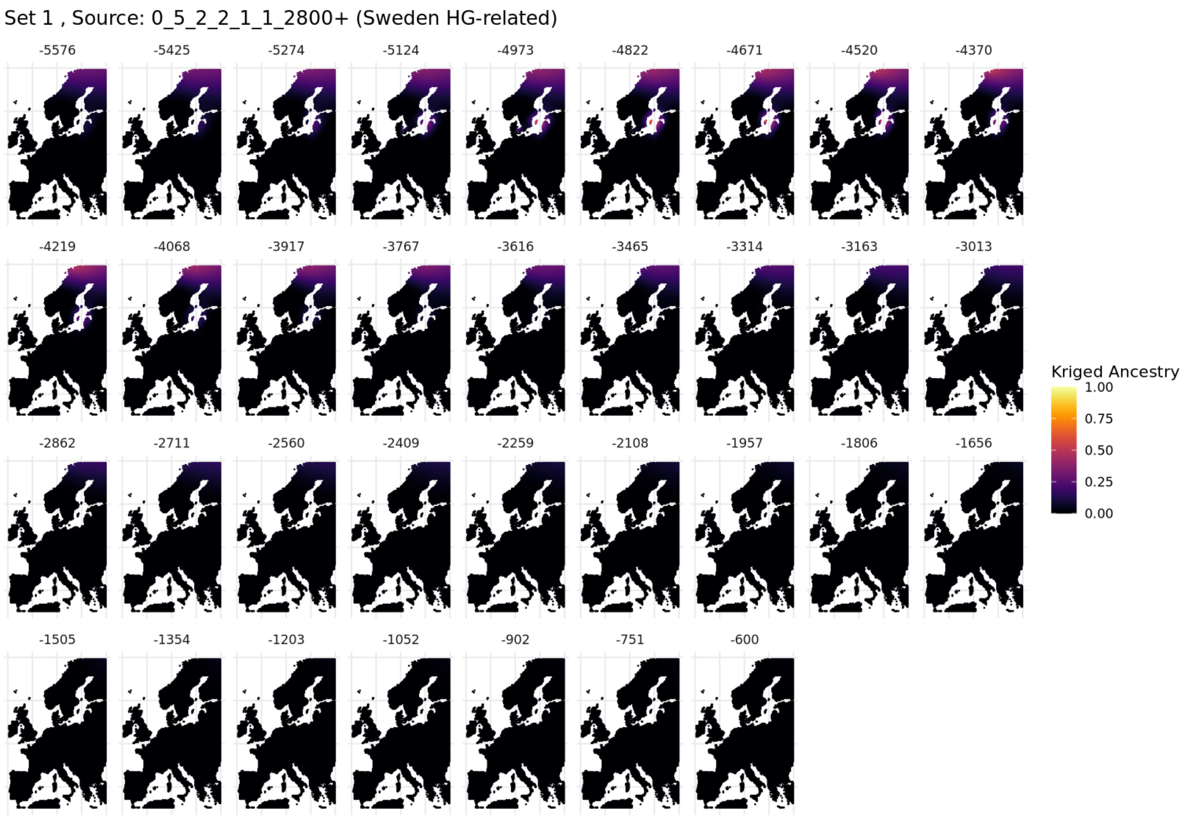

**Fig. S5.52. Spatiotemporal kriging results for Set 1 using 'Europe-Wide' parameters.**

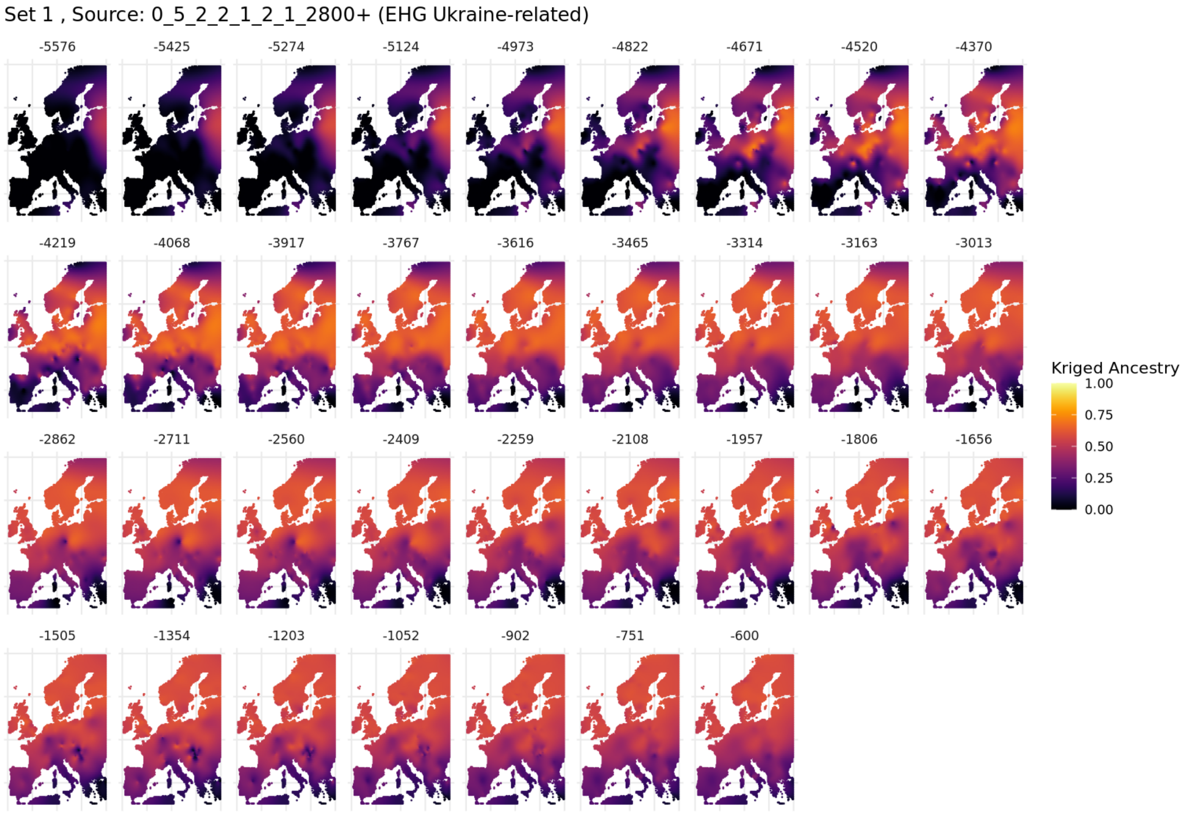

**Fig. S5.53. Spatiotemporal kriging results for Set 1 using 'Europe-Wide' parameters.**

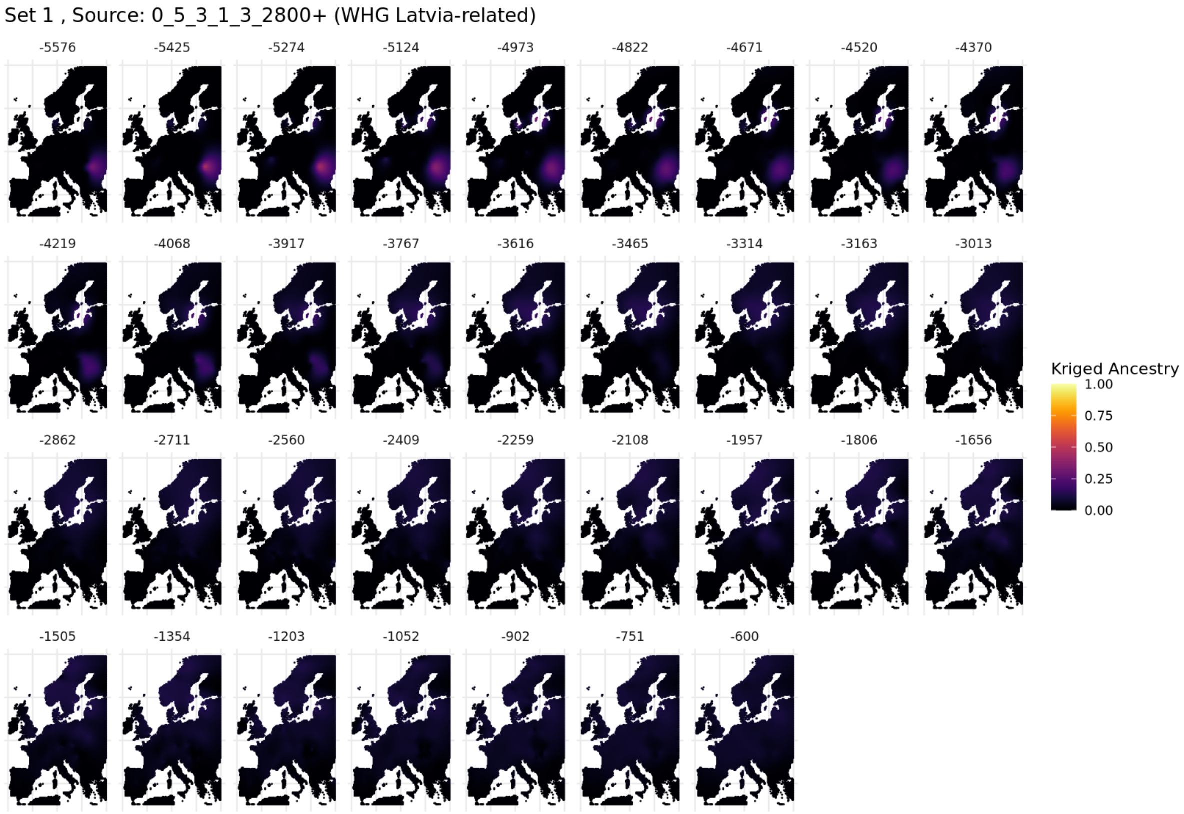

**Fig. S5.54. Spatiotemporal kriging results for Set 1 using 'Europe-Wide' parameters.**

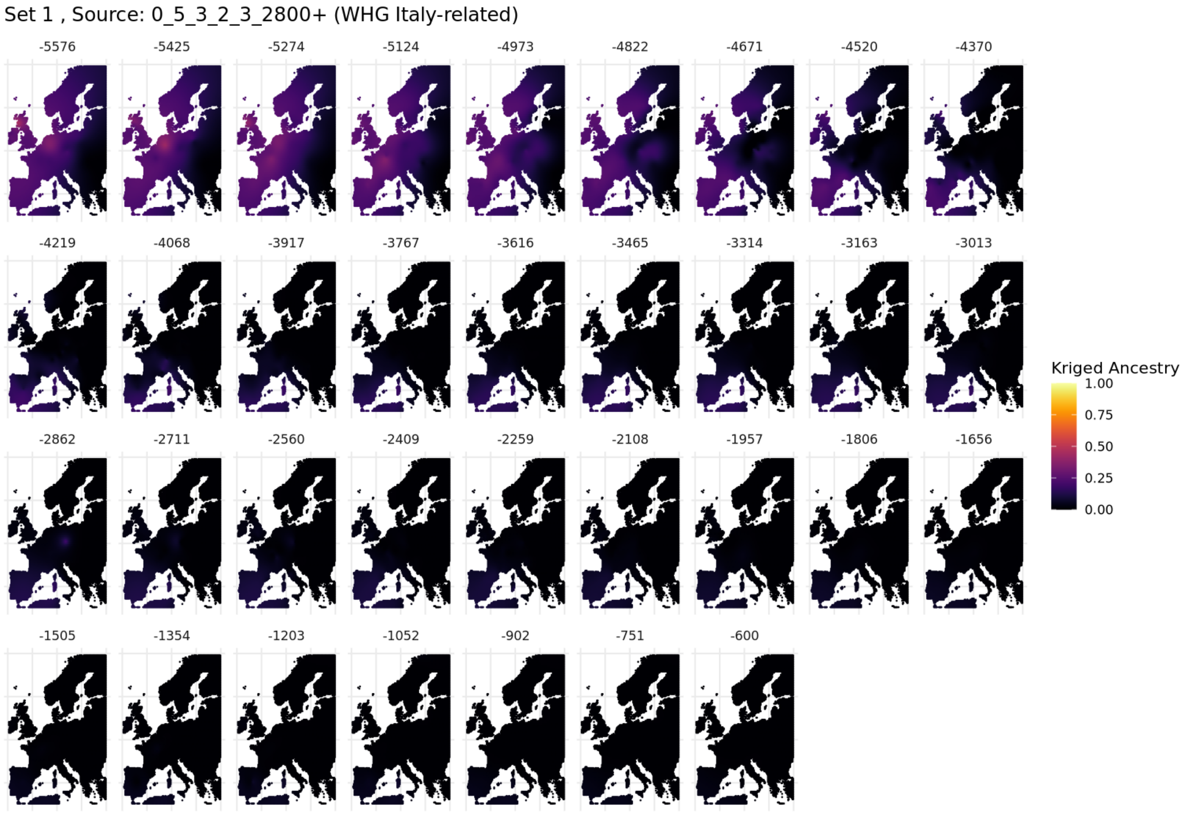

**Fig. S5.55. Spatiotemporal kriging results for Set 1 using 'Europe-Wide' parameters.**

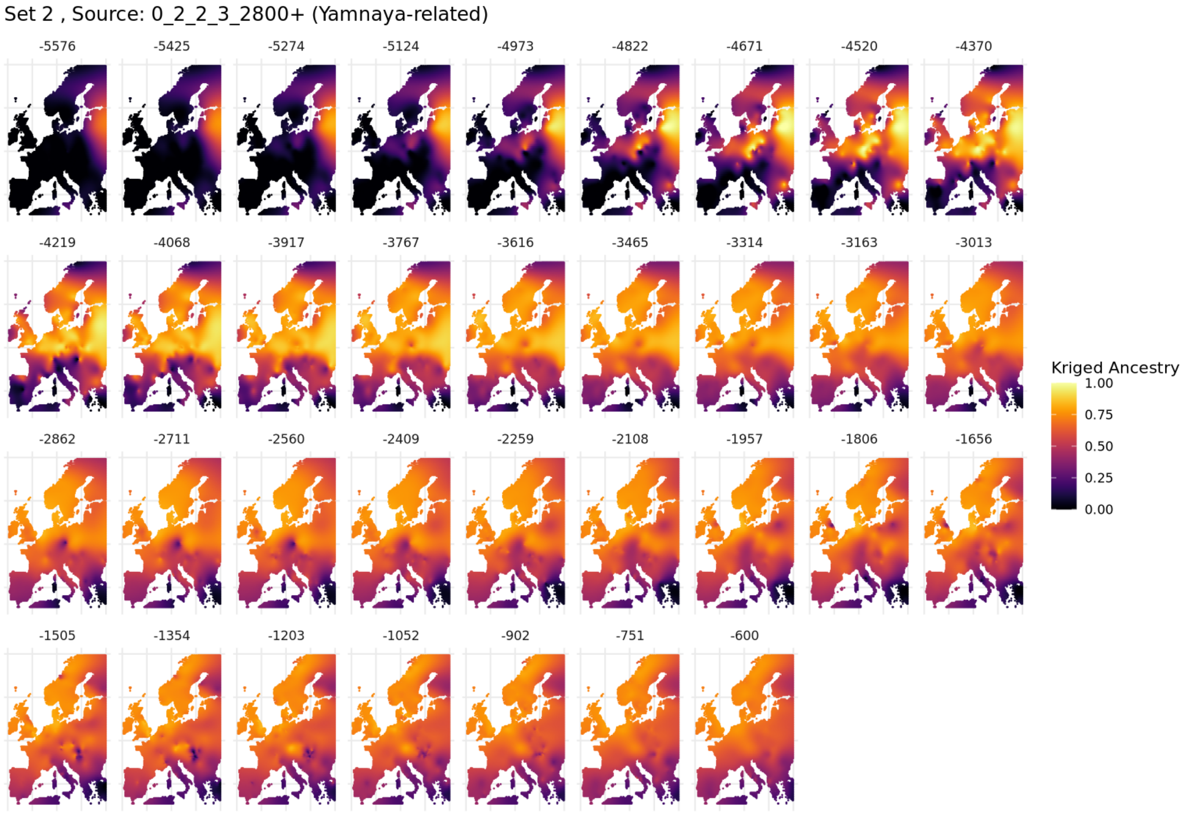

**Fig. S5.56. Spatiotemporal kriging results for Set 2 using 'Europe-Wide' parameters.**

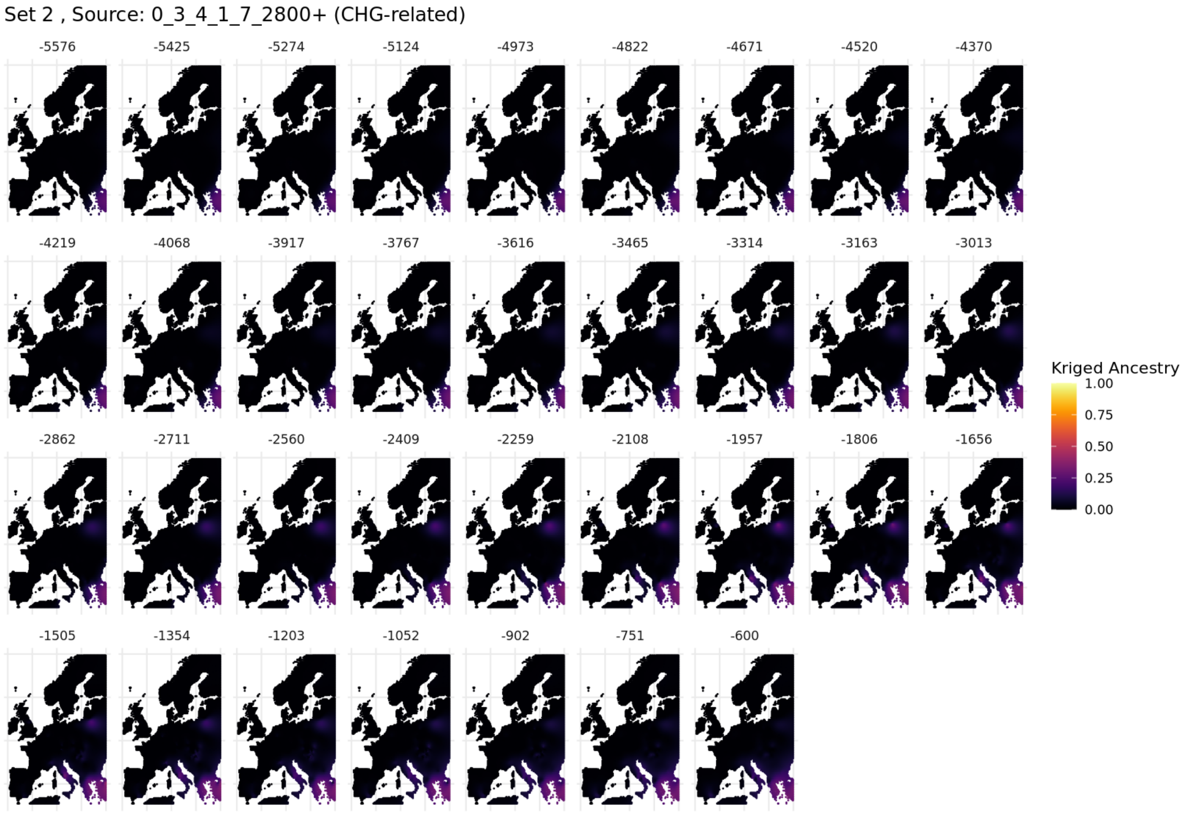

**Fig. S5.57. Spatiotemporal kriging results for Set 2 using 'Europe-Wide' parameters.**

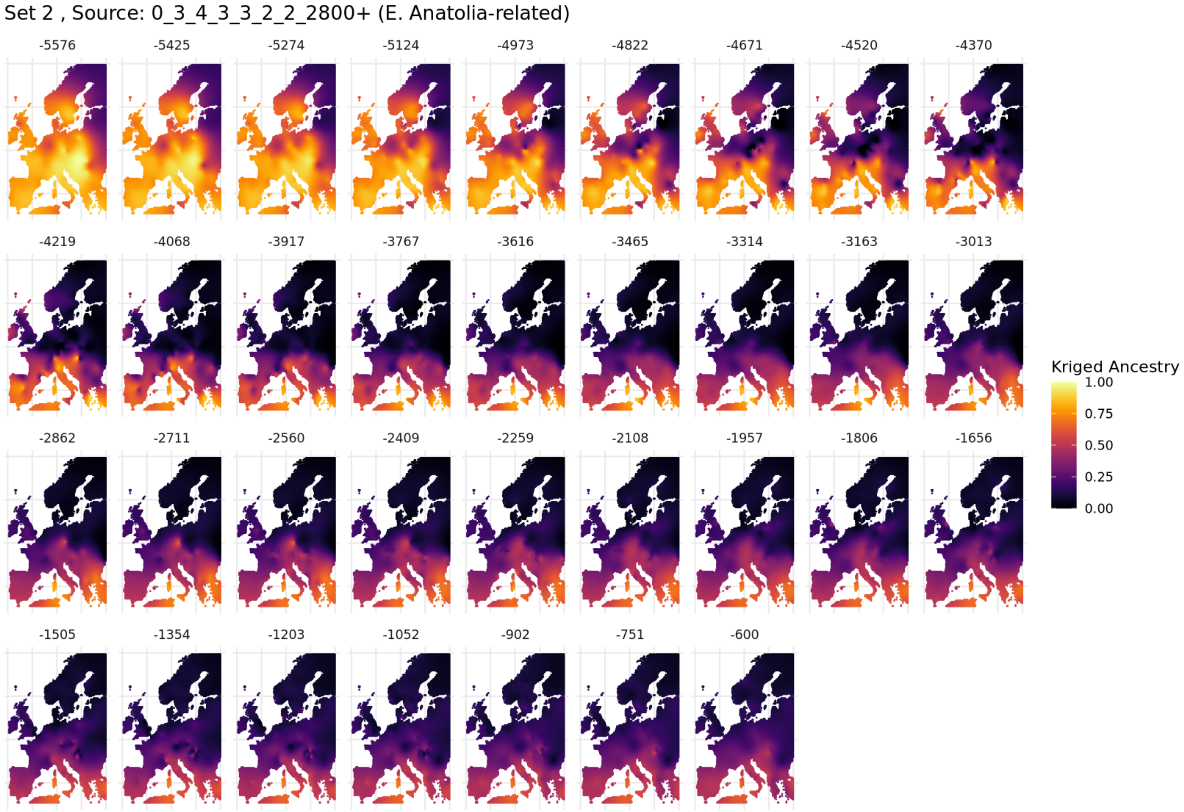

**Fig. S5.58. Spatiotemporal kriging results for Set 2 using 'Europe-Wide' parameters.**

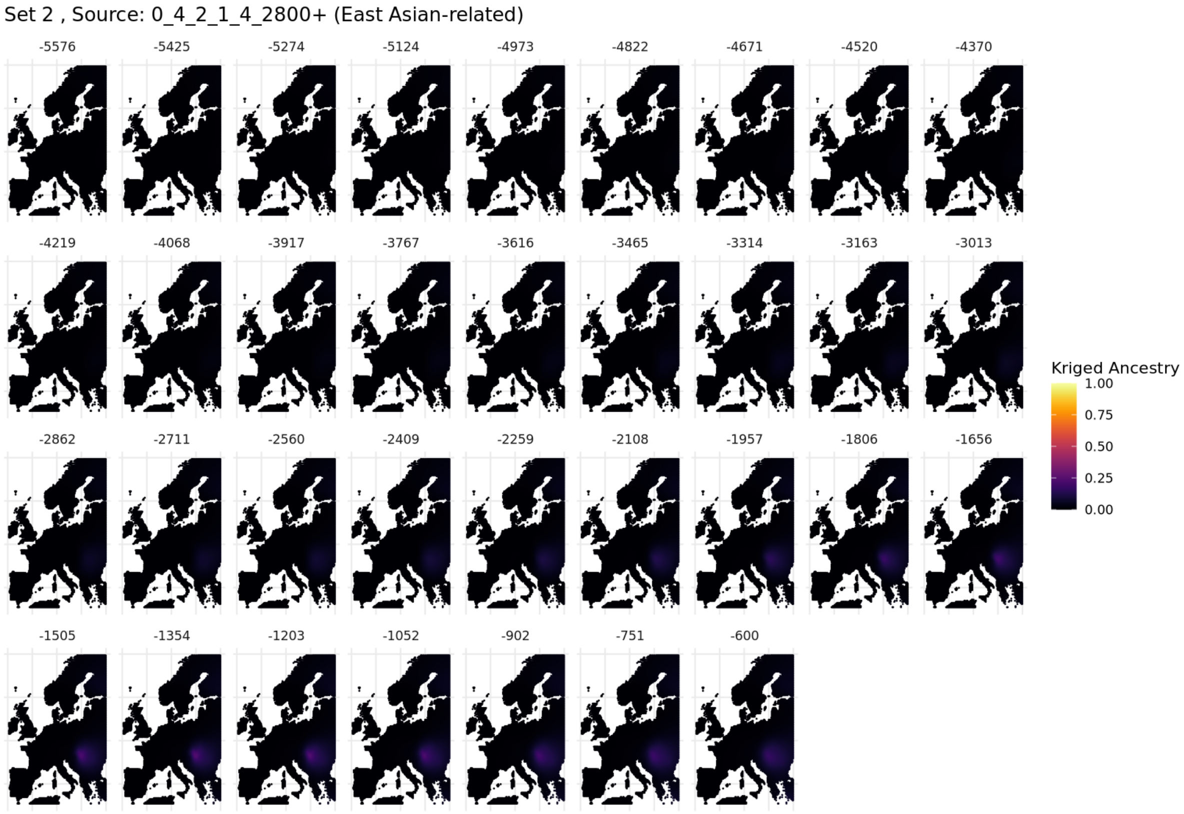

**Fig. S5.59. Spatiotemporal kriging results for Set 2 using 'Europe-Wide' parameters.**

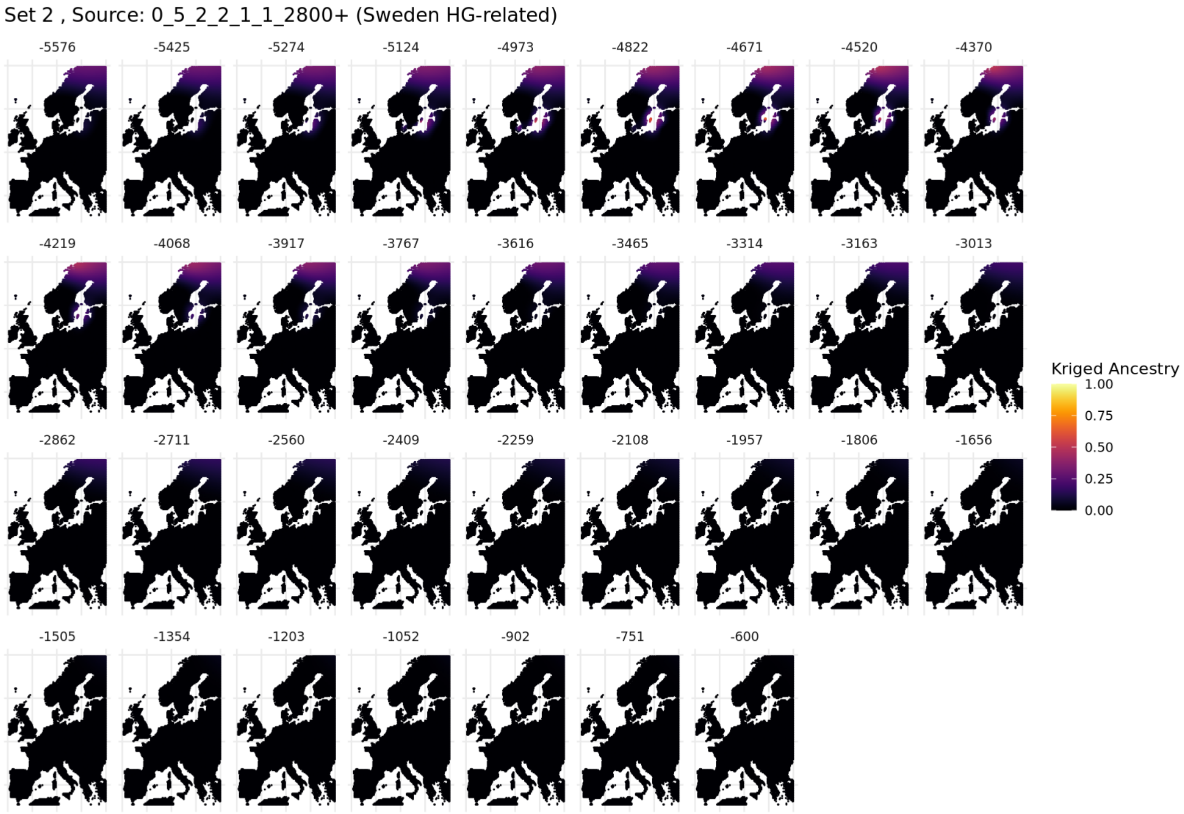

**Fig. S5.60. Spatiotemporal kriging results for Set 2 using 'Europe-Wide' parameters.**

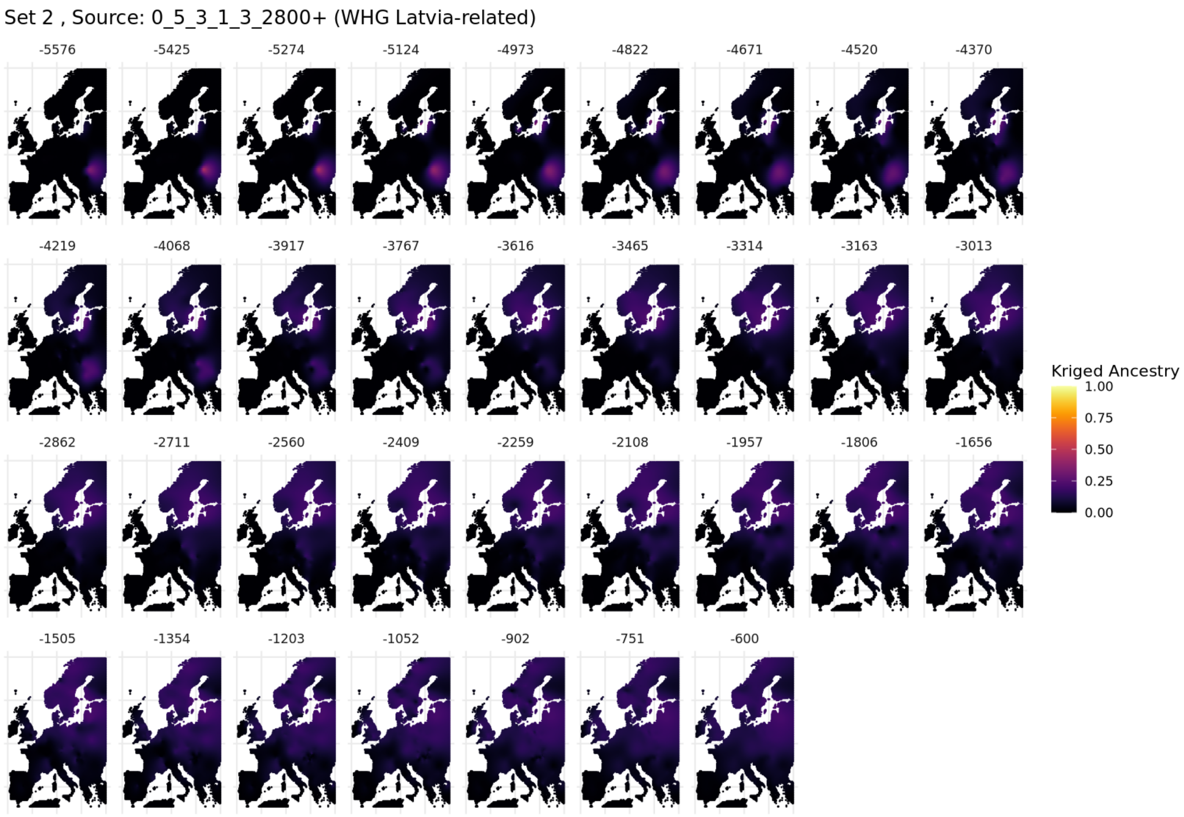

**Fig. S5.61. Spatiotemporal kriging results for Set 2 using 'Europe-Wide' parameters.**

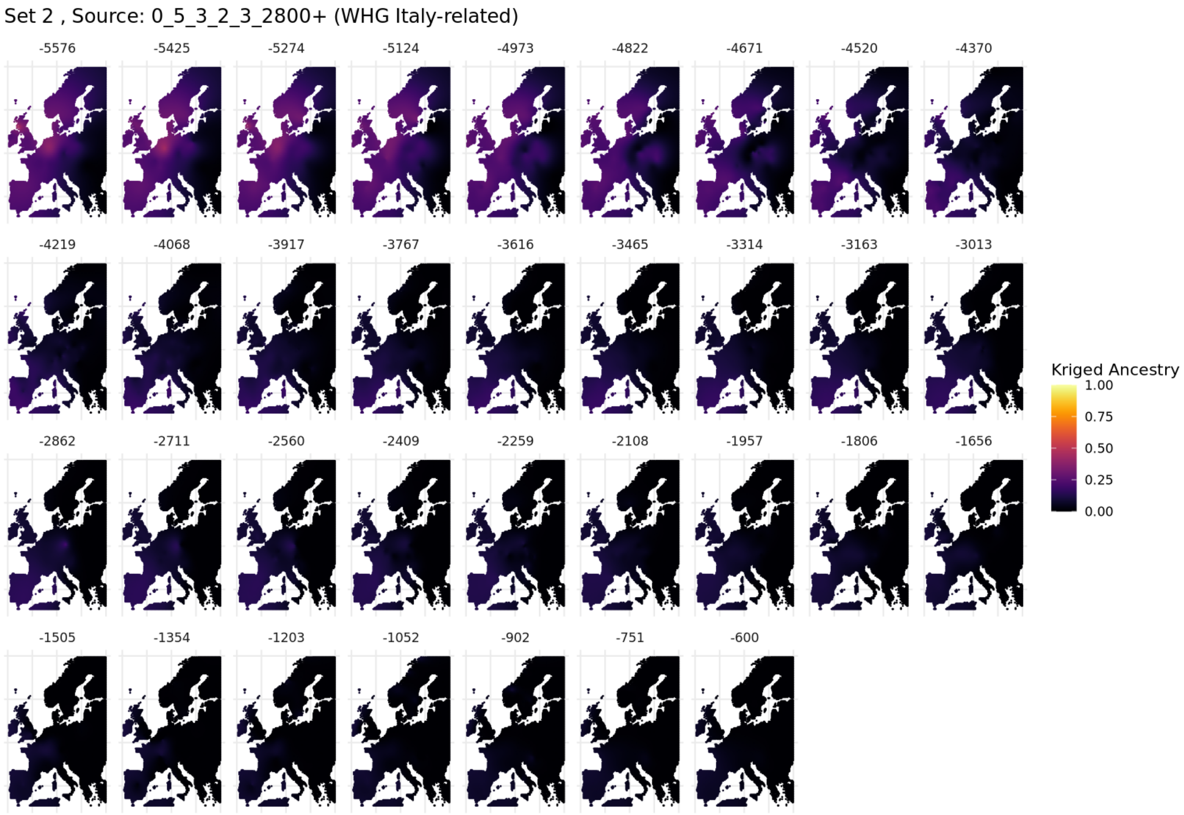

**Fig. S5.62. Spatiotemporal kriging results for Set 2 using 'Europe-Wide' parameters.**

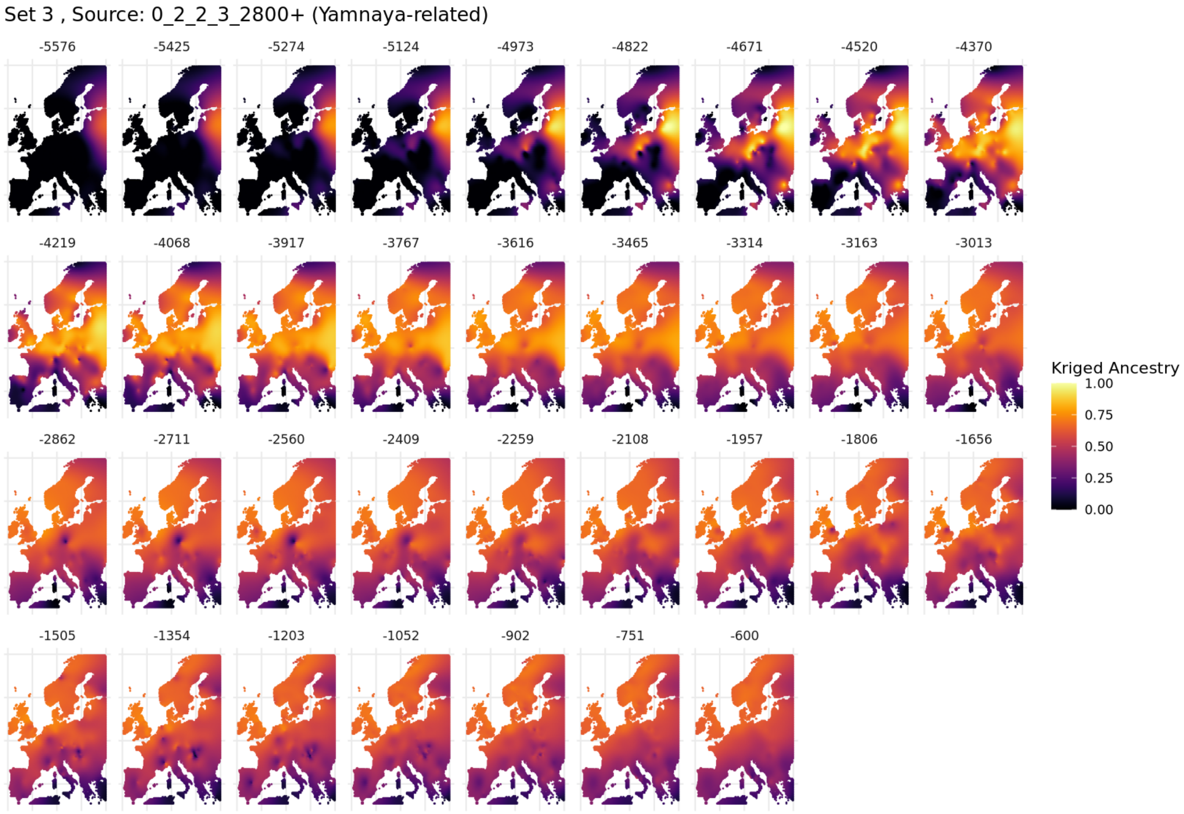

**Fig. S5.63. Spatiotemporal kriging results for Set 3 using 'Europe-Wide' parameters.**

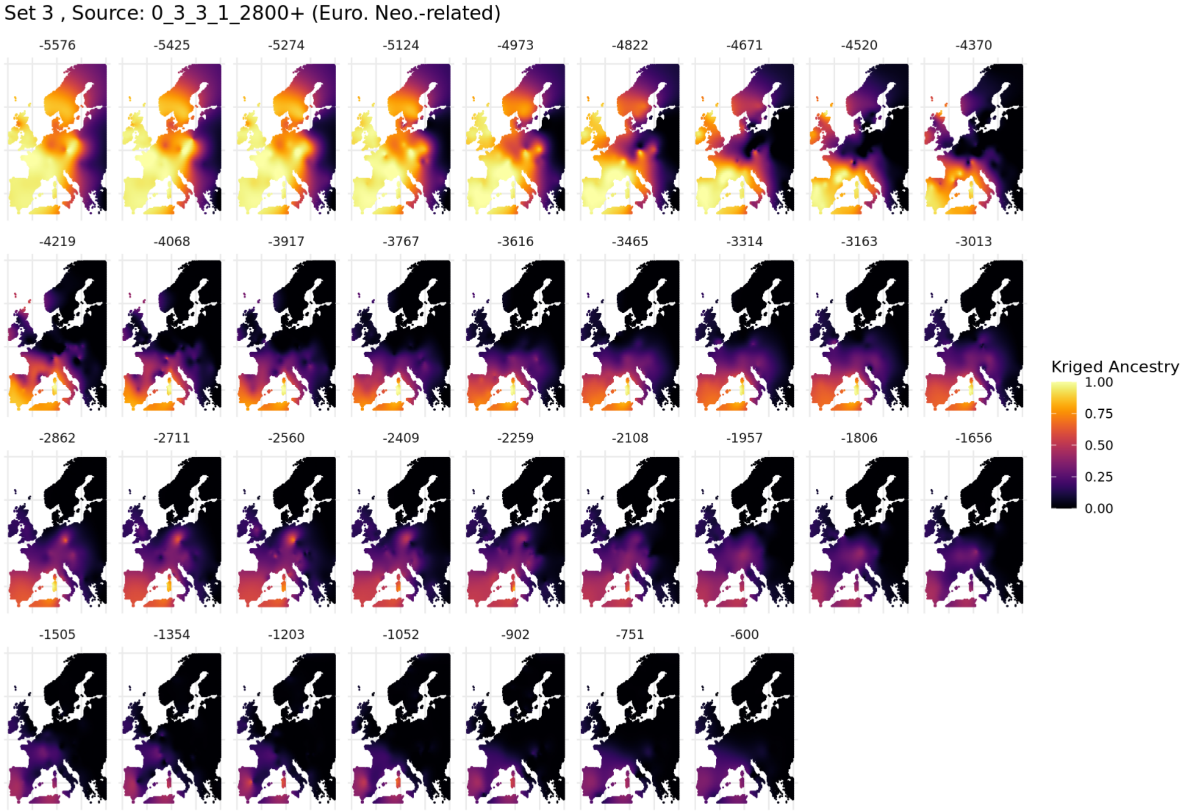

**Fig. S5.64. Spatiotemporal kriging results for Set 3 using 'Europe-Wide' parameters.**

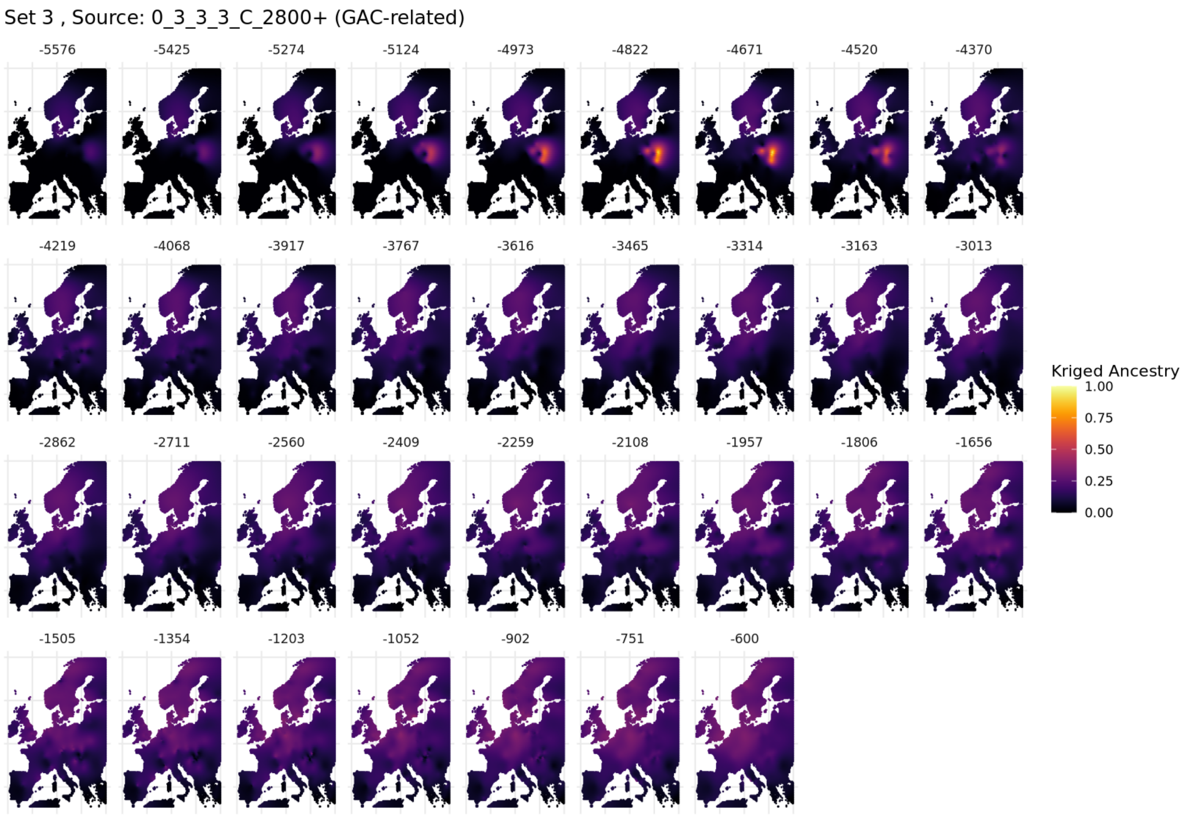

**Fig. S5.65. Spatiotemporal kriging results for Set 3 using 'Europe-Wide' parameters.**

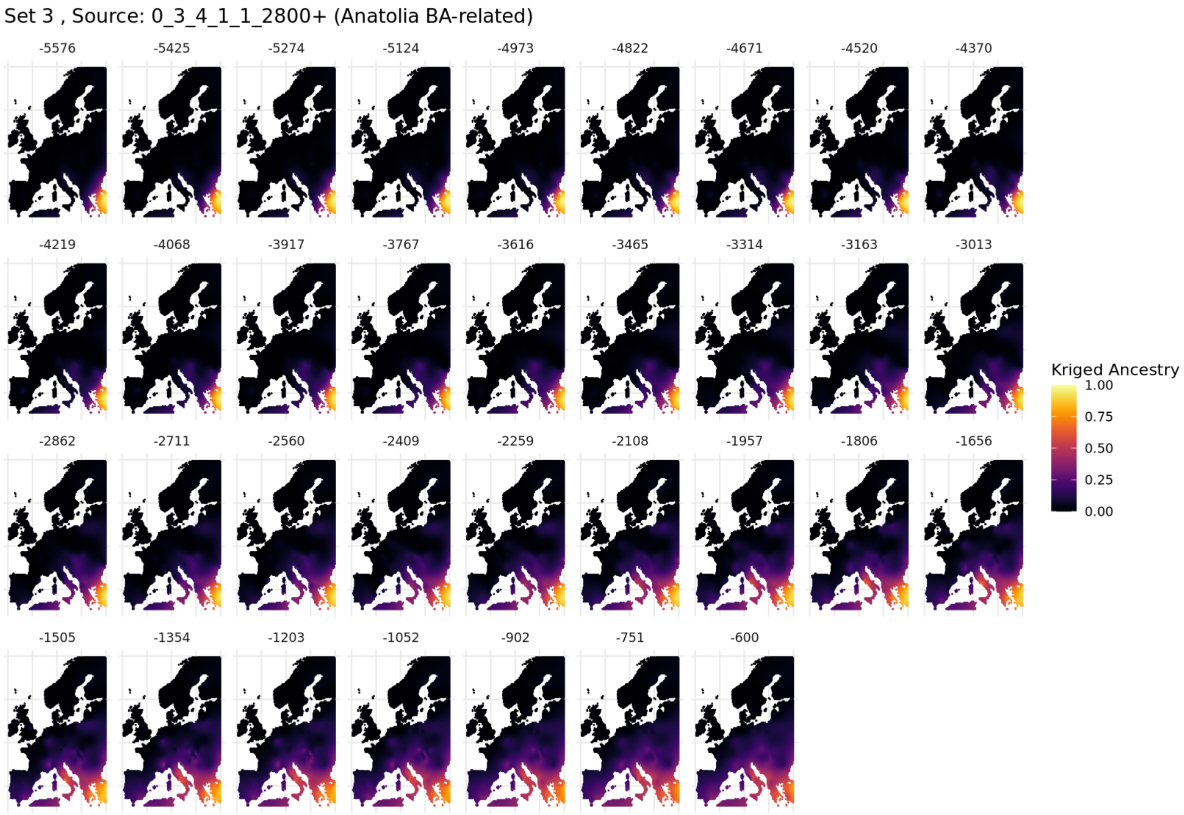

**Fig. S5.66. Spatiotemporal kriging results for Set 3 using 'Europe-Wide' parameters.**

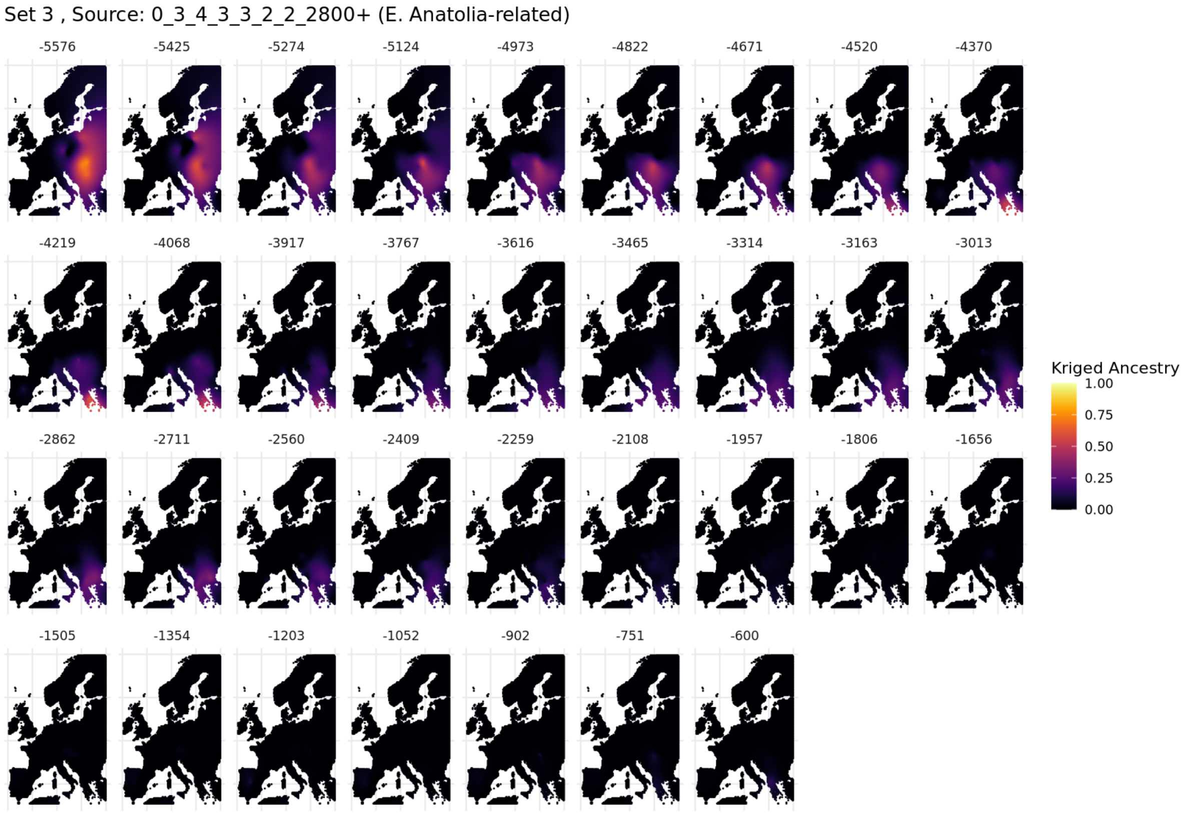

**Fig. S5.67. Spatiotemporal kriging results for Set 3 using 'Europe-Wide' parameters.**

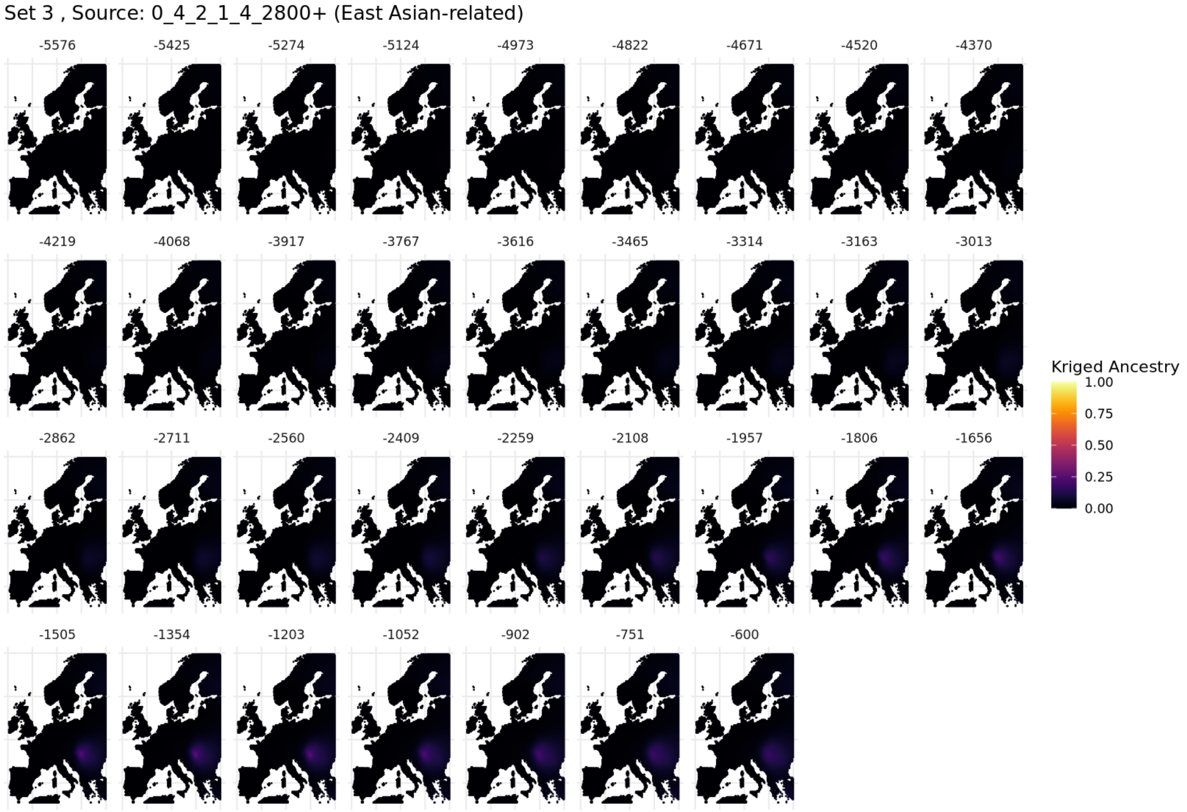

**Fig. S5.68. Spatiotemporal kriging results for Set 3 using 'Europe-Wide' parameters.**

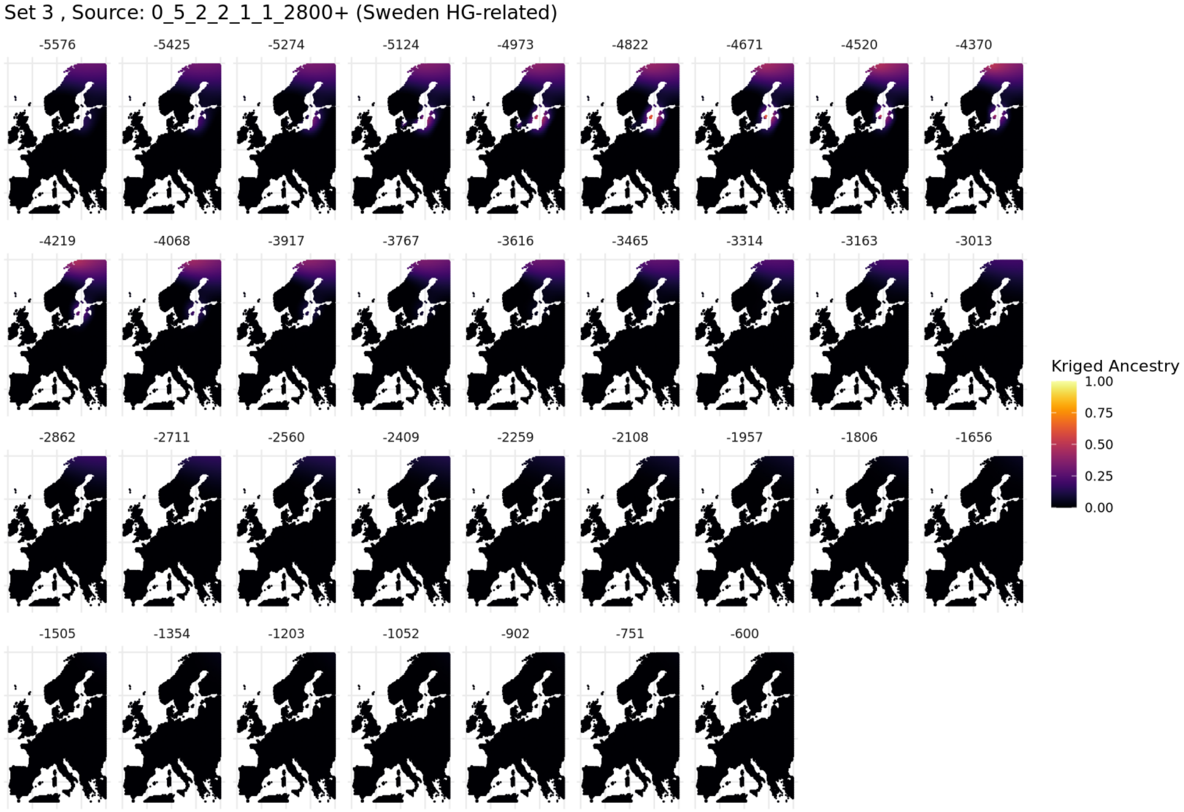

**Fig. S5.69. Spatiotemporal kriging results for Set 3 using 'Europe-Wide' parameters.**

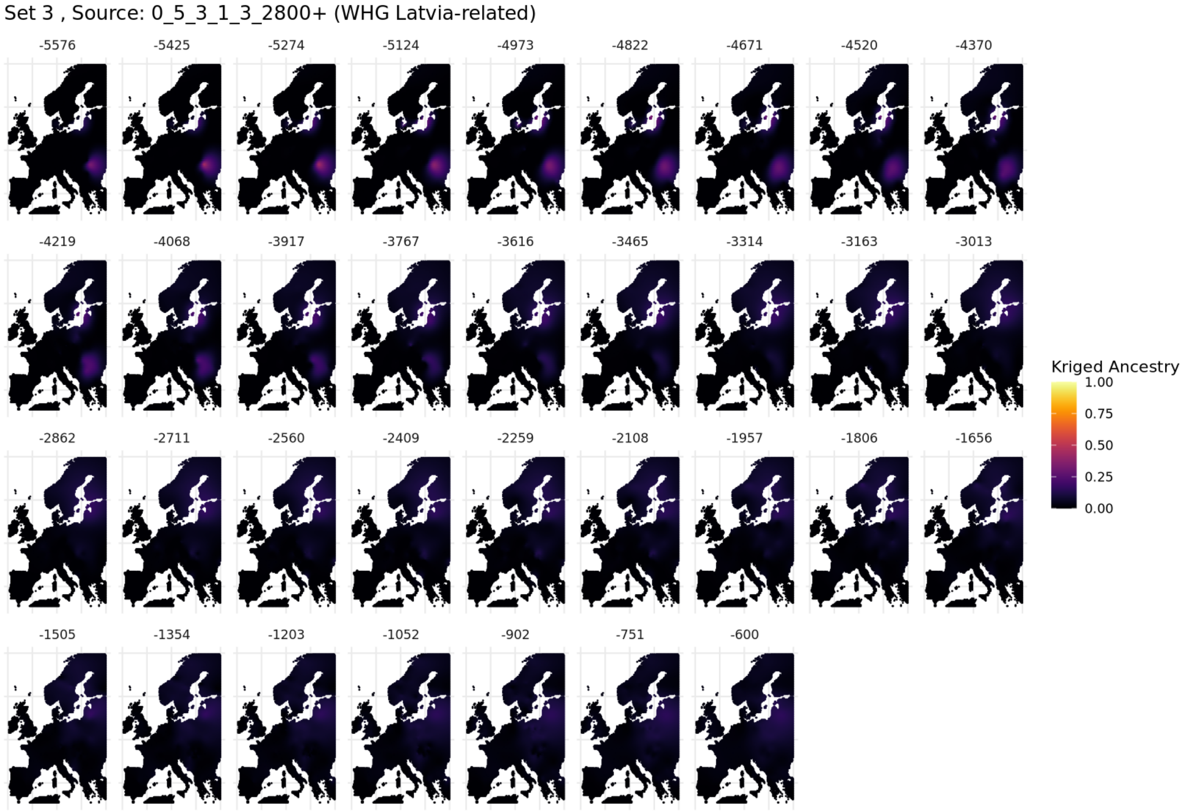

**Fig. S5.70. Spatiotemporal kriging results for Set 3 using 'Europe-Wide' parameters.**

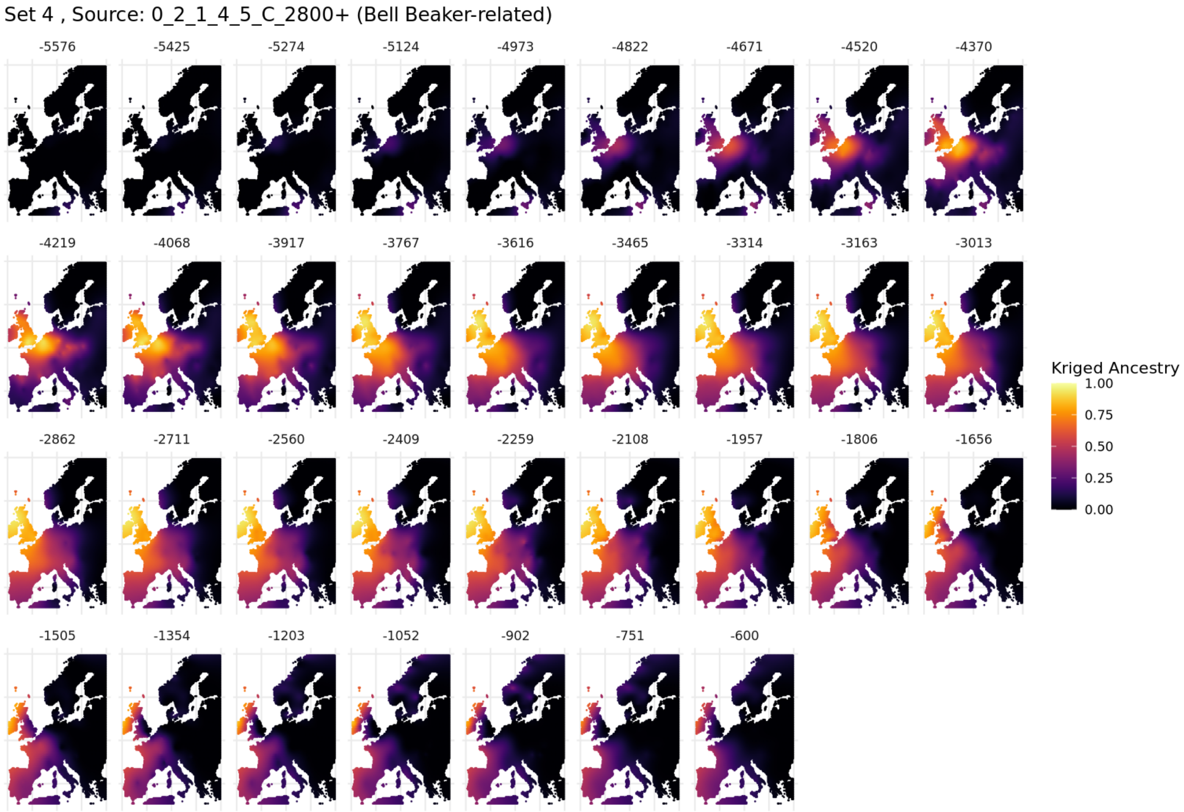

**Fig. S5.71. Spatiotemporal kriging results for Set 4 using 'Europe-Wide' parameters.**

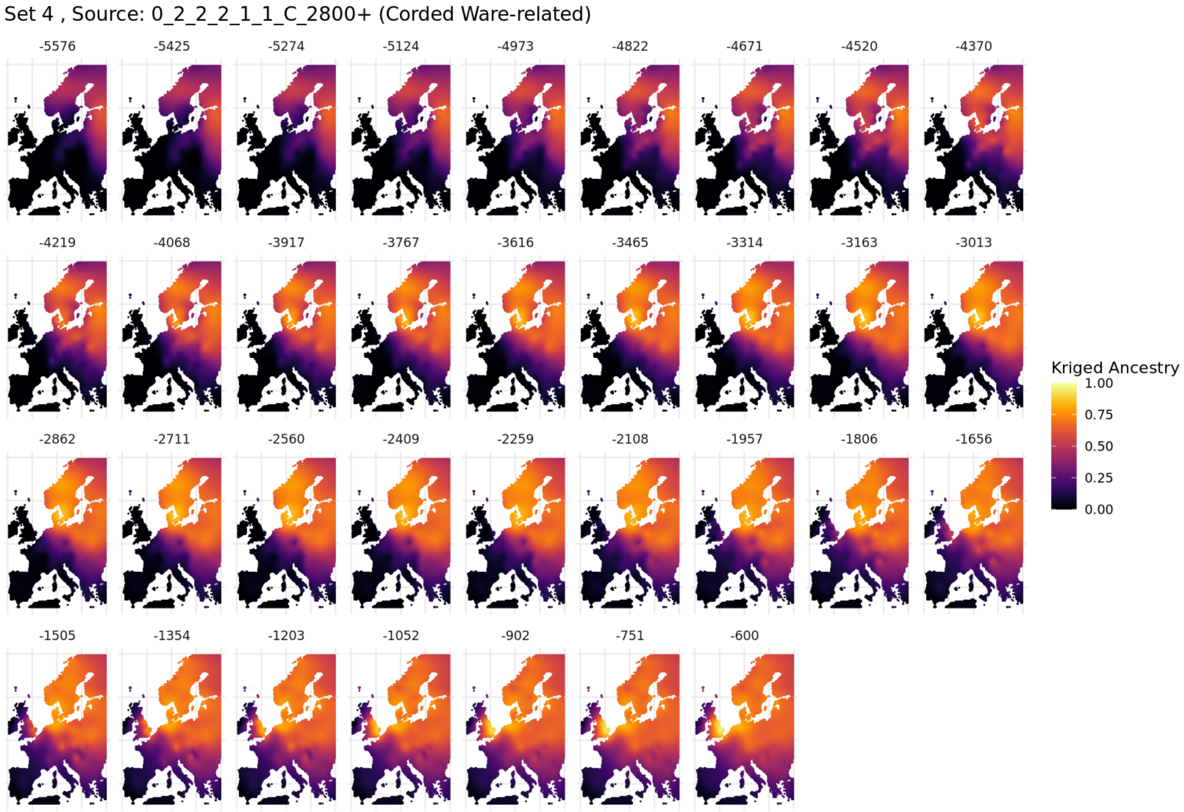

**Fig. S5.72. Spatiotemporal kriging results for Set 4 using 'Europe-Wide' parameters.**

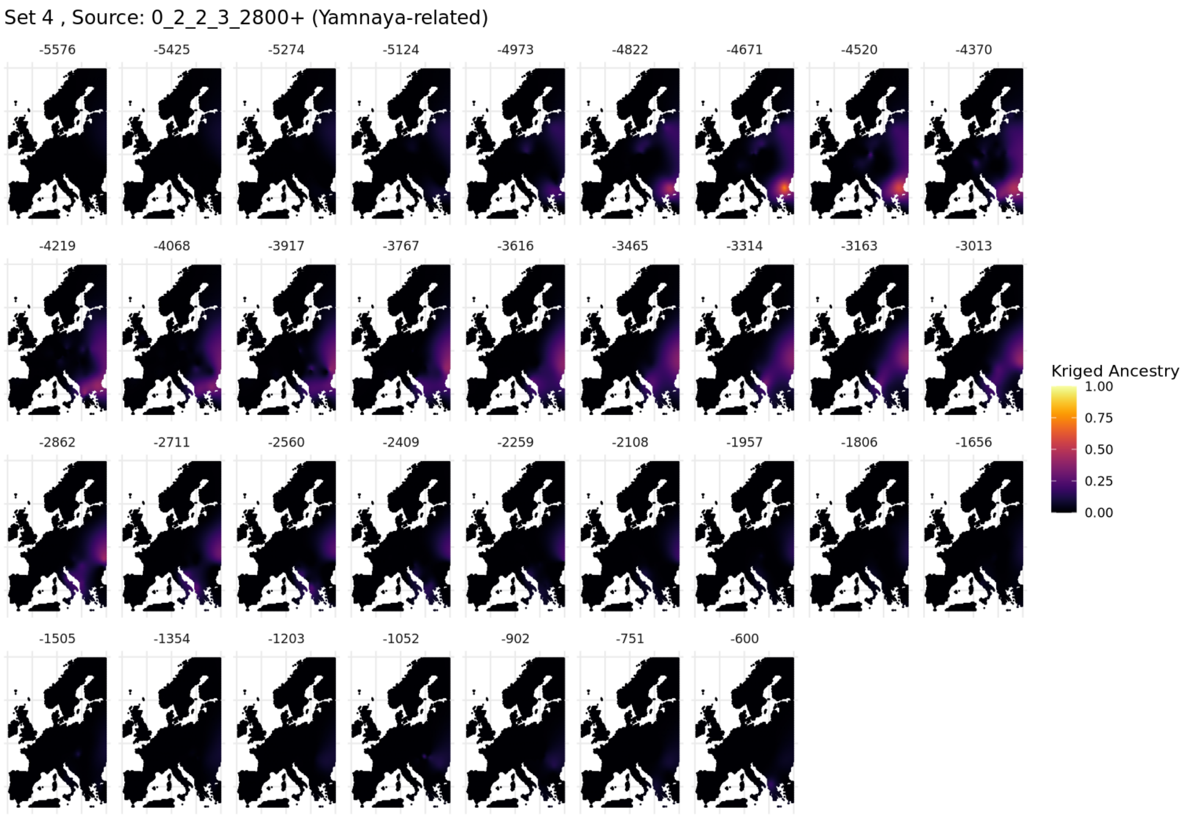

**Fig. S5.73. Spatiotemporal kriging results for Set 4 using ‘Europe-Wide’ parameters.**

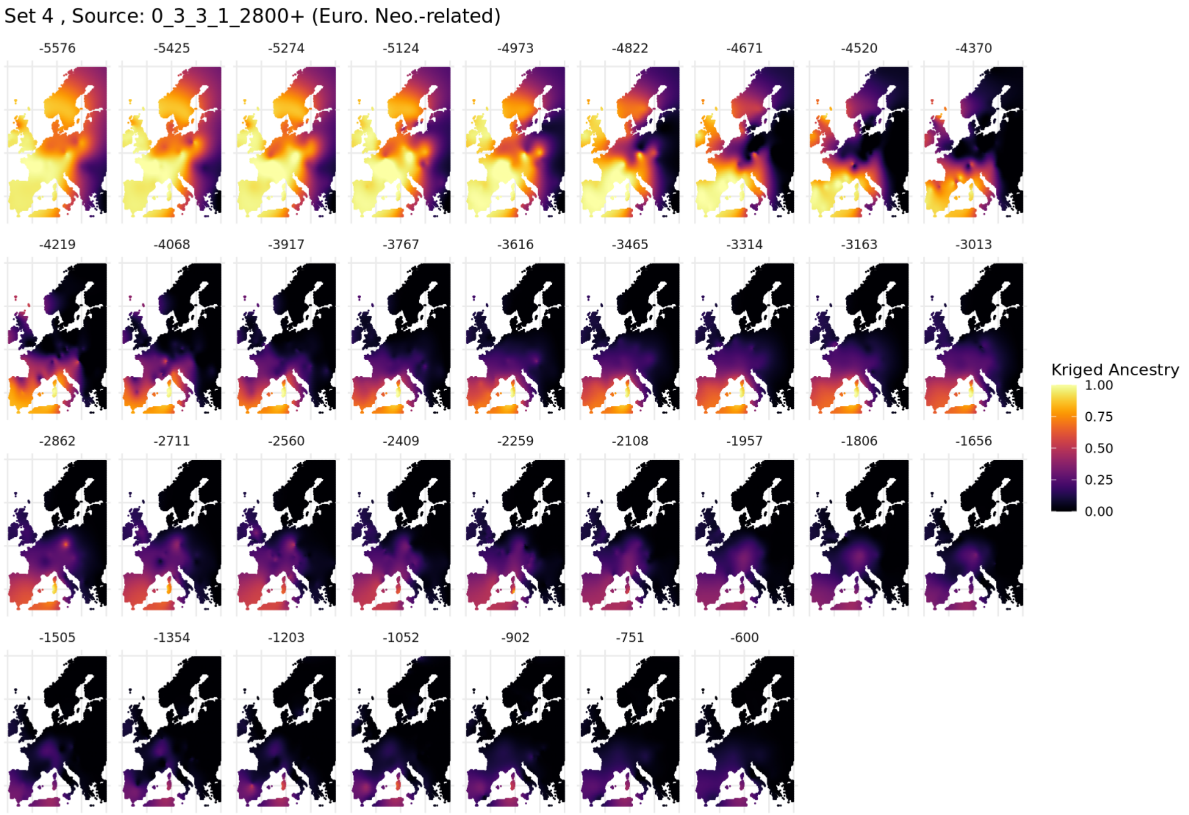

**Fig. S5.74. Spatiotemporal kriging results for Set 4 using ‘Europe-Wide’ parameters.**

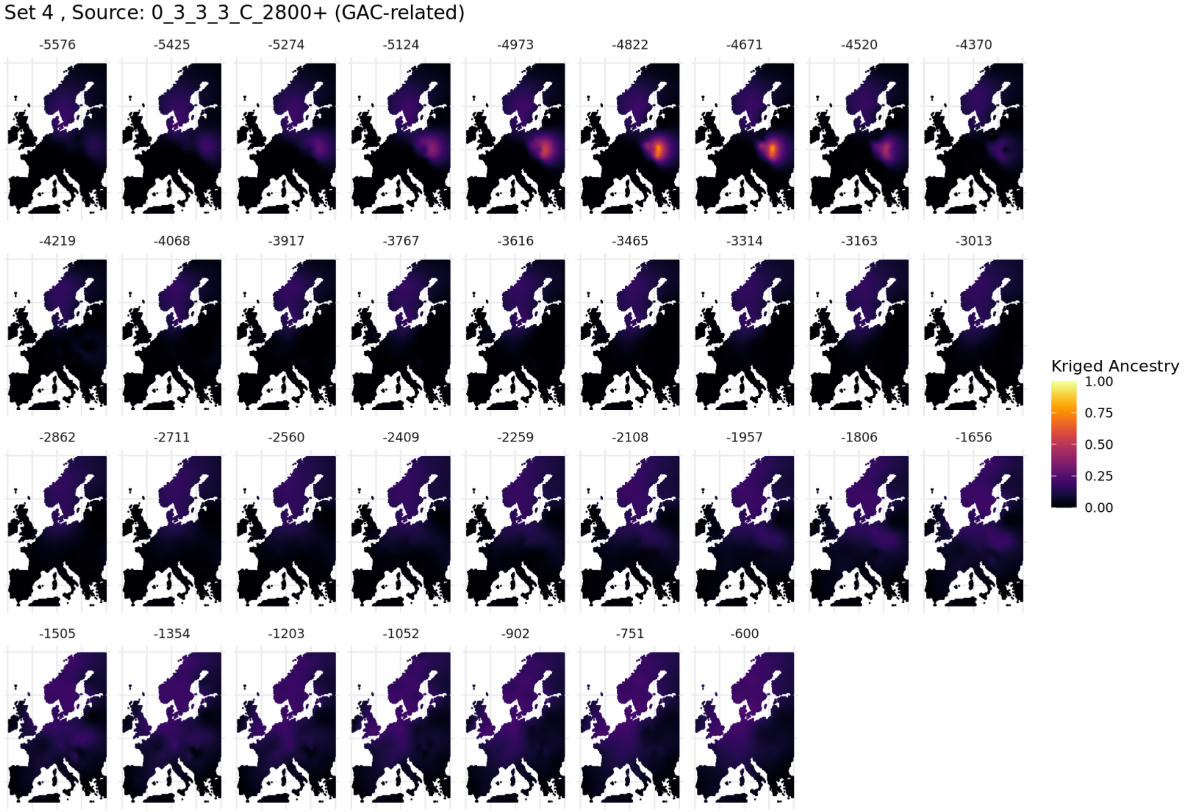

**Fig. S5.75. Spatiotemporal kriging results for Set 4 using 'Europe-Wide' parameters.**

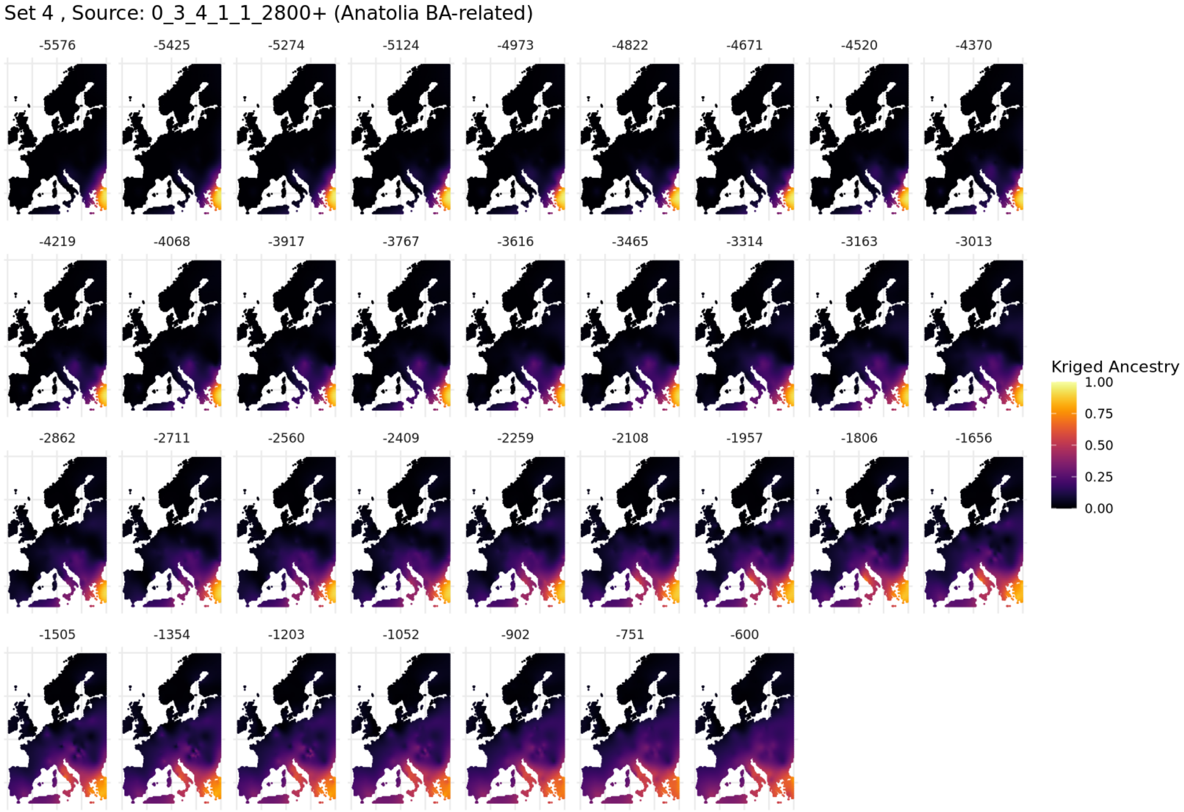

**Fig. S5.76. Spatiotemporal kriging results for Set 4 using 'Europe-Wide' parameters.**

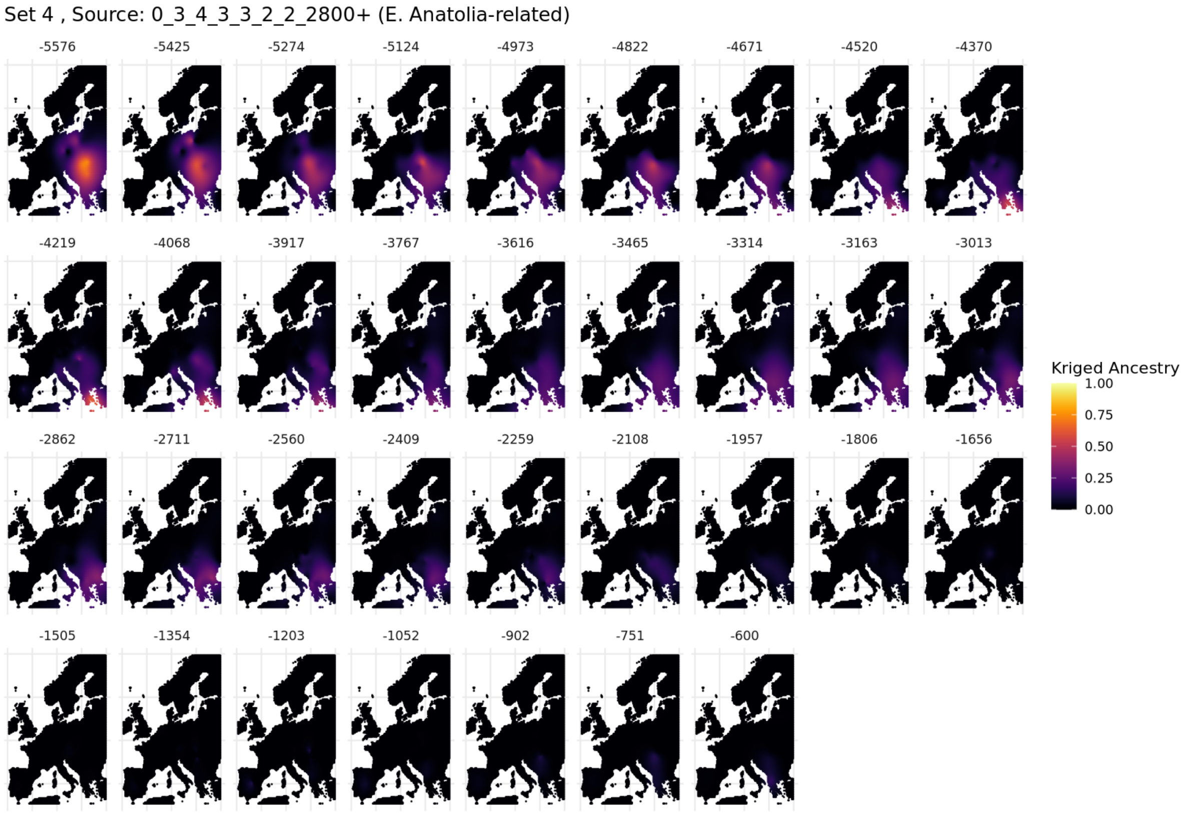

**Fig. S5.77. Spatiotemporal kriging results for Set 4 using 'Europe-Wide' parameters.**

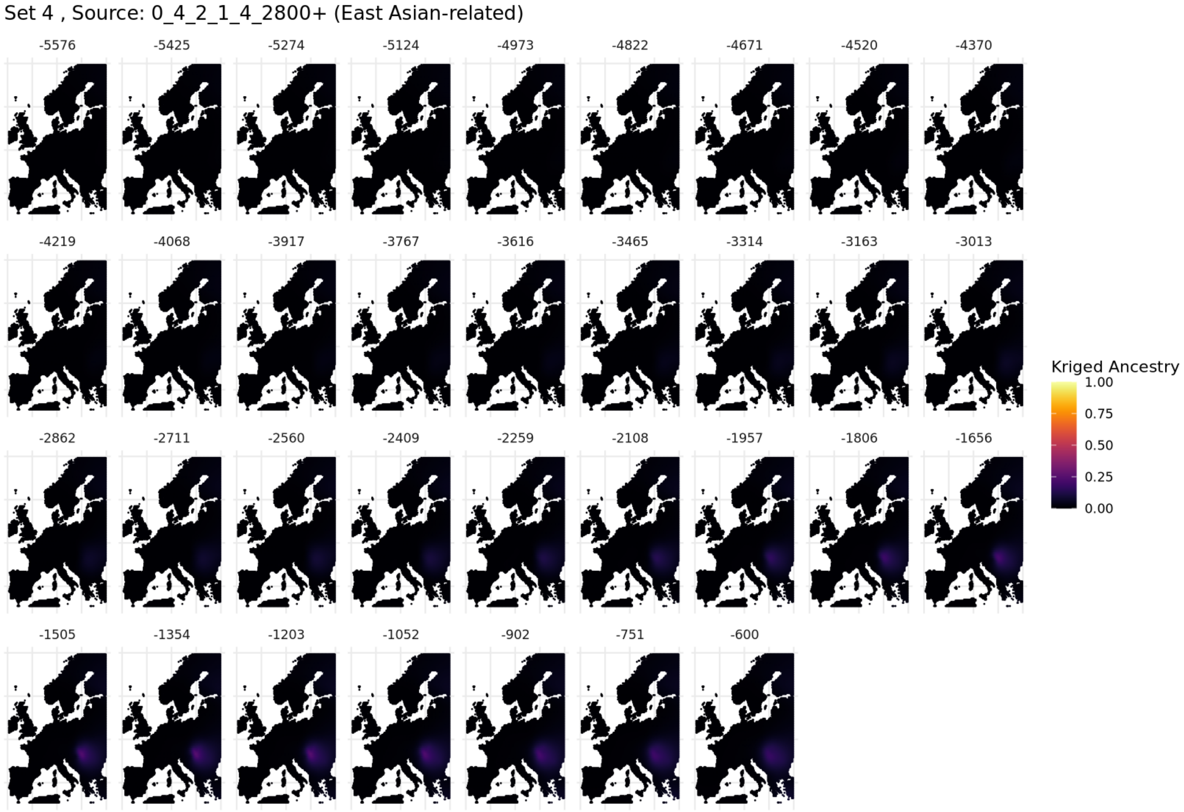

**Fig. S5.78. Spatiotemporal kriging results for Set 4 using 'Europe-Wide' parameters.**

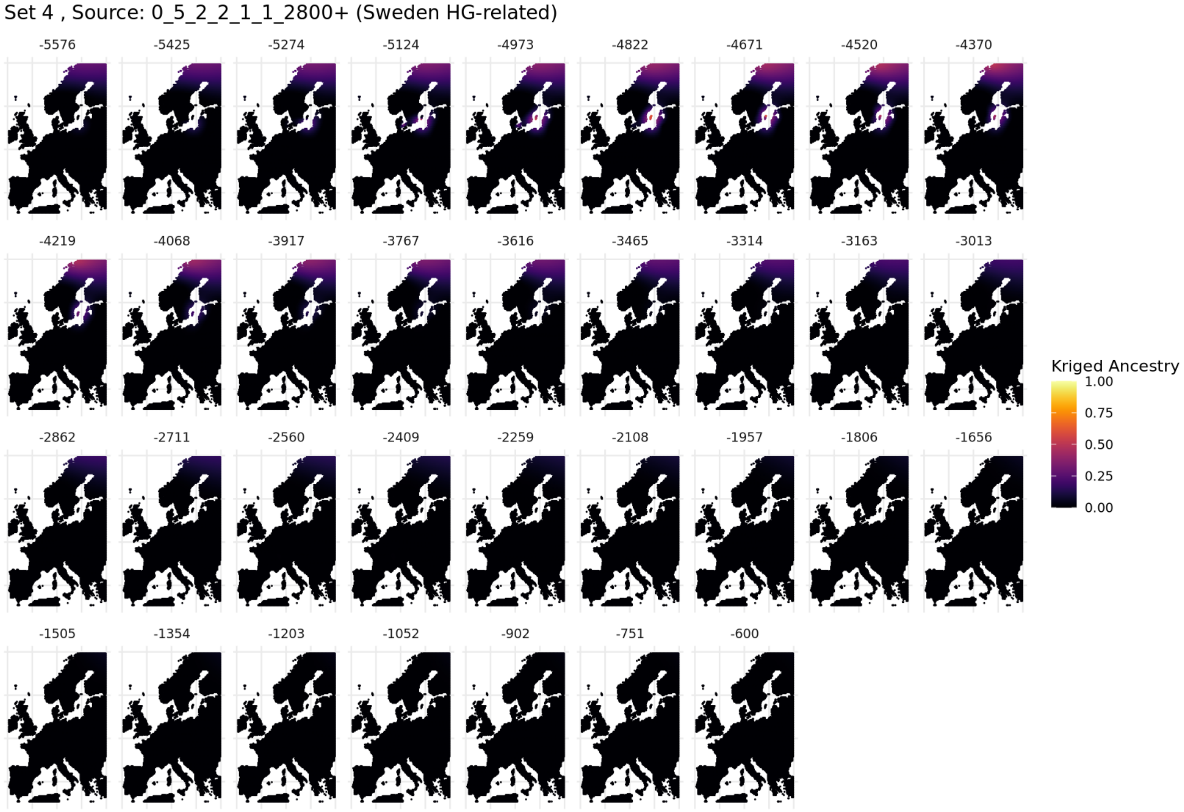

**Fig. S5.79. Spatiotemporal kriging results for Set 4 using 'Europe-Wide' parameters.**

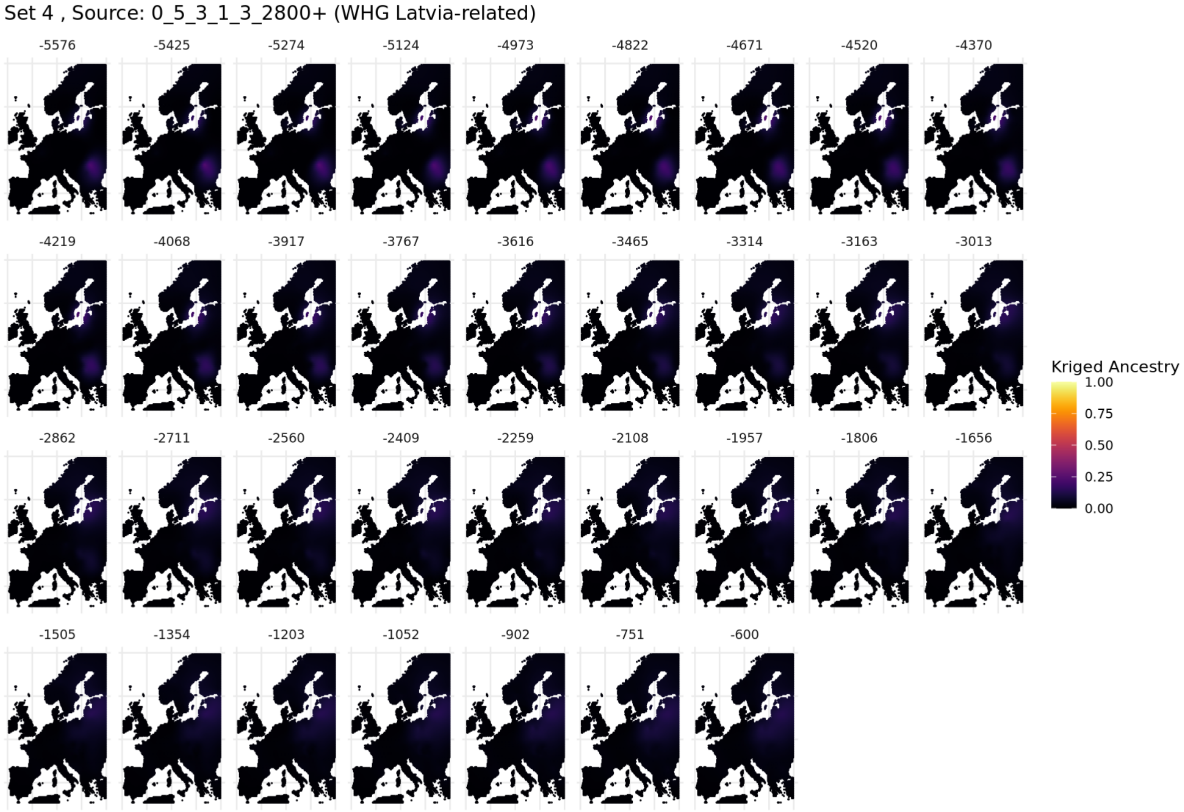

**Fig. S5.80. Spatiotemporal kriging results for Set 4 using 'Europe-Wide' parameters.**

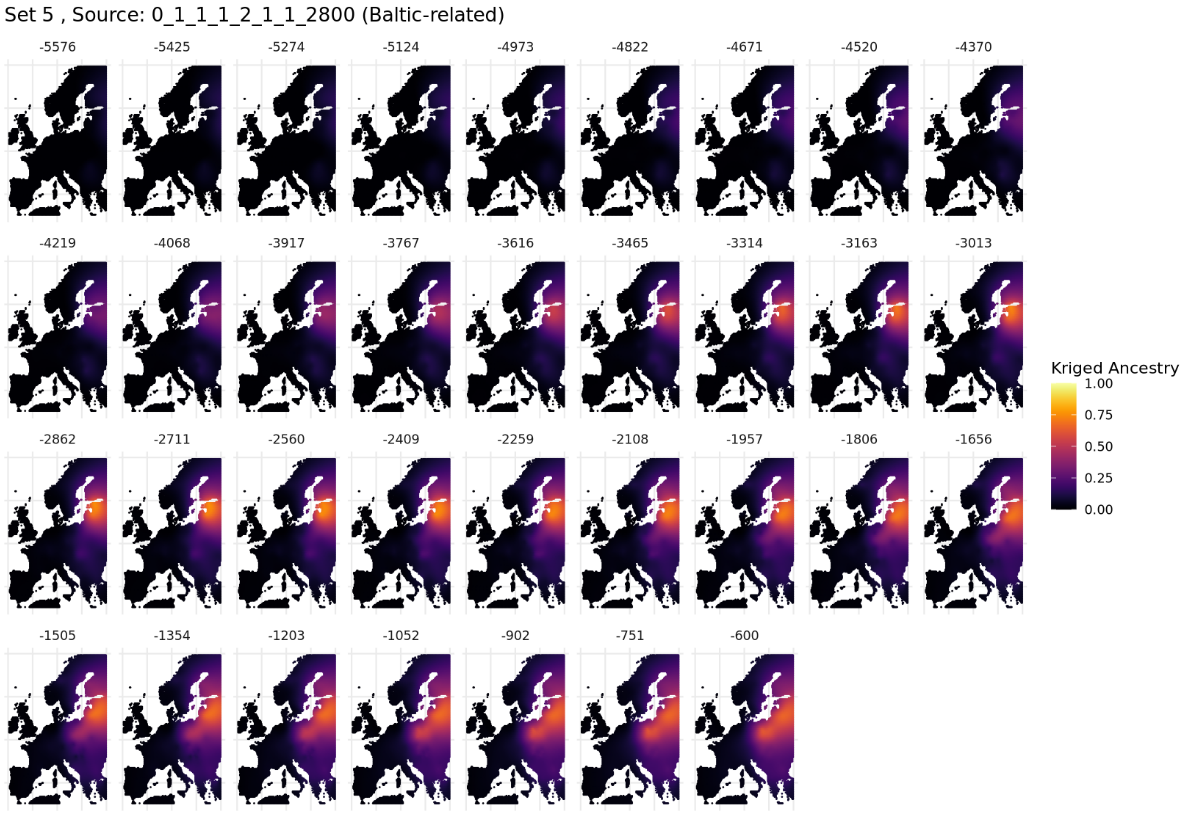

**Fig. S5.81. Spatiotemporal kriging results for Set 5 using 'Europe-Wide' parameters.**

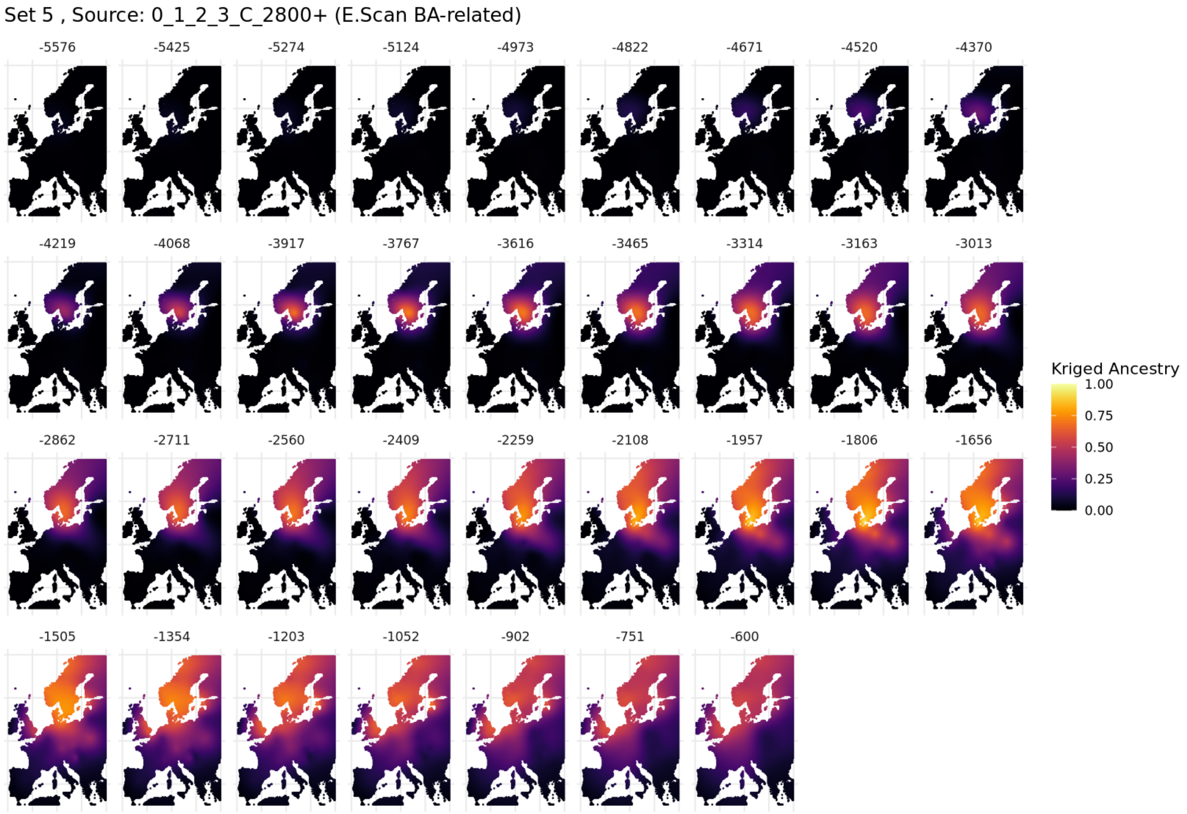

**Fig. S5.82. Spatiotemporal kriging results for Set 5 using 'Europe-Wide' parameters.**

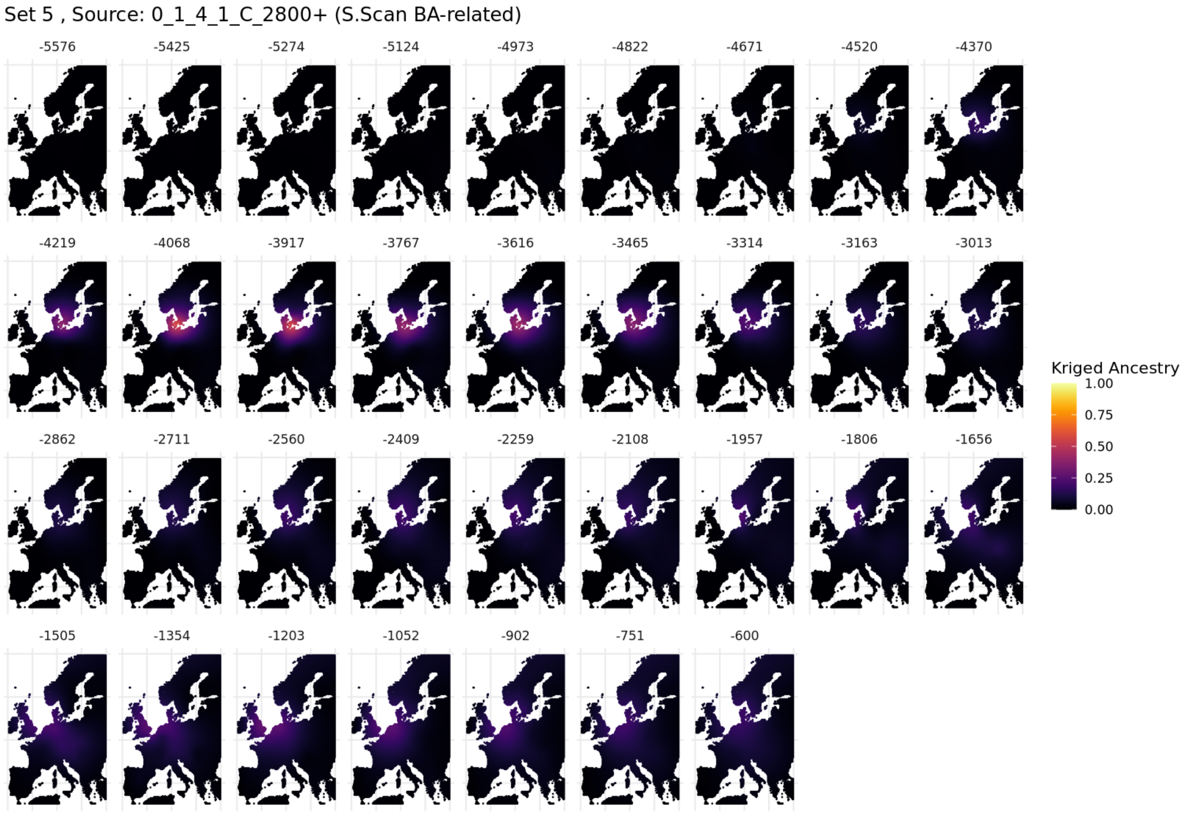

**Fig. S5.83. Spatiotemporal kriging results for Set 5 using 'Europe-Wide' parameters.**

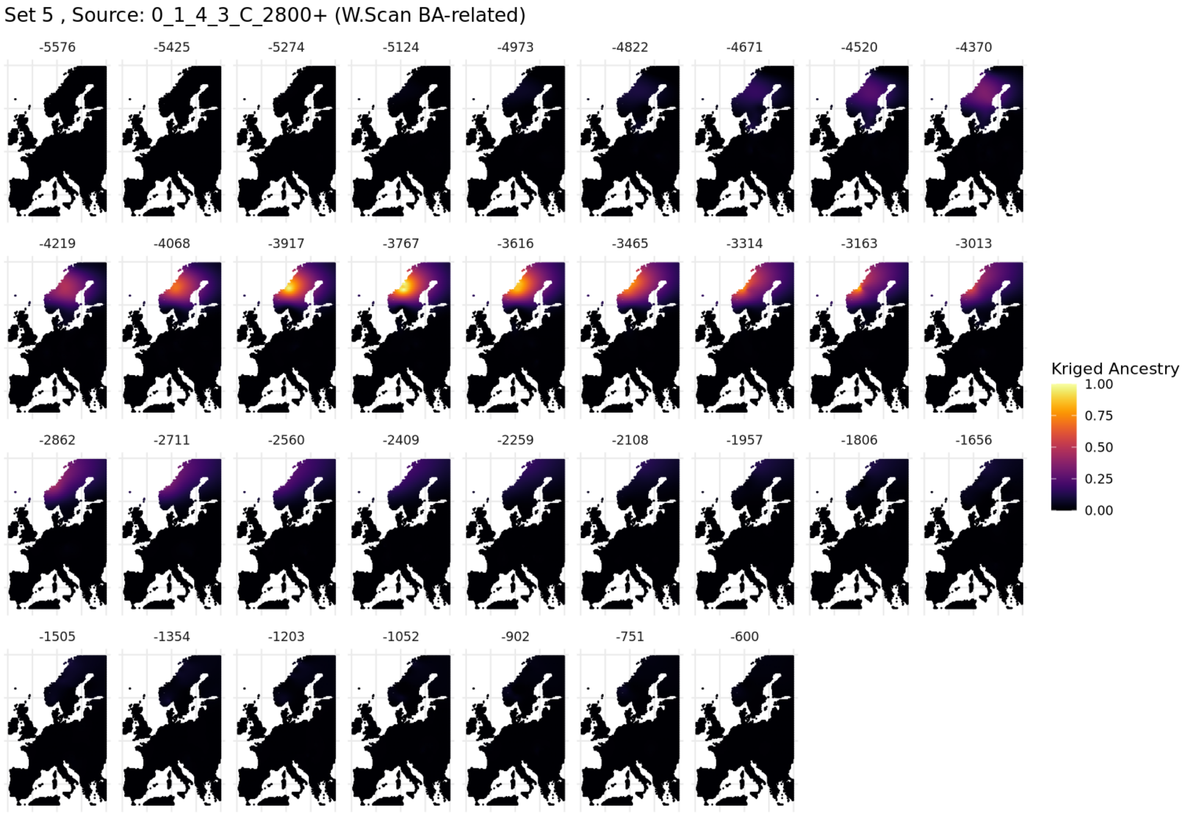

**Fig. S5.84. Spatiotemporal kriging results for Set 5 using 'Europe-Wide' parameters.**

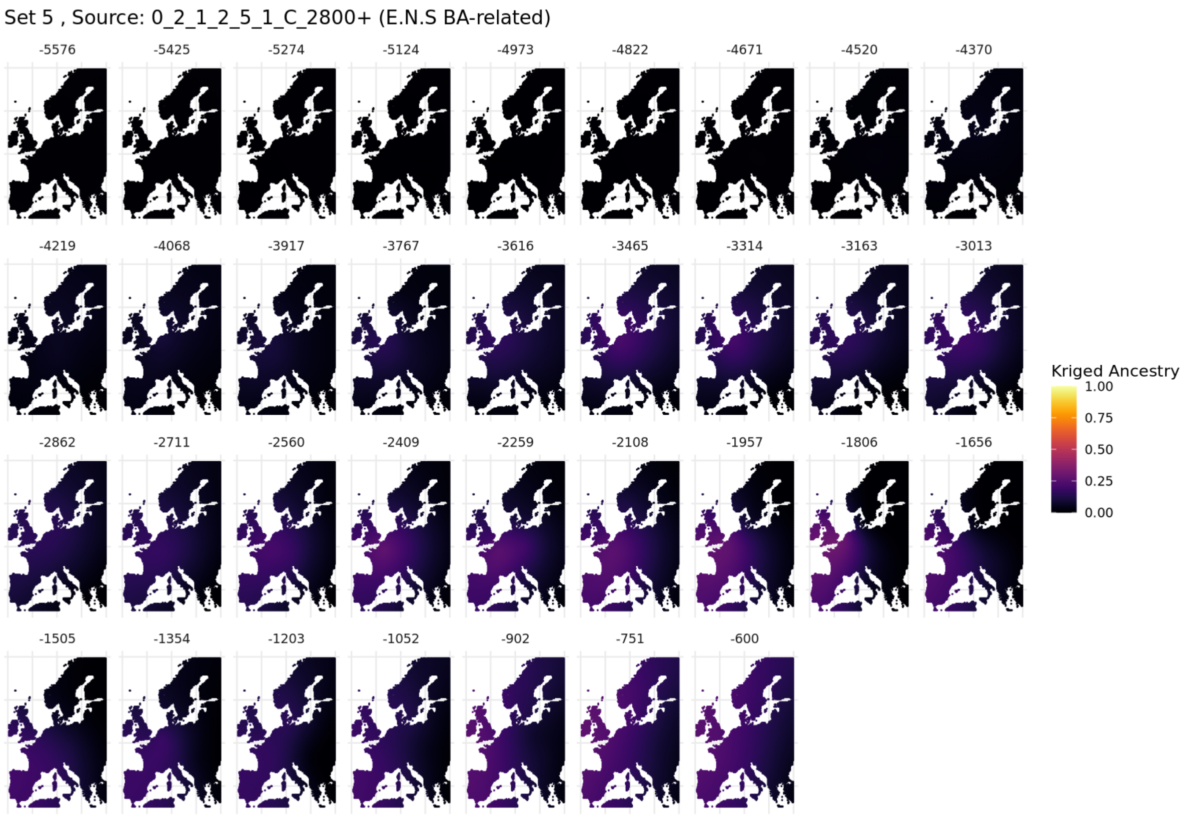

**Fig. S5.85. Spatiotemporal kriging results for Set 5 using 'Europe-Wide' parameters.**

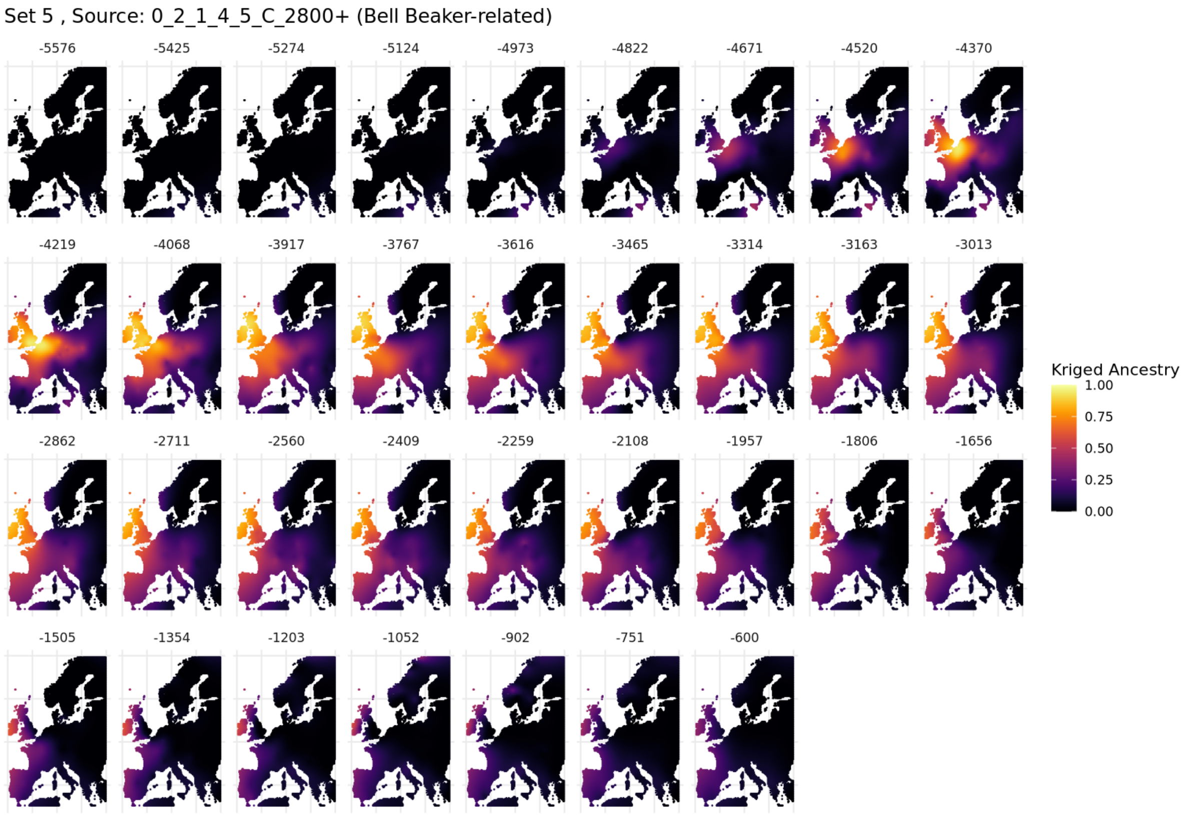

**Fig. S5.86. Spatiotemporal kriging results for Set 5 using 'Europe-Wide' parameters.**

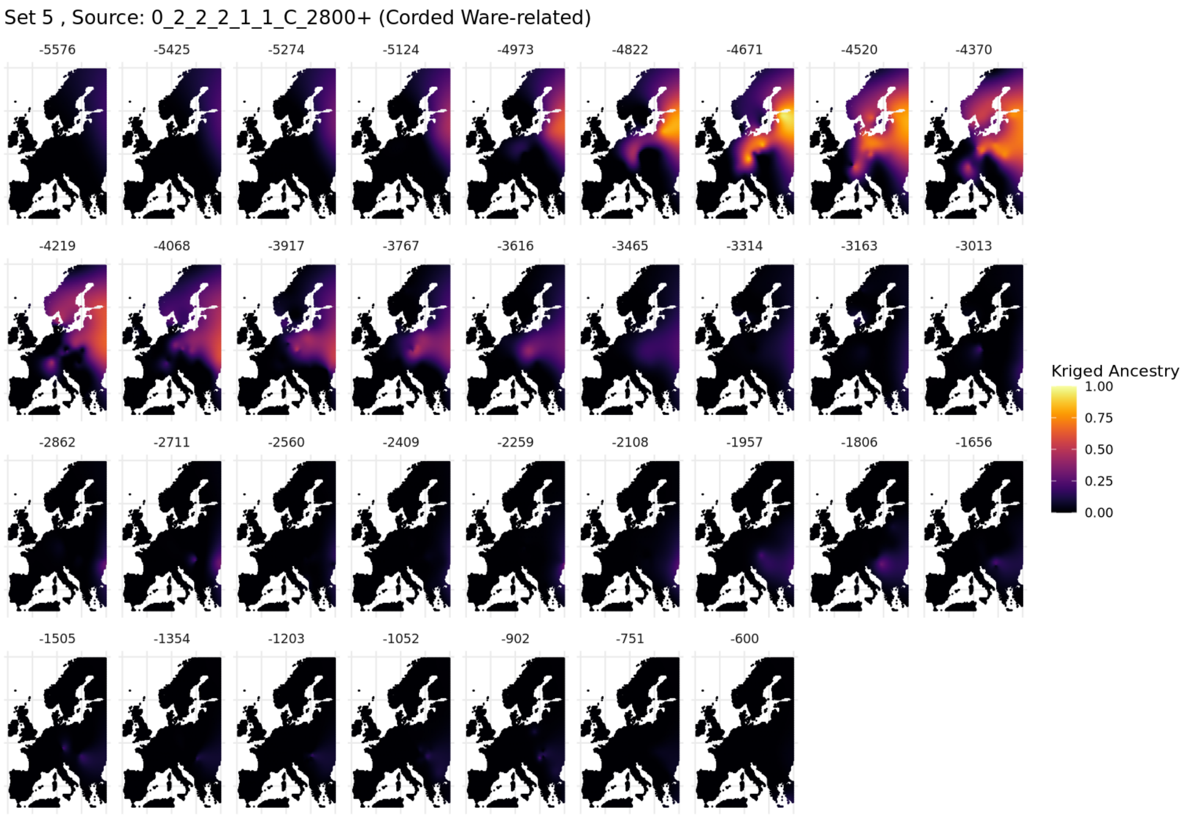

**Fig. S5.87. Spatiotemporal kriging results for Set 5 using 'Europe-Wide' parameters.**

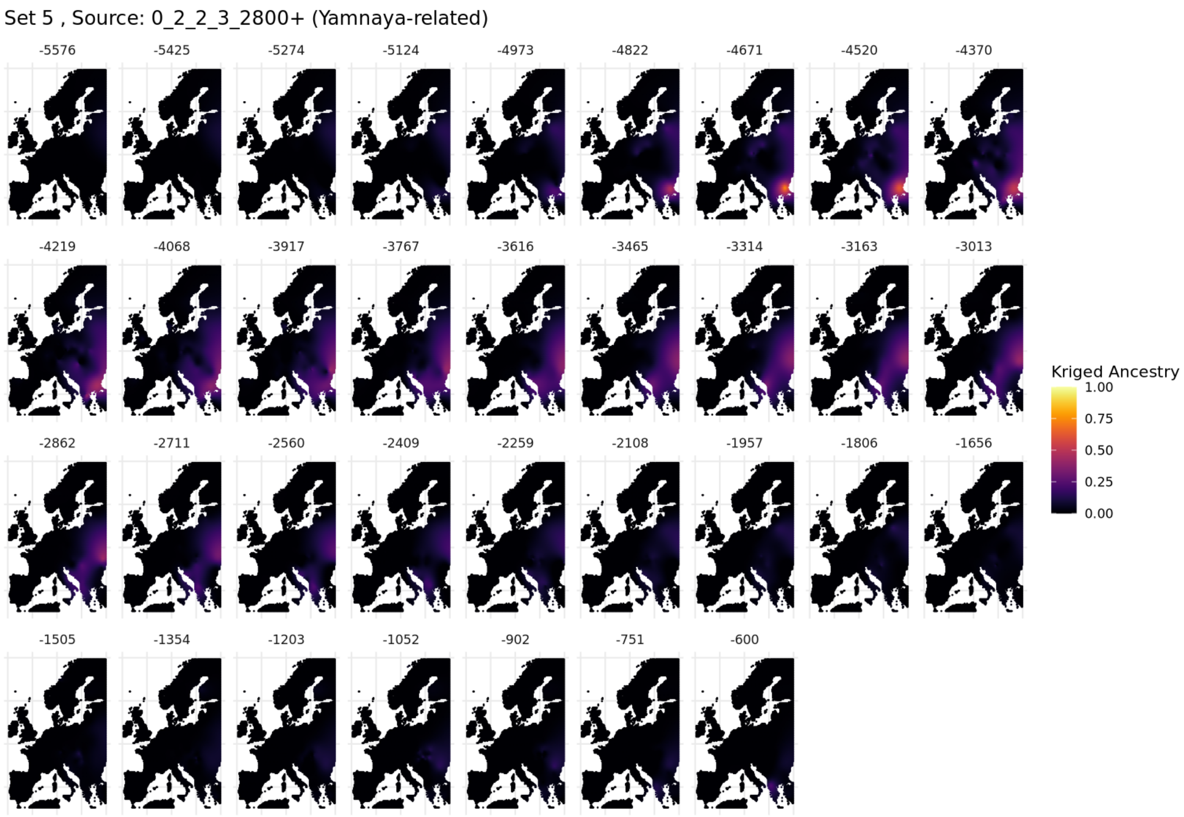

**Fig. S5.88. Spatiotemporal kriging results for Set 5 using 'Europe-Wide' parameters.**

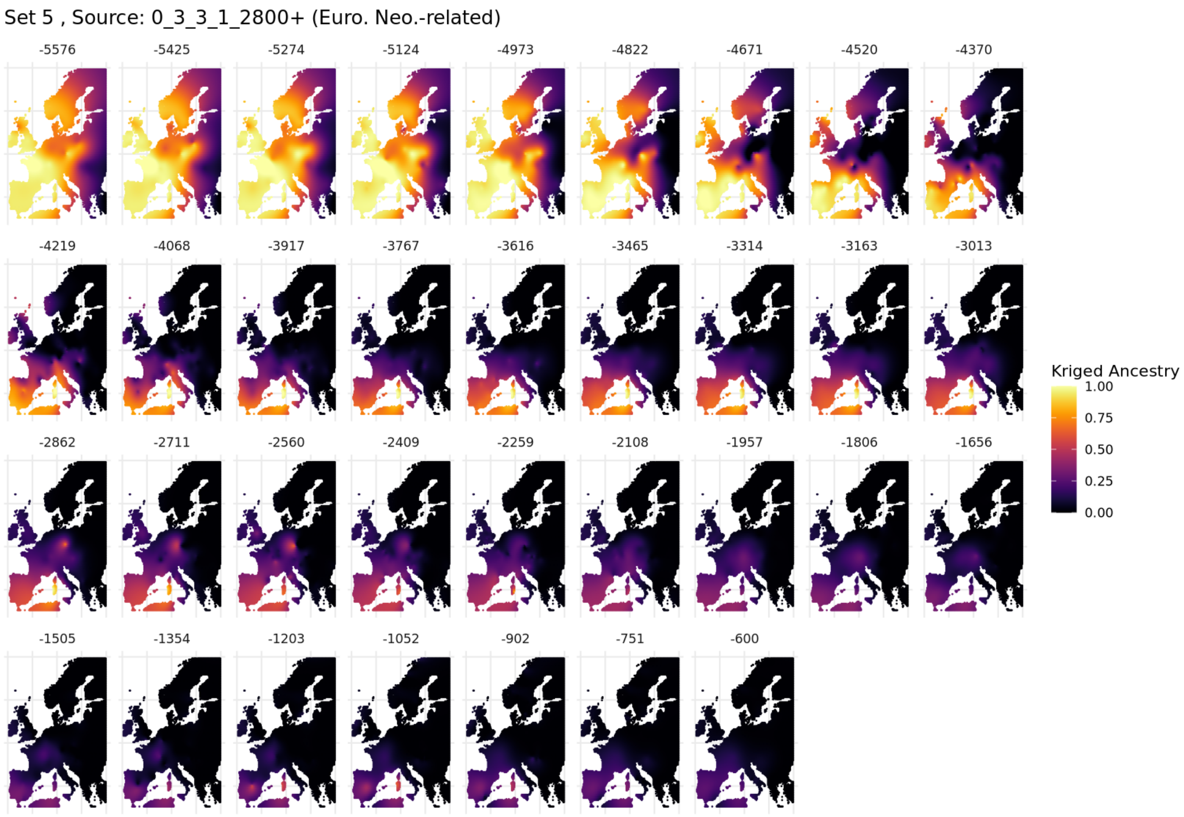

**Fig. S5.89. Spatiotemporal kriging results for Set 5 using 'Europe-Wide' parameters.**

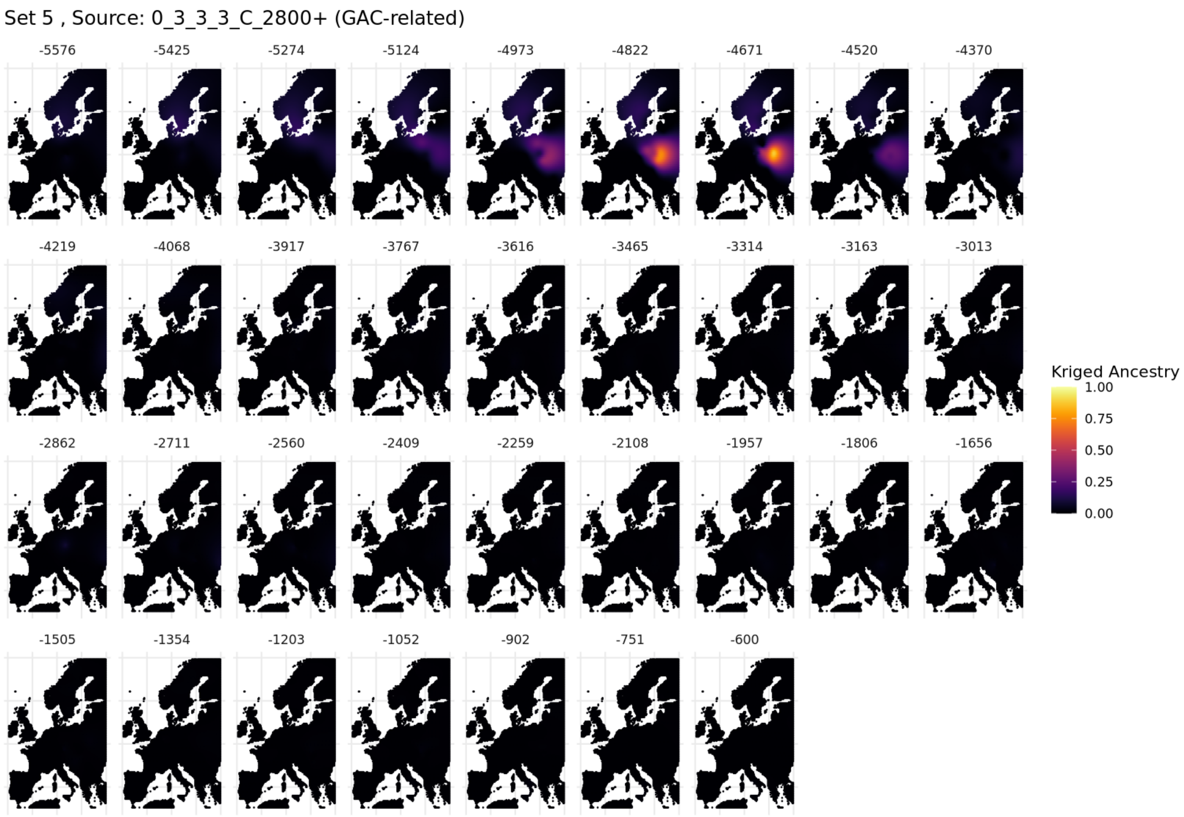

**Fig. S5.90. Spatiotemporal kriging results for Set 5 using 'Europe-Wide' parameters.**

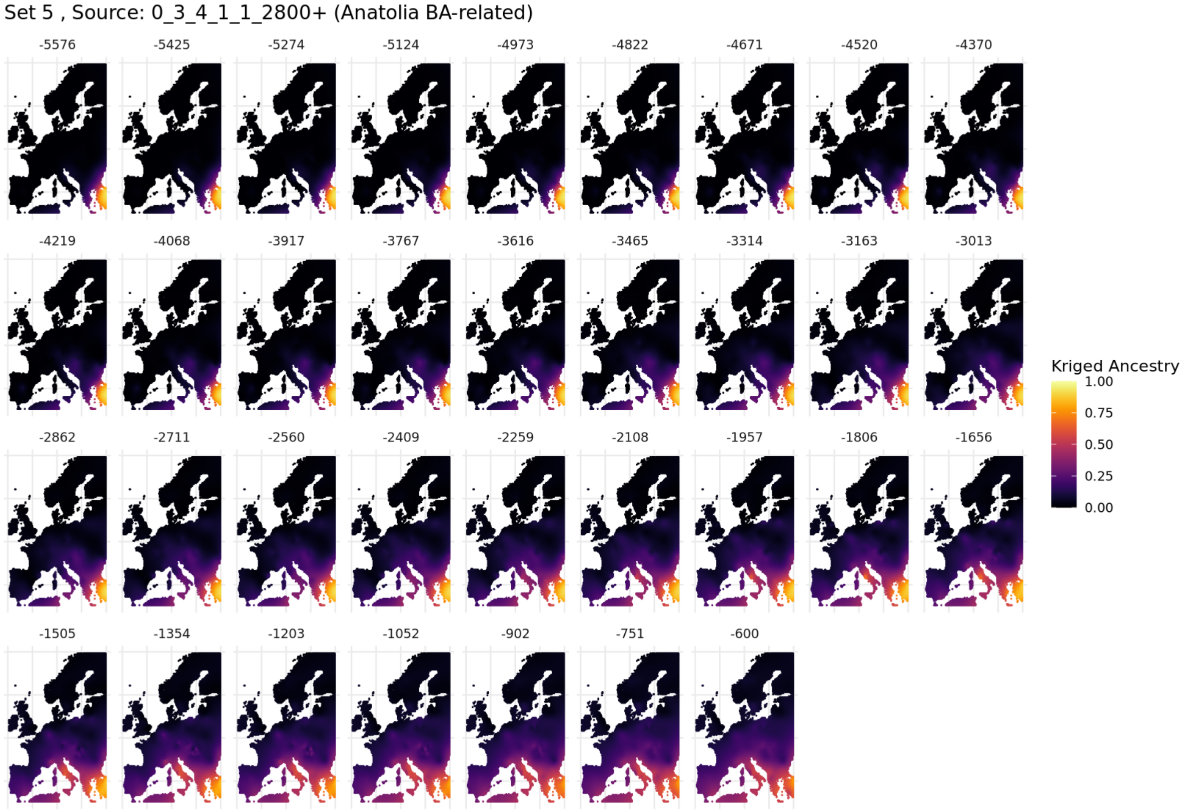

**Fig. S5.91. Spatiotemporal kriging results for Set 5 using 'Europe-Wide' parameters.**

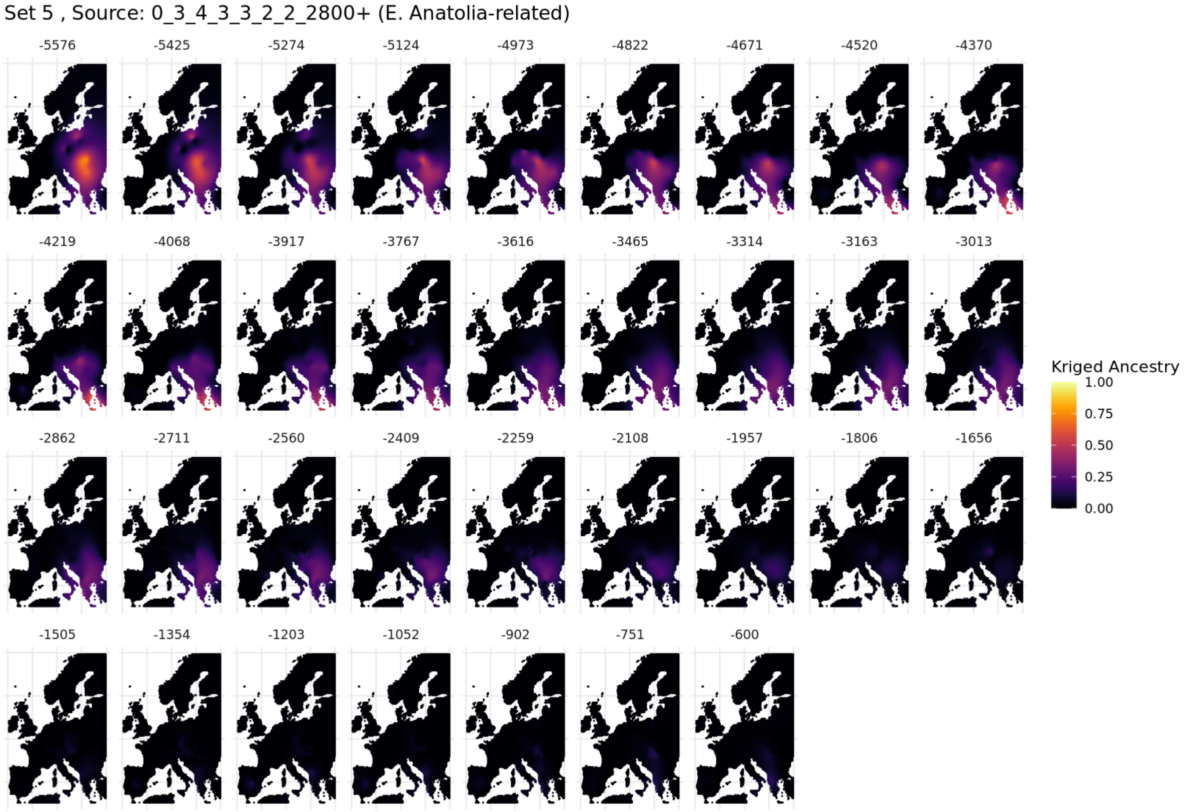

**Fig. S5.92. Spatiotemporal kriging results for Set 5 using 'Europe-Wide' parameters.**

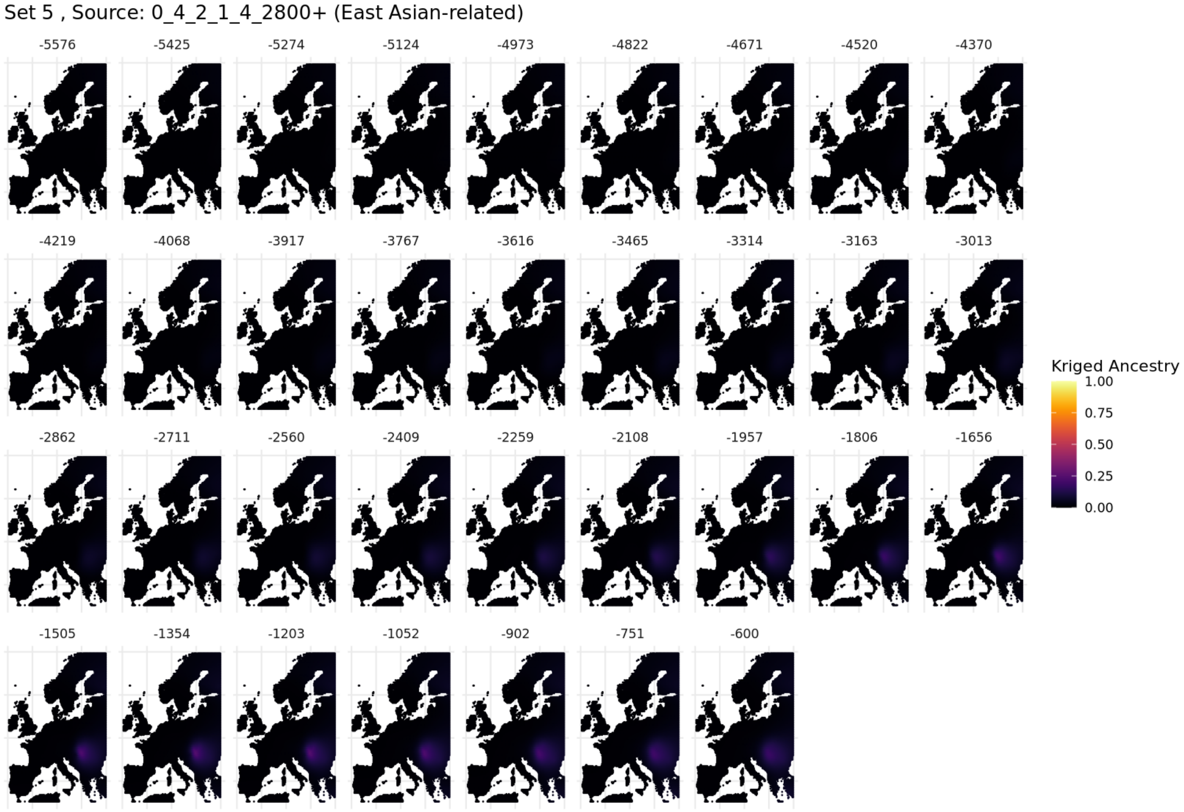

**Fig. S5.93. Spatiotemporal kriging results for Set 5 using 'Europe-Wide' parameters.**

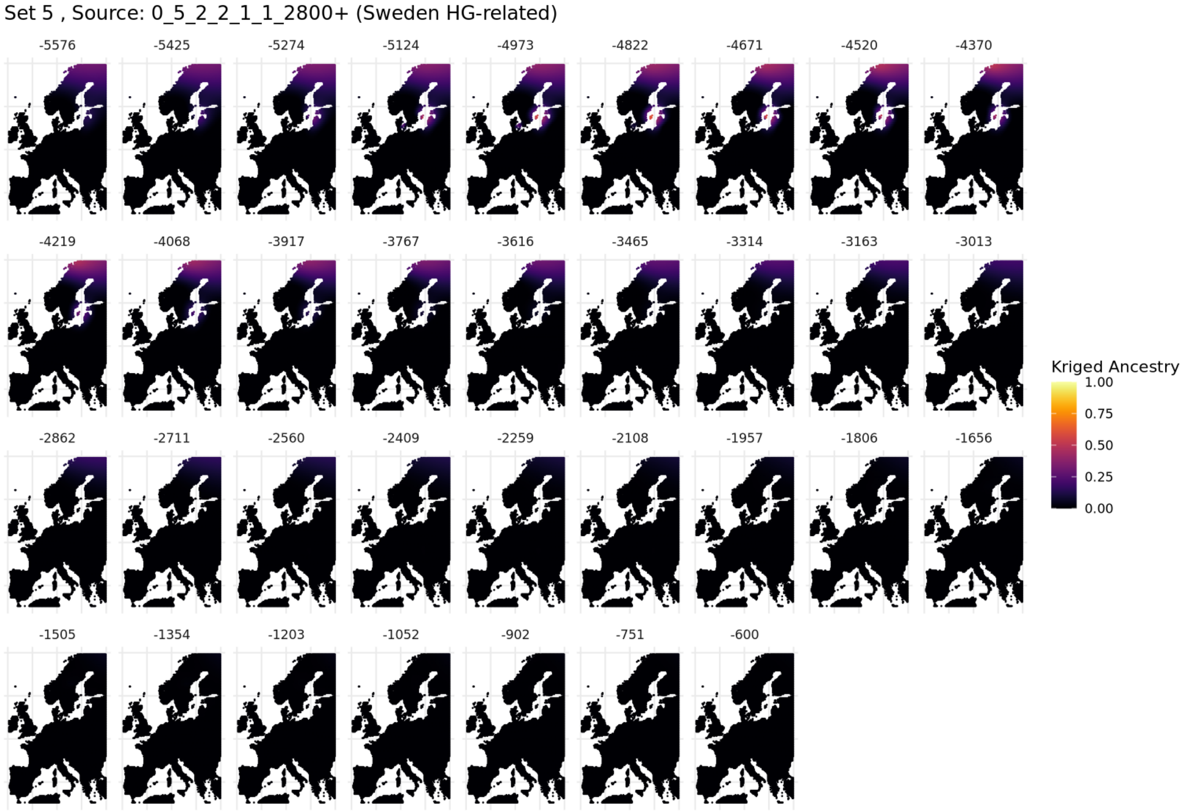

**Fig. S5.94. Spatiotemporal kriging results for Set 5 using 'Europe-Wide' parameters.**

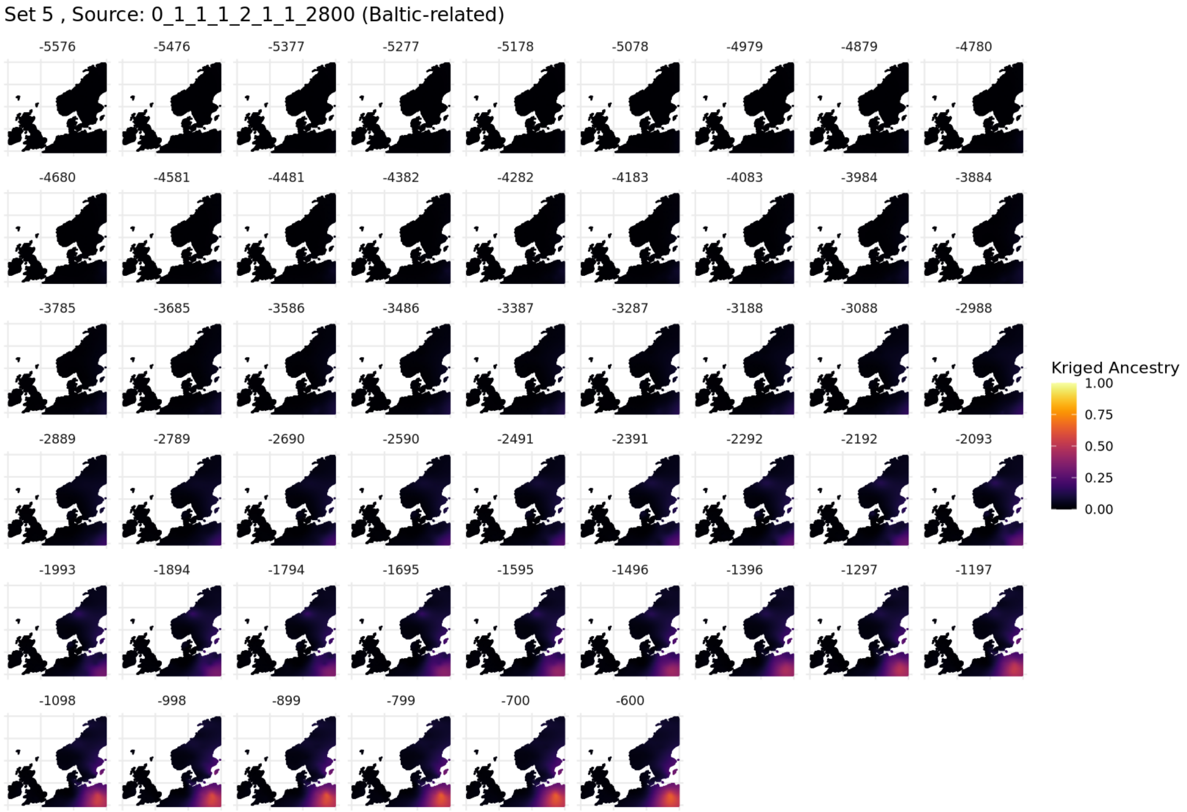

**Fig. S5.95. Spatiotemporal kriging results for Set 5, ‘North/West Europe’ parameters.**

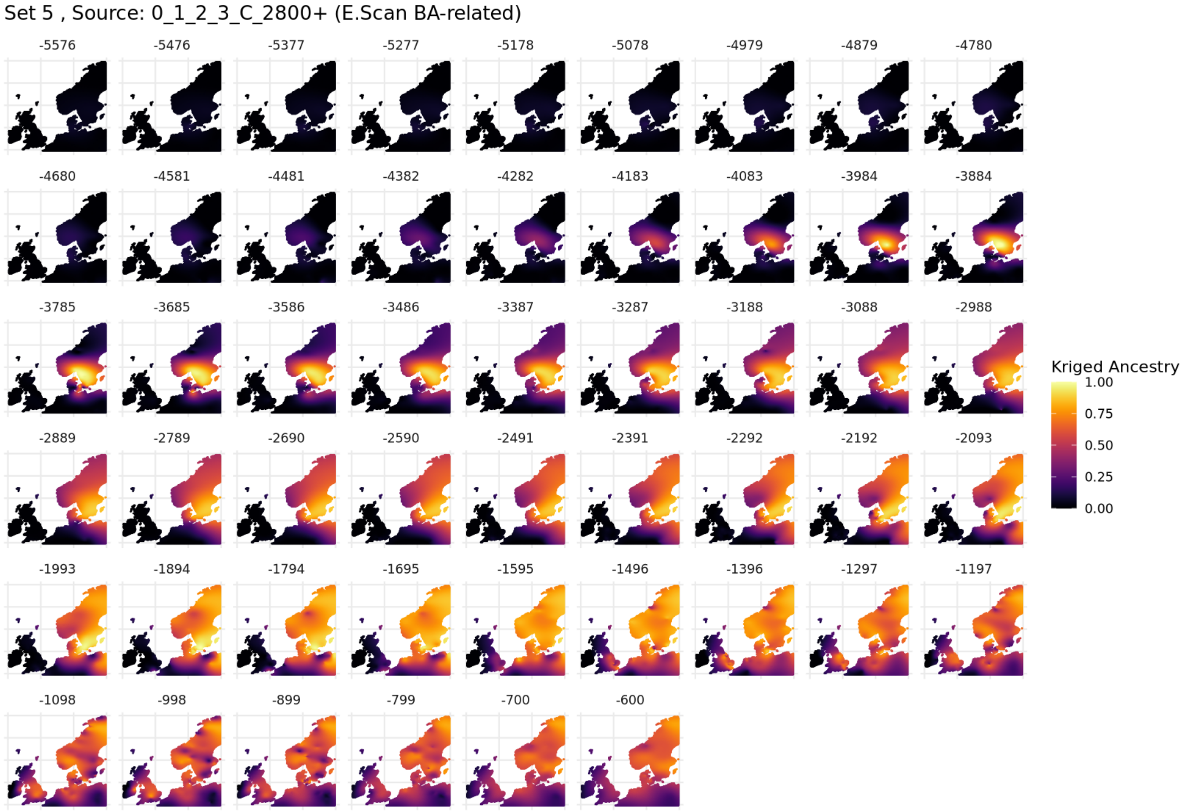

**Fig. S5.96. Spatiotemporal kriging results for Set 5, ‘North / West Europe’ parameters.**

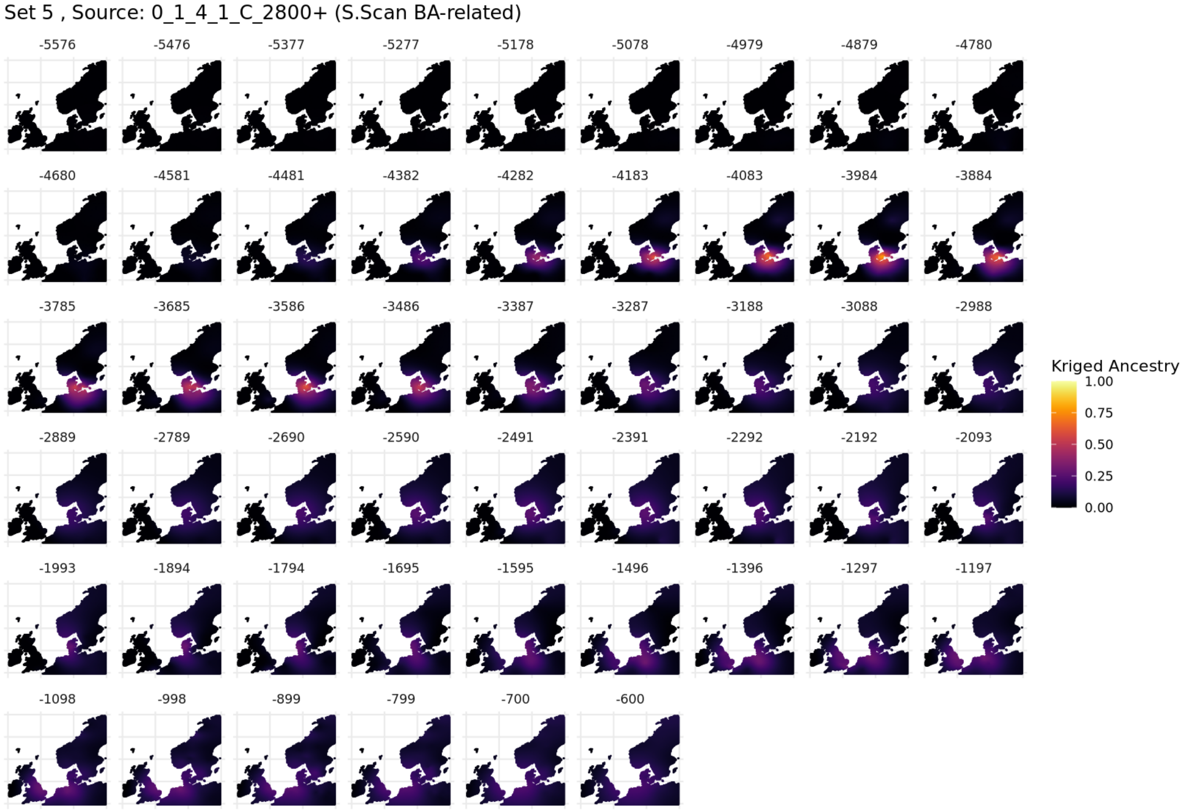

**Fig. S5.97. Spatiotemporal kriging results for Set 5, ‘North / West Europe’ parameters.**

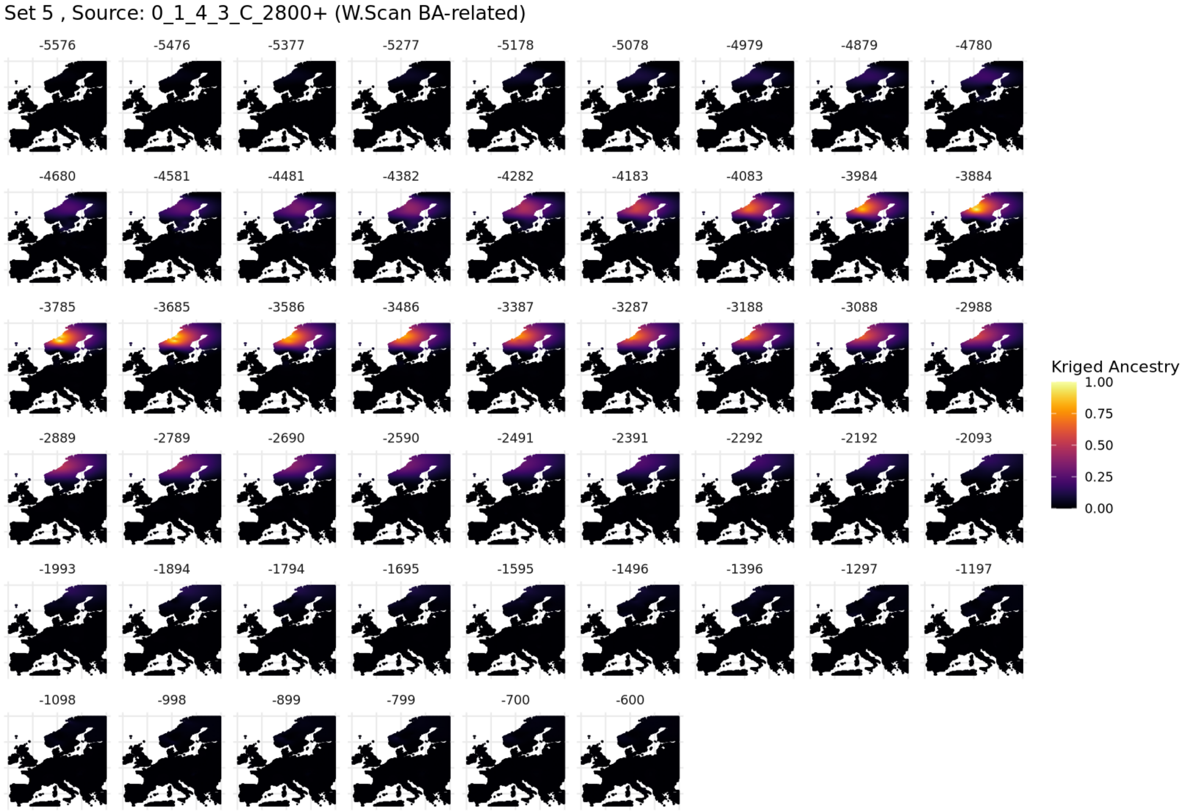

**Fig. S5.98. Spatiotemporal kriging results for Set 5, ‘North / West Europe*’ parameters.**

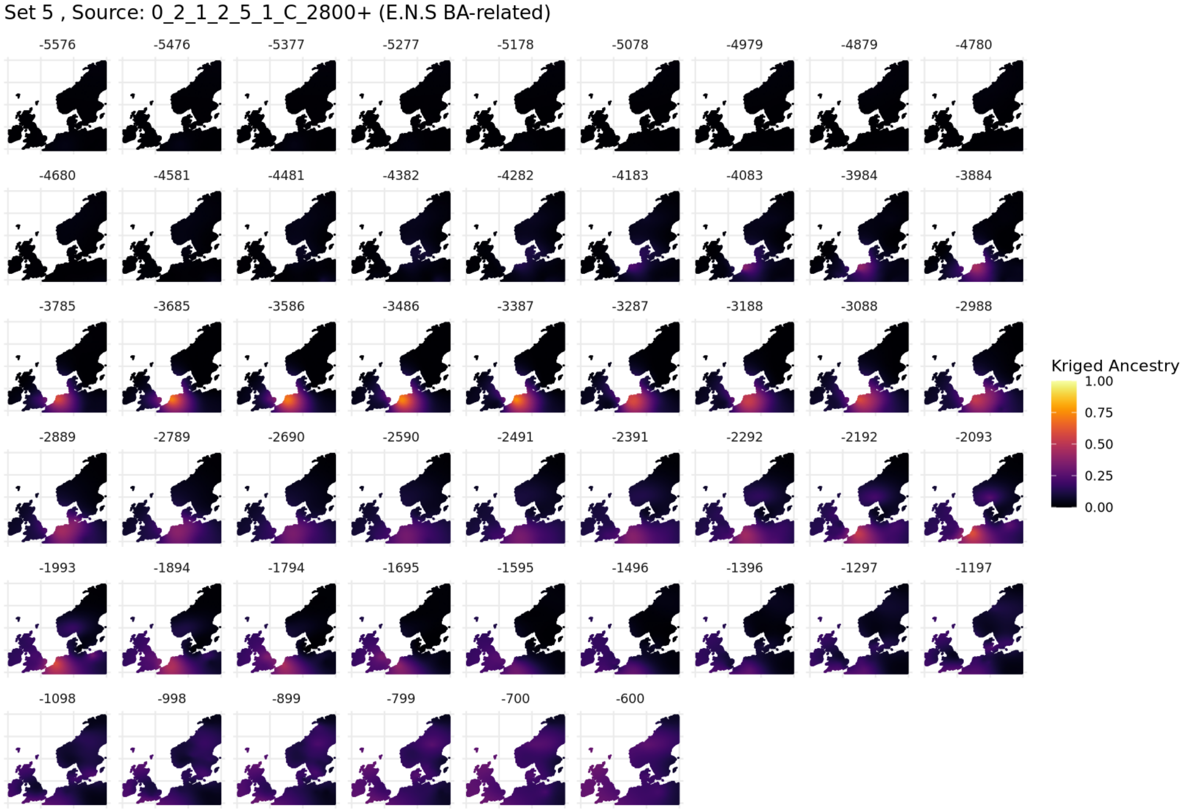

**Fig. S5.99. Spatiotemporal kriging results for Set 5, 'North / West Europe' parameters.**

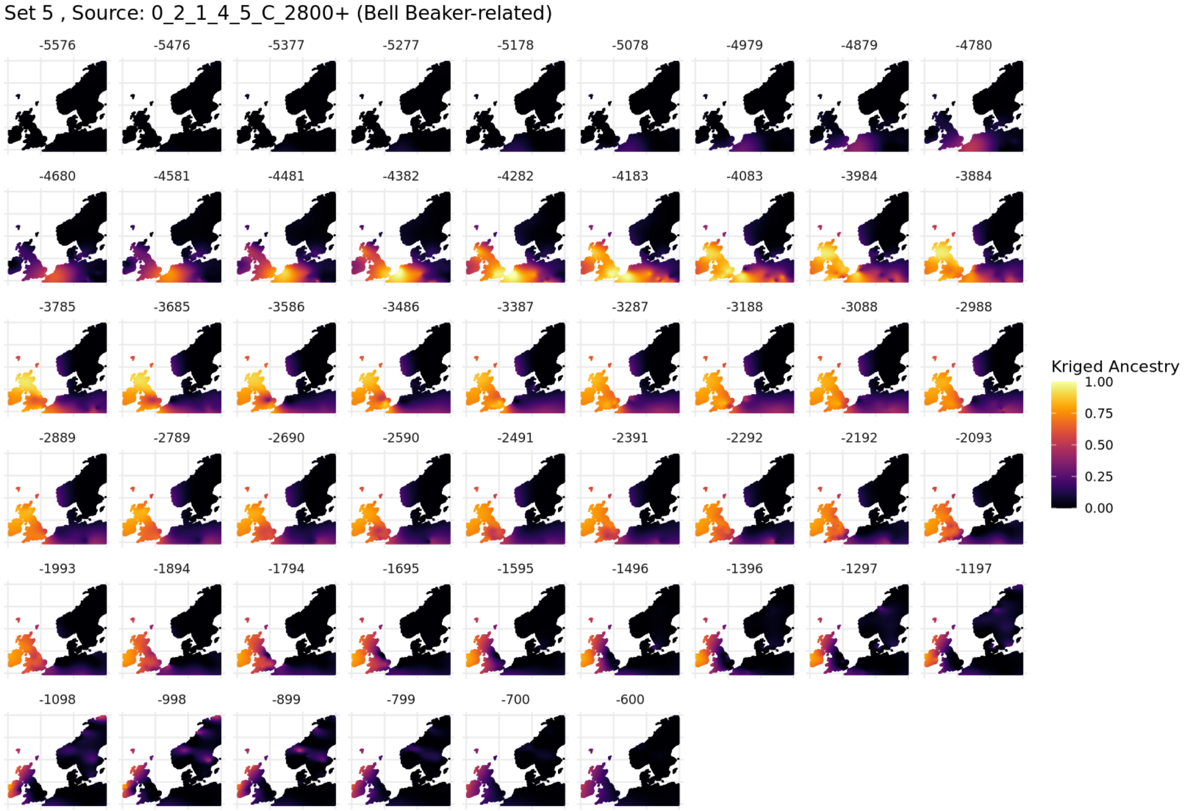

**Fig. S5.100. Spatiotemporal kriging results for Set 5, 'North / West Europe' parameters.**

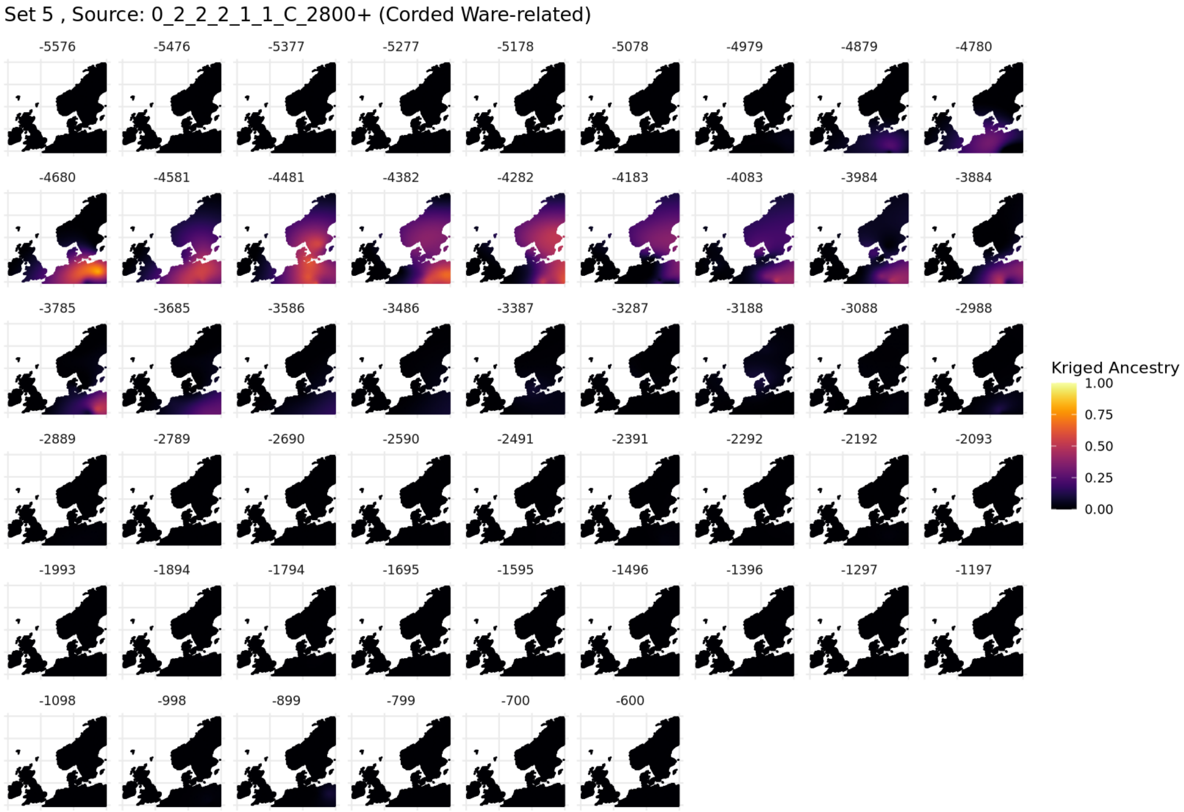

**Fig. S5.101. Spatiotemporal kriging results for Set 5, 'North / West Europe' parameters.**

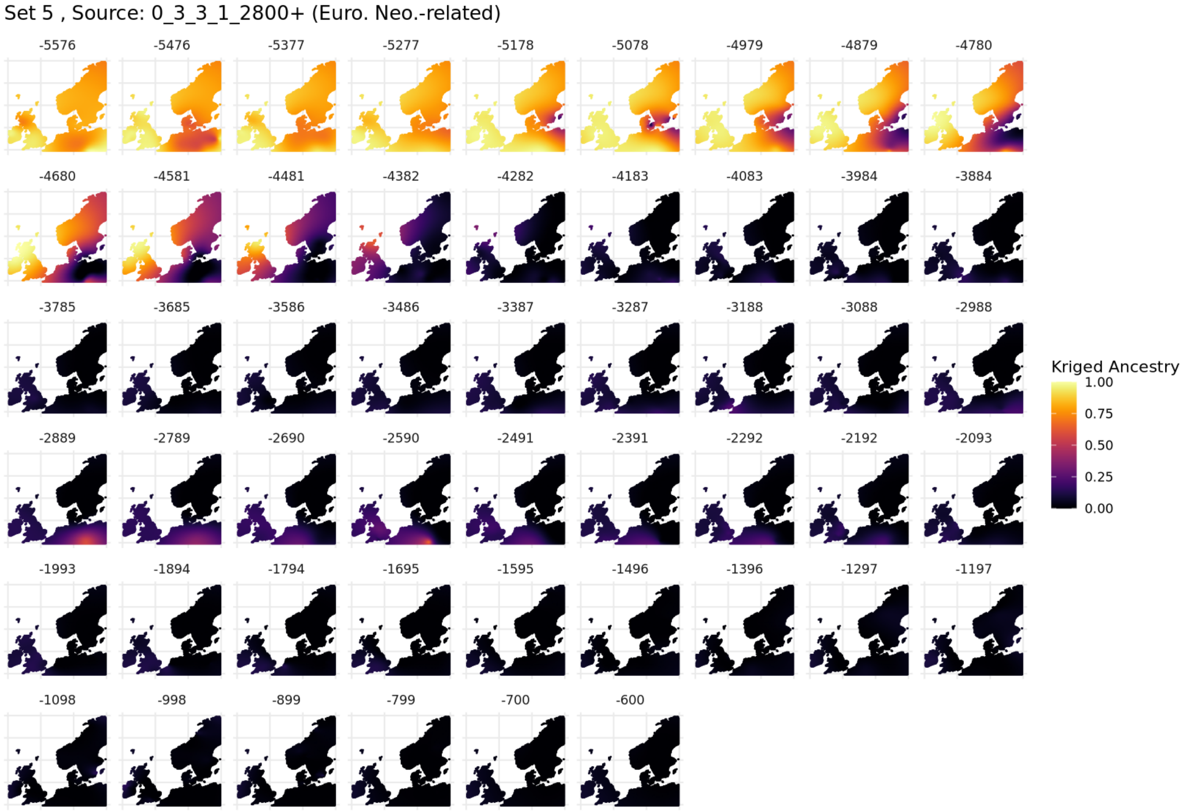

**Fig. S5.102. Spatiotemporal kriging results for Set 5, 'North / West Europe' parameters.**

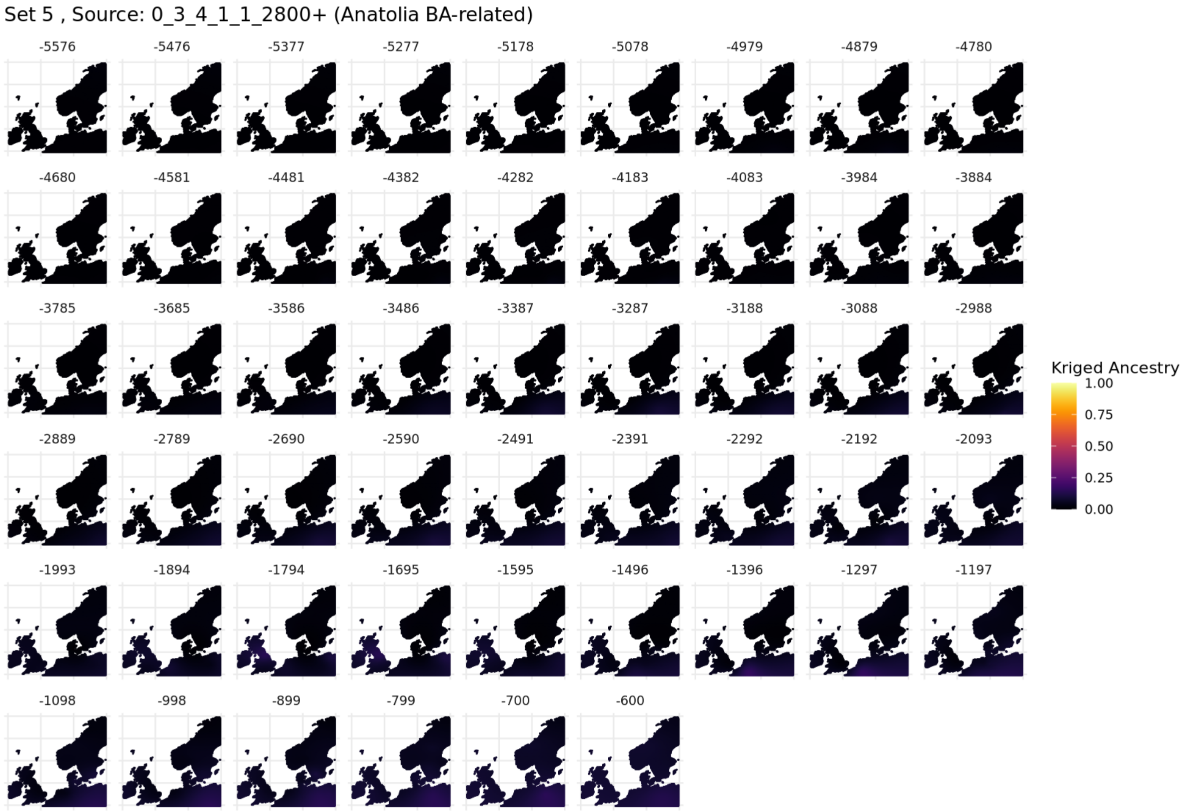

**Fig. S5.103. Spatiotemporal kriging results for Set 5, 'North / West Europe' parameters.**

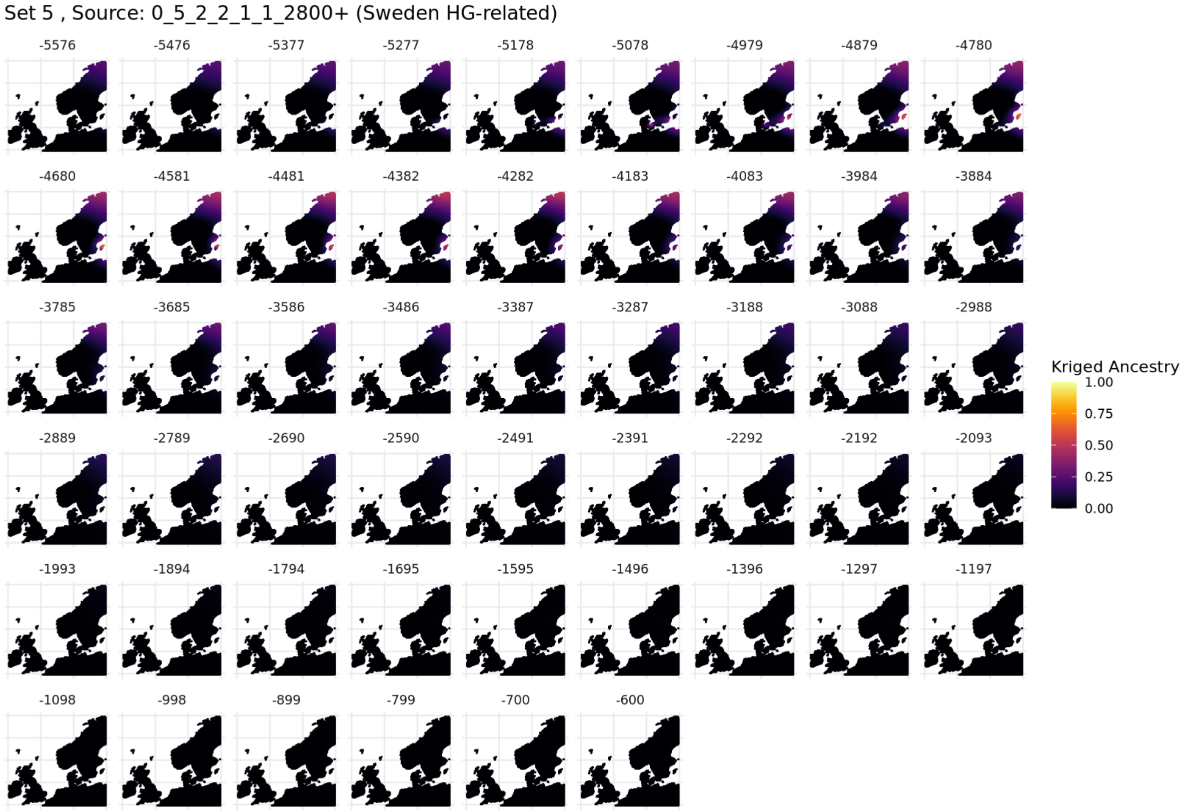

**Fig. S5.104. Spatiotemporal kriging results for Set 5, 'North / West Europe' parameters.**

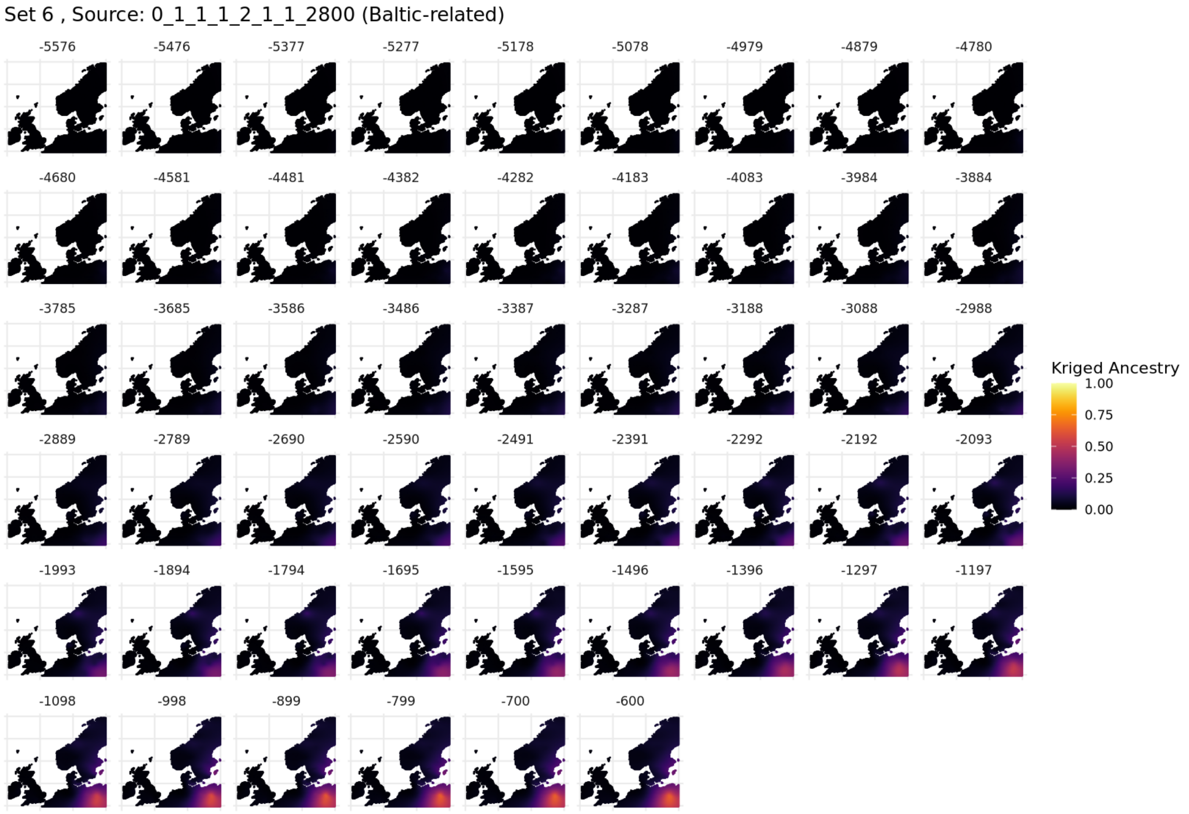

**Fig. S5.105. Spatiotemporal kriging results for Set 6, 'North / West Europe' parameters.**

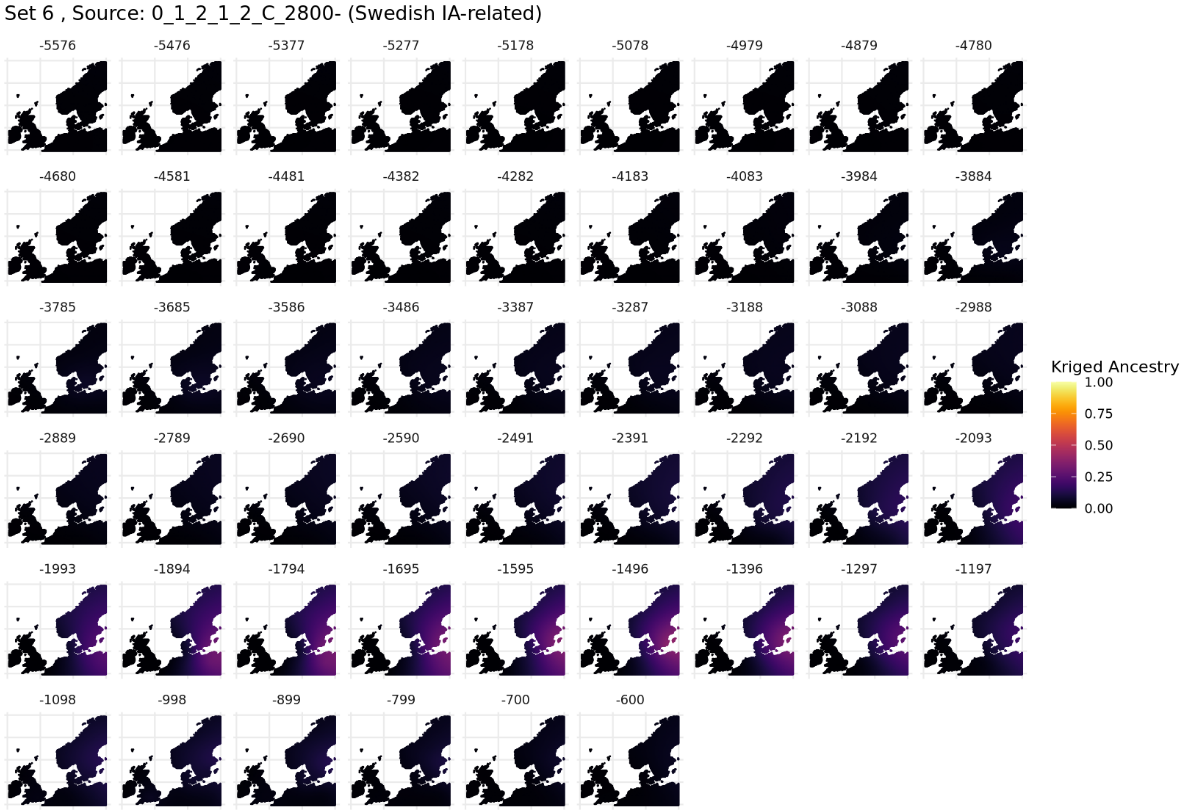

**Fig. S5.106. Spatiotemporal kriging results for Set 6, 'North / West Europe' parameters.**

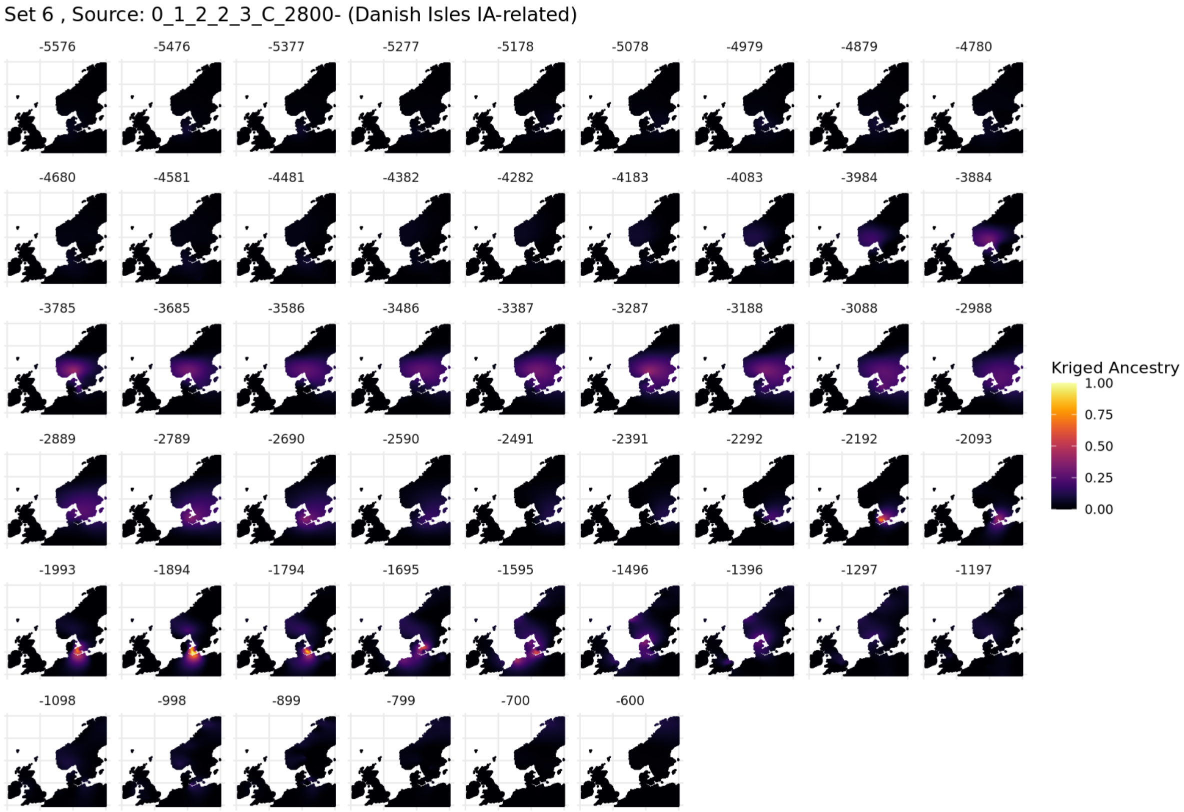

**Fig. S5.107. Spatiotemporal kriging results for Set 6, 'North / West Europe' parameters.**

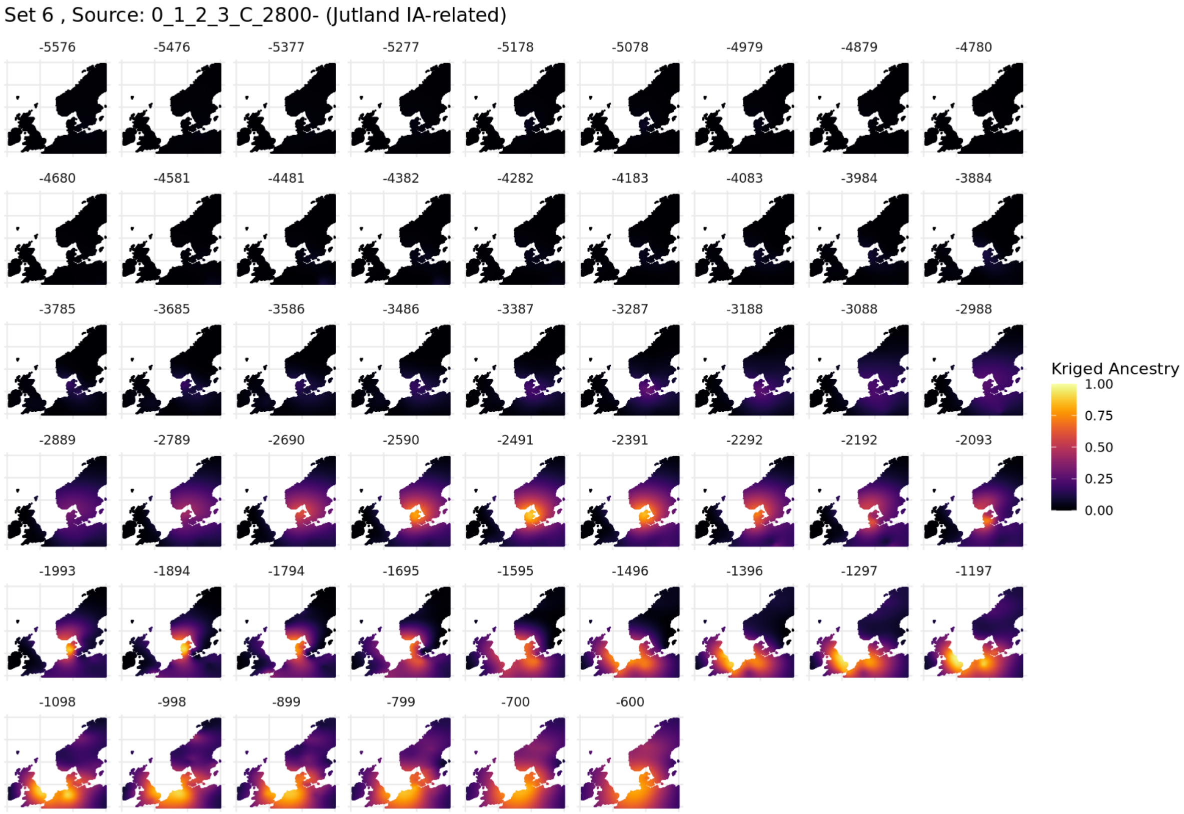

**Fig. S5.108. Spatiotemporal kriging results for Set 6, 'North / West Europe' parameters.**

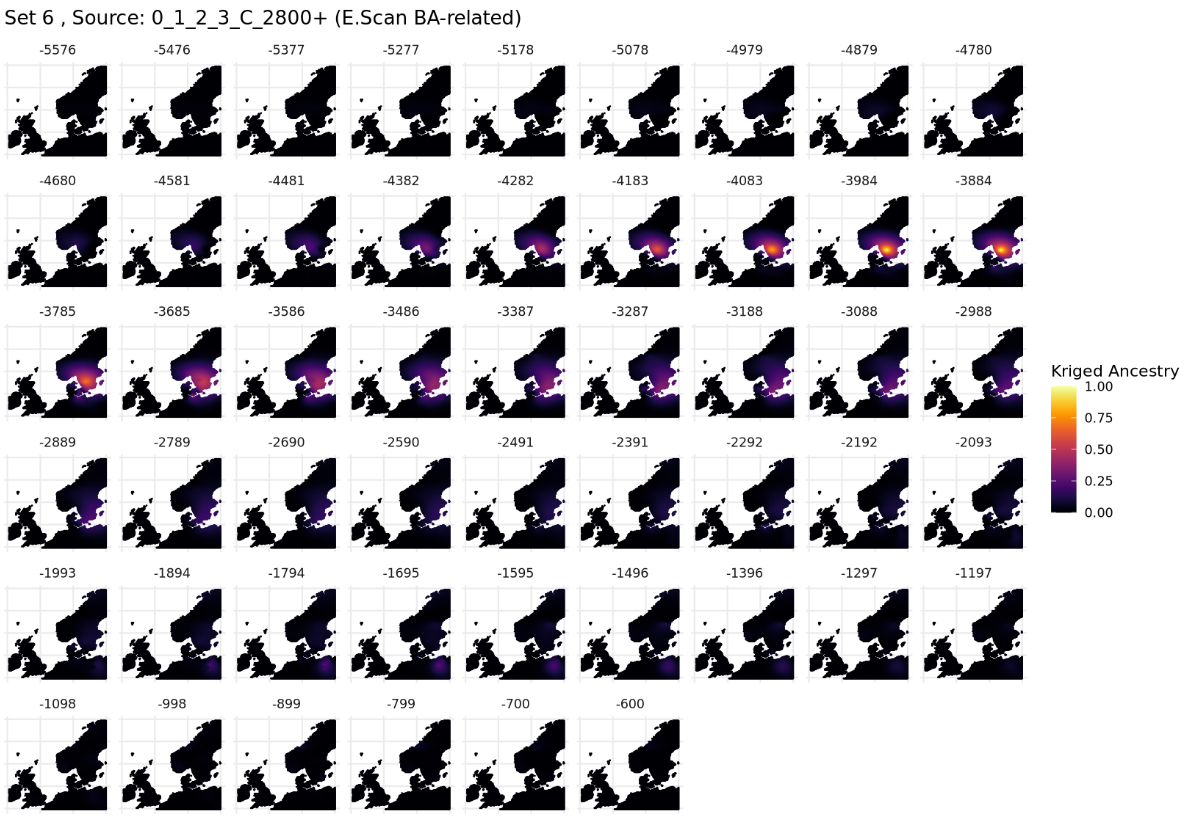

**Fig. S5.109. Spatiotemporal kriging results for Set 6, 'North / West Europe' parameters.**

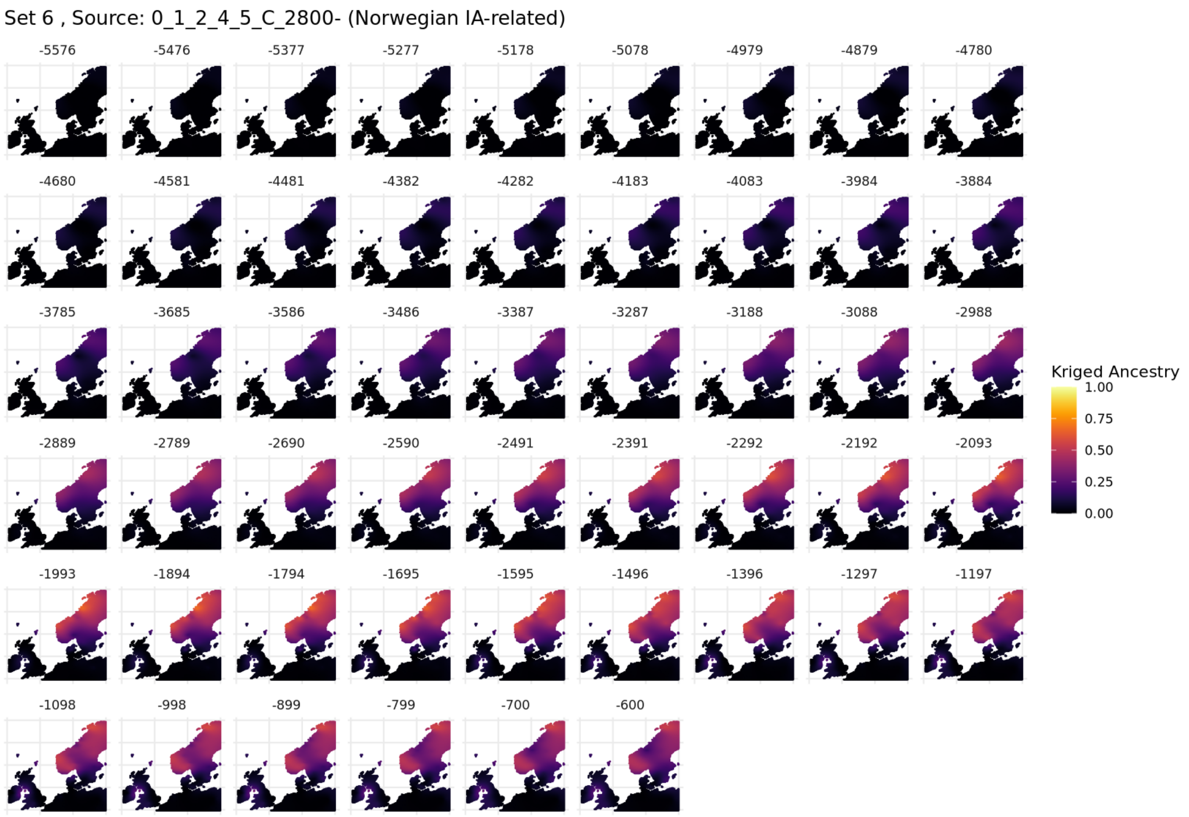

**Fig. S5.110. Spatiotemporal kriging results for Set 6, 'North / West Europe' parameters.**

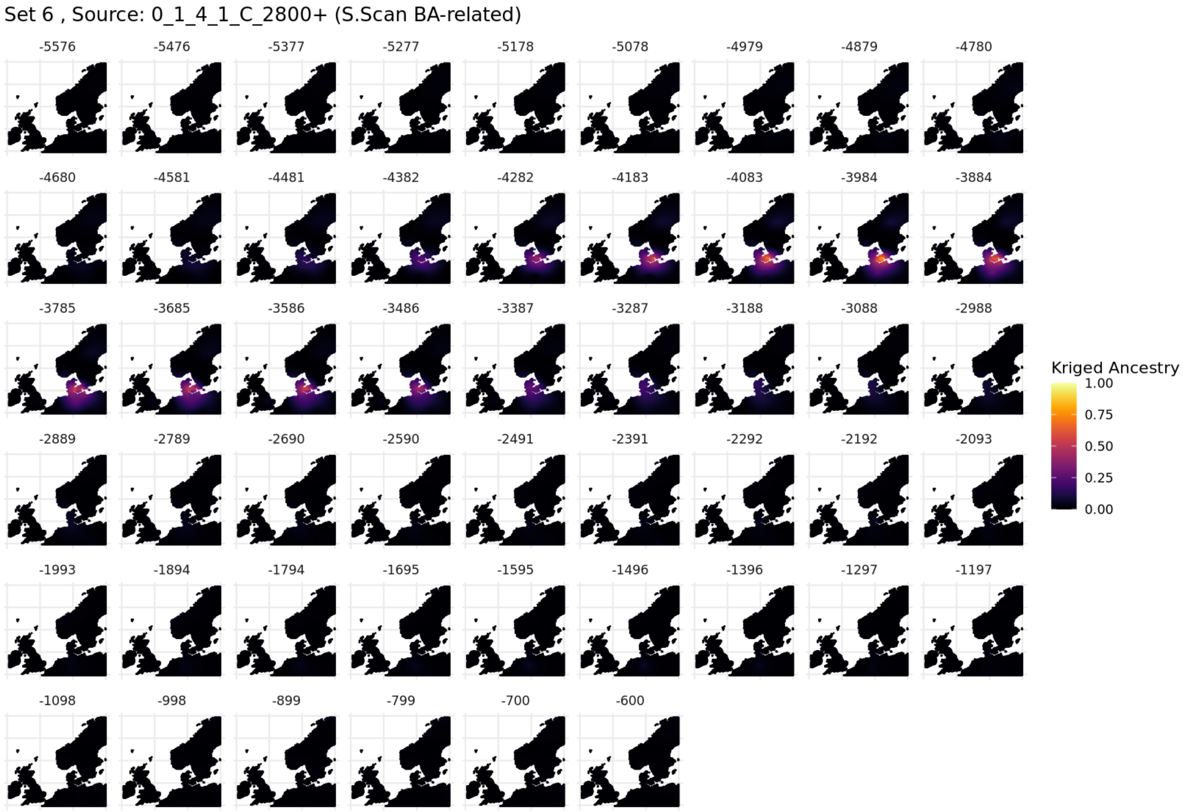

**Fig. S5.111. Spatiotemporal kriging results for Set 6, 'North / West Europe' parameters.**

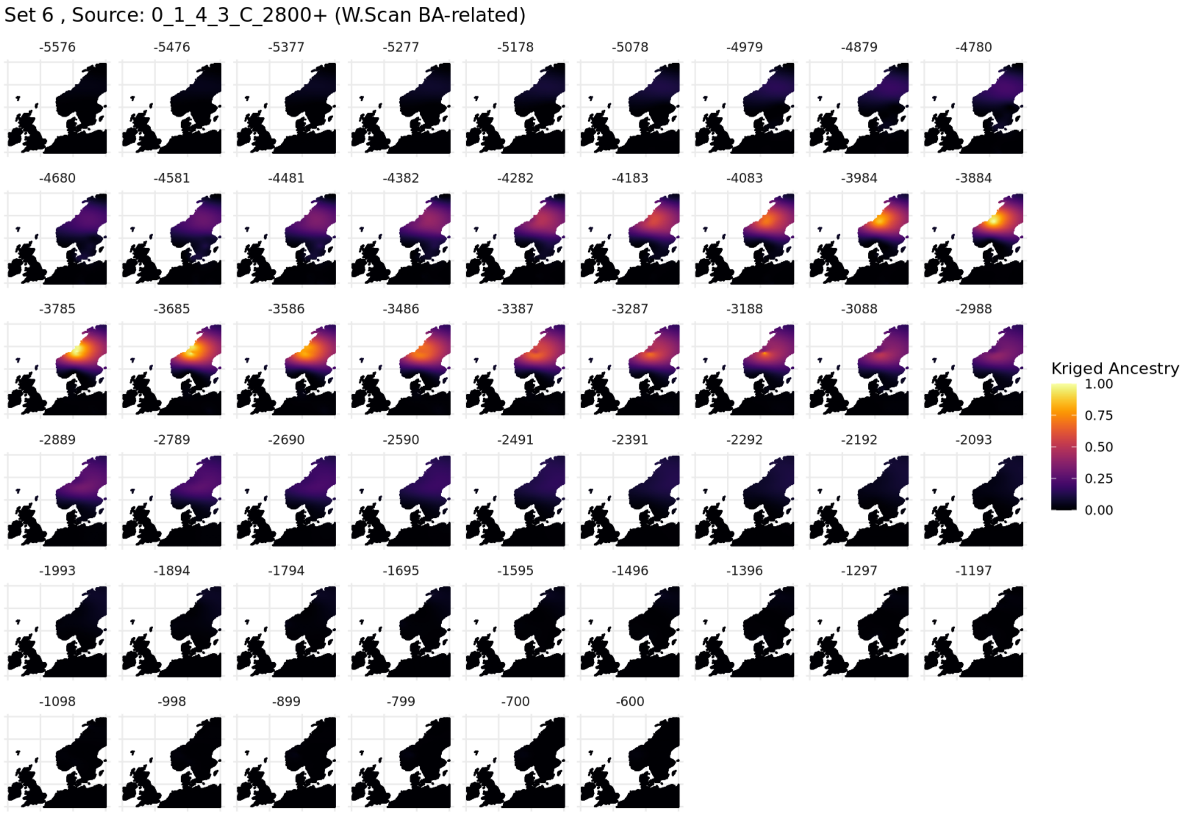

**Fig. S5.112. Spatiotemporal kriging results for Set 6, 'North / West Europe' parameters.**

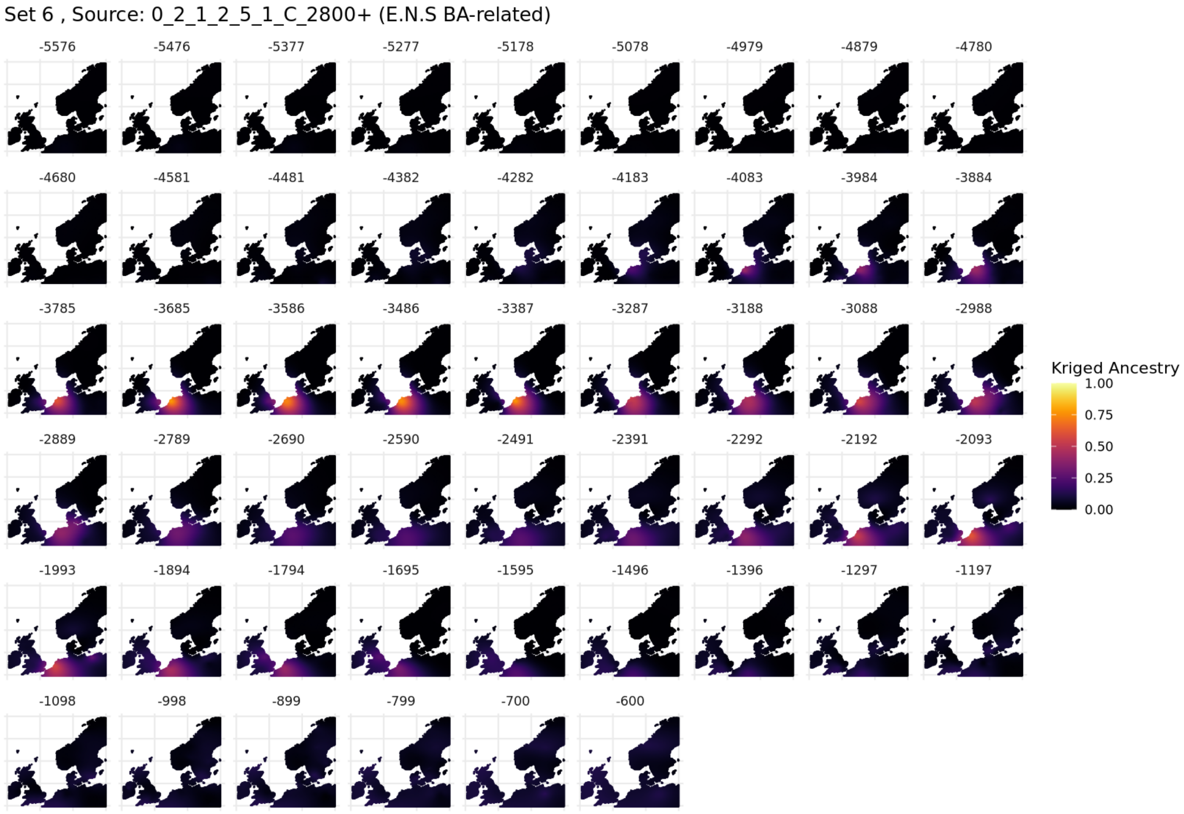

**Fig. S5.113. Spatiotemporal kriging results for Set 6, 'North / West Europe' parameters.**

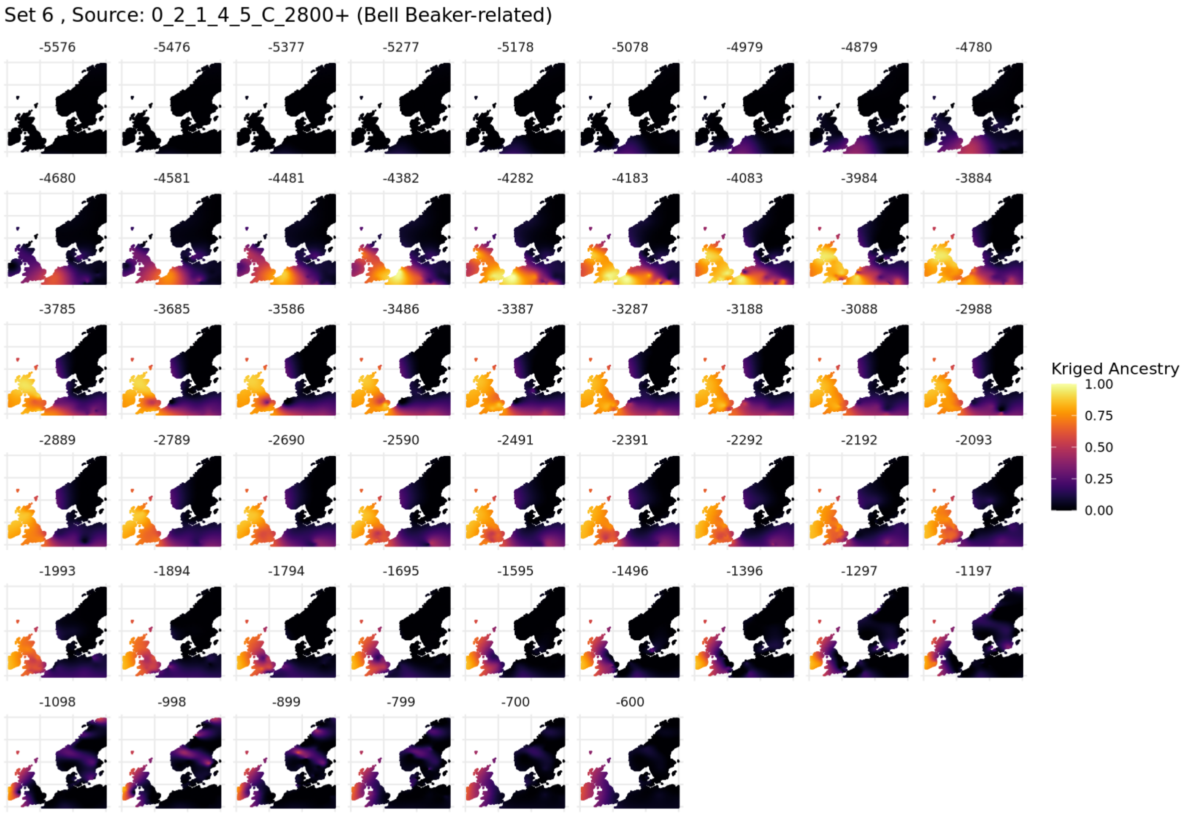

**Fig. S5.114. Spatiotemporal kriging results for Set 6, 'North / West Europe' parameters.**

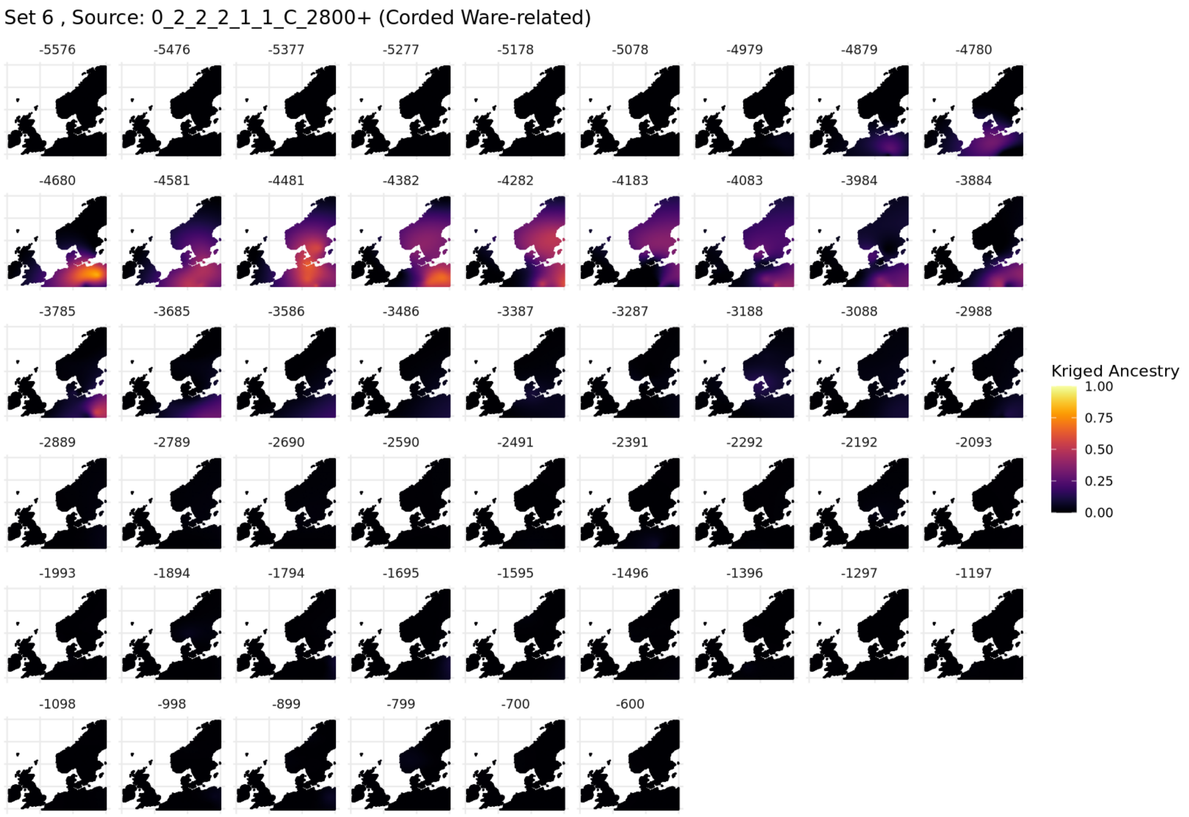

**Fig. S5.115. Spatiotemporal kriging results for Set 6, 'North / West Europe' parameters.**

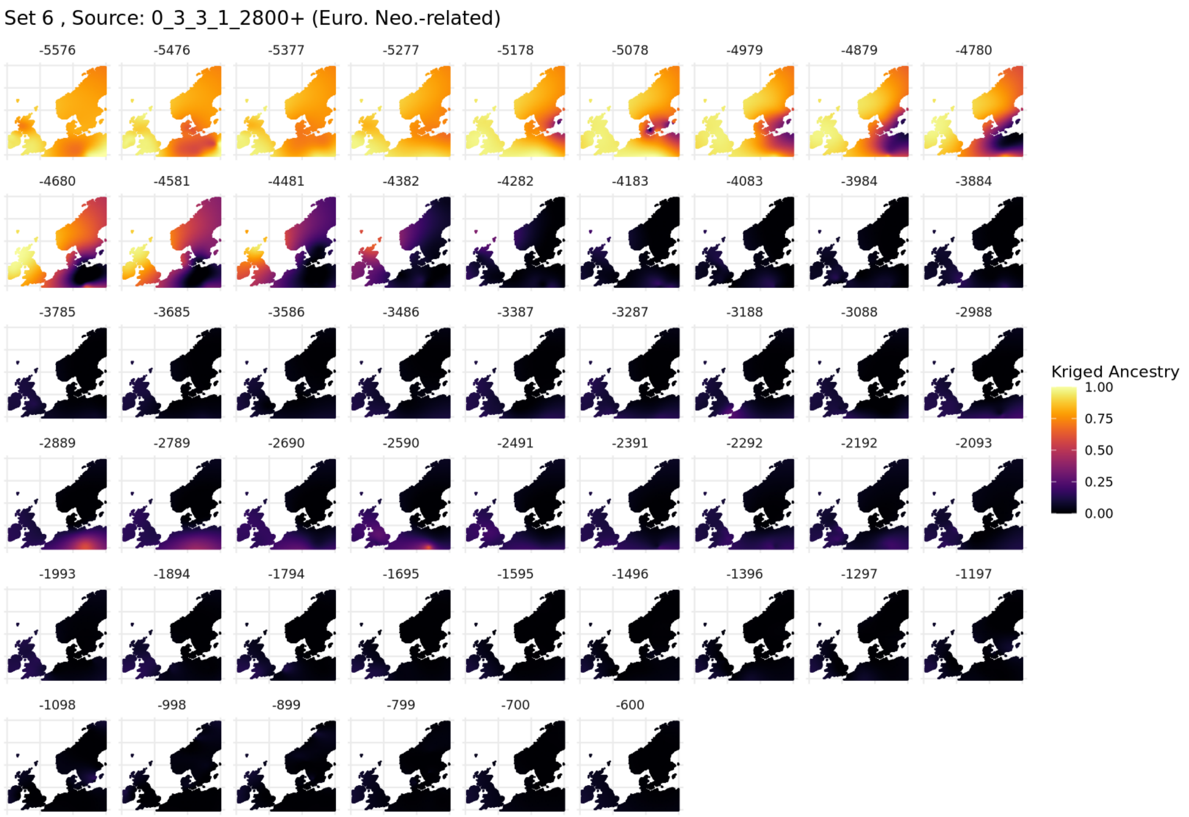

**Fig. S5.116. Spatiotemporal kriging results for Set 6, 'North / West Europe' parameters.**

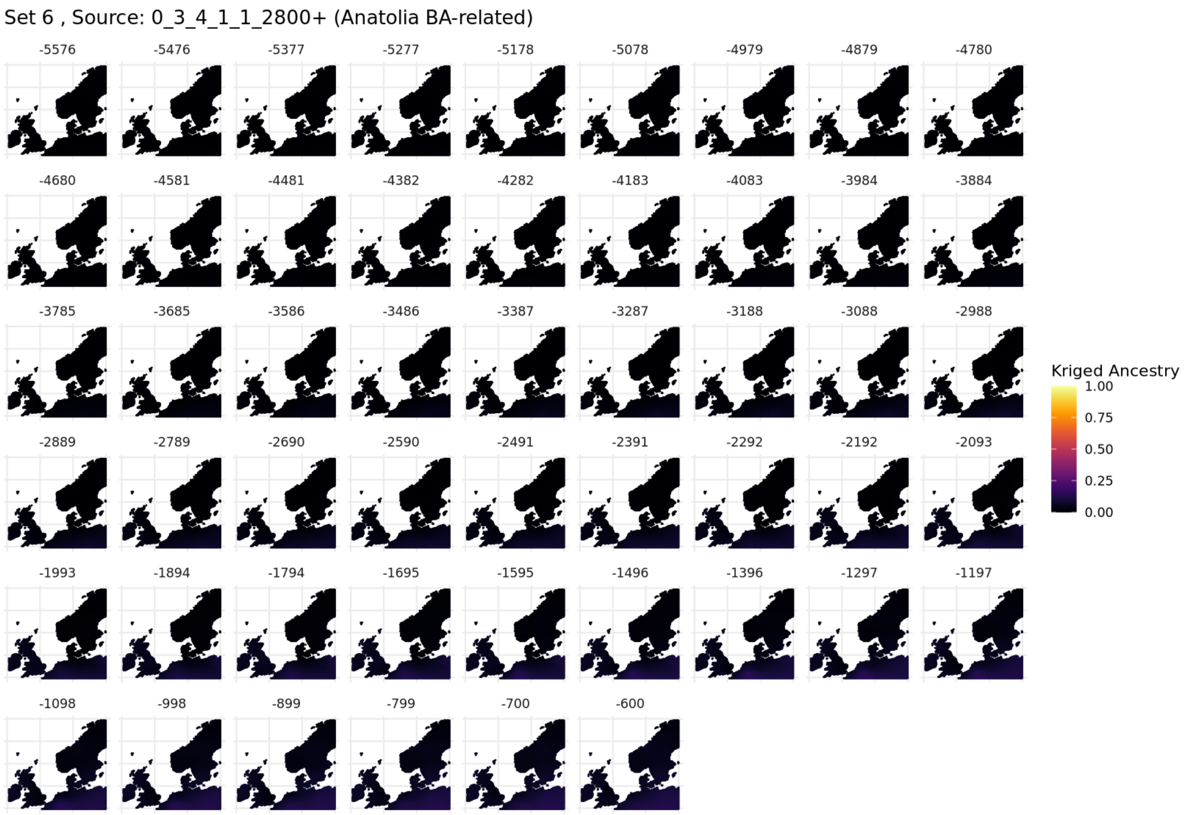

**Fig. S5.117. Spatiotemporal kriging results for Set 6, 'North / West Europe' parameters.**

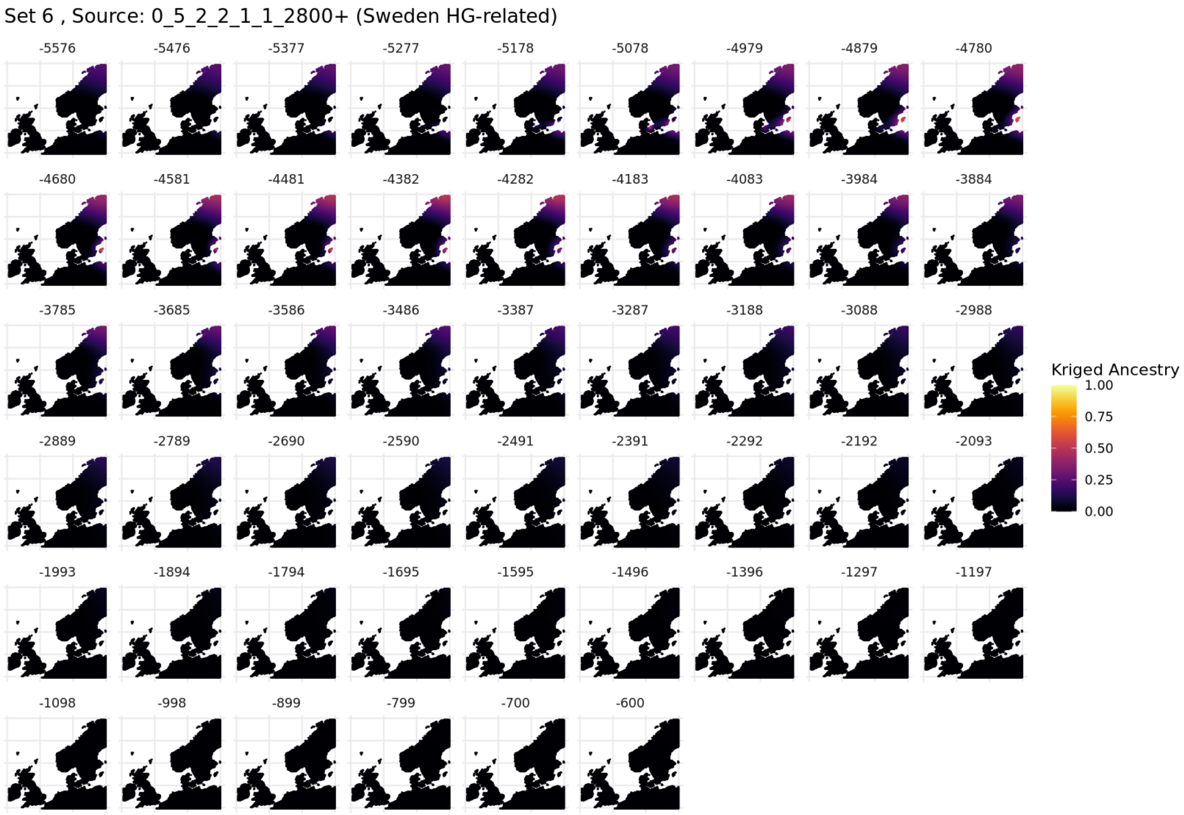

**Fig. S5.118. Spatiotemporal kriging results for Set 6, 'North / West Europe' parameters.**

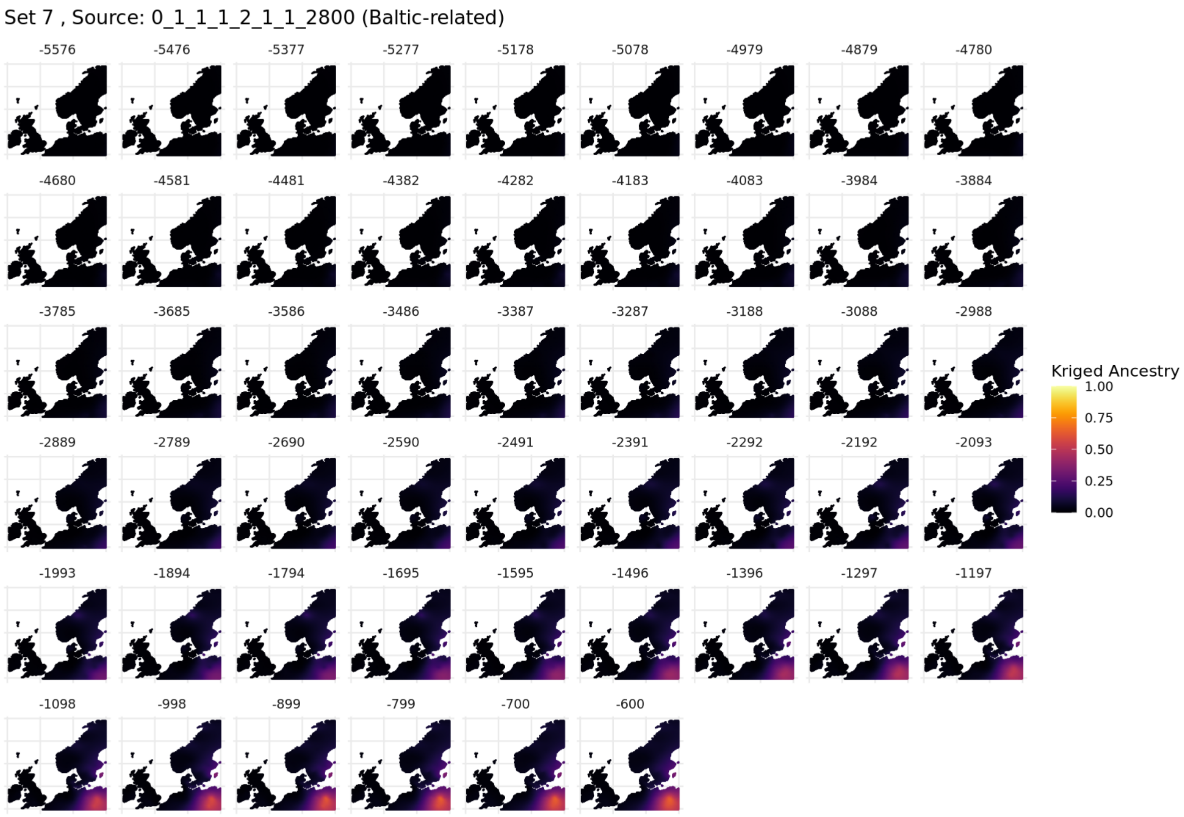

**Fig. S5.119. Spatiotemporal kriging results for Set 7, 'North / West Europe' parameters.**

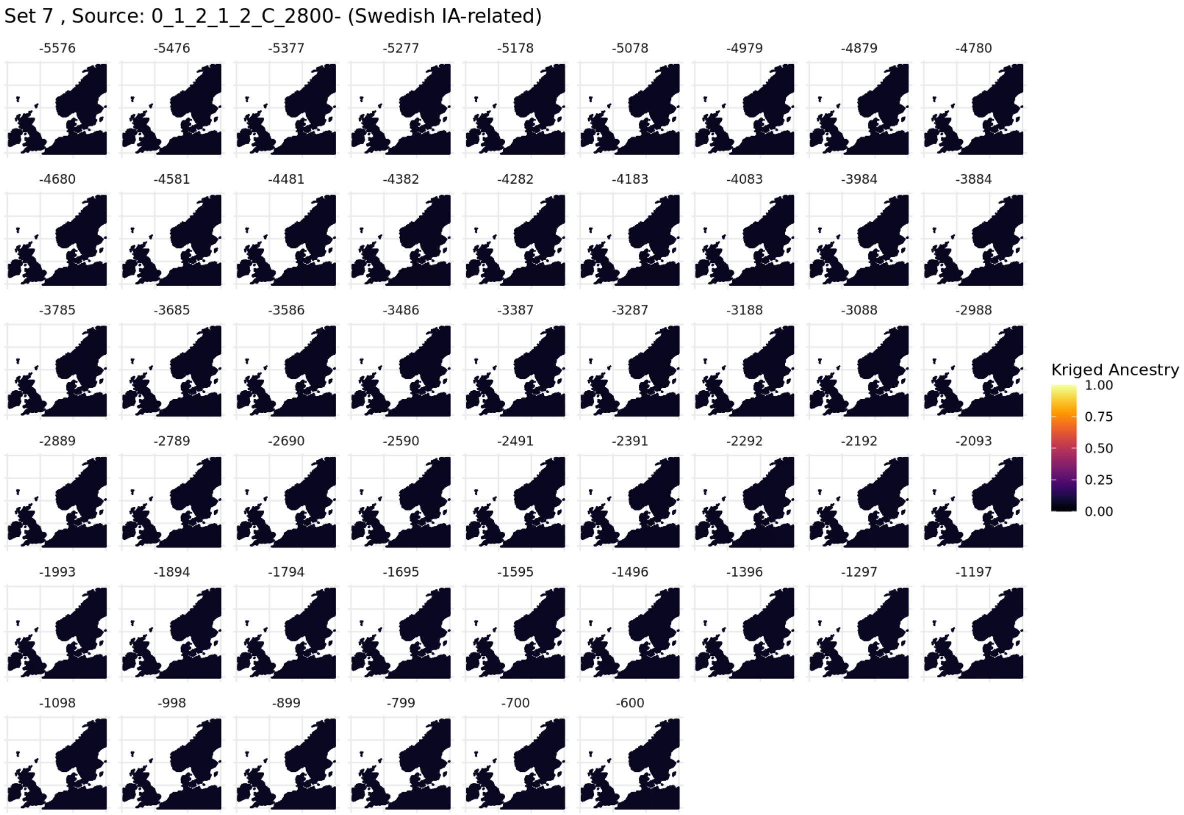

**Fig. S5.120. Spatiotemporal kriging results for Set 7, 'North / West Europe' parameters.**

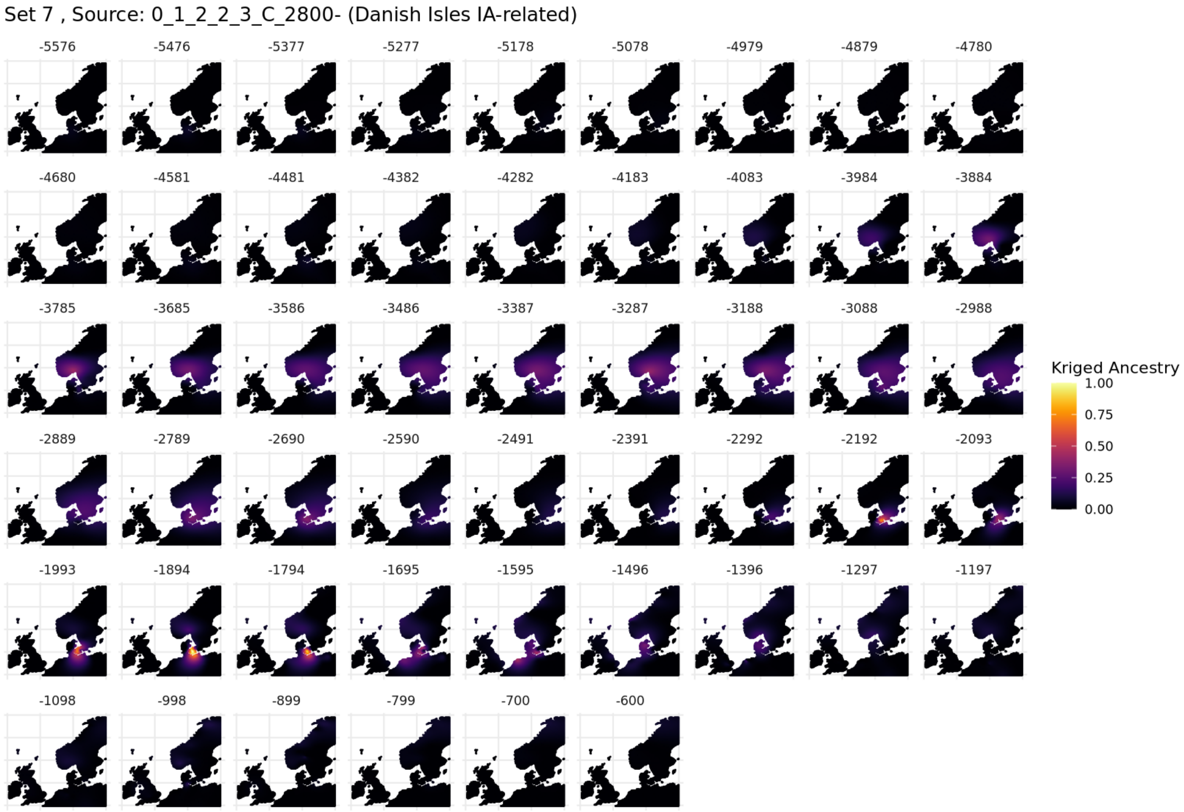

**Fig. S5.121. Spatiotemporal kriging results for Set 7, 'North / West Europe' parameters.**

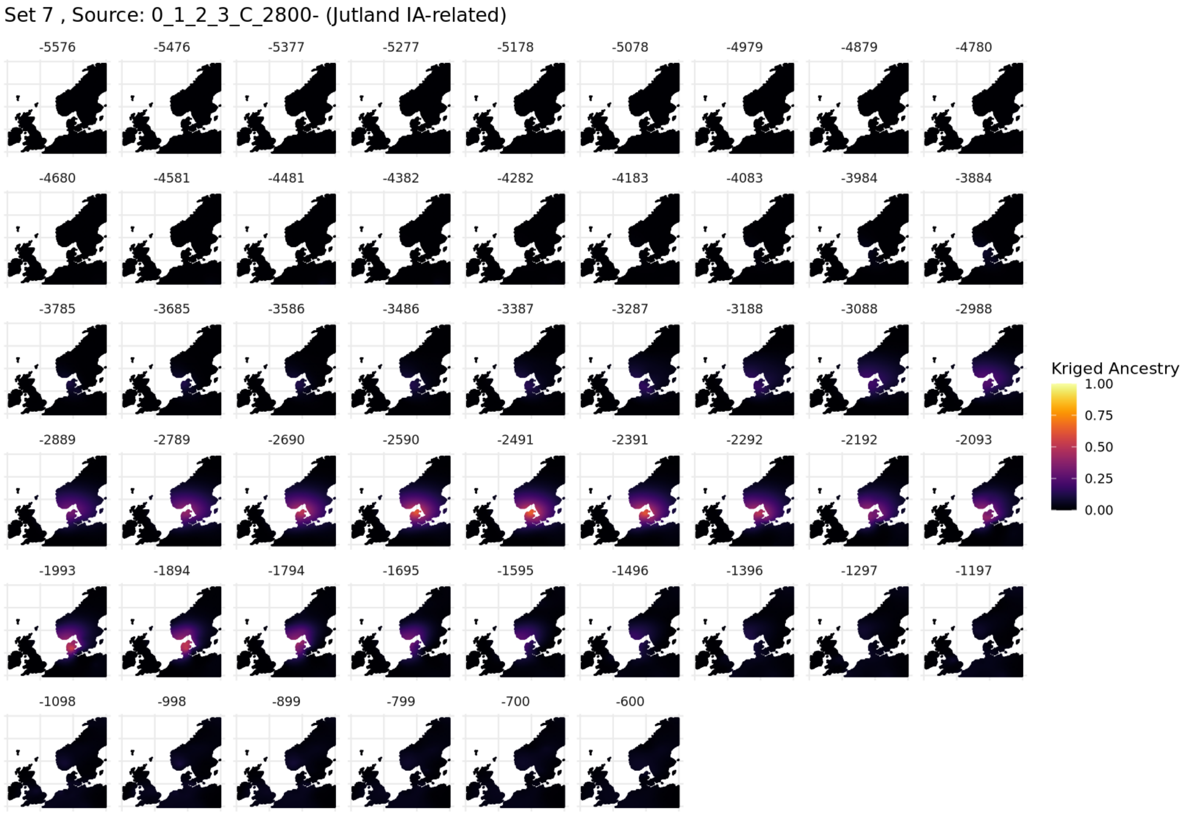

**Fig. S5.122. Spatiotemporal kriging results for Set 7, 'North / West Europe' parameters.**

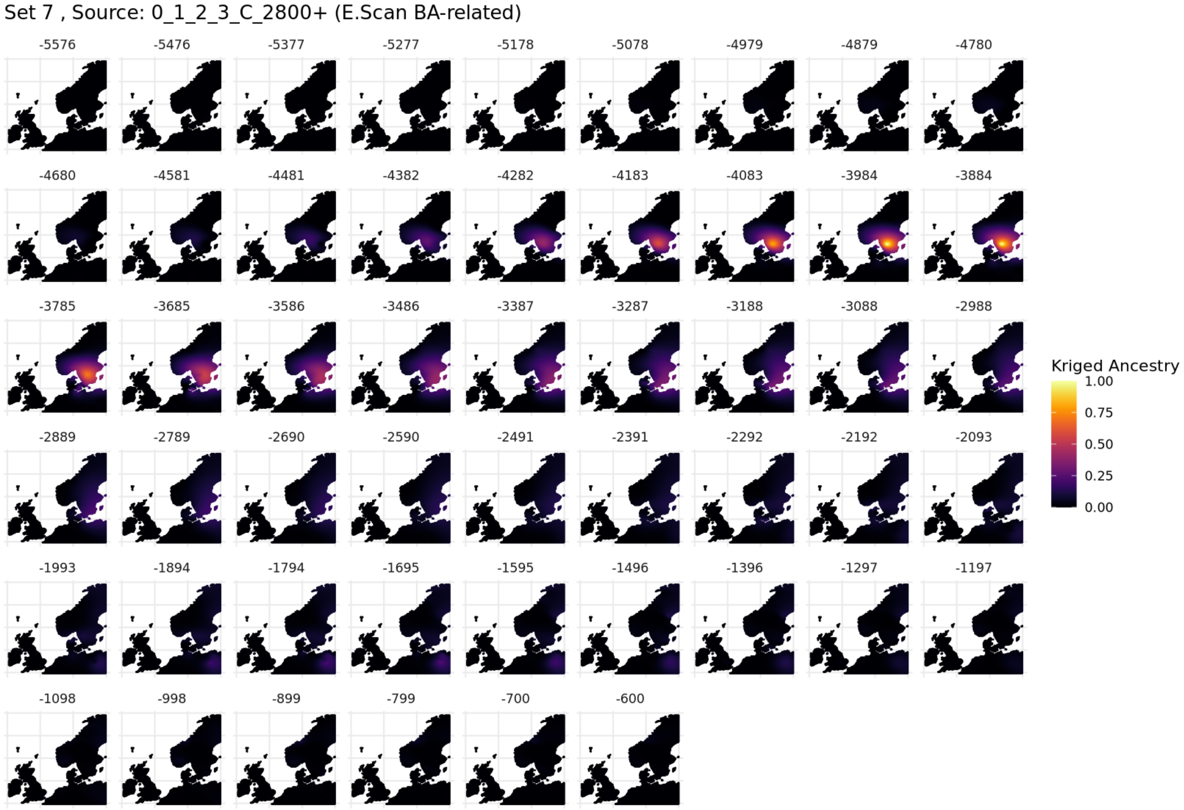

**Fig. S5.123. Spatiotemporal kriging results for Set 7, 'North / West Europe' parameters.**

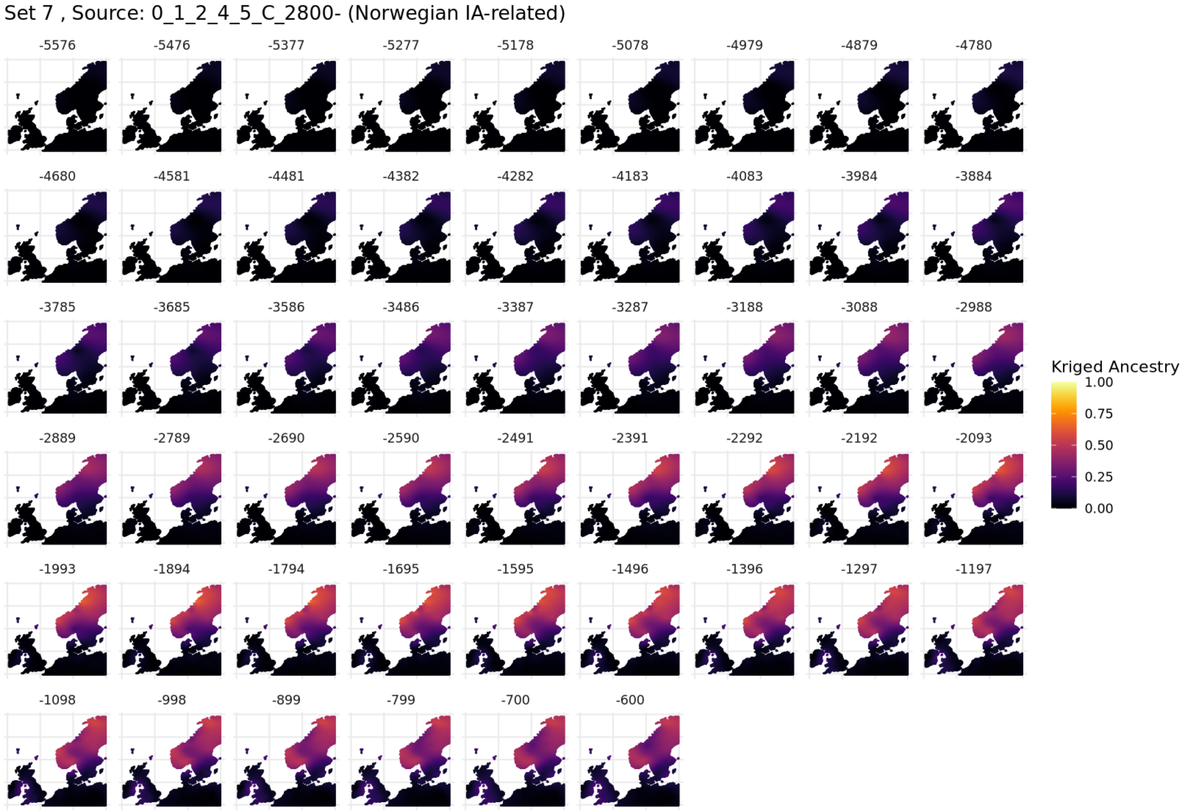

**Fig. S5.124. Spatiotemporal kriging results for Set 7, 'North / West Europe' parameters.**

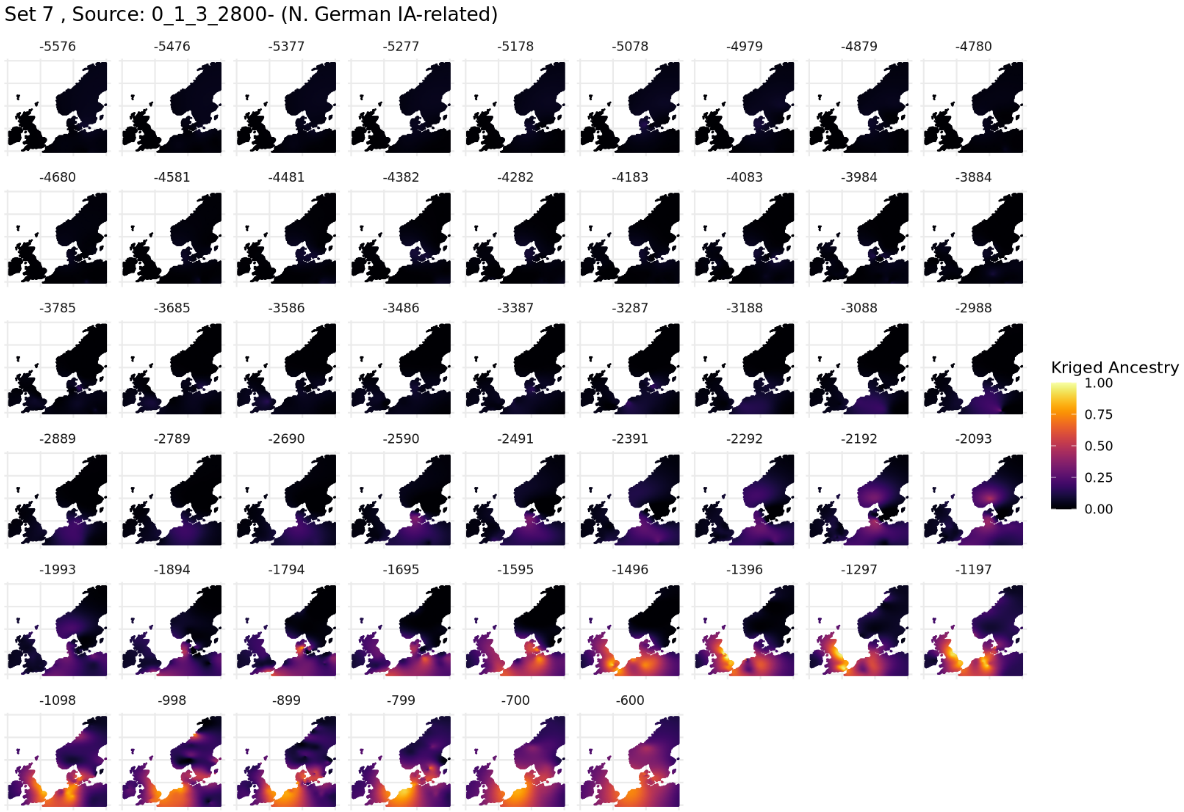

**Fig. S5.125. Spatiotemporal kriging results for Set 7, 'North / West Europe' parameters.**

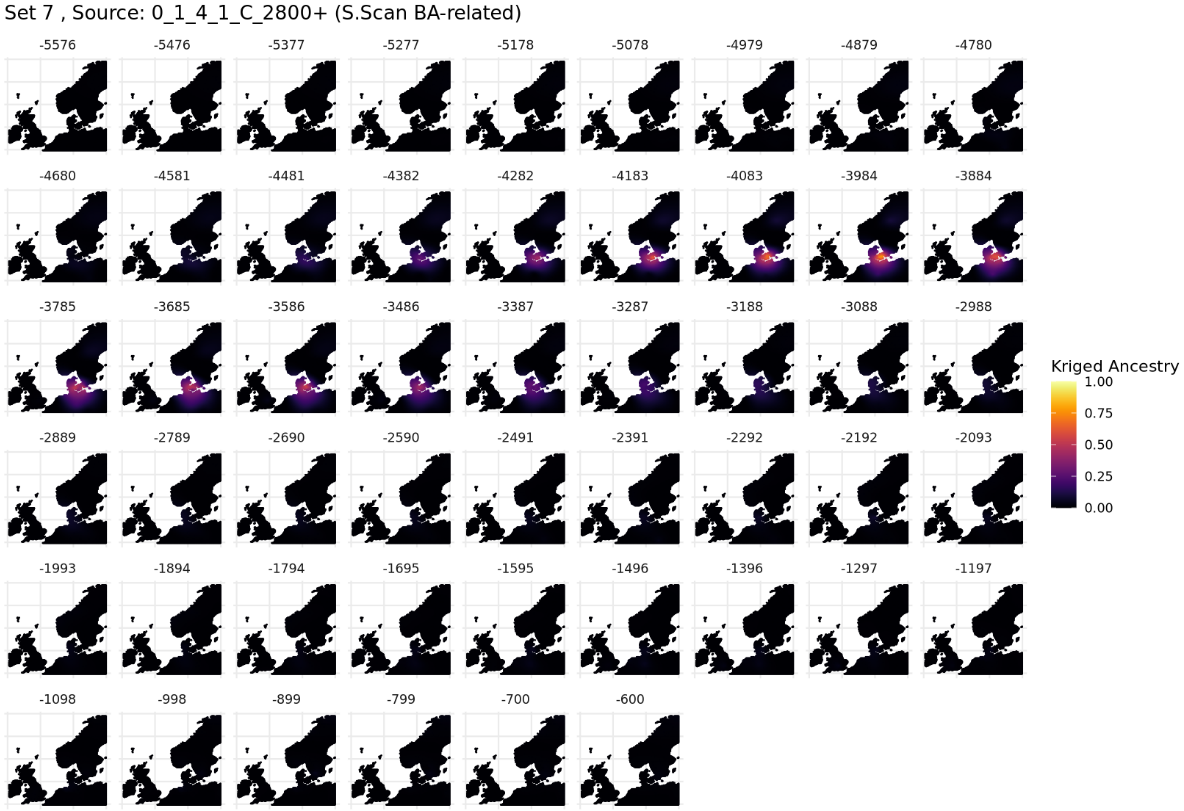

**Fig. S5.126. Spatiotemporal kriging results for Set 7, 'North / West Europe' parameters.**

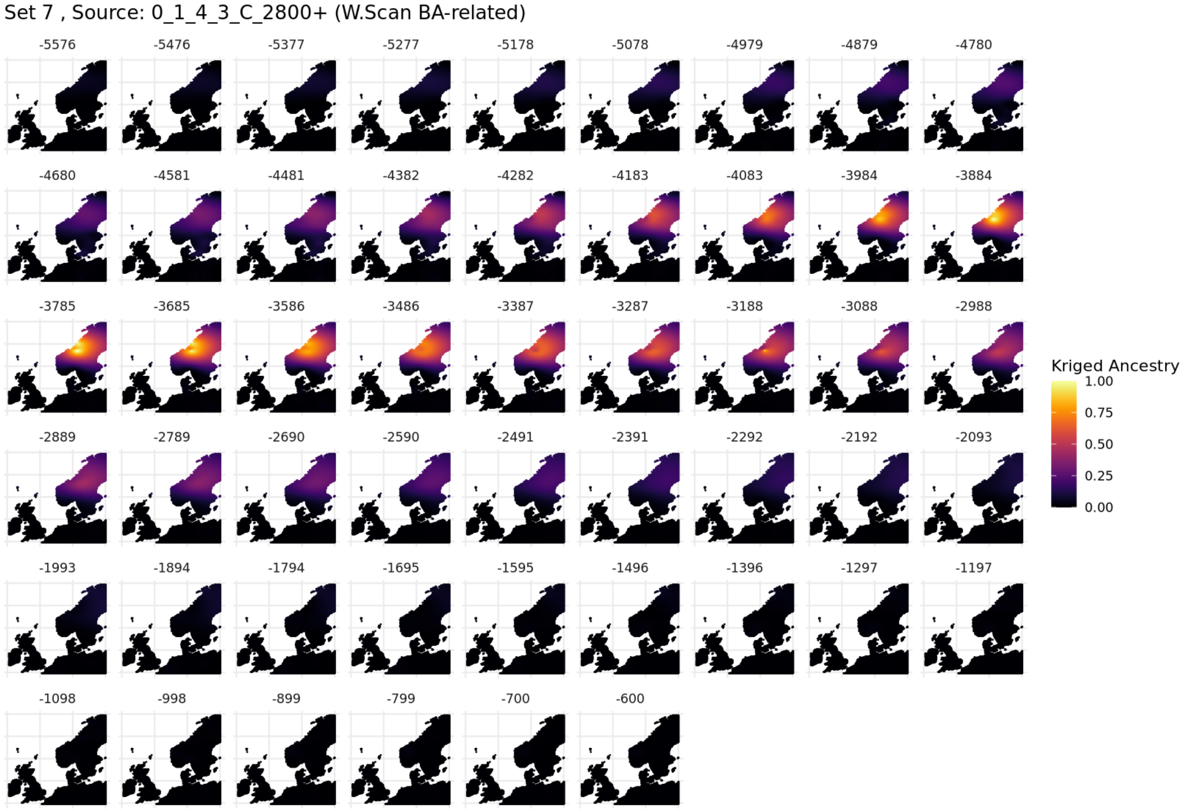

1585

1586 **Fig. S5.127. Spatiotemporal kriging results for Set 7, 'North / West Europe' parameters.**

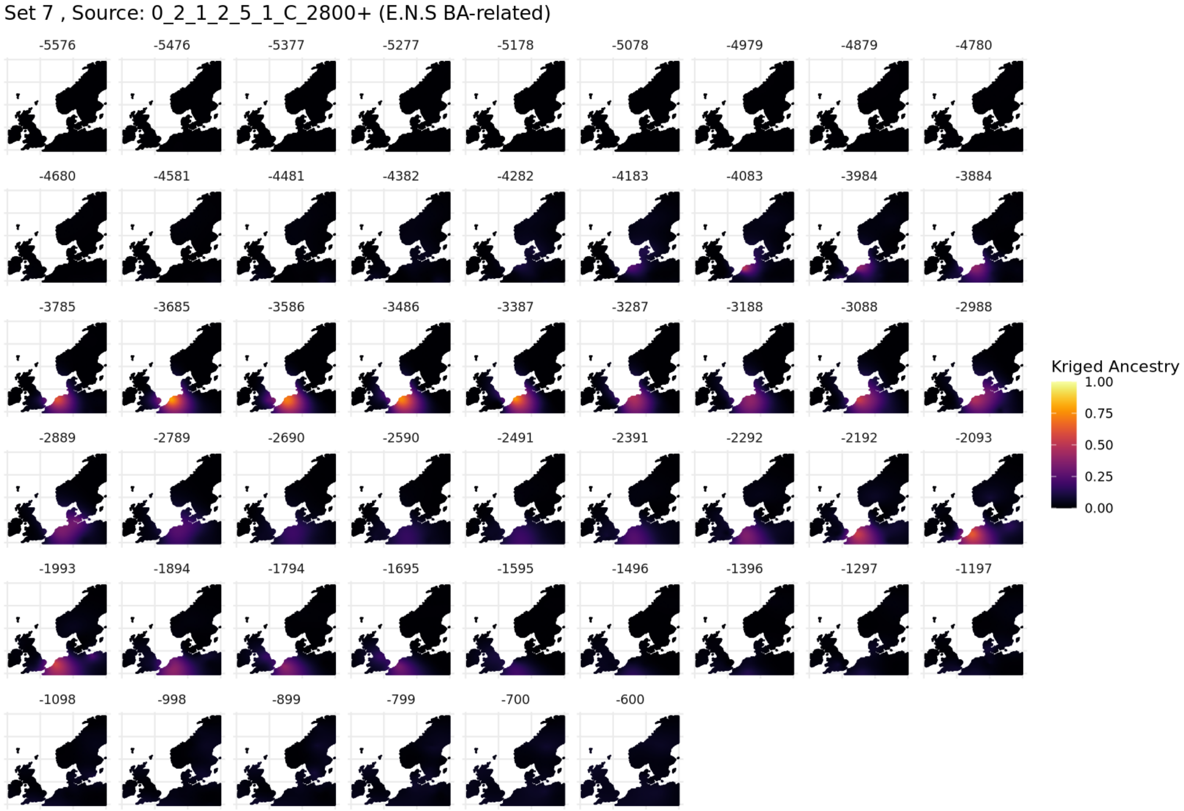

1587

1588 **Fig. S5.128. Spatiotemporal kriging results for Set 7, 'North / West Europe' parameters.**

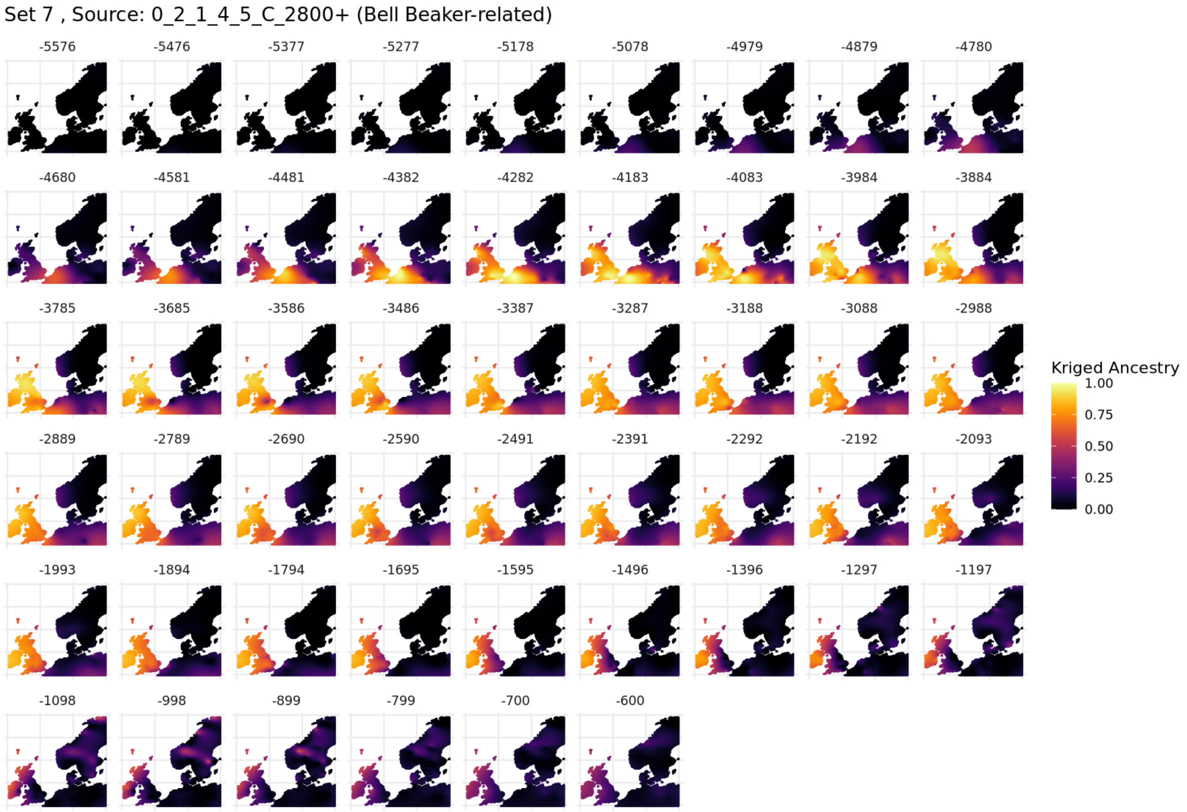

**Fig. S5.129.0 Spatiotemporal kriging results for Set 7, 'North / West Europe' parameters.**

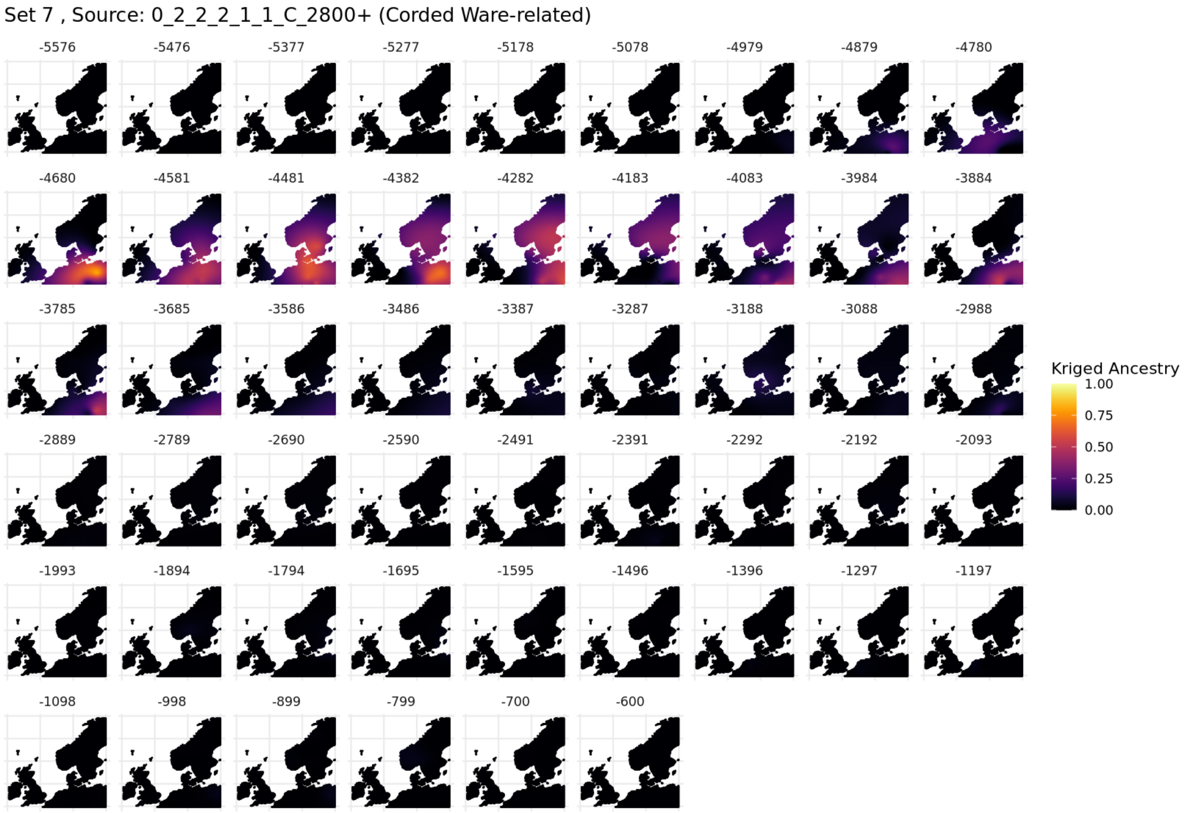

**Fig. S5.130. Spatiotemporal kriging results for Set 7, 'North / West Europe' parameters.**

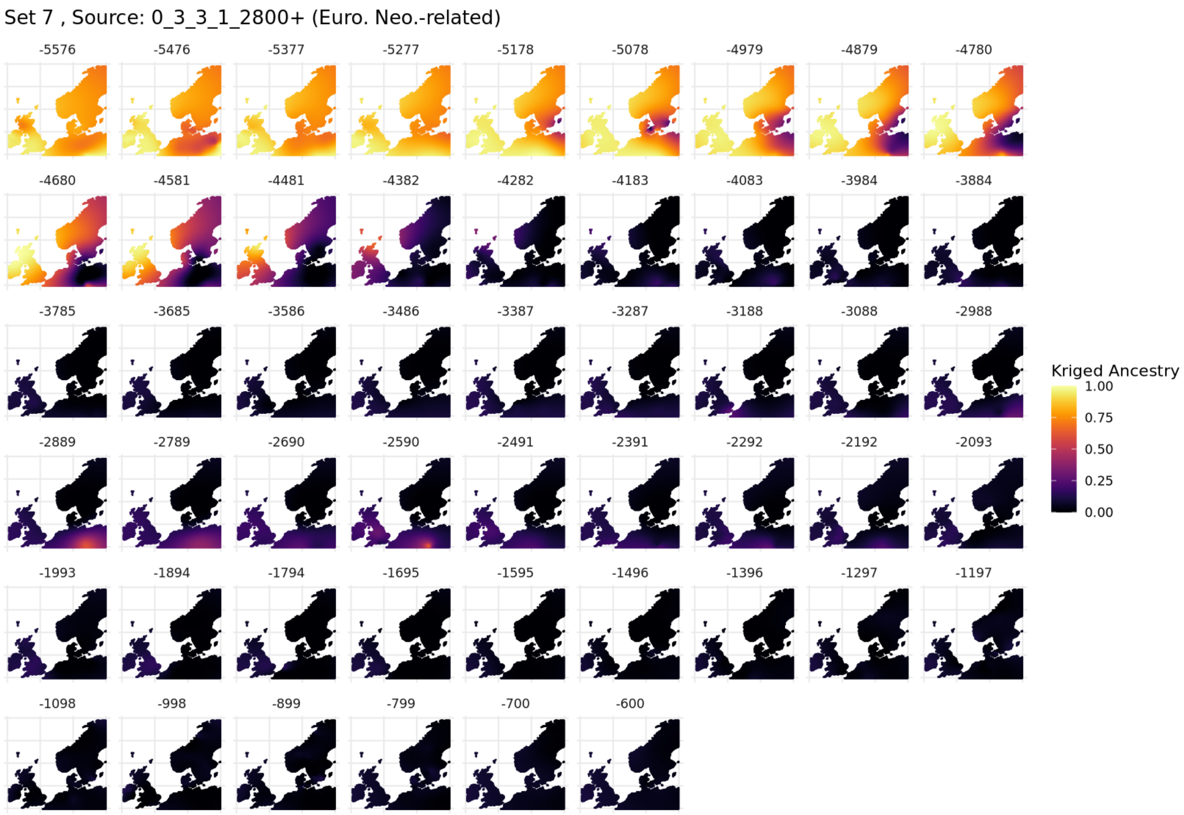

**Fig. S5.131. Spatiotemporal kriging results for Set 7, 'North / West Europe' parameters.**

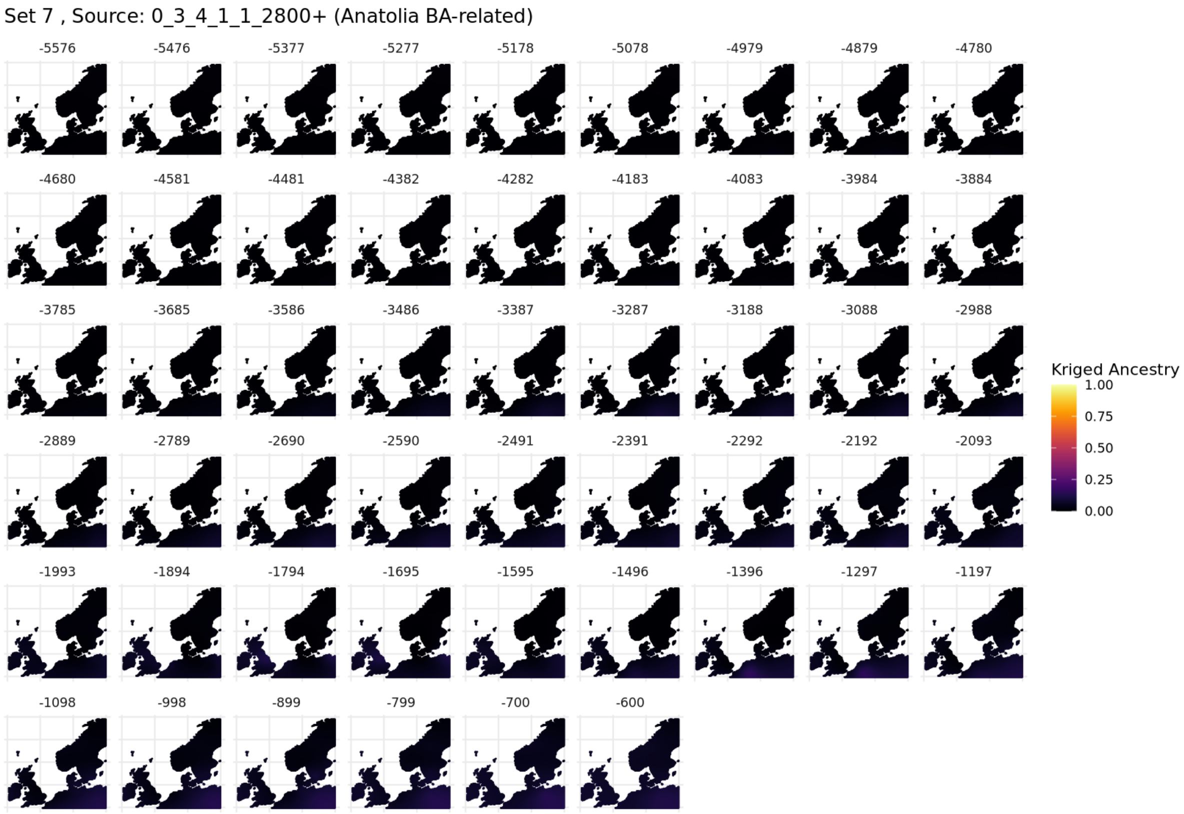

**Fig. S5.132. Spatiotemporal kriging results for Set 7, 'North / West Europe' parameters.**

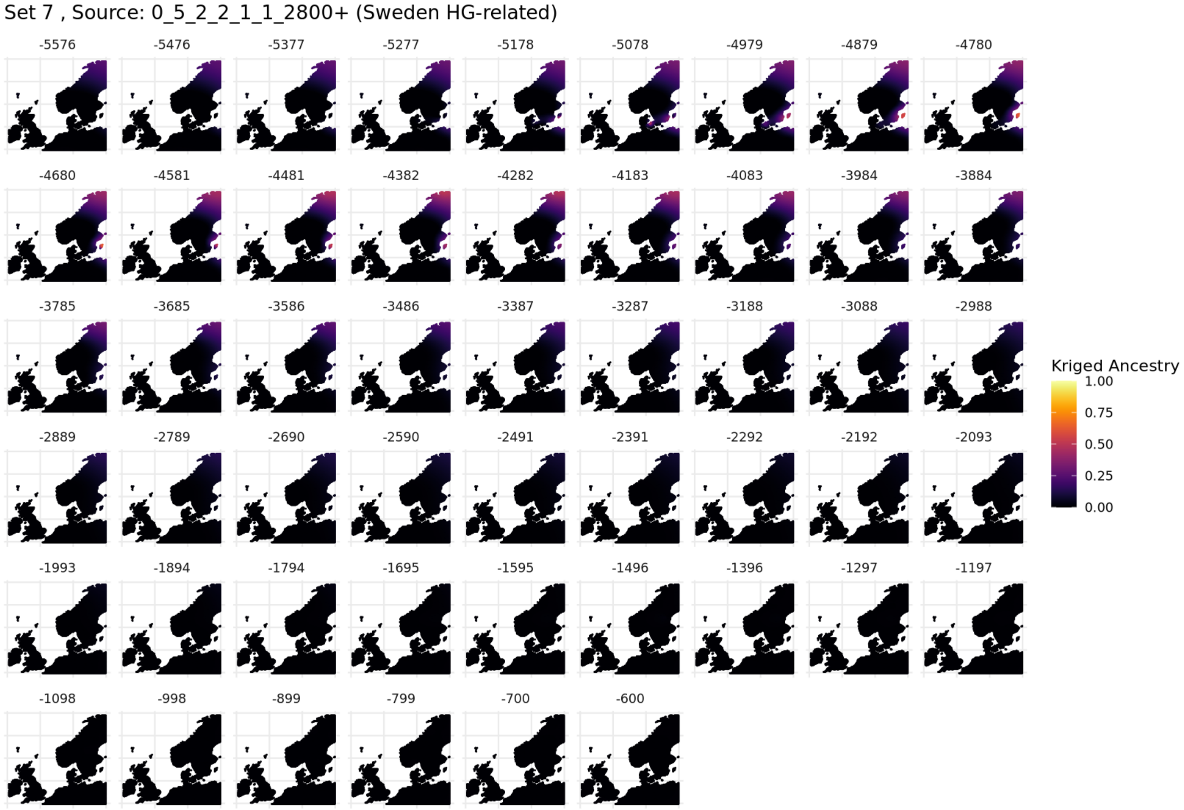

**Fig. S5.133. Spatiotemporal kriging results for Set 7, ‘North / West Europe’ parameters.**

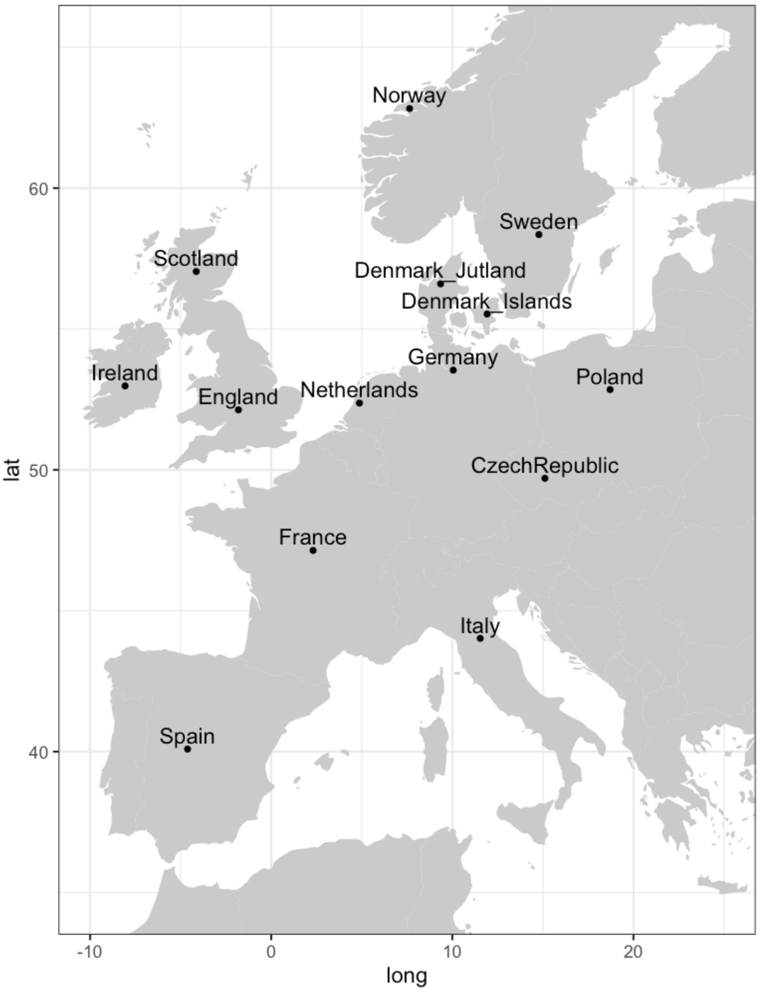

**Fig. S5.134. Coordinates used for 'country' points in spatiotemporal kriging timelines.**

#### S5.3.9. Comparisons of IBD Mixture Modelling and ADMIXTURE

To determine whether similar patterns could be captured using traditional methods, we ran unsupervised ADMIXTURE (v1.3.0) on the 2211 individuals falling within the three major clusters on Bronze Age Europe: the Corded Ware (North) 0_1_x, Bell Beaker (0_2_1) and Corded Ware (East) (0_2_2_2_x). Only transversion sites were considered, resulting in a total of 218,680 sites.

When plotting the proportions at K2 against the IBD Mixture Modelling Set 6 (Fig. S5.135), a clear correspondence between K1 and proportions modelled with Bell Beaker-related ancestry (0_2_1_4_5_C_2800+,), and K2 and the Corded Ware-related ancestry (0_2_2_2_3_C_2800+). There is a more subtle correspondence between K1 and European Farming ancestry (0_3_3_1_2800+), and K2 and Globular Amphora-related ancestry, consistent with our results.

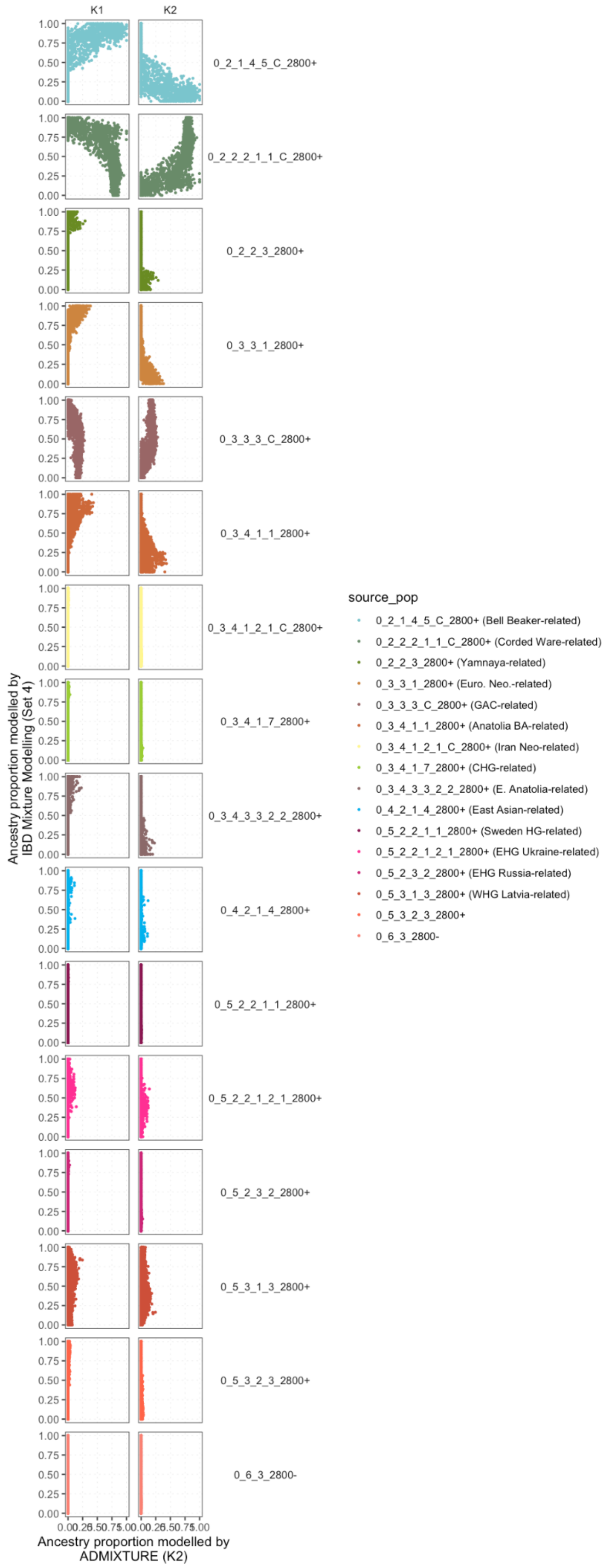

**Fig. S5.135. Mixture modelling results for Admixture results at K2, and IBD Mixture Modelling Results (set 4).**

When plotting the proportions at K7 against the IBD Mixture Modelling Set 7 (Fig. S5.136), a clear correspondence between the following is apparent:

- K1 and the Yamnaya-related ancestry
    - 0_2_2_3_2800+ (Yamnaya-related)
- K2 and farming-related ancestries
    - 0_3_3_1_2800+ (Euro. Neo.-related),
    - 0_3_4_1_1_2800+ (Anatolia BA-related),
    - 0_3_4_3_3_2_2_2800+ (E. Anatolia-related)
- K3 and the Northern German-related ancestry
    - 0_1_3_2800- (N. German IA-related)
- K4 and the Eastern Scandinavian BA enriched Swedish and Danish Isles IA-related ancestries
    - 0_1_2_1_2_C_2800- (Swedish IA-related,
    - 0_1_2_2_3_C_2800- (Danish Isles IA-related)
- K5 and the Bell Beaker-related ancestry
    - 0_2_1_4_5_C_2800+ (Bell Beaker-related), and
- K7 and the Baltic-related ancestry is visible (Fig. X).
    - 0_1_1_1_2_1_1_2800 (Baltic-related)"

For K6, the signal is more subtle, but appears linked to

0_1_2_3_C_2800- (Jutland IA-related) and

0_2_1_2_5_1_C_2800+ (E.N.S BA-related)

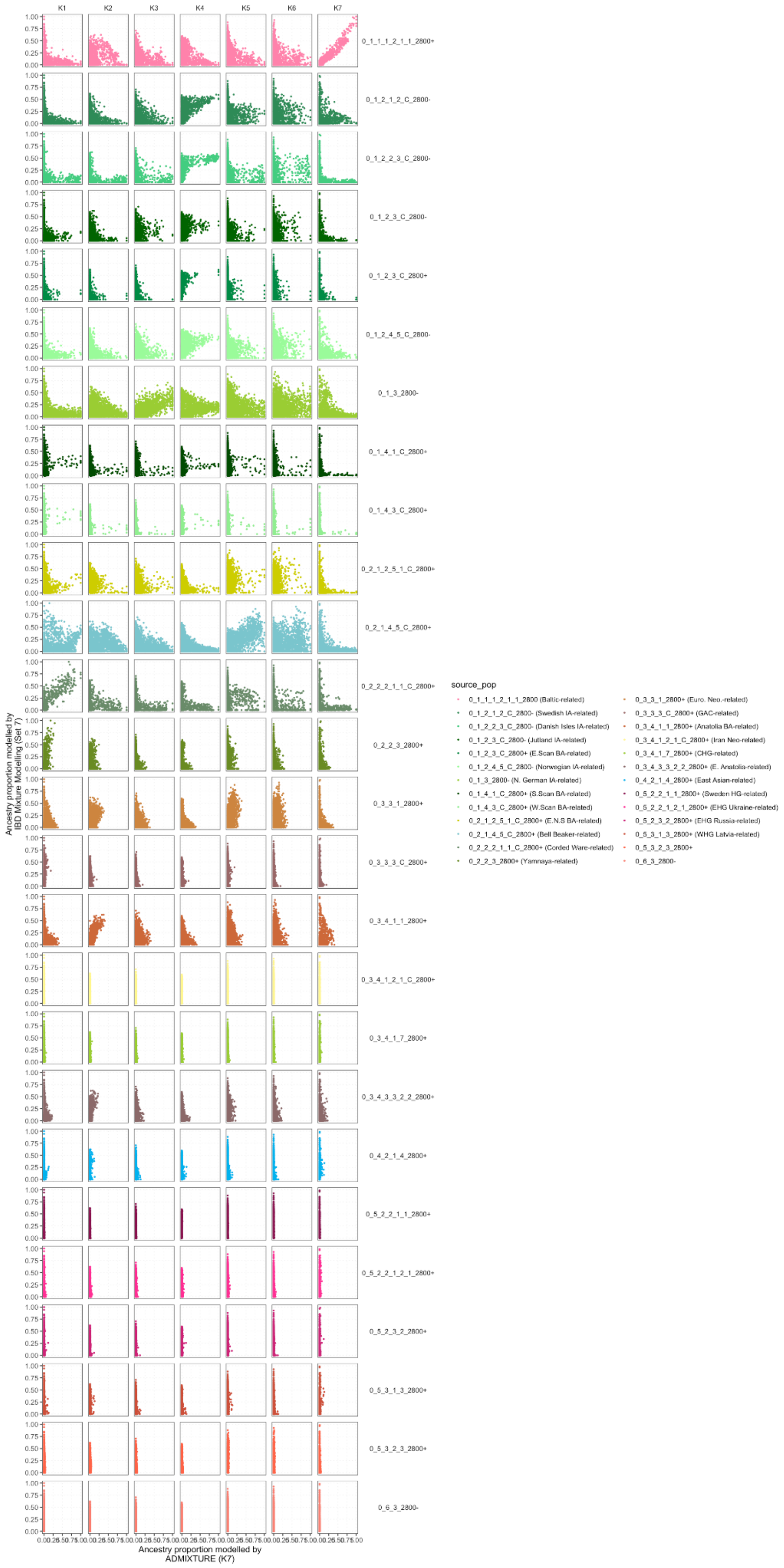

**Fig. S5.136. Mixture modelling results for Admixture results at K7, and IBD Mixture Modelling Results (set 7).**

Some of the best fitting results can be seen in **Fig. S5.137**.

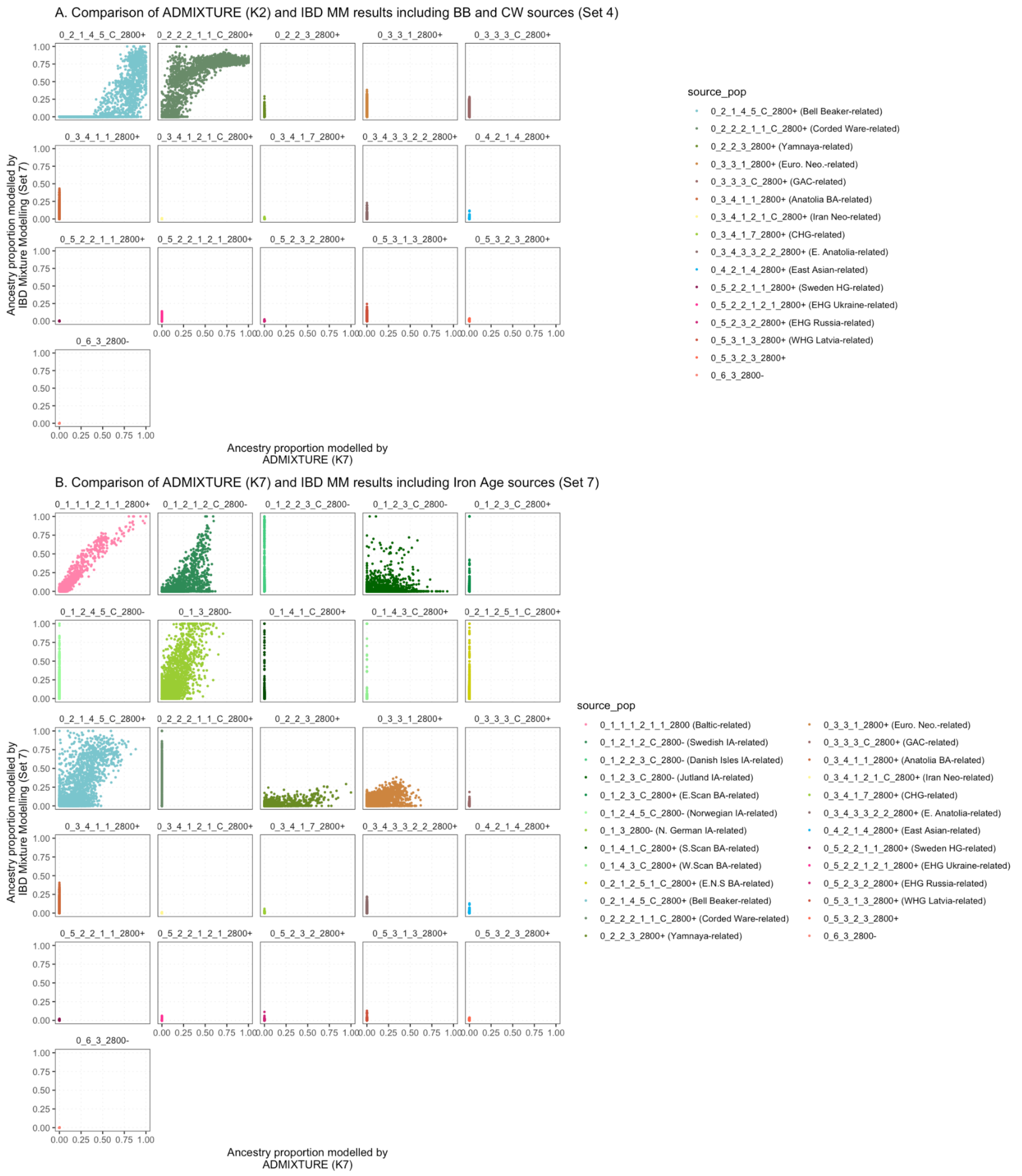

**Fig. S5.137. A comparison of (A) the IBD Mixture Modelling results (Set 4) and unsupervised admixture at K2 and (B) the IBD Mixture Modelling results (Set 7) and unsupervised admixture at K7.**

For the ADMIXTURE set at K2 and IBD Mixture Modelling Set 4 (including the Corded Ware / Bell Beaker-related sources), the correspondence can be visualised, for example, with the transition on the British Isles, coinciding with the arrival of Saxons[33] (Fig. S5.138).

For the ADMIXTURE set at K7 and IBD Mixture Modelling, showing most clearly the associations for Baltic Bronze Age, Swedish IA, N.German IA and Bell Beaker ancestries, the results show similar transitions. On the British Isles, the Saxon transition brings the widespread presence of Northern German and Swedish IA-associated ancestry (Fig. S5.138). The distinction between Northern German and Swedish IA-associated ancestry is more apparent on the Danish Isles and in Norway in the Iron Age to Viking Period transition. On the Danish Isles, we see the decrease of local Swedish IA-associated ancestry, to the widespread appearance of N.German associated ancestry, in support of migrations into Denmark (Fig. S5.139). In Norway, the Viking individuals that are modelled similarly to local Iron Age individuals continue to show only very small proportions of the N.German associated ancestry (Fig. S5.139). The distinction between the Swedish IA ancestry and the Baltic Bronze Age ancestry is clearly apparent in Poland, with the transition from the East Germanic Weilbark period to the Slavic Middle Ages (Fig. S5.140). The results for all samples can be found ordered by country (Fig. S5.141) and by cluster (Fig. S5.142).

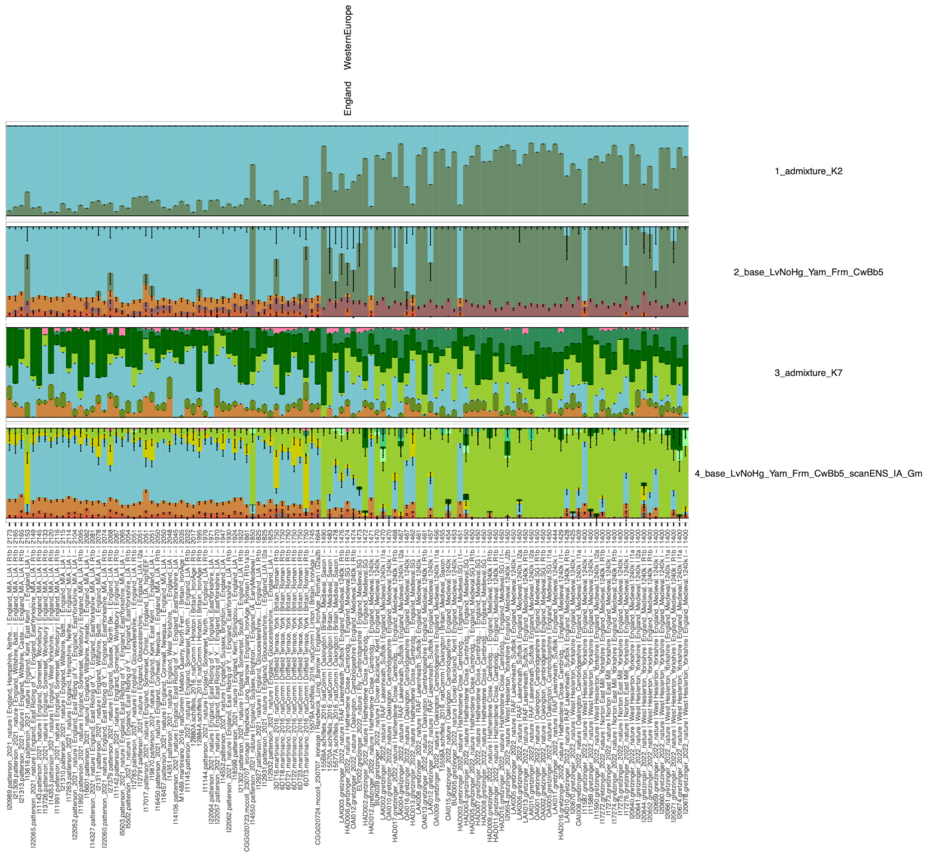

**Fig. S5.138. Mixture Modelling and Admixture results for the British Isles Saxon-associated transition.** The results for all samples can be found ordered by country (Fig. S5.141) and by cluster (Fig. S5.142).

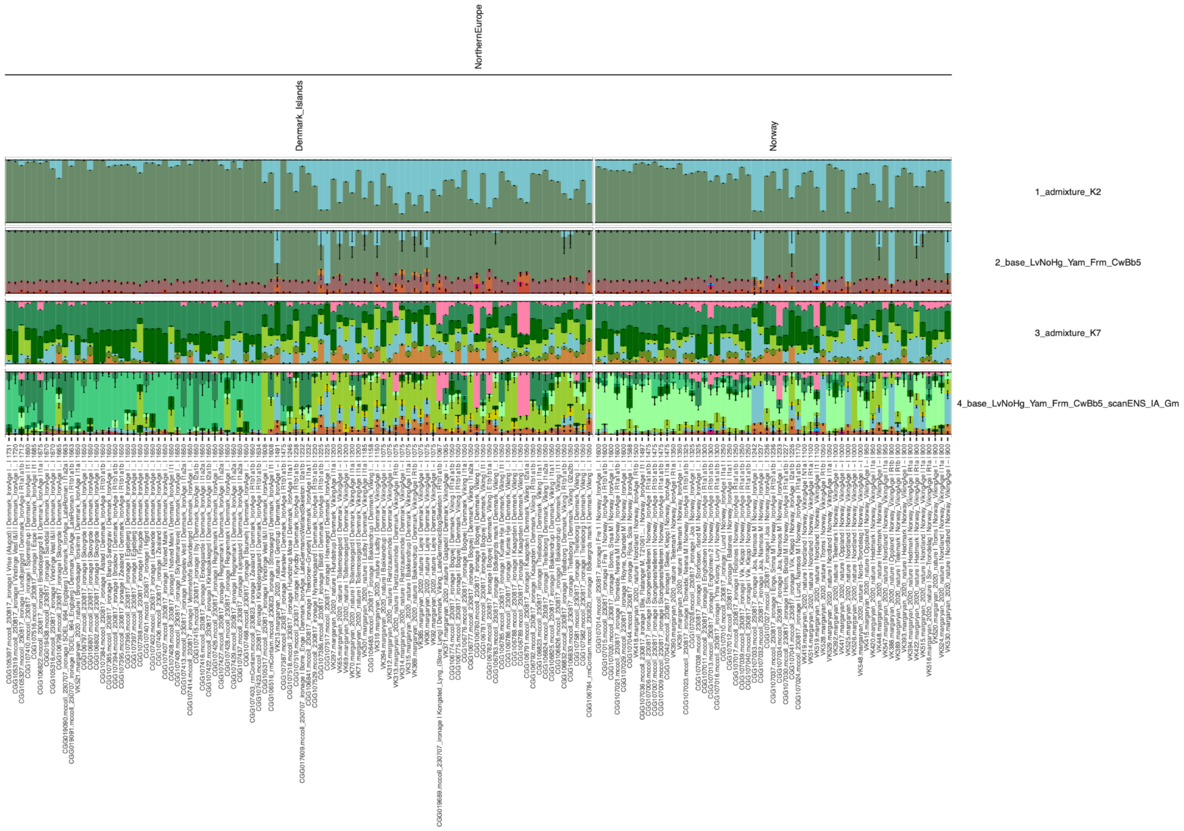

**Fig. S5.139. Mixture Modelling and Admixture results for the Iron Age to Viking Period transition on the Danish Isles and in Norway.** The results for all samples can be found ordered by country (Fig. S5.141) and by cluster (Fig. S5.142).

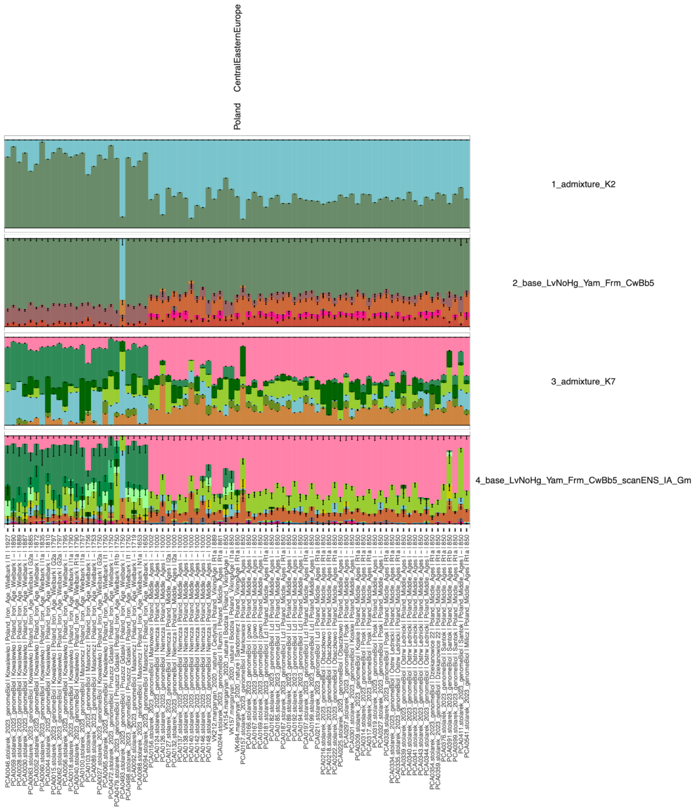

**Fig. S5.140. Mixture Modelling and Admixture results for the Iron Age to Medieval transition in Poland.** The results for all samples can be found ordered by country (Fig. S5.141) and by cluster (Fig. S5.142).

---

FIGURE INCLUDED SEPARATELY

---

**Fig. S5.141. Admixture and IBD Modelling results, ordered by country**

---

FIGURE INCLUDED SEPARATELY

---

**Fig. S5.142. Admixture and IBD Modelling results, ordered by cluster**

1697

1698

#### S5.4. Analysis of uniparental markers

##### S5.4.1. Mitochondrial DNA analysis

Mitogenomes of the newly sequenced individuals were reconstructed using bcftools[198] call and mpileup from reads mapped to rCRS from which we subsequently classified the haplogroups using haplogrep[100]. We then aligned the sequences with mafft[199] and generated a phylogenetic tree with the maximum likelihood (ML)-based tree inference tool raxML[200] using under the GTR+I+G4 model with the options [--all –bs-trees 100 ]. The analysis was restricted to only the coding region ranging from 577 to 16,023 base pairs (bp) (rCRS coordinates).

The phylogenetic analysis and classification of haplogroups suggests that the introduction of, eg., Farmer-related haplogroups such as H, V, T, and J were already widespread across Europe during the Iron Age. We furthermore identify U-haplogroups, which have formerly been commonly observed in Mesolithic Hunter-Gatherers, to be frequently observed among the sampled individuals, suggesting either continuation or a reintroduction of Hunter-Gatherer-related haplogroups into Scandinavia. We see evidence of a broad level of continuity between the global IBD clusters for the European individuals from the period spanning from Bronze Age to Iron Age, and find farmer associated haplogroups such as haplogroup H to be predominantly observed among Western and Northern European populations, suggesting that the migrations occurring in this period might not have caused complete replacement of mitochondrial lineages.

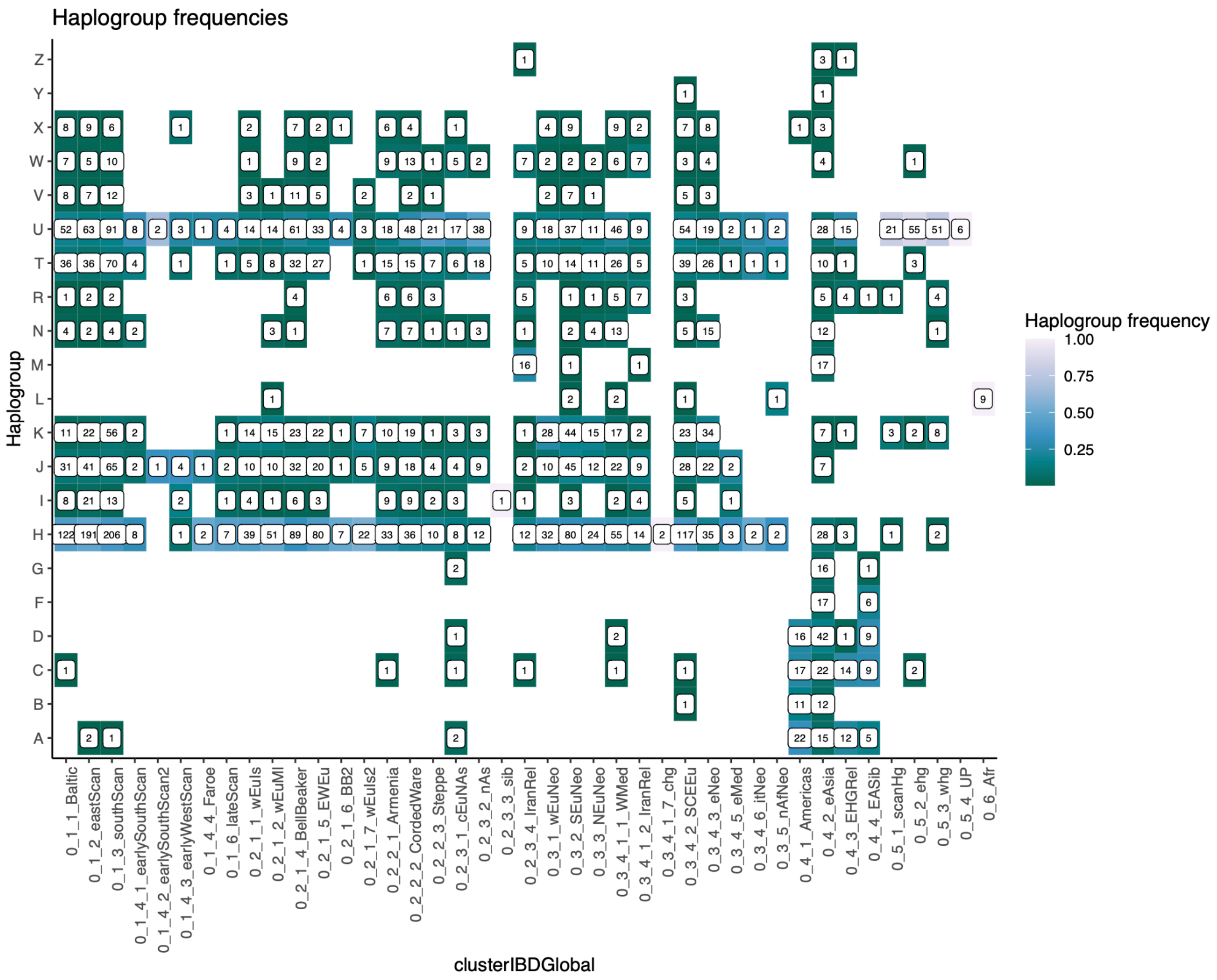

**Fig. S5.143. Haplogroup frequencies across Global IBD clusters.**

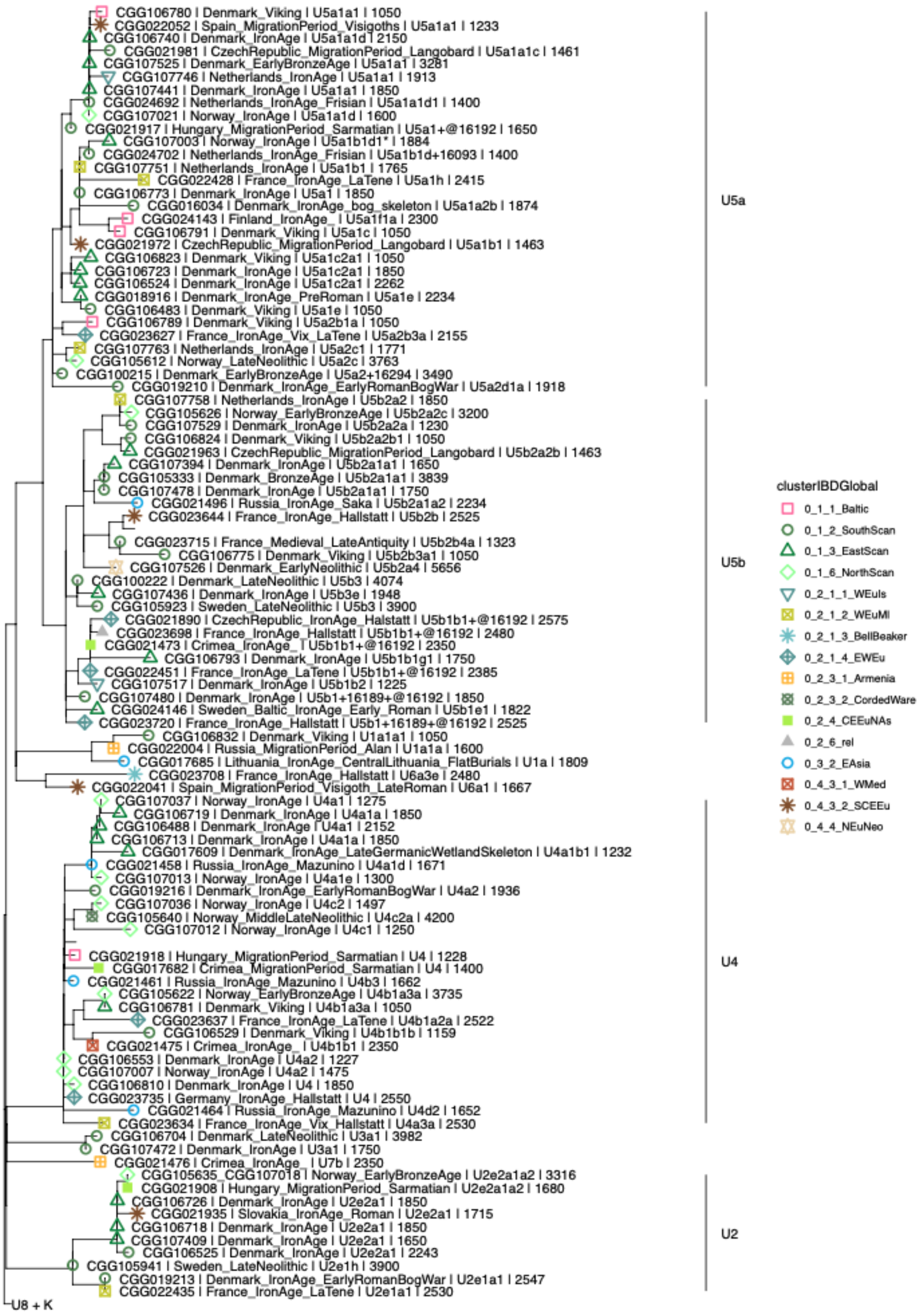

**Fig. S5.144. Maximum likelihood of U haplogroup mitogenomes.**

1727

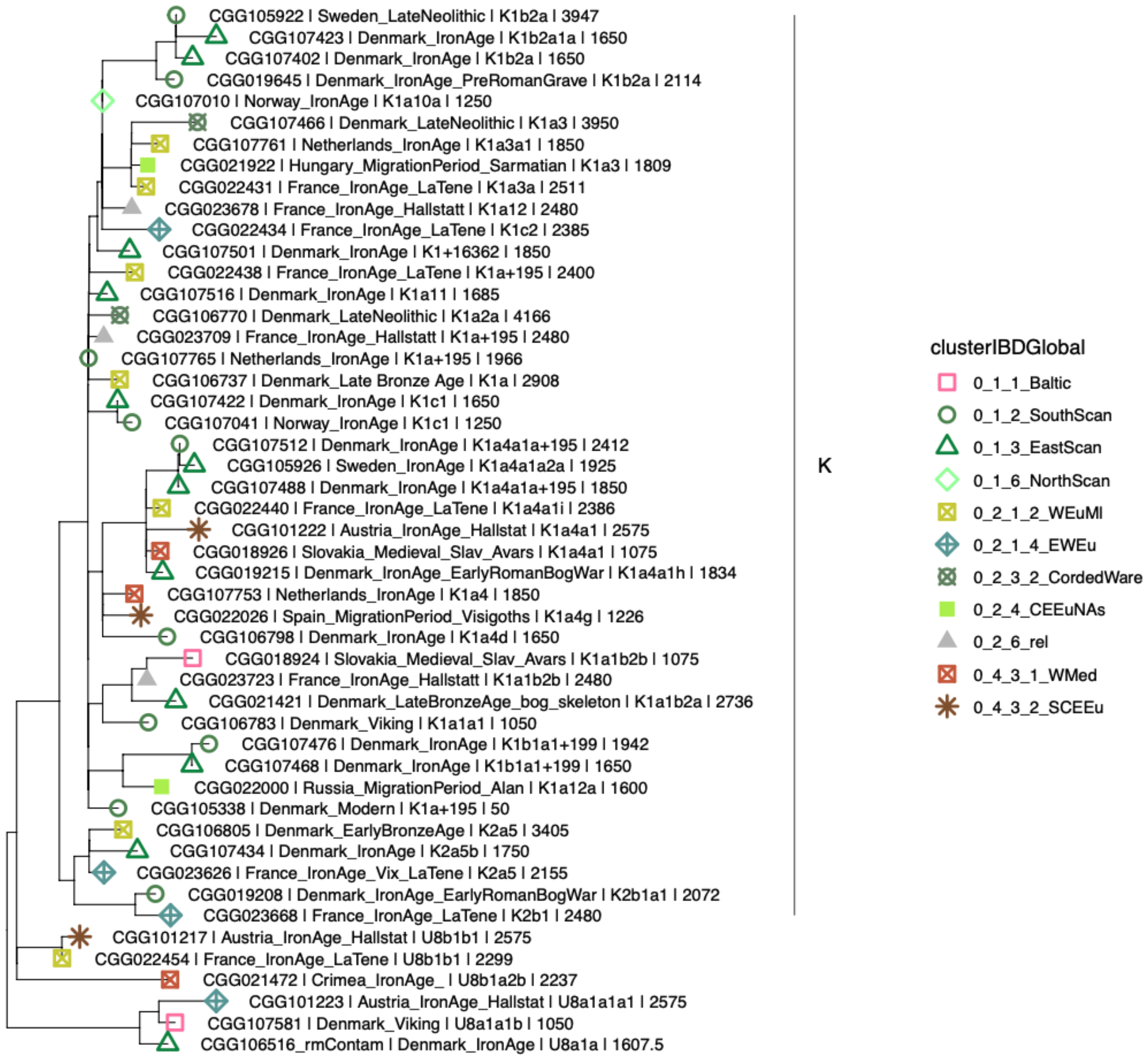

1728 **Fig. S5.145. Maximum likelihood of K haplogroup mitogenomes.**

1729

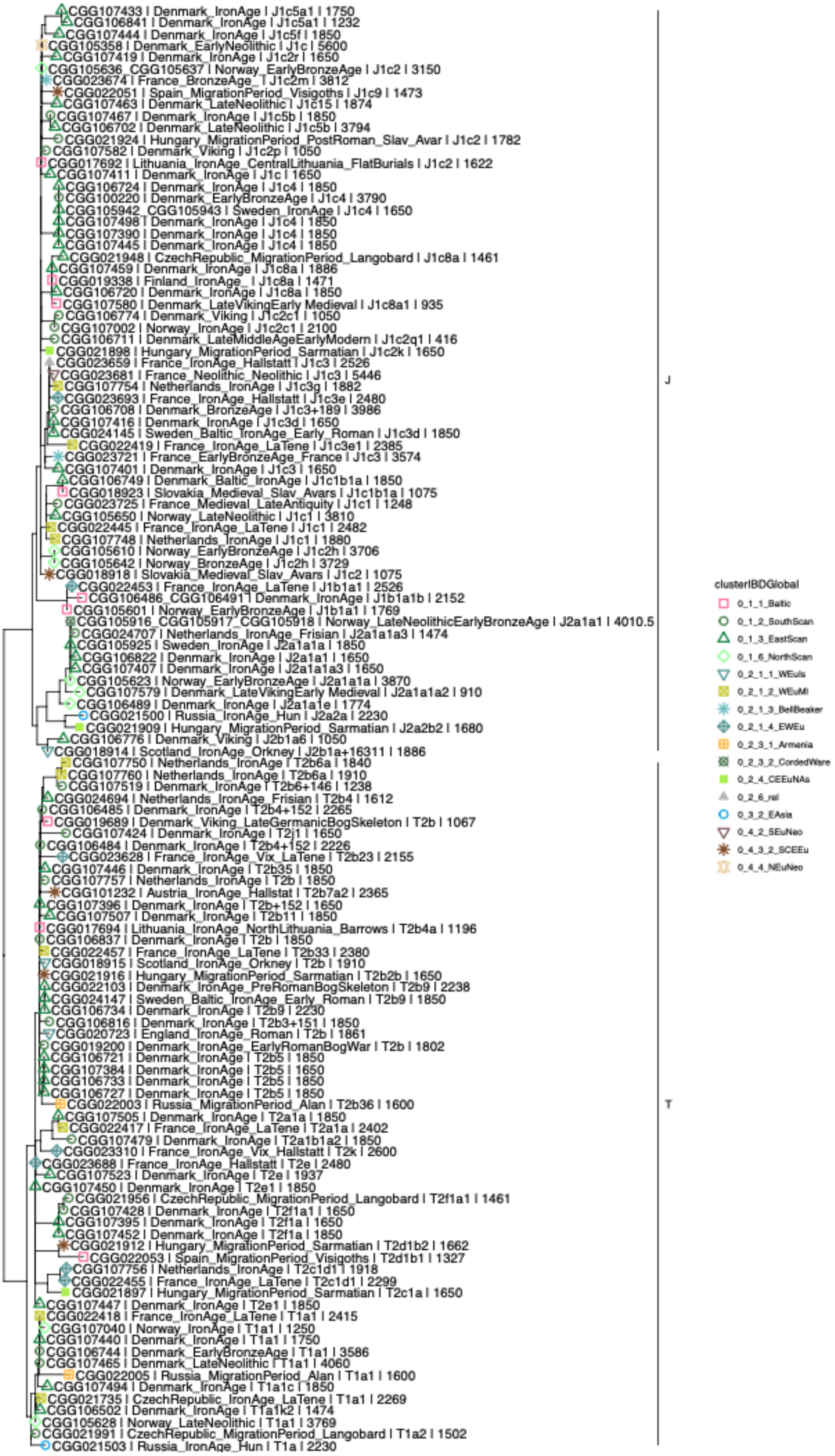
CGG107433 | Denmark_IronAge | J1c5a1 | 1750
CGG106841 | Denmark_IronAge | J1c5a1 | 1232
CGG107444 | Denmark_IronAge | J1c5f | 1850
CGG105358 | Denmark_EarlyNeolithic | J1c | 5600
CGG107419 | Denmark_IronAge | J1c2r | 1650
CGG105636_CGG105637 | Norway_EarlyBronzeAge | J1c2 | 3150
CGG023674 | France_BronzeAge_ | J1c2m | 3812
CGG022051 | Spain_MigrationPeriod_Visigoths | J1c9 | 1473
CGG107463 | Denmark_LateNeolithic | J1c15 | 1874
CGG107467 | Denmark_IronAge | J1c5b | 1850
CGG106702 | Denmark_LateNeolithic | J1c5b | 3794
CGG021924 | Hungary_MigrationPeriod_PostRoman_Slav_Avar | J1c2 | 1782
CGG107582 | Denmark_Viking | J1c2p | 1050
CGG017692 | Lithuania_IronAge_CentralLithuania_FlatBurials | J1c2 | 1622
CGG107411 | Denmark_IronAge | J1c | 1650
CGG106724 | Denmark_IronAge | J1c4 | 1850
CGG100220 | Denmark_EarlyBronzeAge | J1c4 | 3790
CGG105942_CGG105943 | Sweden_IronAge | J1c4 | 1650
CGG107498 | Denmark_IronAge | J1c4 | 1850
CGG107390 | Denmark_IronAge | J1c4 | 1850
CGG107445 | Denmark_IronAge | J1c4 | 1850
CGG021948 | CzechRepublic_MigrationPeriod_Langobard | J1c8a | 1461
CGG107459 | Denmark_IronAge | J1c8a | 1886
CGG019338 | Finland_IronAge_ | J1c8a | 1471
CGG106720 | Denmark_IronAge | J1c8a | 1850
CGG107580 | Denmark_LateVikingEarly Medieval | J1c8a1 | 935
CGG106774 | Denmark_Viking | J1c2c1 | 1050
CGG107002 | Norway_IronAge | J1c2c1 | 2100
CGG106711 | Denmark_LateMiddleAgeEarlyModern | J1c2q1 | 416
CGG021898 | Hungary_MigrationPeriod_Sarmatian | J1c2k | 1650
CGG023659 | France_IronAge_Hallstatt | J1c3 | 2526
CGG023681 | France_Neolithic_Neolithic | J1c3 | 5446
CGG107754 | Netherlands_IronAge | J1c3g | 1882
CGG023693 | France_IronAge_Hallstatt | J1c3e | 2480
CGG106708 | Denmark_BronzeAge | J1c3+189 | 3986
CGG107416 | Denmark_IronAge | J1c3d | 1650
CGG024145 | Sweden_Baltic_IronAge_Early_Roman | J1c3d | 1850
CGG022419 | France_IronAge_LaTene | J1c3e1 | 2385
CGG023721 | France_EarlyBronzeAge_France | J1c3 | 3574
CGG107401 | Denmark_IronAge | J1c3 | 1650
CGG106749 | Denmark_Baltic_IronAge | J1c1b1a | 1850
CGG018923 | Slovakia_Medieval_Slav_Avars | J1c1b1a | 1075
CGG023725 | France_Medieval_LateAntiquity | J1c1 | 1248
CGG105650 | Norway_LateNeolithic | J1c1 | 3810
CGG022445 | France_IronAge_LaTene | J1c1 | 2482
CGG107748 | Netherlands_IronAge | J1c1 | 1880
CGG105610 | Norway_EarlyBronzeAge | J1c2h | 3706
CGG105642 | Norway_BronzeAge | J1c2h | 3729
CGG018918 | Slovakia_Medieval_Slav_Avars | J1c2 | 1075
CGG022453 | France_IronAge_LaTene | J1b1a1 | 2526
CGG106486_CGG106491 | Denmark_IronAge | J1b1a1b | 2152
CGG105601 | Norway_EarlyBronzeAge | J1b1a1 | 1769
CGG105916_CGG105917_CGG105918 | Norway_LateNeolithicEarlyBronzeAge | J2a1a1 | 4010.5
CGG024707 | Netherlands_IronAge_Frisian | J2a1a1a3 | 1474
CGG105925 | Sweden_IronAge | J2a1a1a | 1850
CGG106822 | Denmark_IronAge | J2a1a1 | 1650
CGG107407 | Denmark_IronAge | J2a1a1a3 | 1650
CGG105623 | Norway_EarlyBronzeAge | J2a1a1a | 3870
CGG107579 | Denmark_LateVikingEarly Medieval | J2a1a1a2 | 910
CGG106489 | Denmark_IronAge | J2a1a1e | 1774
CGG021500 | Russia_IronAge_Hun | J2a2a | 2230
CGG021909 | Hungary_MigrationPeriod_Sarmatian | J2a2b2 | 1680
CGG106776 | Denmark_Viking | J2b1a6 | 1050
CGG018914 | Scotland_IronAge_Orkney | J2b1a+16311 | 1886
CGG107750 | Netherlands_IronAge | T2b6a | 1840
CGG107760 | Netherlands_IronAge | T2b6a | 1910
CGG107519 | Denmark_IronAge | T2b6+146 | 1238
CGG024694 | Netherlands_IronAge_Frisian | T2b4 | 1612
CGG106485 | Denmark_IronAge | T2b4+152 | 2265
CGG019689 | Denmark_Viking_LateGermanicBogSkeleton | T2b | 1067
CGG107424 | Denmark_IronAge | T2j1 | 1650
CGG106484 | Denmark_IronAge | T2b4+152 | 2226
CGG023628 | France_IronAge_Vix_LaTene | T2b23 | 2155
CGG107446 | Denmark_IronAge | T2b35 | 1850
CGG107757 | Netherlands_IronAge | T2b | 1850
CGG101232 | Austria_IronAge_Hallstat | T2b7a2 | 2365
CGG107396 | Denmark_IronAge | T2b+152 | 1650
CGG107507 | Denmark_IronAge | T2b11 | 1850
CGG017694 | Lithuania_IronAge_NorthLithuania_Barrows | T2b4a | 1196
CGG106837 | Denmark_IronAge | T2b | 1850
CGG022457 | France_IronAge_LaTene | T2b33 | 2380
CGG018915 | Scotland_IronAge_Orkney | T2b | 1910
CGG021916 | Hungary_MigrationPeriod_Sarmatian | T2b2b | 1650
CGG022103 | Denmark_IronAge_PreRomanBogSkeleton | T2b9 | 2238
CGG024147 | Sweden_Baltic_IronAge_Early_Roman | T2b9 | 1850
CGG106734 | Denmark_IronAge | T2b9 | 2230
CGG106816 | Denmark_IronAge | T2b3+151 | 1850
CGG020723 | England_IronAge_Roman | T2b | 1861
CGG019200 | Denmark_IronAge_EarlyRomanBogWar | T2b | 1802
CGG106721 | Denmark_IronAge | T2b5 | 1850
CGG107384 | Denmark_IronAge | T2b5 | 1650
CGG106733 | Denmark_IronAge | T2b5 | 1850
CGG106727 | Denmark_IronAge | T2b5 | 1850
CGG022003 | Russia_MigrationPeriod_Alan | T2b36 | 1600
CGG107505 | Denmark_IronAge | T2a1a | 1850
CGG022417 | France_IronAge_LaTene | T2a1a | 2402
CGG107479 | Denmark_IronAge | T2a1b1a2 | 1850
CGG023310 | France_IronAge_Vix_Hallstatt | T2k | 2600
CGG023688 | France_IronAge_Hallstatt | T2e | 2480
CGG107523 | Denmark_IronAge | T2e | 1937
CGG107450 | Denmark_IronAge | T2e1 | 1850
CGG021956 | CzechRepublic_MigrationPeriod_Langobard | T2f1a1 | 1461
CGG107428 | Denmark_IronAge | T2f1a1 | 1650
CGG107395 | Denmark_IronAge | T2f1a | 1650
CGG107452 | Denmark_IronAge | T2f1a | 1850
CGG021912 | Hungary_MigrationPeriod_Sarmatian | T2d1b2 | 1662
CGG022053 | Spain_MigrationPeriod_Visigoths | T2d1b1 | 1327
CGG107756 | Netherlands_IronAge | T2c1d1 | 1918
CGG022455 | France_IronAge_LaTene | T2c1d1 | 2299
CGG021897 | Hungary_MigrationPeriod_Sarmatian | T2c1a | 1650
CGG107447 | Denmark_IronAge | T2e1 | 1850
CGG022418 | France_IronAge_LaTene | T1a1 | 2415
CGG107040 | Norway_IronAge | T1a1 | 1250
CGG107440 | Denmark_IronAge | T1a1 | 1750
CGG106744 | Denmark_EarlyBronzeAge | T1a1 | 3586
CGG107465 | Denmark_LateNeolithic | T1a1 | 4060
CGG022005 | Russia_MigrationPeriod_Alan | T1a1 | 1600
CGG107494 | Denmark_IronAge | T1a1c | 1850
CGG021735 | CzechRepublic_IronAge_LaTene | T1a1 | 2269
CGG106502 | Denmark_IronAge | T1a1k2 | 1474
CGG105628 | Norway_LateNeolithic | T1a1 | 3769
CGG021991 | CzechRepublic_MigrationPeriod_Langobard | T1a2 | 1502
CGG021503 | Russia_IronAge_Hun | T1a | 2230
J
T
clusterIBDGlobal
0_1_1_Baltic
0_1_2_SouthScan
0_1_3_EastScan
0_1_6_NorthScan
0_2_1_1_WEuIs
0_2_1_2_WEuMI
0_2_1_3_BellBeaker
0_2_1_4_EWEu
0_2_3_1_Armenia
0_2_3_2_CordedWare
0_2_4_CEEuNAs
0_2_6_rel
0_3_2_EAsia
0_4_2_SEuNeo
0_4_3_2_SCEEu
0_4_4_NEuNeo

**Fig. S5.146. Maximum likelihood of J and T haplogroup mitogenomes.**

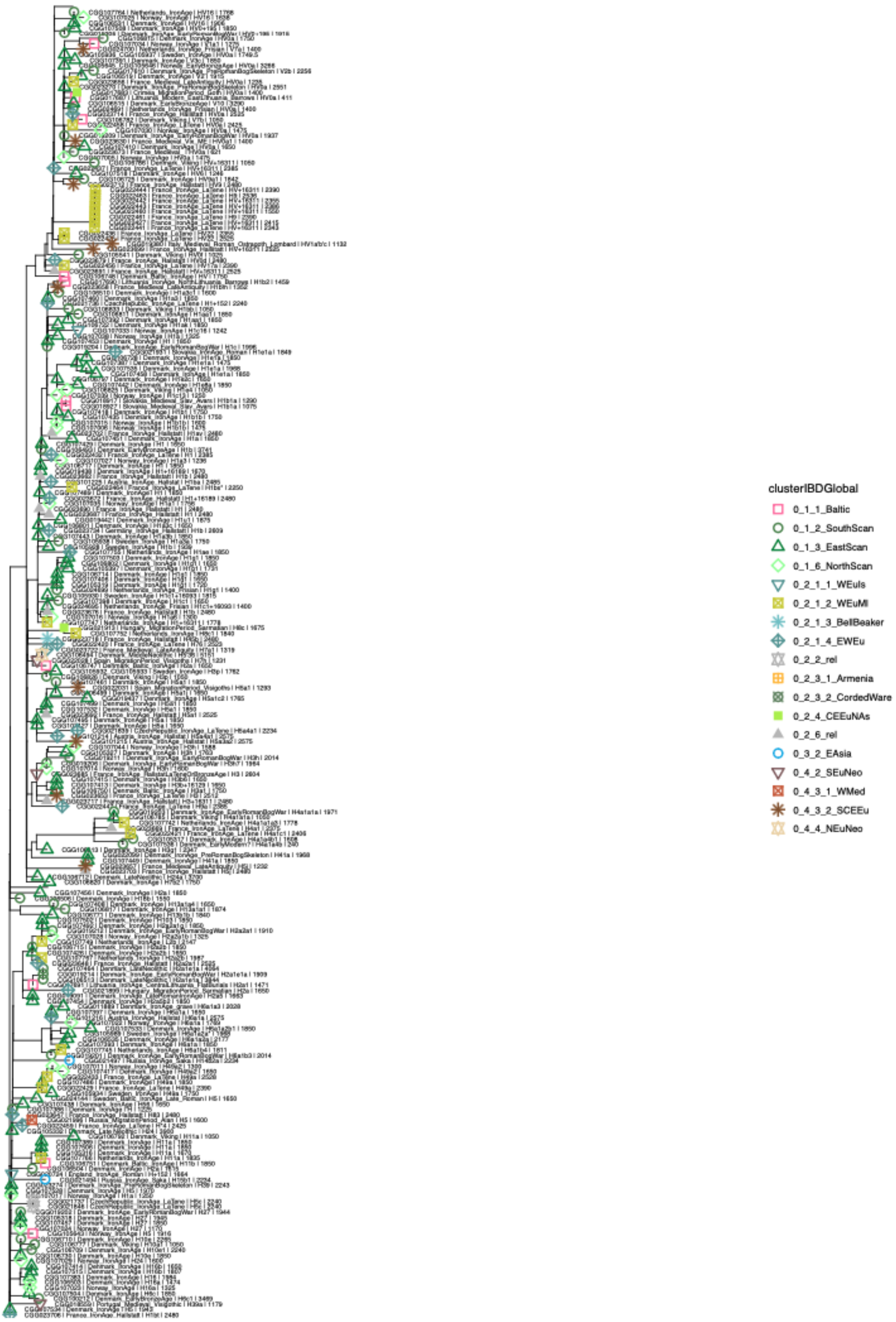

**Fig. S5.147. Maximum likelihood of H and V haplogroup mitogenomes.**

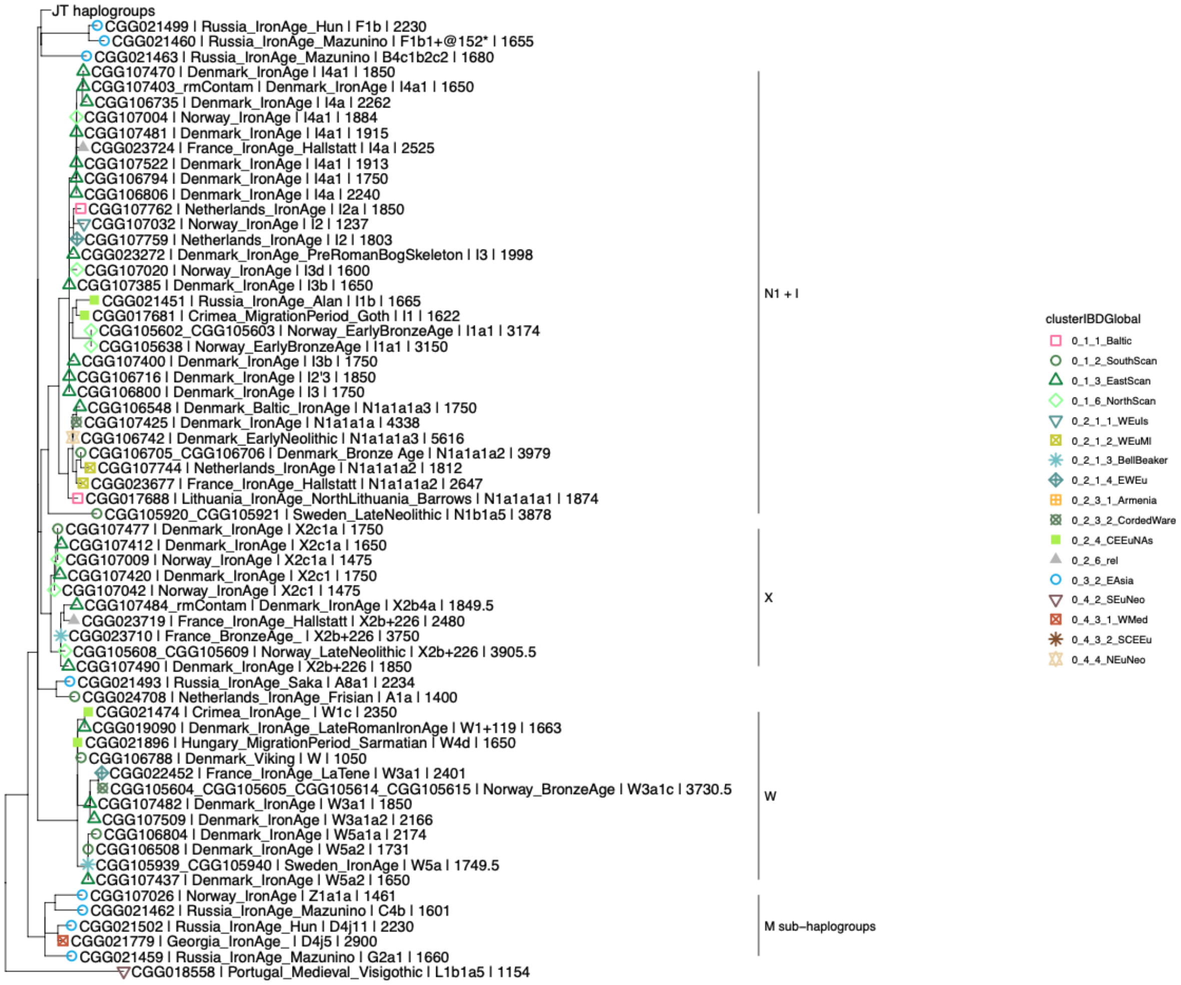

**Fig. S5.148. Maximum likelihood of N, X, W, and M haplogroup mitogenomes.**

#### S5.4.2. Y-chromosome analysis

The Y-chromosome is the largest uniparentally inherited haplotypic block of the human genome, and it has been correlated with many known events in human history[201–203]. Due to its larger size, lower effective population size and because it traces primarily the movements of males, it has been shown to be more sensitive to diagnose past demographic fluctuations, such as migration and depopulation, than mitochondrial DNA[204] . Initially, most of the developments in the Y-chromosome phylogeny were done by large public academic projects of worldwide diversity[202,205–208], but today most of the recent advances were largely first described in forensic or community-based citizen-scientist efforts using private genetic data[203], and often privately owned databases are far more complete than academic ones. While those are immensely biased towards Western Eurasian ancestry (particularly Northwest Europe), this is also true of most genetic data overall[209,210]. Ancient DNA poses particular problems to the study of Y-chromosome, such as pervasive missing data, relatively lower coverage and confounding factors, as miscoding lesions ('damage') and potential contamination[211,212]. Many approaches have been used to address those issues, such as giving different weights to transition and transversion variants[163], but in cases where dense sampling in the haplogroup exists and depth of coverage is not a limiting factor for ancient data, phylogenetic placement algorithms have proven to be the most effective[28,213,214].

**Materials and methods**

We used bcftools[215] mpileup and call function to call genotypes specifying chromosome ploidy in the 10 Mb accessible region of the Y[216] and excluded indels, triallelic positions and genotypes not called in more than 95% of the population of non-clonal reads. Because haplogroup-defining SNPs are defined from the root of the Y-chromosome phylogeny but the Y-chromosome reference is a mosaic of at least two donors of common European haplogroups (R1b and G2a – see also[217]), variants should be considered in terms of ancestral/derived instead of reference/alternate, particularly for individuals belonging to those two haplogroups. We therefore proceeded to call haplogroups by matching ancestral and derived calls to ISOGG 2019-2020 database, using an in-house script which generates haplogroup paths in root-to-tip fashion, sorted by number of supported variants seen, while also distinguishing C>T and G>A variants in the forward and reverse strand, which could arise from damage. To further incorporate results not listed in ISOGG and make use of the wealth of customer-based private datasets, we have also annotated VCFs using bcftools to all biallelic single nucleotide point mutation variants listed in yBrowse (https://ybrowse.org/gbrowse2/gff/) and matched those to the privately-owned Yfull tree (https://github.com/YFullTeam/YTree/blob/master/ytree/tree_10.07.0.zip). To visualise the results and confirm haplogroup placement of low-coverage samples, we used PathPhynder[214], which also takes into consideration the presence of ancestral variants to create a parsimonious phylogenetic path for the samples. Y-chromosome results from the new samples have been tabulated in Table S1.1. As we intended to match Y-chromosome haplogroups with the autosome IBD clusters, we repeated the process above with samples from the literature, but restricting the subhaplogroup definition to informative clades discussed in the text.

**Discussion**

Almost all Scandinavians older than 4500 BP belong to haplogroup I2-P215 (52 out of 57). Only starting at 4500 BP, we see the arrival of lineages more common today in the region, such as I1-M253, R1a1a1 (R1a-M417) and R1b1a1b1a (R1b-L11).

Europeans after 5000 BP were clustered in four distinct groups, with the majority of individuals from Scandinavia belonging to the cluster associated with Corded Ware (North) ancestry (0_1 CordedWareNorth). Notably, fifteen Scandinavian individuals dated between 4600 and 3700 BP instead were placed along other with individuals of similar age in a more eastern group (0_2_2_2 CordedWareEast). The haplogroup frequencies in those is similar to the main cluster, with R1a1a1 (R1a-M417) and R1b1a1b1a (R1b-L11) representing together most of the male lineages in both cases.

In agreement with the finding that those four main clusters are genetically related, when investigating the Y-chromosome lineages of the older members of those groupings, we find them to be dominated by closely related, but clearly distinct, lineages. This can be seen by comparing individuals most associated with early Bell Beakers (0_2_1_4 BellBeaker) and Yamnaya (0_2_2_3 Steppe) clusters. While both are dominated by R1b1a1b (R1b-M269) lineages, the prevalent lineage among the Yamnaya – R1b1a1b1b (R1b-Z2103), found in 17 out of 28 males – is much more common in the Caucasus and Eastern Europe, and is a sister lineage (therefore not a direct source) of the present-day dominant Western European lineage R1b1a1b1a (R1b-L11) (Fig. S5.149 and S5.150). Instead, almost the totality of males in the Bell Beaker cluster (133 out of 153) are downstream of this haplogroup, at R1b1a1b1a1a2 (R1b-P312), agreeing with modelling results of Bell Beakers being a better source for modern populations than Yamnaya.

When investigating the main subgroups in the main cluster containing Scandinavian individuals (0_1 CordedWareNorth), we found no lineage occurring more than five times to be exclusive to either cluster, which reiterates the genetic similarity of those. However, some lineages carry clear geographical correlations when using a time transect, as those clusters are shown with autosomes to have been admixed after 2800 BP.

Haplogroup I1a-DF29 first appears in Scandinavia in the Bronze Age, and its vast number of local descendant lineages seem to indicate an in situ diversification and point of origin (Fig. S5.151). Haplogroup I1a2a (I1a-Z59) is most the common variant of I1a-DF29 among both 0_1_3 SouthScan and 0_1_2 EastScan clusters, but when restricting to individuals older than 2800 BP, all four are in the Eastern Scandinavian cluster (Fig. S5.152). This likely indicates that its presence in the Southern Scandinavian cluster is associated with the later mixture from an Eastern Scandinavian source.

The four more common subhaplogroups of R1a1a1 (R1a-M417) in Northern Europe also show frequency differences among those four clusters. Almost all R1a1a1b1a1 (R1a-M458) and R1a1a1b1a2 (R1a-Z280) individuals belong to the 0_1_1 Baltic cluster which, together with the presence of E1b1b1a1b1 (E1b-L618) individuals, represent a Central and Eastern European affinity of this cluster not seen in the remaining Corded Ware (North) subgroups.

Conversely, haplogroups R1a1a1a1 (R1a-L664) and R1a1a1b1a3a (R1b-Z284) are more frequent in three mainly Scandinavian clusters, with the majority of R1a1a1b1a3a (R1b-Z284) from the whole dataset deriving from 0_1_2_4_x cluster, who tend to be modelled with high proportions of Norwegian Iron Age ancestry (Fig. S5.29), including Viking age individuals from Ireland, Britain, Iceland and Greenland. The oldest evidence of this haplogroup takes place in the Corded Ware (East) cluster (0_2_2_2 CordedWareEast) at around 4500 BP in Sweden. Within Scandinavia, the earliest detection in a Corded Ware (North) cluster is within the Western Scandinavian BA cluster (0_1_4_3_C_2800+), whose ancestry is modelled to a small degree in many Norwegian Iron Age individuals (Fig. S5.153). Therefore, while we can ultimately trace this subhaplogroup to individuals related to eastern Corded Ware ancestry, it likely spread later within Northern Europe, through mixture with individuals carrying small amounts of Western Scandinavian Bronze Age-associated ancestry.

Downstream of R1b1a1b1a (R1b-L11), haplogroup R1b1a1b1a1a1 (R1b-U106) has been previously argued to be related to the expansion of the Germanic languages, due to its high frequency in places where those languages are spoken today (Fig. S5.154). We found most of the individuals of the dataset positive for R1b-U106 to belong to two different downstream sublineages, which have starkly distinct distributions, particularly in the early Iron Age. R1b1a1b1a1a1c (R1b-Z19) is found almost exclusively in Northern Europe (with the only exception being a Langobard from Hungary), and likely represents a local variant of R1b-U106 (Fig. S5.155).

Instead, its sister lineage, R1b1a1b1a1a1b (R1b-S263), is absent in Scandinavia before the Iron Age (Fig. S5.156), where it spreads, likely through an Eastern North Sea source, and becomes dominant in Southern Scandinavia during the Iron Age, before spreading through Northern Europe. This pattern strongly matches the one seen using autosomes, which detect gene flow back into Scandinavia related to the spread of Germanic languages. Another potential signal of this migration is the increase in frequency of the R1b-U106 sister lineage, R1b1a1b1a1a2 (R1b-P312), which has a more continental distribution, and is almost absent in Scandinavia before 2000 BP.

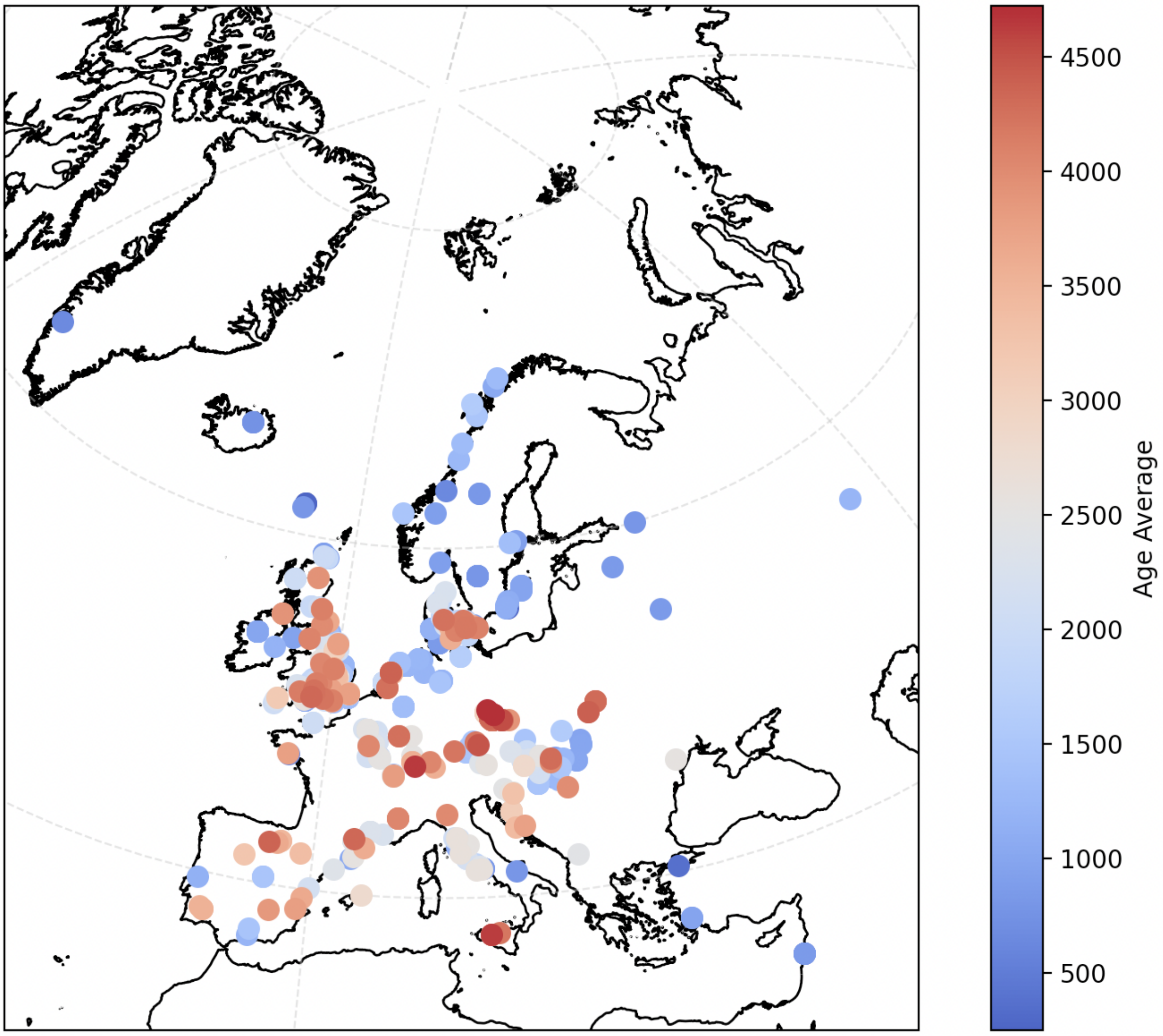

**Fig. S5.149. All individuals from the dataset belonging to haplogroup R1b1a1b1a (R1b-L11), coloured by age BP.**

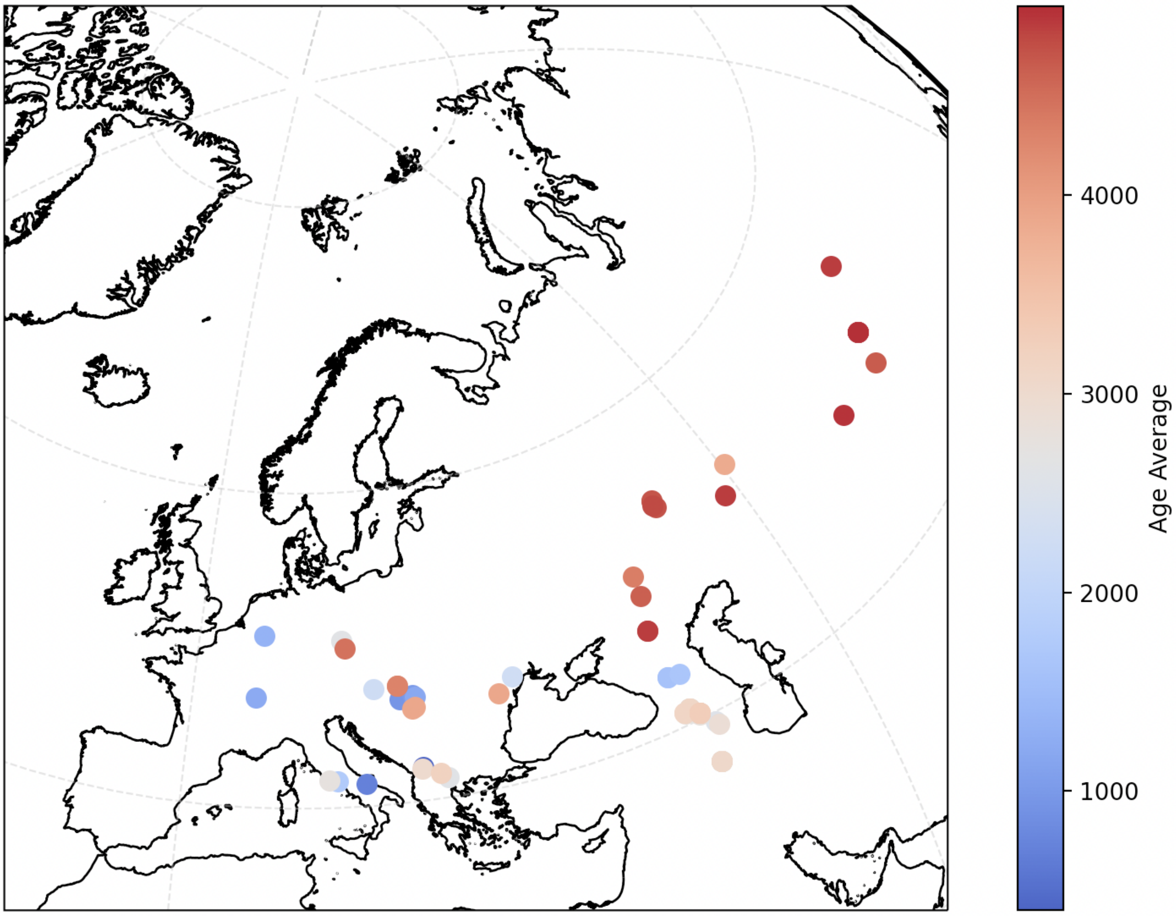

**Fig. S5.150. All individuals from the dataset belonging to haplogroup R1b1a1b1b (R1b-LZ2103), coloured by age BP.**

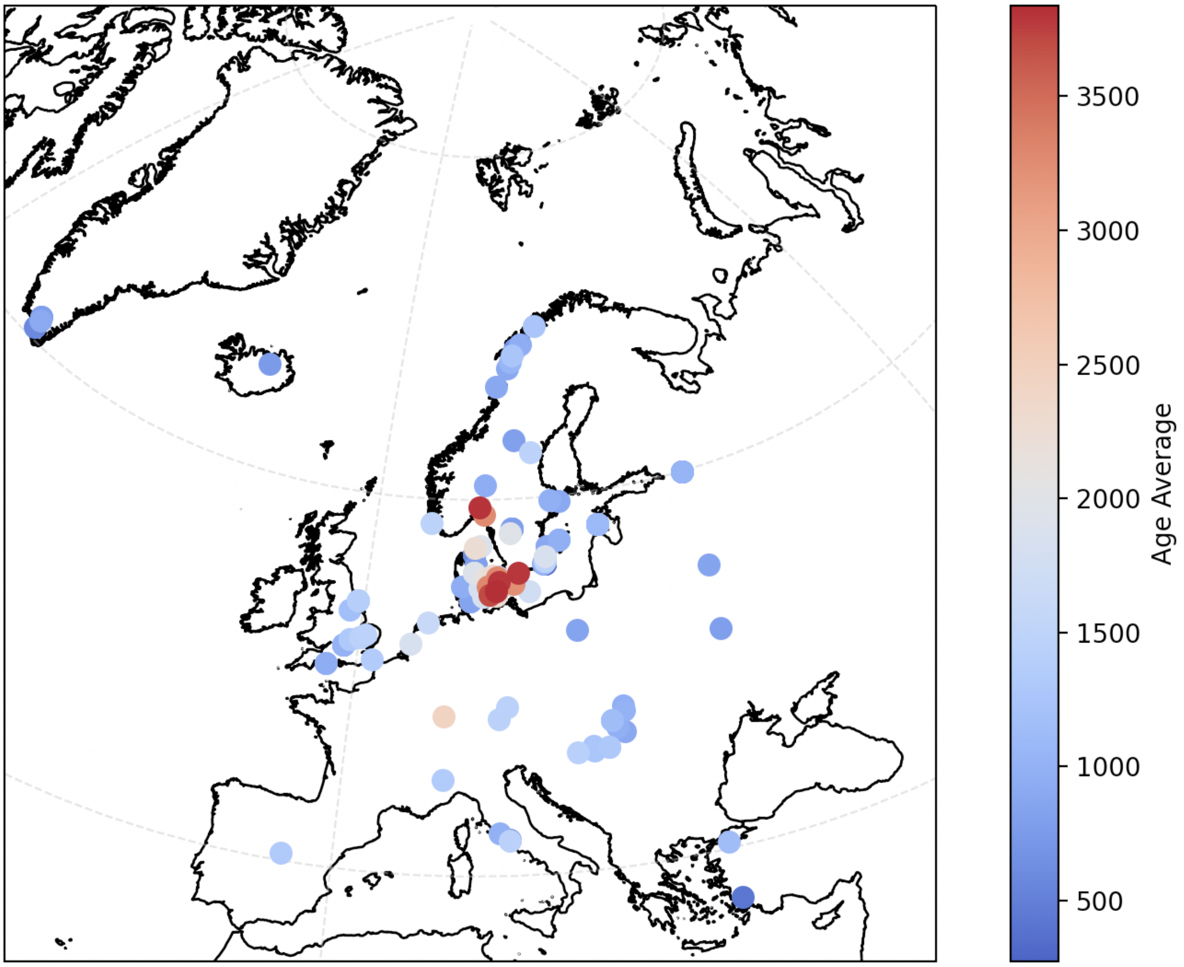

**Fig. S5.151. All individuals carrying haplogroup I1a-DF29 from the dataset, coloured by age BP.**

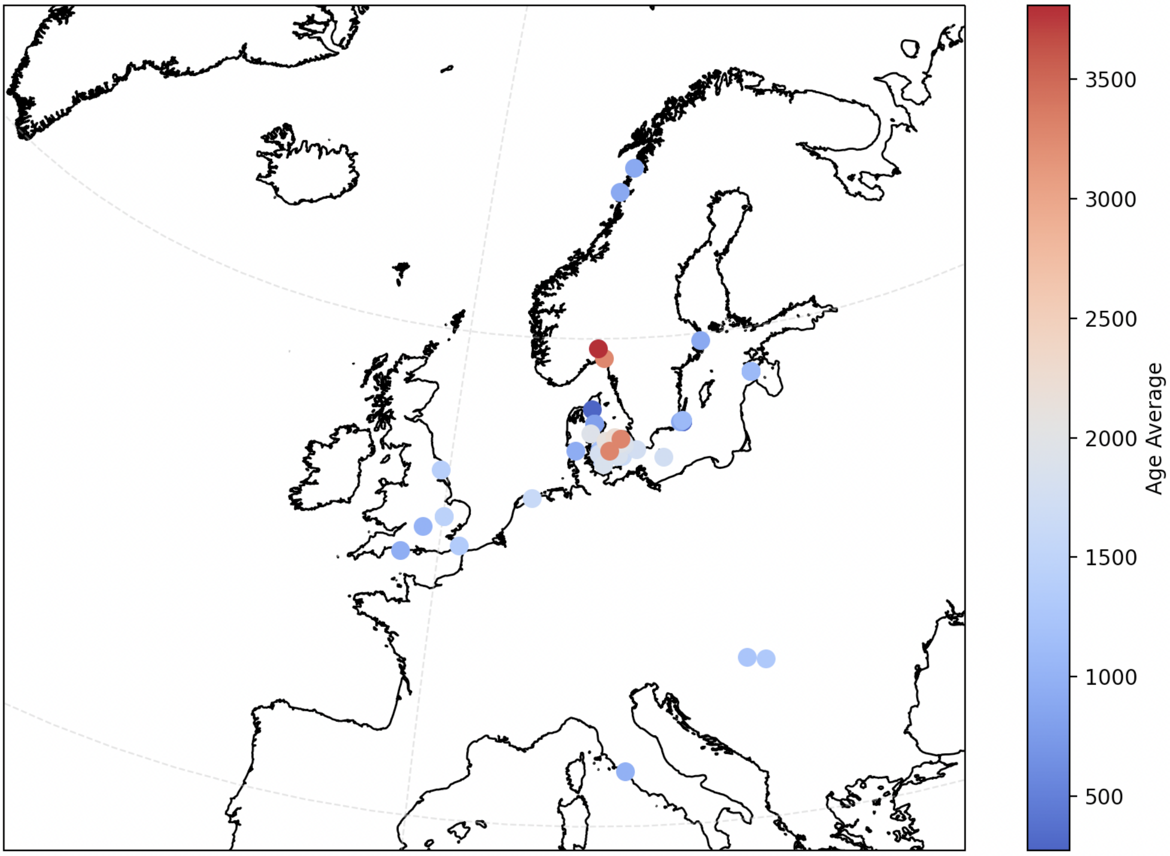

**Fig. S5.152. All individuals carrying haplogroup I1a1a (I1a-Z59) from the dataset, coloured by age BP.**

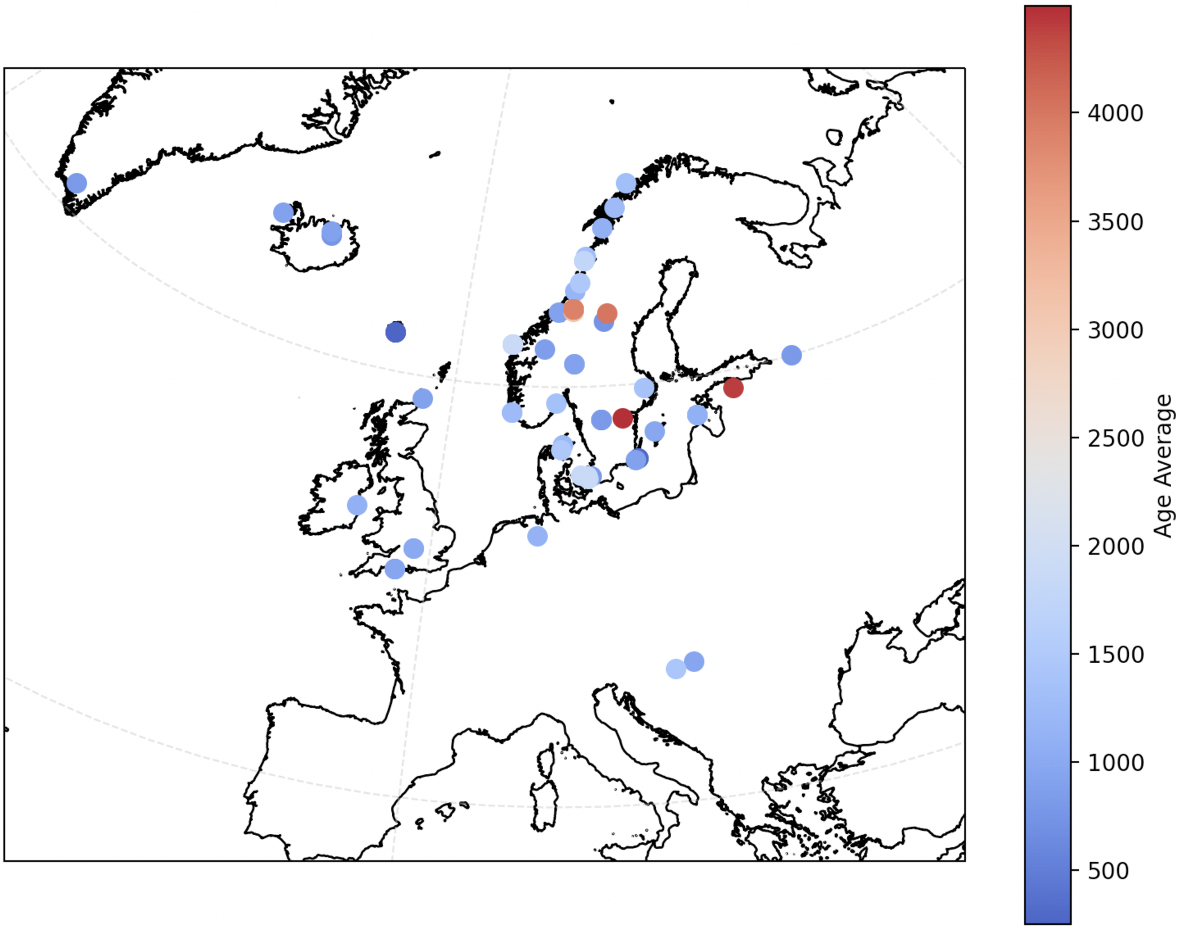

**Fig. S5.153. All individuals carrying haplogroup R1a1a1b1a3a (R1b-Z284) from the dataset, coloured by age BP.**

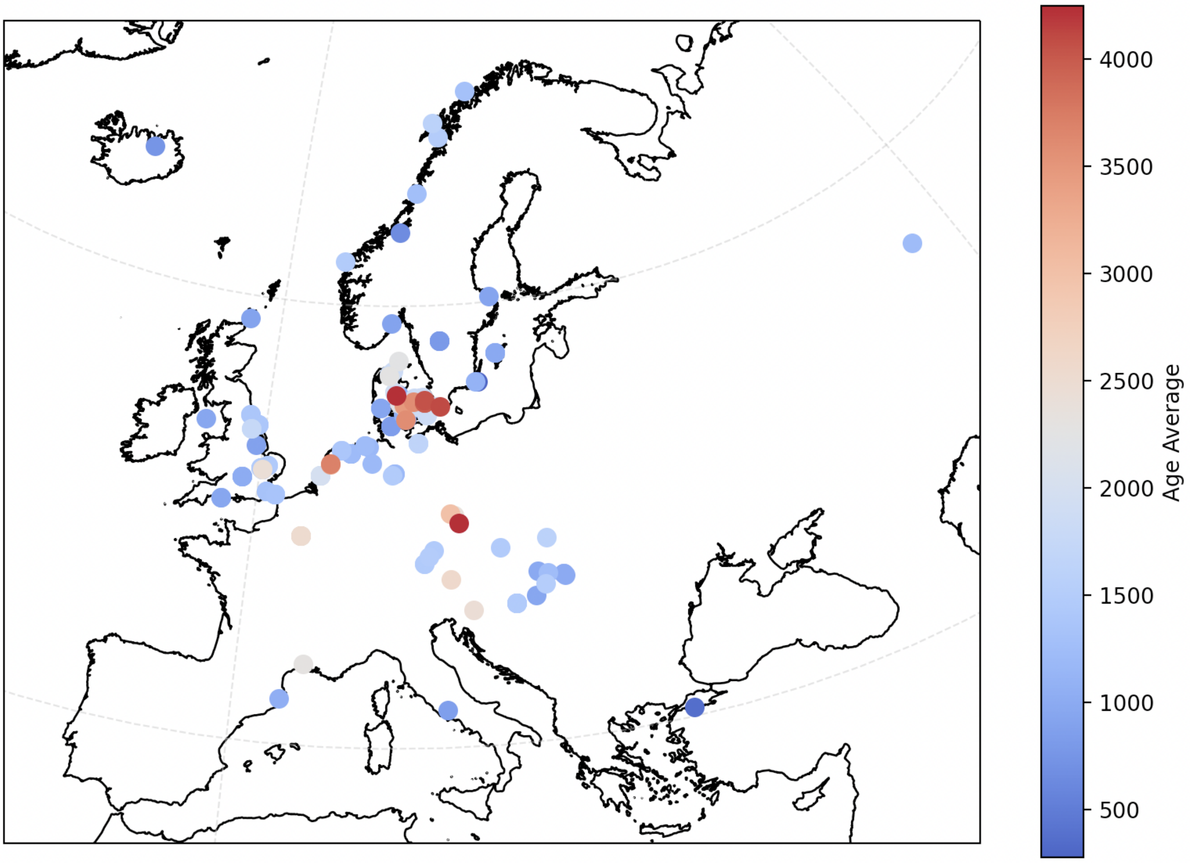

**Fig. S5.154. All individuals carrying haplogroup R1b1a1b1a1a1 (R1b-U106) from the dataset, coloured by age BP.**

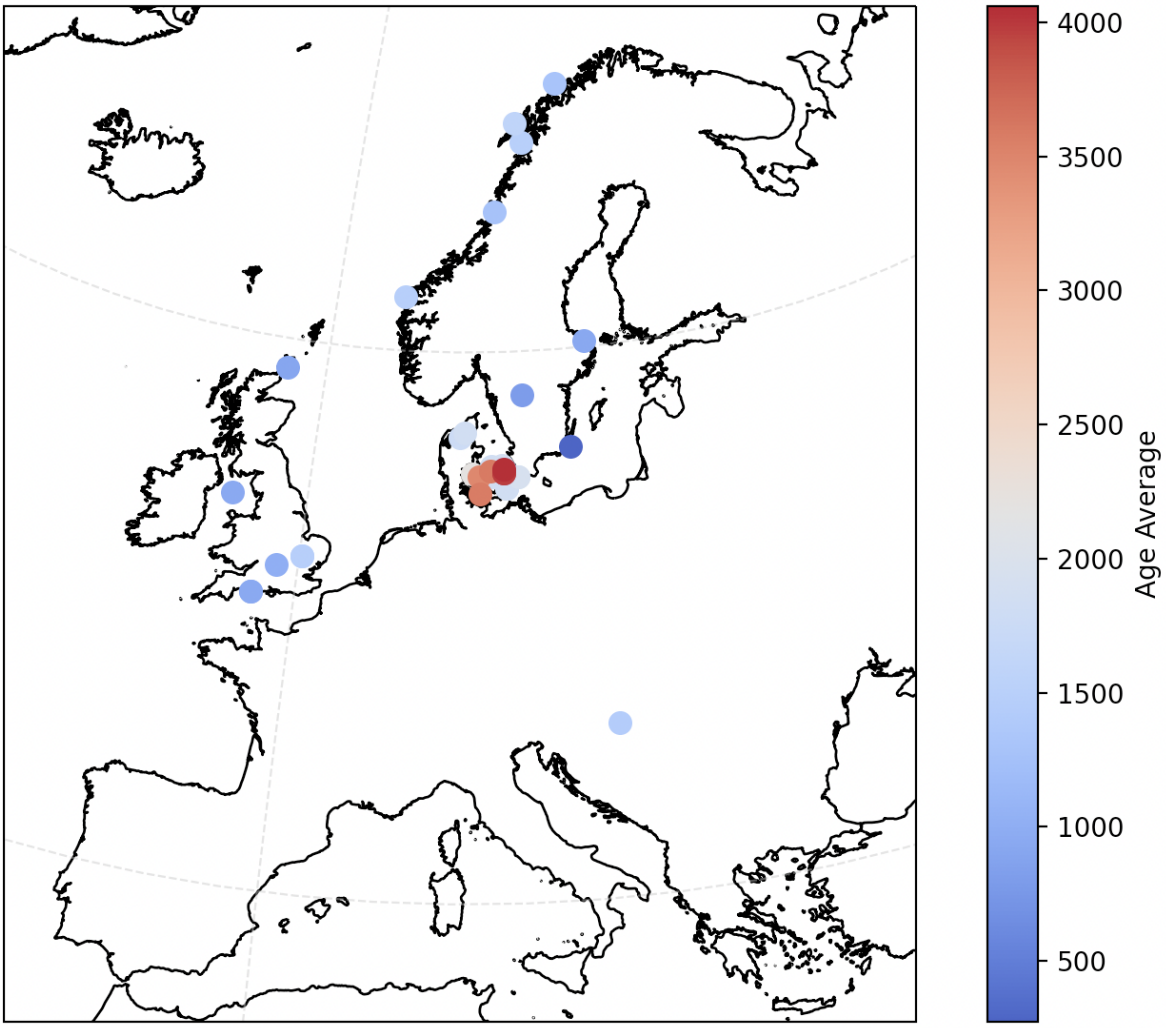

**Fig. S5.155. All individuals carrying haplogroup R1b1a1b1a1a1c (R1b-Z19) from the dataset, coloured by age BP.**

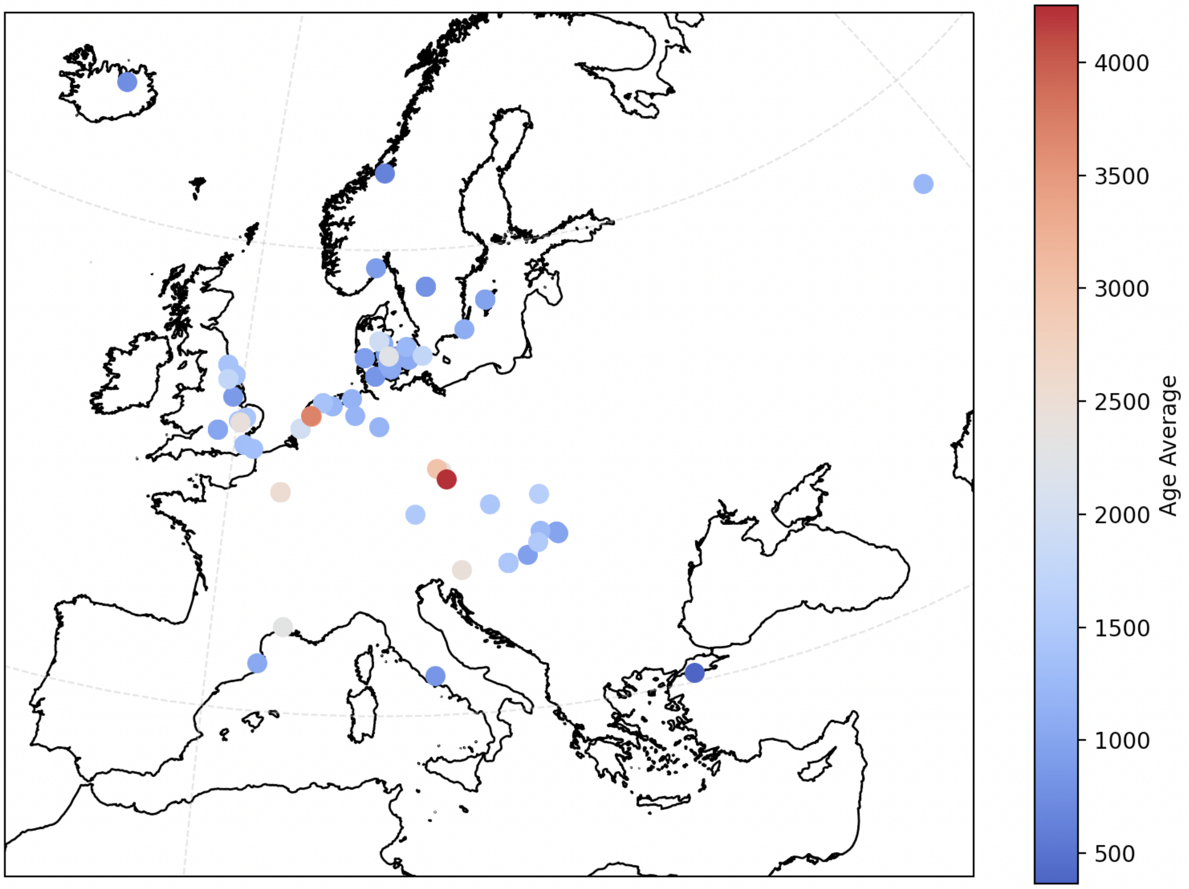

**Fig. S5.156. All individuals carrying haplogroup R1b1a1b1a1a1b (R1b-S263) from the dataset, coloured by age BP.**

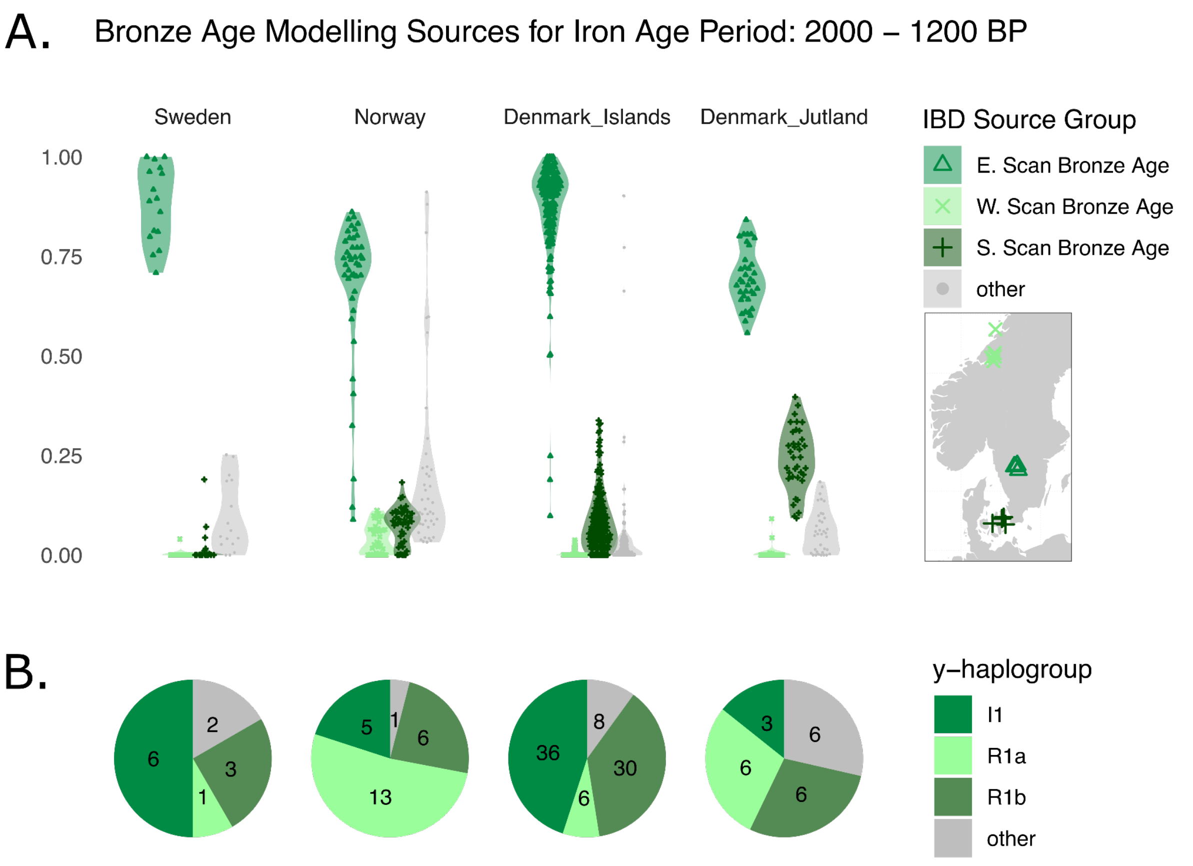

**Fig. 5.157 A) Violin Plots showing the proportion of Bronze Age Scandinavian ancestries modelled for each Iron Age individual from Sweden, Norway and Denmark (Set 5) and B) pie charts showing the proportions of the Y-haplogroups for the Iron Age regions.** The highest proportions of Eastern, Western and Southern Scandinavian Bronze Age ancestry are in the local region, despite Eastern Scandinavian Bronze Age being the highest on average in all regions. Despite the low proportion of Western Scandinavian Bronze Age ancestry in the Norwegians, the proportion of the corresponding R1a haplogroup is high.

#### S5.4.3. Genetic sex determination

We used pysam to estimate depth of coverage for chromosomes 1 to 22, X and Y, by counting and measuring the length of reads with mapping quality above 30 and dividing this sum by the according reference contig length. In the case of the Y-chromosome, due to pervasive repetitive regions not amenable to short-read mapping[203,216,218], we restricted our analyses to the 10 Mb single-copy region defined in[216]. Males have both their X and Y chromosomes at theoretically half the coverage of the autosomes, so we calculated X/auto and Y/auto ratios to determine the genetic sex of the individuals (Fig. 6.4.3.1). Two individuals were found to be strong outliers (CGG023676, DoC: 6.31x; CGG107480, DoC: 0.96x), so we further estimated the ratio between the coverage of the Y- and X-chromosome (Fig. S5.159 and Fig. S5.160). While both individuals possess Y-chromosomes, one individual was found to have two times more dosage at the X than the Y (CGG023676, karyotype: 47,XXY), while the other instead has double the amount of Y than X (CGG107480, karyotype: 47,XYY). Aneuploidies have been previously detected in the archaeogenetic record, with at least eight previous cases of a 47,XXY karyotype[78,154,219–222] and one other case of 47,XYY[222]. Individual CGG107480, reported here, is context dated to 1850 BP, therefore being the oldest 47,XYY case known.

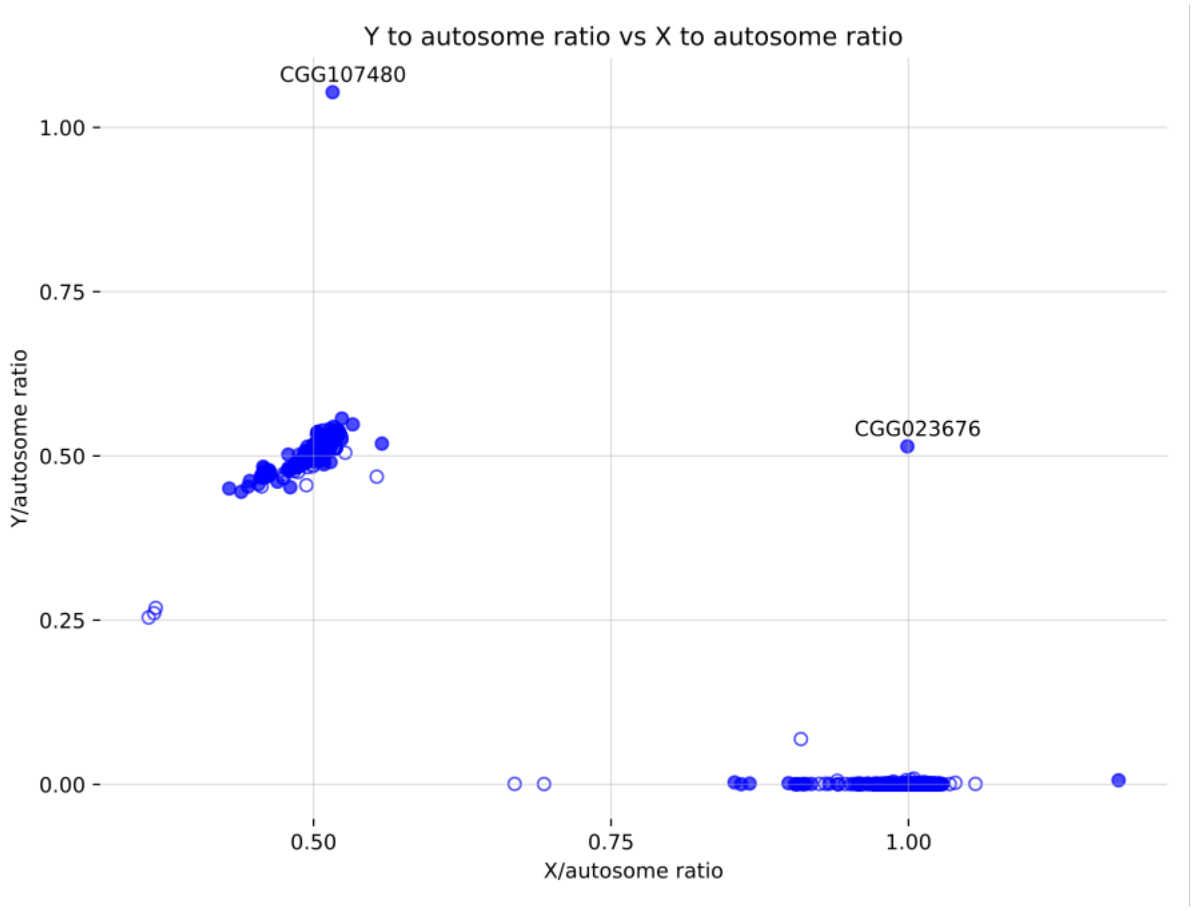

**Fig. S5.158. Y-chromosome/autosome and X-chromosome/autosome depth of coverage ratio obtained for all individuals in the dataset**. Unfilled circles represent low-coverage individuals and/or with signal of contamination. Individuals in the centre left are karyotypically males, while individuals in the bottom right are karyotypically females.

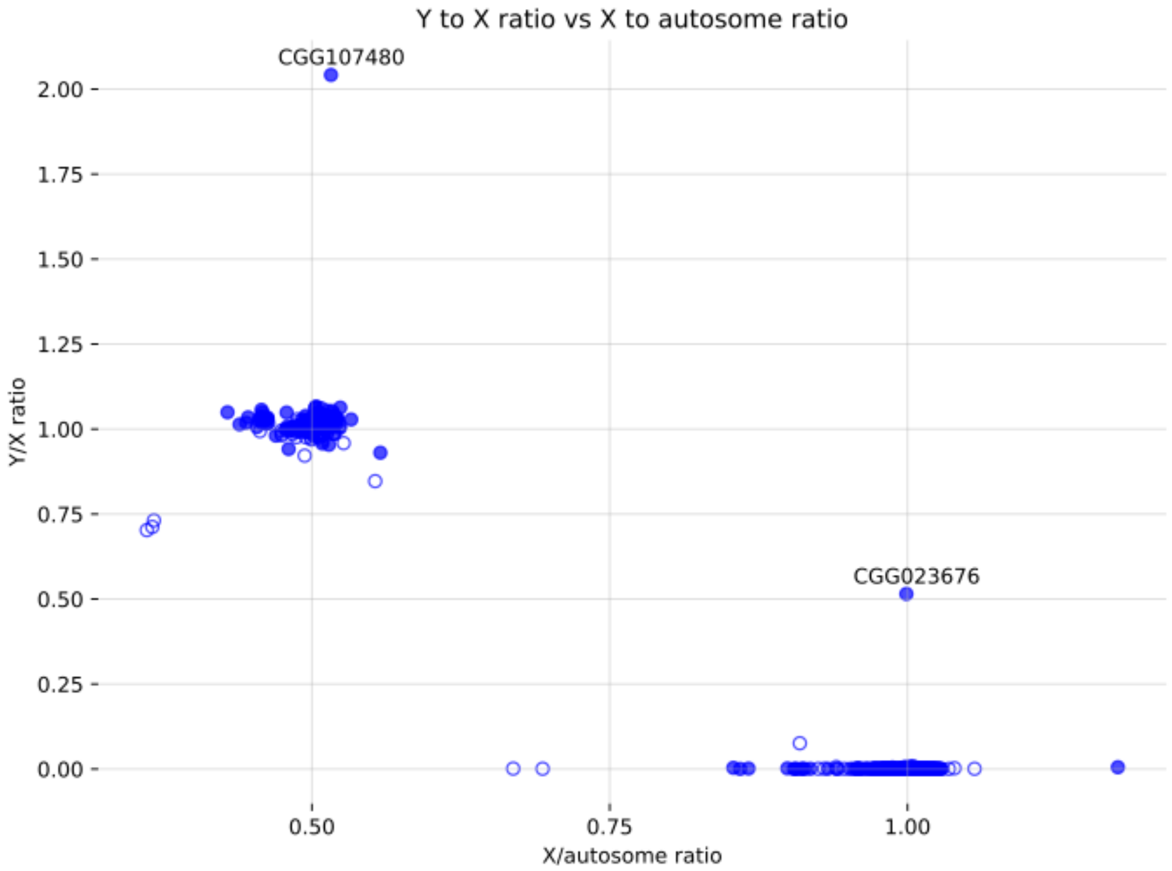

**Fig. S5.159. Y-chromosome/X-chromosome and X-chromosome/autosome depth of coverage ratio obtained for all individuals in the dataset.** Unfilled circles represent low-coverage individuals and/or with signal of contamination. Individuals in the centre left are karyotypically males, while individuals in the bottom right are karyotypically females.

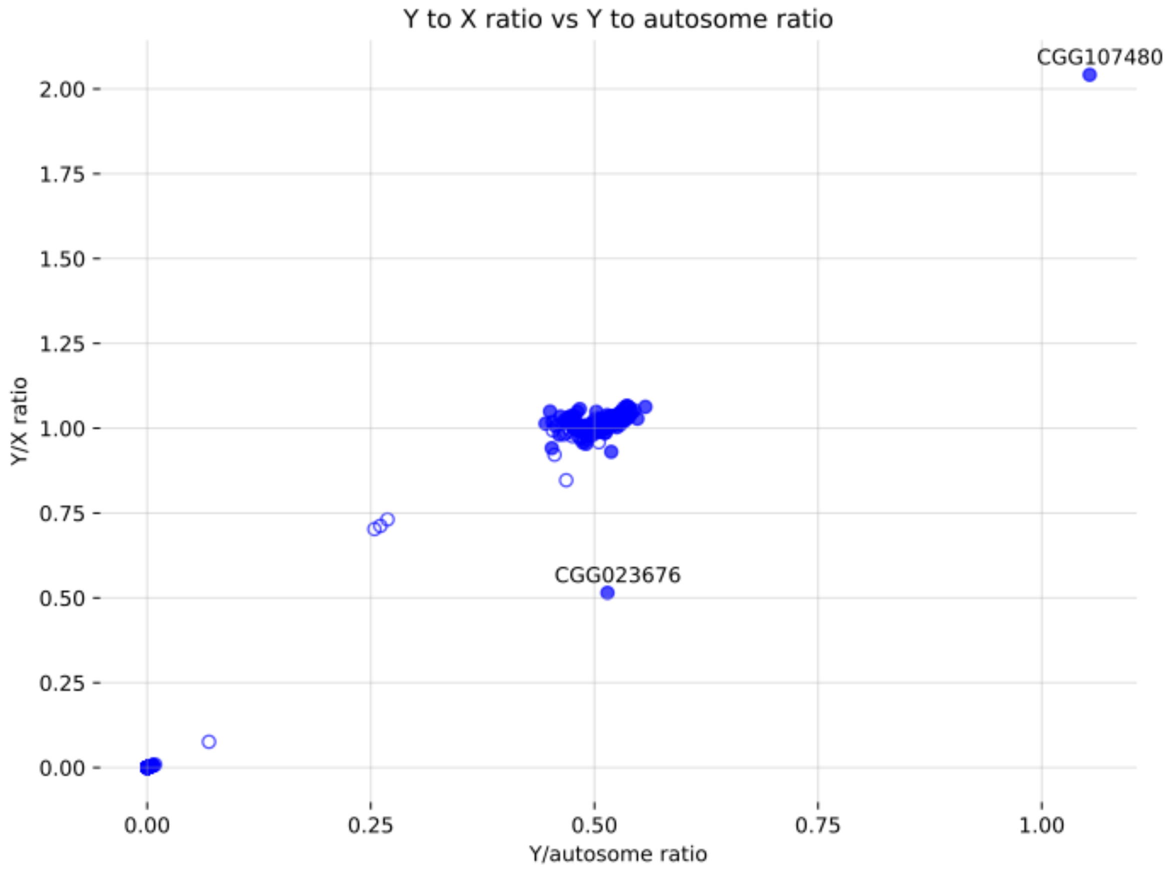

**Fig. S5.160. Y-chromosome/X-chromosome and Y-chromosome/autosome depth of coverage ratio obtained for all individuals in the dataset.** Unfilled circles represent low-coverage individuals and/or with signal of contamination. Individuals in the centre left are karyotypically females, while individuals in the centre of the plot are karyotypically males.

### S5.5. DATES

#### S5.5.1. Method

We carried out DATES[58] analyses to estimate the timing of relevant admixture events in this study. For each set of target/source populations listed in Table S5.10, we first subset plink files to relevant individuals and to the 690211 SNPs used for imputation analyses. Next, we converted plink files to eigenstrat format using convertf[195,223] and ran DATES using default settings. Lastly, we calculated the absolute admixture date using the relative age of the admixture event from DATES (in generations ago), a mean generation time of 25 years, and the midpoint of the mean radiocarbon ages of all dated individuals in the target population (see Fig. S5.161 and Fig. S5.162).

The sets of ancient individuals used in each run can be found in Table S5.10. Here, the 'Dates_group' lists the groups used for each set. Their role as target or source is indicated under the 'role' column. The number of target individuals in each target cluster is indicated in the final field of the 'set' name.

To assess how nonsensical source population would compare to the sources we identify using IBD mixture modelling, we modelled a set of Iron Age Jutlandic individuals as a mixture of Eastern Scandinavian and Africans, Eastern Scandinavian and East Asians, Eastern Scandinavian and Bell Beakers, and Eastern Scandinavian and Southern Scandinavians.

In addition, we modelled the admixture events between Yamnaya and Globular Amphora that occurred around 4800 BP in Bronze Age Southern Scandinavians, Eastern Scandinavians and Western Scandinavians.

Finally, we modelled a series of Iron Age populations from Jutland, the Danish Isles and Norway, as well as early Bronze Age admixed individuals from the Danish Isles and Norway.

#### S5.5.2. Results

For the test sets, including African, East Asians, and Bell Beakers as a source together with Eastern Scandinavian BA, to model Jutlandic Iron Age individuals, we see the weighted LD value curves flatten out sooner and the highest values for the weighted LD to be lower than the source identified by the mixture modelling: the Southern Scandinavian Bronze Age source (Supplementary Fig. S5.161).

By modelling the Eastern, Southern and Western Bronze Age Scandinavians as mixtures of Yamnaya and Globular Amphora, we can ascertain how curves appear with sufficiently distant source populations (Fig. S5.165). For the majority of curves, with the Scandinavian sources, we find curves similar to that of the Yamnaya and Globular Amphora sources for the Iron Age target sets (Fig. S5.162A,B, Fig. S5.163A,B Fig. S5.164A,B ). However, we find the two Bronze Age admixed target sets to be unreliable, which appear very noisy (Fig. S5.162C, Fig. S5.164C), likely due to the small number of individuals in these groups.

In Fig S5.166, we plot the admixture times with 95% confidence intervals, relative to the mean time the target individuals are dated. For the sources from Africa, East Asia, and a target from the Danish Isles and from Norway, the confidence interval intersects zero. As a result, there is no statistical significance of admixture looking based on LD for these sets. Although the error bars for the Bell Beaker sources do not intersect zero when modelling Jutlandic Iron Age sources, based on the curve flattening at a similar time Fig. S5.161C to the African Fig. S5.161A and East Asian Fig. S5.161B sources and the very large error bars, we find the DATES result to be unreliable. Unreliable results are indicated by a dashed line in Fig S5.166.

For the Iron Age Norwegian and Danish Isles sets that do not intersect zero, we find the admixture date to precede the 2500 BP transition from Palaeo-Germanic to Proto-Germanic. For the Jutlandic Iron Age sets the error bars slightly overlap with the 2500 BP time of transition, however for all Iron Age sets, the earliest admixed individuals coincide with the point estimates from DATES.

The output from DATES including confidence intervals can be found in Supplementary Table S5.10.

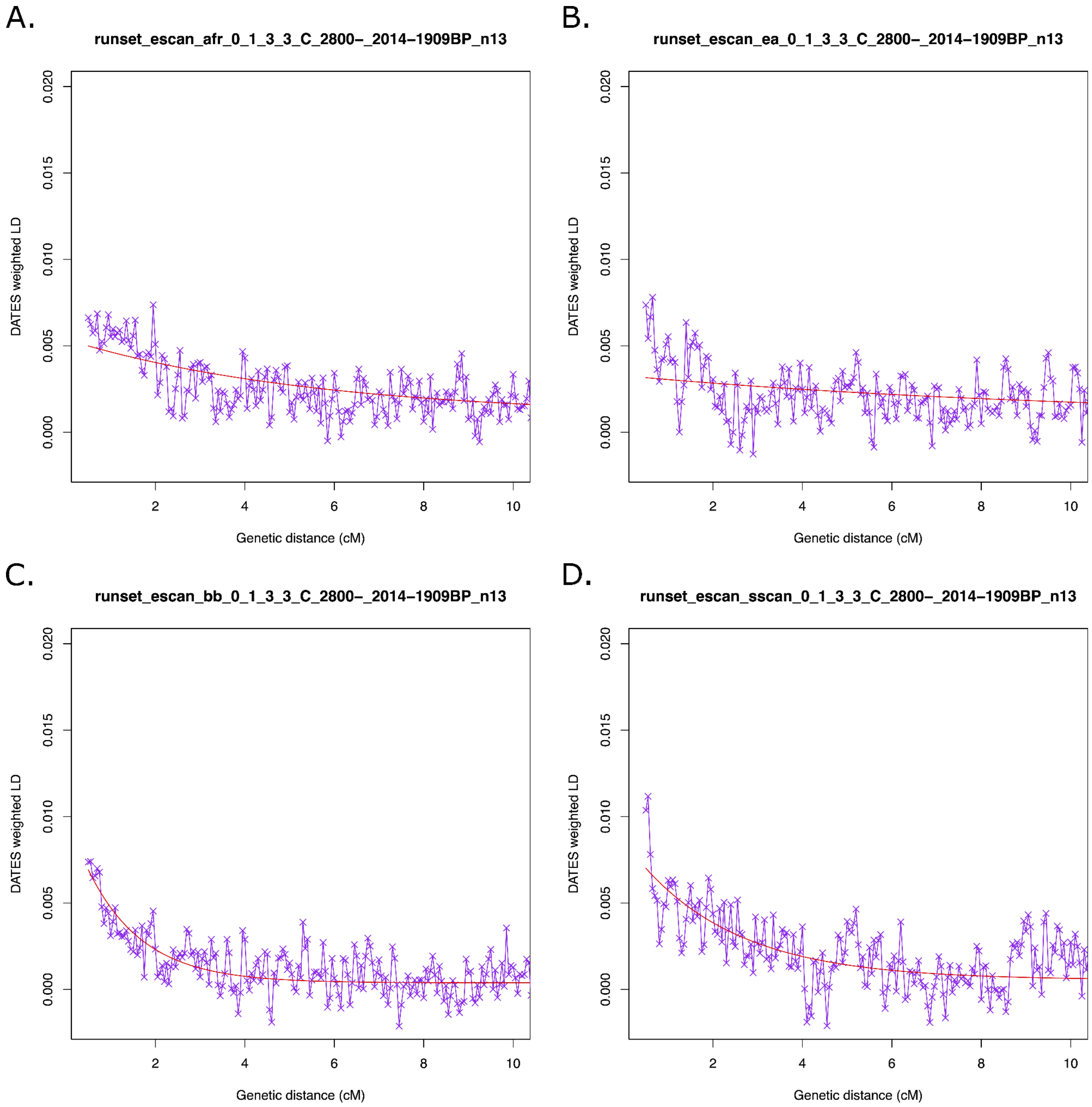

**Fig. S5.161. DATES curves when modelling Jutlandic Iron Age using Eastern Scandinavian Bronze Age and (A) African, (B) East Asian, (C) Bell Beaker and (D) Southern Scandinavian Bronze Age.**

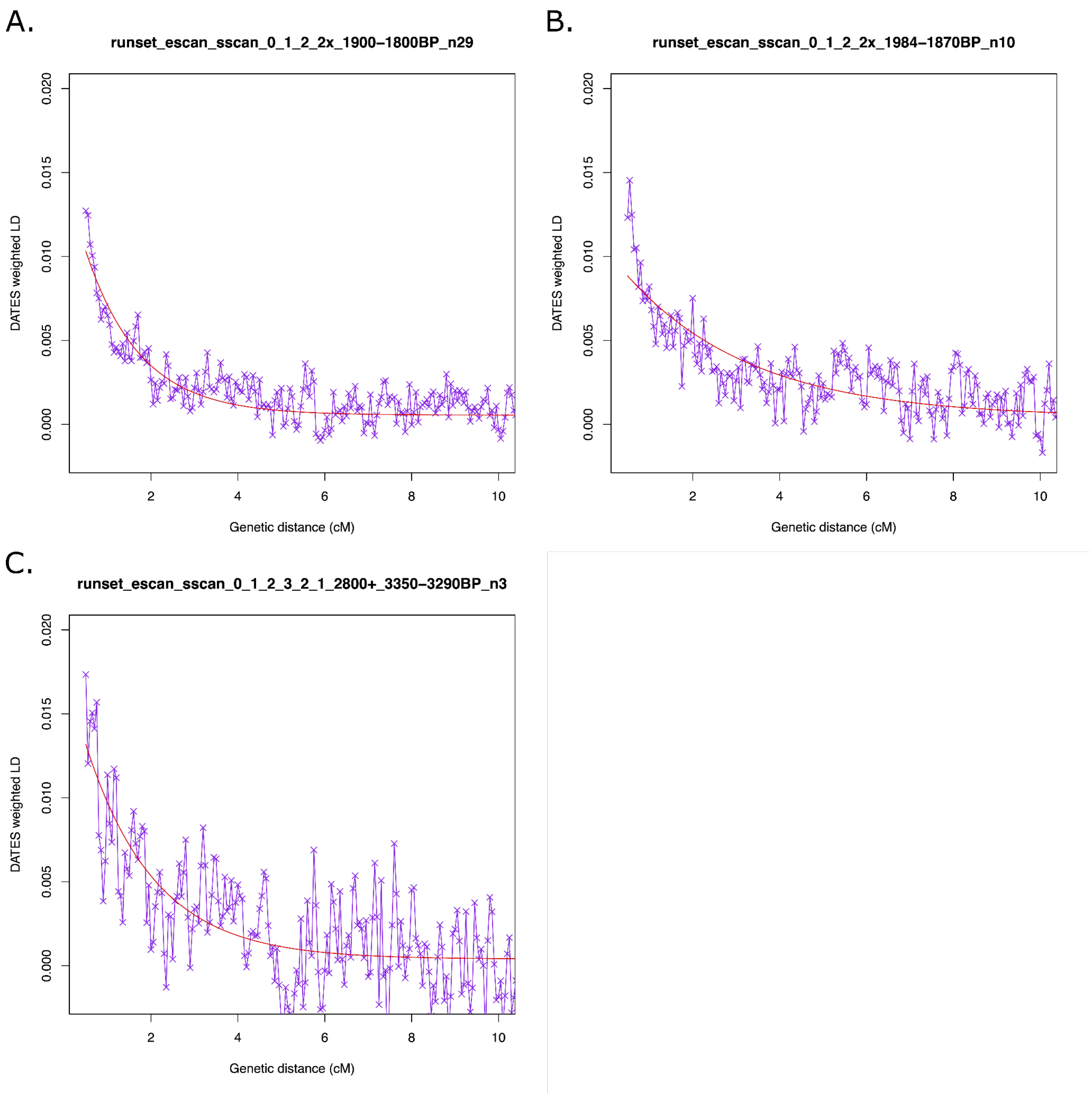

**Fig. S5.162. DATES curves when modelling Danish Isles Iron Age sets (A, B) and Danish Isles admixed Bronze Age set (C) using Eastern Scandinavian Bronze Age and Southern Scandinavian Bronze Age sources.**

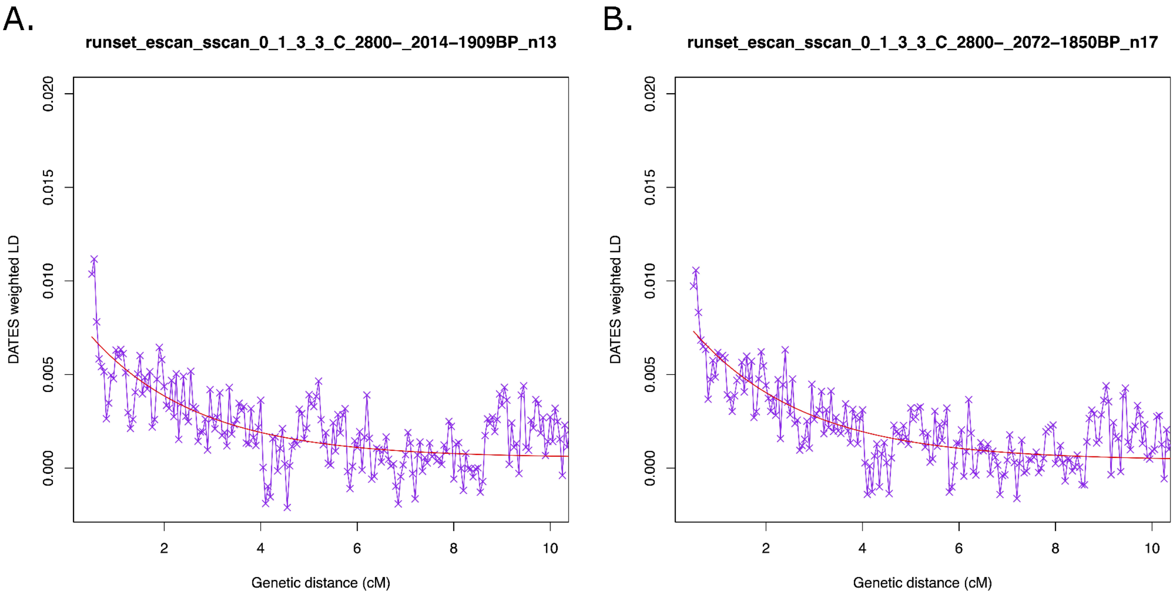

**Fig. S5.163 DATES curves when modelling Jutlandic Iron Age sets (A, B) using Eastern Scandinavian Bronze Age and Southern Scandinavian Bronze Age sources.**

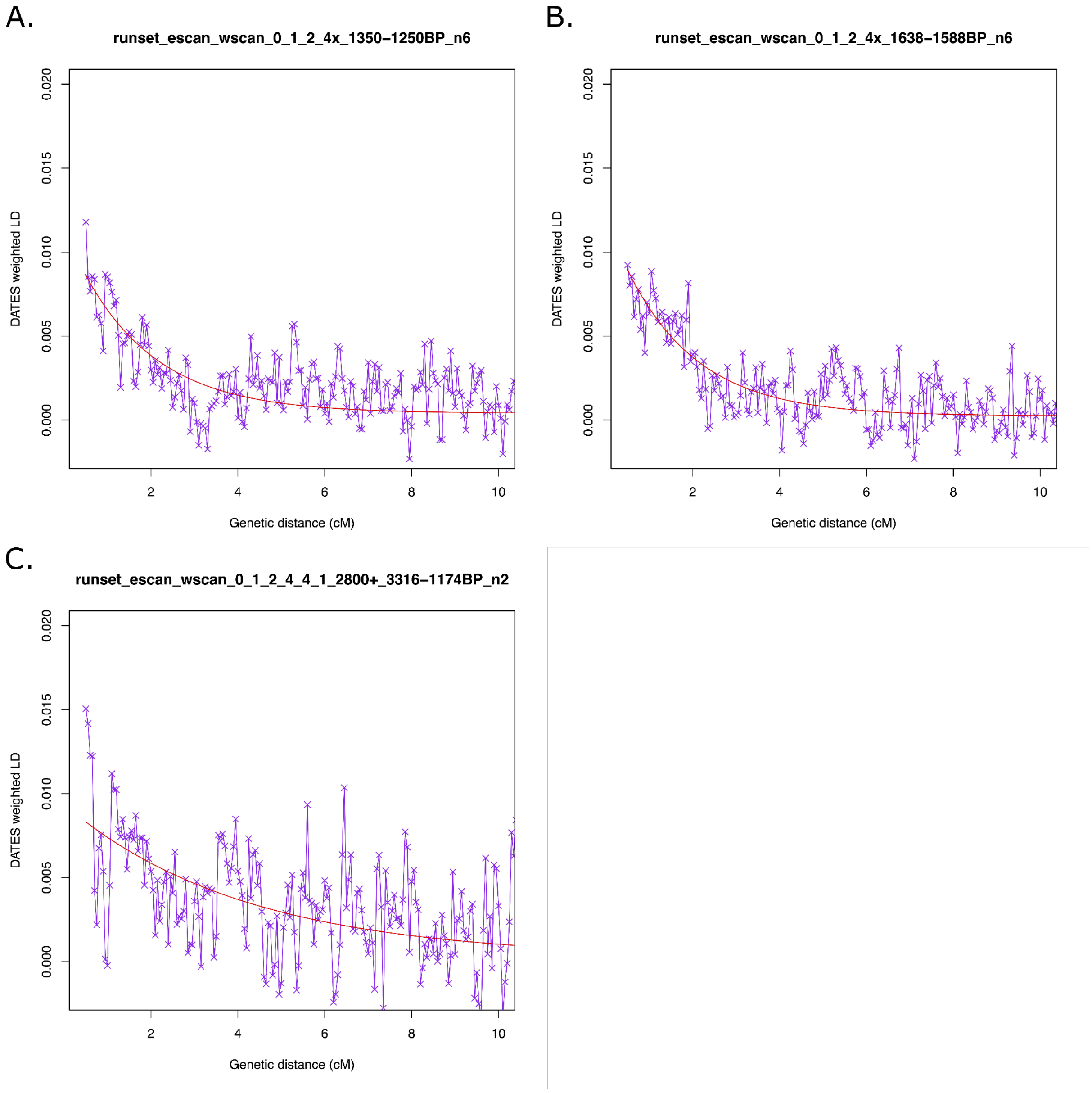

**Fig. S5.164 DATES curves when modelling Norwegian Iron Age sets (A, Bt) and Norwegian admixed Bronze Age set (C) using Eastern Scandinavian Bronze Age and Western Scandinavian Bronze Age sources.**

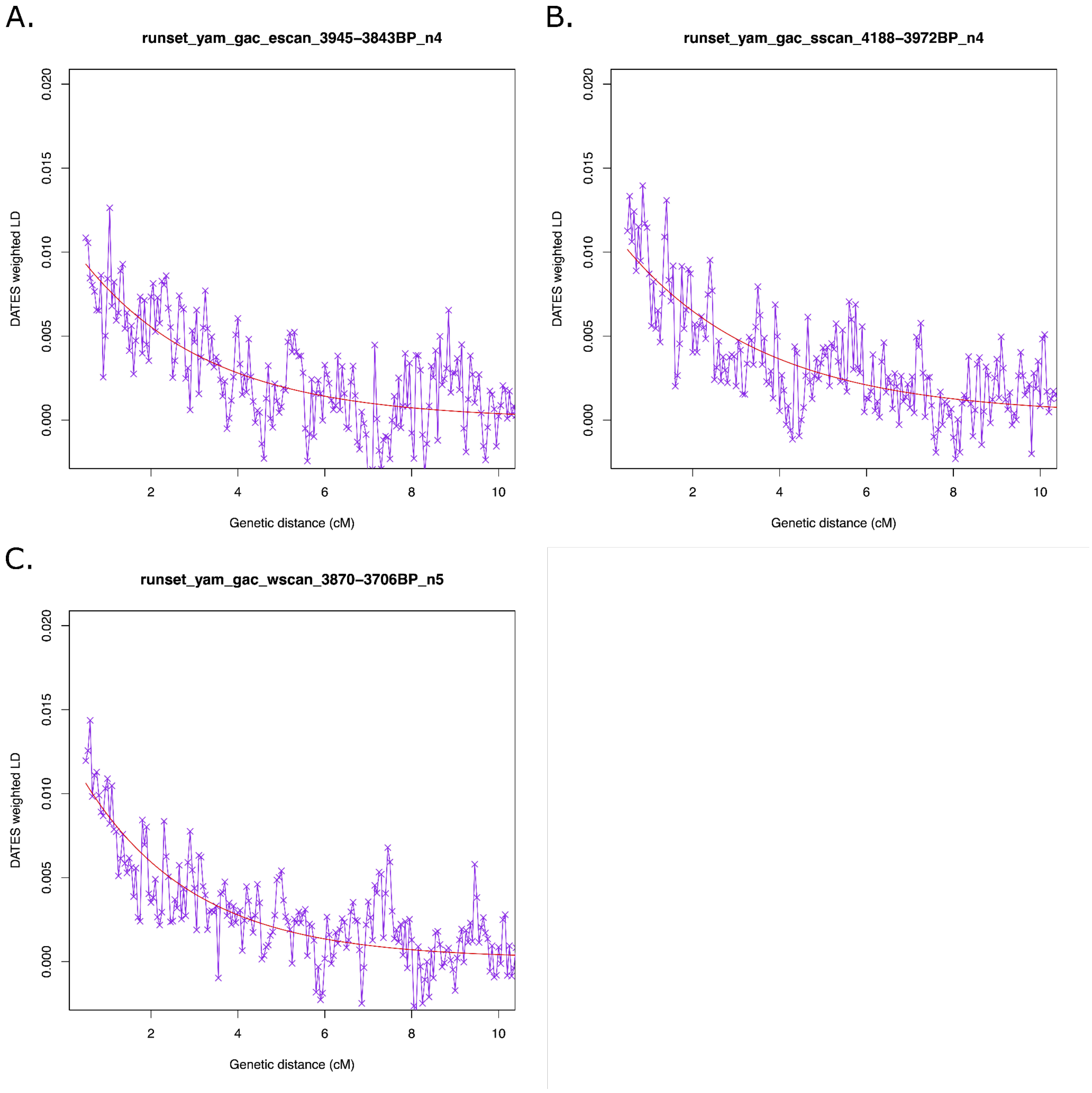

**Fig. S5.165 DATES curves when modelling Eastern Scandinavian BA (A), Southern Scandinavian BA (B), and Western Scandinavian Bronze Age set (C) using Yamnaya and Globular Amphora sources.**

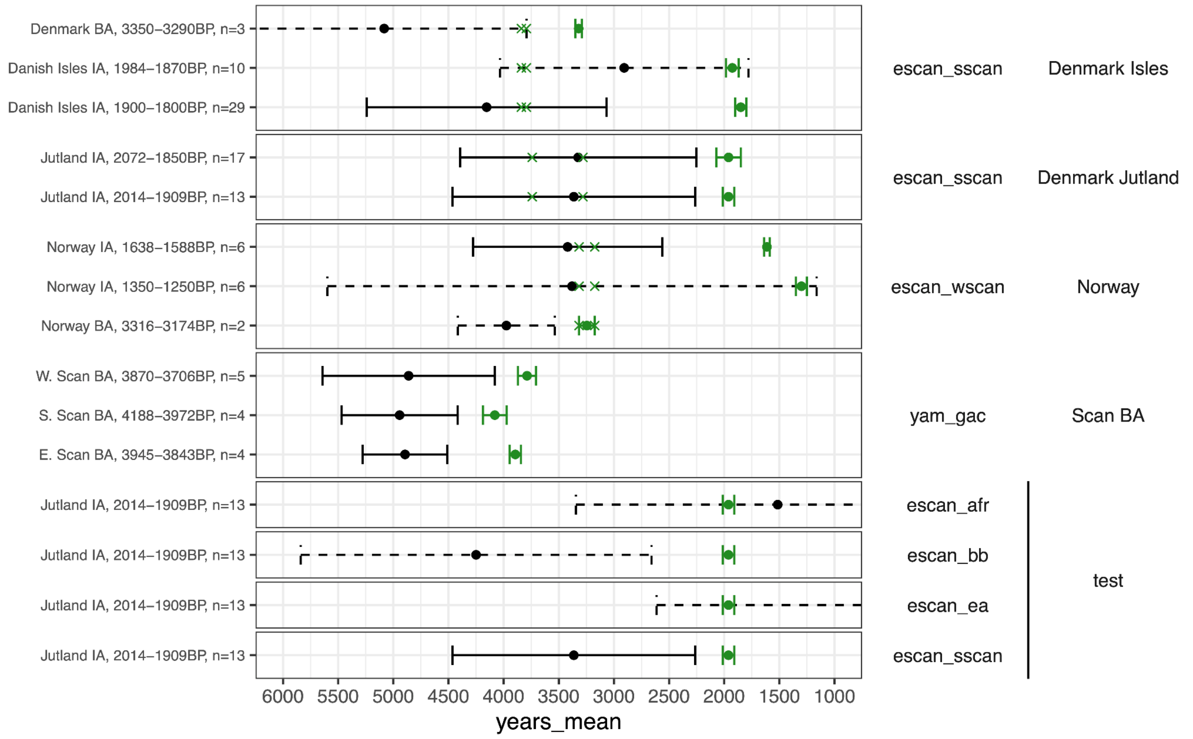

**Fig. S5.166. DATES admixture times. The green error bars represent the time range that the target individuals fall within. The black error bars indicate the 95% confidence intervals for the admixture time, with unreliable dates as dashes. The earliest admixed individuals from each region are marked with a green 'x'.**

#### S5.6. Kinship

In order to identify pairs of close relatives in our dataset, we ran ngsRelate (v2)[97] on the imputed dataset (see section S5) subset to individuals of European ancestry. We used the pairwise relatedness estimate (rab) from ngsRelate to categorise pairs of individuals into degrees of relatedness (0, 1, 2 or unrelated). We define cut-offs for each degree as the midpoint between the theoretical value of the following and preceding degrees of relatedness as follows: 0 degree [1, 0.75), 1st degree [0.75, 0.375), 2nd degree [0.375, 0.1875), unrelated [0.1875,0]. Furthermore, for first degree relatives we categorise each pair of individuals as either parent-offspring or siblings based on the R0 estimate from ngsRelate as follows: PO [0,2-2), sib [2-2,1].

We first merged data from 31 samples that we determined originated from 14 unique individuals. After merging the data, we find in total 82 pairs of close relatives among our samples sequenced for this study, represented by 110 individuals. Among the pairs of close relatives, we identify 26 parent-offspring relationships, 13 pairs of full siblings and 43 pairs of second degree relationships (grandparent-grandchild, avuncular relationship or half-siblings). We identify several clusters where multiple individuals are related. Where possible, we resolved these clusters into pedigrees (Fig. S5.167, Fig. S5.168, Fig. S5.169, Fig. S5.170, Fig. S5.171). Generally, owing to a relatively low number of samples from each site, we do not find any large extended pedigrees. We do, however, identify multiple examples of relatives coming from different sites. Most striking is the finding of two pairs of siblings and one pair of parent-offspring, all of which have one individual buried at the site Les Moidons and the other at Parançot 2.4 km away (Fig. S5.62).

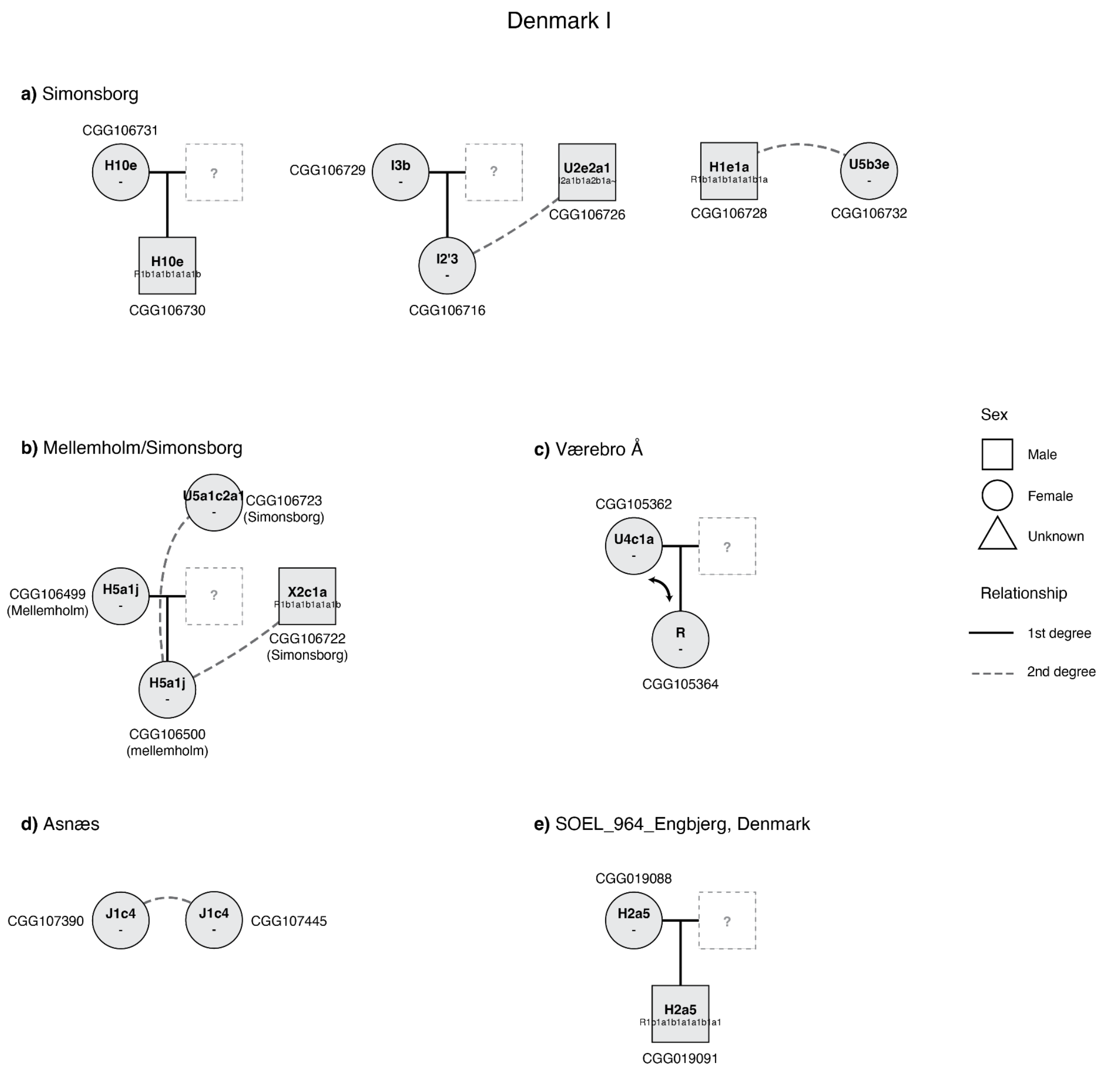

**Fig. S5.167. Pedigrees from Denmark (part 1).** Pedigrees were constructed manually, based on results from ngsRelate. Solid black line indicates first-degree relationships, whereas stippled line indicates unknown second-degree relationships. Shape for each individual indicates sex. Mitochondrial haplogroup is denoted inside the shape for each individual.

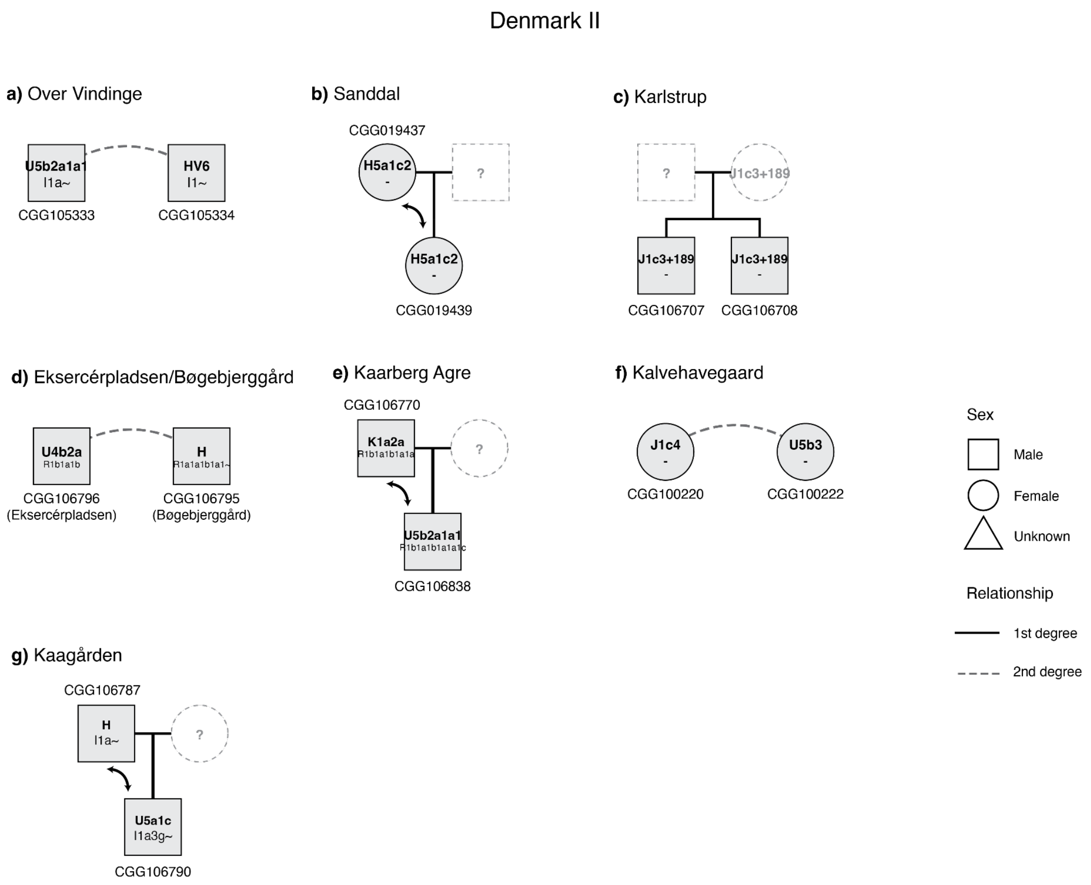

**Fig. S5.168. Pedigrees from Denmark (part 2).** Pedigrees were constructed manually, based on results from ngsRelate. Solid black line indicates first-degree relationships, whereas stippled line indicates unknown second-degree relationships. Shape for each individual indicates sex. Mitochondrial haplogroup is denoted inside the shape for each individual.

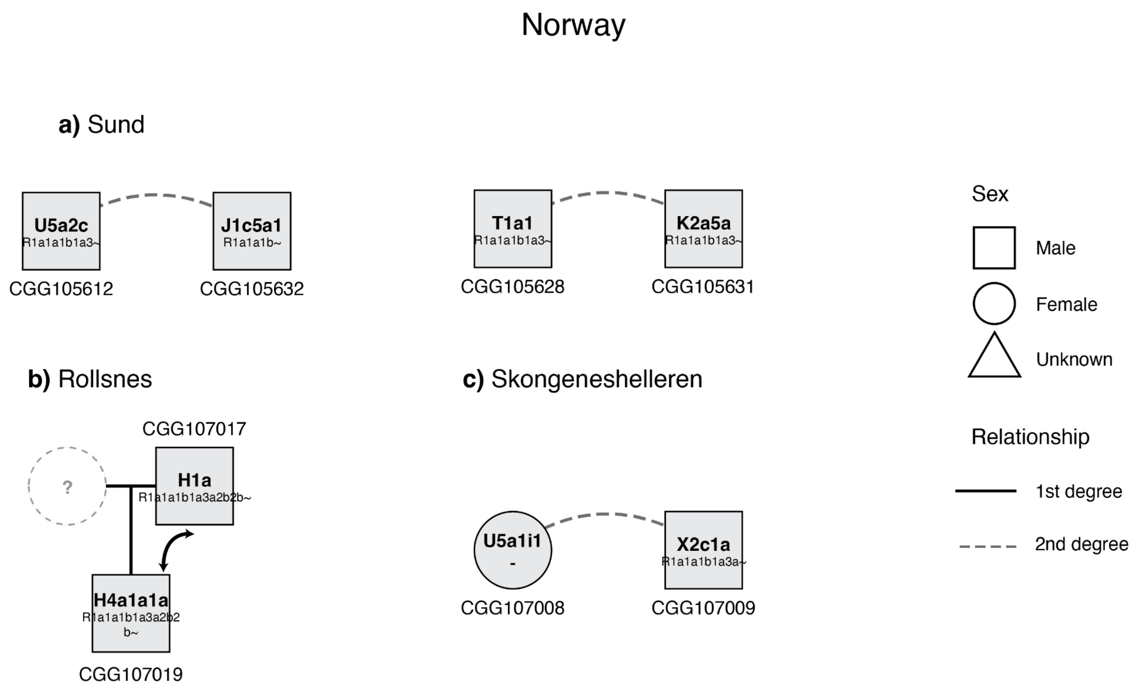

**Fig. S5.169. Pedigrees from Norway.** Pedigrees were constructed manually, based on results from ngsRelate. Solid black line indicates first-degree relationships, whereas stippled line indicates unknown second-degree relationships. The shape for each individual indicates sex. Mitochondrial haplogroup is denoted inside the shape for each individual. The incompatibility of the context dates between the Rollsnes and Toldnes individuals suggest C14-dating these individuals is required before the context can be properly understood.

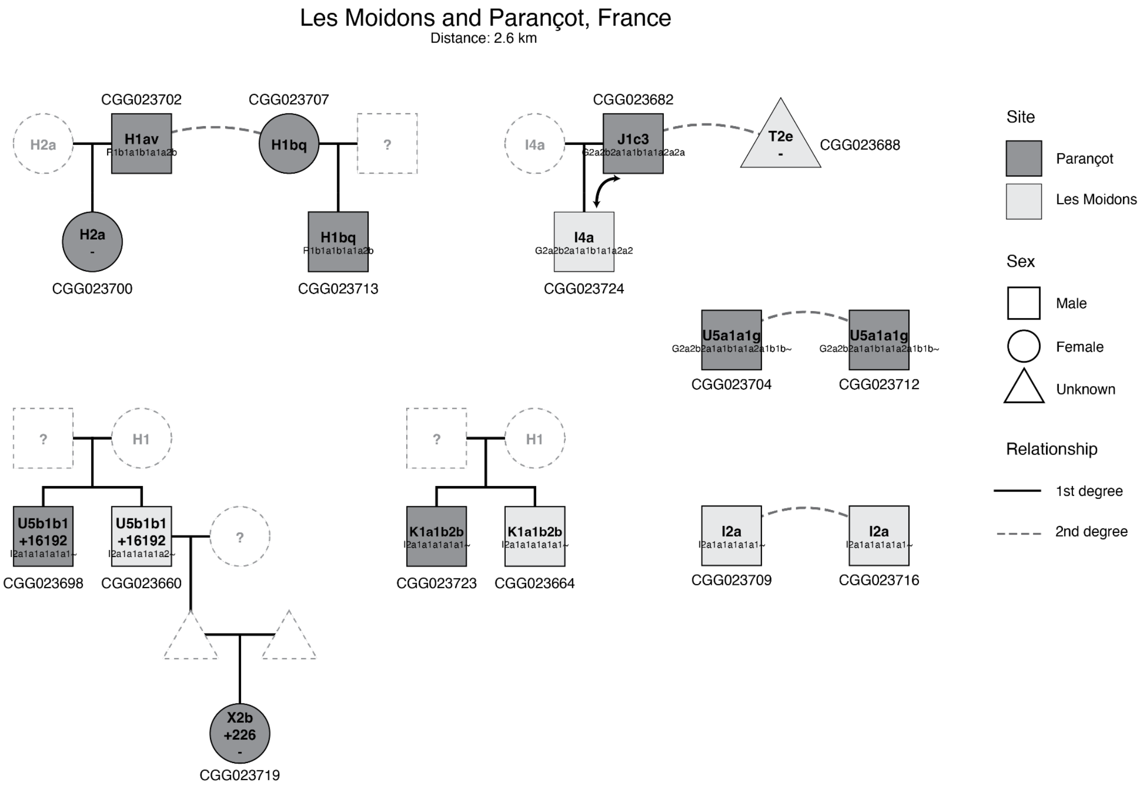

**Fig. S5.170. Pedigrees from Les Moidons and Parançot, France.** Pedigrees were constructed manually, based on results from ngsRelate. Solid black line indicates first-degree relationships, whereas stippled line indicates unknown second-degree relationships. Colour and shape for each individual indicate site and sex, respectively. Mitochondrial haplogroup is denoted inside the shape for each individual.

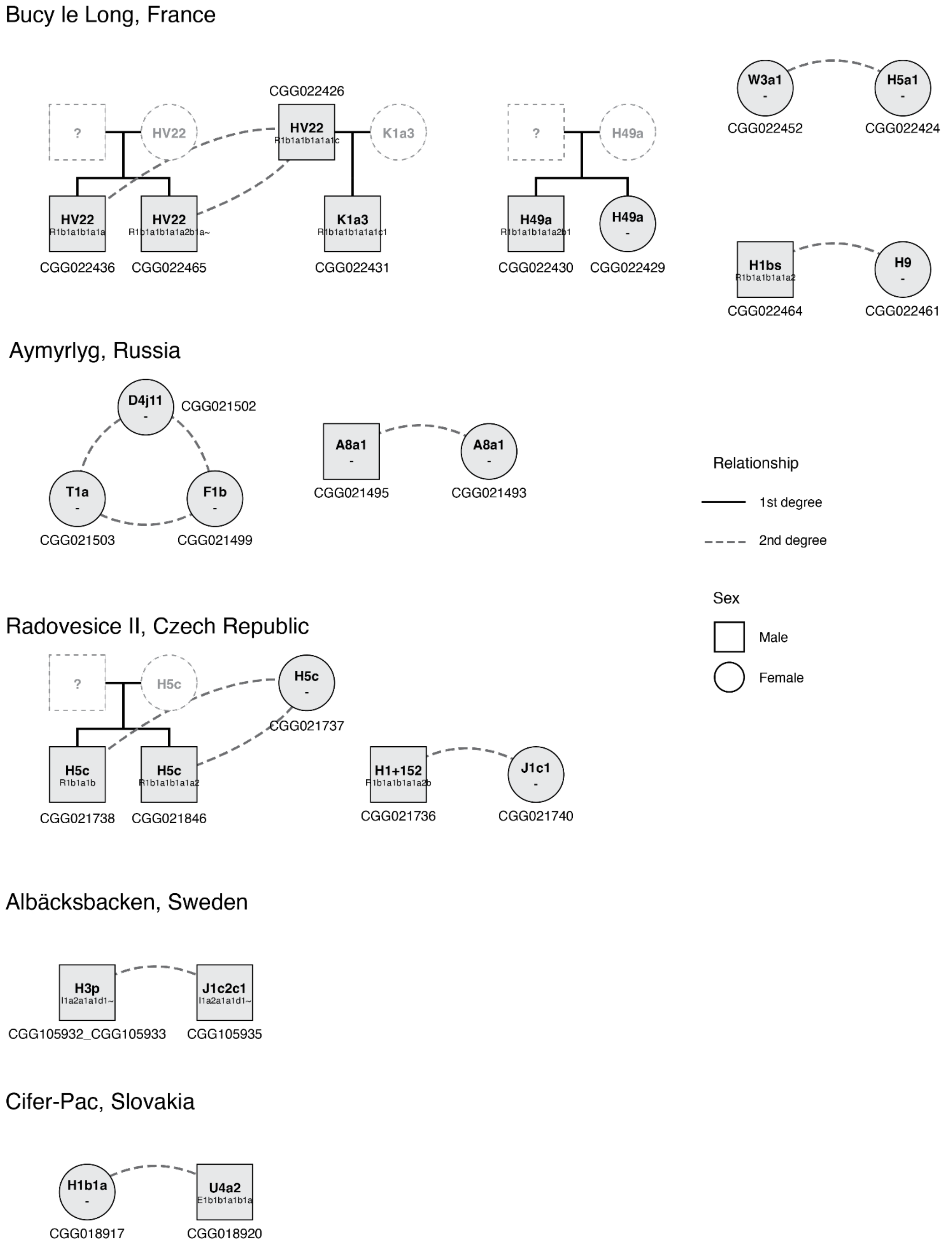

**Fig. S5.171. Smaller pedigrees from various sites.** Pedigrees were constructed manually, based on results from ngsRelate. Solid black line indicates first-degree relationships, whereas stippled line indicates unknown second-degree relationships. Shape for each individual indicate sex. Mitochondrial haplogroup is denoted inside the shape for each individual.

#### S5.7. Extended discussion

##### S5.7.1 Viking Age genetic outliers

The combination of dense sampling and high-resolution IBD modelling from the Iron Age sources also allows for the identification of outliers and their origins within Scandinavia (Fig. S5.29). Some notable examples are individuals from the Salme Viking burial on the Island of Saaremaa cluster together with and are modelled with similar profiles to those of Iron Age of Central Sweden rather than the later Vikings, supporting origins in the Mälaren valley of Sweden. At Kalmargården, two beheaded individuals appear to be non-locals, one (CGG107579, 2654/79, East) modelled with a large proportion of Norwegian IA ancestry, and the second (CGG107580, 2654/79, West) with a large proportion of Baltic Bronze Age-related ancestry. In pre-Viking Norway, as early as ~1240 BP, we find individuals with Celtic British Isles-associated ancestry, representing the earliest genetic evidence of the Viking migrations between the two regions.

##### S5.7.2 The Netherlands and Bohemia

In the Netherlands and Bohemia, on the border regions between the Corded Ware and Bell Beaker cultures, dense sampling provides insight into the Bronze to Iron Age transition. In the Netherlands, we see a large degree of continuity distinct from the surrounding areas, whereas in Bohemia we detect a series of transitions.

The Bell Beaker subcluster 0_2_1_5_x located primarily from the Eastern North Sea region (present day the Netherlands) is unique in its high Lithuanian/Latvian Hunter-Gatherer ancestry, low European Farmer and inability to be modelled primarily as Bell Beaker ancestry, like most others from Bell Beaker subclusters (0_2_1_x) (Fig. S5.172).

When using early representatives of this cluster as a source, we see a large degree of genetic continuity from 3700 to 1700 BP. From Valkenburg (ZH), however, there are a number of individuals that do not fit the profile. The Roman cemetery Valkenburg Marktveld, located south of the auxiliary fort, was used between 1950 and 1650 BP for the entire military community that consisted of men, women and children, who lived in the vicinity of the auxiliary fort. Over 650 individuals were recovered, 145 of which are inhumations (41 adults, 104 children and infants); an extraordinary number, as cremation dominates the Roman burial record in the Netherlands. The individuals included in this study are possibly associated with different departments of the Roman army, so the presence of non-local individuals is not unexpected (DeCoster et al., in prep.). For the individuals that do not fit the local profile, most are Celtic (similar to contemporaneous people from British Isles or France) and one is similar to people deriving their ancestry from the Bronze Age Eastern Mediterranean. A single individual is modelled with North East European ancestry.

A transition by at least 1612 BP is apparent, where Frisian individuals are modelled primarily as Southern Scandinavian ancestries, but many possess small amounts of the local Eastern North Sea ancestry.

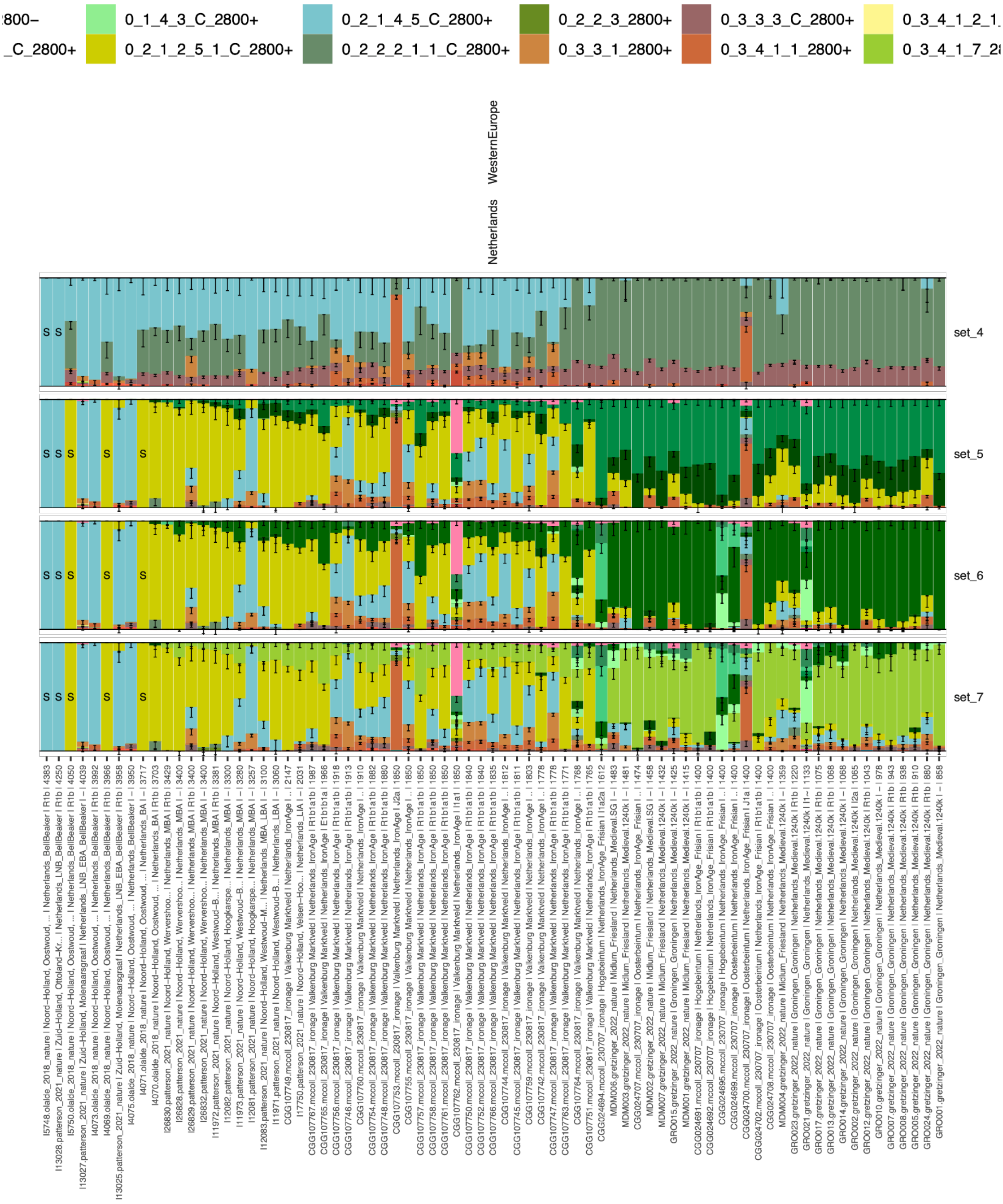

**Fig. S5.172. Subset of Mixture modelling results from Fig. S5.24, for the Netherlands.**

Rows 1-4 correspond to sets 5-8 of the mixture modelling.

This contrasts with Bohemia, where we replicate previous findings[184], and provide additional insight into the cultural transition to the Knoviz cultures, previously suggested to have links with Celtic Europe[26]. As visible in the top panel in **Fig. S5.173**, we see the increased Western Hunter-Gatherer ancestry during the Neolithic after 7000 BP. We see the transition from Corded Ware to Bell Beaker around 4250 BP, reflected in the increased Farming ancestry visible in the top panel. The specific farming ancestry can be identified in the second panel as the introduction of European Farming ancestry, rather than just Globular Amphora Farming ancestry that was present during the Corded Ware period. In the third panel, using Corded Ware and Bell Beaker related sources, we can identify the transition directly. The transition to the Únětice period around 4100 BP is reflected again by an influx of Corded Ware-related ancestry. From the lower panel, we can see this ancestry is not modelled by the Scandinavian BA sources. At this time, ancestry to the west and south is primarily Bell-Beaker related, and to the north is Scandinavian or Baltic, the trends of which can be seen in the spatio-temporal kriging results in Fig S5.174. Hence the most likely source region for this Corded Ware related ancestry is further east, supporting previous suggestions of genetic links to the northeast during the Únětice period[184].

By the time of the first ancient genomes associated with the Tumulus Culture in Bohemia, we find another transition has occurred, where individuals are modelled with the highest yet proportions of Farming-related ancestry (top panel), reflecting another migration into the region. From the middle panel, we see that this farming ancestry is related to European Farmers. In the lower we find the steppe ancestry to be Bell-Beaker related, and that this period is marked by the appearance of Bronze Age Anatolian-related ancestry. We see a general pattern of continuity from the Tumulus to Knoviz to Hallstatt. We also note this later period as one of diversity, in which we find unadmixed outliers during this period with genetic origins in Scandinavia, the Baltic, and individuals modelled entirely with Neolithic Farming ancestry (Fig. S5.173).

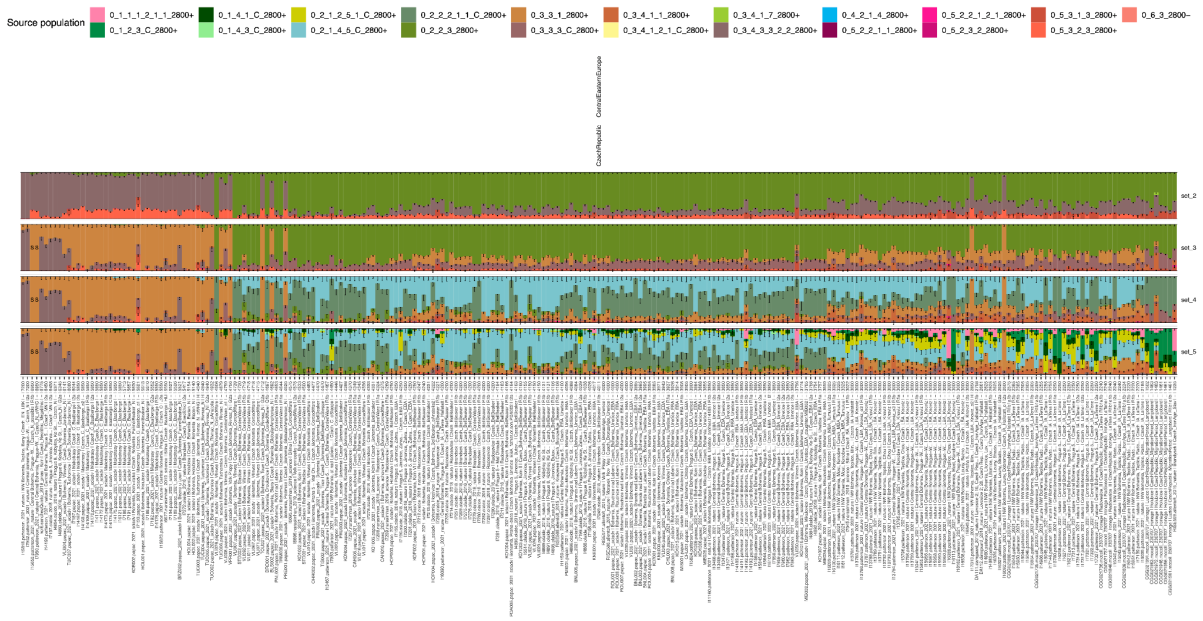

**Fig. S5.173. Mixture Modelling Results for the Czech Republic showing the shifts in Hunter-Gatherer, Anatolian Farmer and Yamnaya-related ancestry in the top panel, additional resolution into Farming ancestries (European Farmer, Globular Amphora, Bronze Age Mediterranean) in the second panel, into Corded Ware and Bell Beaker-related ancestry in the third, and Eastern North Sea and Scandinavian Bronze Age ancestries in the lower panel.**

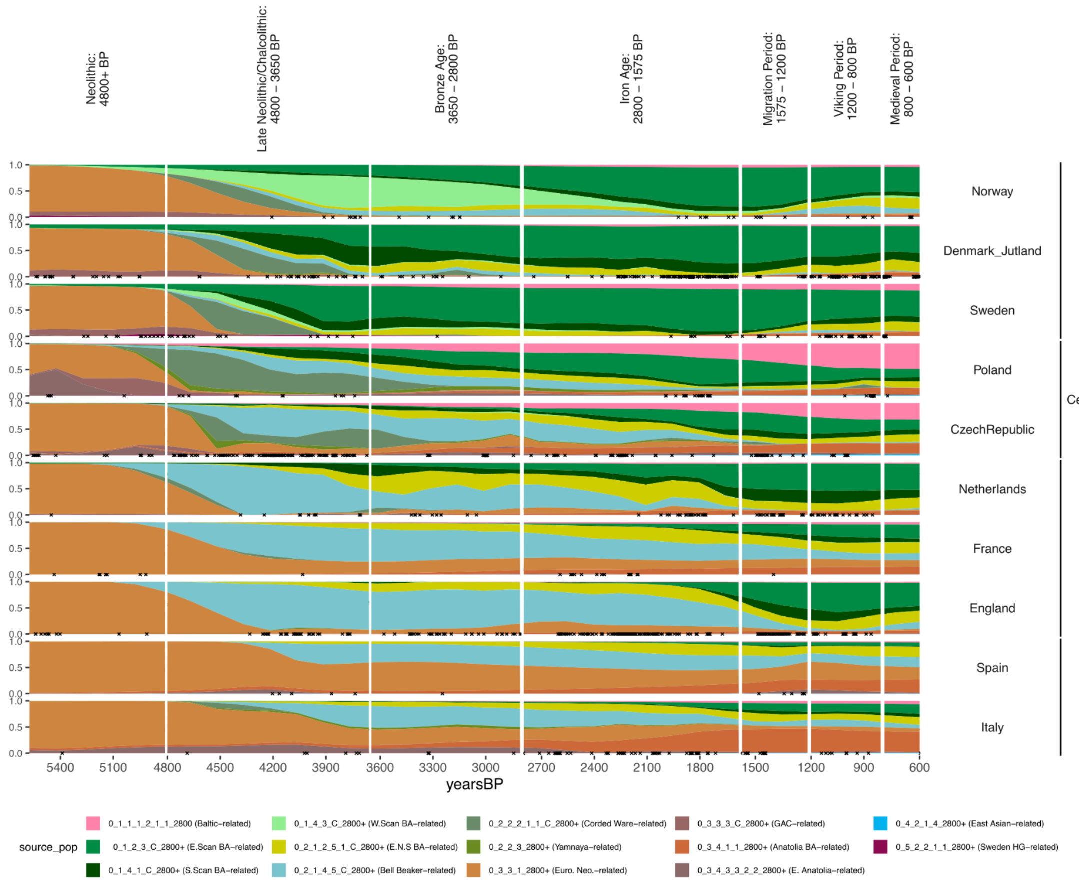

**Fig. S5.174. Spatiotemporal Kriging results showing the general trends of the proportion of ancestries when including additional Bronze Age Scandinavian and Baltic-related sources (Set 5).** The 'Europe-wide' kriging parameters are applied here. Individuals within 250 km of the point used for each country are indicated with an 'x'.

#### S5.7.3 Norway

From the pre2800BP clustering (S5.2.2), we see Norwegian individuals from between 4200 - 3200 BP clustering together with some of the earliest Steppe-related people from across Sweden (4630-4200 BP) and Denmark (4622 BP). In contrast, by 4200 BP in Denmark and 4,000 BP in Sweden, clusters associated with Southern Scandinavian and Eastern Scandinavian Bronze Age ancestry have formed.

As discussed in Supplementary Note S5.2.2 and shown in Fig. S5.29, from the Bronze Age, the expansions of Eastern Scandinavians had a large impact on the population structure of Norway. After 3200 BP in Norway, all individuals are modelled as a mix of Western and Eastern Scandinavian ancestry (Set 5). Initially, proportions are roughly equal, but by the Iron Age, only a small proportion of this Western Scandinavian BA ancestry is detected.

In contrast to Denmark and Southern Sweden, there is a large degree of genetic continuity from the Iron Age to the Viking Period of Norway, with little impact of the migrations from Northern Germany/Southern Jutland. We do, however, find multiple individuals who derive all their ancestry from Bell Beaker-related sources, reflecting migrations from the British Isles as early as ~1240 BP.  In addition, we find early evidence of Baltic Bronze Age related ancestry at 1916 BP (CGG105643, Følstad) and 1769 BP (CGG105601, Toldnes).

From Northern Jutland during the Iron Age, there are multiple instances of individuals modelled with large proportions of Norwegian Iron Age ancestry. Of particular note is CGG106489 (1774 BP, Sondrup Østergaard, Ulstrup sogn) who is modelled almost entirely by Norwegian Iron Age ancestry. Other individuals with high proportions include CGG106810 (1781 BP, Mellemholm), CGG107417 (1650 BP, Gammel Hasseris Grusgrav) and CGG106533 (1227BP,  Hellevad). These individuals likely represent the contact between the two regions at a time in which crossing the sea to the North was relatively easy compared to travelling south on land[224].

##### S5.7.4 Langobards and Goths

Of the Langobards in the dataset from Hungary, the Czech Republic and Italy, 26 are modelled primarily by Scandinavian IA-related ancestry, whereas 15 are modelled by ancestries more common in the region: Bell Beaker-related and Farming-related (Fig. S5.175). A single Langobard is modelled with some East Asian ancestry, however this individual has been flagged as contaminated and should be interpreted with caution. The dense sampling from Italy around this time reveals that the Langobard individuals with Scandinavian ancestry are genetic outliers (Fig. S5.176); only three other individuals cluster as Southern Scandinavians (R31, R106 and R108 from Antonio et al.[167])

This contrasts with the Ostrogoths and Visigoths of Southern Europe, in which the majority of individuals (12) are modelled primarily with Farming-related ancestries. Notably, two individuals from are modelled with a large amount of Northern European Bronze Age ancestry, however these are associated with Southern Scandinavia and the Baltic cline, rather than Eastern Scandinavia. Finally, two individual from Crimea are modelled with a large amount of Corded Ware ancestry rather than the more temporally close Iron Age ancestries, similar to the more local Sarmatian individuals that they cluster with.

To compare the southern European Ostrogoth and Visigoth individuals with the recently published Polish Wielbark genomes by Stolarek et al.[34], we first modelled all Polish individuals with the Corded Ware (East) and Bell Beaker sources (Set 5, Fig. S5.177 - top row). Transitions from predominantly Corded Ware ancestry (grey) in the Corded Ware Period to Bell Beaker-related ancestry (blue) in the Bell Beaker Period, and back to Corded Ware-related ancestry during the Gothic Iron Age and Middle Ages are apparent. The transition to the Middle Ages is accompanied by the appearance of additional EHG and

Anatolian Bronze Age-related ancestry. When including the Bronze Age Scandinavians sources (Set 5, Fig. S5.177 - bottom row), the Gothic Iron Age individuals are modelled as Eastern Scandinavian BA ancestry (light green), whereas by the Middle Ages individuals are modelled as the ‘Baltic’ Bronze Age ancestry (pink), much like Slavic-related groups in the region (Supplementary Note S5.7.6), with a small degree of continuity, consistent with previous studies[34] . This result provides further support that the Wielbark individuals had ancestral origins in Eastern Scandinavian and therefore likely were Goths.

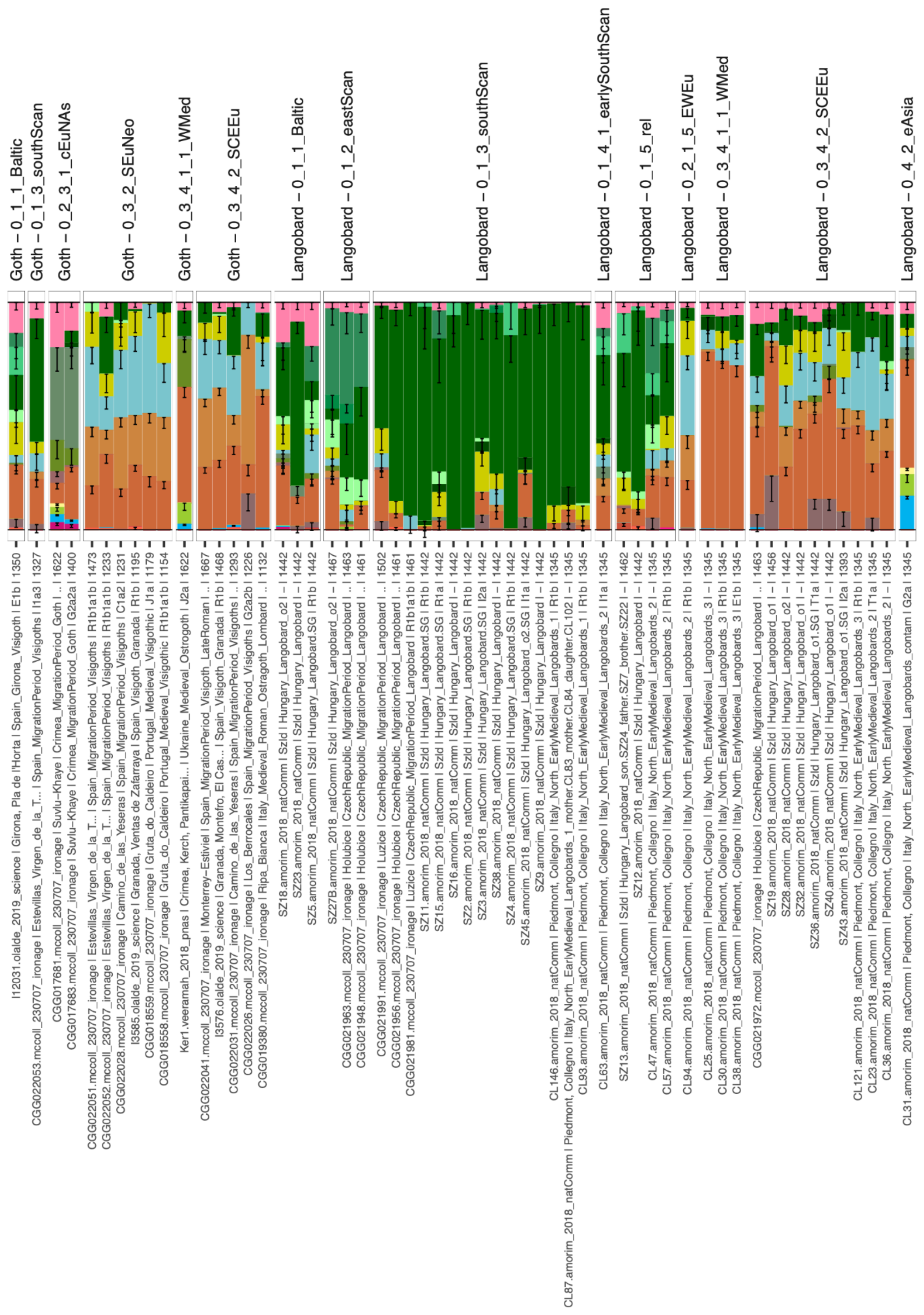

**Fig. S5.175. Mixture modelling results (Set 6) for Langobards and Goths, faceted by Culture and IBD clusters.**

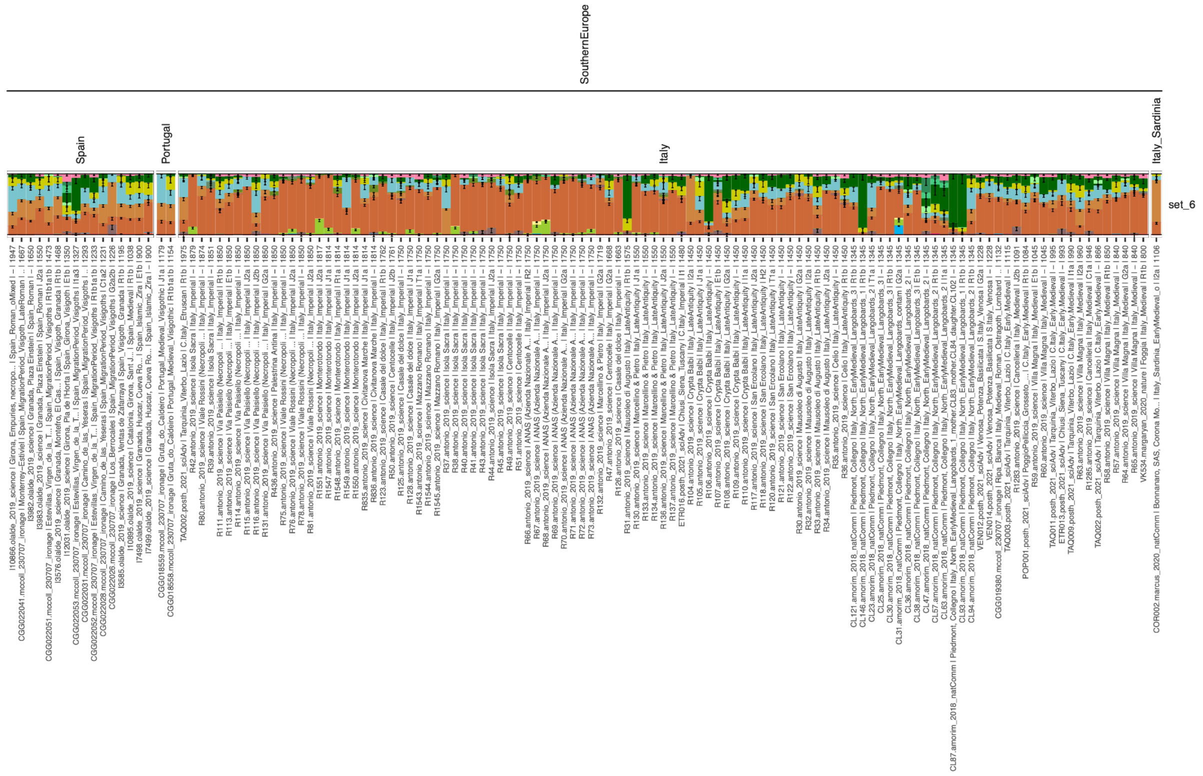

**Fig. S5.176. Mixture modelling results (Set 6) for Italy and Iberia from 2000 - 800 BP.** Excluding the Langobards, only two individuals are modelled with a large degree of Jutlandic IA ancestry (R31, R106 from Antonio et al.[167]).

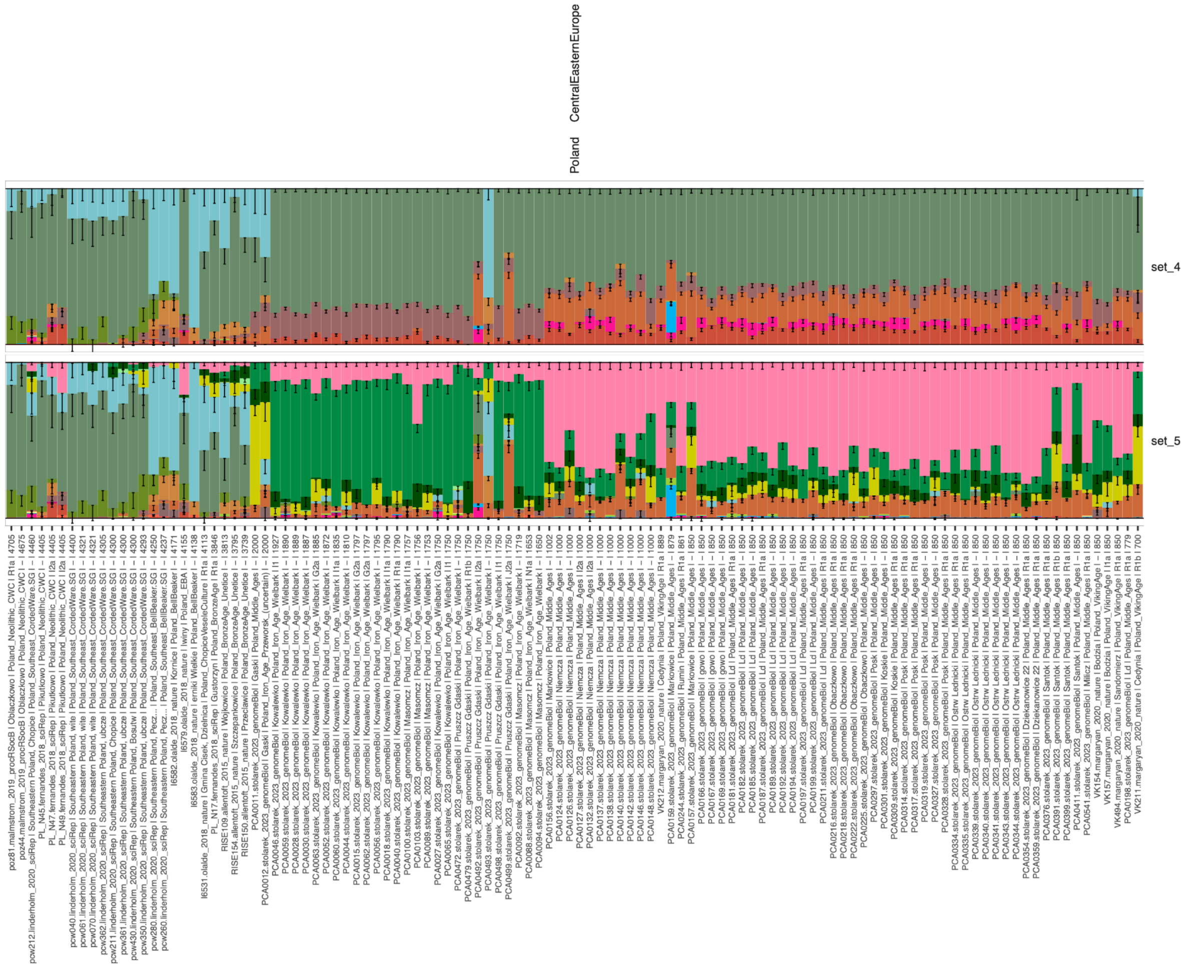

**Fig. S5.177. Mixture modelling results for Poland individuals.** Set 4 includes the Corded Ware (grey) and Bell Beaker (blue) sources. Set 5 includes the Corded Ware (North) Bronze Age sources, including Eastern Scandinavian (light green) and Southern Scandinavian (dark green) and Estonian Bronze Age (pink). The transitions from Corded Ware to Bell Beaker to Iron Age to Middle Ages indicate a series of population shifts over the last 4800 years.

##### S5.7.5. Britain and Ireland

After the arrival of Southern Scandinavian ancestry to Britain around 1500 BP, the majority of samples are modelled with North Germanic (Mecklenburg) ancestry. Notably, we find some outliers modelled with North Jutlandic IA rather than North German IA ancestry (Extended Data Fig.-12). Although bias in sampling may mean that the specific region and timing of the arrival of individuals with this profile cannot be identified, the heterogeneity present is expected due to the various homelands of the Angles, Saxons and Jutes along the Eastern North Sea coast migrating to Britain during this period.

The revelation of a migration into Denmark and Southern Sweden from a region similar to that of the Saxon migrations also has impacts on the interpretations of Gretzinger et al.[33], which found a source range for the Saxon migrations into Britain ranging from the Netherlands to Southern Sweden, but included samples post-dating the migration into Southern Scandinavia as potential sources. Similarly, the major influx of Danish Viking Age ancestry into England detected by Margaryan et al.[78] is likely in part impacted by the closely related sources for both populations.

By the Viking Age, we detect Eastern Scandinavian and Western Scandinavian ancestries across Britain and its Islands, representing Viking migrations from Sweden and Norway. Although migration from Denmark is likely during this period, the close relation between and heterogeneity within the Anglo-Saxons and the Danish Vikings limits our ability to detect this migration.

##### S5.7.6. The Baltic cline

The Baltic cluster is distinct from other Corded Ware (North) subclusters due to its high proportion of Eastern Hunter-Gatherer ancestry (Fig. S5.178, Supplementary Note S5.3.4). Later individuals from this cluster include populations related to the spread of Slavic populations  and are to be related to the Baltic Bronze Age ancestry originating in North East Europe. With the current sampling, determining a more precise homeland of the Slavic migrations is not possible.

From the mixture modelling with the Baltic Bronze Age as a source (Fig. S5.178: Set 5), the Slavic-associated populations from Slovakia, Czech Republic and Poland are modelled primarily as this Baltic-related ancestry.

Throughout the Iron Age, individuals with ancestral origins across the Baltic were found in Norway and Denmark (Fig. S5.29, Extended Data Fig.-13). By the late Iron Age, the presence of this ancestry into Scandinavia becomes more common, associated with Slavic populations, who migrated into areas south of the Baltic Sea formerly settled by East Germanic speakers.

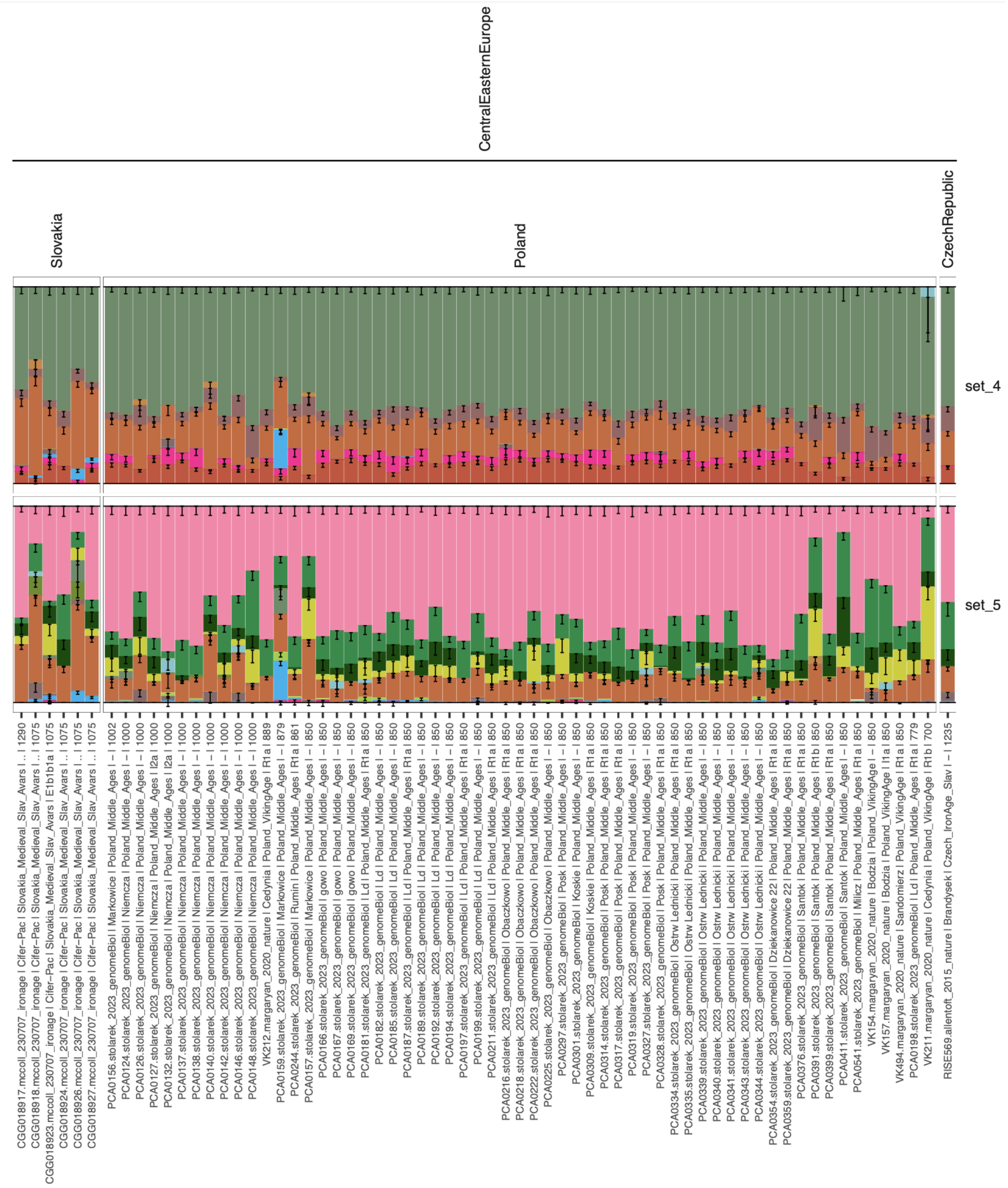

**Fig. S5.178. Mixture modelling results for Slavic populations, including the Stolarek et al.[34] individuals.** The relevant sources for Set 4 includes the Corded Ware (grey-green) and the EHG ancestry (bright pink). Set 5 includes the Corded Ware (North) Bronze Age sources, including Eastern Scandinavian (light green) and Southern Scandinavian (dark green) and Estonian Bronze Age (light pink).

#### S5.7.7. Aymyrlyg

The site of Ayrmyrlyg in Russia around 2200 BP is home to individuals from both the Saka and Hun cultures. Under IBD mixture modelling Set 4, the Saka individuals are modelled with similar ancestry proportions to the Sakas of Kazakhstan and Kyrgyzstan, with ~50% Steppe ancestry, smaller proportions of Iranian, Neolithic Farming and Caucausus Hunter-Gatherer and Eastern Hunter-Gatherer ancestry. Of the Huns, two are modelled here entirely as East Asian ancestry (light blue), similar to the Hun from Kazakhstan. The third Hun is modelled similar to the Saka (CGG021500), and the last has proportions expected from an individual deriving 50% of their ancestry from an East Asian source, and 50% from a Saka source.

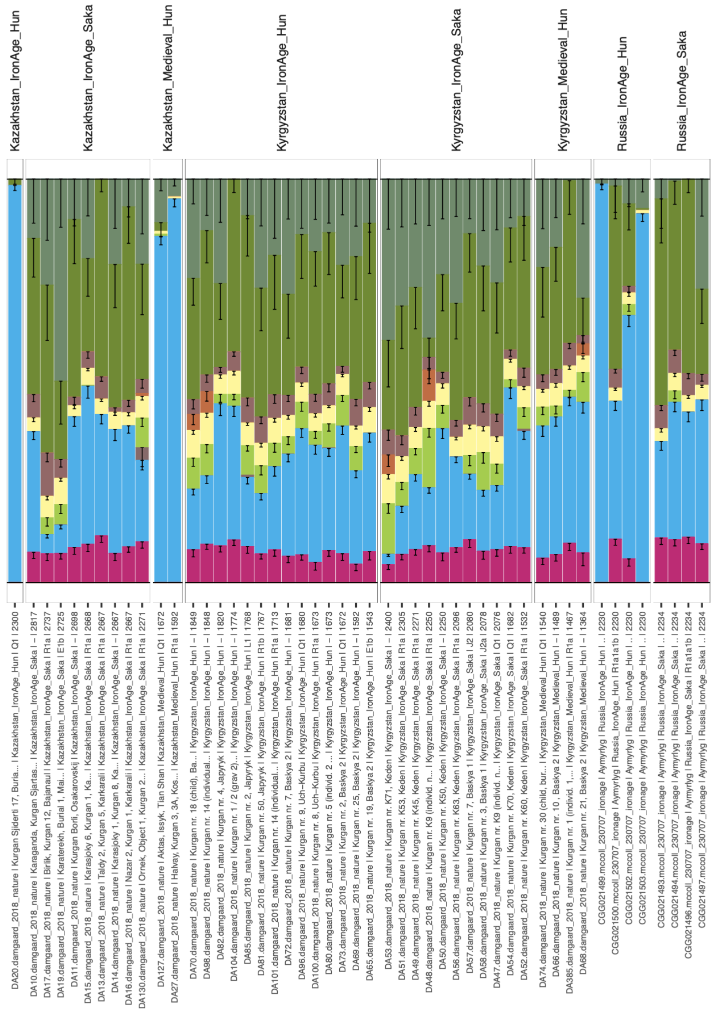

**Fig. S5.179. Mixture modelling results (Set 4) for Han and Saka individuals from Kazakhstan, Kyrgyzstan and Russia.** Here, East Asian ancestry is widespread (blue), in varying amounts, broadly corresponding to the cultural label.

### S6. Pre-Viking migration into Scandinavia

#### S6.1. Summary

*Ralph Fyfe, Marie-José Gaillard*

Large-scale demographic trends are reflected in the history of land use, and informed by data from dendrochronology and palynology. This evidence suggests profound changes across wide parts of Scandinavia between 1600 and 1200 BP.

Data from dendrochronology shows an abrupt cooling episode after c. 1415 BP (535 CE), a trend associated with the global Late Antique Little Ice Age (LALIA). Across Scandinavia, the impact has been shown to vary regionally[225], however, studies of tree-rings from Denmark show the most significant decrease of growth in 1411 BP (539 CE), compared to 1415 BP (535 CE)[226], and transitions in land use over this time. By reconstructing the vegetation through pollen records, we are able to gain insight into human activity during this period. The expectation for a gradual population decline followed by gradual immigration would be a decrease and discontinuity in land use resulting in the recovery of vegetation over an extended period of time, with heterogeneity in the forms of land use occurring over time. In contrast, the expectation for a rapid replacement would be a transition from heterogeneity in the region that has developed over time to homogeneity of land use of the incoming population.

O'Dwyer et al.[227] used pollen data from Skåne, Halland, Blekinge and Småland to quantify local vegetation cover for a number of time windows from 7500 BP to present using the

Landscape Reconstruction Algorithm (REVEALS and LOVE models[228,229]). The published maps of land cover indicate a clear increase in broadleaf deciduous woodland and a decrease in open land in the time window 1400–1500 BP (450–550 CE) compared to the previous and following time windows, 1900–2000 BP (50 BC–50 CE) and 1000–1100 BP (850–950 CE), respectively. The changes are most pronounced in southernmost Scania, the western coast of Scania and Halland, the eastern coast of Scania and Blekinge. In the time span 1100 - 1000 BP (850–950 CE), it is mainly southernmost Scania that regains its large landscape openness, while northern Småland becomes for the first time as deforested as southern Scania. Deforestation of Småland at that time is explained by the development of iron production (e.g. Lindström et al.[230]). For the purpose of the present study, we re-examined the pollen data from Skåne (Supplementary Note S6.3, Supplementary Table S6.1 and Supplementary Fig. S7.4 and Supplementary Fig. S7.5 ), looking at the pollen percentage changes of broadleaved trees (as a group of taxa) and four indicators of deforestation and agriculture. It shows a clear decrease in agriculture in lowland Scania (southern and northern hummocky regions) c. 1400 - 1300 BP (550–650 CE), while the more marginal sites at higher locations in the inland (Bökesjön), in NW Scania and at the northwestern coast show a decrease in grazing land already from 1800 - 1700 BP (150–250 CE) and very little or no cultivation of cereals all through the studied period until 1150 BP (800 CE). In lowland Scania, agriculture increased again from c. 1300 BP (650 CE) (southern hummocky area; e.g. Gaillard et al.[231]) or slightly later, and the same is valid for grazing in the marginal areas. It is clear that land use both decreased and underwent reorganisation from the time of the large ‘dynasty shift’ for Scania, from being an independent political community since 1500 BP (400 CE) to belonging to the realm of the Danes from 1410 BP (540 CE)[230]. From 1300 BP (650 CE), lowland areas were reorganised or reoccupied, and marginal areas were settled for the first time c. 1150 BP (800 CE), even though they were used for grazing earlier. The study by

Vinogradova et al.[232] using 5 pollen records and the Landscape Reconstruction Algorithm to quantify land-cover changes along the western coast of Småland N of Kalmar also indicates a decrease in landscape openness and a reorganisation of land use during the Migration Period. Without estimates of population change, it is assumed that the pollen records are primarily showing a major change in the land-use strategies and land management, as well as in the location of settlements, rather than changes in the intensity of human impact on the landscape, in accordance with assumptions made earlier by, e.g., Berglund et al.[233,234].

#### S6.2. The impact of the volcanic double event in 1414 BP (536 CE) and 1411/1410 BP (539/540 CE) on tree-ring growth

*Hanne Larsen, Morten Fischer Mortensen*

Dendrochronology provides a year-to-year record of growth conditions and hence constitutes a unique method to reveal past climate variation. The method's ability to capture the speed and regional impact of these changes makes it a powerful tool for understanding environmental shifts over time.

We examined dendrochronological data consisting of tree-ring measurements from 654 wood samples of *Quercus* sp. from 42 archaeological sites in Denmark covering the late Iron Age. Most sites are located in Jutland. The samples collected from various types of wooden constructions mainly originated from wells (54%) and bridge constructions (26%), but also pole bars from the seabed, timber, house constructions, canals and caissons. The dendrochronological data is believed to cover most of all wooden material from Denmark between 1650 and 1150 BP (300 CE and 800 CE). This time interval of 500 years (Fig. 1) was chosen to emphasise that no such remarkable decrease in tree-ring formation seemed to lie within the normal range of growth variation over an extended timeframe. The tree-ring measurements were combined into an average growth curve for Denmark by use of TSAP-Win$^{TM}$ Rinntech (version 4.82b2).

The average growth curve of the 654 samples of *Quercus* sp. showed a marked decrease in growth from 1415 and 1414 BP (535 CE to 536 CE) with a total reduction of 33% (Fig. 2), which was also expressed by very narrow growth rings on the wood samples (Fig. 4). Apart from a growth increment in 1413 BP (537 CE), the growth continued to decrease and reached a minimum in 1411 BP (539 CE) with a total growth reduction of 53% compared with 1415 BP (535 CE).

The average growth curve indicates that the climate changes towards much colder conditions. Climate modelling from Southern Norway indicates a temperature reduction of up to 3.5°C during the mid-sixth century[235]. The marked reduction in tree-ring growth in the years around 1414 BP (536 CE) and 1411/1410 BP (539/540 CE) is known from all over the Northern Hemisphere and is associated with major volcanic eruption followed by a global volcanic dust veil that reduced the solar radiation and lowered the global temperature[236].

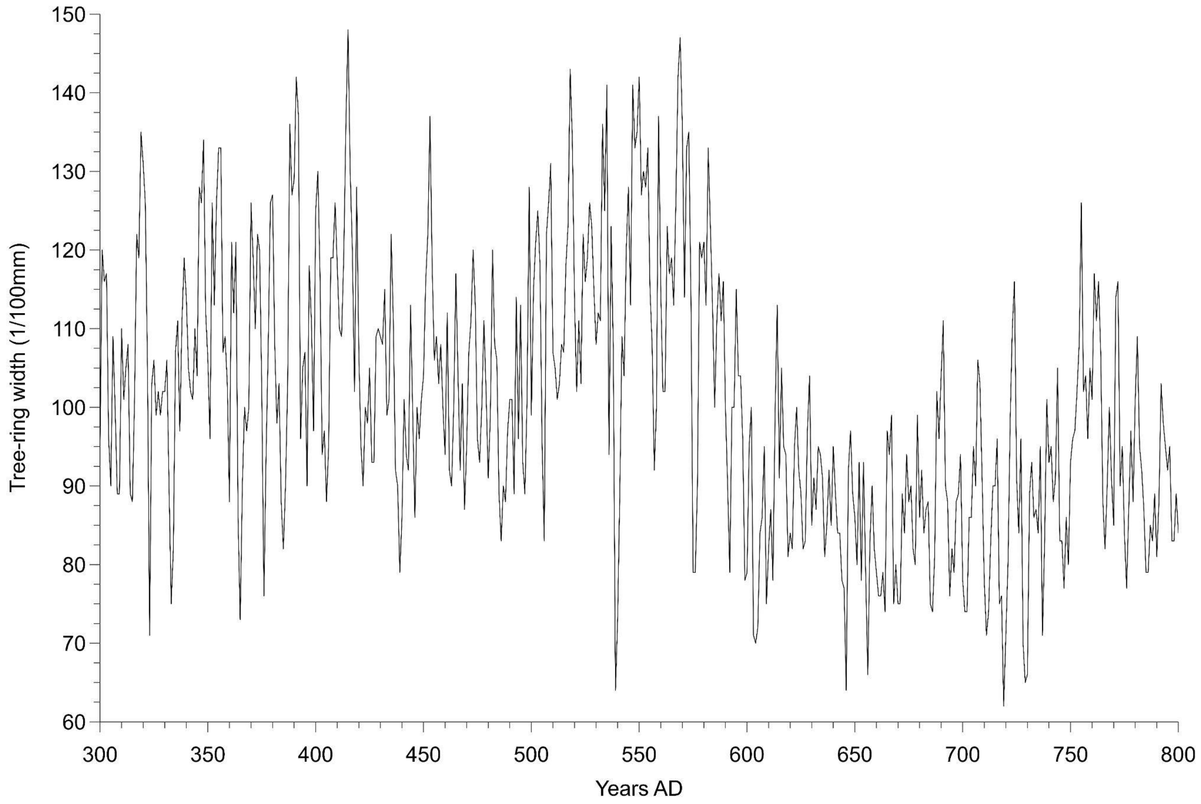

**Fig. S6.1. Average growth curve of *Quercus* sp. in Denmark from 300 to 800 CE (1650–1150 BP).**

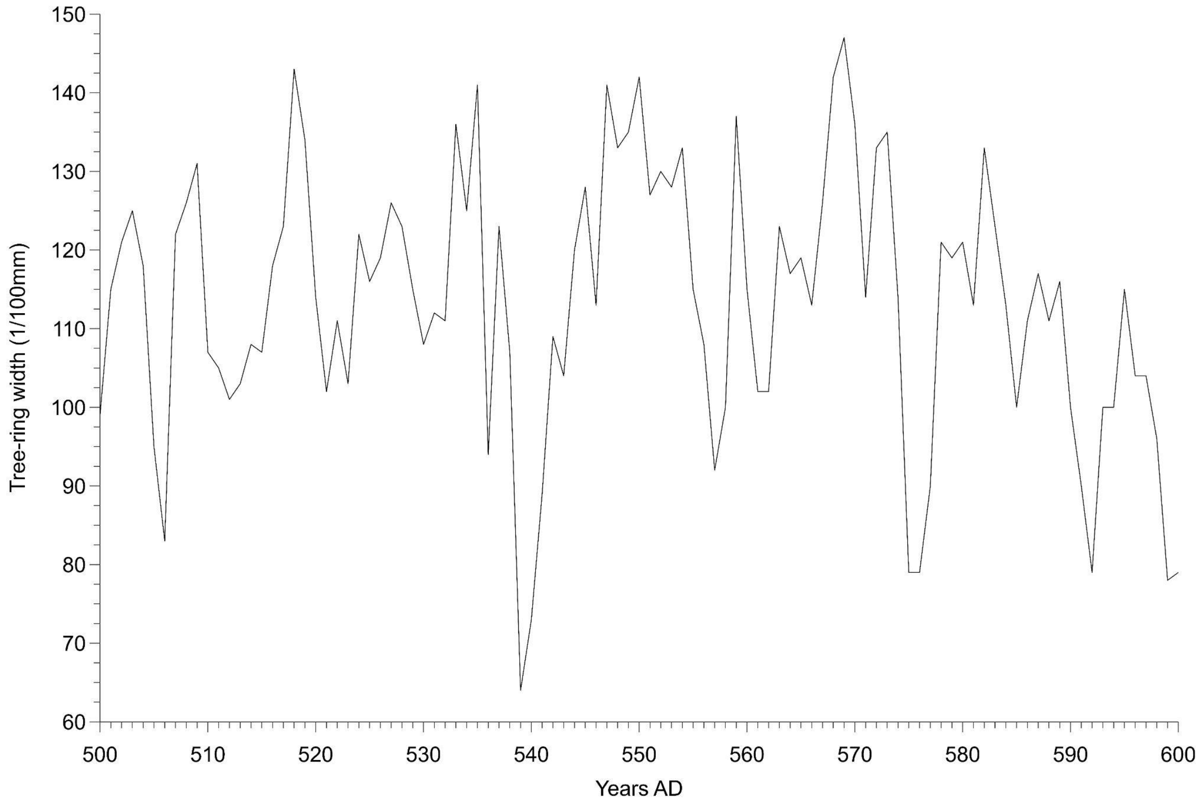

**Fig. S6.2. Average growth curve of *Quercus* sp. in Denmark from 500-600 CE (1450–1350 BP).**

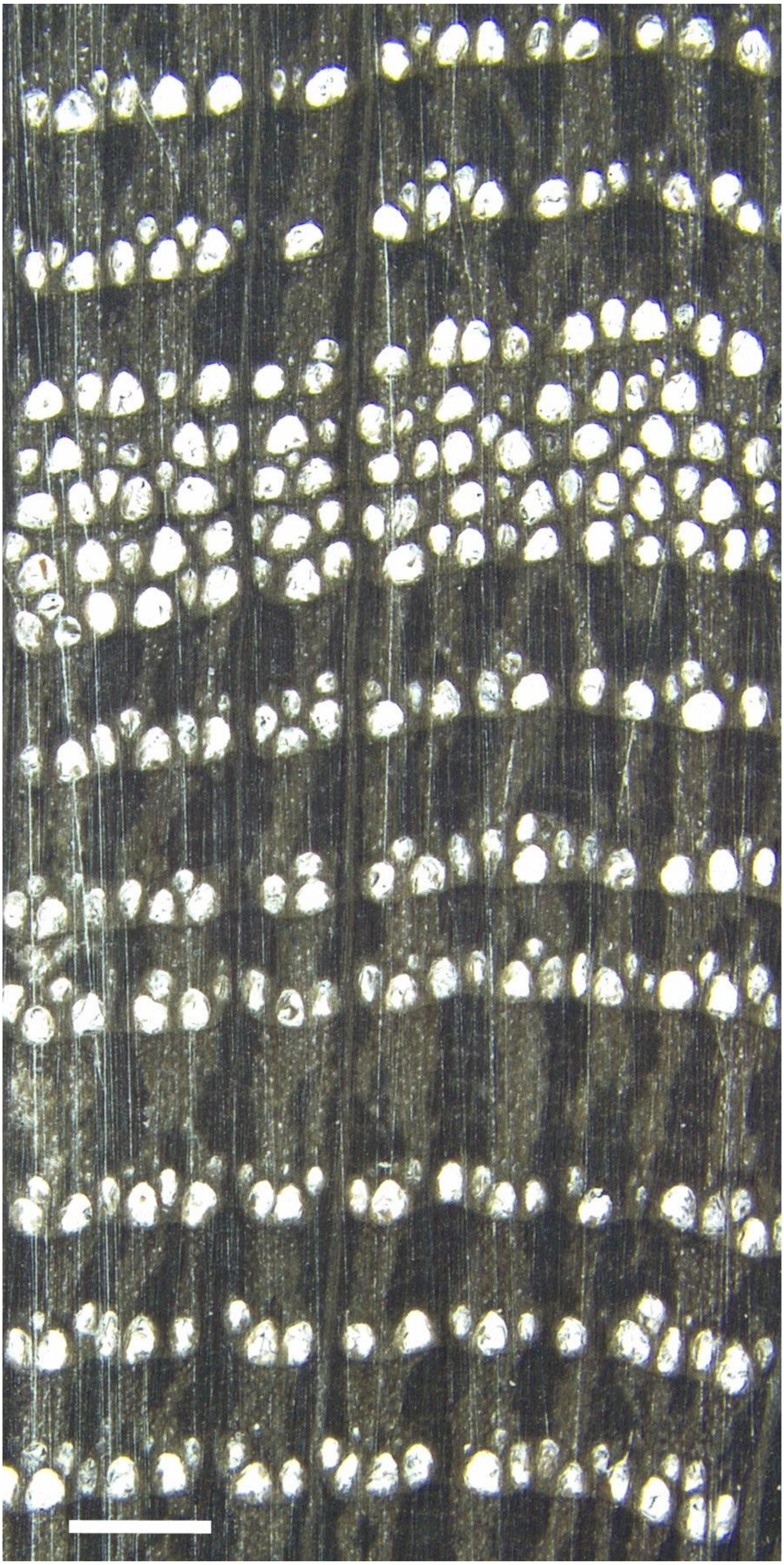

AD 542

AD 539

AD 536

**Fig. S6.3. The impact of the climate changes is shown by the formation of narrow growth rings between 539 and 542 CE (1411–1408 BP).**

### S6.3 Vegetation change 4000–800 cal BP in Scania, Southern Sweden

*Ralph Fyfe, Marie-José Gaillard*

**Datasets and analytical approach**

Twenty sequences were drawn from the LANDCLIMII project data archive for Scania [237] (Table S6.1), which was compiled using sequences available within the European Pollen Database[238,239] and provided by individual data contributors. The sequences were grouped into four geographical regions: southern hummocky landscapes, northern hummocky landscapes, and upland sites (i.e., marginal areas at slightly higher altitudes on horst ridges or Precambrian bedrock). In this way, we can identify major changes for three distinctive areas of Scania in terms of geology and soils; akaline soils for the S hummocky area, slightly acidic soils or acidic soils for the N hummocky area and acidic soils on eskers and NW Scania. Sequences were prepared for analysis by standardising the pollen nomenclature to a set of 87 of the most common taxa (those that were present in each sequence at at least 2% total land pollen) and samples dated to 4000–800 cal BP (2050 cal BC–1150 cal AD), extracted using the chronologies established in the LANDCLIMII project. A simplified indicator-species approach was used to explore the intensity of land-use practice over time by summarising the proportion of pollen types directly associated with cereal production (Cerealia types, *Secale t.* (rye)), open ground indicators including grasses (Poaceae) and heather (*Calluna vulgaris*) and deciduous trees (the sum of *Corylus*, *Betula*, *Quercus*, *Tilia* and *Ulmus*). In some regions of southern Sweden, heather can reflect the development of cultural heathlands used for grazing.

Pollen proportions were summarised in 100-year time intervals between 4000 and 800 cal BP.

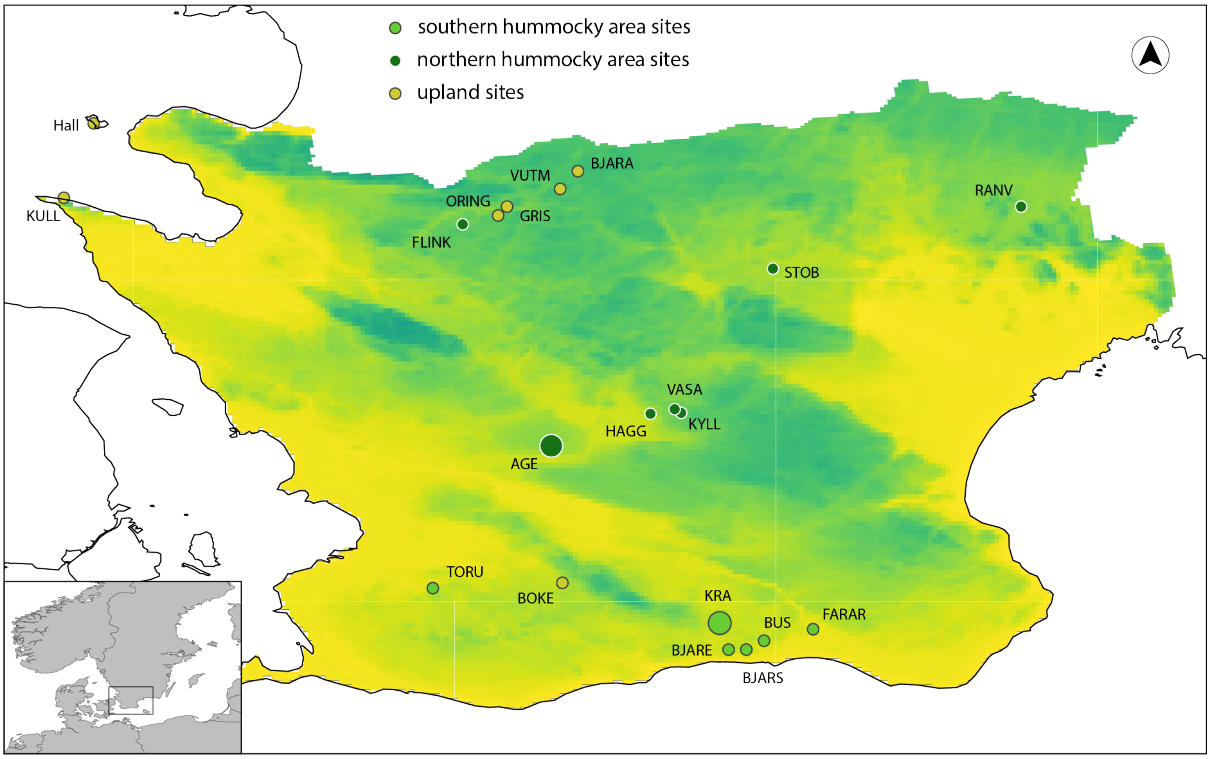

**Fig. S6.4. Location of pollen sequences used to assess intensity and nature of land use in southernmost Sweden, 4000–800 BP.** Larger circles indicate sites with a more regional character; small circles indicate local pollen sequences. Background colour indicates general pattern of elevation. Details of sites are in Table S6.1.

| Site name | Code | Grouping | References |
|---|---|---|---|
| Bjäresjösjön | BJARE | Southern | [231,240–242] |

| | | | |
|---|---|---|---|
| | | hummocky[240] | |
| Bjärsjöholmssjön | BJARS | Southern hummocky | [241,243] |
| Bussjösjön | BUS | Southern hummocky | [244] |
| Torup | TORU | Southern hummocky | [245] |
| Fararpsmosse | FARAR | Southern hummocky | [246] |
| Krageholmssjön | KRA | Southern hummocky | [247,248] |
| Flinkasjön | FLINK | Northern hummocky | [249] |
| Fulltofta/Häggenäs | HAGG | Northern hummocky | [250,251] |
| Fulltofta/Kyllingahus | KYLL | Northern hummocky | [250,251] |
| Fulltofta/Vasahus | VASA | Northern hummocky | [250,251] |

| | | | |
|---|---|---|---|
| Ranviken | RANV | Northern hummocky | [252] |
| Stoby | STOB | Northern hummocky | [253] |
| Ageröds Mosse | AGE | Northern hummocky | [254] |
| Kullaberg | KULL | Upland | [255] |
| Hälledammen | Hall | Upland | [251,256] |
| Bjärabygget | BJARA | Upland | [253] |
| Grisavad | GRIS | Upland | [253] |
| Östra Ringarp | ORING | Upland | [253] |
| Värsjö Utmark | VUTM | Upland | [253] |
| Bökesjön | BOKE | Upland | Gaillard, M.-J. unpublished |

**Table S6.1. Details of pollen sequences from Scania, taken from Githumbi et al.[237]**

**Results and interpretation**

The compilation of datasets from Scania shown in Fig. 7.3.1 shows the impact of agricultural groups since the early Bronze Age. Taking the region as a whole (Fig. 7.3.2A), landscape openness across the region increases around 3000 BP (reflected in both the Poaceae and *Calluna vulgaris* values), with a second increase (Poaceae and cereals) at 2600 BP indicating further opening of the landscape and increase in agriculture. Although h trends in landscape openness can be established using percentage pollen data, the degree of openness is likely strongly under-estimated in these data. Quantified vegetation openness, using the pollen record from Krageholmssjön (Fig. 7.3.1) and the REVEALS model[229], confirms the broad trends in changes, and indicates that tree cover drops from 75% to 35% in Scania between 2800 and 2400 BP[227,257,258]. The grouping of sites into upland, northern and southern hummocky areas demonstrates regional variability in land-cover openness. Sites to the south (Fig. 7.3.2C) show earlier increases in openness (around 3000 cal BP) and higher levels of openness, reflecting the earlier increase in agriculture with very large areas used for grazing[246]. Opening of the landscape in the northern hummocky landscape does not occur until several hundred years later (Fig. 7.3.2B). Cereal pollen types are recorded from as early as 4000 cal BP in the southern hummocky landscape; quantified regional vegetation estimates suggest that just under 1% of the landscape was under cereals between 2700 and 2200 BP[237]. Evidence for cereals is scattered in the northern hummocky region, and very limited or absent from the uplands.

The period between 1600 and 1200 BP (400–800 CE) is of particular interest for this study, and the evidence from the pollen record can be used to assess the changes in extent of forest recovery, and cultivation, across southern Sweden through this period. Fig. S6.5 demonstrates sub-regional trends within Scania, based on pollen proportion data. Cover of broadleaf trees increases in marginal landscapes at 1600 BP (Fig. S6.5D), and pollen proportions remain higher until 1300 BP. There is no evidence of cultivation throughout the period in the uplands. In the northern hummocky area, broadleaf tree pollen proportions initially increase (from, on average, 52 to 58% total land pollen); however, this woodland recovery is short-lived, with tree pollen proportions dropping again in the interval 1500–1400 BP. There is evidence for continued cereal cultivation during the period, but *Secale* is cultivated first from

c. 1200 BP. In the southern hummocky region, tree pollen proportions declined across the period 1600–1200 BP, with no evidence of woodland regeneration. The southern hummocky sites also show a gradual increase in intensity of land use between 1600 and 1200 BP, characterised by increased representation of cereals (Cerealia type and *Secale*).

The sub-regional picture that emerges from the careful examination of the pollen sequences is thus one of decrease in grazing of marginal land at 1600 cal BP, as reflected in the upland sites, decrease in land use (both grazing and agriculture) in the northern and southern hummocky areas, however rather a stagnation in the south c. 1600–1400 BP with no regeneration of broadleaved trees. Recovery in the northern hummocky regions is short-lived, with apparent reoccupation seen after around 100 years (by 1500 BP). Cultivation of cereals in the upland sites does not occur until after 1300–1200 BP.

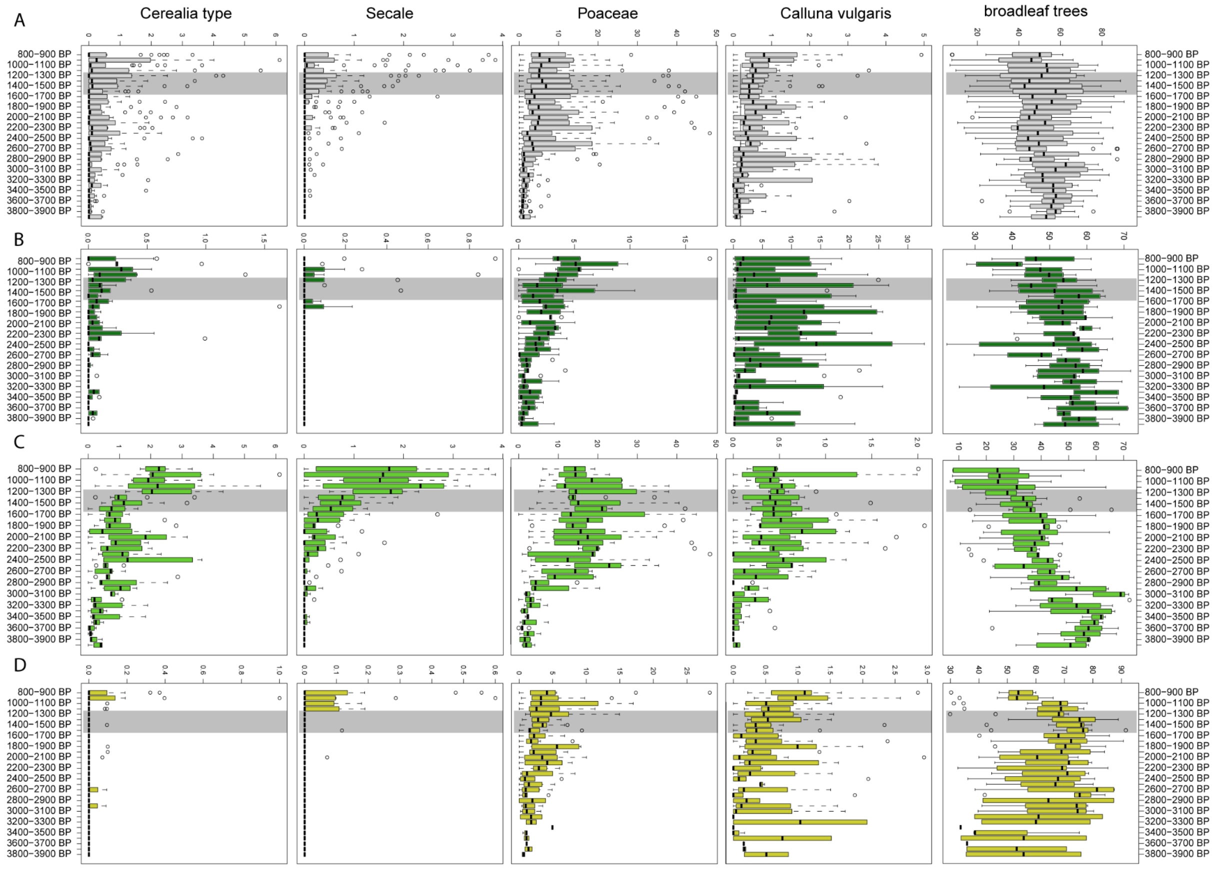

**Fig. S6.5. Summary of key pollen indicator types, southernmost Sweden.** Pollen data have been aggregated into 100-year time intervals. Broadleaf trees include *Corlyus*, *Betula*, *Quercus*, *Tilia* and *Ulmus*. Panel A: all sites (n=20); Panel B: northern hummocky region sites (7); Panel C: southern hummocky region sites (6); Panel D: upland sites (7). Locations of sites are shown on Fig. S6.4.

*Changes in Coastline, Climate and Human Settlement* (eds. Yanko-Hombach, V., Gilbert, A. S., Panin, N. & Dolukhanov, P. M.) (Springer Science & Business Media, 2006).

10. Kristiansen, K. *et al.* Re-theorising mobility and the formation of culture and language among the Corded Ware Culture in Europe. *Antiquity* **91**, 334–347 (2017).
11. Iversen, R. & Kroonen, G. Talking Neolithic: Linguistic and Archaeological Perspectives on How Indo-European Was Implemented in Southern Scandinavia. *Am. J. Archaeol.* **121**, 511–525 (2017).
12. Glob, P. V. Studier over den jyske enkeltgravskultur. in *Fortid Og Nutid* 298–302 (Gyldendal, 1945).
13. Ringe, D. Proto-Germanic. in *From Proto-Indo-European to Proto-Germanic* (ed. Ringe, D.) 0 (Oxford University Press, 2017). doi:10.1093/oso/9780198792581.003.0004.
14. Salmons, J. 59. The evolution of Germanic. in *Handbook of Comparative and Historical Indo-European Linguistics* vol. 2 1002–1027 (De Gruyter Mouton, 2017).
15. Hansen, B. S. S. & Kroonen, G. J. Germanic. in *The Indo-European Language Family* (ed. Olander, T.) 152–172 (Cambridge University Press, Cambridge, 2022). doi:10.1017/9781108758666.010.
16. Agee, J. R. Using Historical Glottometry to Subgroup the Early Germanic Languages. *J. Ger. Linguist.* **33**, 319–357 (2021).
17. Allentoft, M. E. *et al.* Population genomics of Bronze Age Eurasia. *Nature* **522**, 167–172 (2015).
18. Haak, W. *et al.* Massive migration from the steppe was a source for Indo-European languages in Europe. *Nature* **522**, 207–211 (2015).
19. *The Indo-European Puzzle Revisited: Integrating Archaeology, Genetics, and Linguistics*. (Cambridge University Press, 2023).

20. Prescott, C. Dramatic beginnings of Norway's history? Archaeology and Indo-Europeanization. in *Language and Prehistory of the Indo- European Peoples: A Cross-disciplinary Perspective* (ed. Hyllested, A.) (Museum Tusculanum Press, 2017).

21. Kroonen, G. Questions concerning the Indo-Europeanization of Scandinavia: The Germanic languages. in *Oxford Handbook of Scandinavian Archaeology* (eds. Hedeager, L., Prescott, C. & Kristiansen, K.) (Oxford University Press, Oxford, in press).

41. Hachmann, R., Kossack, G. & Kuhn, H. *Völker zwischen Germanen und Kelten: Schriftquellen, Bodenfunde und Namengut zur Geschichte des nördlichen Westdeutschlands um Christi Geburt*. (Wachholtz, Rhine River Valley, 1962).
42. Saag, L. *et al.* The Arrival of Siberian Ancestry Connecting the Eastern Baltic to Uralic Speakers further East. *Curr. Biol.* **29**, 1701-1711.e16 (2019).
43. Lazaridis, I. *et al.* The genetic history of the Southern Arc: A bridge between West Asia and Europe. *Science* **377**, eabm4247 (2022).
44. Mittnik, A. *et al.* The genetic prehistory of the Baltic Sea region. *Nat. Commun.* **9**, 442 (2018).
45. Saag, L. *et al.* Genetic ancestry changes in Stone to Bronze Age transition in the East European plain. *Sci. Adv.* **7**, eabd6535 (2021).
46. Posth, C. *et al.* Palaeogenomics of Upper Palaeolithic to Neolithic European hunter-gatherers. *Nature* **615**, 117–126 (2023).
47. Blank, M. Mobility, Subsistence and Mortuary practices. An interdisciplinary study of Neolithic and Early Bronze Age megalithic populations of southwestern Sweden. (University of Gothenburg, Gothenburg, 2021).
48. Blank, M. *et al.* Mobility patterns in inland southwestern Sweden during the Neolithic and Early Bronze Age. *Archaeol. Anthropol. Sci.* **13**, 64 (2021).
49. Larsson, Å. M. Breaking and Making Bodies and Pots : Material and Ritual Practices in Sweden in the Third Millennium BC. (Uppsala University, Uppsala, 2009).
50. Brorsson, T., Lucenius, J. & Stenbäck, N. Changing perspectives Thin section and ICP analysis of Neolithic pottery from the Åland Islands. in *Helsinki Harvest: Proceedings of the 11th Nordic Conference on the Application of Scientific Methods in Archaeology. MASF 7* vol. 7 (MASF (Monographs of the Archaeological Society of Finland), 2021).

51. Østmo, E. *Krigergraver: en dokumentarisk studie av senneolittiske hellekister i Norge*. (Kulturhistorisk Museum, Universtitet i Oslo, 2011).

52. Austvoll, K. I. *Seaways to Complexity: A Study of Sociopolitical Organisation Along the Coast of Northwestern Scandinavia in the Late Neolithic and Early Bronze Age*. (Equinox Publishing Limited, 2021).

53. Johannsen, J. W. The Late Neolithic Expansion: Ancient and new traditions 2350-1700 BC. *Dan. J. Archaeol.* **12**, 1–22 (2023).

54. Larsson, M. N. *et al.* Ancient Sheep Genomes reveal four Millennia of North European Short-Tailed Sheep in the Baltic Sea region. 2023.06.26.544912 Preprint at https://doi.org/10.1101/2023.06.26.544912 (2023).

55. Vandkilde, H. *The Metal Hoard from Pile in Scania, Sweden: Place, Things, Time, Metals, and Worlds around 2000 BCE*. (Aarhus Universitetsforlag, 2017).

56. Kristiansen, K. & Larsson, T. B. 5:3 The iconography of ruling elites: The Kivik burial and the origins of the Nordic Bronze Age. in *The rise of Bronze Age society: travels, transmissions and transformations* (Cambridge University Press, Cambridge ; New York, 2005).

57. Kristiansen, K. *Europe Before History. Cambridge*. (Cambridge University Press, Cambridge, 1998).

58. Narasimhan, V. M. *et al.* The formation of human populations in South and Central Asia. *Science* **365**, eaat7487 (2019).

59. Stang, C. S. *Lexikalische Sonderübereinstimmungen zwischen dem Slavischen, Baltischen und Germanischen*. (Univ.-Forl., 1972).

60. Hedeager, L. *Iron-Age Societies: From Tribe to State in Northern Europe, 500 BC to AD 700*. (Blackwell, 1992).

61. Fulk, R. D. *A Comparative Grammar of the Early Germanic Languages*. (John Benjamins, 2018). doi:10.1075/sigl.3.
62. Kümmel, M. J. Reconstructing Proto-Germanic. in *Oxford Research Encyclopedia of Linguistics* (2023). doi:10.1093/acrefore/9780199384655.013.931.
63. Polomé, E. C. Methodological Aspects of Glotto-and Ethnogenesis of the Germanic Peoples. *Entsteh. Von Sprachen Volkern Giotto- Ethnogenetische Aspekte Eur. Sprachen Akten Des* **6**, 45–70 (1985).
64. Hirt, H. Die Urheimat der Germanen: Nachwort. in *Neue Jahrbücher für das klassische Altertum, Geschichte und deutsche Litteratur* vol. 3 570–572 (1899).
65. Ringe, D. A. *From Proto-Indo-European to Proto-Germanic*. (Oxford University Press, 2017).
66. Nielsen, H. F. *The Germanic Languages: Origins and Early Dialectal Interrelations*. (University of Alabama Press, 1989).
67. Schwantes, G. Die Jastorf-Zivilisation. in *Reinecke Festschrift* (eds. Behrens, G. & Werner, J.) (Mainz, 1950).
68. Hedeager, L. *Iron Age Myth and Materiality: An Archaeology of Scandinavia AD 400-1000*. (Routledge, 2011).
69. Hines, J. *The Scandinavian Character of Anglian England in the Pre-Viking Period*. (University of Oxford, 1983).
70. Knol, E. & IJssennagger, N. Palaeogeography and People. in *Frisians and Their North Sea Neighbours: From the Fifth Century to the Viking Age* (eds. Hines, J. & IJssennagger, N.) 5–24 (Boydell & Brewer, 2017).
71. Hutterer, C. J. *Die germanischen Sprachen: ihre Geschichte in Grundzügen*. (Akadémiai Kiadó, 1975).
72. Diaconus, P. *History of the Lombards*. (University of Pennsylvania Press, 1974).

73. Wolfram, H. *History of the Goths*. (University of California Press, 1990).

74. Eiddon, I., Edwards, S. & Heather, P. Goths & Huns. in *The Cambridge Ancient History* (eds. Garnsey, P. & Cameron, A.) vol. 13: The Late Empire (Cambridge University Press, 1998).

75. Birkmann, T. 79. From Ancient Nordic to Old Nordic: Definition and delimitation of the period. in *79 . From Ancient Nordic to Old Nordic: Definition and delimitation of the period* 691–698 (De Gruyter Mouton, 2008). doi:10.1515/9783110197051-079.

76. Barnes, M. P. *Runes: A Handbook*. (Boydell Press, 2012).

85. Price, N. Climate change and the Fimbulwinter. in *Oxford Handbook of Scandinavian Archaeology* (Oxford University Press, Oxford, in press).

86. Jordanes. Chapter 1:23. in *The Gothic History of Jordanes in English Version* (Princeton University Press, 1915).

87. Procopius. *De Bellis [History of the Wars Book I-VIII]*. (Harvard University Press, 1918).

106. Schubert, M. *et al.* Improving ancient DNA read mapping against modern reference genomes. *BMC Genomics* **13**, 178 (2012).
107. Daley, T., Helus, V. & Smith, A. The preseq Manual. (2014).
108. Bushnell, B. BBMap: A Fast, Accurate, Splice-Aware Aligner. (2014).
109. Fu, Q. *et al.* A revised timescale for human evolution based on ancient mitochondrial genomes. *Curr. Biol.* **23**, 553–559 (2013).
110. Rasmussen, M. *et al.* Ancient human genome sequence of an extinct Palaeo-Eskimo. *Nature* **463**, 757–762 (2010).
111. Rasmussen, M. *et al.* The genome of a Late Pleistocene human from a Clovis burial site in western Montana. *Nature* **506**, 225–229 (2014).
112. Rasmussen, M. *et al.* The ancestry and affiliations of Kennewick Man. *Nature* vol. 523 455–458 (2015).
113. Fu, Q. *et al.* The genetic history of Ice Age Europe. *Nature* vol. 534 200–5 (2016).
114. Gamba, C. *et al.* Genome flux and stasis in a five millennium transect of European prehistory. *Nat. Commun.* **5**, 5257 (2014).
115. Lazaridis, I. *et al.* Ancient human genomes suggest three ancestral populations for present-day Europeans. *Nature* **513**, 409–413 (2014).
116. Olalde, I. *et al.* Derived immune and ancestral pigmentation alleles in a 7,000-year-old Mesolithic European. *Nature* **507**, 225–228 (2014).
117. Olalde, I. *et al.* A Common Genetic Origin for Early Farmers from Mediterranean Cardial and Central European LBK Cultures. *Mol. Biol. Evol.* **32**, 3132–3142 (2015).
118. Olalde, I. *et al.* The Beaker phenomenon and the genomic transformation of northwest Europe. *Nature* **555**, 190–196 (2018).
119. Olalde, I. *et al.* The genomic history of the Iberian Peninsula over the past 8000 years. *Science* **363**, 1230–1234 (2019).

197. Maier, R. & Patterson, N. admixtools: Inferring demographic history from genetic data. (2024).
198. Li, H. A statistical framework for SNP calling, mutation discovery, association mapping and population genetical parameter estimation from sequencing data. *Bioinformatics* **27**, 2987–2993 (2011).
199. Katoh, K. & Standley, D. M. MAFFT Multiple Sequence Alignment Software Version 7: Improvements in Performance and Usability. *Mol. Biol. Evol.* **30**, 772–780 (2013).
200. Kozlov, A. M., Darriba, D., Flouri, T., Morel, B. & Stamatakis, A. RAxML-NG: a fast, scalable and user-friendly tool for maximum likelihood phylogenetic inference. *Bioinformatics* **35**, 4453–4455 (2019).
201. Diamond, J. & Bellwood, P. Farmers and Their Languages: The First Expansions. *Science* **300**, 597–603 (2003).
202. Poznik, G. D. *et al.* Punctuated bursts in human male demography inferred from 1,244 worldwide Y-chromosome sequences. *Nat. Genet.* **48**, 593–599 (2016).
203. Jobling, M. A. & Tyler-Smith, C. Human Y-chromosome variation in the genome-sequencing era. *Nat. Rev. Genet.* **18**, 485–497 (2017).
204. A recent bottleneck of Y chromosome diversity coincides with a global change in culture. https://genome.cshlp.org/content/early/2015/03/13/gr.186684.114.abstract.
205. Y. Chromosome Consortium. A nomenclature system for the tree of human Y-chromosomal binary haplogroups. *Genome Res.* **12**, 339–348 (2002).
206. Karafet, T. M. *et al.* New binary polymorphisms reshape and increase resolution of the human Y chromosomal haplogroup tree. *Genome Res.* **18**, 830–838 (2008).
207. Hallast, P. *et al.* The Y-Chromosome Tree Bursts into Leaf: 13,000 High-Confidence SNPs Covering the Majority of Known Clades. *Mol. Biol. Evol.* **32**, 661–673 (2015).

225. Gundersen, I. M. Iron Age Vulnerability. The Fimbulwinter hypothesis and the archaeology of the inlands of eastern Norway. (Faculty of Humanities, University of Oslo, 2022).

226. Ellegård Larsen, H. M. Træernes årringe afslører fortidens klima: Nordisk dendrokonference 2023. (2023).

227. O'Dwyer, R., Marquer, L., Trondman, A.-K. & Jönsson, A. M. Spatially Continuous Land-Cover Reconstructions Through the Holocene in Southern Sweden. *Ecosystems* **24**, 1450–1467 (2021).

228. Sugita, S. Theory of quantitative reconstruction of vegetation II: all you need is LOVE. *The Holocene* **17**, 243–257 (2007).

229. Sugita, S. Theory of quantitative reconstruction of vegetation I: pollen from large sites REVEALS regional vegetation composition. *The Holocene* **17**, 229–241 (2007).

230. Lindström, J. *Sveriges långa historia*. (Norstedts, 2022).

231. Gaillard, M. J., Olausson, D. & Skansjö, S. The Bjäresjö area. Conclusions. in *The Cultural Landscape During 6000 Years in Southern Sweden: The Ystad Project* (ed. Berglund, B. E.) (Munksgaard, Denmark, 1991).

232. Vinogradova, O. *et al.* 3000 Years of past regional and local land-use and land-cover change in the southeastern Swedish coastal area: Early human-induced increases in landscape openness as a potential nutrient source to the Baltic Sea coastal waters. *The Holocene* **34**, 56–73 (2024).

233. Berglund, B. E., Larsson, L., Lewan, N., Olsson, G. A. & Stensjö, S. Ecological and Social factors behind the landscape changes. Cultural landscape - patterns and processes. in *The Cultural Landscape During 6000 Years in Southern Sweden: The Ystad Project* (ed. Berglund, B. E.) 167–174 (Munksgaard, Denmark, 1991).

234. Berglund, B. E., Hjelmroos, M. & Kolstrup, E. The Köpinge area. Vegetation and landscape through time. in *The Cultural Landscape During 6000 Years in Southern Sweden: The Ystad Project* (ed. Berglund, B. E.) 109–112 (Munksgaard, Denmark, 1991).

235. van Dijk, E. *et al.* Climatic and societal impacts in Scandinavia following the 536 and 540 CE volcanic double event. *Clim. Past* **19**, 357–398 (2023).

236. Gundersen, I. M. Iron Age Vulnerability. The Fimbulwinter hypothesis and the archaeology of the inlands of eastern Norway. (University of Oslo, Oslo, 2021).

237. Githumbi, E. *et al.* European pollen-based REVEALS land-cover reconstructions for the Holocene: methodology, mapping and potentials. *Earth Syst. Sci. Data* **14**, 1581–1619 (2022).

238. Giesecke, T. *et al.* Towards mapping the late Quaternary vegetation change of Europe. *Veg. Hist. Archaeobotany* **23**, 75–86 (2014).

239. Fyfe, R. M. *et al.* The European Pollen Database: past efforts and current activities. *Veg. Hist. Archaeobotany* **18**, 417–424 (2009).

240. Gaillard, M.-J. & Berglund, B. E. *Land-Use History during the Last 2700 Years in the Area of Bjäresjö, Southern Sweden*. (Lunds universitet, 1988).

241. Gaillard, M. J. & Göransson, H. The Bjäresjö area - Vegetation and landscape through time. in *The Cultural Landscape During 6000 Years in Southern Sweden: The Ystad Project* (ed. Berglund, B. E.) 67–173 (Munksgaard, Denmark, 1991).

242. Gaillard, M. J., Dearing, J. A., El-Daoushy, F., Enell, M. & H□kansson, H. A multidisciplinary study of the lake Bj□resj□sj□n (S Sweden): land-use history, soil erosion, lake trophy and lake-level fluctuations during the last 3000 years. *Hydrobiologia* **214**, 107–114 (1991).

243. Göransson, H. *Vegetation and Man Around Lake Bjärsjöholmssjön During Prehistoric Time: A Contribution to the Project" Kulturlandskapet Under 6000 År"(The Cultural Landscape During the Last 6000 Years in Southern Sweden)*. (Lund University, Department of Quaternary Geology, 1991).

244. Regnéll, J. *Vegetation and Land Use During 6000 Years: Palaeoecology of the Cultural Landscape at 2 Lake Sites in Southern Skåne, Sweden*. (Lund Univ., Department of Quaternary Geology, 1989).

245. Hultberg, T., Brunet, J., Broström, A. & Lindbladh, M. Forest in a cultural landscape - the vegetation history of Torup in southernmost Sweden. *Ecol. Bull.* 141–154 (2010).

246. *The Cultural Landscape During 6000 Years in Southern Sweden: The Ystad Project*. (Munksgaard, Denmark, 1991).

247. Gaillard, M.-J. Water-Level Changes, Climate and Human Impact: A Palaeohydrological Study of Krageholm Lake (Scania, Southern Sweden). in *Climatic Changes on a Yearly to Millennial Basis: Geological, Historical and Instrumental Records* (eds. Mörner, N.-A. & Karlén, W.) 147–154 (Springer Netherlands, Dordrecht, 1984). doi:10.1007/978-94-015-7692-5_14.

248. Gaillard, M.-J. *A Palaeohydrological Study of Krageholmssjön (Scania, South Sweden). Regional Vegetation History and Water-Level Changes*. (Lunds Universitet, 1984).

249. Björkman, L. Detaljerad pollenanalytisk undersökning av tre lagerföljder (Flinkasjön, E4:13, E4:16) från Örkelljungatrakten i nordvästra Skåne. *LUNDQUA Uppdrag* **54**, 1–20 (2004).

250. Lindbladh, M. *et al.* Forest History as a Basis for Ecosystem Restoration—A Multidisciplinary Case Study in a South Swedish Temperate Landscape. *Restor. Ecol.* **15**, 284–295 (2007).

251. Lindbladh, M. & Foster, D. R. Dynamics of long-lived foundation species: the history of *Quercus* in southern Scandinavia. *J. Ecol.* **98**, 1330–1345 (2010).

252. Digerfeldt, G. The Post-Glacial development of the Ranviken bay in Lake Immeln; I. The history of the regional vegetation, and II. The water-level changes. *Geol. Fören. Stockh. Förh.* **96**, 3–32 (1974).

253. Lagerås, P. *The Ecology of Expansion and Abandonment: Medieval and Post-Medieval Agriculture and Settlement in a Landscape Perspective*. (Riksantikvarieämbetet forlag, 2007).

254. Nilsson, T. Standardpollendiagramme und C14-Datierungen aus dem Ageröds Mosse im mittleren Schonen. *LUNDS Univ. ÄRSSKRIFT NF Avd 2 Bd 59 Nr 7* (1964).

255. Björkman, L. The role of human disturbance in Late Holocene vegetation changes on Kullaberg, southern Sweden: *Veg. Hist. Archaeobotany* **10**, 201–210 (2001).

256. Molinari, C. 2500 years of forest dynamics at Söndre Skog, a semi-natural forest on Hallands Väderö island, Southern Sweden. (Turin University, Italy, 2002).

257. Sugita, S., Gaillard, M.-J., Hellman, S. & Bröstrom, A. Model-based reconstruction of vegetation and landscape using fossil pollen. (2008).

258. Gaillard, M.-J. *et al.* Holocene land-cover reconstructions for studies on land cover-climate feedbacks. *Clim. Past* **6**, 483–499 (2010).
